# Systematic discovery of directional regulatory motifs associated with human insulator sites

**DOI:** 10.1101/2024.01.20.573595

**Authors:** Naoki Osato, Michiaki Hamada

## Abstract

Insulator proteins function as barriers to enhancer–promoter interactions (EPIs), thereby regulating gene expression. The primary insulator protein in vertebrates is CTCF, a DNA-binding protein (DBP); however, the roles of other DBPs in EPI insulation are not fully understood. To address this, we developed a systematic and comprehensive deep learning–based approach to identify DNA motifs of DBPs associated with insulator function. Applying this method to human fibroblast cells, we identified 97 directional motifs and a smaller number of non-directional motifs. These motifs were mapped to 23 DBPs previously linked to insulator activity, CTCF, and/or other forms of chromosomal transcriptional regulation. We found that the estimated orientation bias of CTCF was consistently proportional to the orientation bias observed in chromatin interaction data. Furthermore, these motifs showed significant enrichment at insulator sites that separate repressive and active chromatin regions, at chromatin interaction–defined boundaries, and at splice sites, compared to motifs of other DBPs. For instance, we observed that the key regulator MyoD-binding site is located at an insulator site near a gene involved in skeletal muscle differentiation and function. Importantly, our findings support the previously proposed insulator-pairing model, which suggests that insulator–insulator interactions are orientation-dependent, and highlight the involvement of multiple DNA-binding proteins beyond CTCF. Together, these results provide new insights into transcriptional regulatory mechanisms mediated by insulator-associated DBPs.

## Introduction

Enhancer ‒ promoter interactions (EPIs), which are organized by chromatin architecture, regulate gene transcription ^1^. Chromatin interactions establish loop structures stabilized by transcription factors (TFs) and DNA-binding proteins (DBPs), such as CTCF and components of the cohesin complex (e.g., RAD21 and SMC3). CTCF typically acts as an insulator by anchoring chromatin loops and restricting enhancer activity within these loops ^2^. Pairs of CTCF-binding sites display a characteristic directional bias, favoring a forward‒reverse (FR) orientation over forward ‒forward (FF), reverse‒reverse (RR), or reverse‒forward (RF) orientations ^3^. The forward and reverse orientations of DNA-binding sites are complementary in sequence and can be readily distinguished based on reference genome sequences. Moreover, although FR and RF orientations are sequence complements, their frequencies at chromatin loop anchors differ significantly ^3^, suggesting that the directional bias of DNA-binding sites is associated with chromatin loop formation.

While CTCF and cohesin are recognized as the primary insulator proteins in vertebrates ^4^, regional micro-capture (Micro-C) analyses have demonstrated that microcompartment interactions are largely independent of cohesin and transcription ^5^. These findings imply that other DBPs may function as insulators or be associated with insulator activity. However, experimentally identifying insulator-associated DBPs among the numerous DBPs present in the genome remains labor-intensive and technically challenging when relying on in situ assays.

One potential approach to identifying insulator-associated DBPs is to analyze the enrichment of DNA-binding sites for chromatin- and insulator-associated proteins at chromatin interaction sites. For example, DBPs such as ZNF143 and YY1 have been shown to participate in chromatin interactions and enhancer‒promoter interactions (EPIs) ^1,6^. However, while the binding sites of ZNF143 are indeed enriched at chromatin interaction sites, those of YY1 are not significantly enriched ^6^. Moreover, CTCF, ZNF143, and YY1 participate in a variety of biological functions beyond chromatin organization. This multifunctionality likely reduces motif enrichment at chromatin interaction sites, complicating the identification of DBPs that are specifically associated with insulator functions.

Another potential approach for predicting insulator-associated DBPs is to focus on the directional bias of CTCF DNA-binding sites. The orientation of a DBP’s binding site may not be relevant when the protein binds to a single genomic location ^7^. However, when a DBP or a complex of DBPs binds to multiple genomic loci through chromatin interactions, their binding is often structurally organized, contributing to the observed directional bias of DNA-binding sites ^8–10^. Some DBPs involved in chromatin interactions, such as YY1, are multifunctional proteins, with certain binding sites participating in both chromatin interactions and insulator functions. As a result, the directional bias of DNA-binding sites for multifunctional DBPs may be diminished. This multifunctionality complicates the identification of their specific insulator-associated roles.

Computational methods have been developed to predict chromatin interactions and their associated DBPs ^1112^. However, these methods are not designed to identify the binding motifs of DBPs specifically associated with insulator function. Regarding DBP binding motifs, Xie et al. noted that a key challenge is developing systematic approaches for determining their specific functions on a genome-wide scale ^13^. In terms of gene regulation, insulators influence EPIs and gene expression; therefore, incorporating gene expression data may improve the identification of insulator-associated DBPs compared to approaches that rely solely on enrichment analysis or the directional bias of DNA-binding sites. Recently, a novel deep learning–based method was introduced to estimate gene expression levels in human tissues based on DNA-binding sites within promoter regions ^14^. However, this method does not account for DNA-binding sites associated with EPIs as input features and was not intended to identify DBPs linked to insulator function.

Therefore, we aimed to identify insulator-associated DBPs using a computational approach that integrates and analyzes genome-wide experimental data. In this study, we refined an existing deep learning method by incorporating EPI information based on eQTL data and enhancer‒promoter association rules (McLean et al., Supplementary Fig. 2) ^15^. We analyzed the directional bias of DNA-binding sites for CTCF and insulator-associated DBPs, including known factors such as RAD21 and SMC3, using contribution scores generated by the deep learning model. Our analysis demonstrated that the directional bias of CTCF binding sites closely matches the orientation bias observed in chromatin interaction data. Furthermore, statistical testing of contribution scores between directional and non-directional DNA-binding sites of insulator-associated DBPs showed that directional sites contribute more significantly to gene expression prediction than non-directional sites. Overall, we identified 23 insulator-associated DBPs related to insulator function, CTCF-associated activity, and/or other transcriptional regulatory mechanisms affecting EPIs, as previously reported. To validate these findings, we showed that the DNA-binding sites of the identified DBPs are located within putative insulator regions, and that some of these sites exist independently of nearby CTCF or cohesin binding sites. Finally, consistent with the insulator-pairing model proposed from experimental studies in flies, we confirmed that both homologous and heterologous insulator–insulator pairing interactions are orientation-dependent. Our approach facilitates the identification of insulator- and chromatin-associated DNA-binding sites that modulate EPIs and uncovers novel functional roles and molecular mechanisms of DBPs involved in transcriptional condensation, phase separation, and transcriptional regulation.

## Results

### Predictive modeling of gene expression from directional DBP binding

Accurate gene expression prediction can be achieved by integrating both distal and proximal DNA-binding protein (DBP) binding sites as features within a machine learning framework. By combining ChIP-seq data from the Gene Transcription Regulation Database (GTRD v20.06) with expression quantitative trait locus (eQTL) data from GTEx v8 ^16^, our model was able to account for enhancer–promoter interactions (EPIs) by linking single nucleotide polymorphisms (SNPs) to gene expression outcomes (**Fig. 1a-c**, **Fig. 2a**, **b**). Specifically, DBP binding sites within ±50 bp of an eQTL SNP were assumed to influence the associated transcript. Focusing on four human cell types—HFFs, monocytes, HMECs, and neural progenitor cells—the model achieved strong concordance with observed expression levels: Spearman’s *ρ* values reached 0.74, 0.71, 0.74, and 0.73 for all 27,428 genes, and 0.70, 0.67, 0.69, and 0.70 for 2,705 test genes (**Fig. 3a**, **Fig. S2a**). Importantly, this performance was obtained without incorporating RNA-binding protein or miRNA data as required in previous approaches ^14^.

**Fig. 1.**
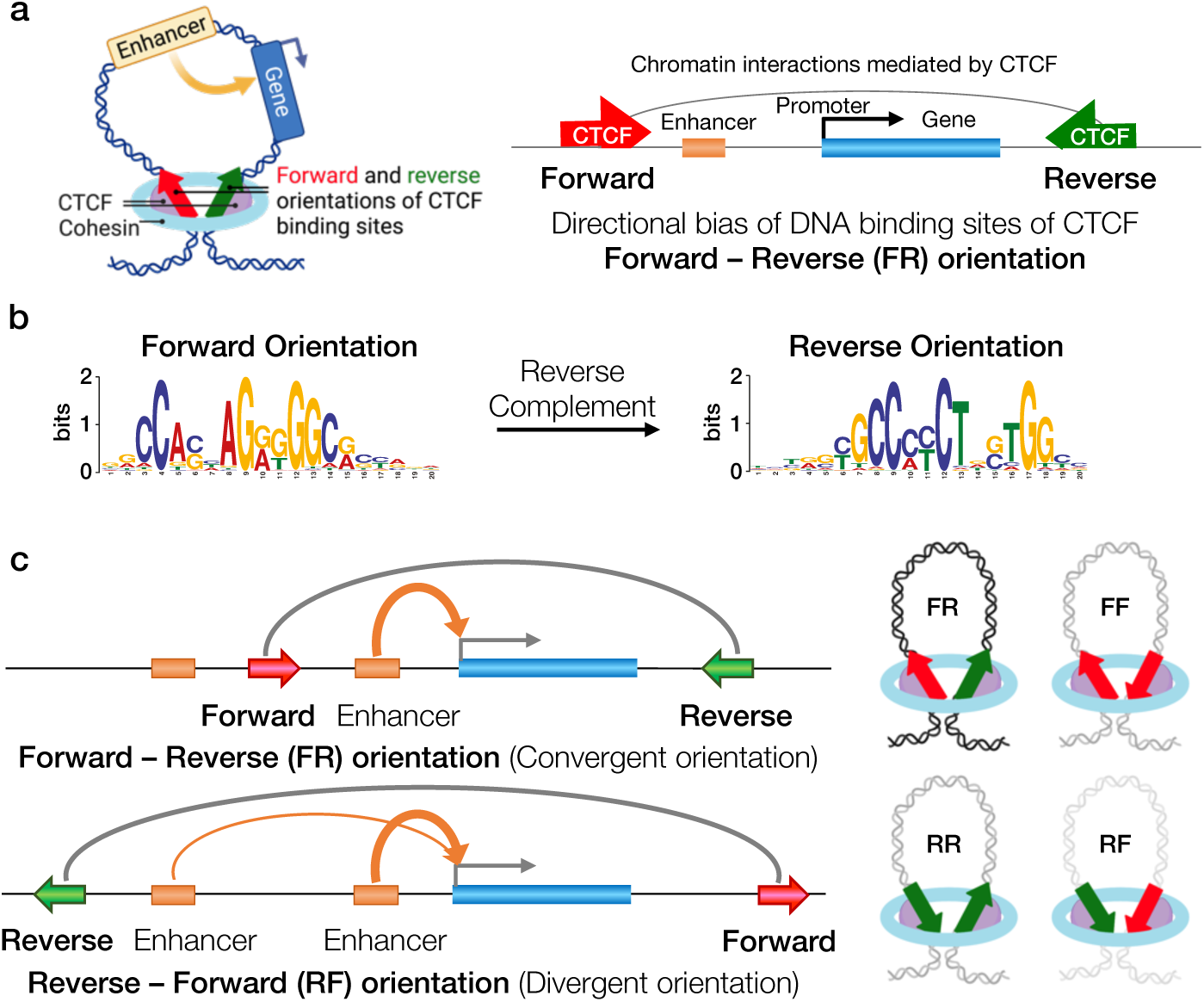
CTCF insulator function mediates enhancer–promoter interactions. **a** CTCF DNA-binding sites with a forward–reverse (FR) orientation are frequently observed at chromatin interaction sites. The forward and reverse orientations correspond to complementary sequences, which are clearly distinguishable in the reference genome. **b** Position weight matrices (PWMs) of CTCF binding motifs. The forward and reverse motifs are complementary on opposite DNA strands. A sequence logo is a graphical representation of nucleotide conservation, where the height of each letter reflects its frequency at that position (adapted from Wikipedia ‘Sequence logo’). **c** CTCF can block interactions between an enhancer and a promoter (or the transcription start site [TSS] of a gene), thereby restricting enhancer activity within a defined chromatin domain (region between red and green arrows). The red and green arrows indicate the DNA-binding sites of a putative insulator-associated DBP, such as the FR orientation of CTCF sites illustrated here. The numbers of DNA-binding sites with FR, RF, FF, and RR orientations differ, as reflected by the intensity of chromatin loops (right panel). Images of double-stranded DNA loops were adapted from “Transcriptional Regulation by CTCF and Cohesin” at BioRender.com (2023).

**Fig. 2.**
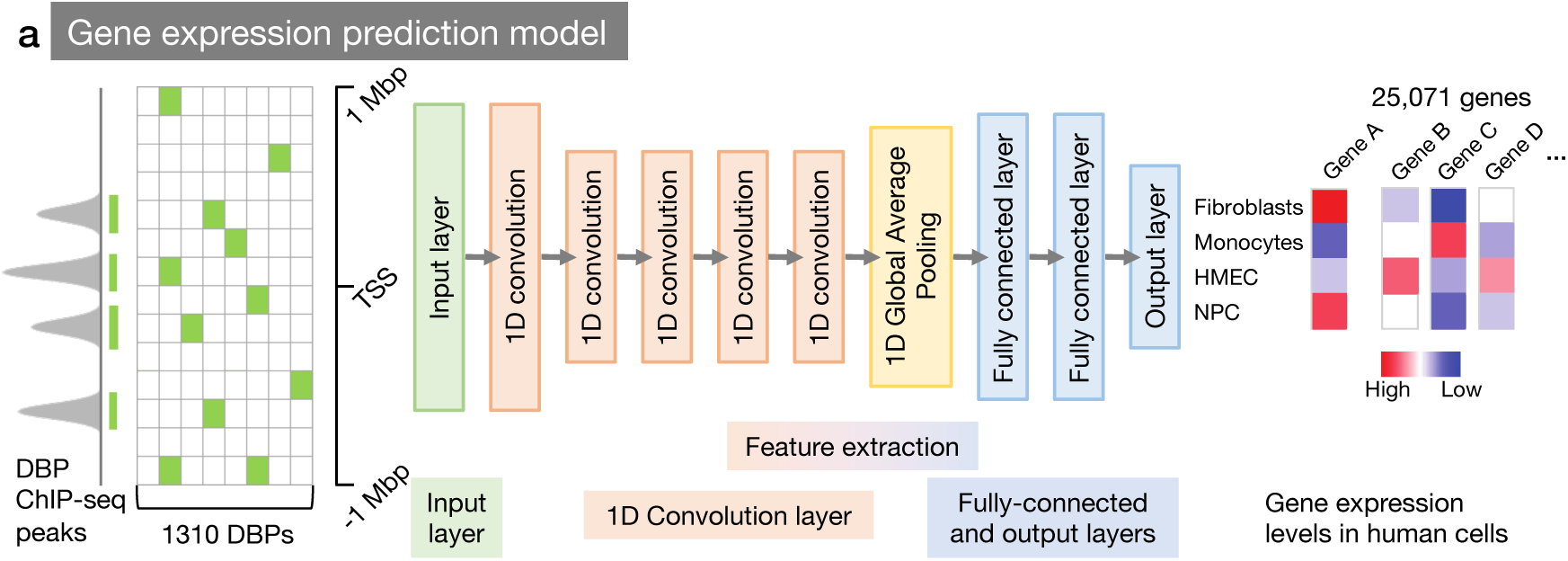
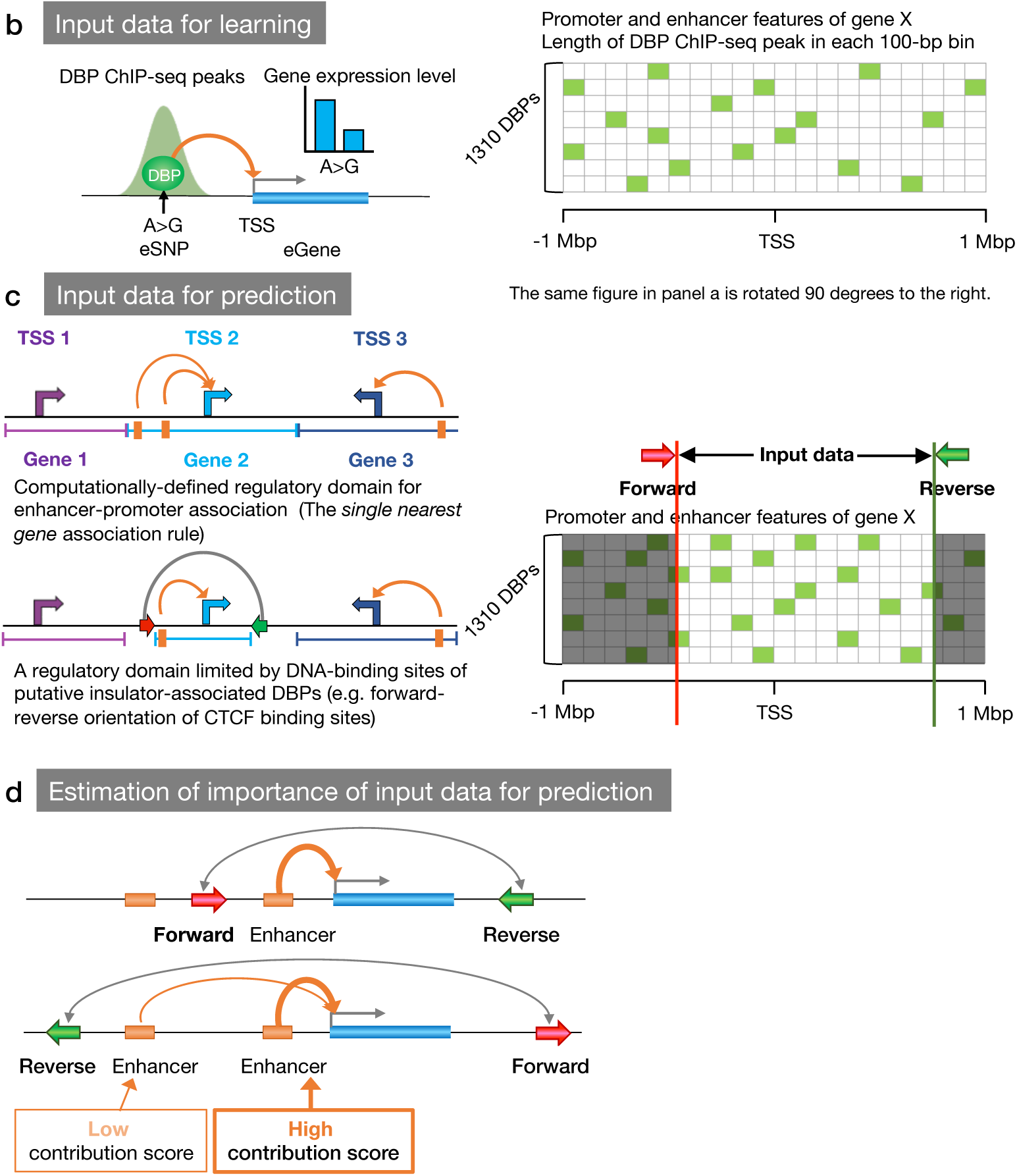
Overview of the prediction pipeline for insulator-associated DBPs. **a** We employed a machine learning method, DEcode, with modifications. Gene expression levels in human cells were predicted using the genomic locations of DBP binding sites at promoters and putative enhancers as input features. **b** In the machine learning phase, a DBP bound to a putative enhancer overlapping an eSNP (expression quantitative trait locus SNP) was linked to a gene (eGene) potentially regulated by the DBP, based on eQTL data. For each gene, DBP ChIP-seq peak locations were binned (green boxes) and summarized in a matrix as input: Promoters were represented by 30 bins of 100 bp each spanning the region from -2 kbp to +1 kbp relative to the TSS. Enhancer–gene regulatory regions were covered by 200 bins of 10 kbp each, spanning -1 Mbp to +1 Mbp from the TSS, excluding the promoter. **c** For prediction of putative insulator-associated DBPs, a DBP bound to a putative enhancer was associated with a gene based on a computationally defined regulatory domain, independent of eQTL data. The single nearest gene association rule defines a regulatory domain as the region between the TSS of a gene and the midpoints to the TSS of the nearest upstream and downstream genes. The TSSs and regulatory domains for each gene are indicated with purple, light blue, or blue arrows and bracketed lines, respectively. The regulatory domain for enhancer–promoter associations was further constrained by DNA-binding sites (red and green arrows) of putative insulator-associated DBPs. For each gene, DBP ChIP-seq peak locations were binned (green boxes in non-gray regions) and summarized in a matrix as input data. **d** Estimation of the directional effect of DNA-binding sites (red and green arrows) of putative insulator-associated DBPs on gene expression prediction. A DBP whose binding (orange box) is highly informative for gene expression prediction receives a high contribution score.

**Fig. 3.**
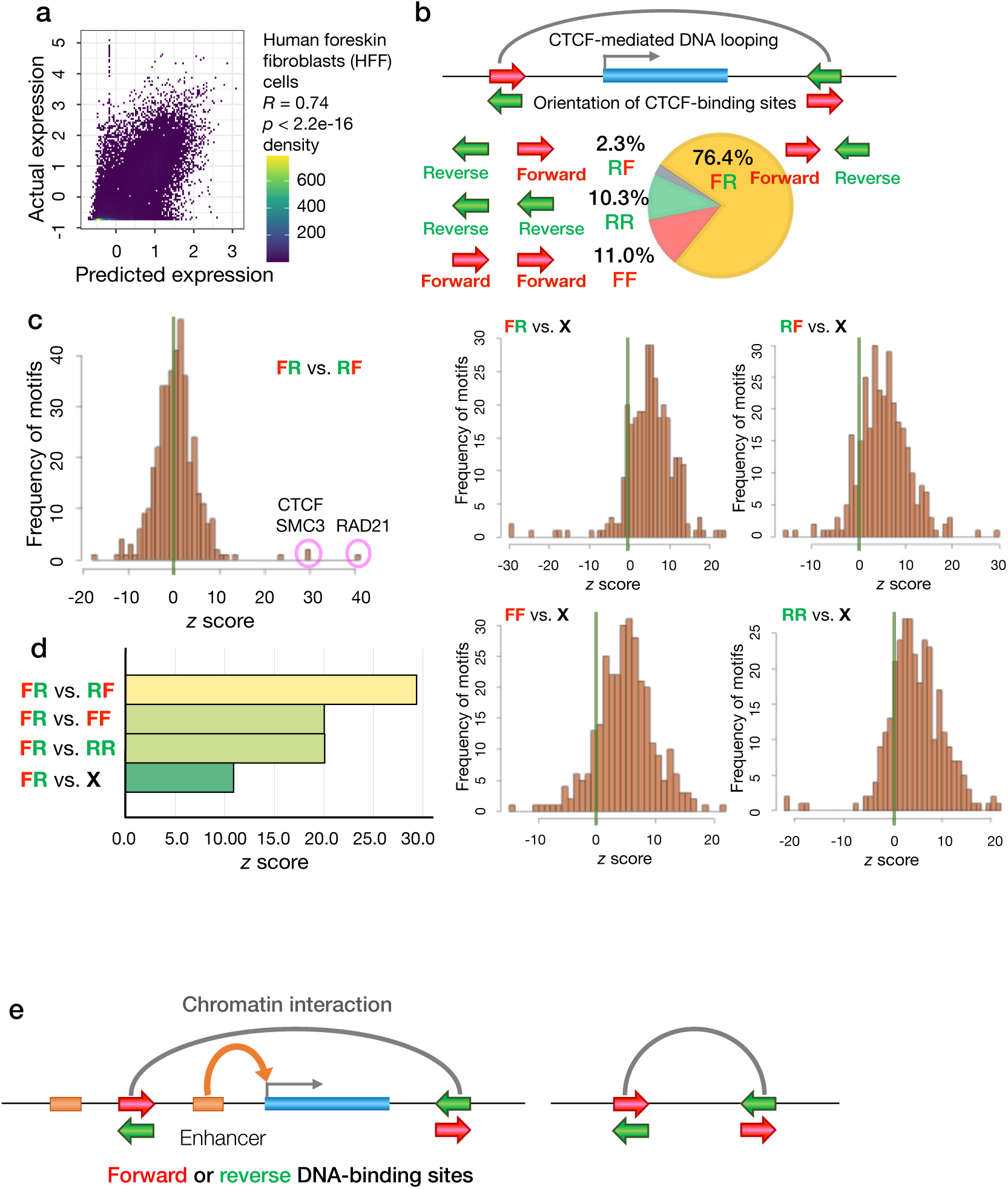
Estimation of the effect of directional regulatory motifs on gene expression. **a** The predicted log₂-transcripts per million (log₂-TPM) values of genes were compared with actual expression levels in human foreskin fibroblast (HFF) cells using Spearman’s rank correlation coefficient. **b** The forward–reverse (FR) orientation of CTCF-binding sites is predominant (76.4%) at chromatin interaction sites, while forward–forward (FF), reverse–reverse (RR), and reverse–forward (RF) orientations are less frequent (11.0%, 10.3%, and 2.3%, respectively) (Guo Y et al., Cell 2015) ^3^. **c** Distribution of z-scores from the Mann–Whitney U test for DeepLIFT scores comparing FR and RF orientations of protein DNA-binding sites. The right panel displays the distribution of z-scores comparing non-directional (X) sites with directional orientations (FR, RF, FF, or RR). Results are shown according to Selection Criteria S2 in Supplementary Fig. 1 and Note. **d** *Z*-scores from the Mann–Whitney U test for DeepLIFT scores in pairwise comparisons of CTCF-binding sites with four different orientations (FR, RF, FF, RR) and non-directional (X) sites. The differences in *z*-scores mirrored the percentages of CTCF-binding site orientations at chromatin interaction sites. **e** The DNA-binding sites of insulator-associated proteins were identified at both sites or at either site within pairs of long-range chromatin interactions.

To build a robust foundation for regulatory modeling, we comprehensively identified binding motif sequences for 715 out of 1,310 DBPs using ChIP-seq peaks from the Gene Transcription Regulation Database (GTRD, v20.06 release) and the Protein Interaction Quantitation (PIQ) tool, which integrates DNase-seq data and a curated library of motifs from TRANSFAC (2019.2), JASPAR (2018), UniPROBE (2018), high-throughput SELEX, ENCODE ChIP-seq–derived motifs, and HOCOMOCO (versions 9 and 11) ^1718–23^. Only motif occurrences overlapping ChIP-seq peaks for each DBP were retained, thereby minimizing false positives and enabling the high-resolution prediction of DBP binding in open chromatin regions.

We next examined how the directionality and genomic position of insulator-associated DBP binding sites affect gene expression prediction. Using DeepLIFT, which quantifies each feature’s contribution to the model by measuring its deviation from a reference, we demonstrated that both the orientation and the position of DBP binding sites have substantial regulatory effects ^24^. Directional differences in putative insulator-associated DBP sites emerged as critical determinants of EPI strength and gene expression outcomes (**Fig. 2d**). This systematic approach allowed us to dissect how the regulatory effects of insulator-associated DBPs depend on both their chromatin context and strand orientation.

In particular, we found that CTCF binding site orientation is a key determinant of its regulatory impact on gene expression. CTCF sites in the forward–reverse (FR, convergent) orientation—accounting for 76.4% of distal CTCF-mediated chromatin interaction sites identified by ChIA-PET (**Fig. 3b**)—exhibited significantly different DeepLIFT score distributions compared to the reverse–forward (RF) orientation (Mann–Whitney U test, FDR-adjusted *p* < 0.05; *p* < 10⁻³; **Fig. 3c**, **d**; **Table S1**) ^3^. This difference was robustly detected when comparing FR and RF CTCF-binding sites located in predicted enhancer and open chromatin regions both upstream and downstream of TSSs (**Fig. 2c**). Further analyses using alternative DNA-binding site selection criteria supported these findings (**Supplementary** Fig. 1 and **Note**), demonstrating that our approach captures the regulatory significance of CTCF binding site orientation.

Extending this analysis, we observed that directional orientation is a widespread and biologically meaningful predictor of gene expression among putative insulator-associated DBPs. DBPs such as RAD21, SMC3, and CTCF exhibited strong differences in DeepLIFT score distributions between FR and RF orientations (**Fig. 3c**, **Table S1**), and this effect was not limited to these proteins. Among 228 DBPs with at least 1,000 binding sites in HFF cells, 92—including ZNF143 and YY1—showed significant differences between sites with different orientations (e.g., FR vs. RF), while 79—including YY1—also showed differences between same-orientation sites (e.g., FF vs. RR) (**Fig. 3c**, **Fig. S2b**, **Table S1**, sheet **“**DBP List 1A”). Comparisons between directional (FR, RF, FF, RR) and non-directional (X) binding sites consistently produced higher *z*-scores for the former, highlighting the greater contribution of directional binding sites to gene expression prediction. The most pronounced effects were observed for established insulator- and chromatin-associated proteins, underlining the biological importance and sensitivity of our model. For increased stringency, we restricted the analysis to DBPs with *p* < 10⁻³, confirming all such cases overlapped with those defined by the broader FDR-adjusted *p* < 0.05 threshold (**Table S1**). These findings prompted us to systematically classify insulator-associated DBPs by their directional or non-directional binding patterns for further analysis.

### Identification and classification of insulator-associated DBPs

Insulator-associated DBPs demonstrate widespread and significant orientation-dependent as well as orientation-independent DNA-binding preferences, reflecting their diverse roles in chromatin regulation. Our analysis revealed that established insulator DBPs—including CTCF, RAD21, and SMC3—exhibit a predominant forward–reverse (FR) orientation in their binding site distributions. Pairwise comparisons of DeepLIFT scores across four orientations (FR, RF, FF, RR) and non-directional sites showed significant differences for 23 DBPs with different orientations and for 17 DBPs with the same orientation. Additionally, non-directional binding sites showed significant score differences for six DBPs (*p* < 10⁻³; **Table 1** and **Table S2**, sheet “DBP List 1A”; FDR-adjusted *p* < 0.05; **Table S3**, “DBP List 1A”). Beyond these established factors, we identified several novel DBPs that displayed both directional and non-directional binding preferences, expanding the pool of proteins potentially involved in insulator function.

**Table 1.**
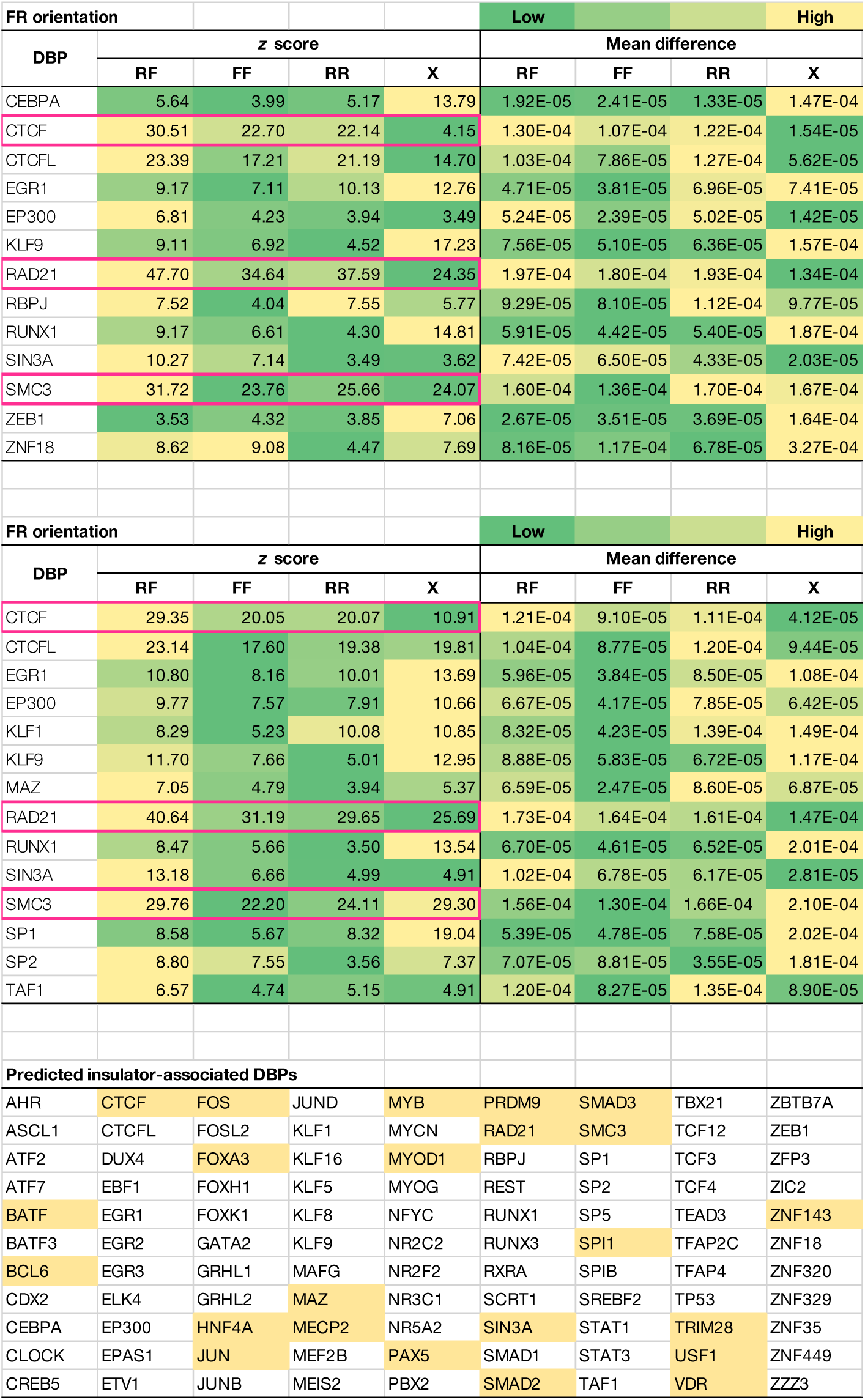
Predicted insulator-associated DNA-binding proteins (DBPs) with directional or nondirectional binding site preferences.

Importantly, these widespread orientation biases were supported by chromatin interaction data. Our deep learning–based predictions accurately recapitulated the characteristic orientation bias of CTCF and RAD21 DNA-binding sites at chromatin interaction sites, as validated by ChIA-PET data. Specifically, among CTCF binding sites overlapping distal interaction anchors, 76.4% displayed an FR orientation, with substantially lower frequencies of FF (11.0%), RR (10.3%), and RF (2.3%) (**Fig. 3b**; Guo Y et al., *Cell* 2015 ^3^, Fig. 4). DeepLIFT scores revealed that the difference between FR and RF orientations was significantly greater than that between FR and FF or RR (Mann–Whitney U test), consistent with experimental distributions (**Fig. 3c**, **d**). Similarly, RAD21 ChIA-PET data revealed 76.3% FR, 3.4% RF, 9.3% FF, and 11.1% RR orientations, and DeepLIFT scores for RAD21 binding sites closely mirrored these results (**Fig. 3c**; **Table 1**). These findings confirm that our framework robustly predicts the experimentally observed orientation biases of insulator-associated DBPs at chromatin interaction sites.

**Fig. 4.**
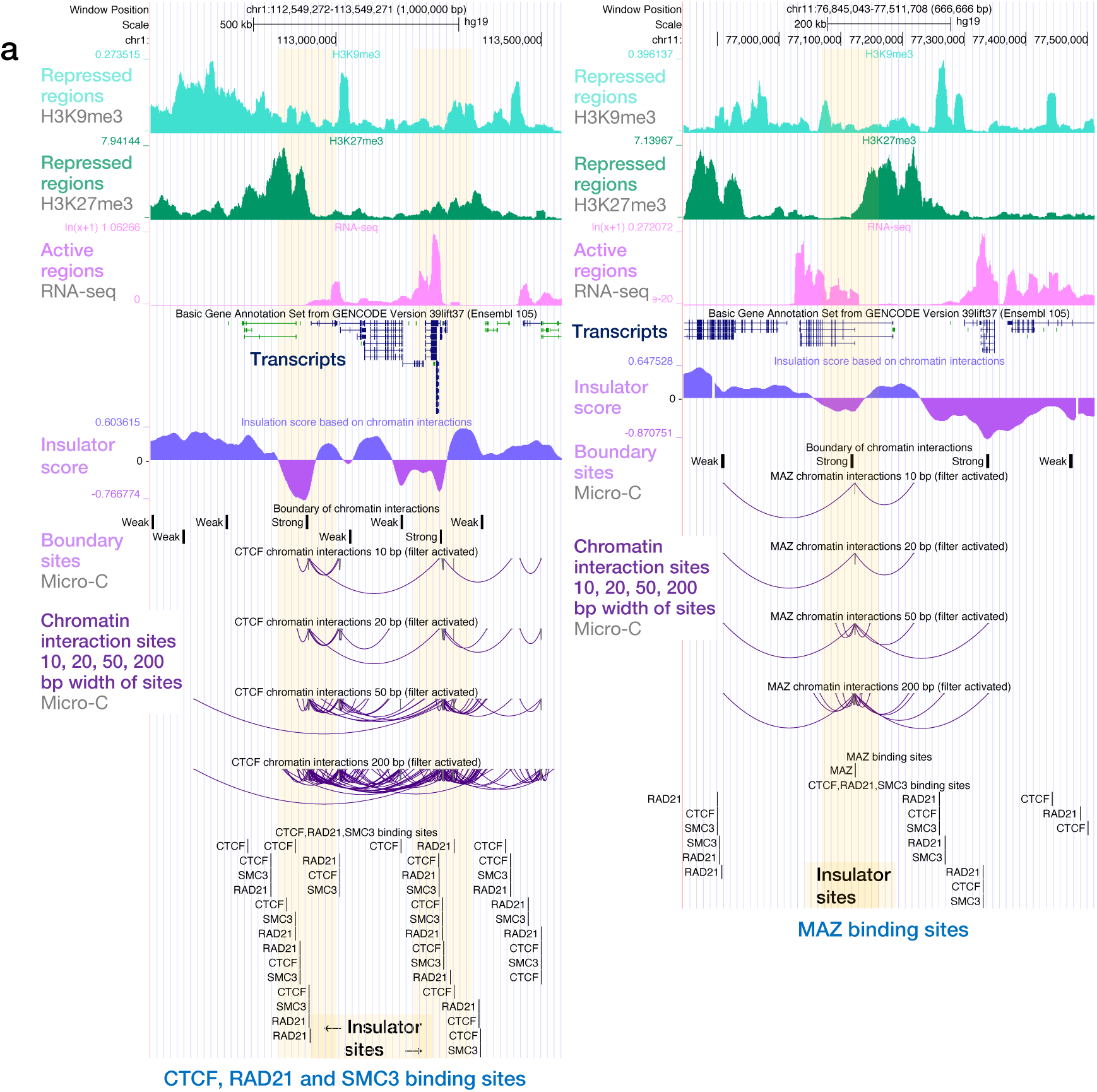
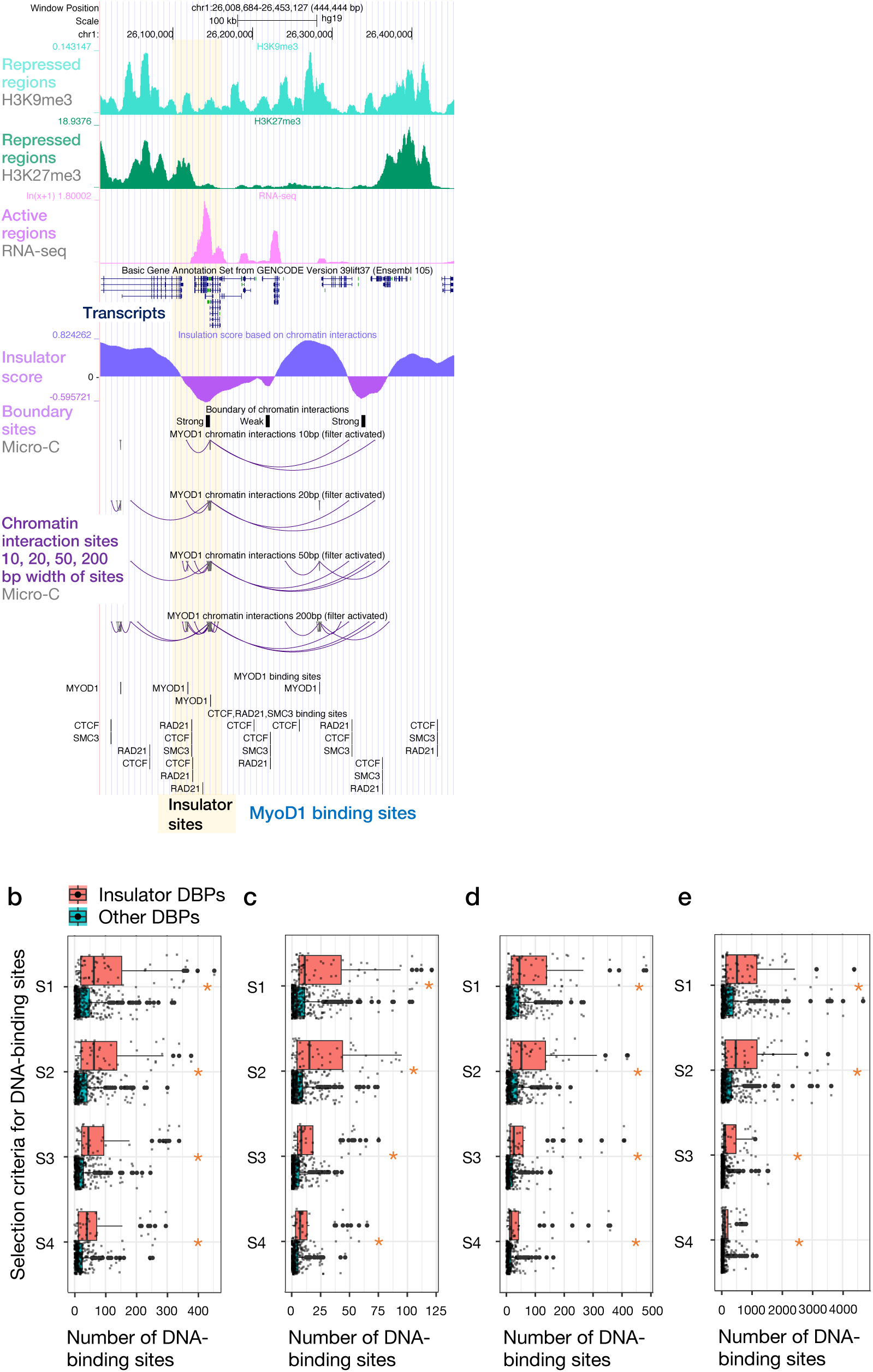
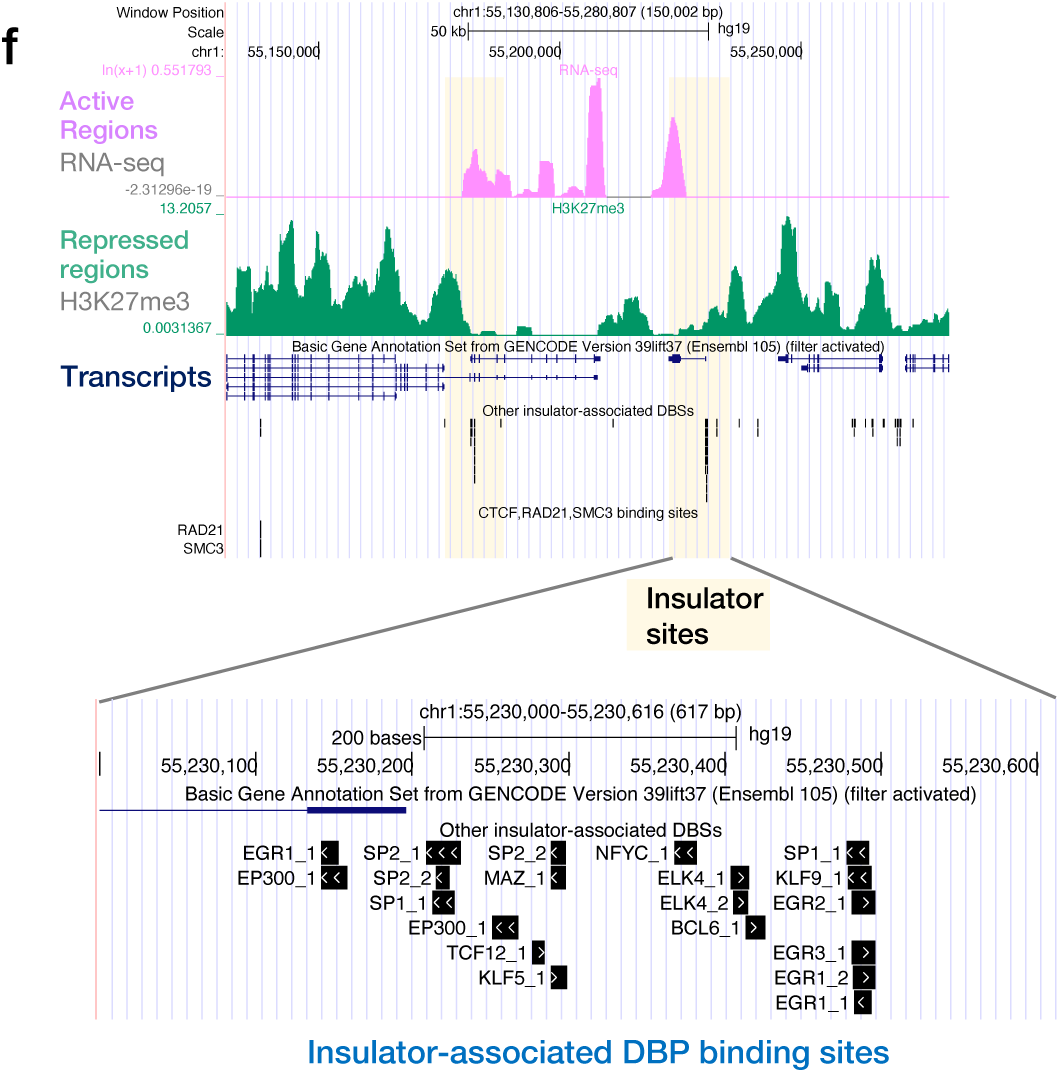
DNA-binding sites of predicted insulator-associated DBPs at potential insulator sites. **a** Distribution of histone modification marks H3K27me3 (green) and H3K9me3 (turquoise), along with transcript levels (pink), in the upstream and downstream regions of a potential insulator site (light orange). These histone modification marks are characteristic of transcriptionally repressed regions. Insulator sites are positioned at the boundary between transcriptionally active and repressed regions (indicated by light orange vertical bars). Blue and green lines in the GENCODE track denote gene transcripts. Insulator scores and boundary regions were estimated from chromatin interaction data available via the 4D Nucleome Data Portal. Dark blue arcs represent chromatin interactions within 10 bp, 20 bp, 50 bp, and 200 bp of the binding sites of CTCF (upper left panel), MAZ (upper right panel), and MyoD1 (lower left panel), respectively. The binding sites of CTCF, RAD21, SMC3 (upper left), MAZ (upper right), and MyoD1 (lower left) are displayed as black bars. Analyses were performed using the Homo sapiens genome assembly GRCh37 (hg19) from the Genome Reference Consortium. **b** Number of DNA-binding sites of predicted insulator-associated DBPs where H3K27me3 signal intensities differed between upstream and downstream regions. * indicates *p* < 0.01 (Mann–Whitney U test) (Table 2). Boxplots show the median (centerline), interquartile range (boxes), whiskers (up to 1.5× IQR), and all outliers (points). All data points are shown. **c** Number of DNA-binding sites of predicted insulator-associated DBPs where both H3K27me3 signal intensities and transcript levels differed between upstream and downstream regions. **d** Number of DNA-binding sites of predicted insulator-associated DBPs located within boundary regions assigned by chromatin interaction data from human foreskin fibroblast (HFF) cells. **e** Number of DNA-binding sites of predicted insulator-associated DBPs overlapping within ±200 bp of splice sites. Insulator-associated DBP binding sites were found within this window significantly more frequently than for other DNA-binding proteins. **f** Cluster of DNA-binding sites of insulator-associated DBPs.

Applying stringent orientation criteria, we identified a comprehensive set of 97 insulator-associated DBPs that directly interact with DNA and exhibit either strong directional or non-directional binding preferences. Most of these DBPs exhibited clear directional biases, while only six—including FOS, FOSL2, JUN, JUNB, JUND (all with palindromic motifs), and SPI1 (their interacting partner)—were characterized by non-directional binding. Notably, 23 of these DBPs have been previously implicated in insulator activity, CTCF association, or chromatin regulatory functions affecting enhancer–promoter interactions (EPIs), with 13 of them directly interacting with CTCF (**Table S7**). These proteins are multifunctional, serving as transcriptional regulators, repressors, chromatin regulators, or cofactors for epigenetic modifier enzymes.

Collectively, our findings indicate that CTCF coordinates EPI regulation through a network of diverse DBPs and that both directional and non-directional binding preferences underlie the regulatory capacity of these proteins. The involvement of these multifunctional DBPs in chromatin architecture and long-range genomic interactions emphasizes the complex and modular organization of chromatin regulatory networks.

Extending these analyses to the spatial organization of chromatin, we found that insulator-associated DBPs are frequently co-localized at both ends of chromatin interaction sites, supporting their role in mediating long-range chromatin architecture. Across 30,369 chromatin interactions defined by 400-bp regions (separated by >5 kb and excluding TSS-proximal areas), DNA-binding sites for 44 (45%) and 97 (100%) insulator-associated DBPs were identified at both ends or at either end of an interaction pair, respectively. For 10,384 interactions defined by 200-bp regions, binding sites for 30 (31%) and 97 (100%) DBPs were detected at both or either end. These analyses, based on orientation-aware pairings (FR, RF, FF, RR) (**Fig. 3e**), likely underestimate the full extent of insulator DBP involvement due to incomplete coverage of chromatin interaction data ^25^. Canonical insulator proteins such as CTCF, RAD21, and SMC3 exhibited strong directional binding within these regions, consistent with ChIA-PET results. To further validate the predictions of candidate insulator-associated DBPs, we next sought independent lines of evidence—including histone modification signatures characteristic of insulator function—to corroborate their roles in chromatin boundary and architecture.

### Epigenomic validation of insulator activity

Epigenomic analyses provide independent support for the predicted regulatory roles of insulator-associated DBPs. We identified insulator-associated DBP binding sites by detecting asymmetry in H3K27me3 signal profiles across their upstream and downstream regions, reflecting boundaries between active and repressed chromatin domains. Specifically, loci where the mean H3K27me3 signal intensity differed by more than twofold between upstream and downstream regions, with the higher value exceeding the first quartile of the genome-wide distribution, were selected to ensure robust discrimination and to filter out low-intensity noise. This approach builds on prior observations that CTCF and USF1 occupy the edges of repressive chromatin domains (marked by H3K27me3) versus active regions (marked by H2AK5ac) ^2627^, making H3K27me3 a suitable marker for broad transcriptional repression.

DNA-binding sites of predicted insulator-associated DBPs were significantly enriched at chromatin boundaries defined by asymmetric H3K27me3 signals. Known insulator-associated factors, such as CTCF, RAD21, and SMC3, showed the highest number of such sites (**Table S4**). Quantitatively, the median and mean counts of binding sites with greater than twofold H3K27me3 signal differences between upstream and downstream regions were significantly higher for predicted insulator-associated DBPs than for other DBPs (Mann–Whitney U test, *p* < 0.05; **Table 2**, **Table S4**, **Fig. 4a**, **b**). These results demonstrate that insulator-associated DBPs preferentially occupy genomic positions flanked by distinct chromatin states, providing further evidence for their function in defining regulatory domains.

**Table 2.** Number of DNA-binding sites for predicted insulator-associated DBPs at insulator sites and splice sites.

|  |  | Selection Criteria |  | Median number of DNA sites | Mean number of DNA sites | Number of DBP | p value |
| --- | --- | --- | --- | --- | --- | --- | --- |
| Experimental data for identifying insulator sites | H3K27me3 | S1 | Insulator-associated DBPs | 66 | 115.66 | 58 | 2.35E-08 |
|  |  |  | Other DBPs | 16 | 38.14 | 354 |  |
|  |  | S2 | Insulator-associated DBPs | 65 | 102.00 | 50 | 3.33E-08 |
|  |  |  | Other DBPs | 14 | 31.73 | 346 |  |
|  |  | S3 | Insulator-associated DBPs | 45 | 93.43 | 40 | 3.20E-08 |
|  |  |  | Other DBPs | 12 | 25.98 | 298 |  |
|  |  | S4 | Insulator-associated DBPs | 40 | 69.65 | 37 | 1.13E-06 |
|  |  |  | Other DBPs | 11 | 22.45 | 274 |  |
|  | H3K27me3, Gene expression | S1 | Insulator-associated DBPs | 12 | 27.95 | 58 | 4.26E-06 |
|  |  |  | Other DBPs | 4 | 10.04 | 354 |  |
|  |  | S2 | Insulator-associated DBPs | 16 | 27.30 | 50 | 2.14E-06 |
|  |  |  | Other DBPs | 4 | 8.28 | 346 |  |
|  |  | S3 | Insulator-associated DBPs | 8 | 18.05 | 40 | 9.89E-06 |
|  |  |  | Other DBPs | 2 | 5.09 | 298 |  |
|  |  | S4 | Insulator-associated DBPs | 7 | 13.97 | 37 | 9.17E-05 |
|  |  |  | Other DBPs | 2 | 4.46 | 274 |  |
|  | Boundary of interactions | S1 | Insulator-associated DBPs | 48 | 96.36 | 58 | 6.43E-07 |
|  |  |  | Other DBPs | 14 | 30.88 | 354 |  |
|  |  | S2 | Insulator-associated DBPs | 54 | 91.58 | 50 | 5.39E-07 |
|  |  |  | Other DBPs | 13 | 26.70 | 346 |  |
|  |  | S3 | Insulator-associated DBPs | 26 | 74.33 | 40 | 4.57E-07 |
|  |  |  | Other DBPs | 7 | 15.17 | 298 |  |
|  |  | S4 | Insulator-associated DBPs | 16 | 58.08 | 37 | 1.27E-05 |
|  |  |  | Other DBPs | 7 | 13.75 | 274 |  |
| Overlap with splice sites |  | S1 | Insulator-associated DBPs | 535 | 774.45 | 58 | 1.02E-06 |
|  |  |  | Other DBPs | 132 | 350.11 | 354 |  |
|  |  | S2 | Insulator-associated DBPs | 472 | 748.72 | 50 | 6.28E-06 |
|  |  |  | Other DBPs | 122 | 290.21 | 346 |  |
|  |  | S3 | Insulator-associated DBPs | 120 | 305.7 | 40 | 2.13E-07 |
|  |  |  | Other DBPs | 35 | 99.23 | 298 |  |
|  |  | S4 | Insulator-associated DBPs | 103 | 198.24 | 37 | 1.45E-05 |
|  |  |  | Other DBPs | 30 | 81.01 | 274 |  |

Furthermore, predicted insulator-associated DBPs are localized to transcriptional transition regions and chromatin boundaries. Analysis of RNA-seq data from HFF cells showed that these DBPs are significantly enriched at sites where transcript signal intensities differ by more than 1.5-fold between upstream and downstream regions, with the higher value exceeding the first quartile of the transcriptome-wide distribution (**Table 2**, **Table S4**, **Fig. 4a**, **c**). Additionally, DNA-binding sites of these DBPs more frequently overlapped with chromatin boundary regions, as defined by insulation scores from chromatin interaction data ^28^, than those of non-insulator DBPs (**Table 2**, **Fig. 4d**). These results underscore the role of insulator-associated DBPs in partitioning active and inactive chromatin domains and functionally positioning at regulatory boundaries. To explore the fine-scale genomic context of these sites, we next conducted genome browser inspections.

Genomic inspection using the UCSC Genome Browser revealed that predicted insulator-associated DBPs consistently occupy sites with insulator-like features, including chromatin boundary regions identified by Micro-C data and transitional regions between H3K27me3-marked and transcriptionally active domains. Many of these binding sites were located at the edges of H3K27me3-enriched domains and were frequently distant from canonical insulator proteins such as CTCF, RAD21, and SMC3 (**Fig. 4a**, **Fig. S3**). For each DBP, up to three representative binding sites in insulator-associated contexts were manually selected (**Fig. S3a**, **S3f**). Notably, MyoD1 was found at a chromatin boundary near the SELENON (SEPN1) gene—both involved in skeletal muscle differentiation ^29^ —suggesting a possible regulatory role for MyoD1 via insulator mechanisms (**Fig. 4a**).

Expanding this analysis to alternative repressive histone modifications, insulator-associated DBPs were also enriched at the edges of H3K9me3-enriched domains (**Fig. S3b**), supporting their ability to define chromatin boundaries beyond those marked by H3K27me3. The distribution of H3K9me3 differed from that of H3K27me3 in certain regions, highlighting the context-dependent nature of insulator activity. Additionally, predicted DBP binding sites frequently formed spatial clusters at insulator regions and were often associated with chromatin interaction anchors (**Fig. 4f**, **Fig. S3c**, **S3d**). This clustering aligns with the role of architectural proteins in organizing TAD boundaries, while discrete insulator motifs contribute to intra-TAD looping ^30^. To further clarify the relationship between these epigenetically defined insulator sites and chromatin architecture, we compared their locations with chromatin boundaries identified by chromatin interaction data.

Comparative analyses showed that histone- and transcription-defined insulator sites often coincided with, or were located near, chromatin boundaries identified by interaction data. In HFF cells, 29% of H3K27me3-based insulator sites (type 1) and 37% of H3K27me3 plus transcription-based sites (type 2) overlapped with 25-kb boundary regions, whereas only 6% and 7% overlapped with 5-kb boundaries, respectively, reflecting differences in boundary resolution (**Fig. 4f**, **Fig. S3**). Genome-wide, DNA-binding sites of the 97 predicted insulator-associated DBPs were found at type 1 insulator sites assigned to 3,170 genes, at type 2 sites assigned to 1,044 genes, and at chromatin interaction–defined boundaries assigned to 6,275 genes. Among these, 1,212 genes (38%) harbored both type 1 insulator and boundary sites, and 389 genes (37%) contained both type 2 insulator and boundary sites. These data indicate that while chromatin interaction boundaries and epigenetically or transcriptionally defined insulator sites are not completely overlapping, they frequently target overlapping gene sets and are co-located within the same regulatory landscapes, supporting their complementary roles in chromatin compartmentalization.

### Independent and cooperative clustering of insulator-associated DBPs at chromatin regulatory landmarks

Predicted insulator-associated DBPs are enriched at critical transcript regulatory regions, such as transcription start sites (TSSs), transcription termination sites (TTSs), and splice sites, suggesting potential roles in shaping transcript structure. Statistical analyses showed that these DBPs are significantly more abundant at splice sites than other DBPs (Mann–Whitney U test, *p* < 0.05; **Fig. 4e**, **Table 2**), with pronounced enrichment at the first and last introns compared to internal introns (**Table 2**, **Table S9**).

This finding is consistent with prior reports that variation in TSS/TTS usage is a key driver of tissue-specific isoform diversity ^31^. Examination of **Fig. 4a** and **Fig. S3** further revealed clustering of multiple insulator-associated DBP binding sites within single loci, suggesting potential cooperative functions in establishing insulator elements. Notably, transcribed regions with overlapping H3K27me3 enrichment and alternative transcriptional repression contained DBP sites between active and repressed domains (**Fig. 5**, **Fig. S3e**), indicating that insulator-associated DBPs may delineate boundaries between transcriptional states, thereby influencing selection of TSSs, TTSs, and exons in a context-dependent manner.

**Fig. 5.**
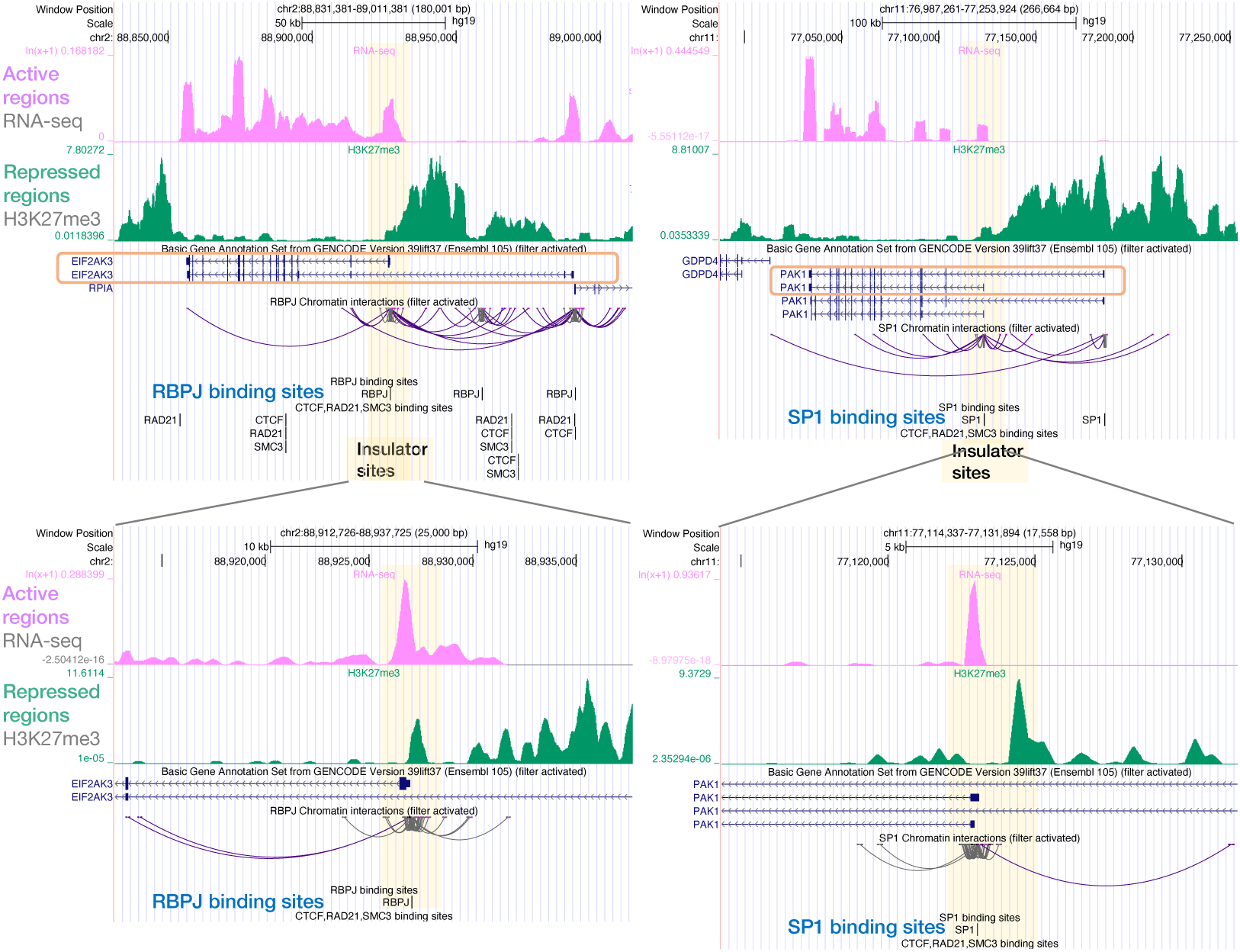
Insulator-associated DBPs potentially regulate alternative transcription. Distribution of the H3K27me3 histone modification mark (green) and transcript levels (pink) is displayed. The H3K27me3 mark is characteristic of transcriptionally repressed regions. Insulator sites are positioned at the boundaries between transcriptionally active and repressed regions (indicated by light orange vertical bars). Blue lines in the GENCODE track indicate gene transcripts. Chromatin interactions overlapping a 200-bp window centered on the binding sites of RBPJ (left panel) and SP1 (right panel) are shown as dark blue arcs. The binding sites of RBPJ (left), SP1 (right), and CTCF, RAD21, SMC3 (both panels) are shown as black bars. The DNA-binding sites of insulator-associated DBPs were frequently observed near transcription start and termination sites. These results suggest that insulator-associated DBPs may differentially regulate transcribed regions that are repressed in alternative transcripts.

Chromatin interaction–associated DBPs, including several predicted insulator factors, are strongly enriched near chromatin interaction centers, emphasizing their potential in supporting chromatin looping. High-resolution Micro-C data from HFF cells identified DBP binding sites located within ±500 bp of interaction anchors ^32^. Canonical organizers CTCF, RAD21, and SMC3 exhibited sharp enrichment at these sites (**Fig. 6a**‒**c**), while YY1 and ZNF143 displayed broader or distinct distributions (**Fig. 6d**, **e**), and MAZ—a DBP implicated in CTCF-independent insulator activity ^33^— demonstrated a unique peak (**Fig. 6f**). Even after removing binding sites overlapping CTCF, RAD21, and SMC3, other predicted insulator-associated DBPs remained consistently enriched at interaction centers, supporting their independent contributions to 3D chromatin structure.

**Fig. 6.**
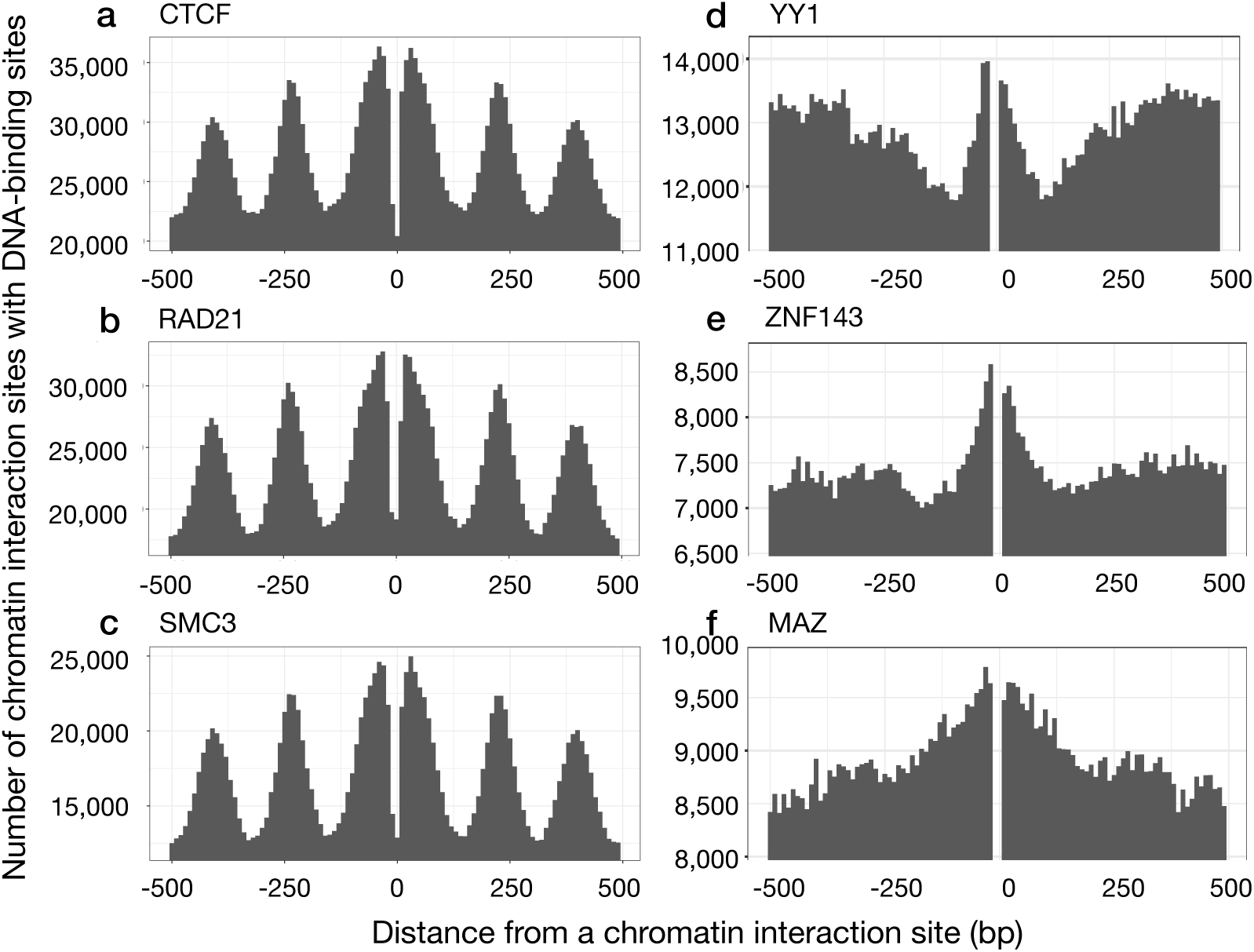
Distribution of DNA-binding sites around chromatin interaction sites. a-f The x-axis represents the distance of a DNA-binding site from the center of a chromatin interaction site, while the y-axis indicates the number of chromatin interaction sites with DNA-binding sites located within a ±500 bp region. For the analyses of YY1, ZNF143, and MAZ, DNA-binding sites overlapping those of CTCF, RAD21, and SMC3 were excluded.

Analysis of 228 DBPs with over 1,000 binding sites in HFF cells revealed that nearly 90% exhibited distinct enrichment near chromatin interaction sites (**Table S8**, **Fig. S4**), displaying sharp, broad, or canonical CTCF-, RAD21-, or SMC3-like peak profiles. Although DBPs such as MeCP2 and PRDM9, with fewer than 1,000 binding sites, did not exhibit this pattern, these findings demonstrate that most DBPs—including many predicted insulator-associated proteins—are spatially positioned to participate in genome architecture regulation.

Co-localization analyses revealed that most predicted insulator-associated DBPs are spatially segregated from canonical insulator proteins. Using 10-bp bins across ±500 bp windows centered at interaction anchors and ChIP-seq–validated binding sites, we identified 245 DBP pairs (involving 135 DBPs) with ≥30% co-localization (Selection Criteria S1 in Fig. S1; **Table S5**). CTCF was highly co-localized with RAD21 (39%) and SMC3 (81%), whereas YY1/ZNF143 had much lower rates (<6%) with canonical factors (**Table 3**). Some DBPs (CTCFL, RXRA, PPARG, NEUROD1, ZBTB7A) exhibited moderate co-localization (>10%), but the majority of predicted insulator-associated DBPs showed minimal overlap (<10%), suggesting functional independence. Spatial clustering of multiple DBP binding sites was also observed at putative insulator loci (**Fig. S3d**), and co-localization was especially strong at 5-kb boundary-associated sites (197 pairs, 101 DBPs, ≥30% overlap; **Table S6**), suggesting that these DBPs, like CTCF and cohesin, may assemble cooperatively at boundaries to mediate domain partitioning.

**Table 3.** Pairs of DBPs that colocalize at equivalent distances from chromatin interaction sites.

| DBP1 | DBP2 | Distance to start (bp) | Distance to end (bp) | # chromatin interactions with DBP1 sites | # chromatin interactions with DBP2 sites | # chromatin interactions with DBP1 and 2 sites | Ratio (DBP1 and 2) / DBP1 | Ratio (DBP1 and 2) / DBP2 | Max ratio (DBP1 or DBP2) |
| --- | --- | --- | --- | --- | --- | --- | --- | --- | --- |
| CTCF | CTCFL | 30 | 39 | 36229 | 27159 | 19743 | 0.54 | 0.73 | 0.73 |
| CTCF | PPARG | 30 | 39 | 36229 | 2577 | 528 | 0.01 | 0.20 | 0.20 |
| CTCF | RAD21 | -10 | -1 | 23108 | 19746 | 7661 | 0.33 | 0.39 | 0.39 |
| CTCF | RXRA | 220 | 229 | 33327 | 13204 | 9102 | 0.27 | 0.69 | 0.69 |
| CTCF | SMC3 | -30 | -21 | 35558 | 24371 | 19701 | 0.55 | 0.81 | 0.81 |
| CTCF | ZBTB7A | 30 | 39 | 36229 | 5190 | 1539 | 0.04 | 0.30 | 0.30 |
| YY1 | CTCF | -50 | -41 | 12717 | 35247 | 291 | 0.02 | 0.01 | 0.02 |
| YY1 | RAD21 | -30 | -21 | 13931 | 32793 | 203 | 0.01 | 0.01 | 0.01 |
| ZNF143 | CTCF | 20 | 29 | 8993 | 35212 | 430 | 0.05 | 0.01 | 0.05 |
| ZNF143 | RAD21 | -30 | -21 | 9064 | 32793 | 500 | 0.06 | 0.02 | 0.06 |
| BACH2 | ATF3 | 290 | 299 | 2496 | 20052 | 634 | 0.25 | 0.03 | 0.25 |
| BACH2 | BACH1 | -280 | -271 | 2471 | 2863 | 730 | 0.30 | 0.25 | 0.30 |
| BACH2 | BATF | -250 | -241 | 2709 | 5924 | 486 | 0.18 | 0.08 | 0.18 |
| BACH2 | FOS | -240 | -231 | 2769 | 39887 | 884 | 0.32 | 0.02 | 0.32 |
| BACH2 | FOSB | -70 | -61 | 2774 | 16108 | 602 | 0.22 | 0.04 | 0.22 |
| BACH2 | FOSL1 | 240 | 249 | 2589 | 19417 | 687 | 0.27 | 0.04 | 0.27 |
| BACH2 | FOSL2 | 260 | 269 | 2649 | 37682 | 837 | 0.32 | 0.02 | 0.32 |
| BACH2 | JUN | 260 | 269 | 2649 | 36745 | 783 | 0.30 | 0.02 | 0.30 |
| BACH2 | JUNB | 440 | 449 | 2519 | 23889 | 535 | 0.21 | 0.02 | 0.21 |
| BACH2 | JUND | 260 | 269 | 2649 | 40876 | 925 | 0.35 | 0.02 | 0.35 |
| BACH2 | MAF | 430 | 439 | 2500 | 1520 | 367 | 0.15 | 0.24 | 0.24 |
| BACH2 | MAFF | -220 | -211 | 2676 | 3302 | 295 | 0.11 | 0.09 | 0.11 |
| BACH2 | MAFG | 180 | 189 | 2514 | 2738 | 385 | 0.15 | 0.14 | 0.15 |
| BACH2 | MAFK | -30 | -21 | 2726 | 7561 | 896 | 0.33 | 0.12 | 0.33 |
| BACH2 | MYC | 90 | 99 | 2674 | 6900 | 417 | 0.16 | 0.06 | 0.16 |
| BACH2 | NFE2 | -10 | -1 | 2399 | 8289 | 1284 | 0.54 | 0.15 | 0.54 |
| BACH2 | NFE2L2 | 440 | 449 | 2519 | 4908 | 928 | 0.37 | 0.19 | 0.37 |

Overlap analysis with ChIP-seq data showed that >98% of CTCF, RAD21, and SMC3 binding sites overlapped their own ChIP-seq peaks under all conditions, thereby validating site definitions. In contrast, nearly all other predicted insulator-associated DBPs showed <50% overlap with CTCF, RAD21, or SMC3 peaks, even when data from various cell types were considered. This low co-occupancy supports the notion that predicted DBPs function largely independently of classical insulator complexes and may establish chromatin boundaries by distinct mechanisms.

Interestingly, a distinct group of predicted insulator-associated DBPs—including BACH2, FOS, ATF3, NFE2, and MAFK—exhibited binding site distributions around chromatin interaction centers similar to canonical insulator proteins, but had less than 1% direct spatial overlap within 10-bp bins (**Fig. S4c**). However, these factors showed strong mutual co-localization (**Table 3**), suggesting the existence of parallel, functionally distinct DBP clusters that independently regulate chromatin structure ^343536^. Thus, multiple DBP clusters with similar positional preferences may exert parallel effects on chromatin remodeling or loop accessibility via independent regulatory pathways.

## Discussion

Here, we present the first, to our knowledge, computational method and genome-wide analysis for the detection of insulator-associated DBPs. Transcriptional regulation mediated by the binding of DBPs at sites located far from TSSs was modeled using our deep learning approach based on eQTL data. Through this framework, the effects of DNA-binding sites of insulator-associated DBPs on EPIs and gene expression were estimated, and insulator-associated DBPs were predicted. We further identified DBPs, including known insulator-associated proteins, that exhibit insulator-associated functions and regulate transcription, thereby affecting EPIs as reported in previous studies. The DNA-binding sites of the predicted insulator-associated DBPs demonstrated orientation bias; unexpectedly, the significant differences in the orientation bias of CTCF DNA-binding sites based on DeepLIFT scores (contribution scores) were found to be consistent with the percent differences in the orientation of CTCF DNA-binding sites at CTCF-mediated chromatin interaction sites. Furthermore, statistical testing of the contribution scores revealed that the directional binding sites of DBPs contributed more significantly to the prediction of gene expression levels than the non-directional sites. The DNA-binding sites of the predicted insulator-associated DBPs were located at potential insulator sites more frequently than those of other DBPs, as indicated by differences in H3K27me3 signal intensities and transcript levels, and by their overlap with boundary sites identified by chromatin interactions. We further observed that only a small percentage of the DNA-binding sites of insulator-associated DBPs co-localized with the DNA-binding sites of known insulator-associated proteins such as CTCF, RAD21, and SMC3 around chromatin interaction sites. Finally, we discovered that DBPs other than the well-established insulator proteins, as well as the directionality of their DNA-binding sites, are associated with insulator function.

Our deep learning analysis accurately recapitulated the percent differences in the FR, FF, RR, and RF orientations of CTCF DNA-binding sites at chromatin interaction sites, by incorporating the genomic locations of DNA-binding sites for putative insulator-associated DBPs. The other predicted insulator-associated DBPs also demonstrated orientation bias in their DNA-binding sites. These results suggest that the directional bias of DNA-binding sites of insulator-associated DBPs may contribute to insulator function and chromatin regulation through structural interactions among DBPs, other proteins, DNA, and RNA. For example, the N-terminal amino acids of CTCF have been shown to interact with RAD21 in chromatin loops ^8–10^. Cohesin complexes that interact with convergent CTCF sites, that is, the N-terminus of CTCF, might be protected from WAPL, whereas those that interact with divergent CTCF sites, i.e., the C-terminus of CTCF, may not be protected from WAPL, which could release cohesin from chromatin and thus disrupt cohesin-mediated chromatin loops ^37^. As another example, although sequence-based predictions of EPIs underestimated actual interactions, an activity-by-contact (ABC) approach, based on a simple distance model, performed well, suggesting that factors in addition to sequence influence EPIs ^38,39^. Furthermore, CTCF-mediated chromatin looping induces the formation of transcriptional condensates and phase separation between the inner and outer regions of the loops, resulting in insulator function ^40–42^. Therefore, structural features that drive phase separation, rather than functional differences in DBPs themselves, may underlie insulator activity. CEBPA, among the identified insulator-associated DNA-binding proteins, has also been reported to participate in transcriptional condensates and phase separation ^43^. The FOXA1 pioneer factor functions as an initial chromatin-binding and chromatin-remodeling factor and has also been shown to form biomolecular condensates ^44^.

Other types of chromatin regulation are also likely to be related to the structural interactions of molecules. In the boundary-pairing (insulator-pairing) model, molecules bound to insulators physically pair with their partners, either head-to-head or head-to-tail, with different degrees of specificity at the termini of TADs in flies (**Fig. 7**). Although experiments have not clarified how partners locate each other, the mechanism is unlikely to require loop extrusion, as is presumed in CTCF- and cohesin-mediated chromatin looping. Both homologous and heterologous insulator–insulator pairing interactions are thought to be central to the architectural functions of insulators. Importantly, the nature of insulator–insulator interactions is orientation-dependent ^45–47^.

**Fig. 7.**
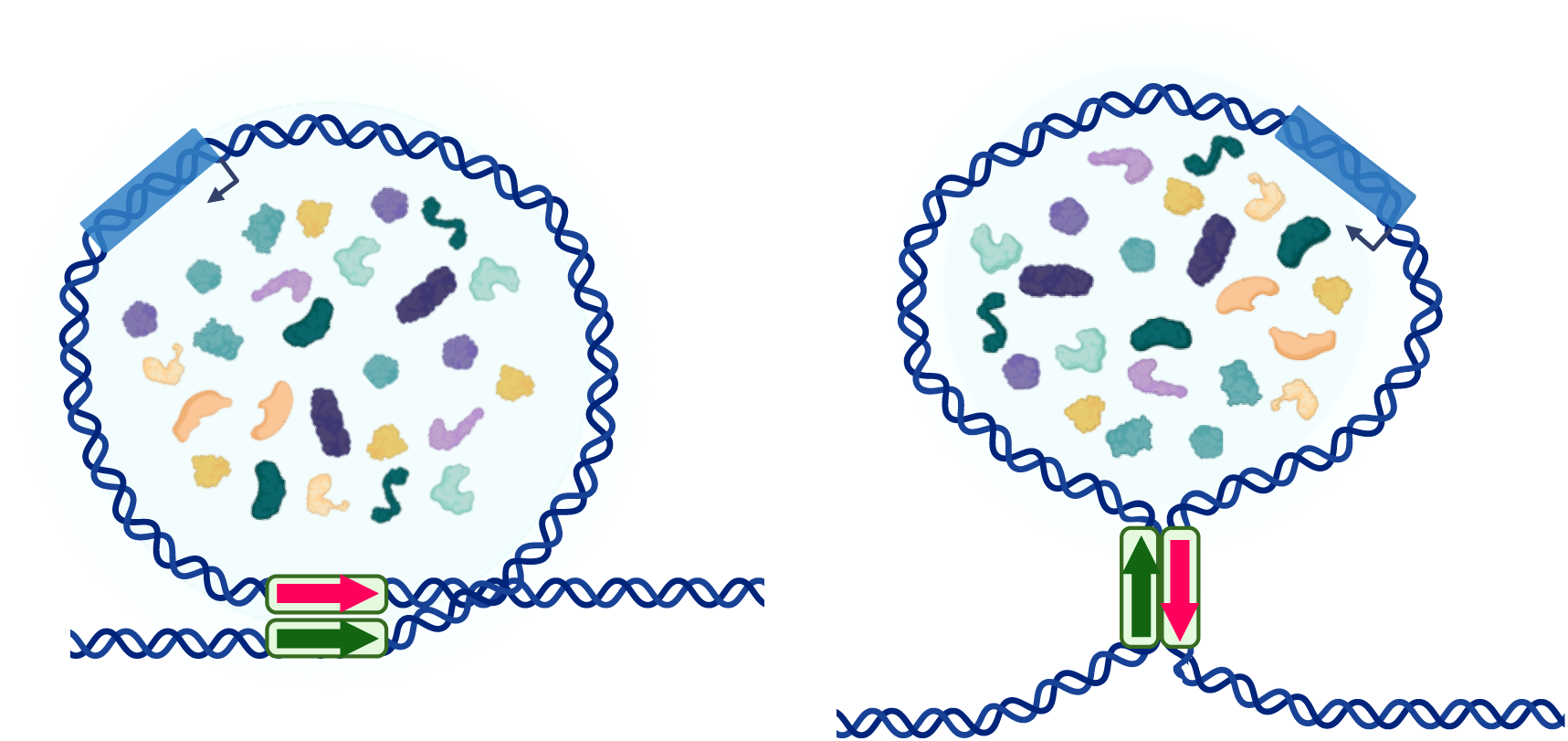
Boundary-pairing (insulator-pairing) model for the regulation of enhancer –promoter interactions. Molecules bound to insulators physically pair with their partners, either head-to-head or head-to-tail, with varying specificity at the termini of topologically associating domains (TADs) in flies ^45–47^. Although the precise mechanism by which partners locate each other remains unclear, it is unlikely to require loop extrusion. Homologous and heterologous insulator–insulator pairing interactions are central to the architectural functions of insulators, and these interactions are orientation-dependent. Homologous and heterologous interactions between insulator-associated DNA-binding proteins may similarly occur in humans. Chromatin looping can induce the formation of transcriptional condensates and phase separation, in addition to cohesin–CTCF-anchored chromatin looping ^40–42^. Figure created with BioRender.com.

Regional micro-capture (Micro-C) analyses have revealed that microcompartment interactions are largely independent of cohesin and transcription ^5^. Depletion of cohesin results in a subtle reduction in longer-range enhancer‒promoter interactions, while CTCF depletion can lead to rewiring of regulatory contacts ^48^. Another group reported that enhancer‒promoter interactions and transcription are largely maintained upon depletion of CTCF, cohesin, WAPL, or YY1; instead, cohesin depletion reduced transcription factor binding to chromatin, suggesting that cohesin may enable transcription factors to find and bind their targets more efficiently ^49^. Furthermore, loop extrusion is not essential for enhancer‒promoter interactions but contributes to their robustness, specificity, and the precise regulation of gene expressio ^48^. The ʻloop extrusion’ hypothesis is motivated by in vitro observations. Experiments in yeast, using cohesin variants unable to extrude DNA loops but capable of topologically entrapping DNA, suggested that in vivo chromatin loops can form independently of loop extrusion. Instead, transcription was shown to promote loop formation and act as an extrinsic motor that extends these loops and defines their final positions ^50^. Maz and MyoD1, among the identified insulator-associated DNA-binding proteins, have been shown to form loops without CTCF ^4,33,51^. Furthermore, protein post-translational modifications (PTMs) facilitate control over the molecular valency and strength of protein‒protein interactions ^52^. O-GlcNAcylation as a PTM has been reported to inhibit CTCF binding to chromatin ^53^. We found that the identified insulator-associated DNA-binding proteins tend to cluster at potential insulator sites (Fig. 4f and Supplementary Fig. 3c). These proteins may interact and actively regulate chromatin interactions, transcriptional condensation, and phase separation via PTMs. Our analyses will contribute to revealing novel functional and structural roles of DBPs in insulator functions, chromatin regulation, and phase separation, and to advancing their experimental analysis.

High-resolution chromatin interaction data from Micro-C assays demonstrated that most of the predicted insulator-associated DBPs exhibited peaks in the DNA-binding site distribution around chromatin interaction sites, suggesting that these DBPs are involved in chromatin interactions and highlighting the high resolution of Micro-C data. Base pair‒level resolution has been reported using Micro Capture-C ^32^. However, due to the high resolution of Micro-C assays, it is challenging to obtain genome-wide chromatin interaction data at a reasonable and accessible cost, whereas Micro Capture-C and Region Capture Micro-C assays focus on specific genomic regions to increase the coverage of chromatin interactions. The extent of chromatin interaction coverage varies depending on the specific Micro-C experimental design. In addition, chromatin interactions include types of interactions that are not associated with EPIs and insulator functions ^54–56^. Therefore, we analyzed and predicted insulator-associated DBPs independently of chromatin interaction data. Our deep learning analysis successfully learned enhancer–promoter interactions (EPIs) that affect gene expression levels and accurately recapitulated the percent differences in the FR, FF, RR, and RF orientations of CTCF DNA-binding sites at chromatin interaction sites.

When comparing insulator sites separated by active and repressive domains with boundary sites identified by chromatin interaction data, we defined insulator sites based on the total signal levels of H3K27me3 histone modification marks and transcript expression levels within a region. Some insulator sites may not display smooth, continuous distributions of histone marks and expression, which could account for differences between insulator and boundary sites. Another possible explanation is that boundary sites are more closely associated with topologically associating domains (TADs) of chromosomes than are insulator sites ^30^. Boundary sites represent regions identified based on the separation of numerous chromatin interactions. In contrast, we found that multiple DNA-binding sites of insulator-associated DNA-binding proteins were frequently clustered at insulator sites and were associated with distinct nested and focal chromatin interactions, as reported by Micro-C assays ^5^. These interactions may be transient and relatively weak, such as tissue-, cell type-, condition-, or lineage-specific interactions ^57^.

AI (Artificial Intelligence) is often regarded as a black box, since the reasons and causes of its predictions are difficult to determine. To address this issue, various tools and methods have been developed to enhance model interpretability, collectively referred to as Explainable AI. DeepLIFT is considered one such tool for Explainable AI. However, DeepLIFT does not provide the specific reasons or causal mechanisms underlying a prediction. Instead, it computes scores representing the contribution of each input feature to the prediction outcome. In this study, analysis of numerous contribution scores (i.e., DeepLIFT scores) for directional DNA-binding sites of putative insulator-associated DBPs in the prediction of transcript expression levels enabled accurate prediction of the percent differences in the FR, FF, RR, and RF orientations of CTCF DNA-binding sites at chromatin interaction sites, based on the contribution scores of DNA-binding sites in putative enhancer and open chromatin regions. It should be noted that some contribution scores may not be accurate, and predictions of insulator-associated DBPs or DNA-binding sites based on a small number of scores may be unreliable. To enable comparison of contribution scores calculated from different input–output pairs within our deep learning model, we normalized the contribution scores for each input‒output pair as previously described ^58^. The analysis of large sets of contribution scores will facilitate the estimation of the effects of other molecules and functions involved in gene regulation.

One of the next computational challenges will be to predict and refine the potential set of insulator-associated DNA-binding sites for multifunctional DBPs. To validate machine learning predictions and identify insulator-associated DNA-binding sites in the genome for subsequent experimental investigation, information on histone modifications such as H3K27me3 and transcript levels will be valuable, and other types of epigenomic data can also be utilized for this purpose. For example, as a reference for an application of this approach, Liu et al. focused on CTCF-binding sites and performed various assays in human and mouse cell cultures, as well as in mice, with the goal of applying these sites in gene therapy ^59^. The analysis of insulator-associated DBPs may ultimately contribute to the development of technologies for precise gene expression control. In summary, our novel computational method has identified insulator-associated DBPs—including previously characterized factors—and revealed that the directional bias of these DBPs is potentially involved in chromatin looping, phase separation, transcriptional condensate formation, and insulator function through the structural interactions of DBPs. These findings will contribute to a more comprehensive and global understanding of transcriptional regulation and gene expression.

## Methods

### Prediction of gene expression levels based on DNA-binding sites Gene expression data as output data for machine learning analysis

To predict DNA-binding sites that may also function as insulator sites in the human genome, we modified a previously developed deep learning method originally designed to predict transcript expression levels across tissues and cell types based on DNA-binding sites of DBPs in promoter regions and other genomic regions (Fig. 2a; Fig. S4) ^14^. To minimize the influence of cellular heterogeneity inherent in tissue samples, we utilized gene expression data from four distinct human cell types for which extensive experimental datasets are publicly available: human foreskin fibroblasts (HFFs; GSM941744), monocytes (Blueprint RNA-seq FPKM data, file ’C0010KB1.transcript_quantification.rsem_grape2_crg.GRCh38.20150622.results’, available at http://dcc.blueprint-epigenome.eu/#/files), human mammary epithelial cells (HMECs; ’wgEncodeCshlLongRnaSeqHmecCellPapAlnRep1.bam’, ENCODE project via UCSC Genome Browser http://genome.ucsc.edu/encode/), and neural progenitor cells (NPCs; GSM915326, ENCFF529SIO). FPKM (Fragments Per Kilobase of exon per Million mapped reads) values representing gene expression levels were retrieved or calculated based on GENCODE v19 annotations from the ENCODE portal ^60^ (https://www.encodeproject.org/) using the RSeQC tool ^61^ (http://rseqc.sourceforge.net/). Following the data processing pipeline described by Tasaki et al. ^14^, we calculated log₂-transformed FPKM values (adding 0.25 to avoid negative infinity), determined median log₂-FPKM values across the four cell types, and computed log₂-fold changes relative to the median expression for use as outputs in the deep learning model.

### DNA-binding sites as input data for machine learning analysis

To investigate the effect of insulator sites on EPIs and gene expression, the DNA-binding sites in putative enhancer and open chromatin regions located away from promoter regions were added as input data in the deep learning method. To identify the DNA-binding sites of DBPs in promoter and enhancer regions, ChIP-seq data of the DBPs were used and were updated to GTRD database release v20.06 (https://gtrd.biouml.org) ^62^. The binding site of a DBP regulating the expression level of each transcript was selected using eQTL data (GTEx_Analysis_v8_eQTL.tar) from GTEx portal release v8 (https://gtexportal.org). If a DNA-binding site was located within a 100-bp region around a single-nucleotide polymorphism (SNP) of an eQTL, we assumed that the DBP regulated the expression of the transcript corresponding to the eQTL. If the SNP of an eQTL was not located on the same chromosome as the transcribed region of the eQTL (i.e., *trans*-eQTL), the eQTL data were not used. The genomic positions of the eQTLs were converted to the corresponding positions in the hg19 version of the human genome using the LiftOver tool (http://hgdownload.soe.ucsc.edu/downloads.html#liftover).

The original model of the input data of DNA-binding sites in promoter regions used for the deep learning analysis consisted of the lengths of a DBP ChIP-seq peak in each 100-base bin in a region between 2 kbp upstream and 1 kbp downstream of the transcription start site (TSS) of each gene or transcript ^14^. To add the input data of DNA-binding sites of DBPs in more upstream and downstream regions of a TSS and to reduce the number of bins, we modified the model and first computed the length of a DBP ChIP-seq peak in a 100-base bin and then selected the maximum length in each interval of 100 bins in a region between 1 Mbp upstream and downstream of a TSS. This step outputted the 200 longest DBP ChIP-seq peaks in the examined regions.

### Training step for machine learning analysis

In the training of the modified deep learning model, we used the same test, training, and validation data as in the original method: all 2,705 genes encoded on chromosome 1 as test data, training data (22,251 genes), and validation data (2,472 genes) randomly selected from the other genes ^14^. The input data for promoters and putative enhancer regions were concatenated into a total of 230 bins of the lengths of 1,310 DBP ChIP-seq peaks, consisting of 30 bins for promoters and 200 bins for putative enhancers because the DeepLIFT tool ’RevealCancel rule on dense layers and the Rescale rule on conv layers (genomics default)’, with high scoring accuracy, ^24^ cannot be applied to a neural network with multiple input layers (Fig. S5). Ubuntu (version 20.04), Python (version 3.8.10), TensorFlow (version 2.6.0), NVIDIA CUDA toolkit (version 11.4.2), GeForce RTX 3080 GPU with 10 GB memory, and Intel Core i7-10700 CPU with 128 GB memory were used for training and prediction, and the original scripts were modified to run on newer versions of the software. The training method was the same as the original, and the training time was two to four hours. In this study, we focused on the analysis of HFF cells and assessed the training process using all the input data as the test data, which slightly improved the prediction of gene expression. Therefore, we used the trained model for the analysis of insulator-associated DBPs.

### Prediction of insulator-associated DBPs affecting EPIs DNA-binding sites of putative insulator-associated DBPs

To identify the direction of DBP-binding motifs in the DNA-binding sites of predicted insulator-associated DBPs, we searched for DBP-binding motifs in open chromatin regions in HFF cells (GSE170964) using the PIQ tool ^17^ and the position frequency matrices of the DBP-binding motifs. The motif sequences were collected from databases and previous studies, specifically, TRANSFAC (2019.2), JASPAR (2018), UniPROBE (2018), DNA motifs from high-throughput SELEX, DNA motifs from ENCODE DBP ChIP-seq data, and HOCOMOCO versions 9 and 11 ^18–23^. To reduce the number of false-positive prediction of DNA-binding sites, only DNA-binding sites that overlapped the ChIP-seq peaks of the same DBP (GTRD database release v20.06 [https://gtrd.biouml.org]) were used for further analyses. The genomic positions of ChIP-seq and DNase-seq data in public databases were converted to those provided in the hg19 version of the human genome using the LiftOver tool.

To confirm that CTCF DNA-binding sites were located at chromatin interaction sites, we examined the overlap between CTCF ChIP-seq peaks and the sites of *in situ* ChIA-PET chromatin interactions mediated by CTCF in human foreskin fibroblast (HFF) cells (4D Nucleome data portal, https://data.4dnucleome.org/files-processed/4DNFI73AFN12/). The directional difference between RAD21 DNA-binding sites located at chromatin interaction sites was also examined using ChIA-PET chromatin interactions mediated by RAD21 in human fibroblast from skin (ENCLB147AJZ). RAD21 DNA-binding sites within 100 bp of chromatin interaction sites were associated with chromatin interactions. To eliminate DNA-binding sites that overlapped EPI, we obtained ChIP-seq peaks of histone modifications, such as H3K4me3, H3K27ac, and H3K4me1, associated with promoter and enhancer regions in HFF cells (4D Nucleome data portal, 4DNFIJOK1BWR, 4DNFI442BQLO, and 4DNFIAGV3X99).

### Preparation of input data considering the DNA-binding sites of an insulator-associated DBP

To prepare the input data of the DNA-binding sites of DBPs in enhancers and promoters for the deep learning analysis, first, a transcript regulated by a DBP bound to enhancers and promoters was predicted based on the single nearest gene association rule (Fig. 2c; McLean et al. ^15^ Fig. S2c), as we found that the association rule predicted EPIs more accurately than the other rules in our previous study ^15^. The maximum length of the regulatory domain for each transcript was set to 1 Mbp upstream and downstream of the TSS. Then, considering the genomic locations and effects of the insulator sites of a DBP on EPIs, the length of a DBP ChIP-seq peak in a 100-base bin was calculated from a TSS to the DNA-binding site of a predicted insulator-associated DBP. To reduce the computational time of the analysis, we first calculated the length of a DBP ChIP-seq peak in the regulatory domains of all transcripts in a 100-base bin. Then, for each interval of 100 bins between a TSS and a DNA-binding site of a predicted insulator-associated DBP in the regulatory domain, we computed the maximum length of each DBP ChIP-seq peak in 100 bins on the basis of the precomputed length of the DBP ChIP-seq peak in each bin. In the case of a set of 100 bins containing the DNA-binding site of a predicted insulator-associated DBP, we computed the maximum lengths from the bin closest to a TSS to the first bin containing the DNA-binding site of the insulator-associated DBP, taking into account the orientation of the DNA-binding site.

### Prediction of insulator-associated DBPs using DeepLIFT tool

The DeepLIFT tool (version 0.6.13.0) was used to calculate the contribution (DeepLIFT score) of each DNA-binding site to the prediction of the expression level of a transcript ^24^. The contribution scores of DNA-binding sites of DBPs in putative enhancers and promoters upstream and downstream of each transcript were normalized by the sum of the absolute values of all contribution scores for the transcript, as the range of contribution scores changes according to the range of the predicted output (i.e., the expression level of a transcript), as described in a previous study ^58^. To estimate the effect of the DNA-binding sites of a predicted insulator-associated DBP on the prediction of transcript levels, positive normalized contribution scores for transcripts with DNA-binding sites in upstream or downstream regions were subjected to comparisons between all combinations of two different orientations (i.e., all combinations of two orientations among FR, RF, forward-forward [FF], and reverse-reverse [RR] orientations, and no orientation [X]: FR vs. RF, FR vs. X, RF vs. X, FF vs. RR, FF vs. X, RR vs. X, FF vs. FR, FF vs. RF, RR vs. FR, and RR vs. RF) among the DNA-binding sites of a predicted insulator-associated DBP by using a statistical test (Mann‒Whitney *U* test, FDR-adjusted *p* value < 0.05 and *p* value < 10^-3^). For EPIs that were common to the predictions based on two different orientations of DNA-binding sites of a predicted insulator-associated DBP, the DNA-binding sites of DBPs in putative enhancers and promoters among the common EPIs were omitted in the comparison of positive normalized contribution scores between the two different orientations. Based on the results of the statistical analysis and the mean differences in contribution scores, we predicted an insulator-associated DBP on the basis of the directional bias or lack of orientation bias of DNA-binding sites; for example, to predict an insulator-associated DBP with a DNA-binding site in the FR orientation, we selected a DBP for which the contribution scores among all combinations of FR vs. RF, FR vs. FF, FR vs. RR, and FR vs. X were significant (Mann‒Whitney *U* test, FDR-adjusted *p* value < 0.05 and *p* value < 10^-3^) and showed positive (high) mean differences among the contribution scores of all the combinations.

### Search for insulator-associated DNA-binding sites based on histone modifications and gene expression levels

The genomic positions in the BAM files of H3K27me3 and H3K9me3 ChIP-seq data (ENCFF267JBO, ENCFF528UZJ) from HFF cells were converted to the corresponding positions in the hg19 version of the human genome using the CrossMap tool ^63^ and then converted to WIG files using bam2wig.pl from Bio::ToolBox (https://github.com/tjparnell/biotoolbox) to visualize the data on the UCSC Genome Browser through custom tracks (http://www.genome.ucsc.edu/cgi-bin/hgCustom) and a track hub (https://genome.ucsc.edu/cgi-bin/hgHubConnect). The BAM file of RNA-seq data (GSM941744) of HFF cells was converted to a WIG file using bam2wig.pl in Bio::ToolBox. The genomic positions in the BED file of the boundaries and the distribution of insulation scores identified through a Micro-C assay (4D Nucleome data portal, https://data.4dnucleome.org/files-processed/4DNFI5OZ7TBE/ and 4DNFIPMHBA41) were converted to the corresponding positions in the hg19 version of the human genome.

### Distribution of DNA-binding sites around chromatin interaction sites

We examined the distribution of the DNA-binding sites of predicted insulator-associated DBPs in regions within ±500 bp of the center of a chromatin interaction site using chromatin interaction data from HFF cells obtained via a Micro-C assay (4D Nucleome data portal, https://data.4dnucleome.org/files-processed/4DNFI1MCF9GZ/). For the insulator-associated DBPs other than CTCF, RAD21, and SMC3, the DNA-binding sites that do not overlap with those of CTCF, RND21, and SMC3 were used to examine their distribution around interaction sites. The number of DNA-binding sites of predicted insulator-associated DBPs that cooccurred with DNA-binding sites of known insulator-associated DBPs, such as CTCF, RAD21, and SMC3, was counted in each 10-bp bin in the regions within ±500 bp of the center of a chromatin interaction site.

### Insulator-associated DNA-binding sites near splice sites

Insulator-associated DNA-binding sites were searched within 200 bp of the center of splice sites using the file of the genomic positions of alternative transcripts in the UCSC Genome Browser (https://hgdownload.soe.ucsc.edu/goldenPath/hg19/database/all_mrna.txt.gz ; https://genome.ucsc.edu/cgi-bin/hgTables?db=hg19&hgta_group=rna&hgta_track=mrna&hgta_table=all_mrna&hgta_doSchema=describe+table+schema). We performed the statistical test to estimate the enrichment of insulator-associated DNA-binding proteins at the splicing sites compared with the other DNA-binding proteins (Mann‒Whitney *U* test, *p* value < 0.05).

### Statistical analysis

Comparison of the distribution of DeepLIFT scores between different orientations of DNA-binding sites was performed using the Mann-Whitney *U* test with the scipy.stats.mannwhitneyu library. Two-tailed *p* values were calculated. For multiple comparisons, adjusted *p* values were calculated by false discovery rate (FDR) using the Benjamini-Hochberg procedure. *Z* scores for the Mann-Whitney *U* test were not output by the library. Therefore, we calculated *z* scores by z-standardizing the *U* values. The code is made available on GitHub at https://github.com/reposit2/insulator/tree/main.

## Data availability

Human foreskin fibroblast (HFF) cell RNA-seq data were obtained from GSM941744. Human monocyte RNA-seq data were obtained from the Blueprint DCC portal http://dcc.blueprint-epigenome.eu/#/files at ’C0010KB1.transcript_quantification.rsem_grape2_crg.GRCh38.20150622.results’. The RNA-seq data of human mammary epithelial cells (HMEC) were found on ENCODE at UCSC http://genome.ucsc.edu/encode/ at ’wgEncodeCshlLongRnaSeqHmecCellPapAlnRep1.bam’. Human neural progenitor cell (NPC) RNA-seq data were obtained from GSM915326. The DNase-seq data of HFF cells were obtained from GSE170964. The *in situ* ChIA-PET chromatin interaction data mediated by CTCF in HFF cells were available at the 4D Nucleome data portal, https://data.4dnucleome.org/files-processed/4DNFI73AFN12/. The ChIP-seq data of H3K4me3, H3K27ac, and H3K4me1 of HFF cells were found at the 4D Nucleome data portal, 4DNFIJOK1BWR, 4DNFI442BQLO, and 4DNFIAGV3X99. The ChIP-seq data of H3K27me3 and H3K9me3 in HFF cells were obtained from the ENCODE portal, ENCFF267JBO and ENCFF528UZJ. The chromatin interaction data and boundary data of HFF cells by the Micro-C were obtained from the 4D Nucleome data portal, https://data.4dnucleome.org/files-processed/4DNFI1MCF9GZ/ and https://data.4dnucleome.org/files-processed/4DNFI5OZ7TBE/. ChIP-seq data of TFs were downloaded from the v20.06 release of the GTRD database (https://gtrd.biouml.org). eQTL data (GTEx_Analysis_v8_eQTL.tar) were downloaded from the v8 release of the GTEx portal (https://gtexportal.org). DNA binding motifs of TFs were collected from TRANSFAC (2019.2), JASPAR (2018) (https://jaspar.genereg.net/), UniPROBE (2018) (http://the_brain.bwh.harvard.edu/uniprobe/), high-throughput SELEX (https://doi.org/10.1016/j.cell.2012.12.009), transcription factor binding sequences of ENCODE TF ChIP-seq data (http://compbio.mit.edu/encode-motifs/), and HOCOMOCO version 9 and 11 (https://autosome.org/HOCOMOCOS/) (https://hocomoco11.autosome.org/). The genomic positions of alternative transcripts in the UCSC Genome Browser (https://hgdownload.soe.ucsc.edu/goldenPath/hg19/database/all_mrna.txt.gz).

## Code availability

Demonstration codes are made available at https://github.com/reposit2/insulator/tree/main and https://doi.org/10.5281/zenodo.8216164.

## Acknowledgements

We would like to thank Prof. Martin Frith for reviewing the manuscript. Supercomputing resources were provided by the Human Genome Center of the Institute of Medical Science at the University of Tokyo and ROIS National Institute of Genetics. This work was partially supported by JSPS KAKENHI (Grant Number 16K00387), JST CREST (Grant Number JPMJCR15G1), Osaka University start-up research grant, KIOXIA young scientist research grant at Waseda University, Sumitomo basic science research projects to NO, AMED [grant nos. JP21ae0121049, 21479280], and JSPS KAKENHI [grant nos. 22H04925, 20H00624, 17K20032] to MH.

## Author contributions

N.O. designed and performed the research, and wrote and revised the manuscript. M.H. revised the manuscript.

## Additional information

**Supplementary information** The online version contains supplementary material available at https://doi.org/.

**Correspondence** and requests for materials should be addressed to Naoki Osato.

