## Supplementary material for "Systematic discovery of directional regulatory motifs associated with human insulator sites": Supplementary_Fig1-2.pdf

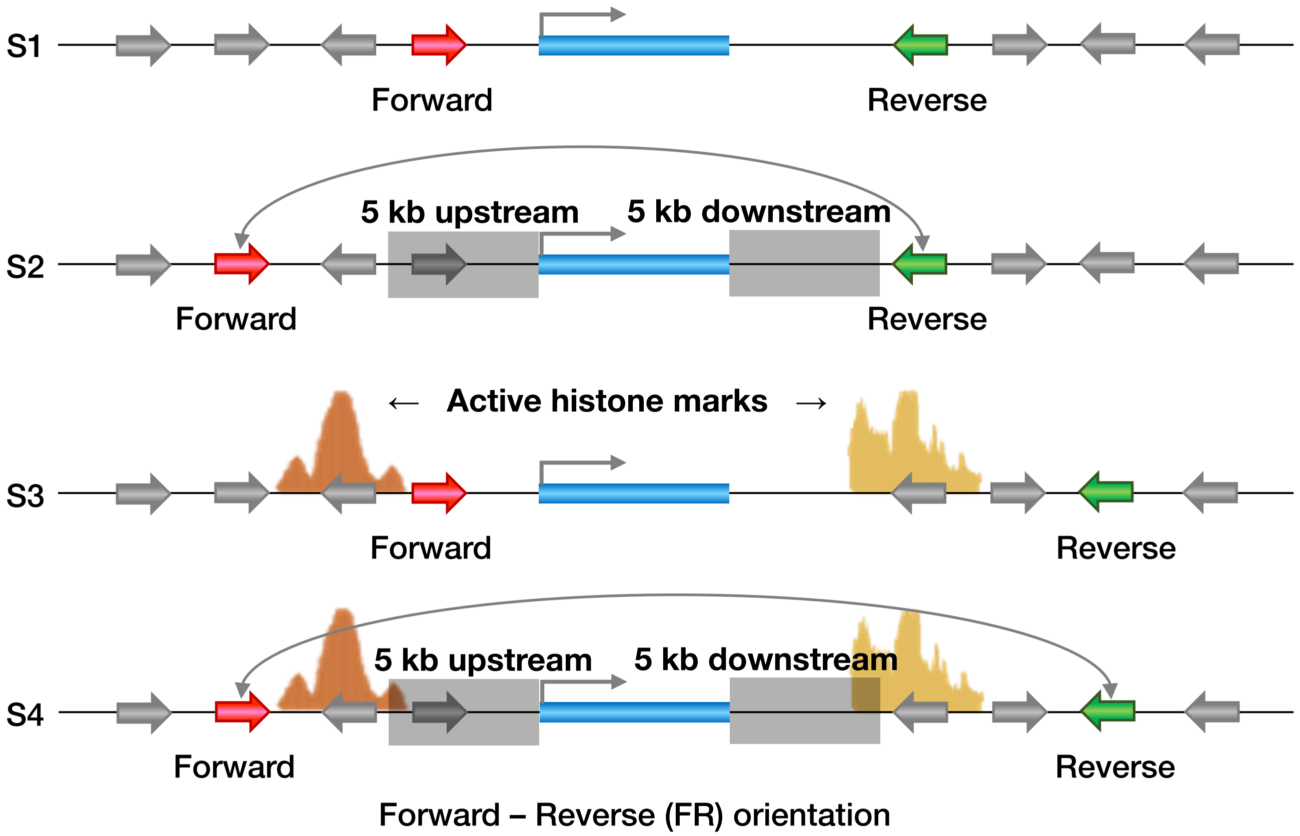


Supplementary Fig. 1. Selection criteria for DNA-binding sites of insulator-associated DBPs.

Selection Criteria S1: DNA-binding sites of predicted insulator-associated DBPs.

S2: DNA-binding sites of predicted insulator-associated DBPs that overlap with chromatin interaction sites located more than 5 kb upstream to 5 kb downstream of a transcribed region.

S3: DNA-binding sites of predicted insulator-associated DBPs that do not overlap with ChIP-seq peaks for H3K4me3, H3K4me1, or H3K27ac.

S4: DNA-binding sites of predicted insulator-associated DBPs that overlap chromatin interaction sites (more than 5 kb upstream to 5 kb downstream of a transcript) and do not overlap with ChIP-seq peaks for H3K4me3, H3K4me1, or H3K27ac.

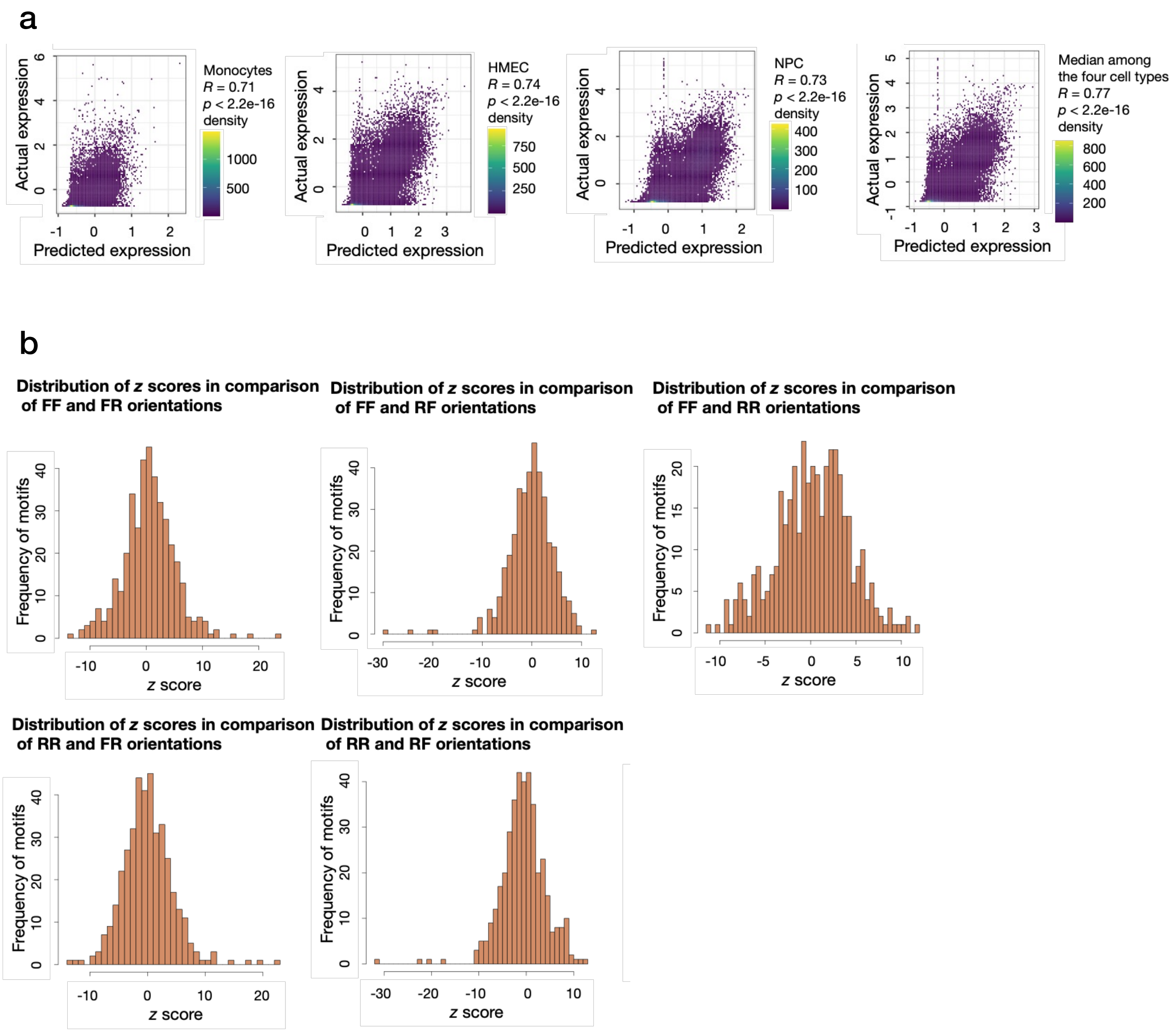


Supplementary Fig. 2. Estimation of the effect of directional regulatory motifs on gene expression.

a Comparison of actual and predicted gene expression levels in monocytes, HMEC, and NPC, as well as comparison of actual and predicted median expression levels across four cell types: HFF, monocytes, HMEC, and NPC.

b Distribution of *z*-scores from the Mann–Whitney U test for DeepLIFT scores, comparing different orientations of DNA-binding sites of proteins.
