## Supplementary material for "Systematic discovery of directional regulatory motifs associated with human insulator sites": Supplementary_Fig3.pdf

Supplementary Fig. 3. DNA-binding sites of predicted insulator-associated DBPs in potential insulator sites.

**a** Insulator sites defined as boundary regions identified by Micro-C chromatin interaction data. Pages 2–97 display UCSC Genome Browser screenshots showing regions around insulator-associated DNA-binding sites. DNA-binding sites selected based on two criteria are shown at the bottom of each figure: the upper panel displays putative insulator-associated DNA-binding sites identified in this study, while the lower panel shows DNA-binding sites for three DBPs—CTCF, RAD21, and SMC3—within open chromatin regions overlapping the corresponding ChIP-seq peaks.

"Chromatin interactions 200 bp" refers to Micro-C-detected chromatin interactions within a 200-bp region centered on the insulator-associated DNA-binding site in HFF cells. "Chromatin interactions 10 bp, 20 bp, and 50 bp" refer to interactions within 10-bp, 20-bp, and 50-bp regions, respectively. A green box around CTCF, RAD21, and SMC3 binding sites indicates proximity to an insulator site.

**b** Insulator-associated DNA-binding sites where H3K9me3 marks differ between the upstream and downstream regions of the site (pages 98–102).

**c** Clusters of DNA-binding sites of insulator-associated DBPs (page 103).

**d** Differences in chromatin interactions between DBPs at the same loci (pages 104–105).

**e** Potential regulation of alternative transcription (pages 106–107).

**f** Insulator sites identified as boundaries between transcriptionally repressed regions (based on H3K27me3 marks) and transcribed regions (based on RNA-seq data). Pages 108–203 show UCSC Genome Browser screenshots around insulator-associated DNA-binding sites.

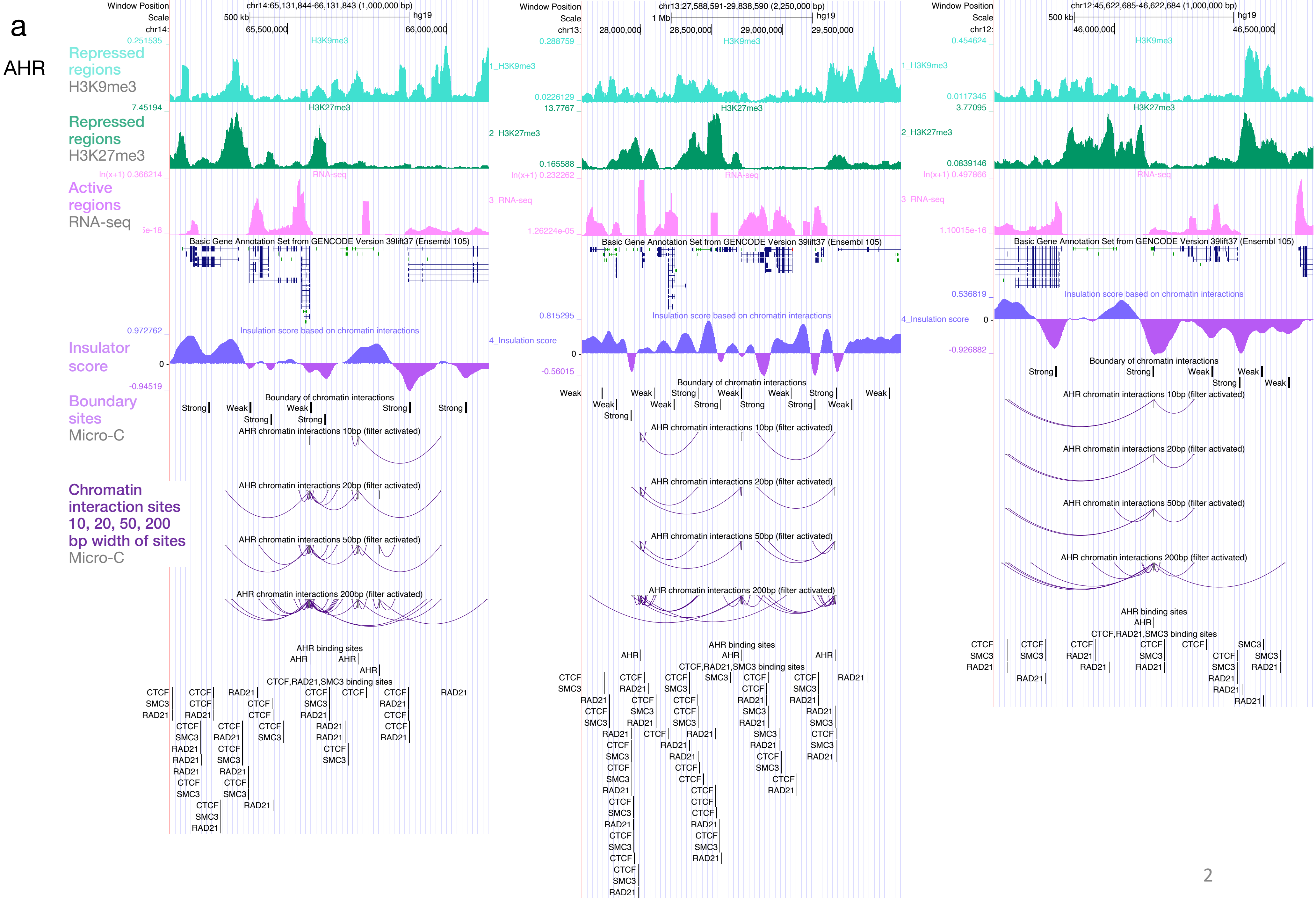

a
AHR
Repressed regions H3K9me3
Repressed regions H3K27me3
Active regions RNA-seq
Insulator score
Boundary sites Micro-C
Chromatin interaction sites 10, 20, 50, 200 bp width of sites Micro-C
Window Position
Scale
chr14:65,131,844-66,131,843 (1,000,000 bp)
500 kb
hg19
65,500,000
66,000,000
chr13:27,588,591-29,838,590 (2,250,000 bp)
1 Mb
28,000,000
28,500,000
29,000,000
29,500,000
chr12:45,622,685-46,622,684 (1,000,000 bp)
46,000,000
46,500,000
H3K9me3
1_H3K9me3
H3K27me3
2_H3K27me3
RNA-seq
3_RNA-seq
Basic Gene Annotation Set from GENCODE Version 39lift37 (Ensembl 105)
Insulation score based on chromatin interactions
4_Insulation score
Boundary of chromatin interactions
Strong
Weak
AHR chromatin interactions 10bp (filter activated)
AHR chromatin interactions 20bp (filter activated)
AHR chromatin interactions 50bp (filter activated)
AHR chromatin interactions 200bp (filter activated)
AHR binding sites
AHR
CTCF,RAD21,SMC3 binding sites
CTCF
SMC3
RAD21

### ASCL1

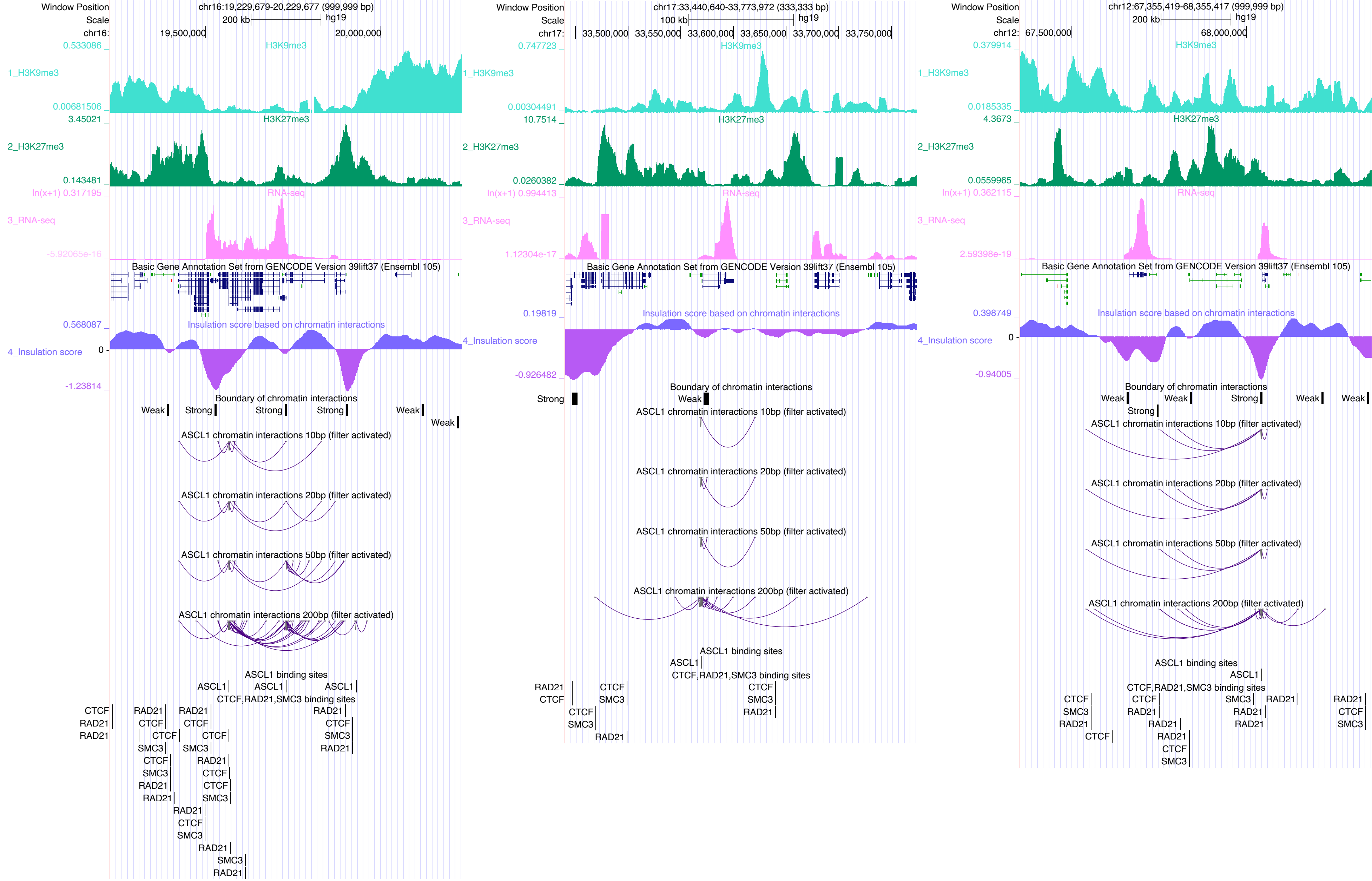

Window Position
chr16:19,229,679-20,229,677 (999,999 bp)
Scale
200 kb
hg19
chr16:
19,500,000
20,000,000
H3K9me3
H3K27me3
RNA-seq
Basic Gene Annotation Set from GENCODE Version 39lift37 (Ensembl 105)
Insulation score based on chromatin interactions
Boundary of chromatin interactions
ASCL1 chromatin interactions 10bp (filter activated)
ASCL1 chromatin interactions 20bp (filter activated)
ASCL1 chromatin interactions 50bp (filter activated)
ASCL1 chromatin interactions 200bp (filter activated)
ASCL1 binding sites
CTCF,RAD21,SMC3 binding sites
chr17:33,440,640-33,773,972 (333,333 bp)
100 kb
chr17:
33,500,000
33,550,000
33,600,000
33,650,000
33,700,000
33,750,000
chr12:67,355,419-68,355,417 (999,999 bp)
chr12:
67,500,000
68,000,000

### ATF2

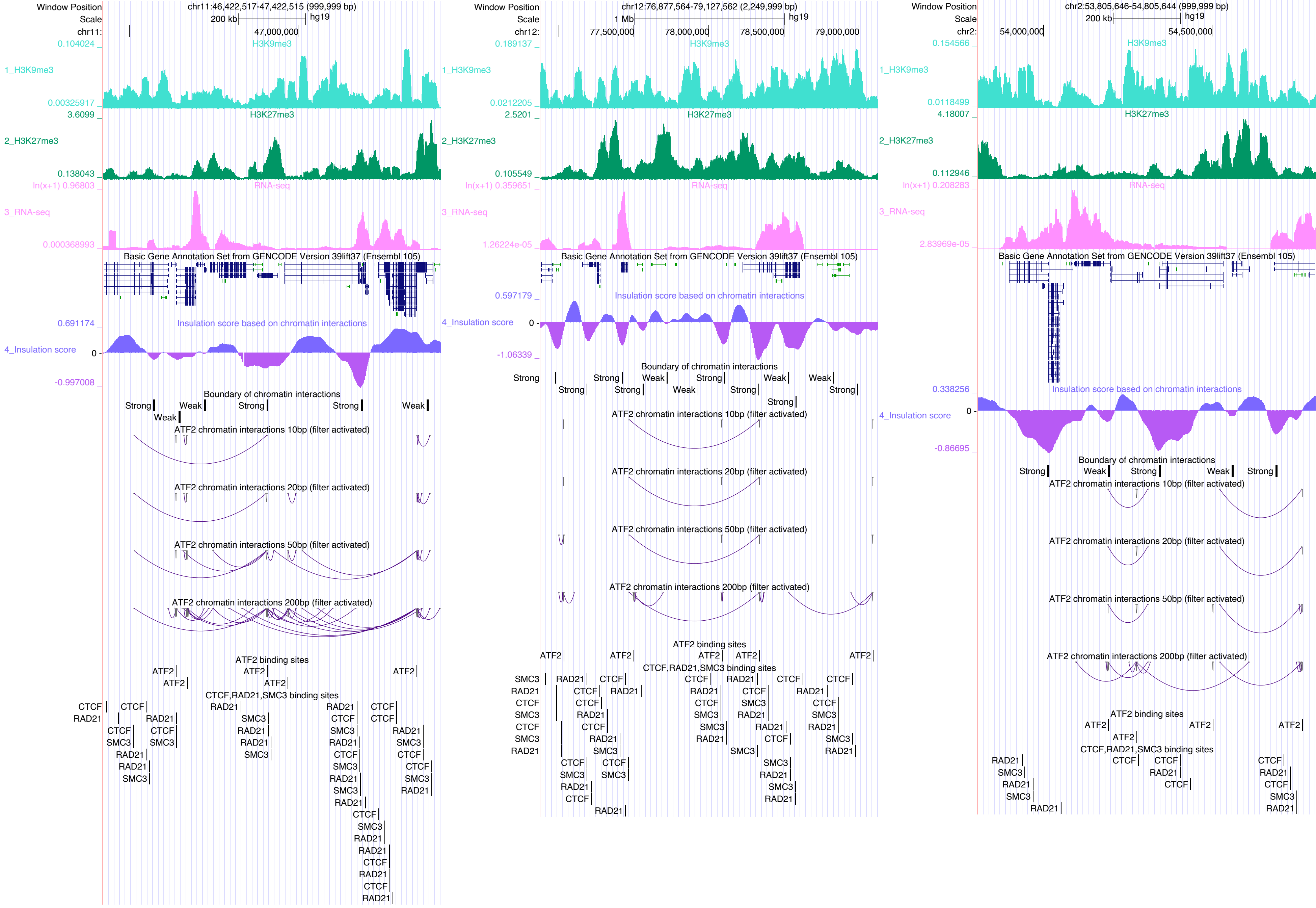
Window Position
chr11:46,422,517-47,422,515 (999,999 bp)
Scale
200 kb
hg19
chr11:
47,000,000
0.104024
H3K9me3
1_H3K9me3
0.00325917
3.6099
H3K27me3
2_H3K27me3
0.138043
ln(x+1) 0.96803
RNA-seq
3_RNA-seq
0.000368993
Basic Gene Annotation Set from GENCODE Version 39lift37 (Ensembl 105)
0.691174
Insulation score based on chromatin interactions
4_Insulation score
0
-0.997008
Boundary of chromatin interactions
Strong
Weak
ATF2 chromatin interactions 10bp (filter activated)
ATF2 chromatin interactions 20bp (filter activated)
ATF2 chromatin interactions 50bp (filter activated)
ATF2 chromatin interactions 200bp (filter activated)
ATF2 binding sites
ATF2
CTCF,RAD21,SMC3 binding sites
CTCF
RAD21
SMC3
Window Position
chr12:76,877,564-79,127,562 (2,249,999 bp)
Scale
1 Mb
hg19
chr12:
77,500,000
78,000,000
78,500,000
79,000,000
0.189137
H3K9me3
1_H3K9me3
0.0212205
2.5201
H3K27me3
2_H3K27me3
0.105549
ln(x+1) 0.359651
RNA-seq
3_RNA-seq
1.26224e-05
Basic Gene Annotation Set from GENCODE Version 39lift37 (Ensembl 105)
0.597179
Insulation score based on chromatin interactions
4_Insulation score
0
-1.06339
Boundary of chromatin interactions
Strong
Weak
ATF2 chromatin interactions 10bp (filter activated)
ATF2 chromatin interactions 20bp (filter activated)
ATF2 chromatin interactions 50bp (filter activated)
ATF2 chromatin interactions 200bp (filter activated)
ATF2 binding sites
ATF2
CTCF,RAD21,SMC3 binding sites
SMC3
RAD21
CTCF
Window Position
chr2:53,805,646-54,805,644 (999,999 bp)
Scale
200 kb
hg19
chr2:
54,000,000
54,500,000
0.154566
H3K9me3
1_H3K9me3
0.0118499
4.18007
H3K27me3
2_H3K27me3
0.112946
ln(x+1) 0.208283
RNA-seq
3_RNA-seq
2.83969e-05
Basic Gene Annotation Set from GENCODE Version 39lift37 (Ensembl 105)
0.338256
Insulation score based on chromatin interactions
4_Insulation score
0
-0.86695
Boundary of chromatin interactions
Strong
Weak
ATF2 chromatin interactions 10bp (filter activated)
ATF2 chromatin interactions 20bp (filter activated)
ATF2 chromatin interactions 50bp (filter activated)
ATF2 chromatin interactions 200bp (filter activated)
ATF2 binding sites
ATF2
CTCF,RAD21,SMC3 binding sites
RAD21
SMC3
CTCF

### ATF7

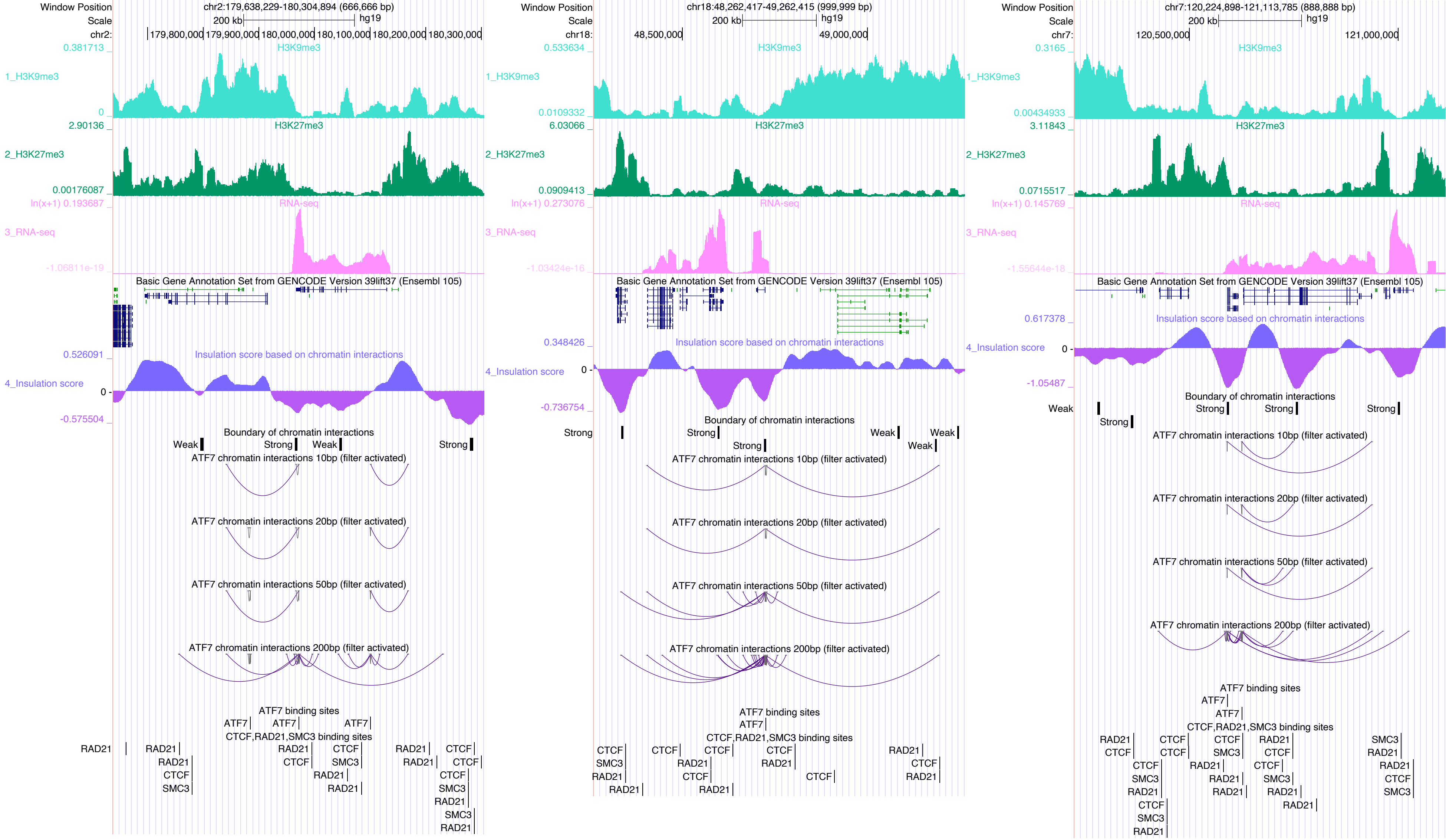
Window Position
chr2:179,638,229-180,304,894 (666,666 bp)
Scale
200 kb
hg19
chr2:
179,800,000
179,900,000
180,000,000
180,100,000
180,200,000
180,300,000
1_H3K9me3
H3K9me3
0.381713
0
2_H3K27me3
H3K27me3
2.90136
0.00176087
3_RNA-seq
RNA-seq
ln(x+1) 0.193687
-1.06811e-19
Basic Gene Annotation Set from GENCODE Version 39lift37 (Ensembl 105)
4_Insulation score
Insulation score based on chromatin interactions
0.526091
0
-0.575504
Boundary of chromatin interactions
Weak
Strong
Weak
Strong
ATF7 chromatin interactions 10bp (filter activated)
ATF7 chromatin interactions 20bp (filter activated)
ATF7 chromatin interactions 50bp (filter activated)
ATF7 chromatin interactions 200bp (filter activated)
ATF7 binding sites
ATF7
ATF7
ATF7
CTCF,RAD21,SMC3 binding sites
RAD21
RAD21
RAD21
CTCF
SMC3
RAD21
CTCF
CTCF
SMC3
RAD21
RAD21
RAD21
RAD21
CTCF
CTCF
CTCF
SMC3
RAD21
SMC3
RAD21
Window Position
chr18:48,262,417-49,262,415 (999,999 bp)
Scale
200 kb
hg19
chr18:
48,500,000
49,000,000
1_H3K9me3
H3K9me3
0.533634
0.0109332
2_H3K27me3
H3K27me3
6.03066
0.0909413
3_RNA-seq
RNA-seq
ln(x+1) 0.273076
-1.03424e-16
Basic Gene Annotation Set from GENCODE Version 39lift37 (Ensembl 105)
4_Insulation score
Insulation score based on chromatin interactions
0.348426
0
-0.736754
Boundary of chromatin interactions
Strong
Strong
Strong
Weak
Weak
Weak
ATF7 chromatin interactions 10bp (filter activated)
ATF7 chromatin interactions 20bp (filter activated)
ATF7 chromatin interactions 50bp (filter activated)
ATF7 chromatin interactions 200bp (filter activated)
ATF7 binding sites
ATF7
CTCF,RAD21,SMC3 binding sites
CTCF
SMC3
RAD21
RAD21
CTCF
CTCF
RAD21
CTCF
RAD21
CTCF
RAD21
RAD21
CTCF
RAD21
CTCF
RAD21
Window Position
chr7:120,224,898-121,113,785 (888,888 bp)
Scale
200 kb
hg19
chr7:
120,500,000
121,000,000
1_H3K9me3
H3K9me3
0.3165
0.00434933
2_H3K27me3
H3K27me3
3.11843
0.0715517
3_RNA-seq
RNA-seq
ln(x+1) 0.145769
-1.55644e-18
Basic Gene Annotation Set from GENCODE Version 39lift37 (Ensembl 105)
4_Insulation score
Insulation score based on chromatin interactions
0.617378
0
-1.05487
Boundary of chromatin interactions
Weak
Strong
Strong
Strong
Strong
ATF7 chromatin interactions 10bp (filter activated)
ATF7 chromatin interactions 20bp (filter activated)
ATF7 chromatin interactions 50bp (filter activated)
ATF7 chromatin interactions 200bp (filter activated)
ATF7 binding sites
ATF7
ATF7
CTCF,RAD21,SMC3 binding sites
RAD21
CTCF
CTCF
SMC3
RAD21
CTCF
SMC3
RAD21
CTCF
CTCF
RAD21
CTCF
SMC3
RAD21
CTCF
RAD21
RAD21
CTCF
SMC3
RAD21
RAD21
SMC3
RAD21
RAD21
CTCF
SMC3

### BATF

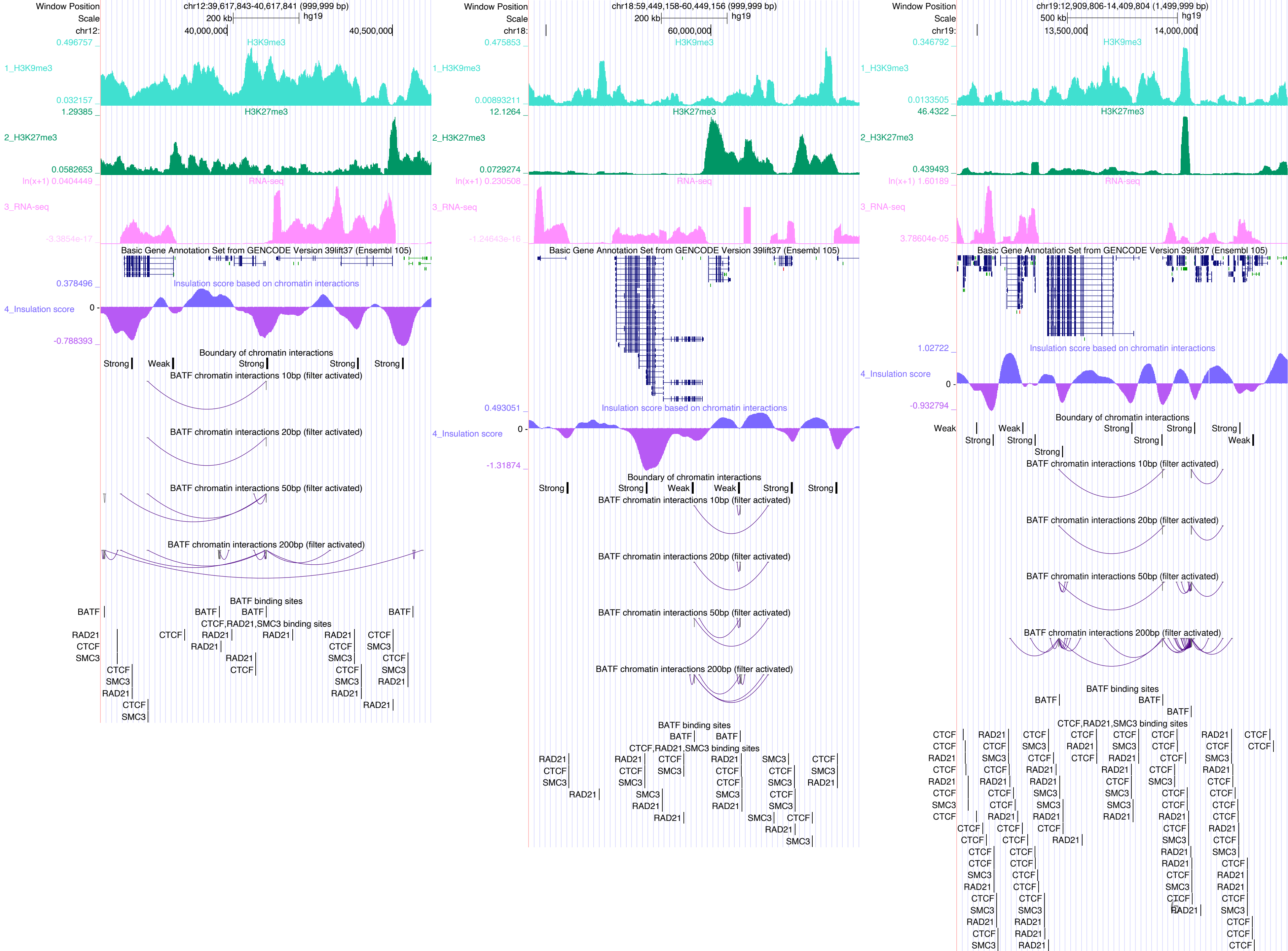

Window Position
Scale
chr12:39,617,843-40,617,841 (999,999 bp)
200 kb
hg19
chr12:
40,000,000
40,500,000
1_H3K9me3
2_H3K27me3
3_RNA-seq
4_Insulation score
H3K9me3
H3K27me3
RNA-seq
Basic Gene Annotation Set from GENCODE Version 39lift37 (Ensembl 105)
Insulation score based on chromatin interactions
Boundary of chromatin interactions
BATF chromatin interactions 10bp (filter activated)
BATF chromatin interactions 20bp (filter activated)
BATF chromatin interactions 50bp (filter activated)
BATF chromatin interactions 200bp (filter activated)
BATF binding sites
CTCF,RAD21,SMC3 binding sites
chr18:59,449,158-60,449,156 (999,999 bp)
chr18:
60,000,000
chr19:12,909,806-14,409,804 (1,499,999 bp)
500 kb
chr19:
13,500,000
14,000,000

### BATF3

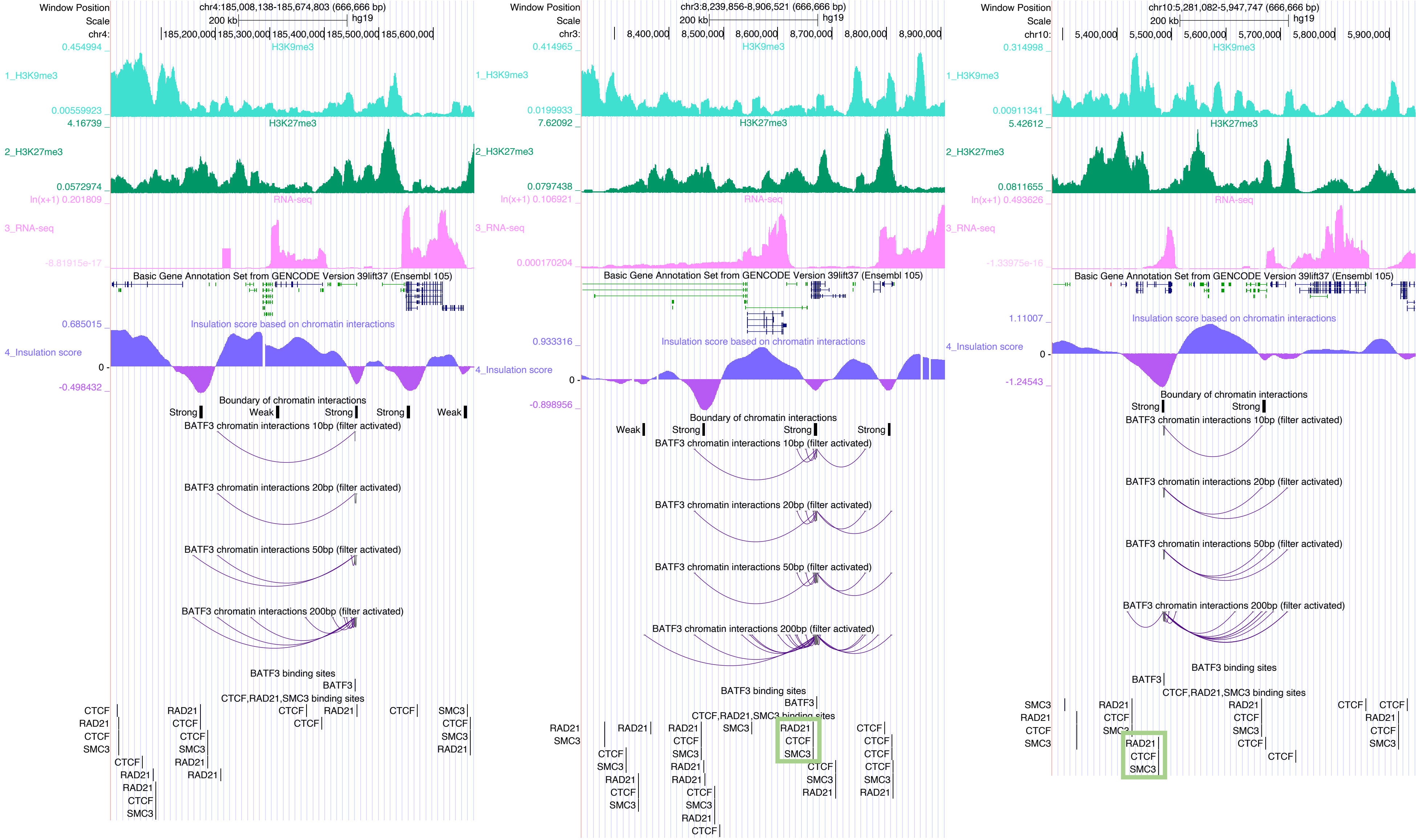

Window Position
chr4:185,008,138-185,674,803 (666,666 bp)
chr3:8,239,856-8,906,521 (666,666 bp)
chr10:5,281,082-5,947,747 (666,666 bp)
200 kb
hg19
1_H3K9me3
2_H3K27me3
3_RNA-seq
4_Insulation score
Basic Gene Annotation Set from GENCODE Version 39lift37 (Ensembl 105)
Insulation score based on chromatin interactions
Boundary of chromatin interactions
Strong
Weak
BATF3 chromatin interactions 10bp (filter activated)
BATF3 chromatin interactions 20bp (filter activated)
BATF3 chromatin interactions 50bp (filter activated)
BATF3 chromatin interactions 200bp (filter activated)
BATF3 binding sites
CTCF,RAD21,SMC3 binding sites
CTCF
RAD21
SMC3

BCL6

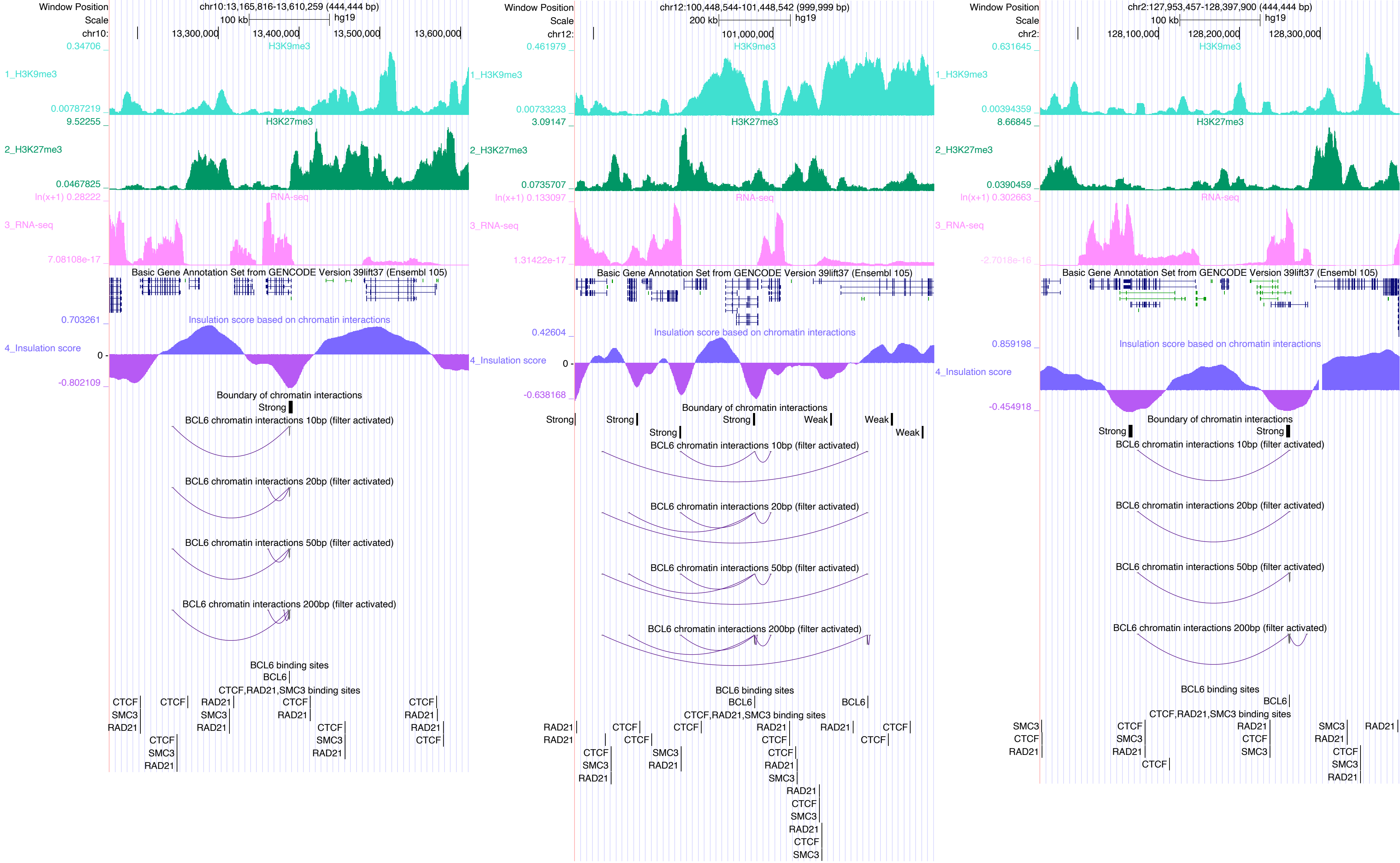

Window Position
Scale
chr10:13,165,816-13,610,259 (444,444 bp)
100 kb
hg19
chr10: 13,300,000 13,400,000 13,500,000 13,600,000
1_H3K9me3
H3K9me3
0.34706
0.00787219
2_H3K27me3
H3K27me3
9.52255
0.0467825
3_RNA-seq
RNA-seq
ln(x+1) 0.28222
7.08108e-17
Basic Gene Annotation Set from GENCODE Version 39lift37 (Ensembl 105)
4_Insulation score
Insulation score based on chromatin interactions
0.703261
0
-0.802109
Boundary of chromatin interactions
Strong
BCL6 chromatin interactions 10bp (filter activated)
BCL6 chromatin interactions 20bp (filter activated)
BCL6 chromatin interactions 50bp (filter activated)
BCL6 chromatin interactions 200bp (filter activated)
BCL6 binding sites
BCL6
CTCF,RAD21,SMC3 binding sites
CTCF
SMC3
RAD21
chr12:100,448,544-101,448,542 (999,999 bp)
200 kb
hg19
chr12: 101,000,000
0.461979
0.00733233
3.09147
0.0735707
ln(x+1) 0.133097
1.31422e-17
0.42604
-0.638168
Weak
chr2:127,953,457-128,397,900 (444,444 bp)
100 kb
hg19
chr2: 128,100,000 128,200,000 128,300,000
0.631645
0.00394359
8.66845
0.0390459
ln(x+1) 0.302663
-2.7018e-16
0.859198
-0.454918

### CDX2

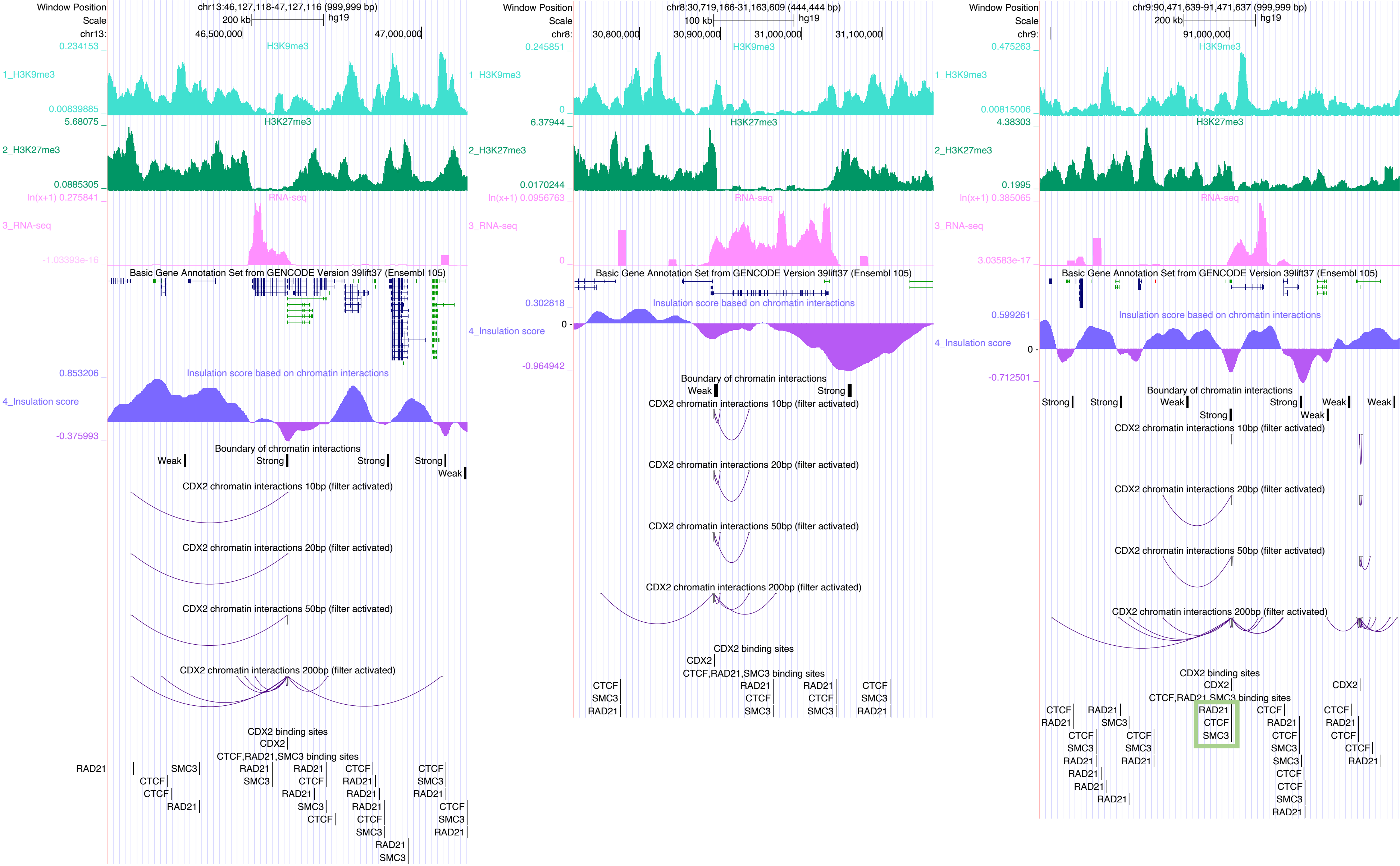

Window Position
Scale
chr13:46,127,118-47,127,116 (999,999 bp)
200 kb
hg19
chr13:
46,500,000
47,000,000
0.234153
H3K9me3
1_H3K9me3
0.00839885
5.68075
H3K27me3
2_H3K27me3
0.0885305
ln(x+1) 0.275841
RNA-seq
3_RNA-seq
-1.03393e-16
Basic Gene Annotation Set from GENCODE Version 39lift37 (Ensembl 105)
0.853206
Insulation score based on chromatin interactions
4_Insulation score
-0.375993
Boundary of chromatin interactions
Weak
Strong
Strong
Strong
Weak
CDX2 chromatin interactions 10bp (filter activated)
CDX2 chromatin interactions 20bp (filter activated)
CDX2 chromatin interactions 50bp (filter activated)
CDX2 chromatin interactions 200bp (filter activated)
CDX2 binding sites
CDX2
CTCF,RAD21,SMC3 binding sites
RAD21
SMC3
CTCF
CTCF
RAD21
RAD21
SMC3
RAD21
CTCF
RAD21
SMC3
CTCF
CTCF
RAD21
RAD21
RAD21
CTCF
SMC3
CTCF
SMC3
RAD21
CTCF
SMC3
RAD21
SMC3
RAD21
Window Position
Scale
chr8:30,719,166-31,163,609 (444,444 bp)
100 kb
hg19
chr8:
30,800,000
30,900,000
31,000,000
31,100,000
0.245851
H3K9me3
1_H3K9me3
0
6.37944
H3K27me3
2_H3K27me3
0.0170244
ln(x+1) 0.0956763
RNA-seq
3_RNA-seq
0
Basic Gene Annotation Set from GENCODE Version 39lift37 (Ensembl 105)
0.302818
Insulation score based on chromatin interactions
4_Insulation score
-0.964942
Boundary of chromatin interactions
Weak
Strong
CDX2 chromatin interactions 10bp (filter activated)
CDX2 chromatin interactions 20bp (filter activated)
CDX2 chromatin interactions 50bp (filter activated)
CDX2 chromatin interactions 200bp (filter activated)
CDX2 binding sites
CDX2
CTCF,RAD21,SMC3 binding sites
CTCF
SMC3
RAD21
RAD21
CTCF
SMC3
RAD21
CTCF
SMC3
CTCF
SMC3
RAD21
Window Position
Scale
chr9:90,471,639-91,471,637 (999,999 bp)
200 kb
hg19
chr9:
91,000,000
0.475263
H3K9me3
1_H3K9me3
0.00815006
4.38303
H3K27me3
2_H3K27me3
0.1995
ln(x+1) 0.385065
RNA-seq
3_RNA-seq
3.03583e-17
Basic Gene Annotation Set from GENCODE Version 39lift37 (Ensembl 105)
0.599261
Insulation score based on chromatin interactions
4_Insulation score
-0.712501
Boundary of chromatin interactions
Strong
Strong
Weak
Strong
Weak
Weak
Strong
Weak
CDX2 chromatin interactions 10bp (filter activated)
CDX2 chromatin interactions 20bp (filter activated)
CDX2 chromatin interactions 50bp (filter activated)
CDX2 chromatin interactions 200bp (filter activated)
CDX2 binding sites
CDX2
CDX2
CTCF,RAD21,SMC3 binding sites
CTCF
RAD21
RAD21
SMC3
CTCF
SMC3
RAD21
RAD21
RAD21
RAD21
CTCF
SMC3
RAD21
RAD21
CTCF
SMC3
CTCF
RAD21
CTCF
SMC3
SMC3
CTCF
CTCF
SMC3
RAD21
CTCF
RAD21
CTCF
CTCF
RAD21

### CEBPA

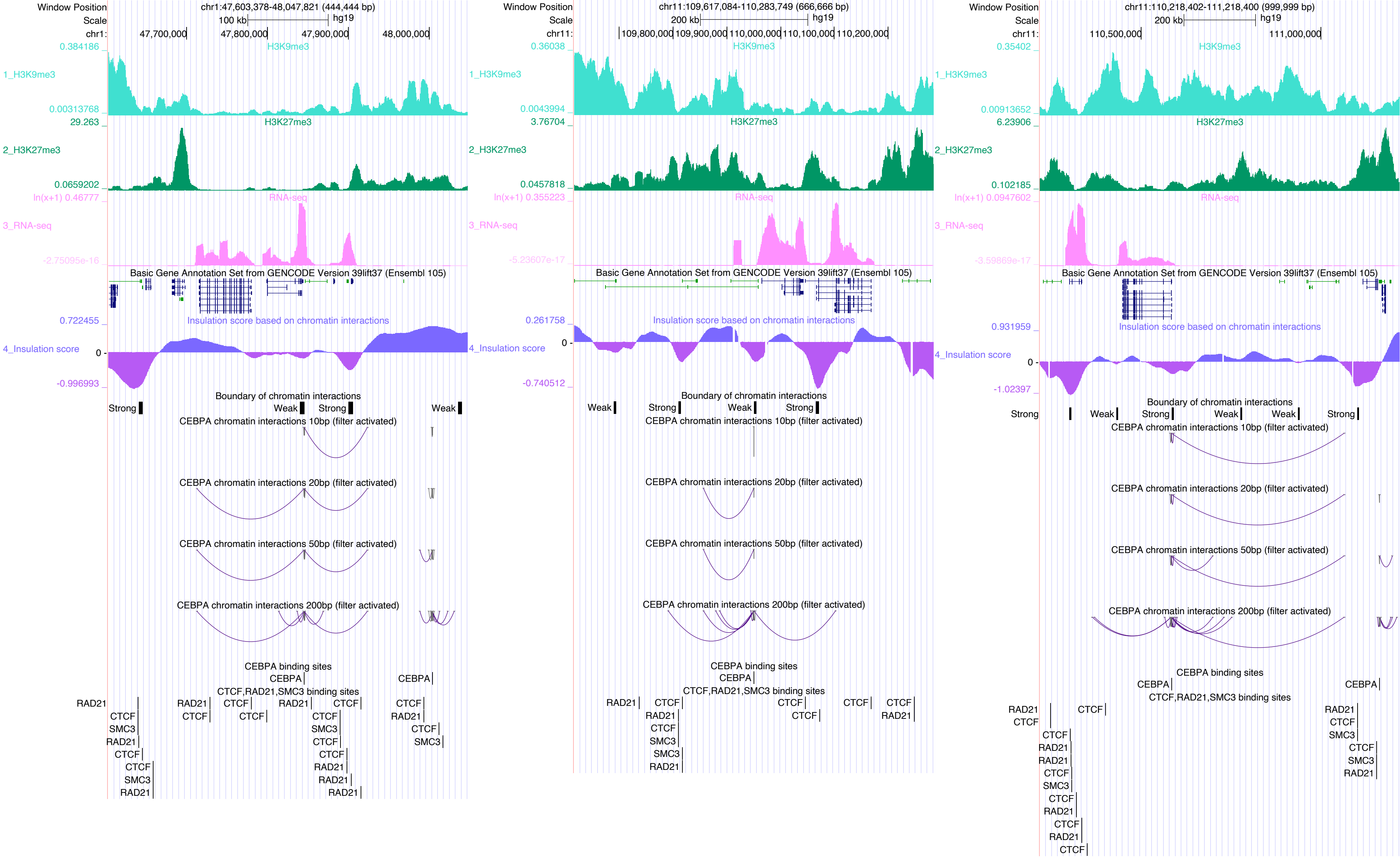
Window Position
Scale
chr1:47,603,378-48,047,821 (444,444 bp)
100 kb
hg19
47,700,000
47,800,000
47,900,000
48,000,000
1_H3K9me3
H3K9me3
0.384186
0.00313768
2_H3K27me3
H3K27me3
29.263
0.0659202
3_RNA-seq
RNA-seq
ln(x+1) 0.46777
-2.75095e-16
Basic Gene Annotation Set from GENCODE Version 39lift37 (Ensembl 105)
4_Insulation score
Insulation score based on chromatin interactions
0.722455
0
-0.996993
Boundary of chromatin interactions
Strong
Weak
CEBPA chromatin interactions 10bp (filter activated)
CEBPA chromatin interactions 20bp (filter activated)
CEBPA chromatin interactions 50bp (filter activated)
CEBPA chromatin interactions 200bp (filter activated)
CEBPA binding sites
CEBPA
CTCF,RAD21,SMC3 binding sites
RAD21
CTCF
SMC3
chr11:109,617,084-110,283,749 (666,666 bp)
200 kb
109,800,000
109,900,000
110,000,000
110,100,000
110,200,000
0.36038
0.0043994
3.76704
0.0457818
ln(x+1) 0.355223
-5.23607e-17
0.261758
-0.740512
chr11:110,218,402-111,218,400 (999,999 bp)
110,500,000
111,000,000
0.35402
0.00913652
6.23906
0.102185
ln(x+1) 0.0947602
-3.59869e-17
0.931959
-1.02397

### CLOCK

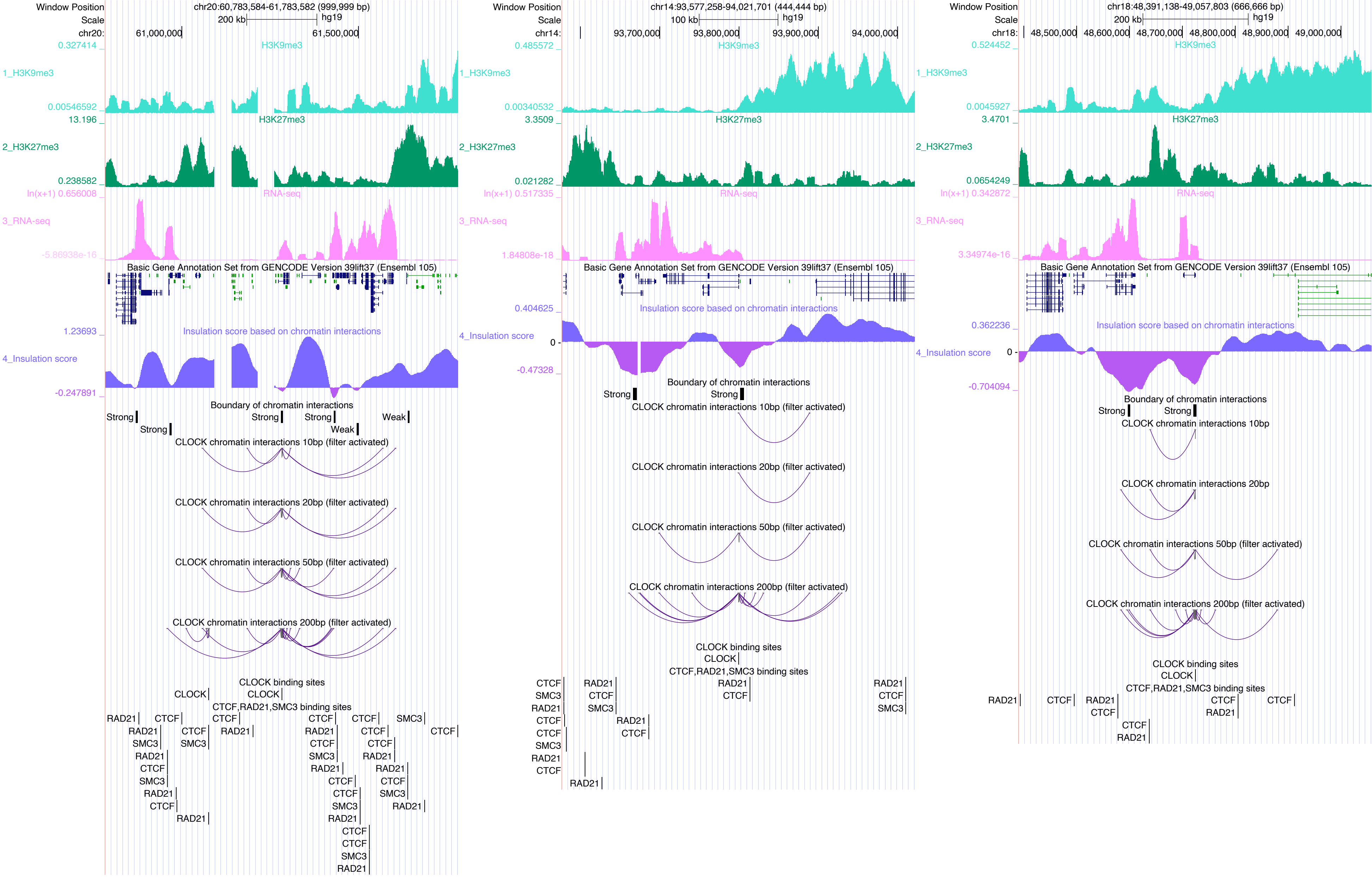
Window Position
Scale
chr20:60,783,584-61,783,582 (999,999 bp)
200 kb
hg19
chr20:
61,000,000
61,500,000
H3K9me3
1_H3K9me3
0.327414
0.00546592
H3K27me3
2_H3K27me3
13.196
0.238582
RNA-seq
3_RNA-seq
ln(x+1) 0.656008
-5.86938e-16
Basic Gene Annotation Set from GENCODE Version 39lift37 (Ensembl 105)
Insulation score based on chromatin interactions
4_Insulation score
1.23693
-0.247891
Boundary of chromatin interactions
Strong
Weak
CLOCK chromatin interactions 10bp (filter activated)
CLOCK chromatin interactions 20bp (filter activated)
CLOCK chromatin interactions 50bp (filter activated)
CLOCK chromatin interactions 200bp (filter activated)
CLOCK binding sites
CLOCK
CTCF,RAD21,SMC3 binding sites
RAD21
CTCF
SMC3
chr14:93,577,258-94,021,701 (444,444 bp)
100 kb
chr14:
93,700,000
93,800,000
93,900,000
94,000,000
0.485572
0.00340532
3.3509
0.021282
ln(x+1) 0.517335
1.84808e-18
0.404625
0
-0.47328
chr18:48,391,138-49,057,803 (666,666 bp)
chr18:
48,500,000
48,600,000
48,700,000
48,800,000
48,900,000
49,000,000
0.524452
0.0045927
3.4701
0.0654249
ln(x+1) 0.342872
3.34974e-16
0.362236
-0.704094
CLOCK chromatin interactions 10bp
CLOCK chromatin interactions 20bp

### CREB5

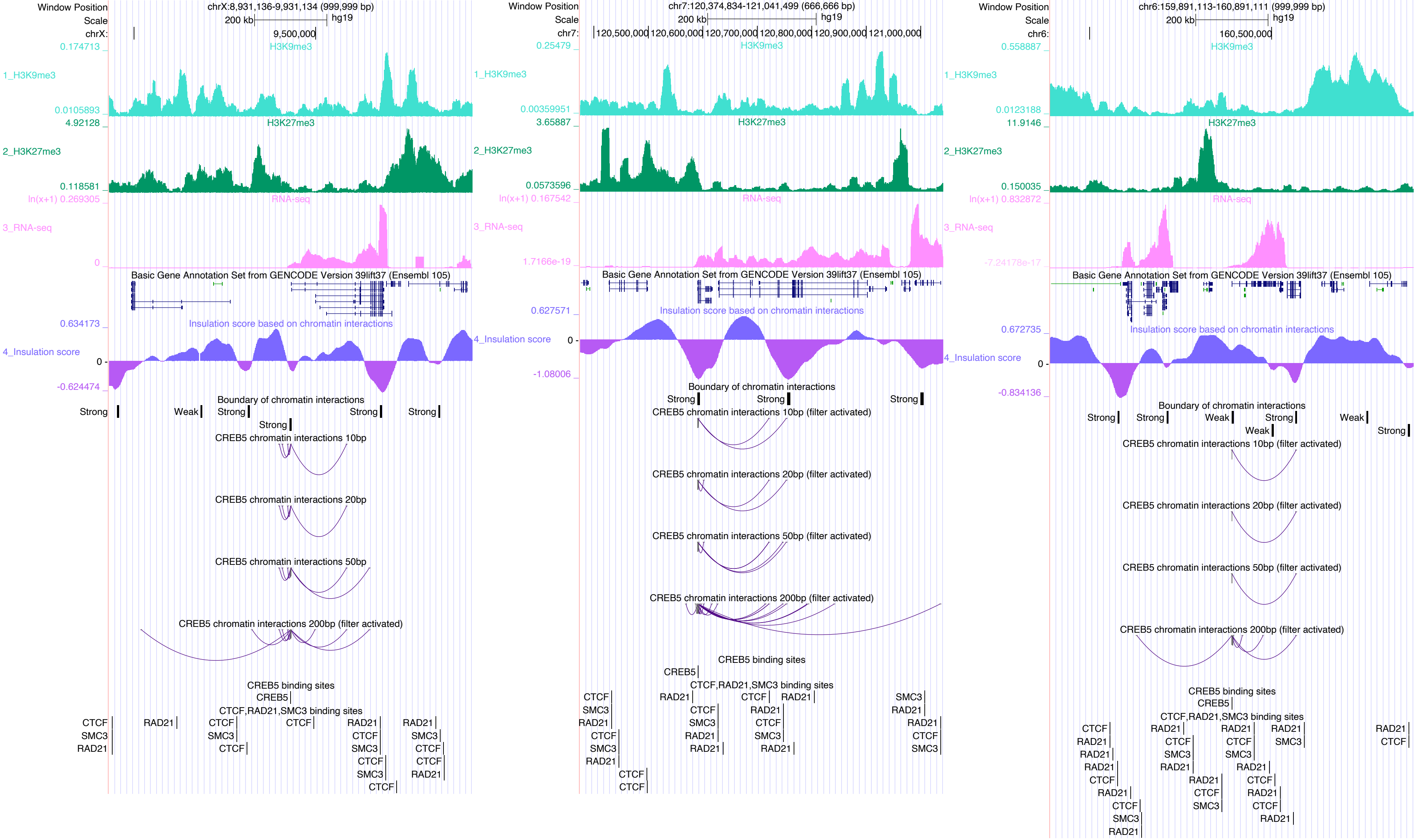

Window Position
Scale
chrX:8,931,136-9,931,134 (999,999 bp)
200 kb
hg19
chrX:
9,500,000
0.174713 _
H3K9me3
1_H3K9me3
0.0105893 _
4.92128 _
H3K27me3
2_H3K27me3
0.118581 _
ln(x+1) 0.269305 _
RNA-seq
3_RNA-seq
0 _
Basic Gene Annotation Set from GENCODE Version 39lift37 (Ensembl 105)
0.634173 _
Insulation score based on chromatin interactions
4_Insulation score
0 -
-0.624474 _
Boundary of chromatin interactions
Strong
Weak
Strong
Strong
Strong
Strong
CREB5 chromatin interactions 10bp
CREB5 chromatin interactions 20bp
CREB5 chromatin interactions 50bp
CREB5 chromatin interactions 200bp (filter activated)
CREB5 binding sites
CREB5
CTCF,RAD21,SMC3 binding sites
CTCF
SMC3
RAD21
Window Position
Scale
chr7:120,374,834-121,041,499 (666,666 bp)
200 kb
hg19
chr7:
120,500,000
120,600,000
120,700,000
120,800,000
120,900,000
121,000,000
0.25479 _
H3K9me3
1_H3K9me3
0.00359951 _
3.65887 _
H3K27me3
2_H3K27me3
0.0573596 _
ln(x+1) 0.167542 _
RNA-seq
3_RNA-seq
1.7166e-19 _
Basic Gene Annotation Set from GENCODE Version 39lift37 (Ensembl 105)
0.627571 _
Insulation score based on chromatin interactions
4_Insulation score
0 -
-1.08006 _
Boundary of chromatin interactions
Strong
Strong
Strong
CREB5 chromatin interactions 10bp (filter activated)
CREB5 chromatin interactions 20bp (filter activated)
CREB5 chromatin interactions 50bp (filter activated)
CREB5 chromatin interactions 200bp (filter activated)
CREB5 binding sites
CREB5
CTCF,RAD21,SMC3 binding sites
Window Position
Scale
chr6:159,891,113-160,891,111 (999,999 bp)
200 kb
hg19
chr6:
160,500,000
0.558887 _
H3K9me3
1_H3K9me3
0.0123188 _
11.9146 _
H3K27me3
2_H3K27me3
0.150035 _
ln(x+1) 0.832872 _
RNA-seq
3_RNA-seq
-7.24178e-17 _
Basic Gene Annotation Set from GENCODE Version 39lift37 (Ensembl 105)
0.672735 _
Insulation score based on chromatin interactions
4_Insulation score
0 -
-0.834136 _
Boundary of chromatin interactions
Strong
Strong
Weak
Strong
Weak
Weak
Strong
CREB5 chromatin interactions 10bp (filter activated)
CREB5 chromatin interactions 20bp (filter activated)
CREB5 chromatin interactions 50bp (filter activated)
CREB5 chromatin interactions 200bp (filter activated)
CREB5 binding sites
CREB5
CTCF,RAD21,SMC3 binding sites

### CTCFL

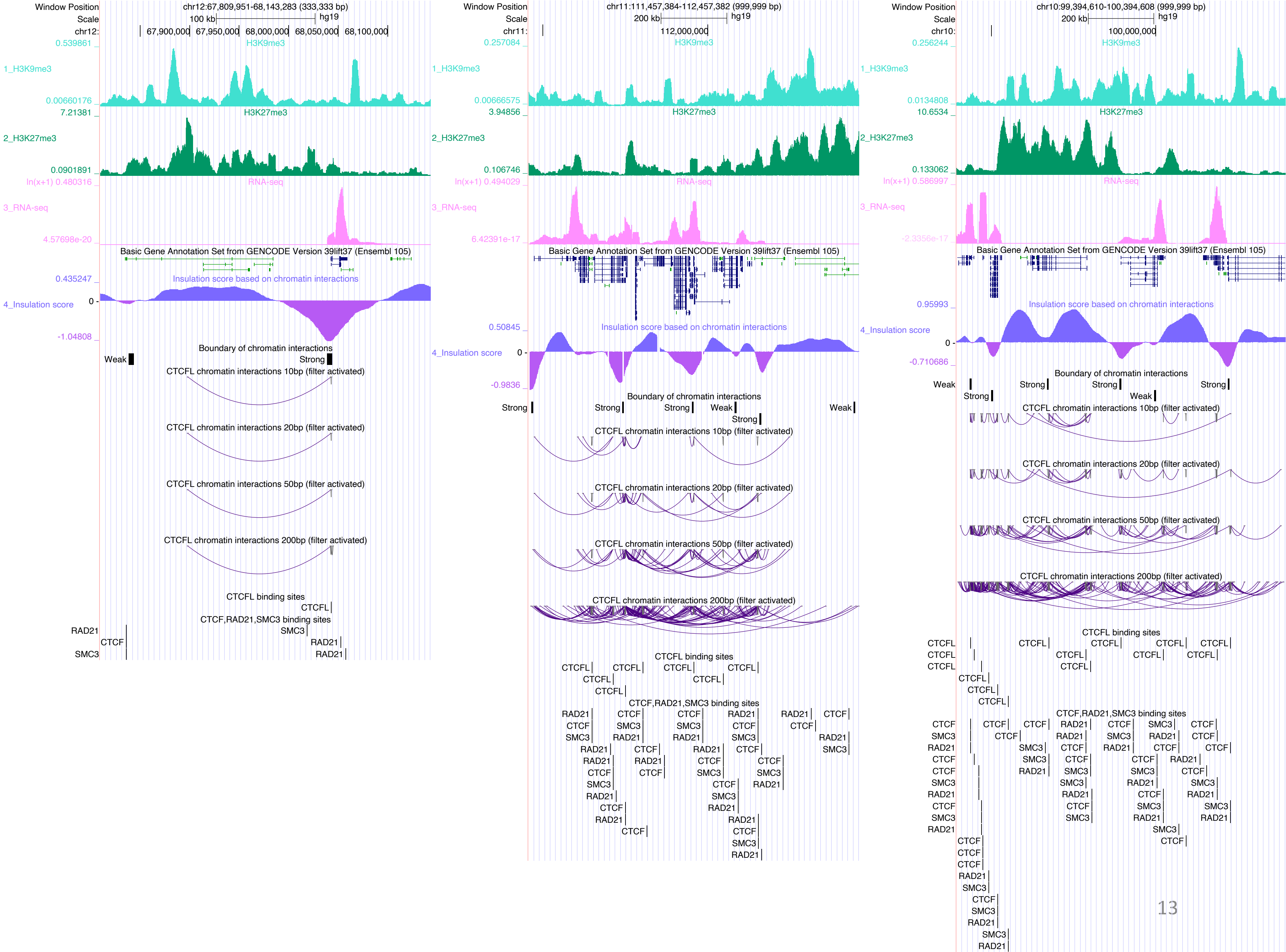

Window Position
chr12:67,809,951-68,143,283 (333,333 bp)
Scale
100 kb
hg19
chr12:
67,900,000
67,950,000
68,000,000
68,050,000
68,100,000
0.539861
H3K9me3
1_H3K9me3
0.00660176
7.21381
H3K27me3
2_H3K27me3
0.0901891
ln(x+1) 0.480316
RNA-seq
3_RNA-seq
4.57698e-20
Basic Gene Annotation Set from GENCODE Version 39lift37 (Ensembl 105)
0.435247
Insulation score based on chromatin interactions
4_Insulation score
0
-1.04808
Boundary of chromatin interactions
Weak
Strong
CTCFL chromatin interactions 10bp (filter activated)
CTCFL chromatin interactions 20bp (filter activated)
CTCFL chromatin interactions 50bp (filter activated)
CTCFL chromatin interactions 200bp (filter activated)
CTCFL binding sites
CTCFL
CTCF,RAD21,SMC3 binding sites
RAD21
CTCF
SMC3
Window Position
chr11:111,457,384-112,457,382 (999,999 bp)
Scale
200 kb
hg19
chr11:
112,000,000
0.257084
H3K9me3
1_H3K9me3
0.00666575
3.94856
H3K27me3
2_H3K27me3
0.106746
ln(x+1) 0.494029
RNA-seq
3_RNA-seq
6.42391e-17
Basic Gene Annotation Set from GENCODE Version 39lift37 (Ensembl 105)
0.50845
Insulation score based on chromatin interactions
4_Insulation score
0
-0.9836
Boundary of chromatin interactions
Strong
Weak
CTCFL chromatin interactions 10bp (filter activated)
CTCFL chromatin interactions 20bp (filter activated)
CTCFL chromatin interactions 50bp (filter activated)
CTCFL chromatin interactions 200bp (filter activated)
CTCFL binding sites
CTCF,RAD21,SMC3 binding sites
Window Position
chr10:99,394,610-100,394,608 (999,999 bp)
Scale
200 kb
hg19
chr10:
100,000,000
0.256244
H3K9me3
1_H3K9me3
0.0134808
10.6534
H3K27me3
2_H3K27me3
0.133062
ln(x+1) 0.586997
RNA-seq
3_RNA-seq
-2.3356e-17
Basic Gene Annotation Set from GENCODE Version 39lift37 (Ensembl 105)
0.95993
Insulation score based on chromatin interactions
4_Insulation score
0
-0.710686
Boundary of chromatin interactions
Weak
Strong
CTCFL chromatin interactions 10bp (filter activated)
CTCFL chromatin interactions 20bp (filter activated)
CTCFL chromatin interactions 50bp (filter activated)
CTCFL chromatin interactions 200bp (filter activated)
CTCFL binding sites
CTCF,RAD21,SMC3 binding sites

### DUX4

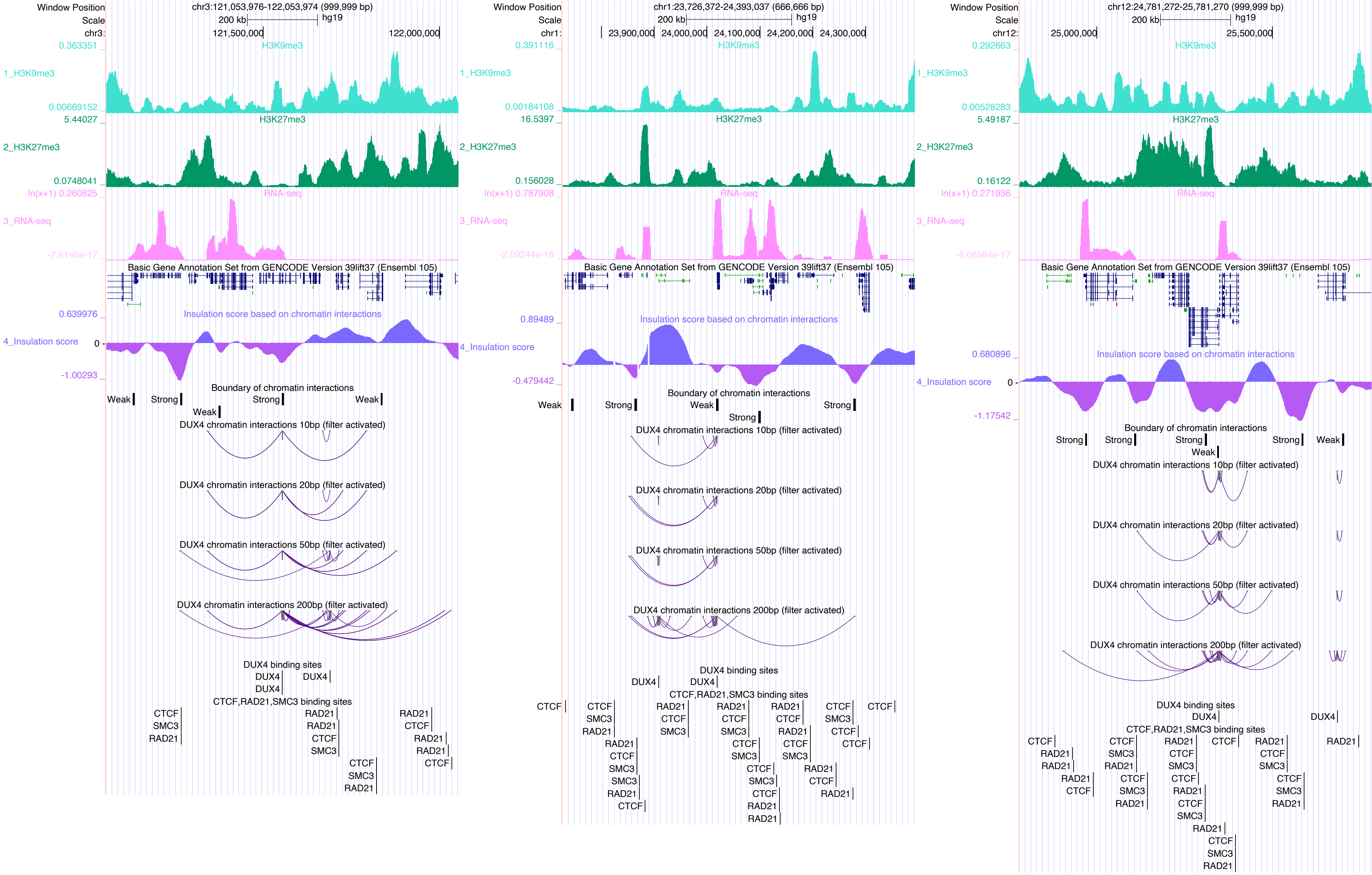

Window Position
Scale
chr3:121,053,976-122,053,974 (999,999 bp)
200 kb
hg19
121,500,000
122,000,000
H3K9me3
1_H3K9me3
H3K27me3
2_H3K27me3
RNA-seq
3_RNA-seq
Basic Gene Annotation Set from GENCODE Version 39lift37 (Ensembl 105)
Insulation score based on chromatin interactions
4_Insulation score
Boundary of chromatin interactions
Weak
Strong
DUX4 chromatin interactions 10bp (filter activated)
DUX4 chromatin interactions 20bp (filter activated)
DUX4 chromatin interactions 50bp (filter activated)
DUX4 chromatin interactions 200bp (filter activated)
DUX4 binding sites
CTCF,RAD21,SMC3 binding sites
chr1:23,726,372-24,393,037 (666,666 bp)
23,900,000
24,000,000
24,100,000
24,200,000
24,300,000
chr12:24,781,272-25,781,270 (999,999 bp)
25,000,000
25,500,000

EBF1

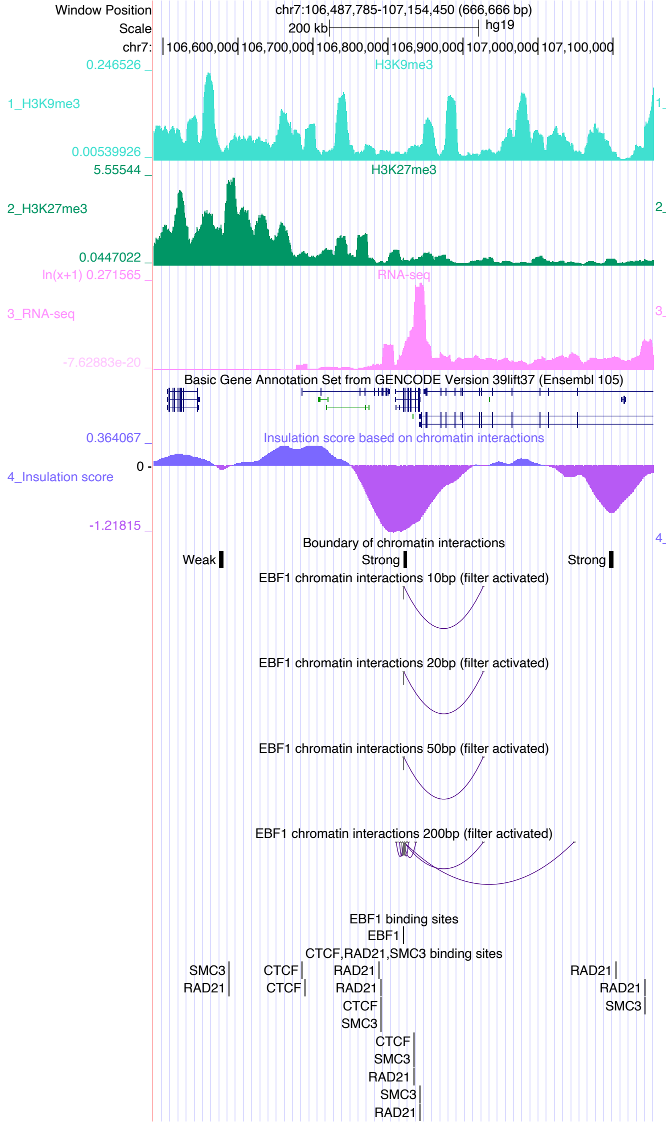

Window Position
chr7:106,487,785-107,154,450 (666,666 bp)
Scale
200 kb
hg19
chr7:
106,600,000
106,700,000
106,800,000
106,900,000
107,000,000
107,100,000
0.246526
H3K9me3
1_H3K9me3
0.00539926
5.55544
H3K27me3
2_H3K27me3
0.0447022
ln(x+1) 0.271565
RNA-seq
3_RNA-seq
-7.62883e-20
Basic Gene Annotation Set from GENCODE Version 39lift37 (Ensembl 105)
0.364067
Insulation score based on chromatin interactions
4_Insulation score
0
-1.21815
Boundary of chromatin interactions
Weak
Strong
Strong
EBF1 chromatin interactions 10bp (filter activated)
EBF1 chromatin interactions 20bp (filter activated)
EBF1 chromatin interactions 50bp (filter activated)
EBF1 chromatin interactions 200bp (filter activated)
EBF1 binding sites
EBF1
CTCF,RAD21,SMC3 binding sites
SMC3
RAD21
CTCF
CTCF
RAD21
RAD21
CTCF
SMC3
CTCF
SMC3
RAD21
SMC3
RAD21
RAD21
RAD21
SMC3


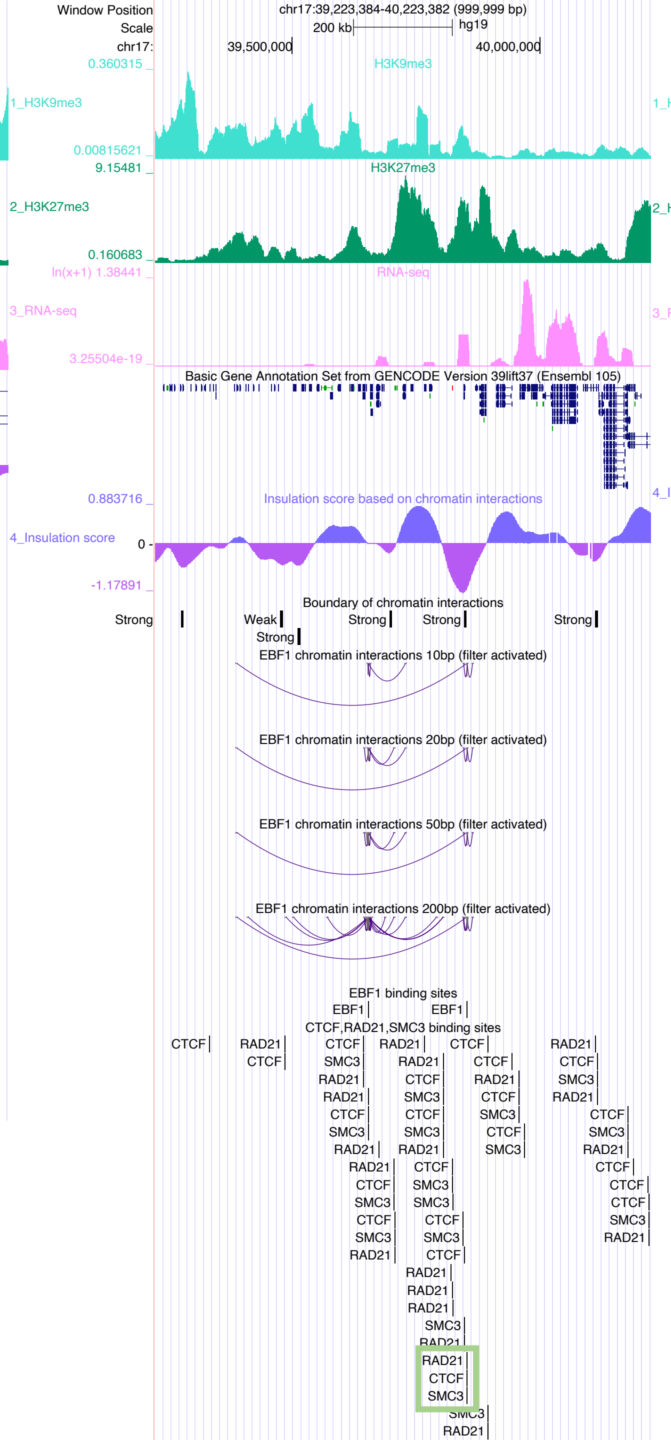

Window Position
chr17:39,223,384-40,223,382 (999,999 bp)
Scale
200 kb
hg19
chr17:
39,500,000
40,000,000
0.360315
H3K9me3
1_H3K9me3
0.00815621
9.15481
H3K27me3
2_H3K27me3
0.160683
ln(x+1) 1.38441
RNA-seq
3_RNA-seq
3.25504e-19
Basic Gene Annotation Set from GENCODE Version 39lift37 (Ensembl 105)
0.883716
Insulation score based on chromatin interactions
4_Insulation score
0
-1.17891
Boundary of chromatin interactions
Strong
Weak
Strong
Strong
Strong
Strong
EBF1 chromatin interactions 10bp (filter activated)
EBF1 chromatin interactions 20bp (filter activated)
EBF1 chromatin interactions 50bp (filter activated)
EBF1 chromatin interactions 200bp (filter activated)
EBF1 binding sites
EBF1
EBF1
CTCF,RAD21,SMC3 binding sites
CTCF
RAD21
CTCF
RAD21
CTCF
RAD21
CTCF


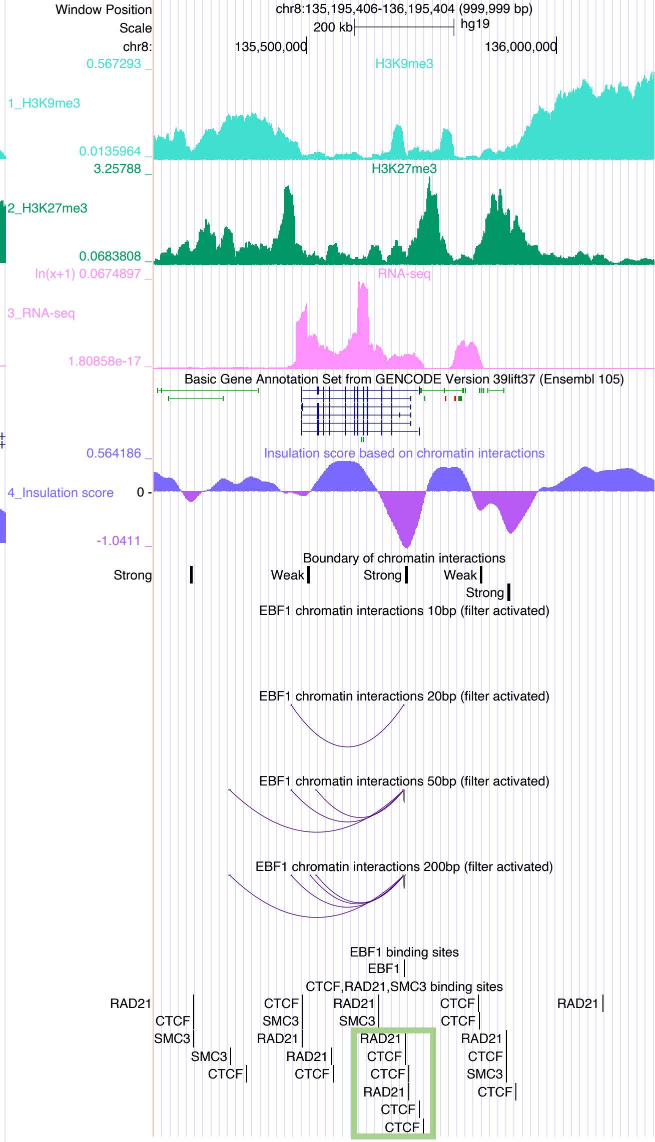

Window Position
chr8:135,195,406-136,195,404 (999,999 bp)
Scale
200 kb
hg19
chr8:
135,500,000
136,000,000
0.567293
H3K9me3
1_H3K9me3
0.0135964
3.25788
H3K27me3
2_H3K27me3
0.0683808
ln(x+1) 0.0674897
RNA-seq
3_RNA-seq
1.80858e-17
Basic Gene Annotation Set from GENCODE Version 39lift37 (Ensembl 105)
0.564186
Insulation score based on chromatin interactions
4_Insulation score
0
-1.0411
Boundary of chromatin interactions
Strong
Weak
Strong
Weak
Strong
EBF1 chromatin interactions 10bp (filter activated)
EBF1 chromatin interactions 20bp (filter activated)
EBF1 chromatin interactions 50bp (filter activated)
EBF1 chromatin interactions 200bp (filter activated)
EBF1 binding sites
EBF1
CTCF,RAD21,SMC3 binding sites
RAD21
CTCF
SMC3
CTCF
SMC3
RAD21
CTCF
RAD21
SMC3
CTCF
RAD21
CTCF
CTCF
RAD21
CTCF
CTCF

### EGR1

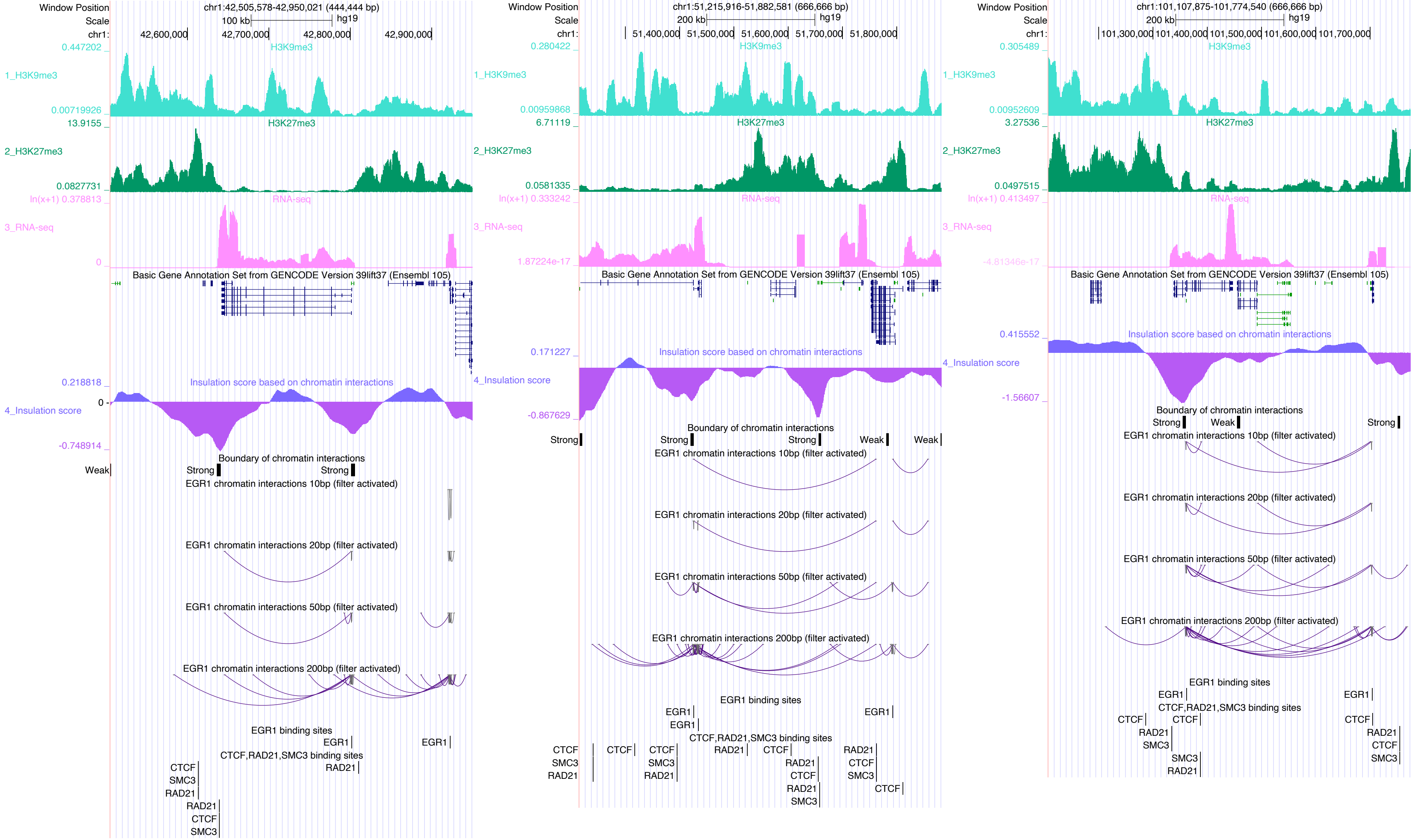

Window Position
Scale
chr1:
chr1:42,505,578-42,950,021 (444,444 bp)
100 kb
hg19
42,600,000
42,700,000
42,800,000
42,900,000
1_H3K9me3
H3K9me3
0.447202
0.00719926
2_H3K27me3
H3K27me3
13.9155
0.0827731
3_RNA-seq
RNA-seq
ln(x+1) 0.378813
0
Basic Gene Annotation Set from GENCODE Version 39lift37 (Ensembl 105)
4_Insulation score
Insulation score based on chromatin interactions
0.218818
0
-0.748914
Boundary of chromatin interactions
Weak
Strong
Strong
EGR1 chromatin interactions 10bp (filter activated)
EGR1 chromatin interactions 20bp (filter activated)
EGR1 chromatin interactions 50bp (filter activated)
EGR1 chromatin interactions 200bp (filter activated)
EGR1 binding sites
EGR1
EGR1
CTCF,RAD21,SMC3 binding sites
CTCF
SMC3
RAD21
RAD21
CTCF
SMC3
RAD21
chr1:51,215,916-51,882,581 (666,666 bp)
200 kb
hg19
51,400,000
51,500,000
51,600,000
51,700,000
51,800,000
0.280422
0.00959868
6.71119
0.0581335
ln(x+1) 0.333242
1.87224e-17
0.171227
-0.867629
Strong
Strong
Strong
Weak
Weak
chr1:101,107,875-101,774,540 (666,666 bp)
101,300,000
101,400,000
101,500,000
101,600,000
101,700,000
0.305489
0.00952609
3.27536
0.0497515
ln(x+1) 0.413497
-4.81346e-17
0.415552
-1.56607
Strong
Weak
Strong

### EGR2

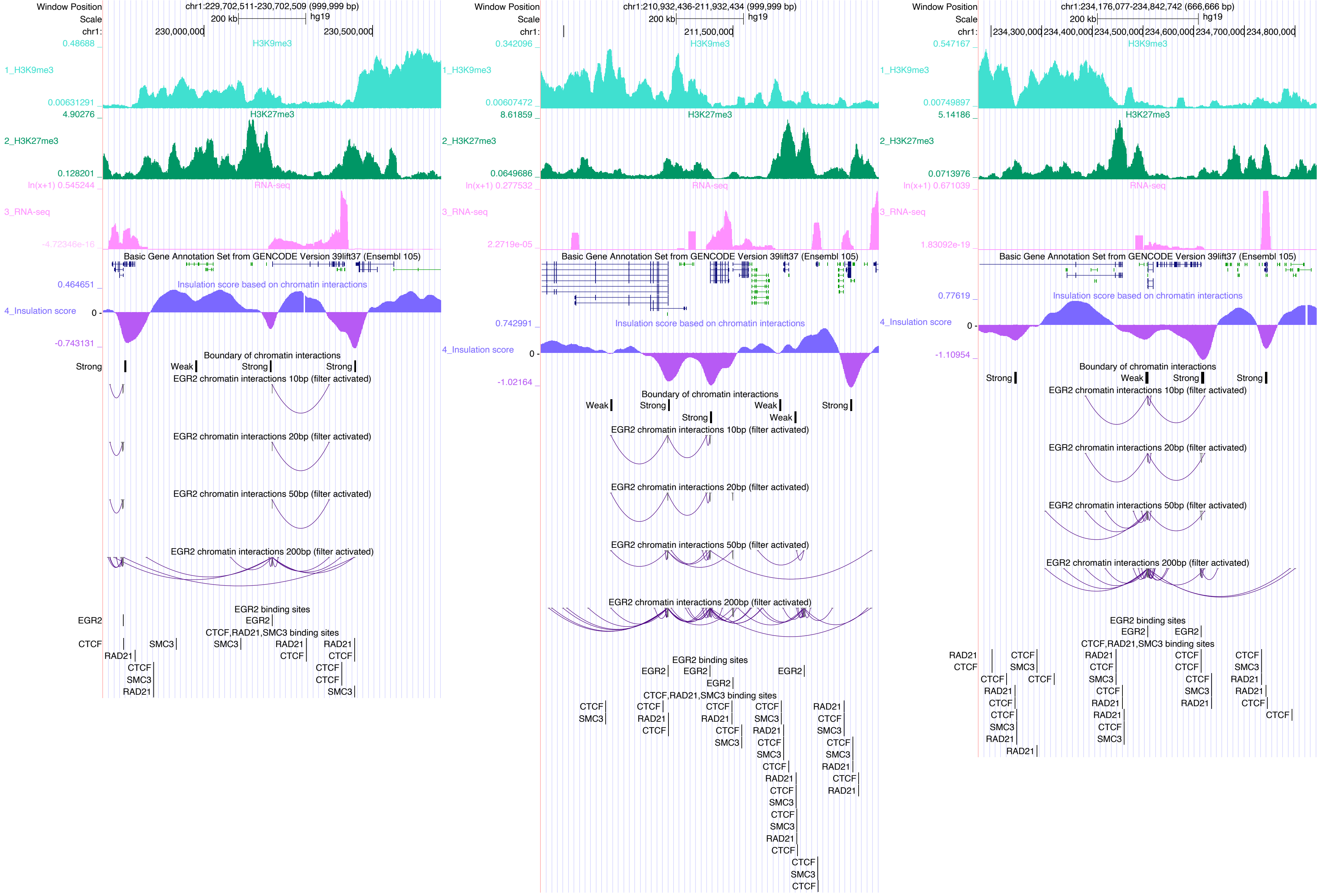

Window Position
Scale
chr1:
chr1:229,702,511-230,702,509 (999,999 bp)
200 kb
hg19
230,000,000
230,500,000
0.48688
H3K9me3
1_H3K9me3
0.00631291
4.90276
H3K27me3
2_H3K27me3
0.128201
ln(x+1) 0.545244
RNA-seq
3_RNA-seq
-4.72346e-16
Basic Gene Annotation Set from GENCODE Version 39lift37 (Ensembl 105)
0.464651
Insulation score based on chromatin interactions
4_Insulation score
0
-0.743131
Boundary of chromatin interactions
Strong
Weak
Strong
Strong
EGR2 chromatin interactions 10bp (filter activated)
EGR2 chromatin interactions 20bp (filter activated)
EGR2 chromatin interactions 50bp (filter activated)
EGR2 chromatin interactions 200bp (filter activated)
EGR2 binding sites
EGR2
EGR2
CTCF,RAD21,SMC3 binding sites
CTCF
RAD21
SMC3
SMC3
RAD21
CTCF
RAD21
CTCF
CTCF
SMC3
RAD21
CTCF
CTCF
SMC3
Window Position
Scale
chr1:
chr1:210,932,436-211,932,434 (999,999 bp)
200 kb
hg19
211,500,000
0.342096
H3K9me3
1_H3K9me3
0.00607472
8.61859
H3K27me3
2_H3K27me3
0.0649686
ln(x+1) 0.277532
RNA-seq
3_RNA-seq
2.2719e-05
Basic Gene Annotation Set from GENCODE Version 39lift37 (Ensembl 105)
0.742991
Insulation score based on chromatin interactions
4_Insulation score
0
-1.02164
Boundary of chromatin interactions
Weak
Strong
Weak
Strong
Strong
Weak
EGR2 chromatin interactions 10bp (filter activated)
EGR2 chromatin interactions 20bp (filter activated)
EGR2 chromatin interactions 50bp (filter activated)
EGR2 chromatin interactions 200bp (filter activated)
EGR2 binding sites
EGR2
EGR2
EGR2
EGR2
CTCF,RAD21,SMC3 binding sites
CTCF
SMC3
CTCF
RAD21
CTCF
CTCF
RAD21
CTCF
SMC3
CTCF
SMC3
RAD21
CTCF
SMC3
CTCF
RAD21
CTCF
SMC3
CTCF
SMC3
RAD21
CTCF
CTCF
SMC3
CTCF
RAD21
CTCF
SMC3
RAD21
CTCF
RAD21
Window Position
Scale
chr1:
chr1:234,176,077-234,842,742 (666,666 bp)
200 kb
hg19
234,300,000
234,400,000
234,500,000
234,600,000
234,700,000
234,800,000
0.547167
H3K9me3
1_H3K9me3
0.00749897
5.14186
H3K27me3
2_H3K27me3
0.0713976
ln(x+1) 0.671039
RNA-seq
3_RNA-seq
1.83092e-19
Basic Gene Annotation Set from GENCODE Version 39lift37 (Ensembl 105)
0.77619
Insulation score based on chromatin interactions
4_Insulation score
0
-1.10954
Boundary of chromatin interactions
Strong
Weak
Strong
Strong
EGR2 chromatin interactions 10bp (filter activated)
EGR2 chromatin interactions 20bp (filter activated)
EGR2 chromatin interactions 50bp (filter activated)
EGR2 chromatin interactions 200bp (filter activated)
EGR2 binding sites
EGR2
EGR2
CTCF,RAD21,SMC3 binding sites
RAD21
CTCF
CTCF
SMC3
CTCF
RAD21
CTCF
CTCF
SMC3
RAD21
CTCF
RAD21
CTCF
SMC3
RAD21
RAD21
CTCF
SMC3
CTCF
CTCF
CTCF
SMC3
RAD21
CTCF
SMC3
RAD21
RAD21
CTCF
CTCF

### EGR3

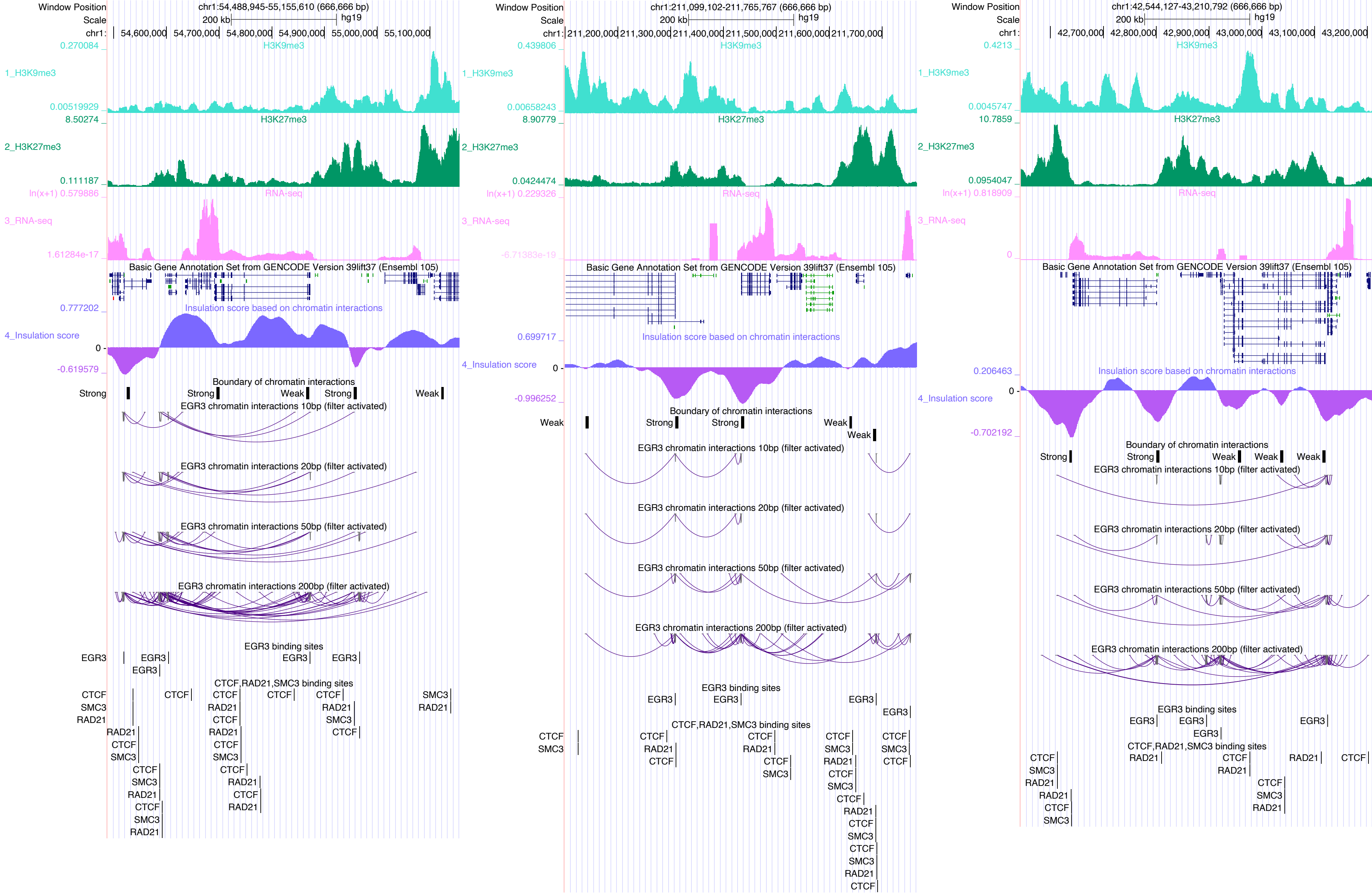
Window Position
Scale
chr1:
chr1:54,488,945-55,155,610 (666,666 bp)
200 kb
hg19
54,600,000
54,700,000
54,800,000
54,900,000
55,000,000
55,100,000
0.270084
1_H3K9me3
0.00519929
H3K9me3
8.50274
2_H3K27me3
0.111187
H3K27me3
ln(x+1) 0.579886
3_RNA-seq
1.61284e-17
RNA-seq
Basic Gene Annotation Set from GENCODE Version 39lift37 (Ensembl 105)
0.777202
4_Insulation score
0
-0.619579
Insulation score based on chromatin interactions
Boundary of chromatin interactions
Strong
Strong
Weak
Strong
Weak
EGR3 chromatin interactions 10bp (filter activated)
EGR3 chromatin interactions 20bp (filter activated)
EGR3 chromatin interactions 50bp (filter activated)
EGR3 chromatin interactions 200bp (filter activated)
EGR3 binding sites
EGR3
EGR3
EGR3
EGR3
EGR3
CTCF,RAD21,SMC3 binding sites
CTCF
SMC3
RAD21
chr1:211,099,102-211,765,767 (666,666 bp)
211,200,000
211,300,000
211,400,000
211,500,000
211,600,000
211,700,000
0.439806
0.00658243
8.90779
0.0424474
ln(x+1) 0.229326
-6.71383e-19
0.699717
0
-0.996252
Weak
Strong
Strong
Weak
Weak
chr1:42,544,127-43,210,792 (666,666 bp)
42,700,000
42,800,000
42,900,000
43,000,000
43,100,000
43,200,000
0.4213
0.0045747
10.7859
0.0954047
ln(x+1) 0.818909
0
0.206463
0
-0.702192
Strong
Strong
Weak
Weak
Weak

ELK4

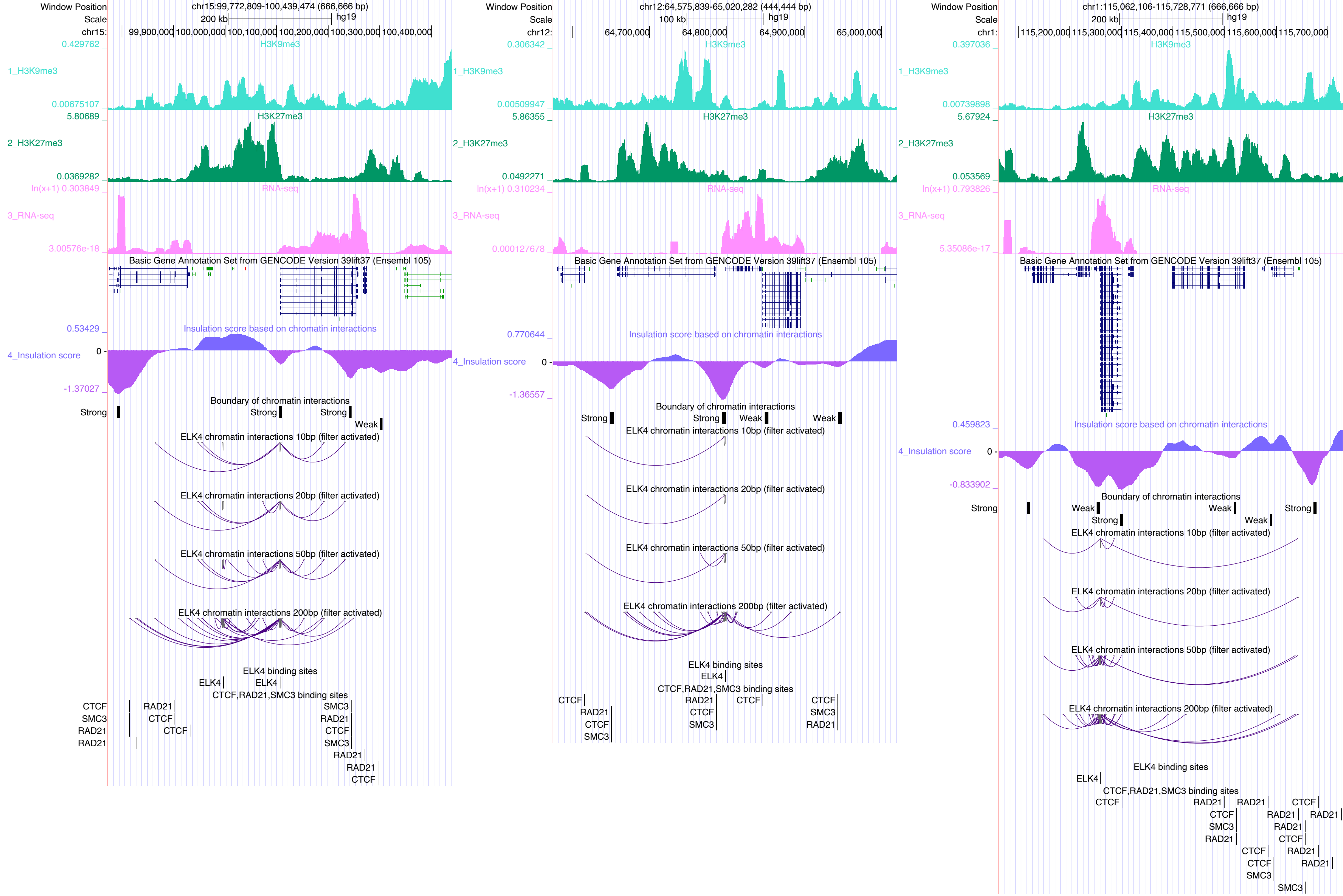
Window Position
chr15:99,772,809-100,439,474 (666,666 bp)
Scale
200 kb
hg19
chr15:
99,900,000
100,000,000
100,100,000
100,200,000
100,300,000
100,400,000
1_H3K9me3
H3K9me3
2_H3K27me3
H3K27me3
3_RNA-seq
RNA-seq
Basic Gene Annotation Set from GENCODE Version 39lift37 (Ensembl 105)
4_Insulation score
Insulation score based on chromatin interactions
Boundary of chromatin interactions
Strong
Weak
ELK4 chromatin interactions 10bp (filter activated)
ELK4 chromatin interactions 20bp (filter activated)
ELK4 chromatin interactions 50bp (filter activated)
ELK4 chromatin interactions 200bp (filter activated)
ELK4 binding sites
ELK4
CTCF,RAD21,SMC3 binding sites
CTCF
SMC3
RAD21
Window Position
chr12:64,575,839-65,020,282 (444,444 bp)
100 kb
chr12:
64,700,000
64,800,000
64,900,000
65,000,000
Window Position
chr1:115,062,106-115,728,771 (666,666 bp)
200 kb
chr1:
115,200,000
115,300,000
115,400,000
115,500,000
115,600,000
115,700,000

# EP300

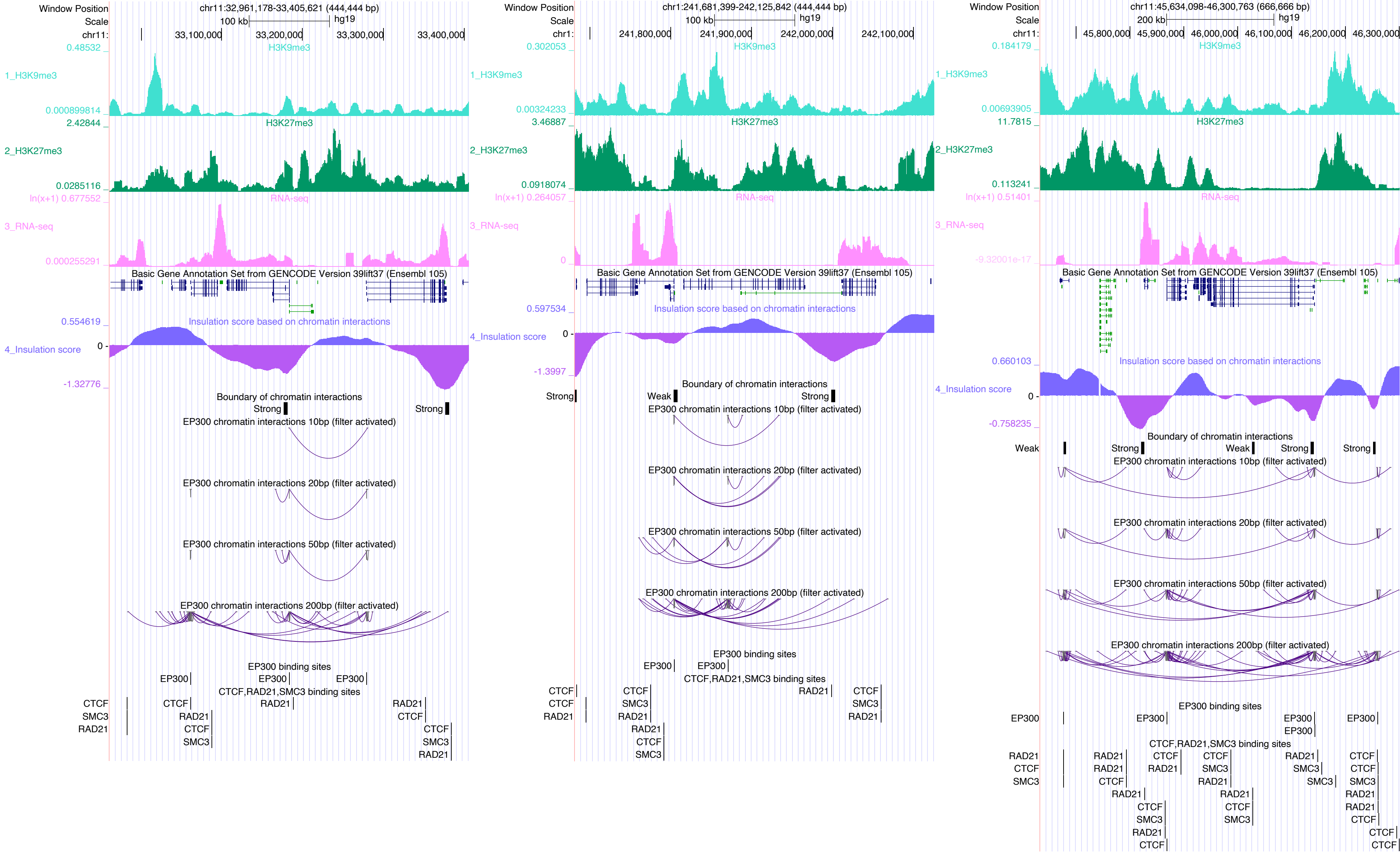

Window Position
Scale
chr11:32,961,178-33,405,621 (444,444 bp)
100 kb
hg19
33,100,000
33,200,000
33,300,000
33,400,000
H3K9me3
1_H3K9me3
H3K27me3
2_H3K27me3
RNA-seq
3_RNA-seq
Basic Gene Annotation Set from GENCODE Version 39lift37 (Ensembl 105)
Insulation score based on chromatin interactions
4_Insulation score
Boundary of chromatin interactions
Strong
Strong
EP300 chromatin interactions 10bp (filter activated)
EP300 chromatin interactions 20bp (filter activated)
EP300 chromatin interactions 50bp (filter activated)
EP300 chromatin interactions 200bp (filter activated)
EP300 binding sites
EP300
EP300
EP300
CTCF,RAD21,SMC3 binding sites
CTCF
SMC3
RAD21
chr1:241,681,399-242,125,842 (444,444 bp)
100 kb
hg19
241,800,000
241,900,000
242,000,000
242,100,000
Weak
Strong
Strong
chr11:45,634,098-46,300,763 (666,666 bp)
200 kb
hg19
45,800,000
45,900,000
46,000,000
46,100,000
46,200,000
46,300,000

### EPAS1

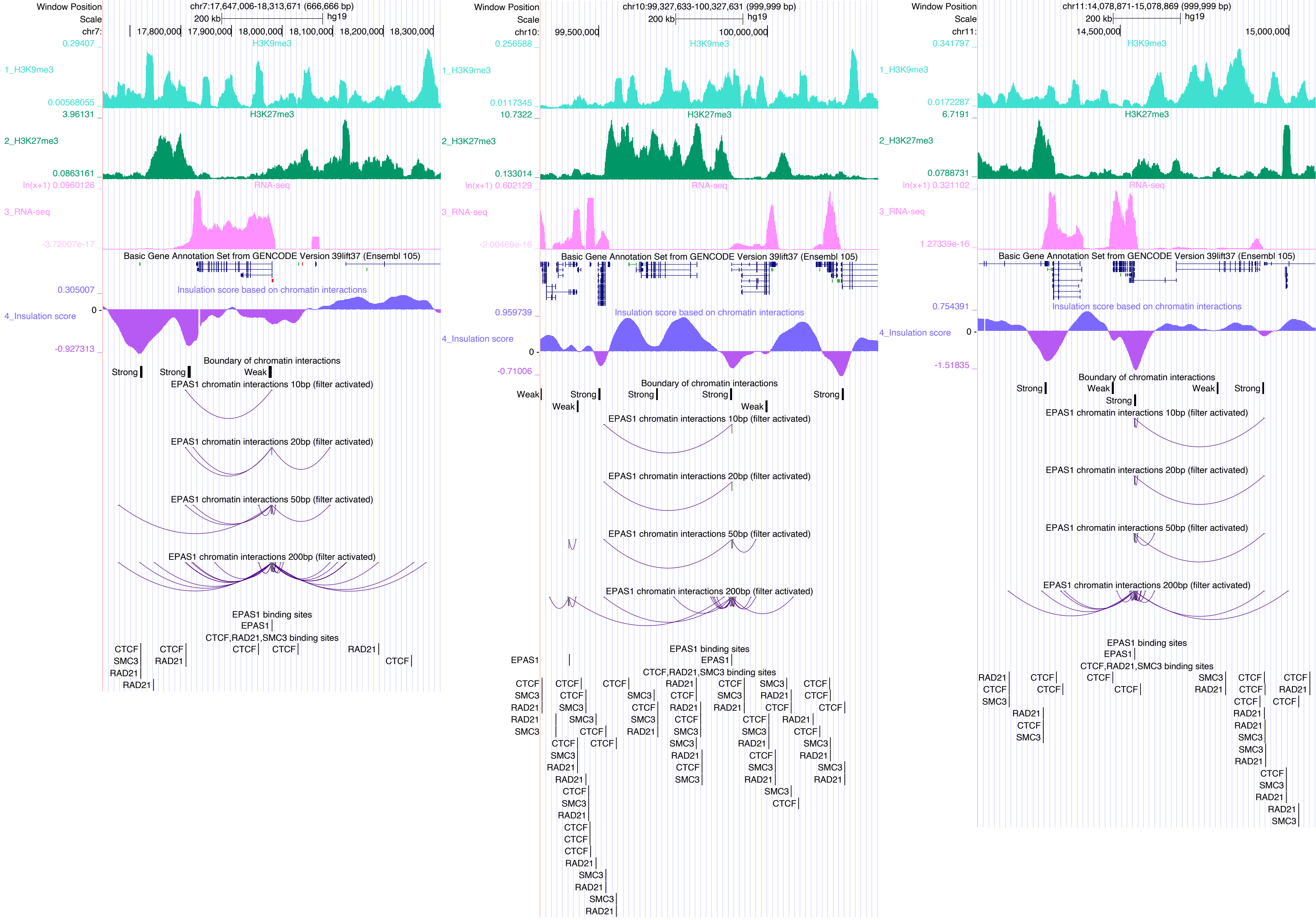
Window Position chr7:17,647,006-18,313,671 (666,666 bp)
Scale 200 kb hg19
chr7: 17,800,000 17,900,000 18,000,000 18,100,000 18,200,000 18,300,000
1_H3K9me3
2_H3K27me3
3_RNA-seq
Basic Gene Annotation Set from GENCODE Version 39lift37 (Ensembl 105)
4_Insulation score
Insulation score based on chromatin interactions
Boundary of chromatin interactions
EPAS1 chromatin interactions 10bp (filter activated)
EPAS1 chromatin interactions 20bp (filter activated)
EPAS1 chromatin interactions 50bp (filter activated)
EPAS1 chromatin interactions 200bp (filter activated)
EPAS1 binding sites
CTCF,RAD21,SMC3 binding sites
Window Position chr10:99,327,633-100,327,631 (999,999 bp)
chr10: 99,500,000 100,000,000
Window Position chr11:14,078,871-15,078,869 (999,999 bp)
chr11: 14,500,000 15,000,000

### ETV1

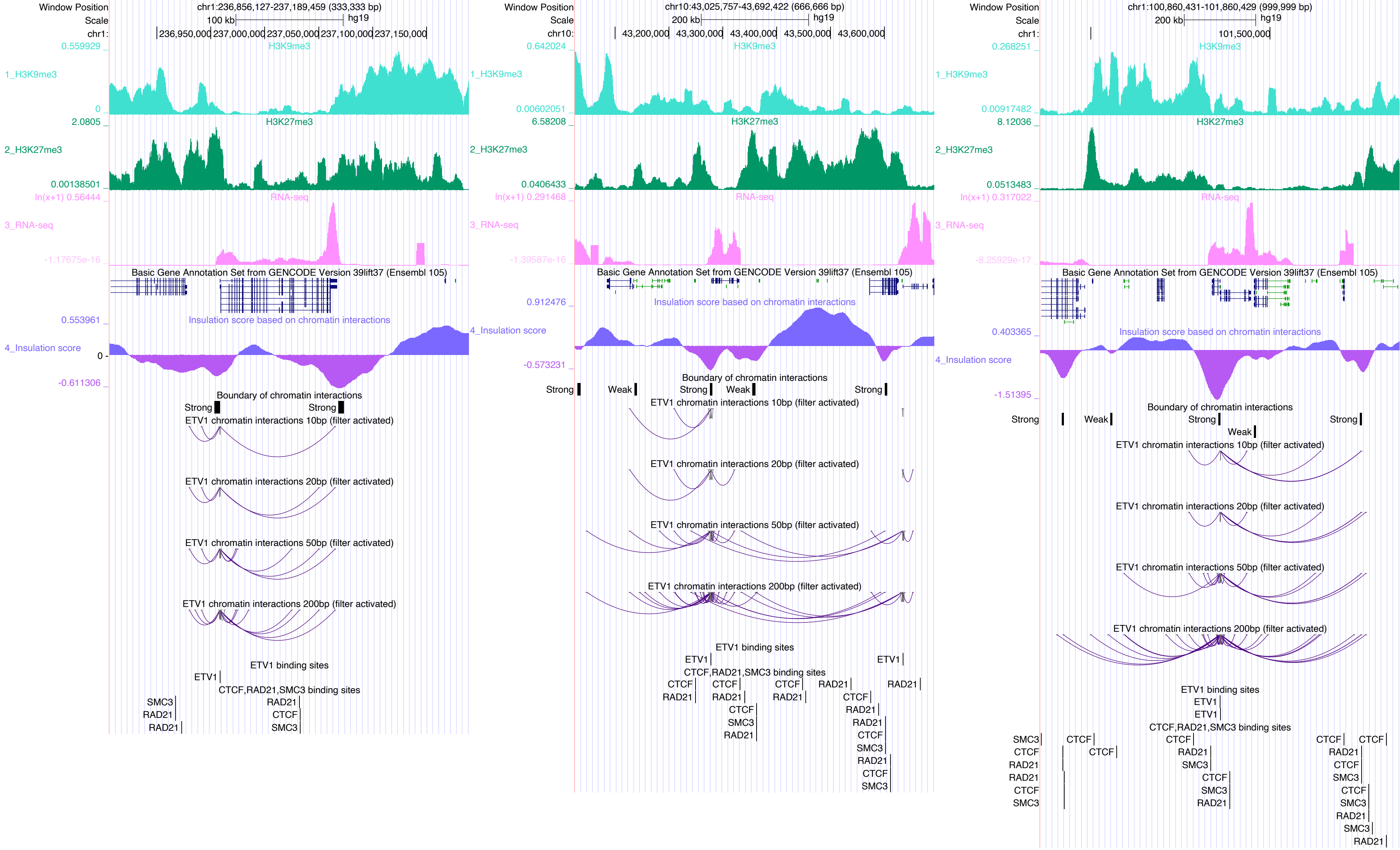

Window Position
Scale
chr1:
chr1:236,856,127-237,189,459 (333,333 bp)
100 kb
hg19
236,950,000 237,000,000 237,050,000 237,100,000 237,150,000
0.559929
1_H3K9me3
H3K9me3
0
2.0805
2_H3K27me3
H3K27me3
0.00138501
ln(x+1) 0.56444
3_RNA-seq
RNA-seq
-1.17675e-16
Basic Gene Annotation Set from GENCODE Version 39lift37 (Ensembl 105)
0.553961
4_Insulation score
Insulation score based on chromatin interactions
0
-0.611306
Boundary of chromatin interactions
Strong
Strong
ETV1 chromatin interactions 10bp (filter activated)
ETV1 chromatin interactions 20bp (filter activated)
ETV1 chromatin interactions 50bp (filter activated)
ETV1 chromatin interactions 200bp (filter activated)
ETV1 binding sites
ETV1
CTCF,RAD21,SMC3 binding sites
SMC3
RAD21
RAD21
RAD21
CTCF
SMC3
Window Position
Scale
chr10:
chr10:43,025,757-43,692,422 (666,666 bp)
200 kb
hg19
43,200,000 43,300,000 43,400,000 43,500,000 43,600,000
0.642024
1_H3K9me3
H3K9me3
0.00602051
6.58208
2_H3K27me3
H3K27me3
0.0406433
ln(x+1) 0.291468
3_RNA-seq
RNA-seq
-1.39587e-16
Basic Gene Annotation Set from GENCODE Version 39lift37 (Ensembl 105)
0.912476
4_Insulation score
Insulation score based on chromatin interactions
-0.573231
Boundary of chromatin interactions
Strong
Weak
Strong
Weak
Strong
ETV1 chromatin interactions 10bp (filter activated)
ETV1 chromatin interactions 20bp (filter activated)
ETV1 chromatin interactions 50bp (filter activated)
ETV1 chromatin interactions 200bp (filter activated)
ETV1 binding sites
ETV1
ETV1
CTCF,RAD21,SMC3 binding sites
CTCF
CTCF
CTCF
RAD21
RAD21
RAD21
RAD21
RAD21
RAD21
CTCF
CTCF
RAD21
SMC3
RAD21
RAD21
CTCF
SMC3
RAD21
CTCF
SMC3
Window Position
Scale
chr1:
chr1:100,860,431-101,860,429 (999,999 bp)
200 kb
hg19
101,500,000
0.268251
1_H3K9me3
H3K9me3
0.00917482
8.12036
2_H3K27me3
H3K27me3
0.0513483
ln(x+1) 0.317022
3_RNA-seq
RNA-seq
-8.25929e-17
Basic Gene Annotation Set from GENCODE Version 39lift37 (Ensembl 105)
0.403365
4_Insulation score
Insulation score based on chromatin interactions
-1.51395
Boundary of chromatin interactions
Strong
Weak
Strong
Weak
Strong
ETV1 chromatin interactions 10bp (filter activated)
ETV1 chromatin interactions 20bp (filter activated)
ETV1 chromatin interactions 50bp (filter activated)
ETV1 chromatin interactions 200bp (filter activated)
ETV1 binding sites
ETV1
ETV1
CTCF,RAD21,SMC3 binding sites
SMC3
CTCF
RAD21
RAD21
CTCF
SMC3
CTCF
CTCF
CTCF
RAD21
SMC3
CTCF
SMC3
RAD21
CTCF
CTCF
RAD21
CTCF
SMC3
CTCF
SMC3
RAD21
SMC3
RAD21

FOS

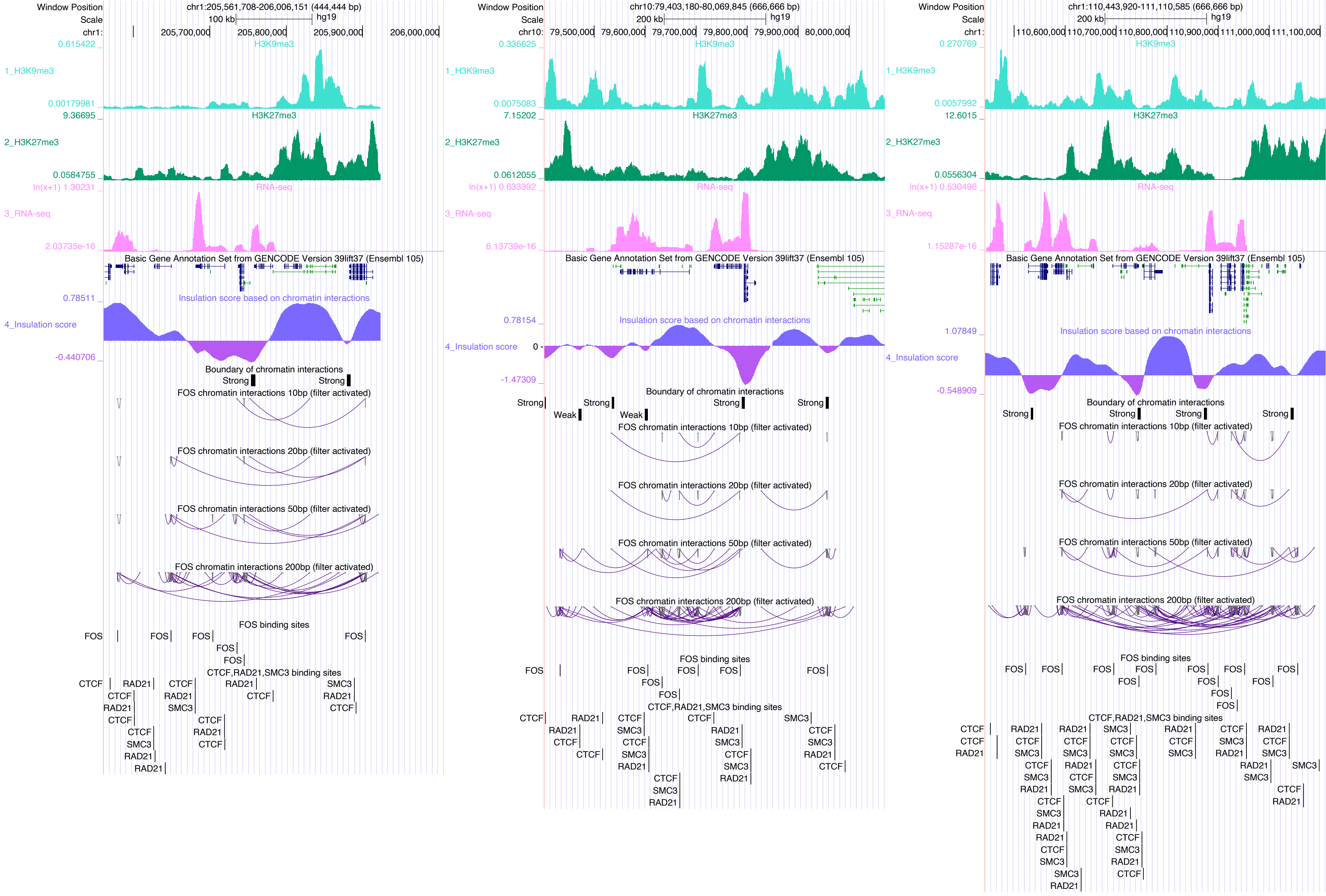

Window Position
Scale
chr1:205,561,708-206,006,151 (444,444 bp)
100 kb
hg19
chr1:
205,700,000
205,800,000
205,900,000
206,000,000
0.615422
H3K9me3
1_H3K9me3
0.00179981
9.36695
H3K27me3
2_H3K27me3
0.0584755
ln(x+1) 1.30231
RNA-seq
3_RNA-seq
2.03735e-16
Basic Gene Annotation Set from GENCODE Version 39lift37 (Ensembl 105)
0.78511
Insulation score based on chromatin interactions
4_Insulation score
-0.440706
Boundary of chromatin interactions
Strong
Strong
FOS chromatin interactions 10bp (filter activated)
FOS chromatin interactions 20bp (filter activated)
FOS chromatin interactions 50bp (filter activated)
FOS chromatin interactions 200bp (filter activated)
FOS binding sites
FOS
CTCF,RAD21,SMC3 binding sites
CTCF
RAD21
SMC3
chr10:79,403,180-80,069,845 (666,666 bp)
200 kb
chr10:
79,500,000
79,600,000
79,700,000
79,800,000
79,900,000
80,000,000
0.336625
0.0075083
7.15202
0.0612055
ln(x+1) 0.633392
6.13739e-16
0.78154
0
-1.47309
Weak
chr1:110,443,920-111,110,585 (666,666 bp)
110,600,000
110,700,000
110,800,000
110,900,000
111,000,000
111,100,000
0.270769
0.0057992
12.6015
0.0556304
ln(x+1) 0.530496
1.15287e-16
1.07849
-0.548909

### FOSL2

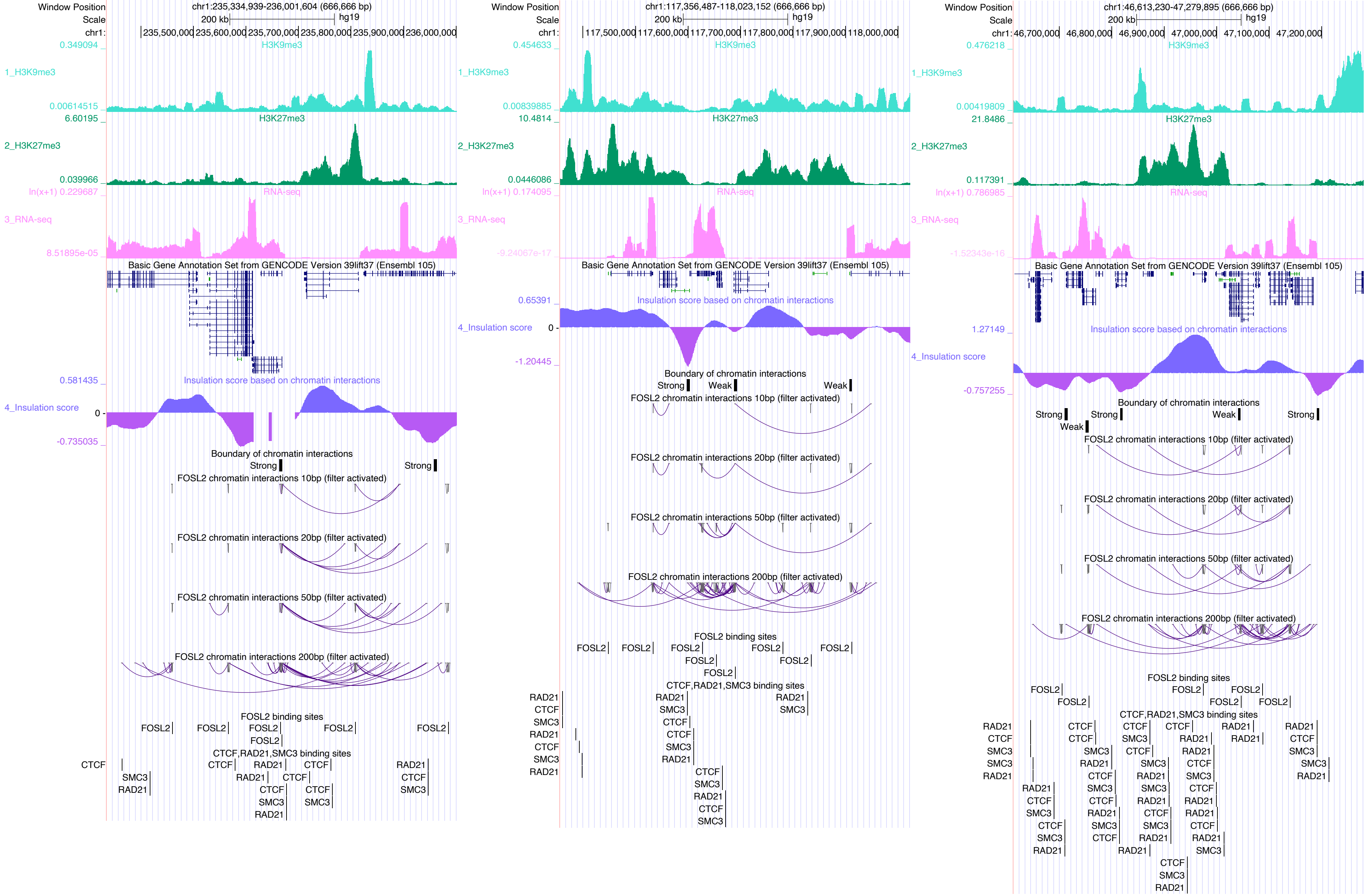
Window Position
Scale
chr1:
chr1:235,334,939-236,001,604 (666,666 bp)
200 kb
hg19
235,500,000
235,600,000
235,700,000
235,800,000
235,900,000
236,000,000
0.349094
1_H3K9me3
0.00614515
H3K9me3
6.60195
2_H3K27me3
0.039966
H3K27me3
ln(x+1) 0.229687
3_RNA-seq
8.51895e-05
RNA-seq
Basic Gene Annotation Set from GENCODE Version 39lift37 (Ensembl 105)
0.581435
4_Insulation score
0
-0.735035
Insulation score based on chromatin interactions
Boundary of chromatin interactions
Strong
Strong
FOSL2 chromatin interactions 10bp (filter activated)
FOSL2 chromatin interactions 20bp (filter activated)
FOSL2 chromatin interactions 50bp (filter activated)
FOSL2 chromatin interactions 200bp (filter activated)
FOSL2 binding sites
FOSL2
FOSL2
FOSL2
FOSL2
FOSL2
FOSL2
CTCF,RAD21,SMC3 binding sites
CTCF
SMC3
RAD21
CTCF
RAD21
RAD21
CTCF
CTCF
CTCF
SMC3
RAD21
CTCF
SMC3
RAD21
CTCF
SMC3
Window Position
Scale
chr1:
chr1:117,356,487-118,023,152 (666,666 bp)
200 kb
hg19
117,500,000
117,600,000
117,700,000
117,800,000
117,900,000
118,000,000
0.454633
1_H3K9me3
0.00839885
H3K9me3
10.4814
2_H3K27me3
0.0446086
H3K27me3
ln(x+1) 0.174095
3_RNA-seq
-9.24067e-17
RNA-seq
Basic Gene Annotation Set from GENCODE Version 39lift37 (Ensembl 105)
0.65391
4_Insulation score
0
-1.20445
Insulation score based on chromatin interactions
Boundary of chromatin interactions
Strong
Weak
Weak
FOSL2 chromatin interactions 10bp (filter activated)
FOSL2 chromatin interactions 20bp (filter activated)
FOSL2 chromatin interactions 50bp (filter activated)
FOSL2 chromatin interactions 200bp (filter activated)
FOSL2 binding sites
FOSL2
FOSL2
FOSL2
FOSL2
FOSL2
FOSL2
FOSL2
FOSL2
CTCF,RAD21,SMC3 binding sites
RAD21
CTCF
SMC3
RAD21
CTCF
SMC3
RAD21
RAD21
SMC3
CTCF
CTCF
SMC3
RAD21
RAD21
SMC3
CTCF
SMC3
RAD21
CTCF
SMC3
Window Position
Scale
chr1:
chr1:46,613,230-47,279,895 (666,666 bp)
200 kb
hg19
46,700,000
46,800,000
46,900,000
47,000,000
47,100,000
47,200,000
0.476218
1_H3K9me3
0.00419809
H3K9me3
21.8486
2_H3K27me3
0.117391
H3K27me3
ln(x+1) 0.786985
3_RNA-seq
-1.52343e-16
RNA-seq
Basic Gene Annotation Set from GENCODE Version 39lift37 (Ensembl 105)
1.27149
4_Insulation score
-0.757255
Insulation score based on chromatin interactions
Boundary of chromatin interactions
Strong
Weak
Strong
Weak
Strong
FOSL2 chromatin interactions 10bp (filter activated)
FOSL2 chromatin interactions 20bp (filter activated)
FOSL2 chromatin interactions 50bp (filter activated)
FOSL2 chromatin interactions 200bp (filter activated)
FOSL2 binding sites
FOSL2
FOSL2
FOSL2
FOSL2
FOSL2
FOSL2
CTCF,RAD21,SMC3 binding sites
RAD21
CTCF
SMC3
SMC3
RAD21
CTCF
CTCF
SMC3
RAD21
RAD21
CTCF
SMC3
CTCF
SMC3
RAD21
CTCF
CTCF
SMC3
RAD21
RAD21
CTCF
SMC3
RAD21
SMC3
CTCF
SMC3
RAD21
CTCF
RAD21
SMC3
CTCF
RAD21
SMC3
CTCF
RAD21
RAD21
SMC3
SMC3
RAD21
CTCF
RAD21
SMC3
CTCF
RAD21
CTCF
SMC3
RAD21
CTCF
SMC3
RAD21
CTCF
RAD21
SMC3
RAD21
CTCF
SMC3
RAD21

### FOXA3

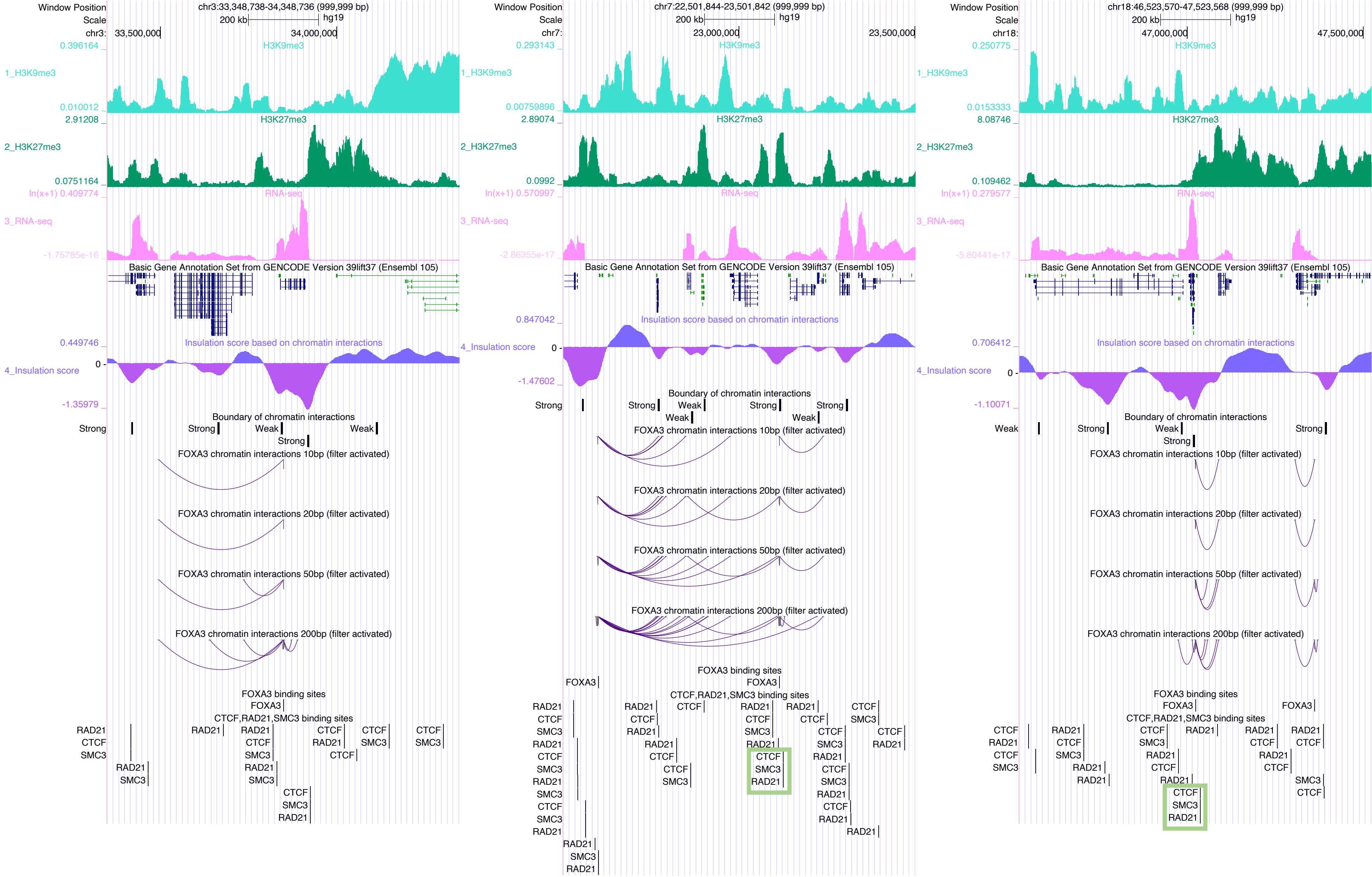

Window Position chr3:33,348,738-34,348,736 (999,999 bp)
Scale 200 kb hg19
chr3: 33,500,000 34,000,000
1_H3K9me3
2_H3K27me3
3_RNA-seq
Basic Gene Annotation Set from GENCODE Version 39lift37 (Ensembl 105)
4_Insulation score
Insulation score based on chromatin interactions
Boundary of chromatin interactions
FOXA3 chromatin interactions 10bp (filter activated)
FOXA3 chromatin interactions 20bp (filter activated)
FOXA3 chromatin interactions 50bp (filter activated)
FOXA3 chromatin interactions 200bp (filter activated)
FOXA3 binding sites
CTCF,RAD21,SMC3 binding sites
Window Position chr7:22,501,844-23,501,842 (999,999 bp)
chr7: 23,000,000 23,500,000
Window Position chr18:46,523,570-47,523,568 (999,999 bp)
chr18: 47,000,000 47,500,000

### FOXH1

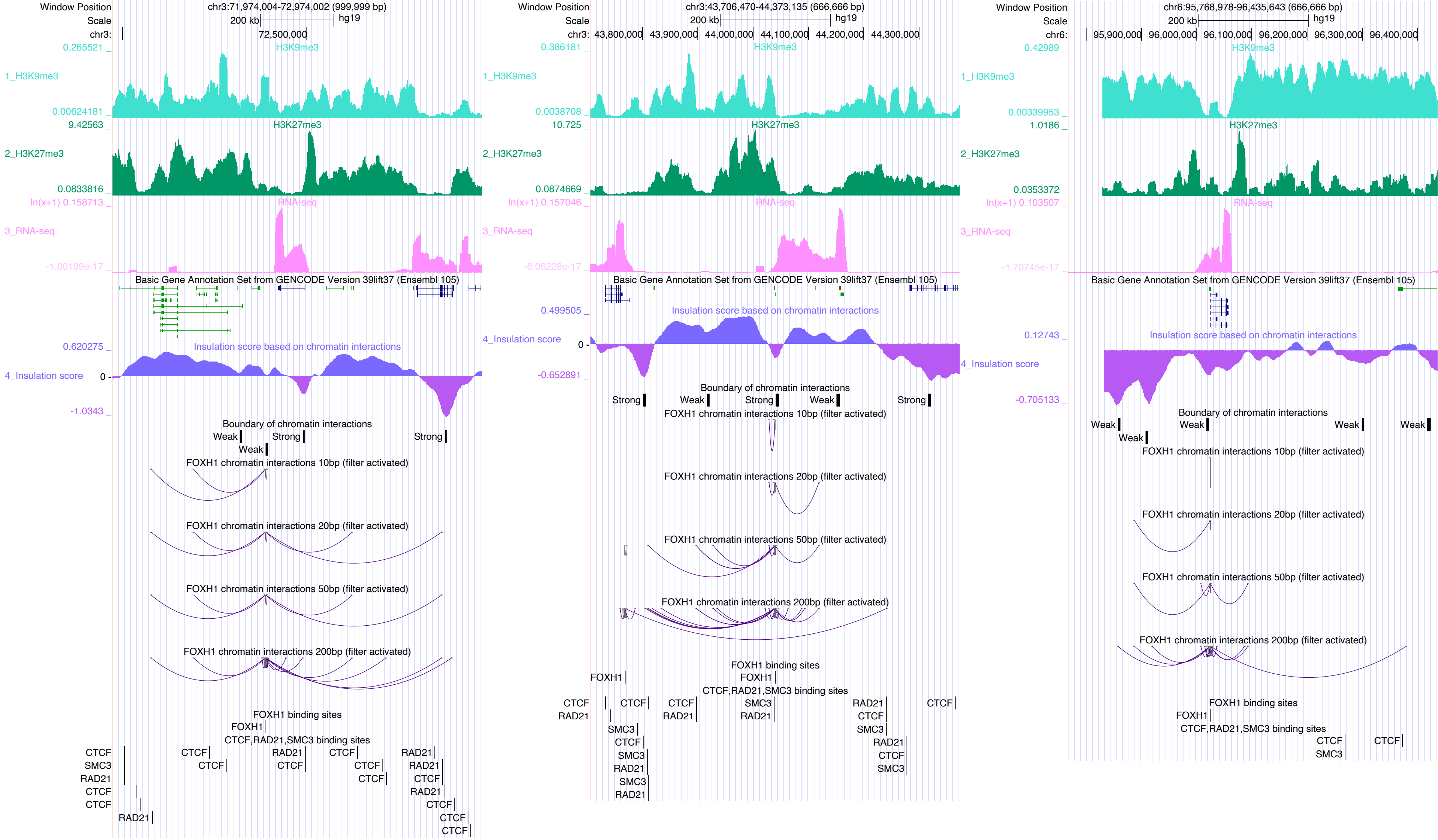
Window Position
Scale
chr3:71,974,004-72,974,002 (999,999 bp)
200 kb
hg19
72,500,000
H3K9me3
1_H3K9me3
0.265521
0.00624181
H3K27me3
2_H3K27me3
9.42563
0.0833816
RNA-seq
3_RNA-seq
ln(x+1) 0.158713
-1.00199e-17
Basic Gene Annotation Set from GENCODE Version 39lift37 (Ensembl 105)
Insulation score based on chromatin interactions
4_Insulation score
0.620275
0
-1.0343
Boundary of chromatin interactions
Weak
Strong
FOXH1 chromatin interactions 10bp (filter activated)
FOXH1 chromatin interactions 20bp (filter activated)
FOXH1 chromatin interactions 50bp (filter activated)
FOXH1 chromatin interactions 200bp (filter activated)
FOXH1 binding sites
FOXH1
CTCF,RAD21,SMC3 binding sites
CTCF
SMC3
RAD21
chr3:43,706,470-44,373,135 (666,666 bp)
43,800,000
43,900,000
44,000,000
44,100,000
44,200,000
44,300,000
0.386181
0.0038708
10.725
0.0874669
ln(x+1) 0.157046
-6.06228e-17
0.499505
-0.652891
chr6:95,768,978-96,435,643 (666,666 bp)
95,900,000
96,000,000
96,100,000
96,200,000
96,300,000
96,400,000
0.42989
0.00339953
1.0186
0.0353372
ln(x+1) 0.103507
-1.70745e-17
0.12743
-0.705133

### FOXK1

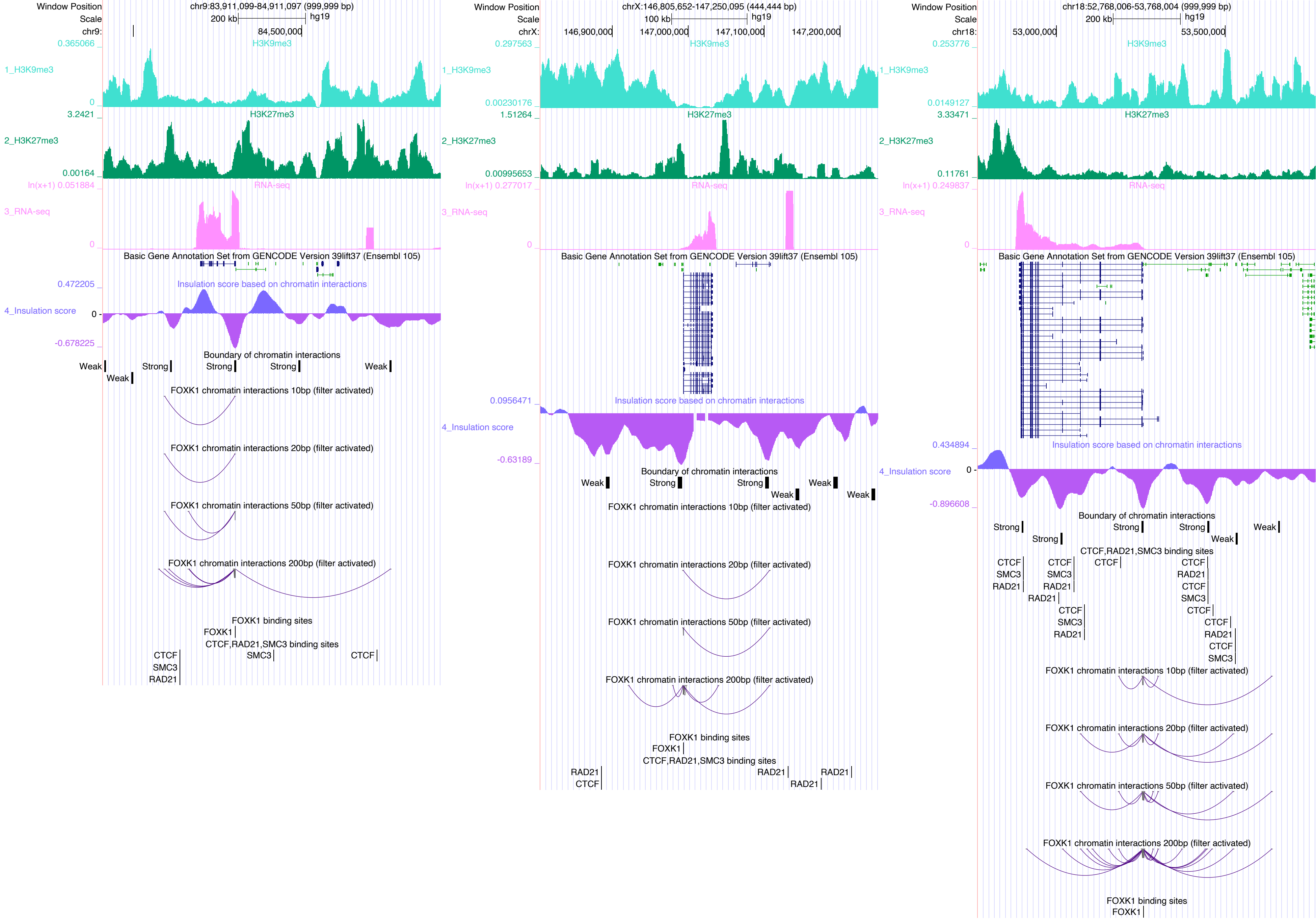

Window Position
chr9:83,911,099-84,911,097 (999,999 bp)
Scale
200 kb
hg19
chr9:
84,500,000
0.365066
H3K9me3
1_H3K9me3
0
3.2421
H3K27me3
2_H3K27me3
0.00164
ln(x+1) 0.051884
RNA-seq
3_RNA-seq
0
Basic Gene Annotation Set from GENCODE Version 39lift37 (Ensembl 105)
0.472205
Insulation score based on chromatin interactions
4_Insulation score
0
-0.678225
Boundary of chromatin interactions
Weak
Strong
Strong
Strong
Weak
Weak
FOXK1 chromatin interactions 10bp (filter activated)
FOXK1 chromatin interactions 20bp (filter activated)
FOXK1 chromatin interactions 50bp (filter activated)
FOXK1 chromatin interactions 200bp (filter activated)
FOXK1 binding sites
FOXK1
CTCF,RAD21,SMC3 binding sites
SMC3
CTCF
CTCF
SMC3
RAD21
Window Position
chrX:146,805,652-147,250,095 (444,444 bp)
Scale
100 kb
hg19
chrX:
146,900,000
147,000,000
147,100,000
147,200,000
0.297563
H3K9me3
1_H3K9me3
0.00230176
1.51264
H3K27me3
2_H3K27me3
0.00995653
ln(x+1) 0.277017
RNA-seq
3_RNA-seq
0
Basic Gene Annotation Set from GENCODE Version 39lift37 (Ensembl 105)
0.0956471
Insulation score based on chromatin interactions
4_Insulation score
-0.63189
Boundary of chromatin interactions
Weak
Strong
Strong
Weak
Weak
Weak
FOXK1 chromatin interactions 10bp (filter activated)
FOXK1 chromatin interactions 20bp (filter activated)
FOXK1 chromatin interactions 50bp (filter activated)
FOXK1 chromatin interactions 200bp (filter activated)
FOXK1 binding sites
FOXK1
CTCF,RAD21,SMC3 binding sites
RAD21
RAD21
RAD21
CTCF
RAD21
Window Position
chr18:52,768,006-53,768,004 (999,999 bp)
Scale
200 kb
hg19
chr18:
53,000,000
53,500,000
0.253776
H3K9me3
1_H3K9me3
0.0149127
3.33471
H3K27me3
2_H3K27me3
0.11761
ln(x+1) 0.249837
RNA-seq
3_RNA-seq
0
Basic Gene Annotation Set from GENCODE Version 39lift37 (Ensembl 105)
0.434894
Insulation score based on chromatin interactions
4_Insulation score
0
-0.896608
Boundary of chromatin interactions
Strong
Strong
Strong
Strong
Weak
Weak
CTCF,RAD21,SMC3 binding sites
CTCF
CTCF
CTCF
CTCF
SMC3
SMC3
RAD21
RAD21
RAD21
RAD21
CTCF
SMC3
CTCF
SMC3
RAD21
CTCF
CTCF
RAD21
CTCF
SMC3
FOXK1 chromatin interactions 10bp (filter activated)
FOXK1 chromatin interactions 20bp (filter activated)
FOXK1 chromatin interactions 50bp (filter activated)
FOXK1 chromatin interactions 200bp (filter activated)
FOXK1 binding sites
FOXK1

### GATA2

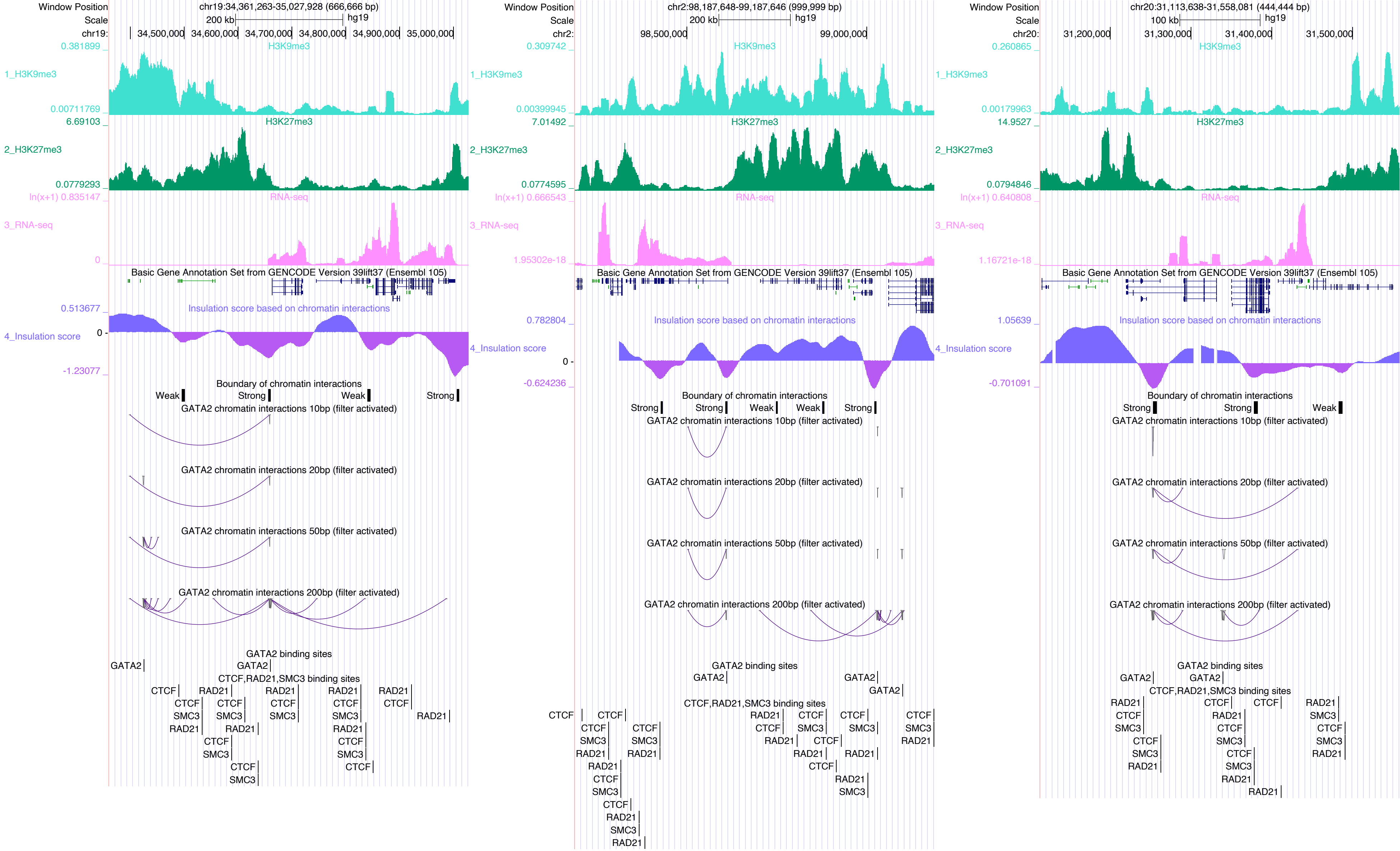

Window Position
Scale
chr19:34,361,263-35,027,928 (666,666 bp)
200 kb
hg19
chr19: 34,500,000 34,600,000 34,700,000 34,800,000 34,900,000 35,000,000
0.381899
H3K9me3
1_H3K9me3
0.00711769
6.69103
H3K27me3
2_H3K27me3
0.0779293
ln(x+1) 0.835147
RNA-seq
3_RNA-seq
0
Basic Gene Annotation Set from GENCODE Version 39lift37 (Ensembl 105)
0.513677
Insulation score based on chromatin interactions
4_Insulation score
0
-1.23077
Boundary of chromatin interactions
Weak Strong Weak Strong
GATA2 chromatin interactions 10bp (filter activated)
GATA2 chromatin interactions 20bp (filter activated)
GATA2 chromatin interactions 50bp (filter activated)
GATA2 chromatin interactions 200bp (filter activated)
GATA2 binding sites
GATA2
GATA2
CTCF,RAD21,SMC3 binding sites
chr2:98,187,648-99,187,646 (999,999 bp)
200 kb
hg19
chr2: 98,500,000 99,000,000
0.309742
0.00399945
7.01492
0.0774595
ln(x+1) 0.666543
1.95302e-18
0.782804
0
-0.624236
Strong Strong Weak Weak Strong
chr20:31,113,638-31,558,081 (444,444 bp)
100 kb
hg19
chr20: 31,200,000 31,300,000 31,400,000 31,500,000
0.260865
0.00179963
14.9527
0.0794846
ln(x+1) 0.640808
1.16721e-18
1.05639
-0.701091
Strong Strong Weak
CTCF
SMC3
RAD21

### GRHL1

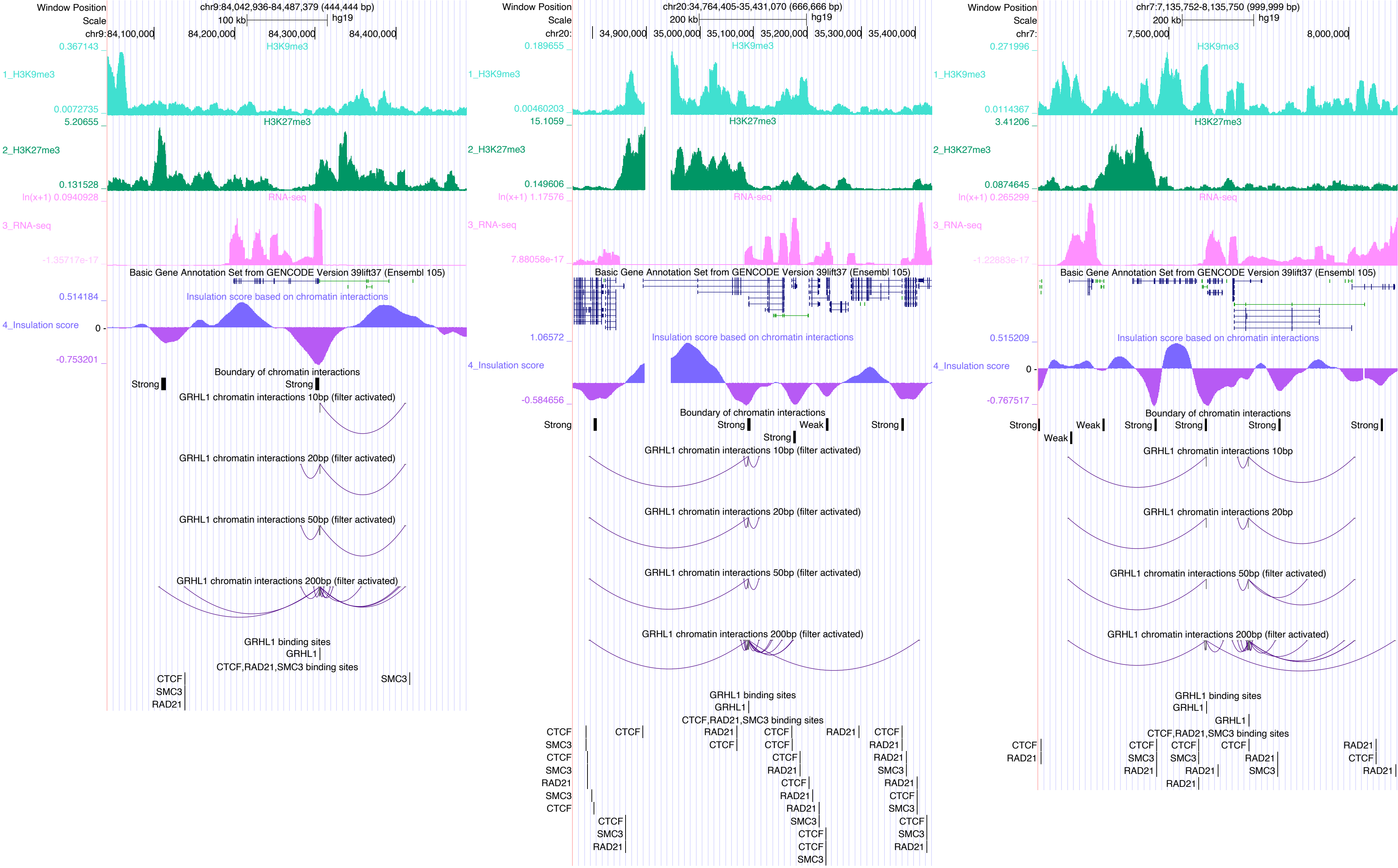
Window Position
chr9:84,042,936-84,487,379 (444,444 bp)
Scale
100 kb
hg19
chr9:84,100,000
84,200,000
84,300,000
84,400,000
0.367143
1_H3K9me3
H3K9me3
0.0072735
5.20655
2_H3K27me3
H3K27me3
0.131528
ln(x+1) 0.0940928
3_RNA-seq
RNA-seq
-1.35717e-17
Basic Gene Annotation Set from GENCODE Version 39lift37 (Ensembl 105)
0.514184
Insulation score based on chromatin interactions
4_Insulation score
0
-0.753201
Boundary of chromatin interactions
Strong
Strong
GRHL1 chromatin interactions 10bp (filter activated)
GRHL1 chromatin interactions 20bp (filter activated)
GRHL1 chromatin interactions 50bp (filter activated)
GRHL1 chromatin interactions 200bp (filter activated)
GRHL1 binding sites
GRHL1
CTCF,RAD21,SMC3 binding sites
CTCF
SMC3
RAD21
SMC3
Window Position
chr20:34,764,405-35,431,070 (666,666 bp)
Scale
200 kb
hg19
chr20:
34,900,000
35,000,000
35,100,000
35,200,000
35,300,000
35,400,000
0.189655
1_H3K9me3
H3K9me3
0.00460203
15.1059
2_H3K27me3
H3K27me3
0.149606
ln(x+1) 1.17576
3_RNA-seq
RNA-seq
7.88058e-17
Basic Gene Annotation Set from GENCODE Version 39lift37 (Ensembl 105)
1.06572
Insulation score based on chromatin interactions
4_Insulation score
-0.584656
Boundary of chromatin interactions
Strong
Strong
Weak
Strong
Strong
GRHL1 chromatin interactions 10bp (filter activated)
GRHL1 chromatin interactions 20bp (filter activated)
GRHL1 chromatin interactions 50bp (filter activated)
GRHL1 chromatin interactions 200bp (filter activated)
GRHL1 binding sites
GRHL1
CTCF,RAD21,SMC3 binding sites
Window Position
chr7:7,135,752-8,135,750 (999,999 bp)
Scale
200 kb
hg19
chr7:
7,500,000
8,000,000
0.271996
1_H3K9me3
H3K9me3
0.0114367
3.41206
2_H3K27me3
H3K27me3
0.0874645
ln(x+1) 0.265299
3_RNA-seq
RNA-seq
-1.22883e-17
Basic Gene Annotation Set from GENCODE Version 39lift37 (Ensembl 105)
0.515209
Insulation score based on chromatin interactions
4_Insulation score
0
-0.767517
Boundary of chromatin interactions
Strong
Weak
Weak
Strong
Strong
Strong
Strong
GRHL1 chromatin interactions 10bp
GRHL1 chromatin interactions 20bp
GRHL1 chromatin interactions 50bp (filter activated)
GRHL1 chromatin interactions 200bp (filter activated)
GRHL1 binding sites
GRHL1
GRHL1
CTCF,RAD21,SMC3 binding sites

### GRHL2

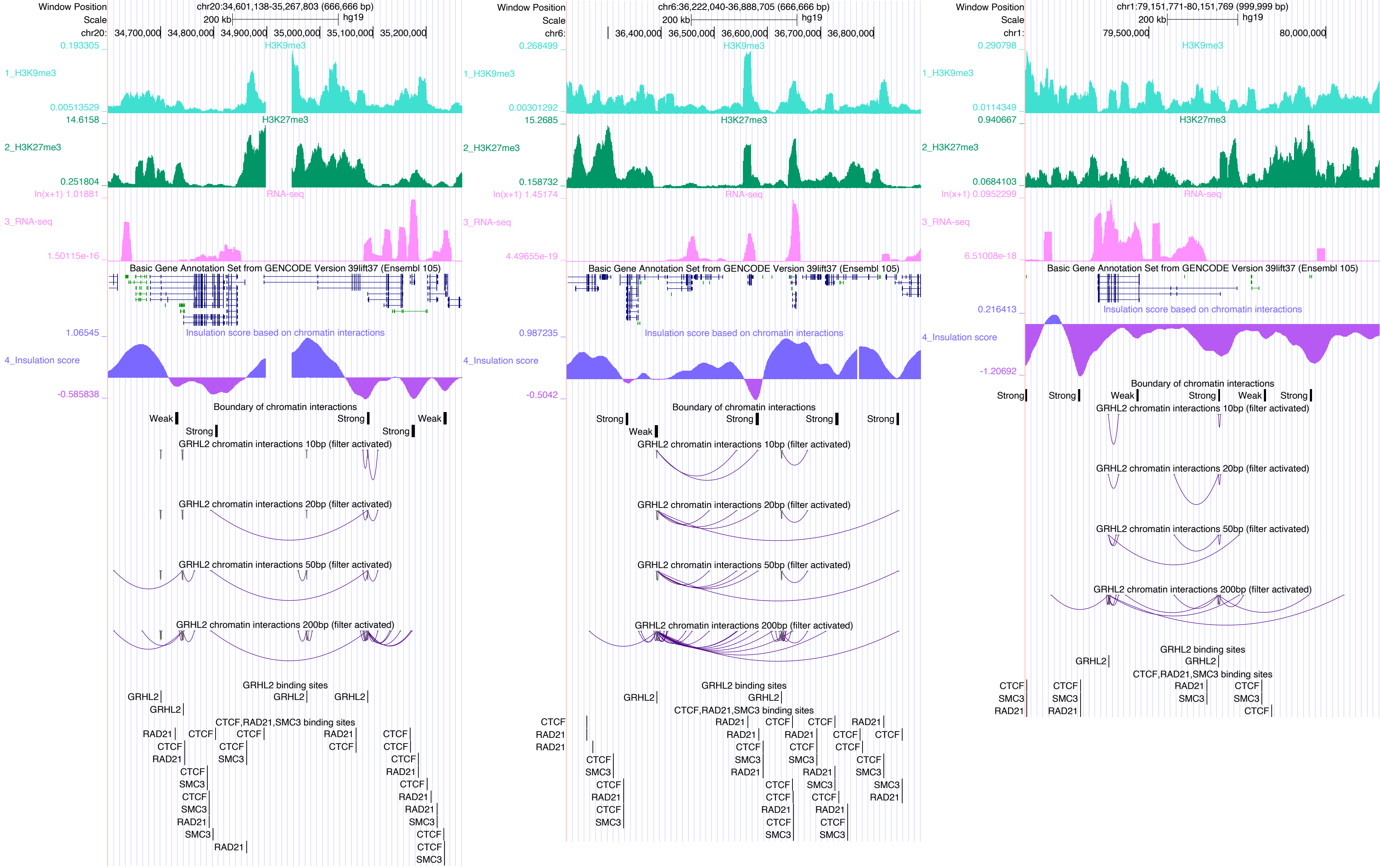

Window Position
Scale
chr20:34,601,138-35,267,803 (666,666 bp)
200 kb
hg19
chr20: 34,700,000 34,800,000 34,900,000 35,000,000 35,100,000 35,200,000
0.193305
H3K9me3
1_H3K9me3
0.00513529
14.6158
H3K27me3
2_H3K27me3
0.251804
ln(x+1) 1.01881
RNA-seq
3_RNA-seq
1.50115e-16
Basic Gene Annotation Set from GENCODE Version 39lift37 (Ensembl 105)
1.06545
Insulation score based on chromatin interactions
4_Insulation score
-0.585838
Boundary of chromatin interactions
Weak
Strong
Strong
Strong
Weak
GRHL2 chromatin interactions 10bp (filter activated)
GRHL2 chromatin interactions 20bp (filter activated)
GRHL2 chromatin interactions 50bp (filter activated)
GRHL2 chromatin interactions 200bp (filter activated)
GRHL2 binding sites
CTCF,RAD21,SMC3 binding sites
chr6:36,222,040-36,888,705 (666,666 bp)
chr6: 36,400,000 36,500,000 36,600,000 36,700,000 36,800,000
0.268499
0.00301292
15.2685
0.158732
ln(x+1) 1.45174
4.49655e-19
0.987235
-0.5042
chr1:79,151,771-80,151,769 (999,999 bp)
chr1: 79,500,000 80,000,000
0.290798
0.0114349
0.940667
0.0684103
ln(x+1) 0.0952299
6.51008e-18
0.216413
-1.20692

### HNF4A

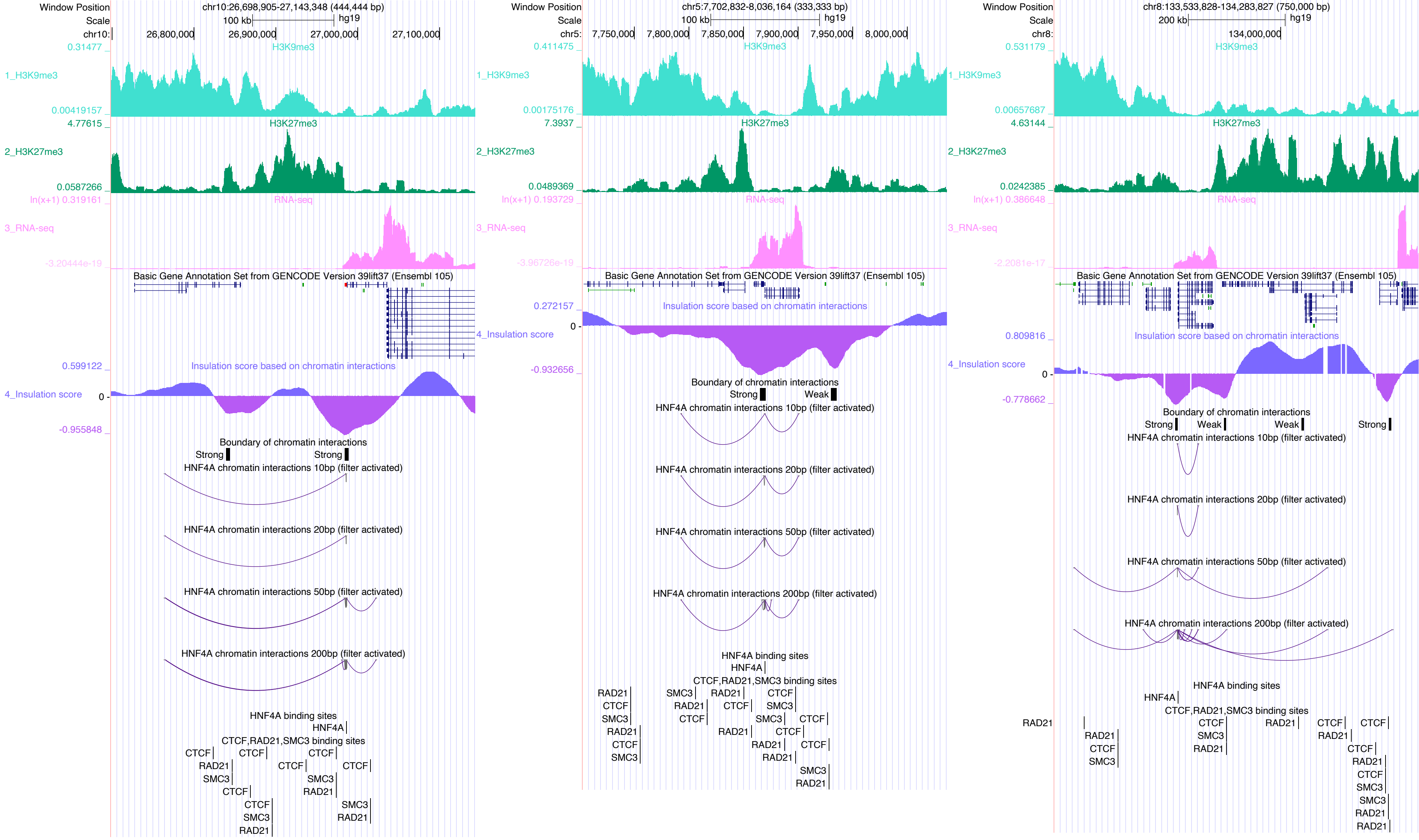

Window Position
chr10:26,698,905-27,143,348 (444,444 bp)
chr5:7,702,832-8,036,164 (333,333 bp)
chr8:133,533,828-134,283,827 (750,000 bp)
hg19
1_H3K9me3
2_H3K27me3
3_RNA-seq
4_Insulation score
Basic Gene Annotation Set from GENCODE Version 39lift37 (Ensembl 105)
Insulation score based on chromatin interactions
Boundary of chromatin interactions
Strong
Weak
HNF4A chromatin interactions 10bp (filter activated)
HNF4A chromatin interactions 20bp (filter activated)
HNF4A chromatin interactions 50bp (filter activated)
HNF4A chromatin interactions 200bp (filter activated)
HNF4A binding sites
CTCF,RAD21,SMC3 binding sites

### JUN

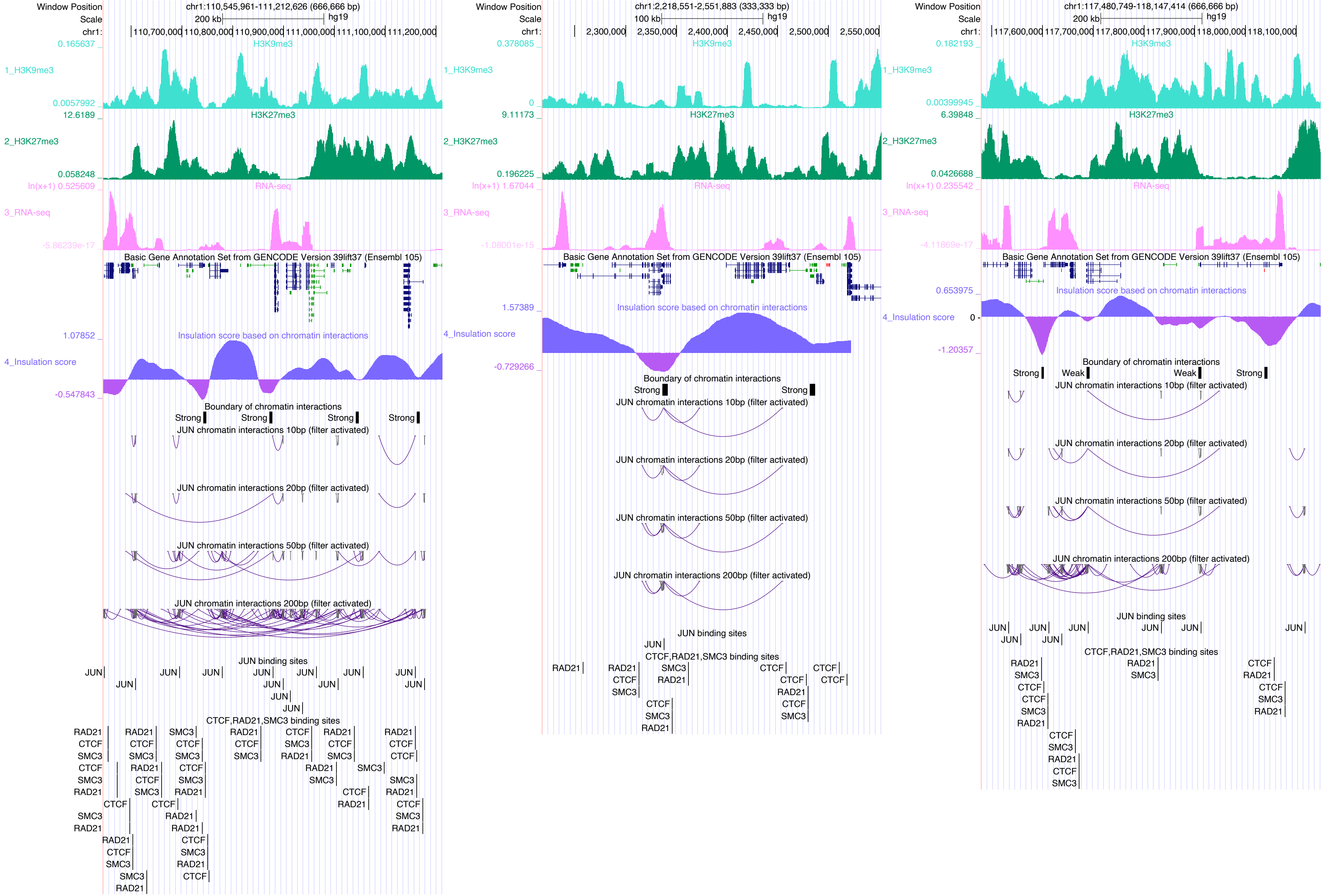

Window Position
chr1:110,545,961-111,212,626 (666,666 bp)
chr1:2,218,551-2,551,883 (333,333 bp)
chr1:117,480,749-118,147,414 (666,666 bp)
hg19
H3K9me3
H3K27me3
RNA-seq
Basic Gene Annotation Set from GENCODE Version 39lift37 (Ensembl 105)
Insulation score based on chromatin interactions
Boundary of chromatin interactions
JUN chromatin interactions 10bp (filter activated)
JUN chromatin interactions 20bp (filter activated)
JUN chromatin interactions 50bp (filter activated)
JUN chromatin interactions 200bp (filter activated)
JUN binding sites
CTCF,RAD21,SMC3 binding sites

### JUNB

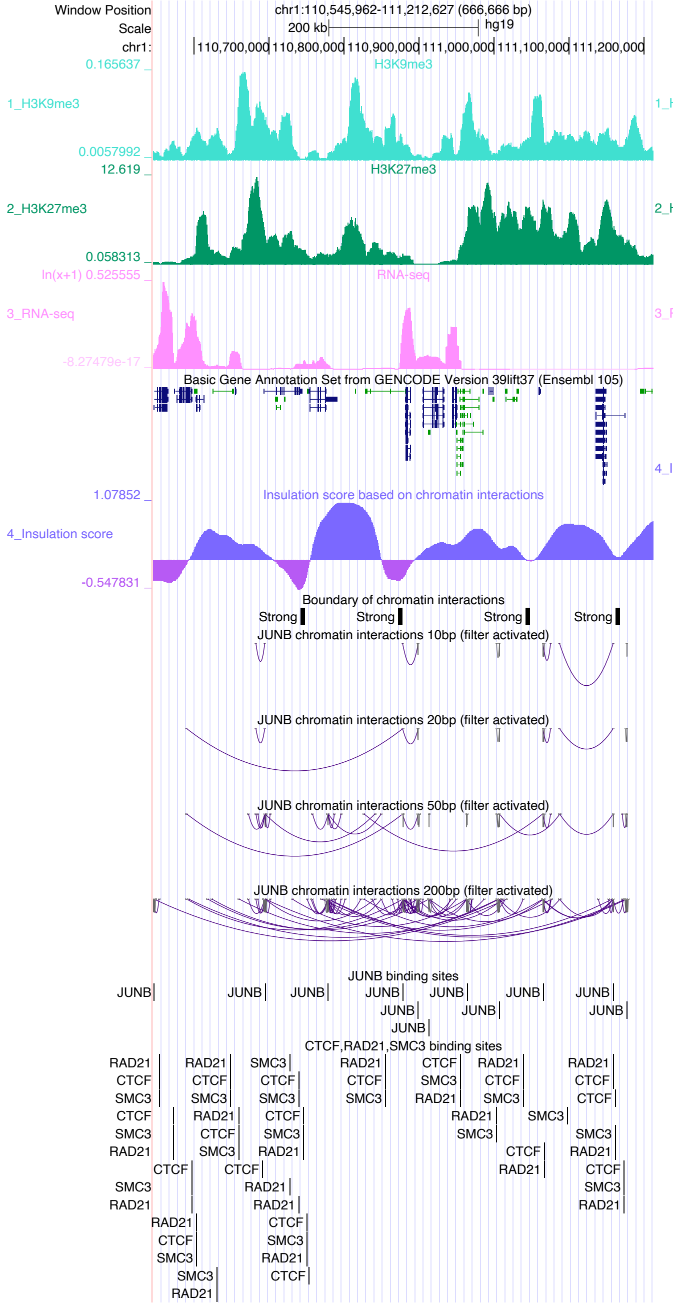

Window Position
chr1:110,545,962-111,212,627 (666,666 bp)
Scale
200 kb
hg19
chr1:
110,700,000
110,800,000
110,900,000
111,000,000
111,100,000
111,200,000
0.165637
H3K9me3
1_H3K9me3
0.0057992
12.619
H3K27me3
2_H3K27me3
0.058313
ln(x+1) 0.525555
RNA-seq
3_RNA-seq
-8.27479e-17
Basic Gene Annotation Set from GENCODE Version 39lift37 (Ensembl 105)
1.07852
Insulation score based on chromatin interactions
4_Insulation score
-0.547831
Boundary of chromatin interactions
Strong
JUNB chromatin interactions 10bp (filter activated)
JUNB chromatin interactions 20bp (filter activated)
JUNB chromatin interactions 50bp (filter activated)
JUNB chromatin interactions 200bp (filter activated)
JUNB binding sites
CTCF,RAD21,SMC3 binding sites


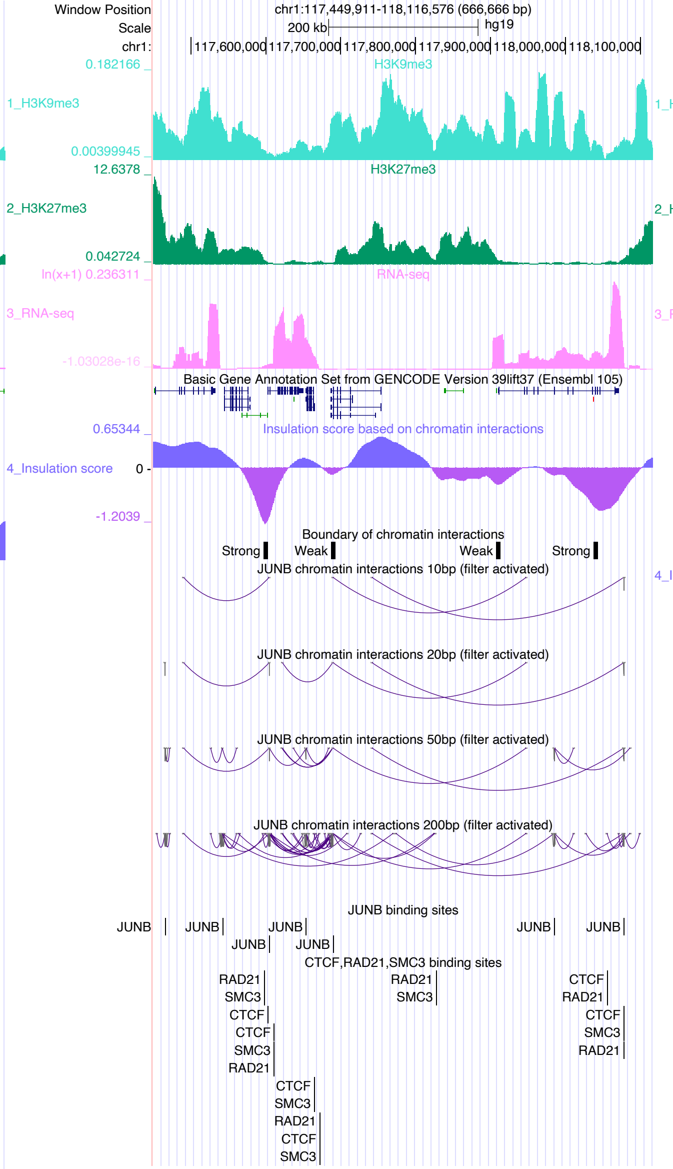

Window Position
chr1:117,449,911-118,116,576 (666,666 bp)
Scale
200 kb
hg19
chr1:
117,600,000
117,700,000
117,800,000
117,900,000
118,000,000
118,100,000
0.182166
H3K9me3
1_H3K9me3
0.00399945
12.6378
H3K27me3
2_H3K27me3
0.042724
ln(x+1) 0.236311
RNA-seq
3_RNA-seq
-1.03028e-16
Basic Gene Annotation Set from GENCODE Version 39lift37 (Ensembl 105)
0.65344
Insulation score based on chromatin interactions
4_Insulation score
-1.2039
Boundary of chromatin interactions
Strong
Weak
JUNB chromatin interactions 10bp (filter activated)
JUNB chromatin interactions 20bp (filter activated)
JUNB chromatin interactions 50bp (filter activated)
JUNB chromatin interactions 200bp (filter activated)
JUNB binding sites
CTCF,RAD21,SMC3 binding sites


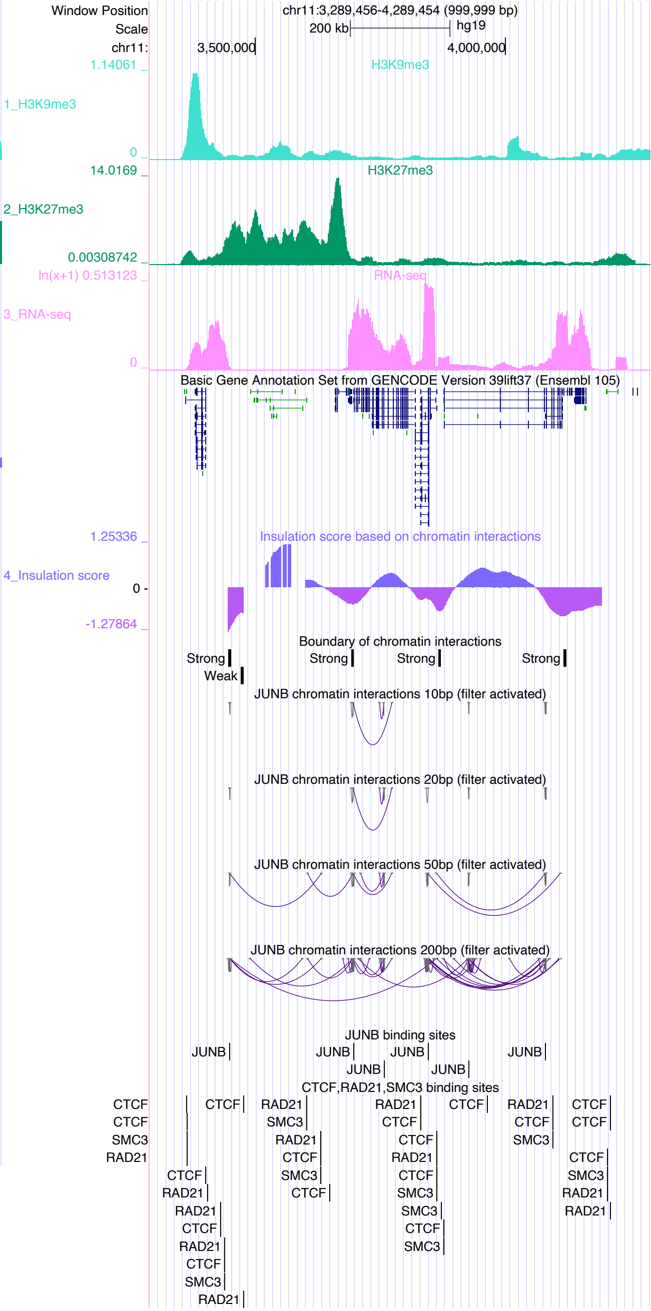

Window Position
chr11:3,289,456-4,289,454 (999,999 bp)
Scale
200 kb
hg19
chr11:
3,500,000
4,000,000
1.14061
H3K9me3
1_H3K9me3
14.0169
H3K27me3
2_H3K27me3
0.00308742
ln(x+1) 0.513123
RNA-seq
3_RNA-seq
Basic Gene Annotation Set from GENCODE Version 39lift37 (Ensembl 105)
1.25336
Insulation score based on chromatin interactions
4_Insulation score
-1.27864
Boundary of chromatin interactions
Strong
Weak
JUNB chromatin interactions 10bp (filter activated)
JUNB chromatin interactions 20bp (filter activated)
JUNB chromatin interactions 50bp (filter activated)
JUNB chromatin interactions 200bp (filter activated)
JUNB binding sites
CTCF,RAD21,SMC3 binding sites

### JUND

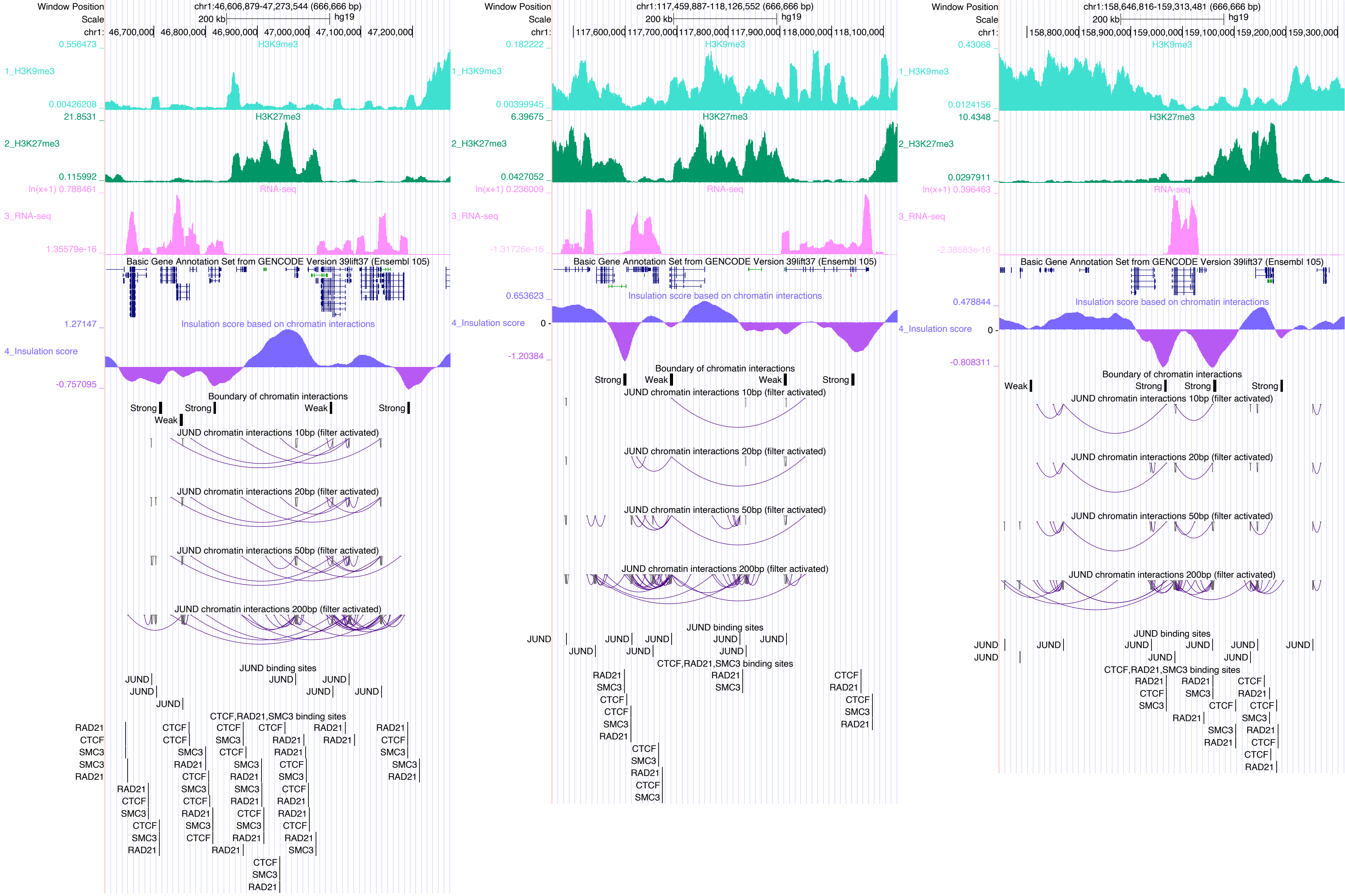

Window Position
chr1:46,606,879-47,273,544 (666,666 bp)
Scale
200 kb
hg19
chr1: 46,700,000 46,800,000 46,900,000 47,000,000 47,100,000 47,200,000
1_H3K9me3
H3K9me3
0.556473
0.00426208
2_H3K27me3
H3K27me3
21.8531
0.115992
3_RNA-seq
RNA-seq
ln(x+1) 0.788461
1.35579e-16
Basic Gene Annotation Set from GENCODE Version 39lift37 (Ensembl 105)
4_Insulation score
Insulation score based on chromatin interactions
1.27147
-0.757095
Boundary of chromatin interactions
Strong
Weak
JUND chromatin interactions 10bp (filter activated)
JUND chromatin interactions 20bp (filter activated)
JUND chromatin interactions 50bp (filter activated)
JUND chromatin interactions 200bp (filter activated)
JUND binding sites
JUND
CTCF,RAD21,SMC3 binding sites
CTCF
RAD21
SMC3
Window Position
chr1:117,459,887-118,126,552 (666,666 bp)
chr1: 117,600,000 117,700,000 117,800,000 117,900,000 118,000,000 118,100,000
0.182222
0.00399945
6.39675
0.0427052
ln(x+1) 0.236009
-1.31726e-16
0.653623
-1.20384
Window Position
chr1:158,646,816-159,313,481 (666,666 bp)
chr1: 158,800,000 158,900,000 159,000,000 159,100,000 159,200,000 159,300,000
0.43068
0.0124156
10.4348
0.0297911
ln(x+1) 0.396463
-2.38583e-16
0.478844
-0.808311

KLF1

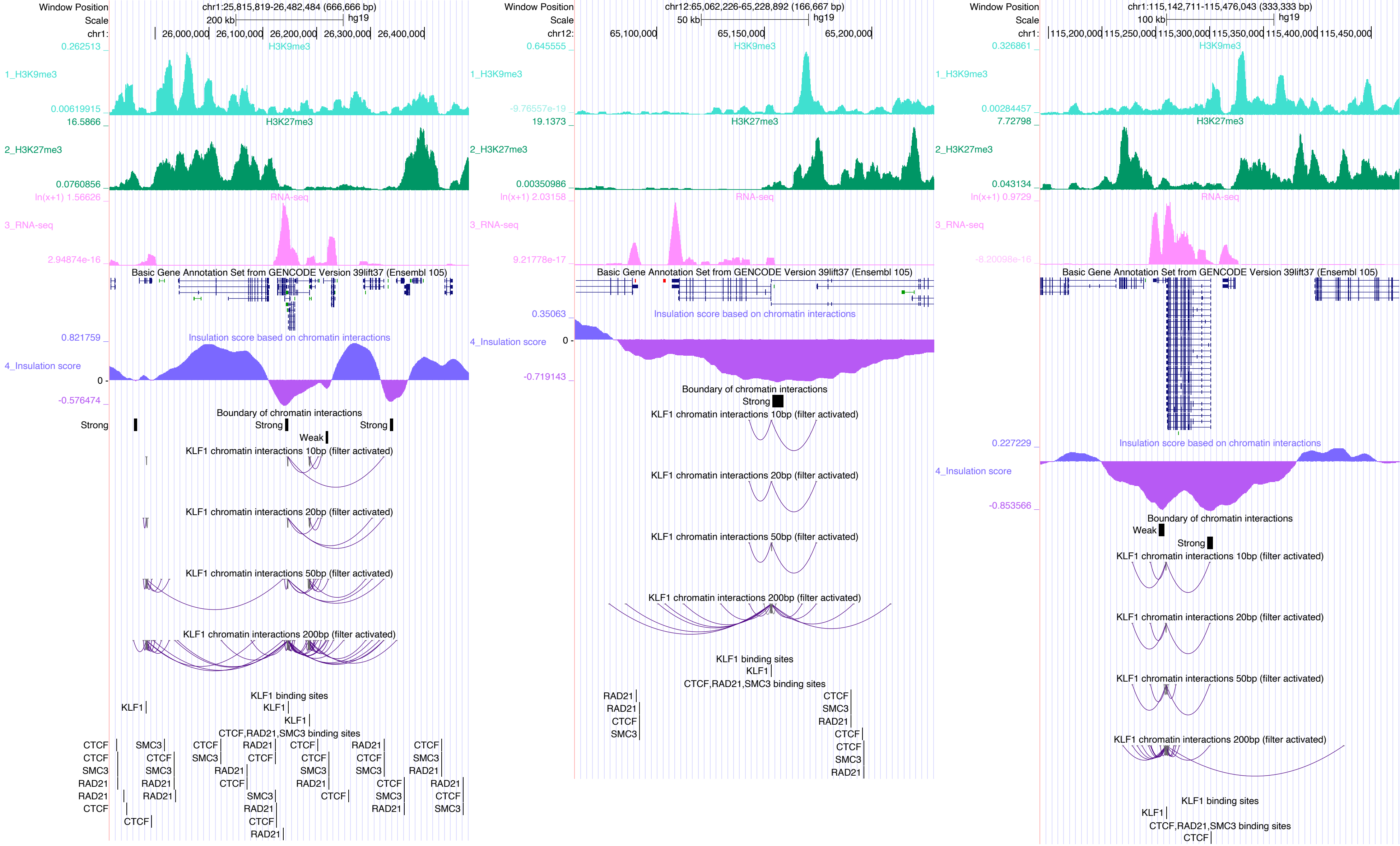
Window Position
chr1:25,815,819-26,482,484 (666,666 bp)
Scale
200 kb
hg19
chr1:
26,000,000
26,100,000
26,200,000
26,300,000
26,400,000
0.262513 _
H3K9me3
1_H3K9me3
0.00619915 _
16.5866 _
H3K27me3
2_H3K27me3
0.0760856 _
ln(x+1) 1.56626 _
RNA-seq
3_RNA-seq
2.94874e-16 _
Basic Gene Annotation Set from GENCODE Version 39lift37 (Ensembl 105)
0.821759 _
Insulation score based on chromatin interactions
4_Insulation score
0 -
-0.576474 _
Boundary of chromatin interactions
Strong
Strong
Strong
Weak
KLF1 chromatin interactions 10bp (filter activated)
KLF1 chromatin interactions 20bp (filter activated)
KLF1 chromatin interactions 50bp (filter activated)
KLF1 chromatin interactions 200bp (filter activated)
KLF1 binding sites
KLF1
KLF1
KLF1
CTCF,RAD21,SMC3 binding sites
CTCF
CTCF
SMC3
RAD21
RAD21
CTCF
SMC3
CTCF
SMC3
RAD21
RAD21
CTCF
CTCF
SMC3
RAD21
CTCF
RAD21
CTCF
SMC3
RAD21
CTCF
RAD21
CTCF
CTCF
SMC3
RAD21
CTCF
RAD21
CTCF
SMC3
CTCF
SMC3
RAD21
CTCF
SMC3
RAD21
CTCF
SMC3
RAD21
RAD21
CTCF
SMC3
Window Position
chr12:65,062,226-65,228,892 (166,667 bp)
Scale
50 kb
hg19
chr12:
65,100,000
65,150,000
65,200,000
0.645555 _
H3K9me3
1_H3K9me3
-9.76557e-19 _
19.1373 _
H3K27me3
2_H3K27me3
0.00350986 _
ln(x+1) 2.03158 _
RNA-seq
3_RNA-seq
9.21778e-17 _
Basic Gene Annotation Set from GENCODE Version 39lift37 (Ensembl 105)
0.35063 _
Insulation score based on chromatin interactions
4_Insulation score
0 -
-0.719143 _
Boundary of chromatin interactions
Strong
KLF1 chromatin interactions 10bp (filter activated)
KLF1 chromatin interactions 20bp (filter activated)
KLF1 chromatin interactions 50bp (filter activated)
KLF1 chromatin interactions 200bp (filter activated)
KLF1 binding sites
KLF1
CTCF,RAD21,SMC3 binding sites
RAD21
RAD21
CTCF
SMC3
CTCF
SMC3
RAD21
CTCF
CTCF
SMC3
RAD21
Window Position
chr1:115,142,711-115,476,043 (333,333 bp)
Scale
100 kb
hg19
chr1:
115,200,000
115,250,000
115,300,000
115,350,000
115,400,000
115,450,000
0.326861 _
H3K9me3
1_H3K9me3
0.00284457 _
7.72798 _
H3K27me3
2_H3K27me3
0.043134 _
ln(x+1) 0.9729 _
RNA-seq
3_RNA-seq
-8.20098e-16 _
Basic Gene Annotation Set from GENCODE Version 39lift37 (Ensembl 105)
0.227229 _
Insulation score based on chromatin interactions
4_Insulation score
-0.853566 _
Boundary of chromatin interactions
Weak
Strong
KLF1 chromatin interactions 10bp (filter activated)
KLF1 chromatin interactions 20bp (filter activated)
KLF1 chromatin interactions 50bp (filter activated)
KLF1 chromatin interactions 200bp (filter activated)
KLF1 binding sites
KLF1
CTCF,RAD21,SMC3 binding sites
CTCF

KLF5

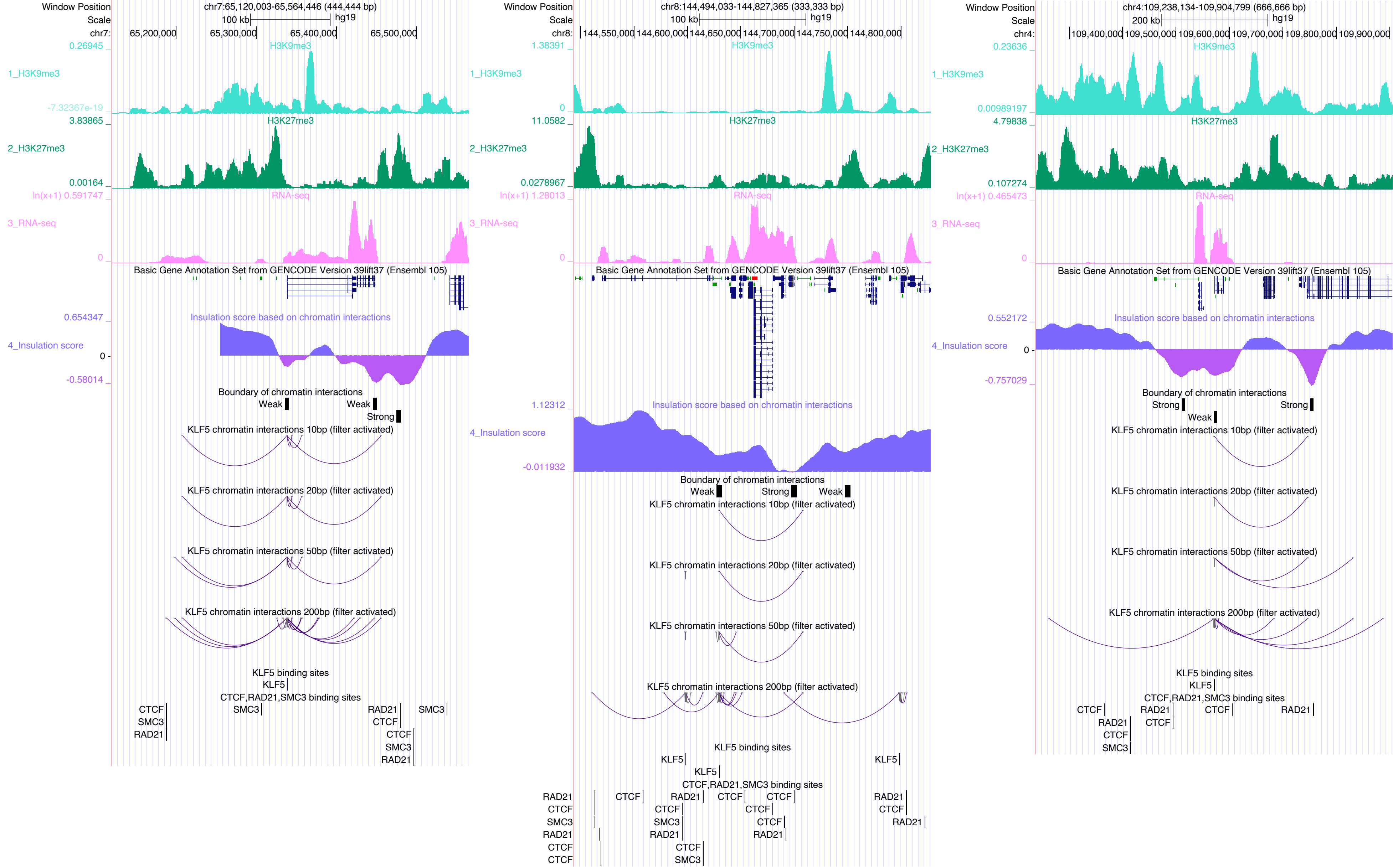

Window Position chr7:65,120,003-65,564,446 (444,444 bp)
Scale 100 kb hg19
chr7: 65,200,000 65,300,000 65,400,000 65,500,000
1_H3K9me3
2_H3K27me3
3_RNA-seq
Basic Gene Annotation Set from GENCODE Version 39lift37 (Ensembl 105)
4_Insulation score
Insulation score based on chromatin interactions
Boundary of chromatin interactions
KLF5 chromatin interactions 10bp (filter activated)
KLF5 chromatin interactions 20bp (filter activated)
KLF5 chromatin interactions 50bp (filter activated)
KLF5 chromatin interactions 200bp (filter activated)
KLF5 binding sites
CTCF,RAD21,SMC3 binding sites
Window Position chr8:144,494,033-144,827,365 (333,333 bp)
chr8: 144,550,000 144,600,000 144,650,000 144,700,000 144,750,000 144,800,000
Window Position chr4:109,238,134-109,904,799 (666,666 bp)
Scale 200 kb hg19
chr4: 109,400,000 109,500,000 109,600,000 109,700,000 109,800,000 109,900,000

### KLF8

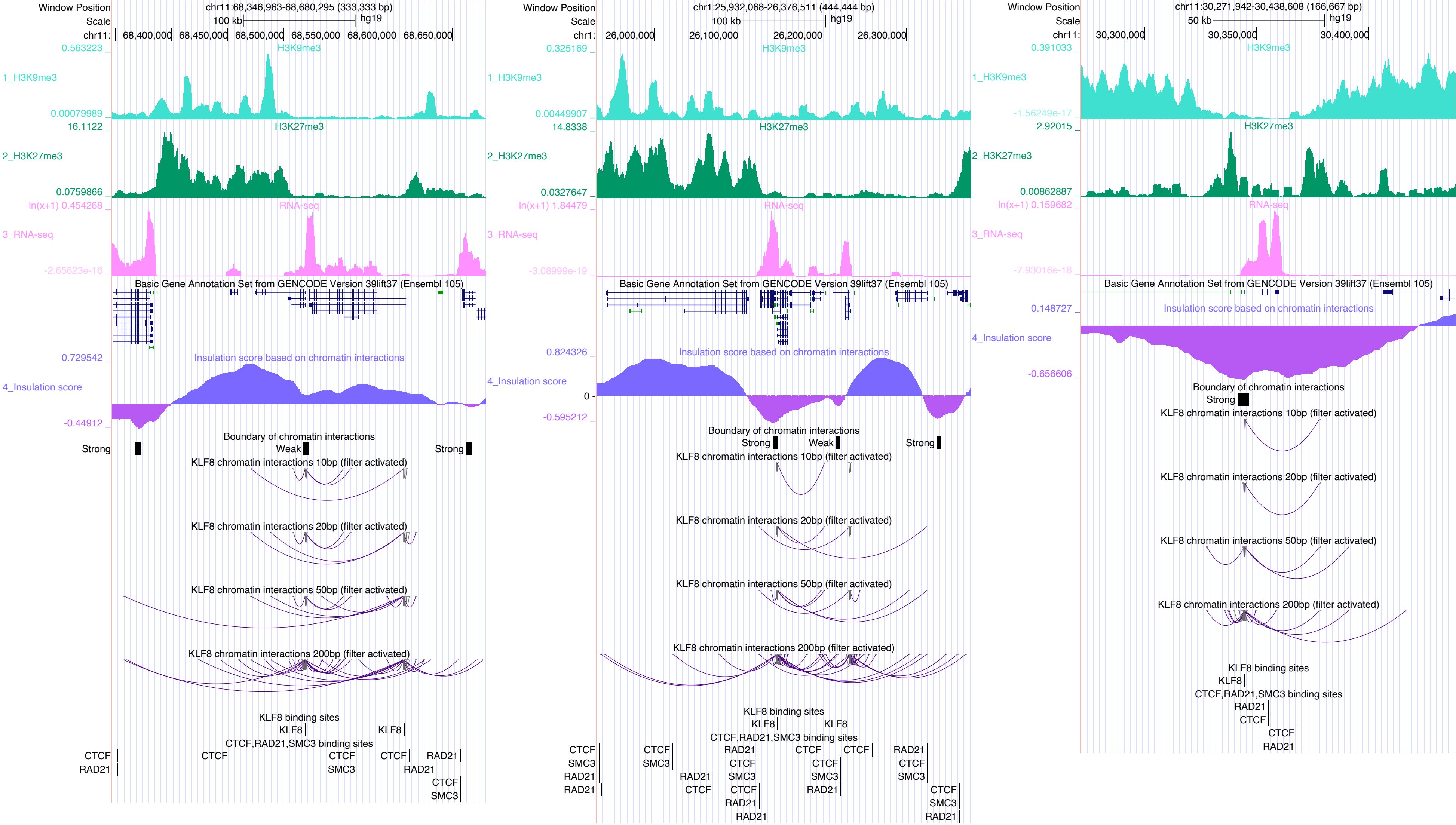

Window Position
chr11:68,346,963-68,680,295 (333,333 bp)
Scale
100 kb
hg19
chr11:
68,400,000
68,450,000
68,500,000
68,550,000
68,600,000
68,650,000
0.563223
H3K9me3
1_H3K9me3
0.00079989
16.1122
H3K27me3
2_H3K27me3
0.0759866
ln(x+1) 0.454268
RNA-seq
3_RNA-seq
-2.65623e-16
Basic Gene Annotation Set from GENCODE Version 39lift37 (Ensembl 105)
0.729542
Insulation score based on chromatin interactions
4_Insulation score
-0.44912
Boundary of chromatin interactions
Strong
Weak
Strong
KLF8 chromatin interactions 10bp (filter activated)
KLF8 chromatin interactions 20bp (filter activated)
KLF8 chromatin interactions 50bp (filter activated)
KLF8 chromatin interactions 200bp (filter activated)
KLF8 binding sites
KLF8
KLF8
CTCF,RAD21,SMC3 binding sites
CTCF
RAD21
CTCF
CTCF
SMC3
CTCF
RAD21
RAD21
CTCF
SMC3
Window Position
chr1:25,932,068-26,376,511 (444,444 bp)
Scale
100 kb
hg19
chr1:
26,000,000
26,100,000
26,200,000
26,300,000
0.325169
H3K9me3
1_H3K9me3
0.00449907
14.8338
H3K27me3
2_H3K27me3
0.0327647
ln(x+1) 1.84479
RNA-seq
3_RNA-seq
-3.08999e-19
Basic Gene Annotation Set from GENCODE Version 39lift37 (Ensembl 105)
0.824326
Insulation score based on chromatin interactions
4_Insulation score
0
-0.595212
Boundary of chromatin interactions
Strong
Weak
Strong
KLF8 chromatin interactions 10bp (filter activated)
KLF8 chromatin interactions 20bp (filter activated)
KLF8 chromatin interactions 50bp (filter activated)
KLF8 chromatin interactions 200bp (filter activated)
KLF8 binding sites
KLF8
KLF8
CTCF,RAD21,SMC3 binding sites
CTCF
SMC3
RAD21
RAD21
CTCF
SMC3
RAD21
CTCF
SMC3
CTCF
RAD21
RAD21
CTCF
CTCF
SMC3
RAD21
CTCF
RAD21
CTCF
SMC3
CTCF
SMC3
RAD21
Window Position
chr11:30,271,942-30,438,608 (166,667 bp)
Scale
50 kb
hg19
chr11:
30,300,000
30,350,000
30,400,000
0.391033
H3K9me3
1_H3K9me3
-1.56249e-17
2.92015
H3K27me3
2_H3K27me3
0.00862887
ln(x+1) 0.159682
RNA-seq
3_RNA-seq
-7.93016e-18
Basic Gene Annotation Set from GENCODE Version 39lift37 (Ensembl 105)
0.148727
Insulation score based on chromatin interactions
4_Insulation score
-0.656606
Boundary of chromatin interactions
Strong
KLF8 chromatin interactions 10bp (filter activated)
KLF8 chromatin interactions 20bp (filter activated)
KLF8 chromatin interactions 50bp (filter activated)
KLF8 chromatin interactions 200bp (filter activated)
KLF8 binding sites
KLF8
CTCF,RAD21,SMC3 binding sites
RAD21
CTCF
CTCF
RAD21

### KLF9

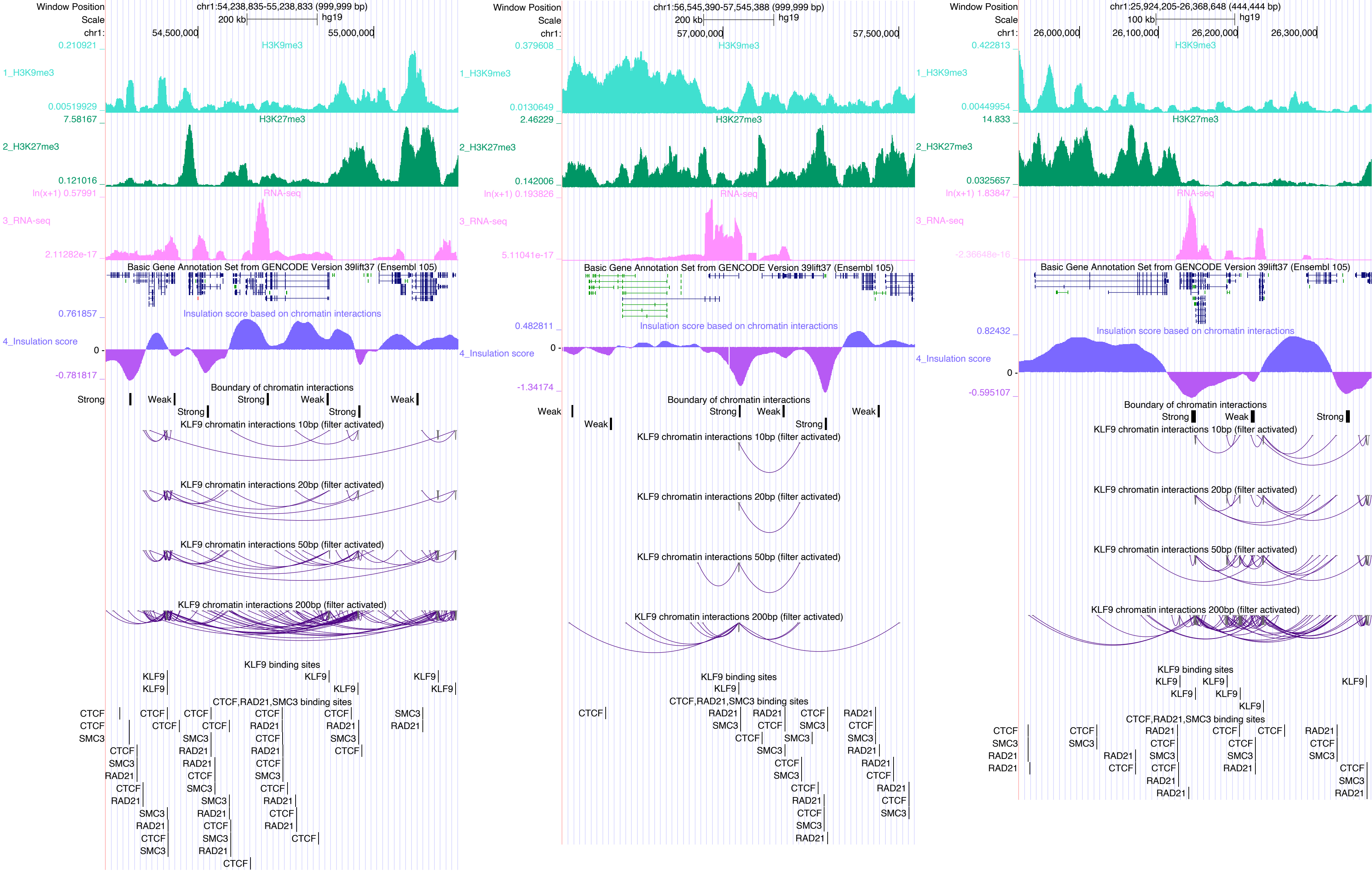

Window Position
Scale
chr1:
chr1:54,238,835-55,238,833 (999,999 bp)
200 kb
hg19
54,500,000
55,000,000
0.210921
1_H3K9me3
H3K9me3
0.00519929
7.58167
2_H3K27me3
H3K27me3
0.121016
ln(x+1) 0.57991
3_RNA-seq
RNA-seq
2.11282e-17
Basic Gene Annotation Set from GENCODE Version 39lift37 (Ensembl 105)
0.761857
4_Insulation score
Insulation score based on chromatin interactions
0
-0.781817
Boundary of chromatin interactions
Strong
Weak
KLF9 chromatin interactions 10bp (filter activated)
KLF9 chromatin interactions 20bp (filter activated)
KLF9 chromatin interactions 50bp (filter activated)
KLF9 chromatin interactions 200bp (filter activated)
KLF9 binding sites
KLF9
CTCF,RAD21,SMC3 binding sites
CTCF
SMC3
RAD21
chr1:56,545,390-57,545,388 (999,999 bp)
57,000,000
57,500,000
0.379608
0.0130649
2.46229
0.142006
ln(x+1) 0.193826
5.11041e-17
0.482811
-1.34174
chr1:25,924,205-26,368,648 (444,444 bp)
100 kb
26,000,000
26,100,000
26,200,000
26,300,000
0.422813
0.00449954
14.833
0.0325657
ln(x+1) 1.83847
-2.36648e-16
0.82432
-0.595107

KLF16

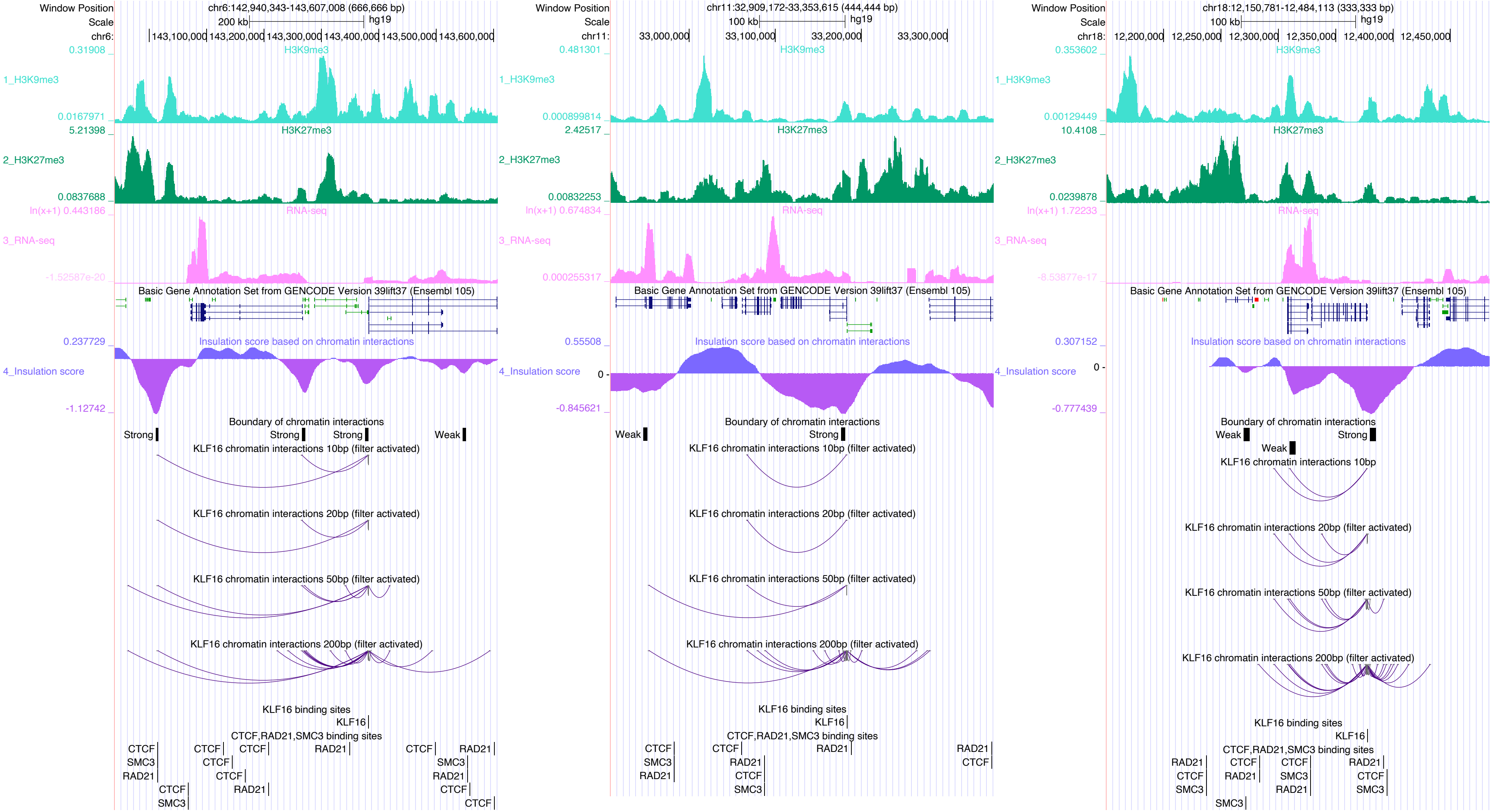

Window Position
chr6:142,940,343-143,607,008 (666,666 bp)
200 kb
hg19
H3K9me3
1_H3K9me3
H3K27me3
2_H3K27me3
RNA-seq
3_RNA-seq
Basic Gene Annotation Set from GENCODE Version 39lift37 (Ensembl 105)
Insulation score based on chromatin interactions
4_Insulation score
Boundary of chromatin interactions
KLF16 chromatin interactions 10bp (filter activated)
KLF16 chromatin interactions 20bp (filter activated)
KLF16 chromatin interactions 50bp (filter activated)
KLF16 chromatin interactions 200bp (filter activated)
KLF16 binding sites
CTCF,RAD21,SMC3 binding sites
chr11:32,909,172-33,353,615 (444,444 bp)
100 kb
chr18:12,150,781-12,484,113 (333,333 bp)
KLF16 chromatin interactions 10bp

### MAFG

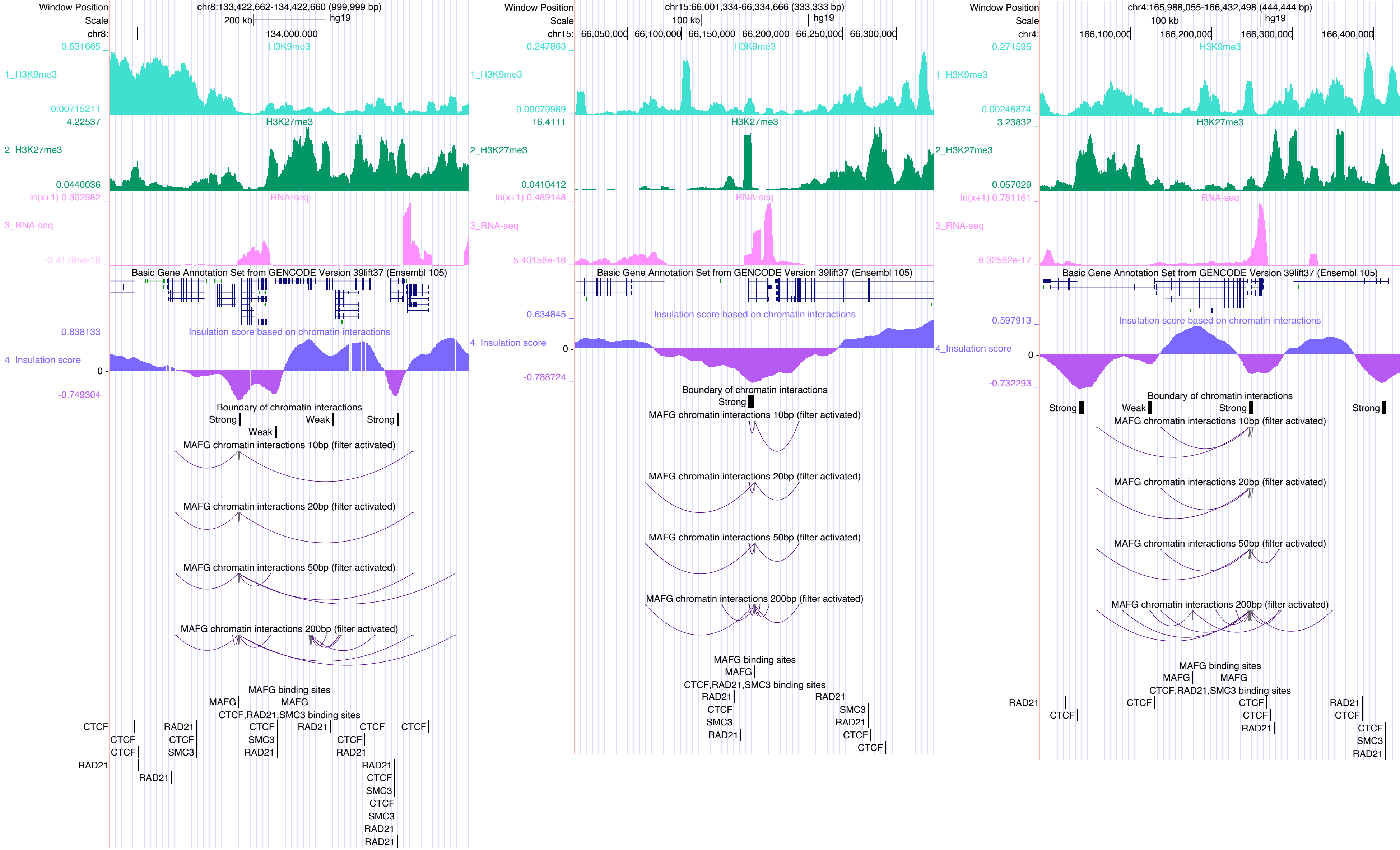
Window Position
Scale
chr8:133,422,662-134,422,660 (999,999 bp)
200 kb
hg19
134,000,000
0.531665
H3K9me3
1_H3K9me3
0.00715211
4.22537
H3K27me3
2_H3K27me3
0.0440036
ln(x+1) 0.302982
RNA-seq
3_RNA-seq
-3.41795e-18
Basic Gene Annotation Set from GENCODE Version 39lift37 (Ensembl 105)
0.838133
Insulation score based on chromatin interactions
4_Insulation score
0
-0.749304
Boundary of chromatin interactions
Strong
Weak
Strong
Weak
MAFG chromatin interactions 10bp (filter activated)
MAFG chromatin interactions 20bp (filter activated)
MAFG chromatin interactions 50bp (filter activated)
MAFG chromatin interactions 200bp (filter activated)
MAFG binding sites
MAFG
MAFG
CTCF,RAD21,SMC3 binding sites
CTCF
RAD21
CTCF
RAD21
CTCF
CTCF
CTCF
CTCF
SMC3
CTCF
SMC3
RAD21
RAD21
RAD21
RAD21
CTCF
SMC3
CTCF
SMC3
RAD21
RAD21
chr15:66,001,334-66,334,666 (333,333 bp)
100 kb
66,050,000
66,100,000
66,150,000
66,200,000
66,250,000
66,300,000
0.247863
0.00079989
16.4111
0.0410412
ln(x+1) 0.489148
5.40158e-18
0.634845
-0.788724
RAD21
CTCF
SMC3
RAD21
SMC3
RAD21
CTCF
CTCF
chr4:165,988,055-166,432,498 (444,444 bp)
100 kb
166,100,000
166,200,000
166,300,000
166,400,000
0.271595
0.00248874
3.23832
0.057029
ln(x+1) 0.781161
6.32582e-17
0.597913
-0.732293
RAD21
CTCF
CTCF
CTCF
CTCF
RAD21
RAD21
CTCF
CTCF
SMC3
RAD21

MAZ

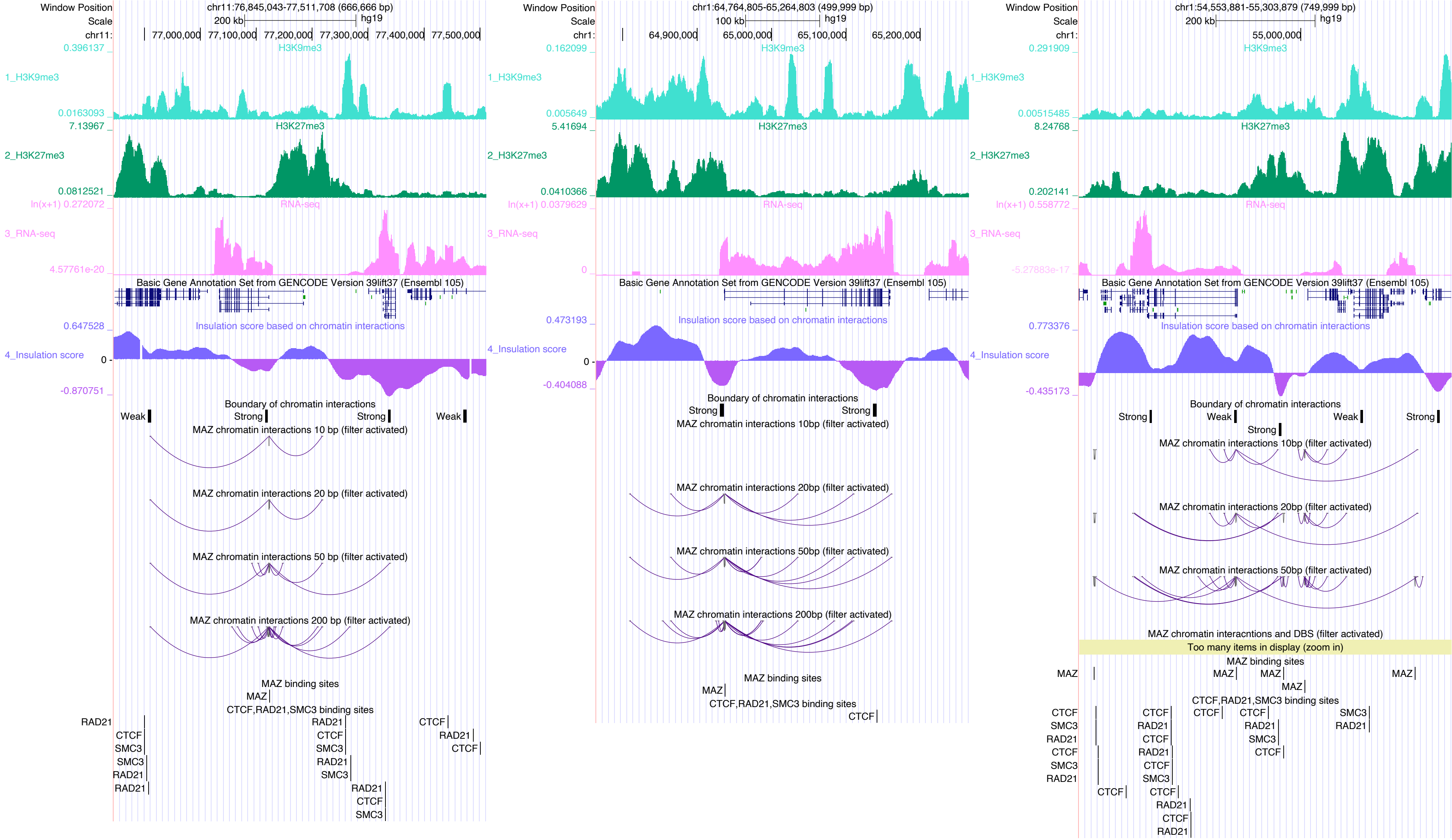
Window Position
chr11:76,845,043-77,511,708 (666,666 bp)
chr1:64,764,805-65,264,803 (499,999 bp)
chr1:54,553,881-55,303,879 (749,999 bp)
Scale
hg19
1_H3K9me3
2_H3K27me3
3_RNA-seq
4_Insulation score
Basic Gene Annotation Set from GENCODE Version 39lift37 (Ensembl 105)
Insulation score based on chromatin interactions
Boundary of chromatin interactions
Strong
Weak
MAZ chromatin interactions 10 bp (filter activated)
MAZ chromatin interactions 20 bp (filter activated)
MAZ chromatin interactions 50 bp (filter activated)
MAZ chromatin interactions 200 bp (filter activated)
MAZ chromatin interacntions and DBS (filter activated)
Too many items in display (zoom in)
MAZ binding sites
CTCF,RAD21,SMC3 binding sites
CTCF
RAD21
SMC3

### MECP2

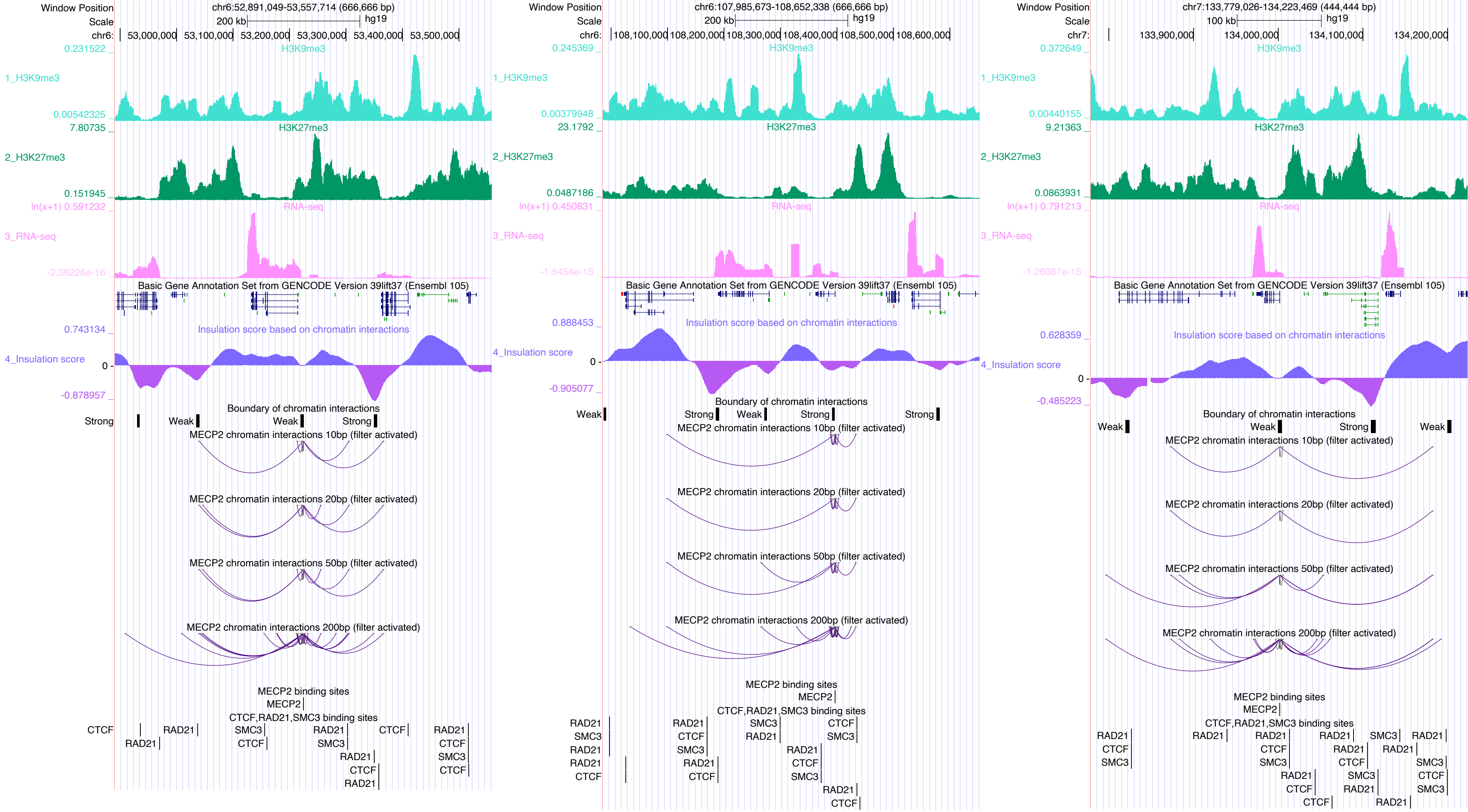
Window Position
Scale
chr6:52,891,049-53,557,714 (666,666 bp)
200 kb
hg19
chr6: 53,000,000 53,100,000 53,200,000 53,300,000 53,400,000 53,500,000
0.231522
H3K9me3
1_H3K9me3
0.00542325
7.80735
H3K27me3
2_H3K27me3
0.151945
ln(x+1) 0.591232
RNA-seq
3_RNA-seq
-2.36226e-16
Basic Gene Annotation Set from GENCODE Version 39lift37 (Ensembl 105)
0.743134
Insulation score based on chromatin interactions
4_Insulation score
0
-0.878957
Boundary of chromatin interactions
Strong
Weak
Weak
Strong
MECP2 chromatin interactions 10bp (filter activated)
MECP2 chromatin interactions 20bp (filter activated)
MECP2 chromatin interactions 50bp (filter activated)
MECP2 chromatin interactions 200bp (filter activated)
MECP2 binding sites
MECP2
CTCF,RAD21,SMC3 binding sites
CTCF
RAD21
RAD21
SMC3
CTCF
RAD21
SMC3
RAD21
CTCF
RAD21
CTCF
RAD21
CTCF
SMC3
CTCF
Window Position
Scale
chr6:107,985,673-108,652,338 (666,666 bp)
200 kb
hg19
chr6: 108,100,000 108,200,000 108,300,000 108,400,000 108,500,000 108,600,000
0.245369
H3K9me3
1_H3K9me3
0.00379948
23.1792
H3K27me3
2_H3K27me3
0.0487186
ln(x+1) 0.450831
RNA-seq
3_RNA-seq
-1.5454e-16
Basic Gene Annotation Set from GENCODE Version 39lift37 (Ensembl 105)
0.888453
Insulation score based on chromatin interactions
4_Insulation score
0
-0.905077
Boundary of chromatin interactions
Weak
Strong
Weak
Strong
Strong
MECP2 chromatin interactions 10bp (filter activated)
MECP2 chromatin interactions 20bp (filter activated)
MECP2 chromatin interactions 50bp (filter activated)
MECP2 chromatin interactions 200bp (filter activated)
MECP2 binding sites
MECP2
CTCF,RAD21,SMC3 binding sites
RAD21
SMC3
RAD21
RAD21
CTCF
RAD21
CTCF
SMC3
RAD21
CTCF
SMC3
RAD21
CTCF
RAD21
CTCF
SMC3
RAD21
CTCF
Window Position
Scale
chr7:133,779,026-134,223,469 (444,444 bp)
100 kb
hg19
chr7: 133,900,000 134,000,000 134,100,000 134,200,000
0.372649
H3K9me3
1_H3K9me3
0.00440155
9.21363
H3K27me3
2_H3K27me3
0.0863931
ln(x+1) 0.791213
RNA-seq
3_RNA-seq
-1.26987e-16
Basic Gene Annotation Set from GENCODE Version 39lift37 (Ensembl 105)
0.628359
Insulation score based on chromatin interactions
4_Insulation score
0
-0.485223
Boundary of chromatin interactions
Weak
Weak
Strong
Weak
MECP2 chromatin interactions 10bp (filter activated)
MECP2 chromatin interactions 20bp (filter activated)
MECP2 chromatin interactions 50bp (filter activated)
MECP2 chromatin interactions 200bp (filter activated)
MECP2 binding sites
MECP2
CTCF,RAD21,SMC3 binding sites
RAD21
CTCF
SMC3
RAD21
RAD21
CTCF
SMC3
RAD21
CTCF
RAD21
RAD21
CTCF
SMC3
RAD21
SMC3
RAD21
RAD21
SMC3
CTCF
SMC3
CTCF
RAD21

### MEF2B

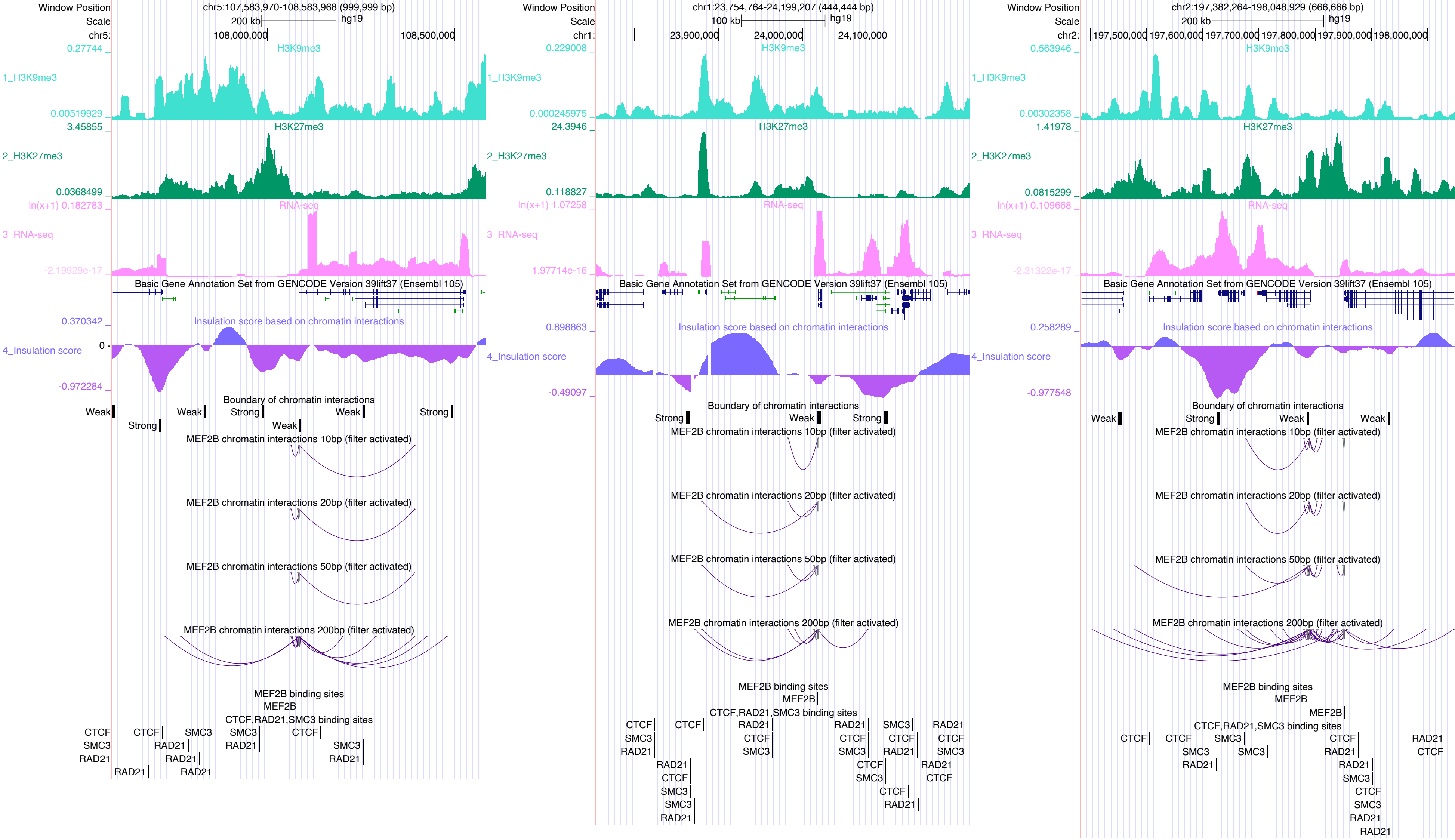

Window Position
chr5:107,583,970-108,583,968 (999,999 bp)
Scale
200 kb
hg19
chr5:
108,000,000
108,500,000
1_H3K9me3
H3K9me3
2_H3K27me3
H3K27me3
3_RNA-seq
RNA-seq
Basic Gene Annotation Set from GENCODE Version 39lift37 (Ensembl 105)
4_Insulation score
Insulation score based on chromatin interactions
Boundary of chromatin interactions
Weak
Strong
MEF2B chromatin interactions 10bp (filter activated)
MEF2B chromatin interactions 20bp (filter activated)
MEF2B chromatin interactions 50bp (filter activated)
MEF2B chromatin interactions 200bp (filter activated)
MEF2B binding sites
MEF2B
CTCF,RAD21,SMC3 binding sites
CTCF
SMC3
RAD21
chr1:23,754,764-24,199,207 (444,444 bp)
100 kb
chr1:
23,900,000
24,000,000
24,100,000
chr2:197,382,264-198,048,929 (666,666 bp)
chr2:
197,500,000
197,600,000
197,700,000
197,800,000
197,900,000
198,000,000

### MEIS2

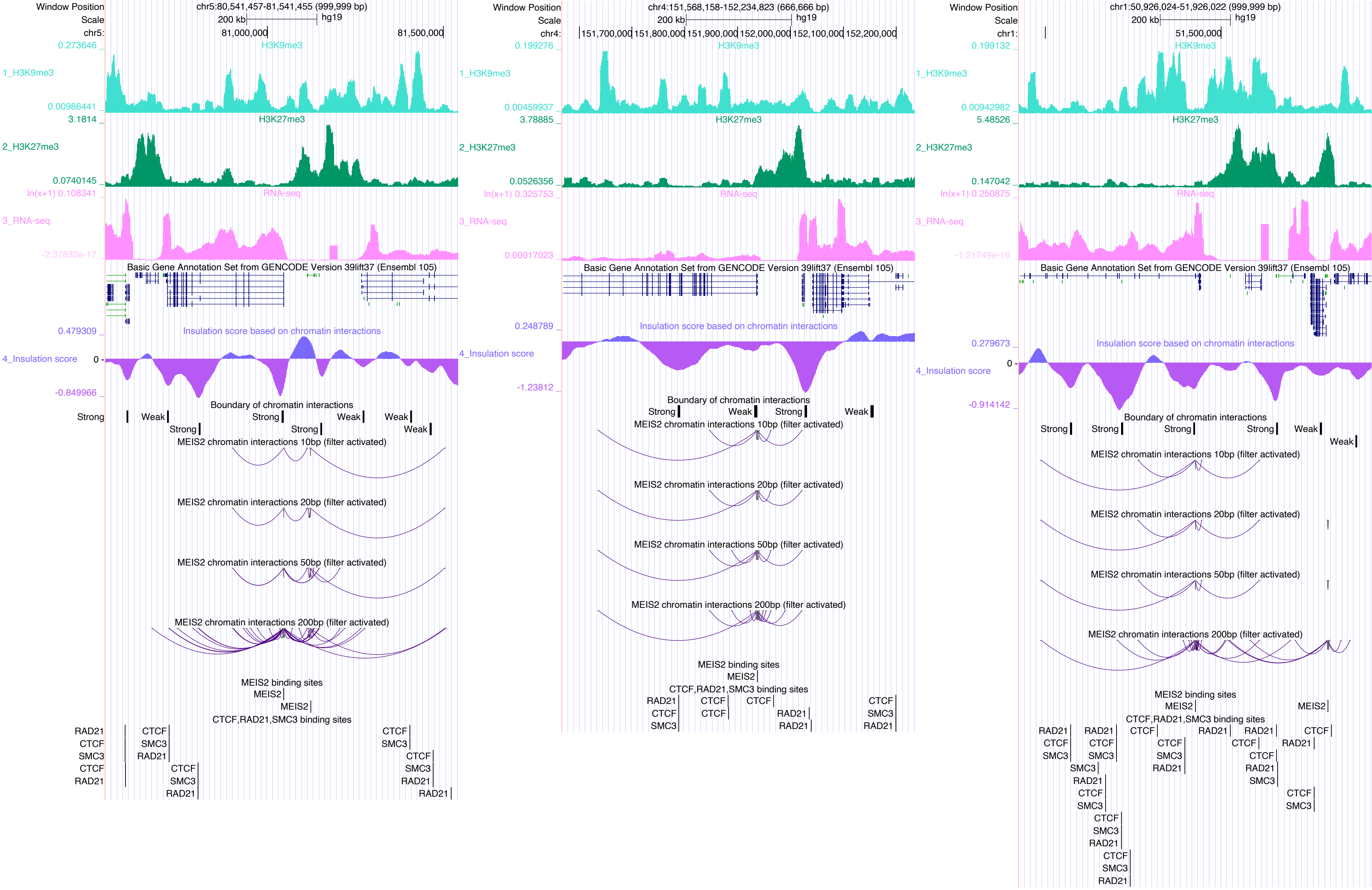

Window Position
Scale
chr5:80,541,457-81,541,455 (999,999 bp)
200 kb
hg19
chr5:
81,000,000
81,500,000
1_H3K9me3
H3K9me3
2_H3K27me3
H3K27me3
3_RNA-seq
RNA-seq
Basic Gene Annotation Set from GENCODE Version 39lift37 (Ensembl 105)
4_Insulation score
Insulation score based on chromatin interactions
Boundary of chromatin interactions
Strong
Weak
MEIS2 chromatin interactions 10bp (filter activated)
MEIS2 chromatin interactions 20bp (filter activated)
MEIS2 chromatin interactions 50bp (filter activated)
MEIS2 chromatin interactions 200bp (filter activated)
MEIS2 binding sites
MEIS2
CTCF,RAD21,SMC3 binding sites
RAD21
CTCF
SMC3
chr4:151,568,158-152,234,823 (666,666 bp)
chr4:
151,700,000
151,800,000
151,900,000
152,000,000
152,100,000
152,200,000
chr1:50,926,024-51,926,022 (999,999 bp)
chr1:
51,500,000

### MYB

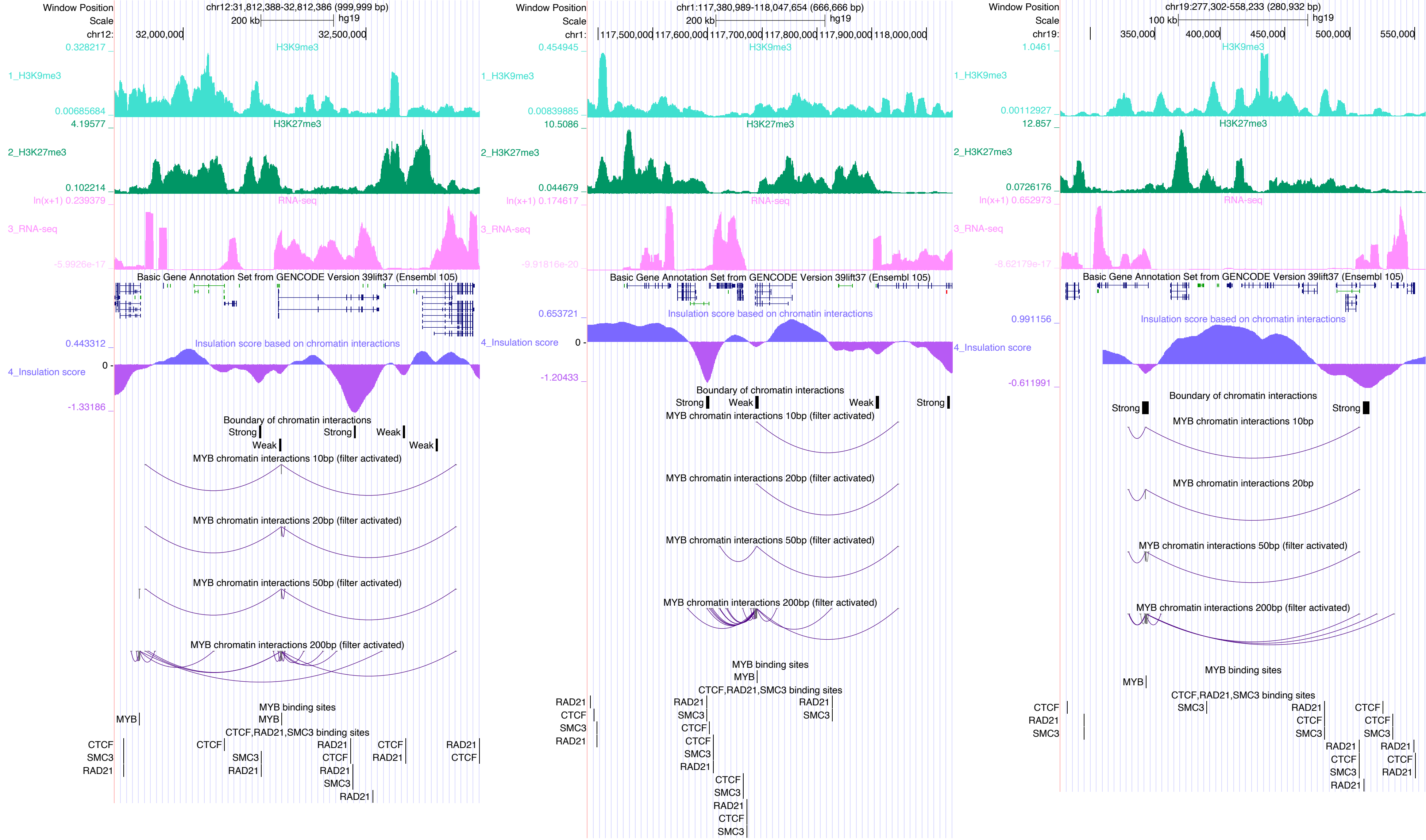

Window Position
chr12:31,812,388-32,812,386 (999,999 bp)
chr1:117,380,989-118,047,654 (666,666 bp)
chr19:277,302-558,233 (280,932 bp)
H3K9me3
H3K27me3
RNA-seq
Basic Gene Annotation Set from GENCODE Version 39lift37 (Ensembl 105)
Insulation score based on chromatin interactions
Boundary of chromatin interactions
MYB chromatin interactions 10bp (filter activated)
MYB chromatin interactions 20bp (filter activated)
MYB chromatin interactions 50bp (filter activated)
MYB chromatin interactions 200bp (filter activated)
MYB binding sites
CTCF,RAD21,SMC3 binding sites

MYCN

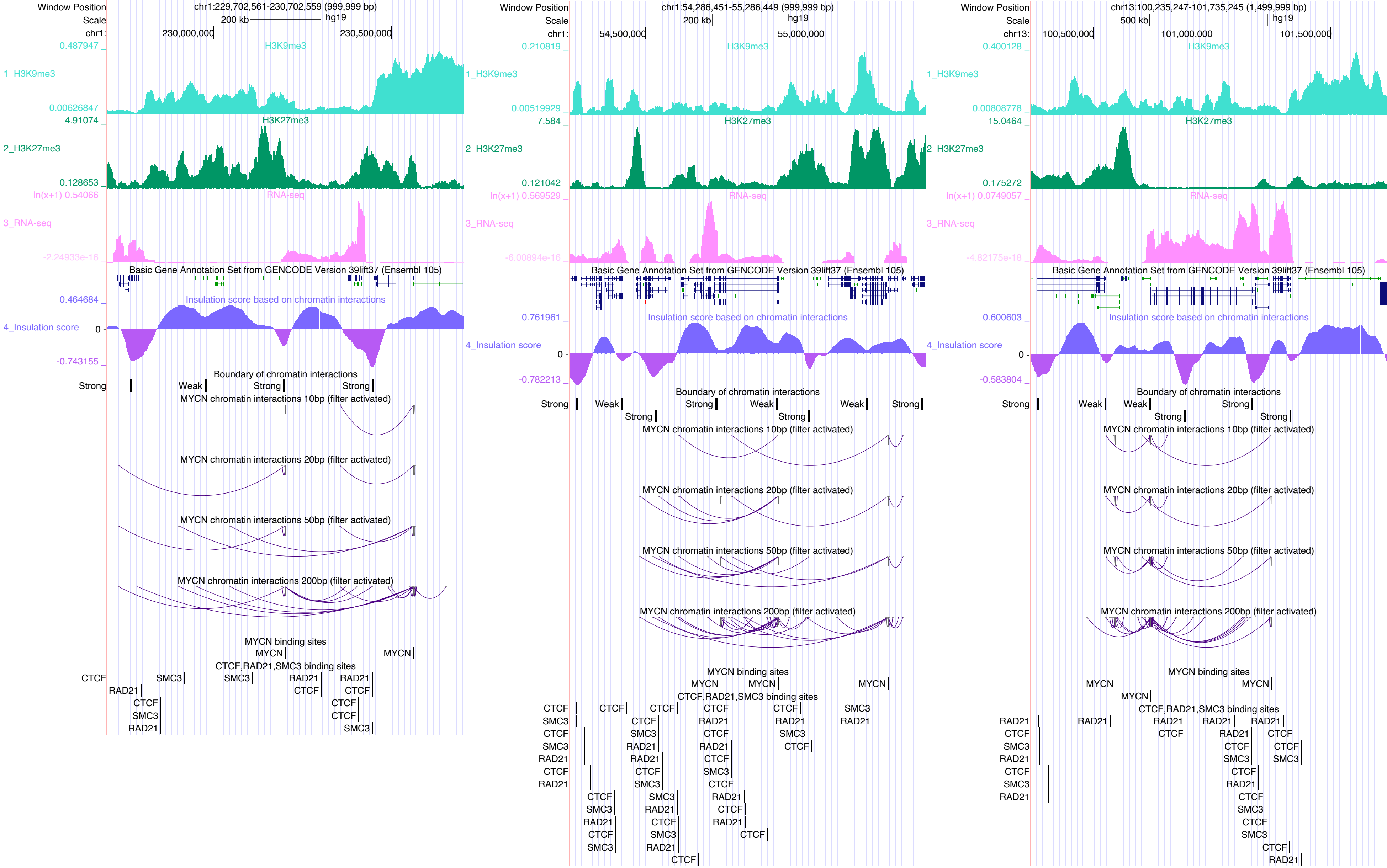

chr1:229,702,561-230,702,559 (999,999 bp)
chr1:54,286,451-55,286,449 (999,999 bp)
chr13:100,235,247-101,735,245 (1,499,999 bp)
1_H3K9me3
2_H3K27me3
3_RNA-seq
4_Insulation score
Basic Gene Annotation Set from GENCODE Version 39lift37 (Ensembl 105)
Insulation score based on chromatin interactions
Boundary of chromatin interactions
MYCN chromatin interactions 10bp (filter activated)
MYCN chromatin interactions 20bp (filter activated)
MYCN chromatin interactions 50bp (filter activated)
MYCN chromatin interactions 200bp (filter activated)
MYCN binding sites
CTCF,RAD21,SMC3 binding sites

MYOD1

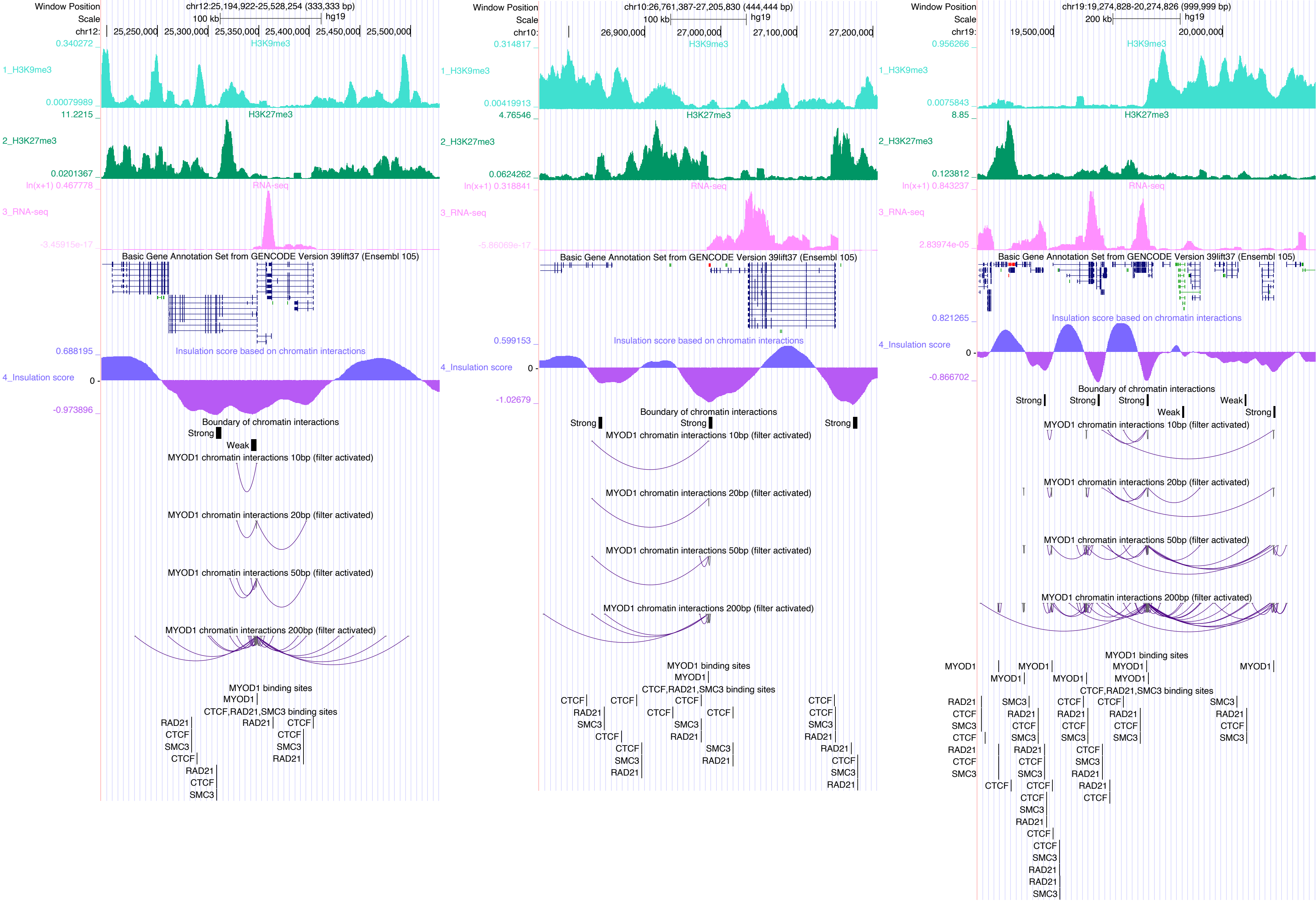

Window Position
chr12:25,194,922-25,528,254 (333,333 bp)
Scale
100 kb
hg19
1_H3K9me3
2_H3K27me3
3_RNA-seq
4_Insulation score
Basic Gene Annotation Set from GENCODE Version 39lift37 (Ensembl 105)
Insulation score based on chromatin interactions
Boundary of chromatin interactions
MYOD1 chromatin interactions 10bp (filter activated)
MYOD1 chromatin interactions 20bp (filter activated)
MYOD1 chromatin interactions 50bp (filter activated)
MYOD1 chromatin interactions 200bp (filter activated)
MYOD1 binding sites
CTCF,RAD21,SMC3 binding sites
chr10:26,761,387-27,205,830 (444,444 bp)
chr19:19,274,828-20,274,826 (999,999 bp)
200 kb

### MYOG

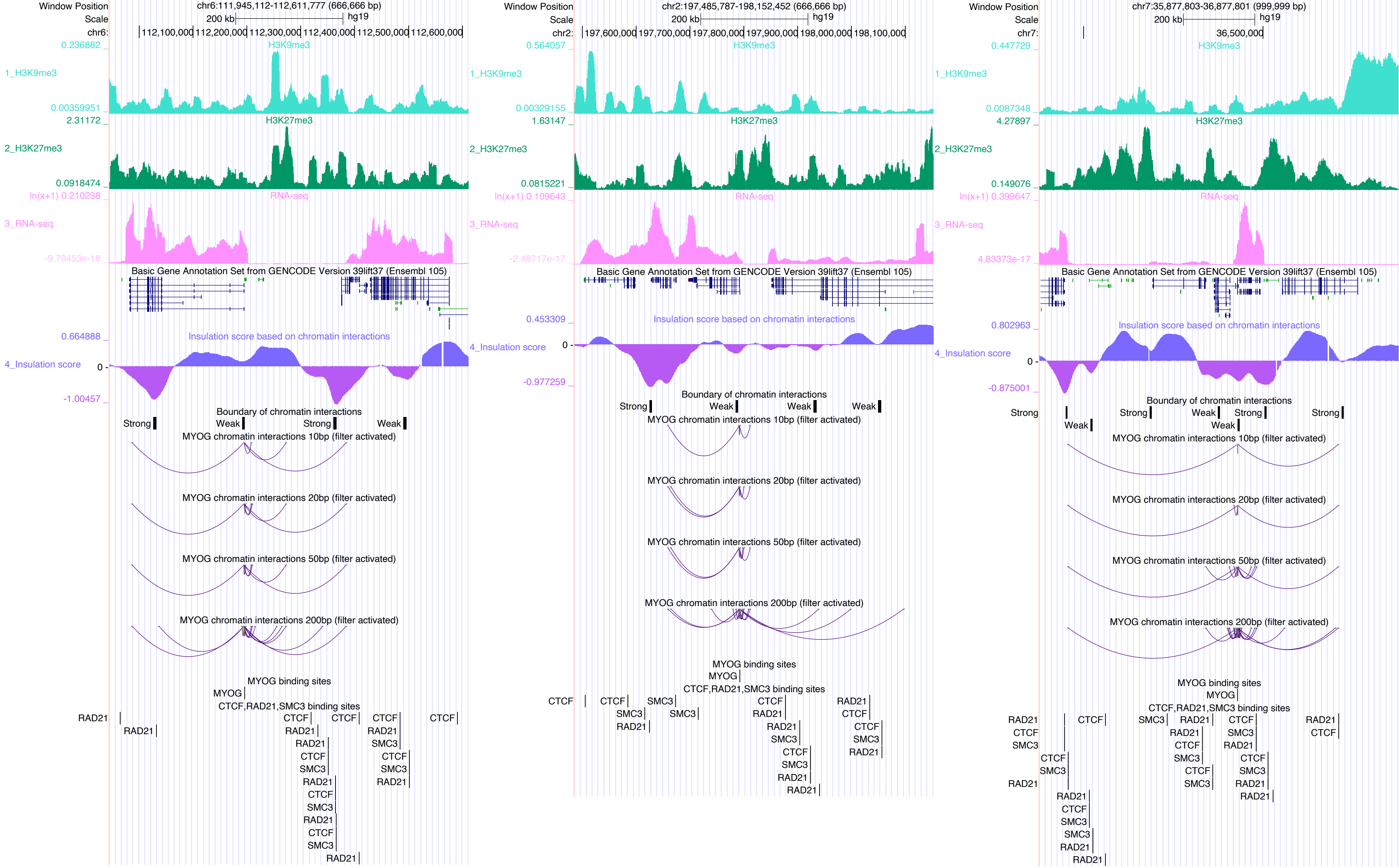

Window Position
Scale
chr6:111,945,112-112,611,777 (666,666 bp)
200 kb
hg19
chr6:
112,100,000
112,200,000
112,300,000
112,400,000
112,500,000
112,600,000
1_H3K9me3
H3K9me3
0.236882
0.00359951
2_H3K27me3
H3K27me3
2.31172
0.0918474
3_RNA-seq
RNA-seq
ln(x+1) 0.210238
-9.70453e-18
Basic Gene Annotation Set from GENCODE Version 39lift37 (Ensembl 105)
4_Insulation score
Insulation score based on chromatin interactions
0.664888
0
-1.00457
Boundary of chromatin interactions
Strong
Weak
Strong
Weak
MYOG chromatin interactions 10bp (filter activated)
MYOG chromatin interactions 20bp (filter activated)
MYOG chromatin interactions 50bp (filter activated)
MYOG chromatin interactions 200bp (filter activated)
MYOG binding sites
MYOG
CTCF,RAD21,SMC3 binding sites
Window Position
Scale
chr2:197,485,787-198,152,452 (666,666 bp)
200 kb
hg19
chr2:
197,600,000
197,700,000
197,800,000
197,900,000
198,000,000
198,100,000
0.564057
0.00329155
1.63147
0.0815221
ln(x+1) 0.109643
-2.48717e-17
0.453309
-0.977259
Window Position
Scale
chr7:35,877,803-36,877,801 (999,999 bp)
200 kb
hg19
chr7:
36,500,000
0.447729
0.0087348
4.27897
0.149076
ln(x+1) 0.399647
4.83373e-17
0.802963
-0.875001

### NFYC

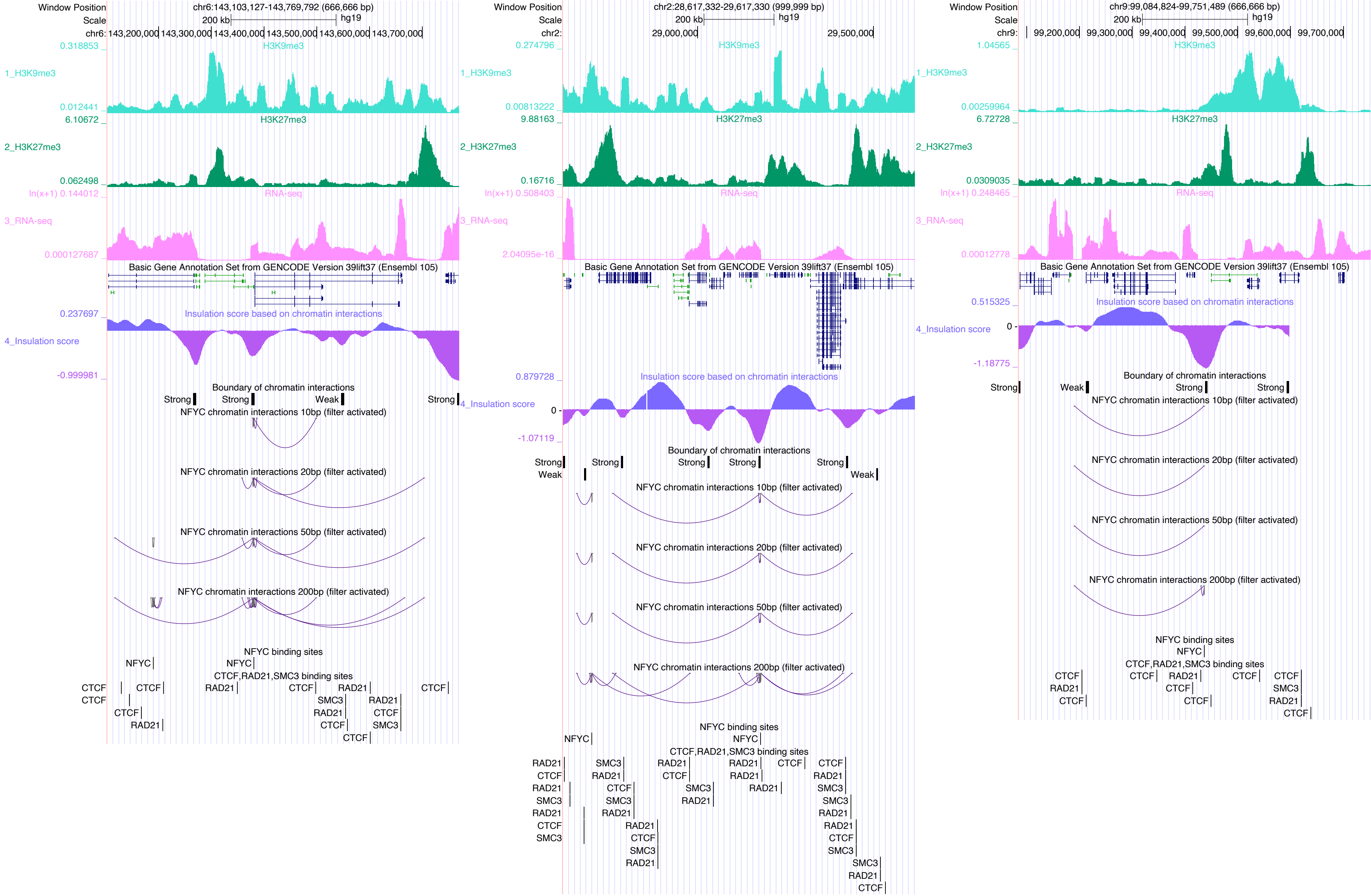

Window Position
Scale
chr6:143,103,127-143,769,792 (666,666 bp)
200 kb
hg19
chr6: 143,200,000 143,300,000 143,400,000 143,500,000 143,600,000 143,700,000
1_H3K9me3
H3K9me3
0.318853
0.012441
2_H3K27me3
H3K27me3
6.10672
0.062498
3_RNA-seq
RNA-seq
ln(x+1) 0.144012
0.000127687
Basic Gene Annotation Set from GENCODE Version 39lift37 (Ensembl 105)
4_Insulation score
Insulation score based on chromatin interactions
0.237697
-0.999981
Boundary of chromatin interactions
Strong
Strong
Weak
Strong
NFYC chromatin interactions 10bp (filter activated)
NFYC chromatin interactions 20bp (filter activated)
NFYC chromatin interactions 50bp (filter activated)
NFYC chromatin interactions 200bp (filter activated)
NFYC binding sites
NFYC
NFYC
CTCF,RAD21,SMC3 binding sites
CTCF
CTCF
CTCF
CTCF
RAD21
RAD21
CTCF
RAD21
SMC3
RAD21
RAD21
CTCF
CTCF
SMC3
CTCF
CTCF
chr2:28,617,332-29,617,330 (999,999 bp)
200 kb
hg19
chr2: 29,000,000 29,500,000
0.274796
0.00813222
9.88163
0.16716
ln(x+1) 0.508403
2.04095e-16
0.879728
0
-1.07119
Boundary of chromatin interactions
Strong
Weak
Strong
Strong
Strong
Strong
Weak
chr9:99,084,824-99,751,489 (666,666 bp)
200 kb
hg19
chr9: 99,200,000 99,300,000 99,400,000 99,500,000 99,600,000 99,700,000
1.04565
0.00259964
6.72728
0.0309035
ln(x+1) 0.248465
0.00012778
0.515325
0
-1.18775
Boundary of chromatin interactions
Strong
Weak
Strong
Strong

NR2C2

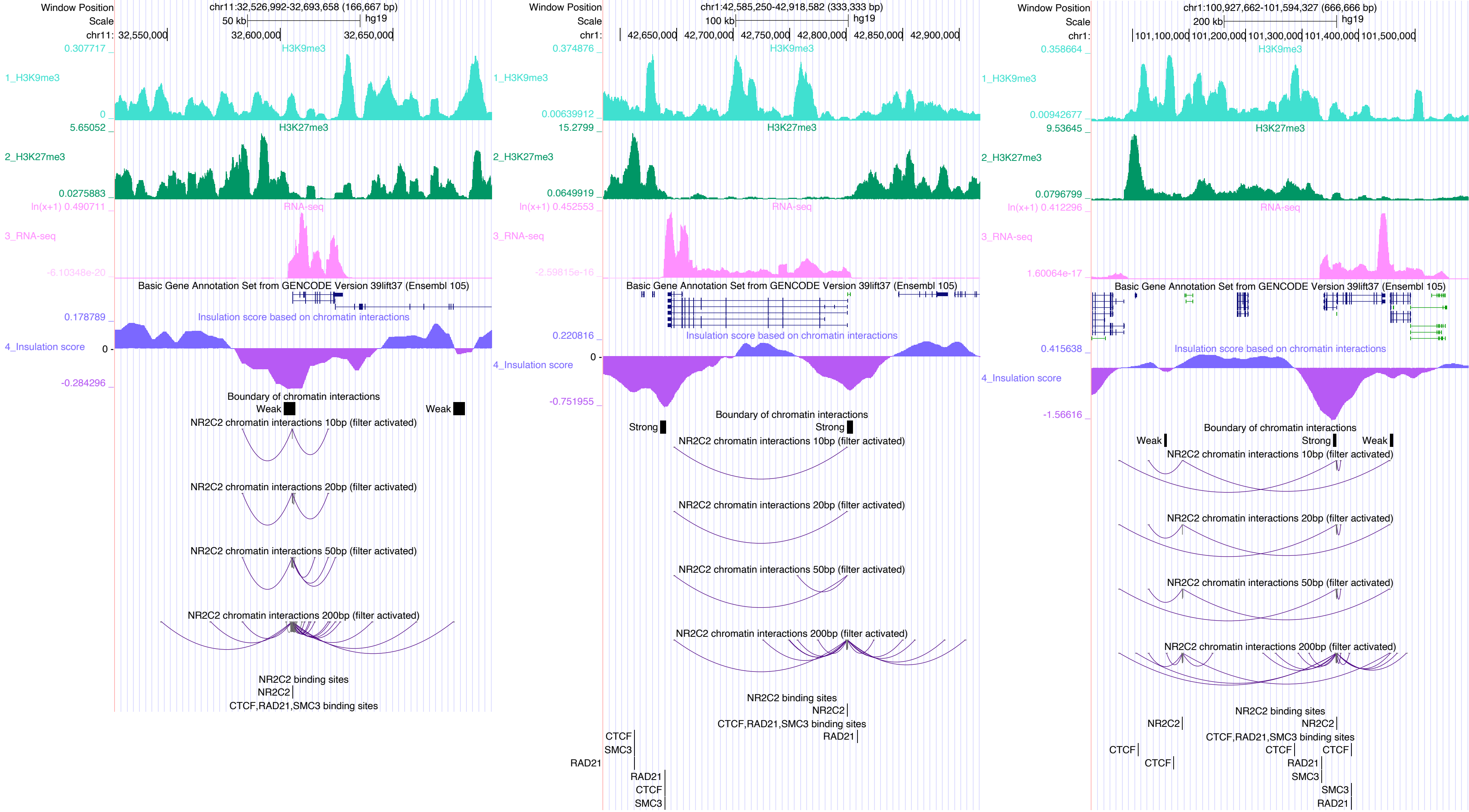
Window Position chr11:32,526,992-32,693,658 (166,667 bp)
Scale 50 kb hg19
1_H3K9me3
2_H3K27me3
3_RNA-seq
Basic Gene Annotation Set from GENCODE Version 39lift37 (Ensembl 105)
4_Insulation score
Insulation score based on chromatin interactions
Boundary of chromatin interactions
Weak
NR2C2 chromatin interactions 10bp (filter activated)
NR2C2 chromatin interactions 20bp (filter activated)
NR2C2 chromatin interactions 50bp (filter activated)
NR2C2 chromatin interactions 200bp (filter activated)
NR2C2 binding sites
CTCF,RAD21,SMC3 binding sites
Window Position chr1:42,585,250-42,918,582 (333,333 bp)
Scale 100 kb hg19
Strong
CTCF
SMC3
RAD21
Window Position chr1:100,927,662-101,594,327 (666,666 bp)
Scale 200 kb hg19

# NR2F2

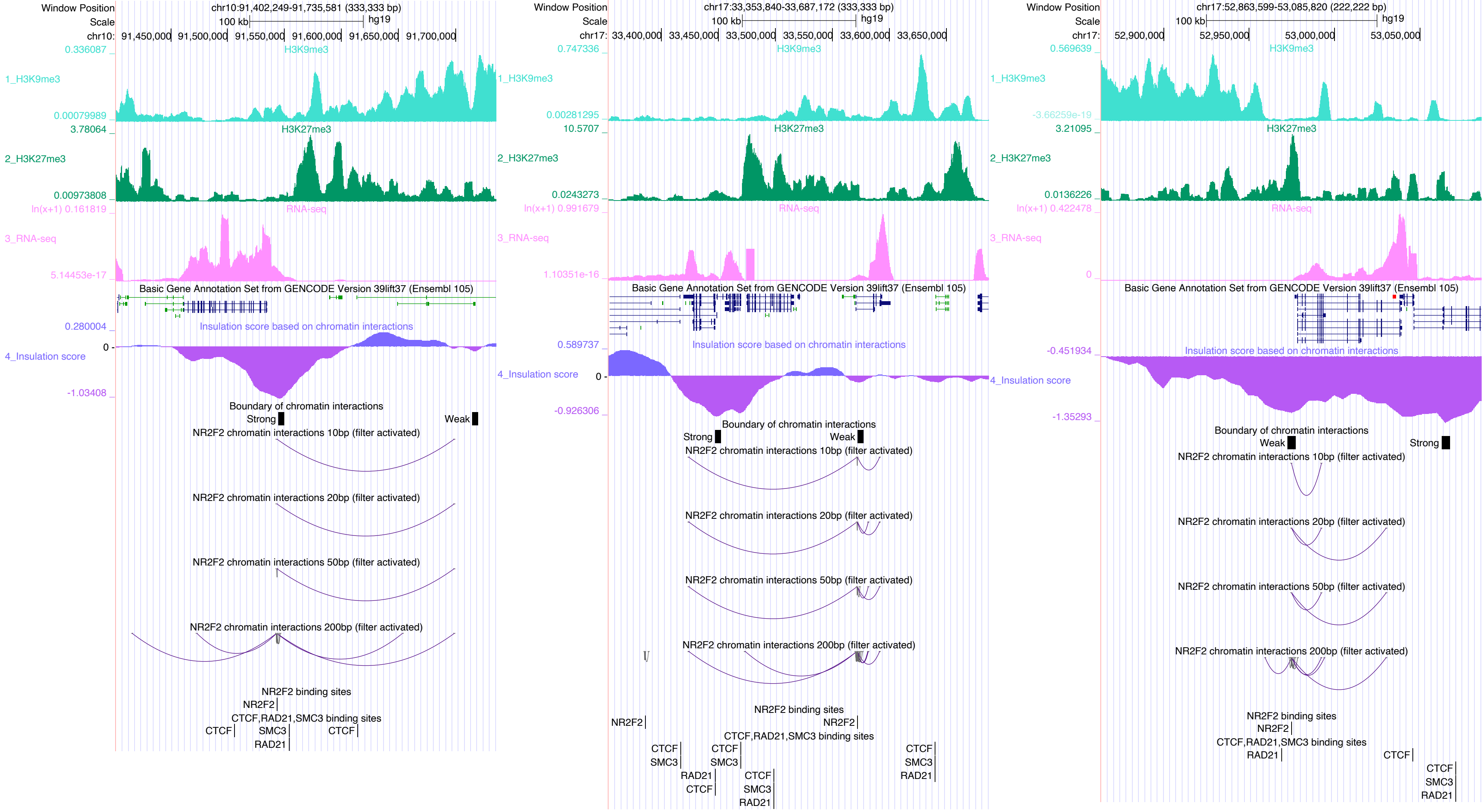

Window Position
chr10:91,402,249-91,735,581 (333,333 bp)
Scale
100 kb
hg19
chr10: 91,450,000 91,500,000 91,550,000 91,600,000 91,650,000 91,700,000
0.336087
H3K9me3
1_H3K9me3
0.00079989
3.78064
H3K27me3
2_H3K27me3
0.00973808
ln(x+1) 0.161819
RNA-seq
3_RNA-seq
5.14453e-17
Basic Gene Annotation Set from GENCODE Version 39lift37 (Ensembl 105)
0.280004
Insulation score based on chromatin interactions
4_Insulation score
0
-1.03408
Boundary of chromatin interactions
Strong
Weak
NR2F2 chromatin interactions 10bp (filter activated)
NR2F2 chromatin interactions 20bp (filter activated)
NR2F2 chromatin interactions 50bp (filter activated)
NR2F2 chromatin interactions 200bp (filter activated)
NR2F2 binding sites
NR2F2
CTCF,RAD21,SMC3 binding sites
CTCF
SMC3
RAD21
CTCF
Window Position
chr17:33,353,840-33,687,172 (333,333 bp)
Scale
100 kb
hg19
chr17: 33,400,000 33,450,000 33,500,000 33,550,000 33,600,000 33,650,000
0.747336
H3K9me3
1_H3K9me3
0.00281295
10.5707
H3K27me3
2_H3K27me3
0.0243273
ln(x+1) 0.991679
RNA-seq
3_RNA-seq
1.10351e-16
Basic Gene Annotation Set from GENCODE Version 39lift37 (Ensembl 105)
0.589737
Insulation score based on chromatin interactions
4_Insulation score
0
-0.926306
Boundary of chromatin interactions
Strong
Weak
NR2F2 chromatin interactions 10bp (filter activated)
NR2F2 chromatin interactions 20bp (filter activated)
NR2F2 chromatin interactions 50bp (filter activated)
NR2F2 chromatin interactions 200bp (filter activated)
NR2F2 binding sites
NR2F2
NR2F2
CTCF,RAD21,SMC3 binding sites
CTCF
SMC3
CTCF
SMC3
RAD21
CTCF
CTCF
SMC3
RAD21
CTCF
SMC3
RAD21
Window Position
chr17:52,863,599-53,085,820 (222,222 bp)
Scale
100 kb
hg19
chr17: 52,900,000 52,950,000 53,000,000 53,050,000
0.569639
H3K9me3
1_H3K9me3
-3.66259e-19
3.21095
H3K27me3
2_H3K27me3
0.0136226
ln(x+1) 0.422478
RNA-seq
3_RNA-seq
0
Basic Gene Annotation Set from GENCODE Version 39lift37 (Ensembl 105)
-0.451934
Insulation score based on chromatin interactions
4_Insulation score
-1.35293
Boundary of chromatin interactions
Weak
Strong
NR2F2 chromatin interactions 10bp (filter activated)
NR2F2 chromatin interactions 20bp (filter activated)
NR2F2 chromatin interactions 50bp (filter activated)
NR2F2 chromatin interactions 200bp (filter activated)
NR2F2 binding sites
NR2F2
CTCF,RAD21,SMC3 binding sites
RAD21
CTCF
CTCF
SMC3
RAD21

# NR3C1

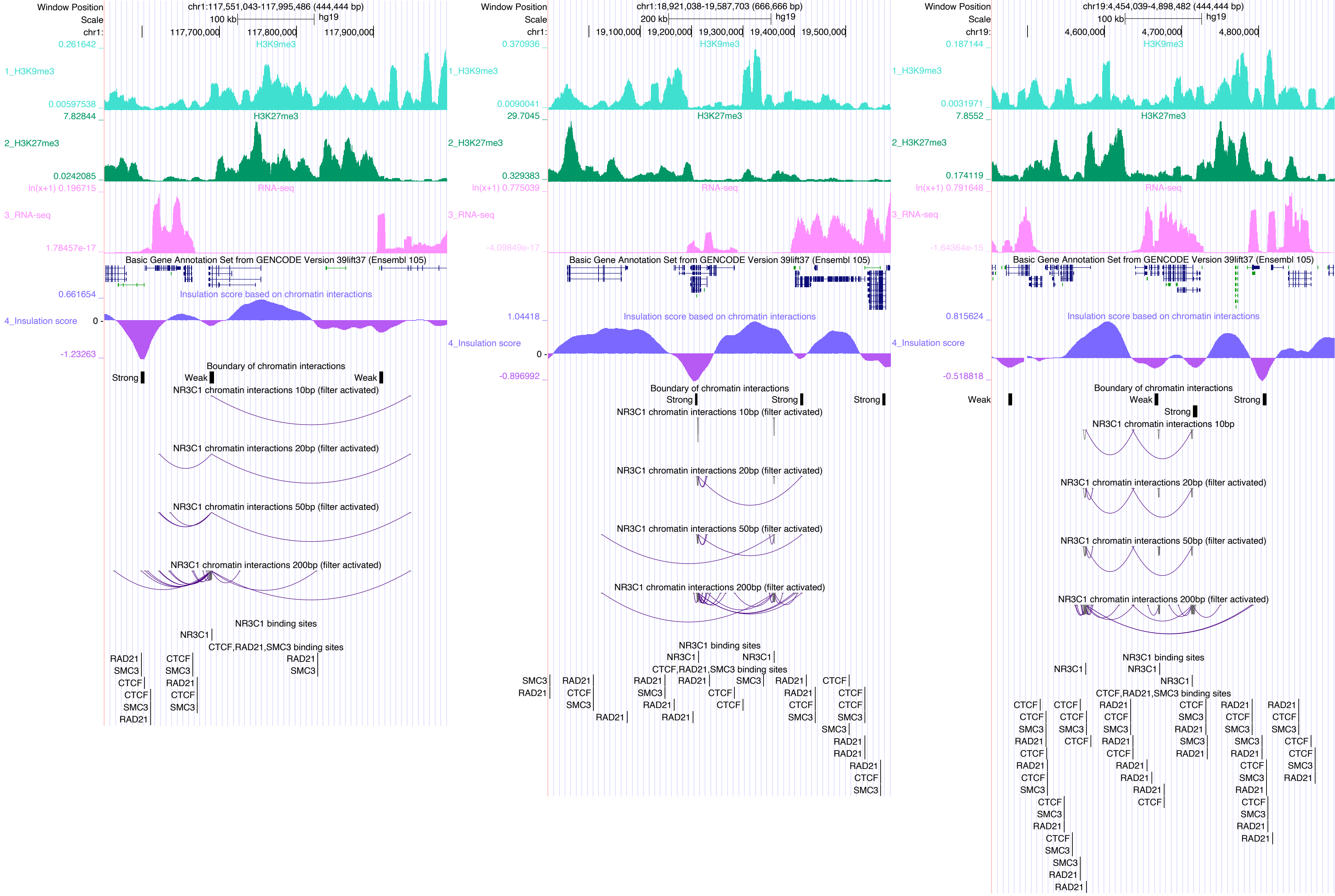

chr1:117,551,043-117,995,486 (444,444 bp)
H3K9me3
H3K27me3
RNA-seq
Basic Gene Annotation Set from GENCODE Version 39lift37 (Ensembl 105)
Insulation score based on chromatin interactions
Boundary of chromatin interactions
NR3C1 chromatin interactions 10bp (filter activated)
NR3C1 chromatin interactions 20bp (filter activated)
NR3C1 chromatin interactions 50bp (filter activated)
NR3C1 chromatin interactions 200bp (filter activated)
NR3C1 binding sites
CTCF,RAD21,SMC3 binding sites
chr1:18,921,038-19,587,703 (666,666 bp)
chr19:4,454,039-4,898,482 (444,444 bp)
NR3C1 chromatin interactions 10bp

# NR5A2

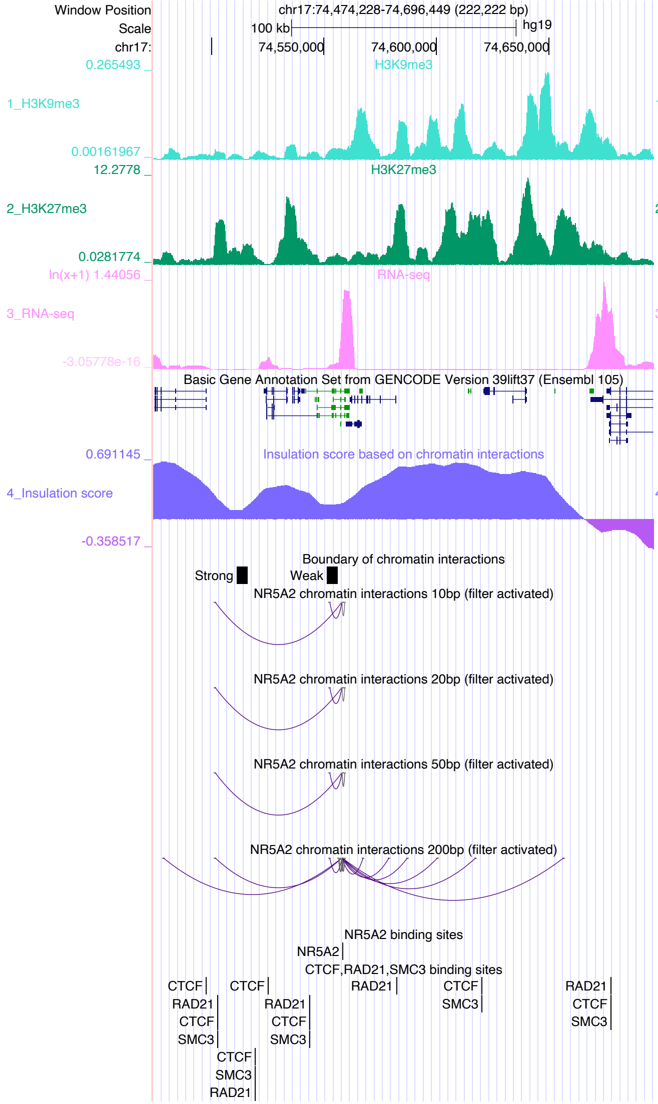

Window Position
chr17:74,474,228-74,696,449 (222,222 bp)
Scale
100 kb
hg19
chr17:
74,550,000
74,600,000
74,650,000
1_H3K9me3
H3K9me3
2_H3K27me3
H3K27me3
3_RNA-seq
RNA-seq
Basic Gene Annotation Set from GENCODE Version 39lift37 (Ensembl 105)
4_Insulation score
Insulation score based on chromatin interactions
Boundary of chromatin interactions
Strong
Weak
NR5A2 chromatin interactions 10bp (filter activated)
NR5A2 chromatin interactions 20bp (filter activated)
NR5A2 chromatin interactions 50bp (filter activated)
NR5A2 chromatin interactions 200bp (filter activated)
NR5A2 binding sites
NR5A2
CTCF,RAD21,SMC3 binding sites
CTCF
RAD21
SMC3


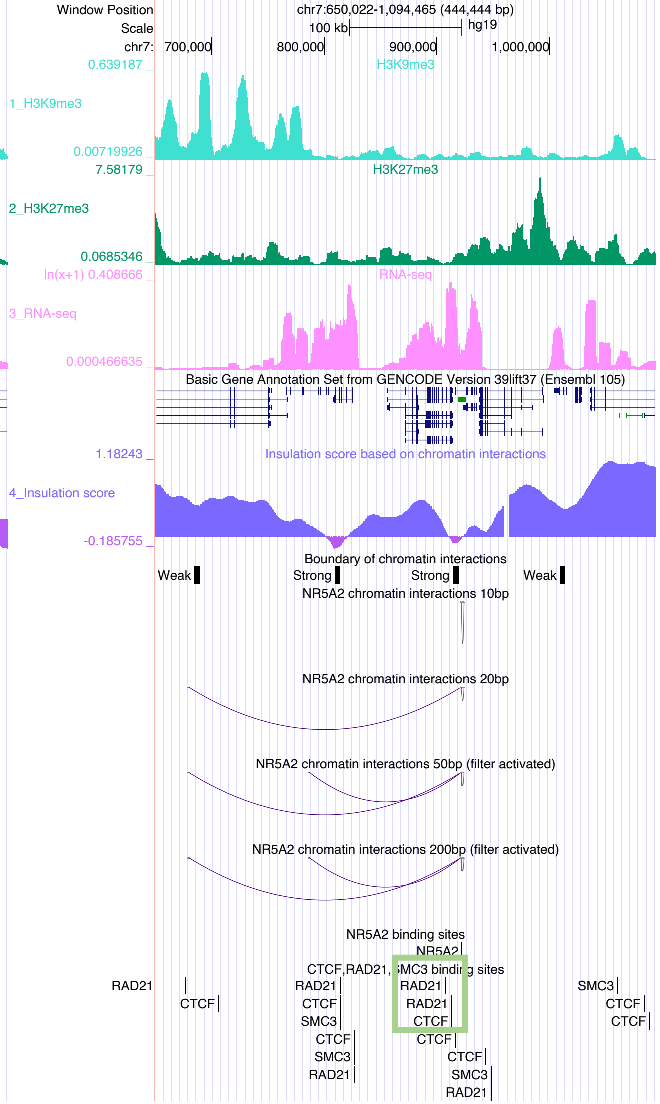

Window Position
chr7:650,022-1,094,465 (444,444 bp)
Scale
100 kb
hg19
chr7:
700,000
800,000
900,000
1,000,000
1_H3K9me3
H3K9me3
2_H3K27me3
H3K27me3
3_RNA-seq
RNA-seq
Basic Gene Annotation Set from GENCODE Version 39lift37 (Ensembl 105)
4_Insulation score
Insulation score based on chromatin interactions
Boundary of chromatin interactions
Weak
Strong
NR5A2 chromatin interactions 10bp
NR5A2 chromatin interactions 20bp
NR5A2 chromatin interactions 50bp (filter activated)
NR5A2 chromatin interactions 200bp (filter activated)
NR5A2 binding sites
NR5A2
CTCF,RAD21,SMC3 binding sites
RAD21
CTCF
SMC3

### PAX5

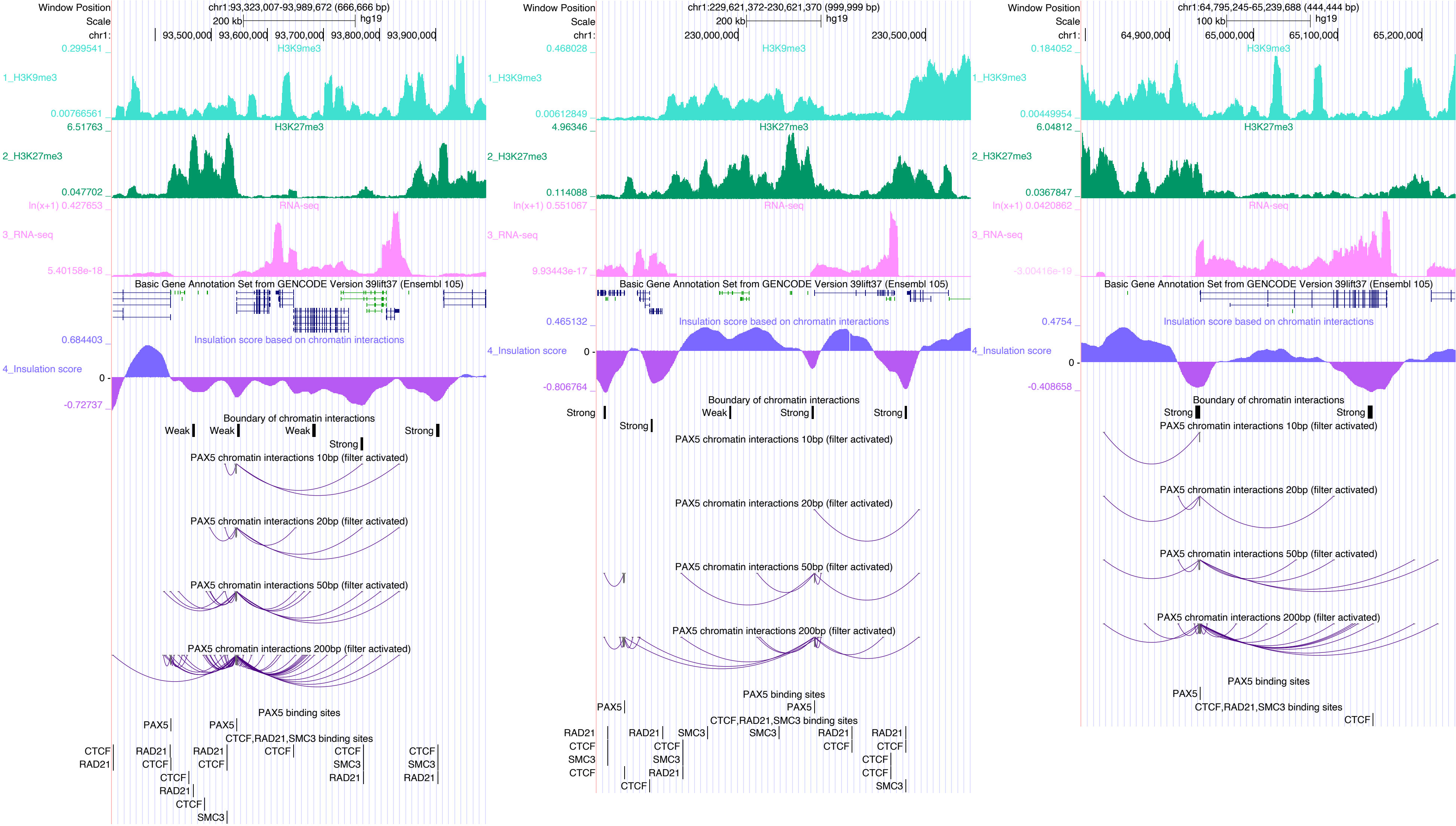

Window Position
chr1:93,323,007-93,989,672 (666,666 bp)
Scale
200 kb
hg19
chr1:
93,500,000
93,600,000
93,700,000
93,800,000
93,900,000
H3K9me3
1_H3K9me3
H3K27me3
2_H3K27me3
RNA-seq
3_RNA-seq
Basic Gene Annotation Set from GENCODE Version 39lift37 (Ensembl 105)
Insulation score based on chromatin interactions
4_Insulation score
Boundary of chromatin interactions
Weak
Strong
PAX5 chromatin interactions 10bp (filter activated)
PAX5 chromatin interactions 20bp (filter activated)
PAX5 chromatin interactions 50bp (filter activated)
PAX5 chromatin interactions 200bp (filter activated)
PAX5 binding sites
PAX5
CTCF,RAD21,SMC3 binding sites
CTCF
RAD21
SMC3
chr1:229,621,372-230,621,370 (999,999 bp)
230,000,000
230,500,000
chr1:64,795,245-65,239,688 (444,444 bp)
100 kb
64,900,000
65,000,000
65,100,000
65,200,000

### PBX2

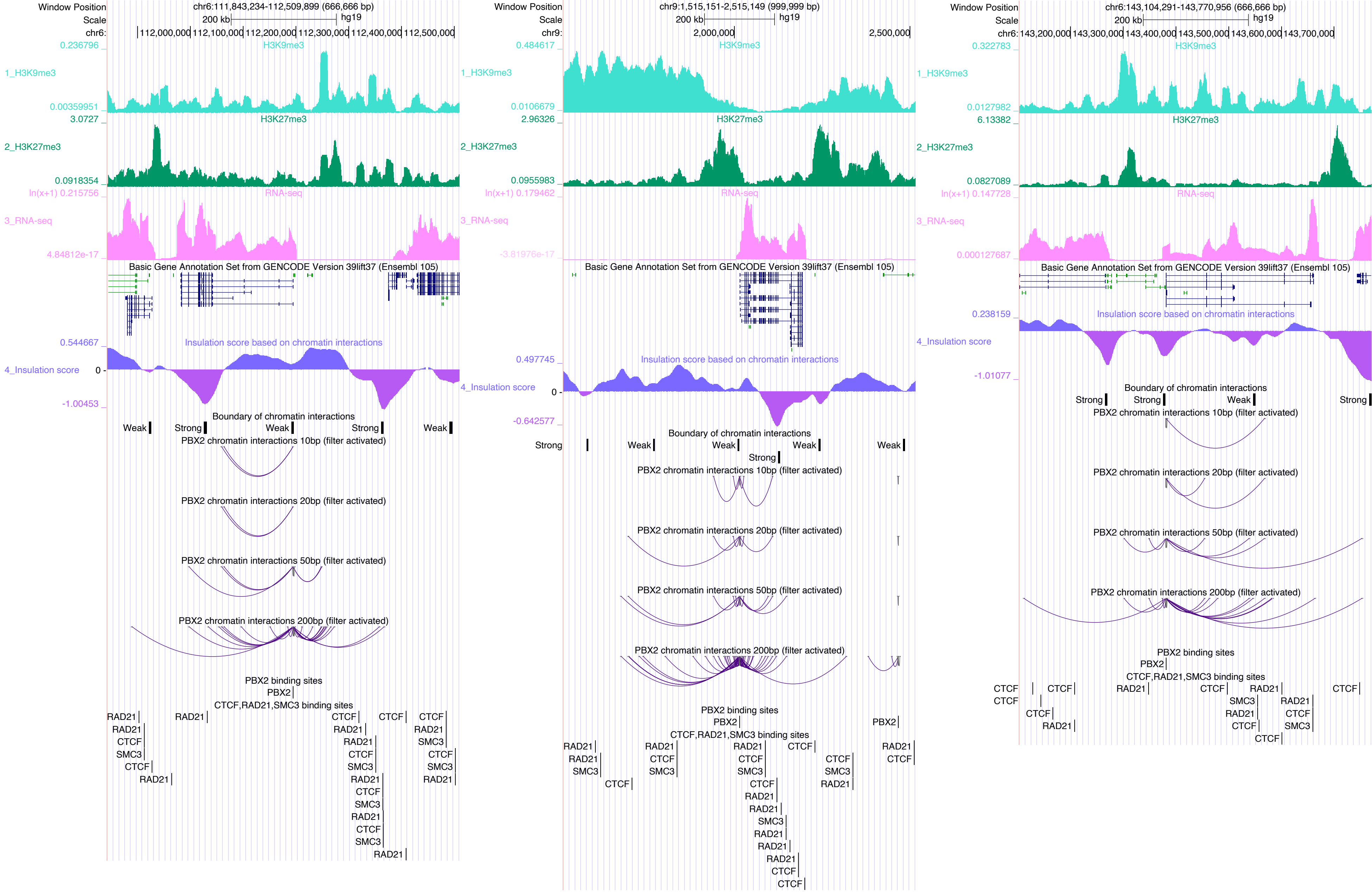

Window Position
Scale
chr6:111,843,234-112,509,899 (666,666 bp)
200 kb
hg19
chr6:
112,000,000 112,100,000 112,200,000 112,300,000 112,400,000 112,500,000
0.236796
H3K9me3
1_H3K9me3
0.00359951
3.0727
H3K27me3
2_H3K27me3
0.0918354
ln(x+1) 0.215756
RNA-seq
3_RNA-seq
4.84812e-17
Basic Gene Annotation Set from GENCODE Version 39lift37 (Ensembl 105)
0.544667
Insulation score based on chromatin interactions
4_Insulation score
0
-1.00453
Boundary of chromatin interactions
Weak
Strong
Weak
Strong
Weak
PBX2 chromatin interactions 10bp (filter activated)
PBX2 chromatin interactions 20bp (filter activated)
PBX2 chromatin interactions 50bp (filter activated)
PBX2 chromatin interactions 200bp (filter activated)
PBX2 binding sites
PBX2
CTCF,RAD21,SMC3 binding sites
RAD21
RAD21
CTCF
SMC3
CTCF
RAD21
RAD21
CTCF
RAD21
RAD21
CTCF
SMC3
RAD21
CTCF
SMC3
RAD21
CTCF
SMC3
RAD21
CTCF
RAD21
SMC3
CTCF
SMC3
RAD21
Window Position
Scale
chr9:1,515,151-2,515,149 (999,999 bp)
200 kb
hg19
chr9:
2,000,000
2,500,000
0.484617
H3K9me3
1_H3K9me3
0.0106679
2.96326
H3K27me3
2_H3K27me3
0.0955983
ln(x+1) 0.179462
RNA-seq
3_RNA-seq
-3.81976e-17
Basic Gene Annotation Set from GENCODE Version 39lift37 (Ensembl 105)
0.497745
Insulation score based on chromatin interactions
4_Insulation score
0
-0.642577
Boundary of chromatin interactions
Strong
Weak
Weak
Weak
Weak
Strong
PBX2 chromatin interactions 10bp (filter activated)
PBX2 chromatin interactions 20bp (filter activated)
PBX2 chromatin interactions 50bp (filter activated)
PBX2 chromatin interactions 200bp (filter activated)
PBX2 binding sites
PBX2
PBX2
CTCF,RAD21,SMC3 binding sites
RAD21
RAD21
SMC3
CTCF
RAD21
CTCF
SMC3
RAD21
CTCF
SMC3
CTCF
RAD21
RAD21
SMC3
RAD21
RAD21
RAD21
CTCF
CTCF
CTCF
CTCF
SMC3
RAD21
RAD21
CTCF
Window Position
Scale
chr6:143,104,291-143,770,956 (666,666 bp)
200 kb
hg19
chr6:
143,200,000 143,300,000 143,400,000 143,500,000 143,600,000 143,700,000
0.322783
H3K9me3
1_H3K9me3
0.0127982
6.13382
H3K27me3
2_H3K27me3
0.0827089
ln(x+1) 0.147728
RNA-seq
3_RNA-seq
0.000127687
Basic Gene Annotation Set from GENCODE Version 39lift37 (Ensembl 105)
0.238159
Insulation score based on chromatin interactions
4_Insulation score
-1.01077
Boundary of chromatin interactions
Strong
Strong
Weak
Strong
PBX2 chromatin interactions 10bp (filter activated)
PBX2 chromatin interactions 20bp (filter activated)
PBX2 chromatin interactions 50bp (filter activated)
PBX2 chromatin interactions 200bp (filter activated)
PBX2 binding sites
PBX2
CTCF,RAD21,SMC3 binding sites
CTCF
CTCF
CTCF
CTCF
RAD21
RAD21
CTCF
RAD21
SMC3
RAD21
RAD21
CTCF
CTCF
SMC3
CTCF
CTCF

### PRDM9

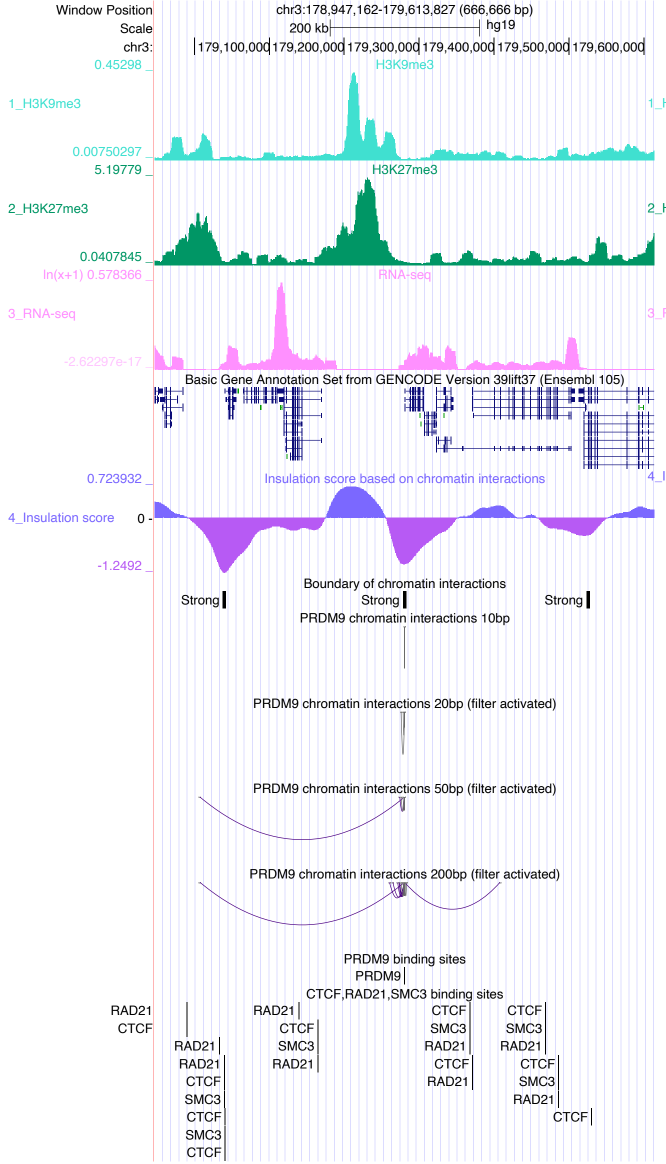

Window Position chr3:178,947,162-179,613,827 (666,666 bp)
Scale 200 kb hg19
H3K9me3
H3K27me3
RNA-seq
Basic Gene Annotation Set from GENCODE Version 39lift37 (Ensembl 105)
Insulation score based on chromatin interactions
Boundary of chromatin interactions
PRDM9 chromatin interactions 10bp
PRDM9 chromatin interactions 20bp (filter activated)
PRDM9 chromatin interactions 50bp (filter activated)
PRDM9 chromatin interactions 200bp (filter activated)
PRDM9 binding sites
CTCF,RAD21,SMC3 binding sites


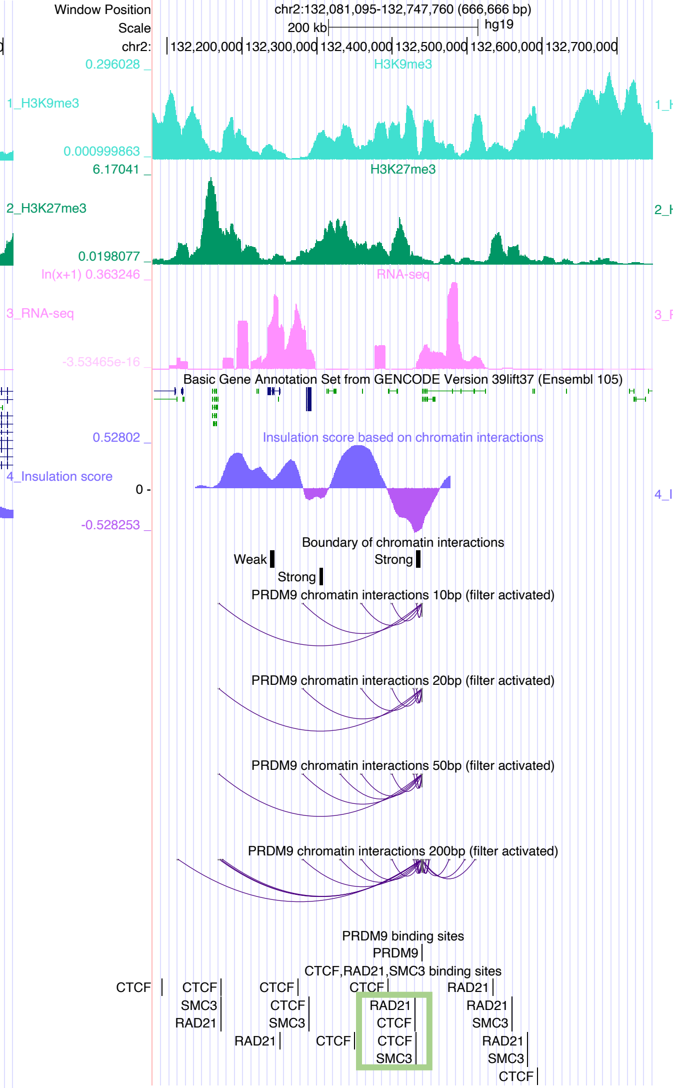

Window Position chr2:132,081,095-132,747,760 (666,666 bp)
Scale 200 kb hg19
H3K9me3
H3K27me3
RNA-seq
Basic Gene Annotation Set from GENCODE Version 39lift37 (Ensembl 105)
Insulation score based on chromatin interactions
Boundary of chromatin interactions
PRDM9 chromatin interactions 10bp (filter activated)
PRDM9 chromatin interactions 20bp (filter activated)
PRDM9 chromatin interactions 50bp (filter activated)
PRDM9 chromatin interactions 200bp (filter activated)
PRDM9 binding sites
CTCF,RAD21,SMC3 binding sites


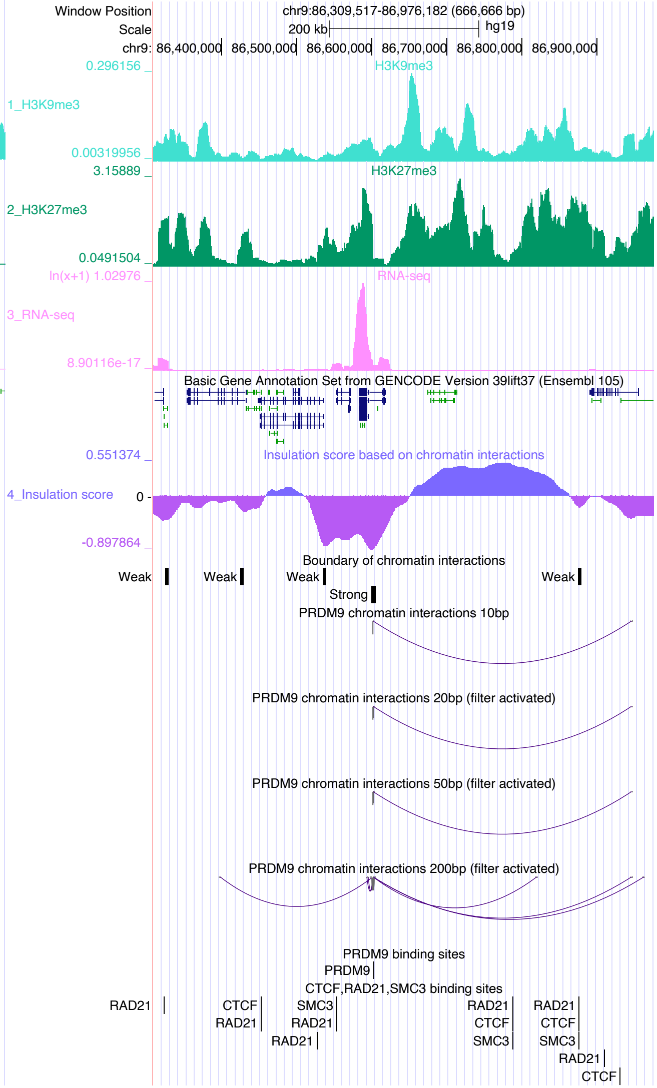

Window Position chr9:86,309,517-86,976,182 (666,666 bp)
Scale 200 kb hg19
H3K9me3
H3K27me3
RNA-seq
Basic Gene Annotation Set from GENCODE Version 39lift37 (Ensembl 105)
Insulation score based on chromatin interactions
Boundary of chromatin interactions
PRDM9 chromatin interactions 10bp
PRDM9 chromatin interactions 20bp (filter activated)
PRDM9 chromatin interactions 50bp (filter activated)
PRDM9 chromatin interactions 200bp (filter activated)
PRDM9 binding sites
CTCF,RAD21,SMC3 binding sites

RBPJ

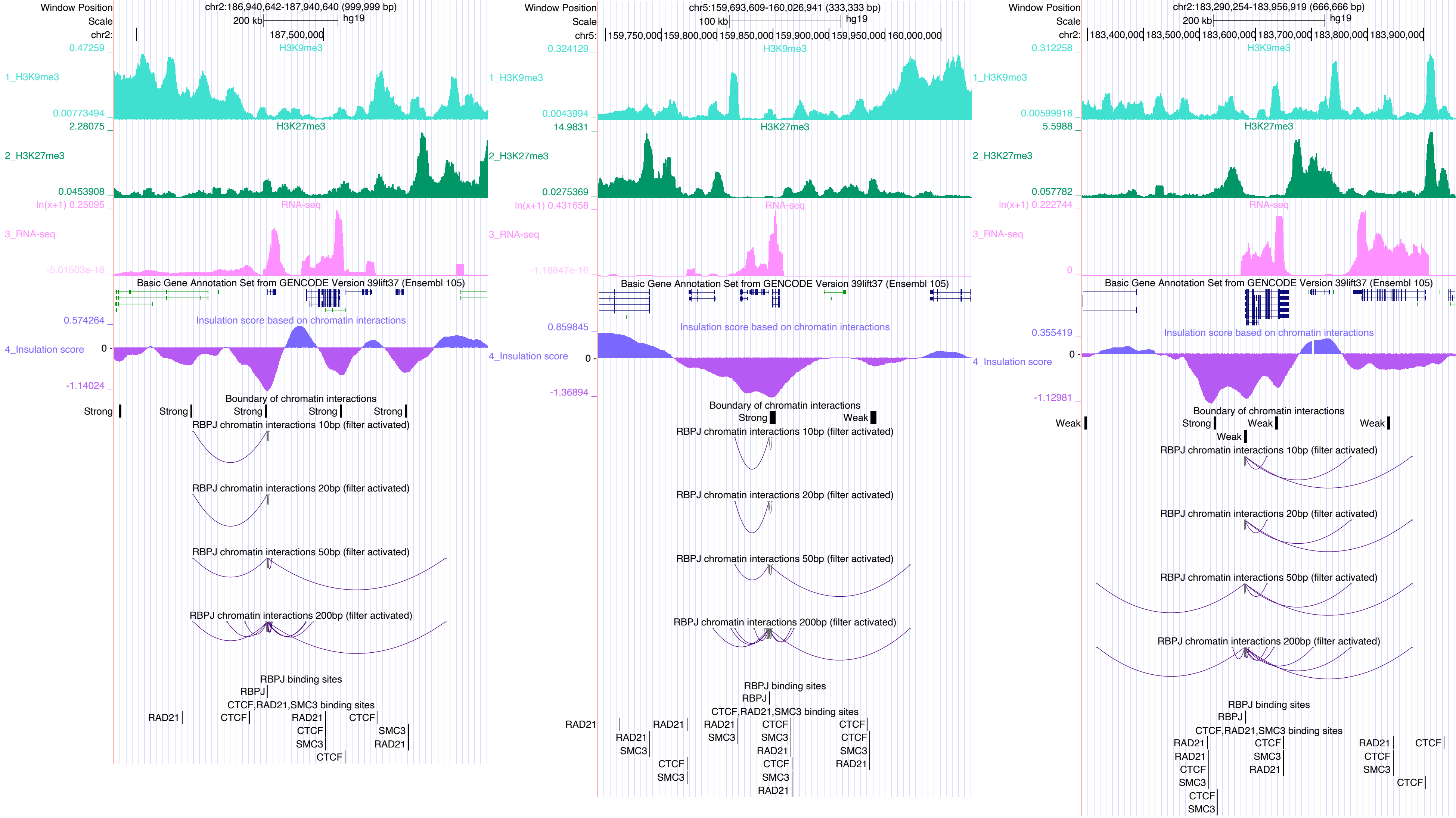

Window Position
chr2:186,940,642-187,940,640 (999,999 bp)
Scale
200 kb
hg19
chr2:
187,500,000
H3K9me3
1_H3K9me3
2_H3K27me3
3_RNA-seq
Basic Gene Annotation Set from GENCODE Version 39lift37 (Ensembl 105)
Insulation score based on chromatin interactions
4_Insulation score
Boundary of chromatin interactions
Strong
RBPJ chromatin interactions 10bp (filter activated)
RBPJ chromatin interactions 20bp (filter activated)
RBPJ chromatin interactions 50bp (filter activated)
RBPJ chromatin interactions 200bp (filter activated)
RBPJ binding sites
RBPJ
CTCF,RAD21,SMC3 binding sites
RAD21
CTCF
SMC3
chr5:159,693,609-160,026,941 (333,333 bp)
100 kb
chr5:
159,750,000
159,800,000
159,850,000
159,900,000
159,950,000
160,000,000
Weak
chr2:183,290,254-183,956,919 (666,666 bp)
183,400,000
183,500,000
183,600,000
183,700,000
183,800,000
183,900,000

### REST

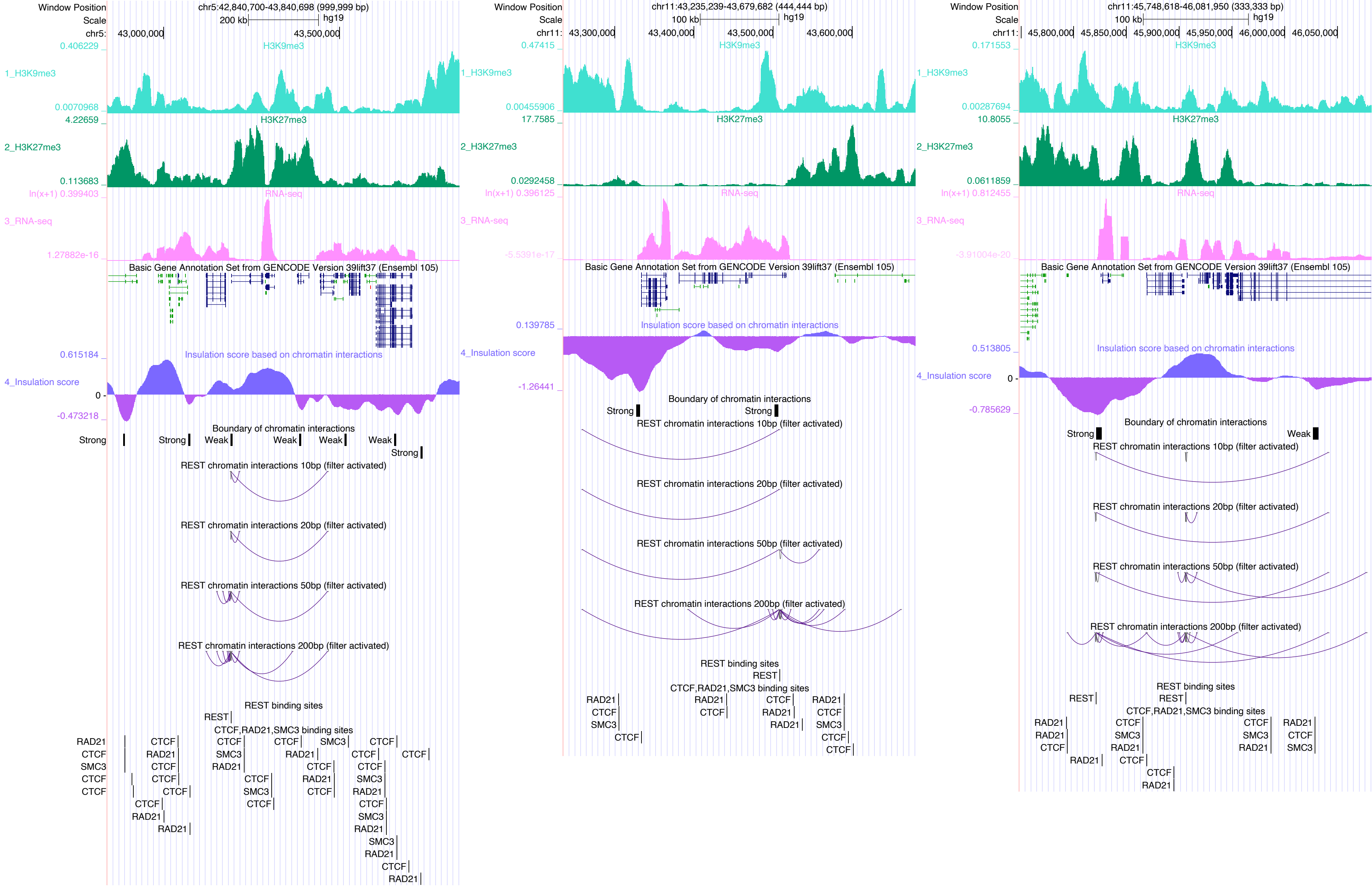

Window Position
Scale
chr5:42,840,700-43,840,698 (999,999 bp)
200 kb
hg19
chr5: 43,000,000 43,500,000
H3K9me3
1_H3K9me3
H3K27me3
2_H3K27me3
RNA-seq
3_RNA-seq
Basic Gene Annotation Set from GENCODE Version 39lift37 (Ensembl 105)
Insulation score based on chromatin interactions
4_Insulation score
Boundary of chromatin interactions
Strong
Weak
REST chromatin interactions 10bp (filter activated)
REST chromatin interactions 20bp (filter activated)
REST chromatin interactions 50bp (filter activated)
REST chromatin interactions 200bp (filter activated)
REST binding sites
REST
CTCF,RAD21,SMC3 binding sites
RAD21
CTCF
SMC3
chr11:43,235,239-43,679,682 (444,444 bp)
100 kb
chr11: 43,300,000 43,400,000 43,500,000 43,600,000
chr11:45,748,618-46,081,950 (333,333 bp)
chr11: 45,800,000 45,850,000 45,900,000 45,950,000 46,000,000 46,050,000

### RUNX1

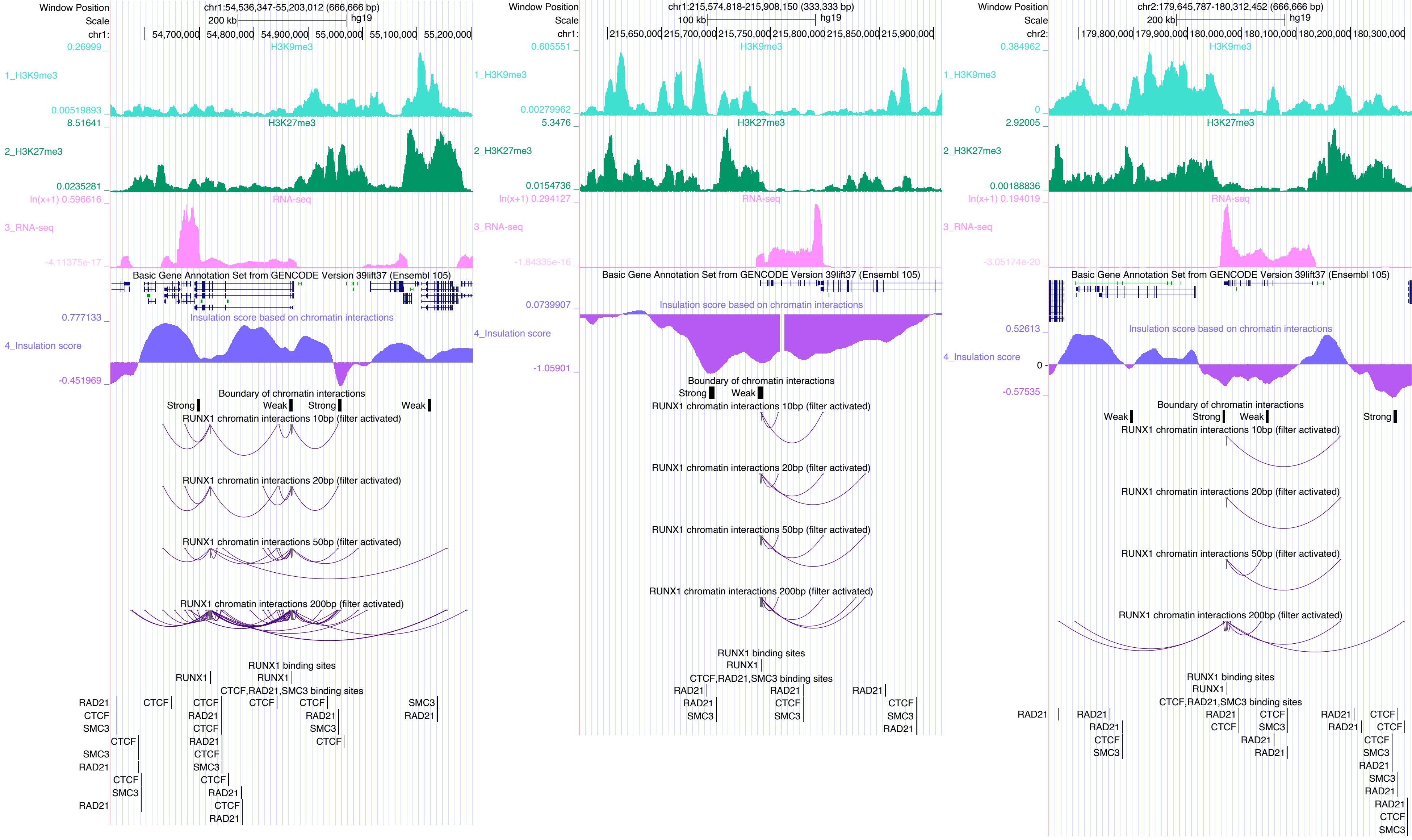

Window Position
Scale
chr1:54,536,347-55,203,012 (666,666 bp)
200 kb
hg19
chr1:
54,700,000
54,800,000
54,900,000
55,000,000
55,100,000
55,200,000
1_H3K9me3
H3K9me3
0.26999
0.00519893
2_H3K27me3
H3K27me3
8.51641
0.0235281
3_RNA-seq
RNA-seq
ln(x+1) 0.596616
-4.11375e-17
Basic Gene Annotation Set from GENCODE Version 39lift37 (Ensembl 105)
4_Insulation score
Insulation score based on chromatin interactions
0.777133
-0.451969
Boundary of chromatin interactions
Strong
Weak
RUNX1 chromatin interactions 10bp (filter activated)
RUNX1 chromatin interactions 20bp (filter activated)
RUNX1 chromatin interactions 50bp (filter activated)
RUNX1 chromatin interactions 200bp (filter activated)
RUNX1 binding sites
RUNX1
CTCF,RAD21,SMC3 binding sites
RAD21
CTCF
SMC3
chr1:215,574,818-215,908,150 (333,333 bp)
100 kb
215,650,000
215,700,000
215,750,000
215,800,000
215,850,000
215,900,000
0.605551
0.00279962
5.3476
0.0154736
ln(x+1) 0.294127
-1.84335e-16
0.0739907
-1.05901
chr2:179,645,787-180,312,452 (666,666 bp)
chr2:
179,800,000
179,900,000
180,000,000
180,100,000
180,200,000
180,300,000
0.384962
0
2.92005
0.00188836
ln(x+1) 0.194019
-3.05174e-20
0.52613
-0.57535

### RUNX3

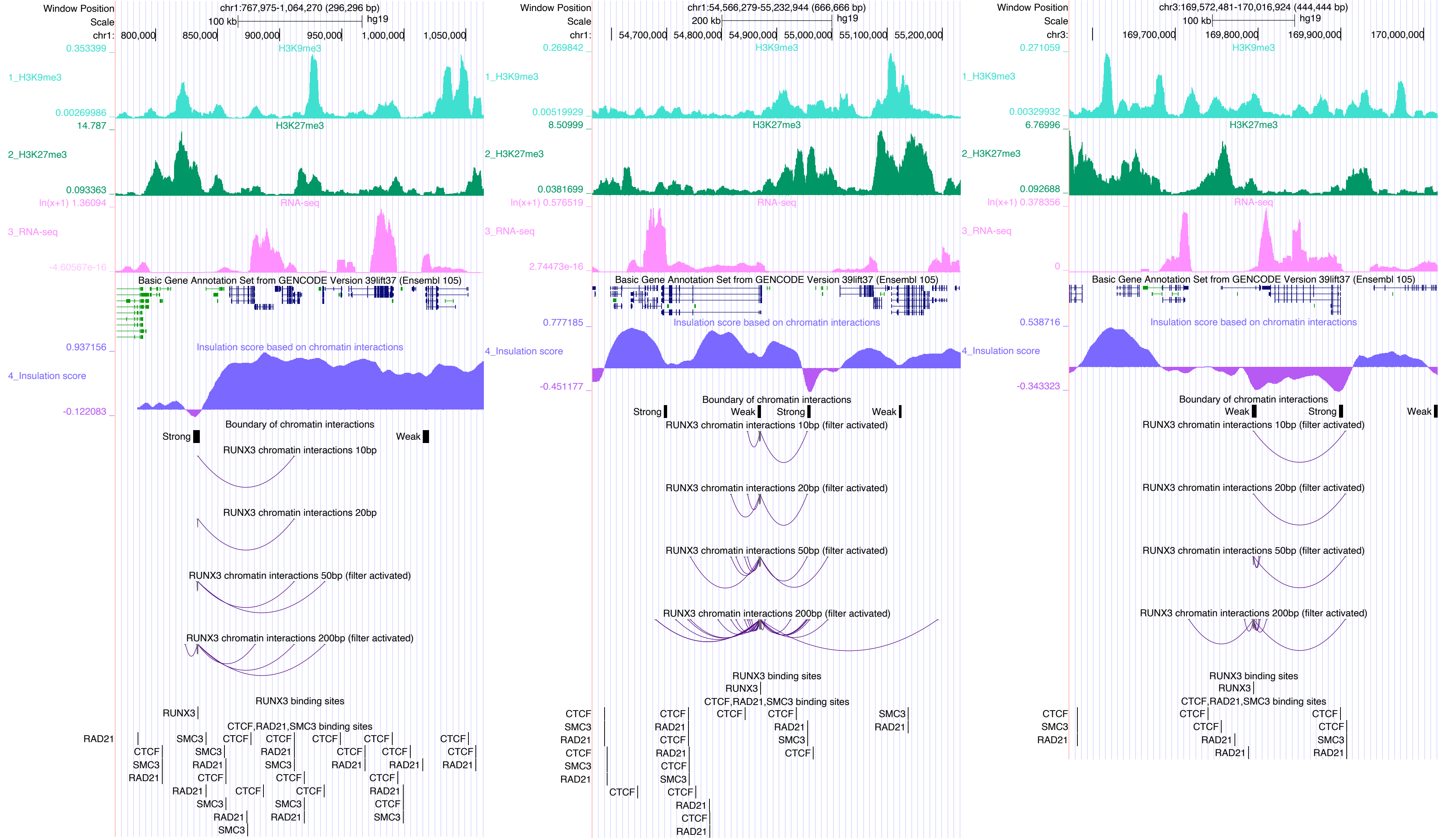

Window Position
chr1:767,975-1,064,270 (296,296 bp)
chr1:54,566,279-55,232,944 (666,666 bp)
chr3:169,572,481-170,016,924 (444,444 bp)
1_H3K9me3
2_H3K27me3
3_RNA-seq
4_Insulation score
Basic Gene Annotation Set from GENCODE Version 39lift37 (Ensembl 105)
Insulation score based on chromatin interactions
Boundary of chromatin interactions
Strong
Weak
RUNX3 chromatin interactions 10bp
RUNX3 chromatin interactions 20bp
RUNX3 chromatin interactions 50bp (filter activated)
RUNX3 chromatin interactions 200bp (filter activated)
RUNX3 binding sites
CTCF,RAD21,SMC3 binding sites

### RXRA

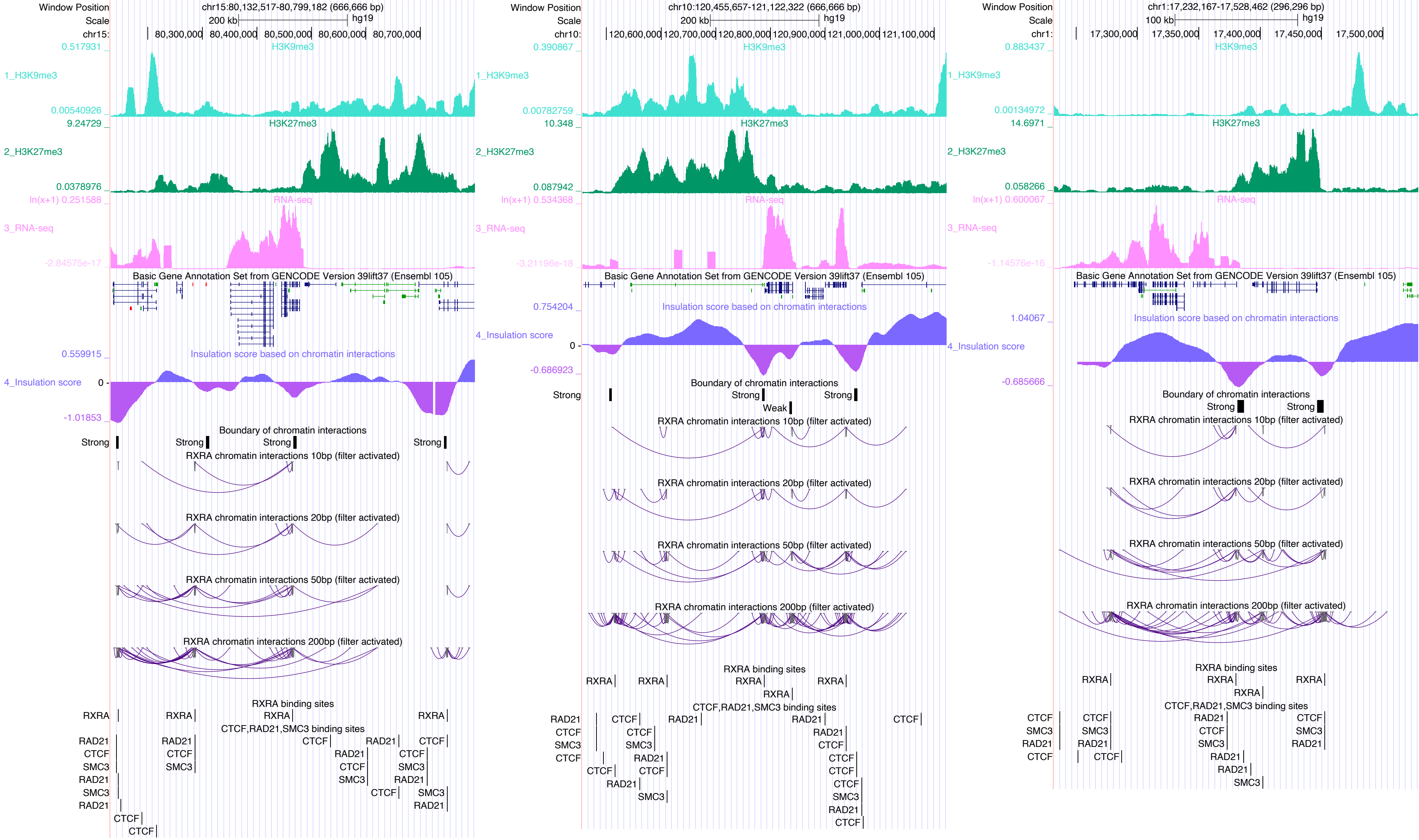

Window Position
Scale
chr15:80,132,517-80,799,182 (666,666 bp)
200 kb
hg19
chr15:
80,300,000
80,400,000
80,500,000
80,600,000
80,700,000
1_H3K9me3
H3K9me3
0.517931
0.00540926
2_H3K27me3
H3K27me3
9.24729
0.0378976
3_RNA-seq
RNA-seq
ln(x+1) 0.251588
-2.84575e-17
Basic Gene Annotation Set from GENCODE Version 39lift37 (Ensembl 105)
4_Insulation score
Insulation score based on chromatin interactions
0.559915
0
-1.01853
Boundary of chromatin interactions
Strong
RXRA chromatin interactions 10bp (filter activated)
RXRA chromatin interactions 20bp (filter activated)
RXRA chromatin interactions 50bp (filter activated)
RXRA chromatin interactions 200bp (filter activated)
RXRA binding sites
RXRA
CTCF,RAD21,SMC3 binding sites
RAD21
CTCF
SMC3
Window Position
Scale
chr10:120,455,657-121,122,322 (666,666 bp)
200 kb
hg19
chr10:
120,600,000
120,700,000
120,800,000
120,900,000
121,000,000
121,100,000
0.390867
0.00782759
10.348
0.087942
ln(x+1) 0.534368
-3.21196e-18
0.754204
-0.686923
Weak
Window Position
Scale
chr1:17,232,167-17,528,462 (296,296 bp)
100 kb
hg19
chr1:
17,300,000
17,350,000
17,400,000
17,450,000
17,500,000
0.883437
0.00134972
14.6971
0.058266
ln(x+1) 0.600067
-1.14576e-16
1.04067
-0.685666

### SCRT1

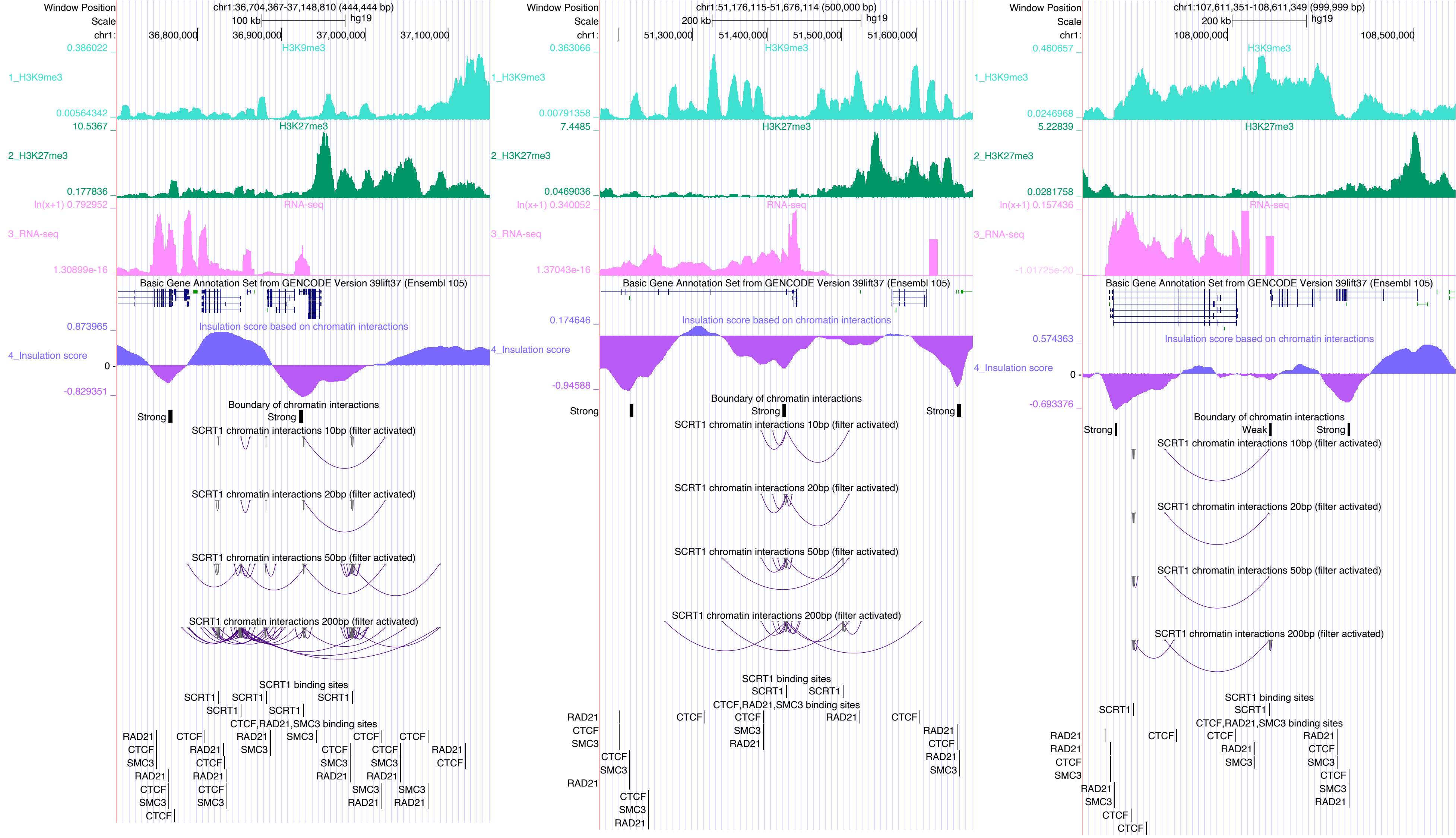
Window Position
chr1:36,704,367-37,148,810 (444,444 bp)
Scale
100 kb
hg19
chr1:
36,800,000
36,900,000
37,000,000
37,100,000
0.386022
H3K9me3
1_H3K9me3
0.00564342
10.5367
H3K27me3
2_H3K27me3
0.177836
ln(x+1) 0.792952
RNA-seq
3_RNA-seq
1.30899e-16
Basic Gene Annotation Set from GENCODE Version 39lift37 (Ensembl 105)
0.873965
Insulation score based on chromatin interactions
4_Insulation score
0
-0.829351
Boundary of chromatin interactions
Strong
Strong
SCRT1 chromatin interactions 10bp (filter activated)
SCRT1 chromatin interactions 20bp (filter activated)
SCRT1 chromatin interactions 50bp (filter activated)
SCRT1 chromatin interactions 200bp (filter activated)
SCRT1 binding sites
CTCF,RAD21,SMC3 binding sites
Window Position
chr1:51,176,115-51,676,114 (500,000 bp)
200 kb
51,300,000
51,400,000
51,500,000
51,600,000
0.363066
0.00791358
7.4485
0.0469036
ln(x+1) 0.340052
1.37043e-16
0.174646
-0.94588
Window Position
chr1:107,611,351-108,611,349 (999,999 bp)
200 kb
108,000,000
108,500,000
0.460657
0.0246968
5.22839
0.0281758
ln(x+1) 0.157436
-1.01725e-20
0.574363
-0.693376
Weak

### SIN3A

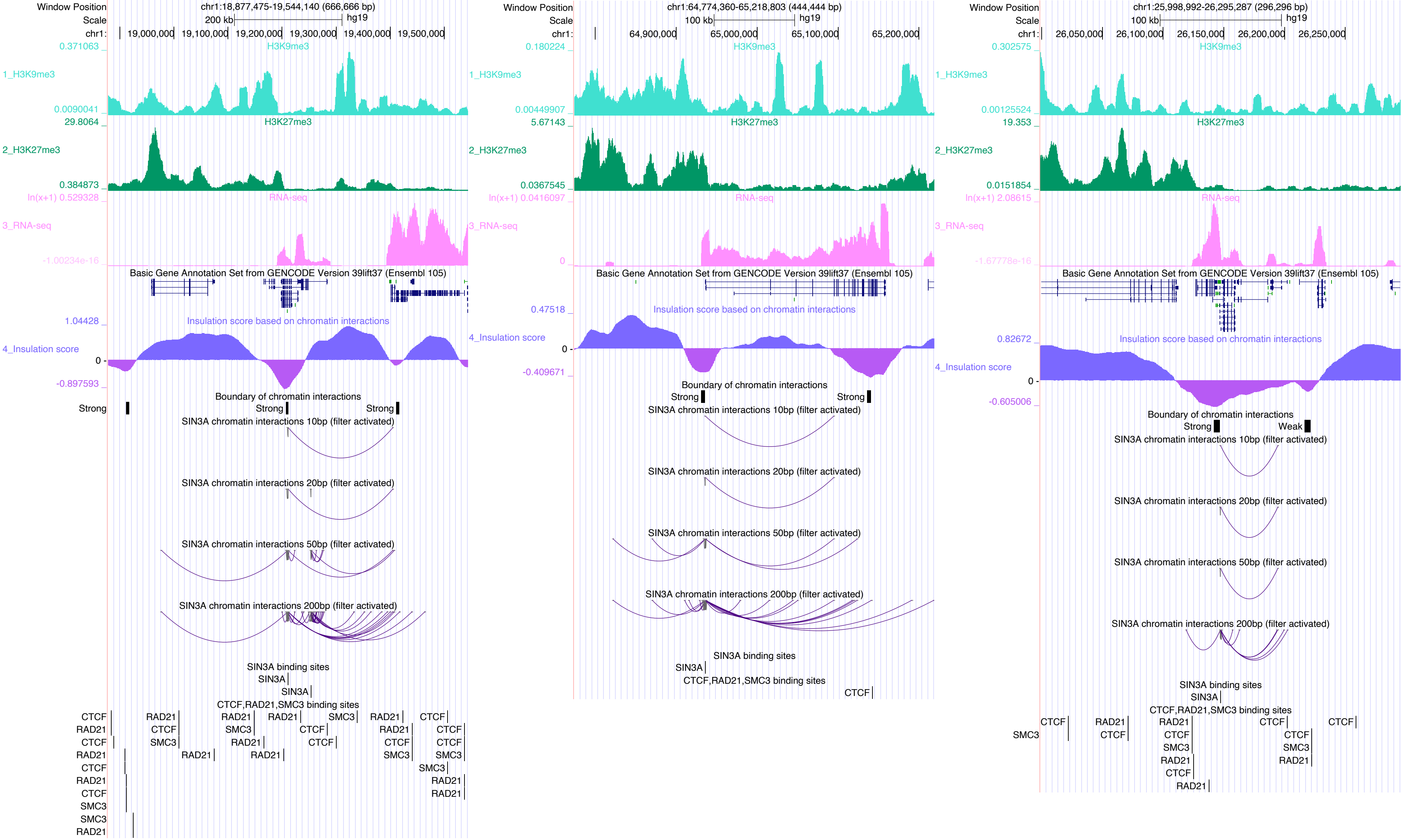
Window Position chr1:18,877,475-19,544,140 (666,666 bp)
Scale 200 kb hg19
chr1: 19,000,000 19,100,000 19,200,000 19,300,000 19,400,000 19,500,000
1_H3K9me3
2_H3K27me3
3_RNA-seq
Basic Gene Annotation Set from GENCODE Version 39lift37 (Ensembl 105)
4_Insulation score
Insulation score based on chromatin interactions
Boundary of chromatin interactions
Strong
SIN3A chromatin interactions 10bp (filter activated)
SIN3A chromatin interactions 20bp (filter activated)
SIN3A chromatin interactions 50bp (filter activated)
SIN3A chromatin interactions 200bp (filter activated)
SIN3A binding sites
CTCF,RAD21,SMC3 binding sites
Window Position chr1:64,774,360-65,218,803 (444,444 bp)
Scale 100 kb hg19
chr1: 64,900,000 65,000,000 65,100,000 65,200,000
Window Position chr1:25,998,992-26,295,287 (296,296 bp)
Scale 100 kb hg19
chr1: 26,050,000 26,100,000 26,150,000 26,200,000 26,250,000
Weak

### SMAD1

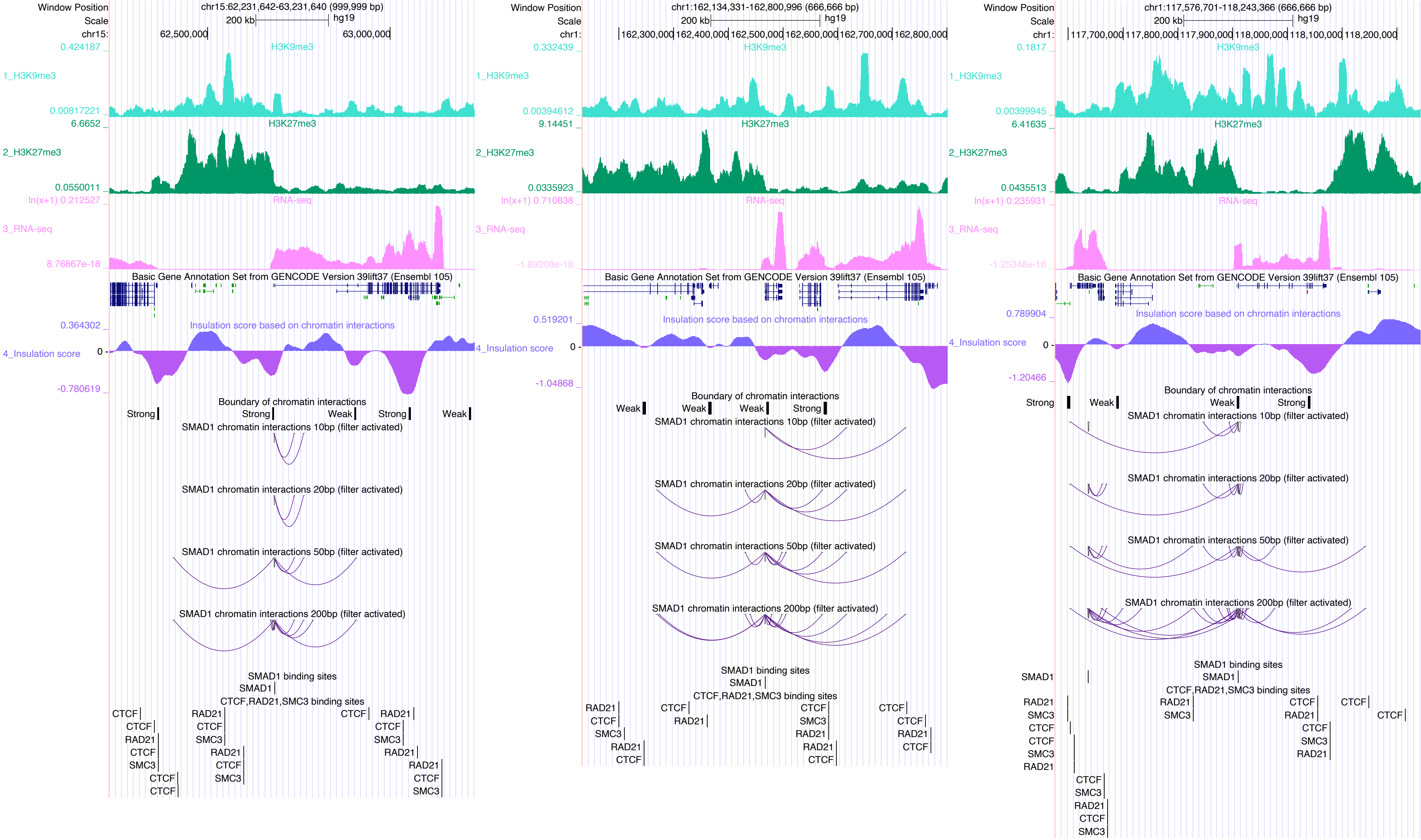

Window Position
chr15:62,231,642-63,231,640 (999,999 bp)
Scale
200 kb
hg19
chr15:
62,500,000
63,000,000
1_H3K9me3
H3K9me3
0.424187
0.00817221
2_H3K27me3
H3K27me3
6.6652
0.0550011
3_RNA-seq
RNA-seq
ln(x+1) 0.212527
8.76867e-18
Basic Gene Annotation Set from GENCODE Version 39lift37 (Ensembl 105)
4_Insulation score
Insulation score based on chromatin interactions
0.364302
0
-0.780619
Boundary of chromatin interactions
Strong
Strong
Weak
Strong
Weak
SMAD1 chromatin interactions 10bp (filter activated)
SMAD1 chromatin interactions 20bp (filter activated)
SMAD1 chromatin interactions 50bp (filter activated)
SMAD1 chromatin interactions 200bp (filter activated)
SMAD1 binding sites
SMAD1
CTCF,RAD21,SMC3 binding sites
Window Position
chr1:162,134,331-162,800,996 (666,666 bp)
chr1:
162,300,000
162,400,000
162,500,000
162,600,000
162,700,000
162,800,000
0.332439
0.00394612
9.14451
0.0335923
ln(x+1) 0.710838
-1.89208e-18
0.519201
-1.04868
Window Position
chr1:117,576,701-118,243,366 (666,666 bp)
117,700,000
117,800,000
117,900,000
118,000,000
118,100,000
118,200,000
0.1817
0.00399945
6.41635
0.0435513
ln(x+1) 0.235931
-1.25348e-16
0.789904
-1.20466

SMAD2

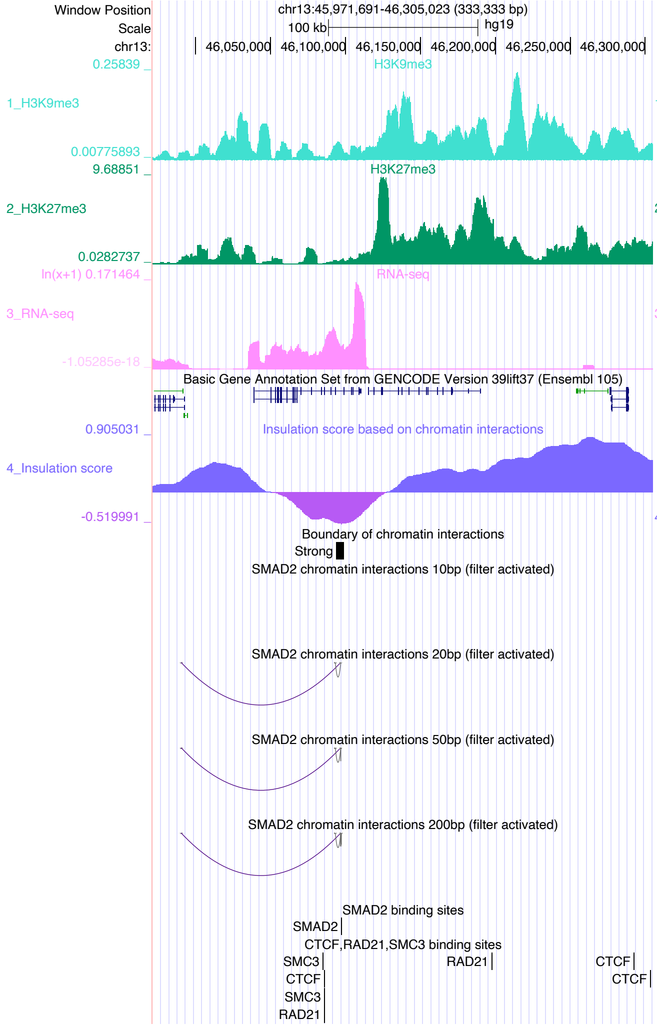

Window Position chr13:45,971,691-46,305,023 (333,333 bp)
Scale 100 kb hg19
chr13: 46,050,000 46,100,000 46,150,000 46,200,000 46,250,000 46,300,000
1_H3K9me3
2_H3K27me3
3_RNA-seq
Basic Gene Annotation Set from GENCODE Version 39lift37 (Ensembl 105)
4_Insulation score
Insulation score based on chromatin interactions
Boundary of chromatin interactions
Strong
SMAD2 chromatin interactions 10bp (filter activated)
SMAD2 chromatin interactions 20bp (filter activated)
SMAD2 chromatin interactions 50bp (filter activated)
SMAD2 chromatin interactions 200bp (filter activated)
SMAD2 binding sites
SMAD2
CTCF,RAD21,SMC3 binding sites
SMC3
CTCF
SMC3
RAD21
RAD21
CTCF
CTCF


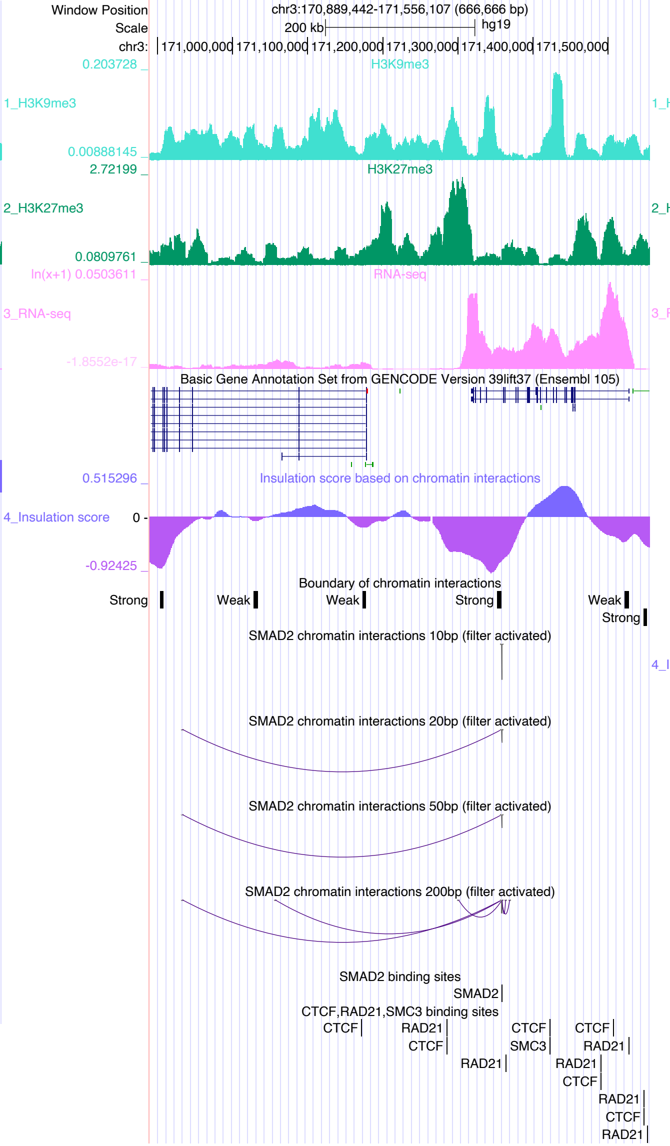

Window Position chr3:170,889,442-171,556,107 (666,666 bp)
Scale 200 kb hg19
chr3: 171,000,000 171,100,000 171,200,000 171,300,000 171,400,000 171,500,000
1_H3K9me3
2_H3K27me3
3_RNA-seq
Basic Gene Annotation Set from GENCODE Version 39lift37 (Ensembl 105)
Insulation score based on chromatin interactions
4_Insulation score
Boundary of chromatin interactions
Strong
Weak
Weak
Strong
Weak
Strong
SMAD2 chromatin interactions 10bp (filter activated)
SMAD2 chromatin interactions 20bp (filter activated)
SMAD2 chromatin interactions 50bp (filter activated)
SMAD2 chromatin interactions 200bp (filter activated)
SMAD2 binding sites
SMAD2
CTCF,RAD21,SMC3 binding sites
CTCF
RAD21
CTCF
RAD21
CTCF
SMC3
CTCF
RAD21
RAD21
CTCF
RAD21
CTCF
RAD21


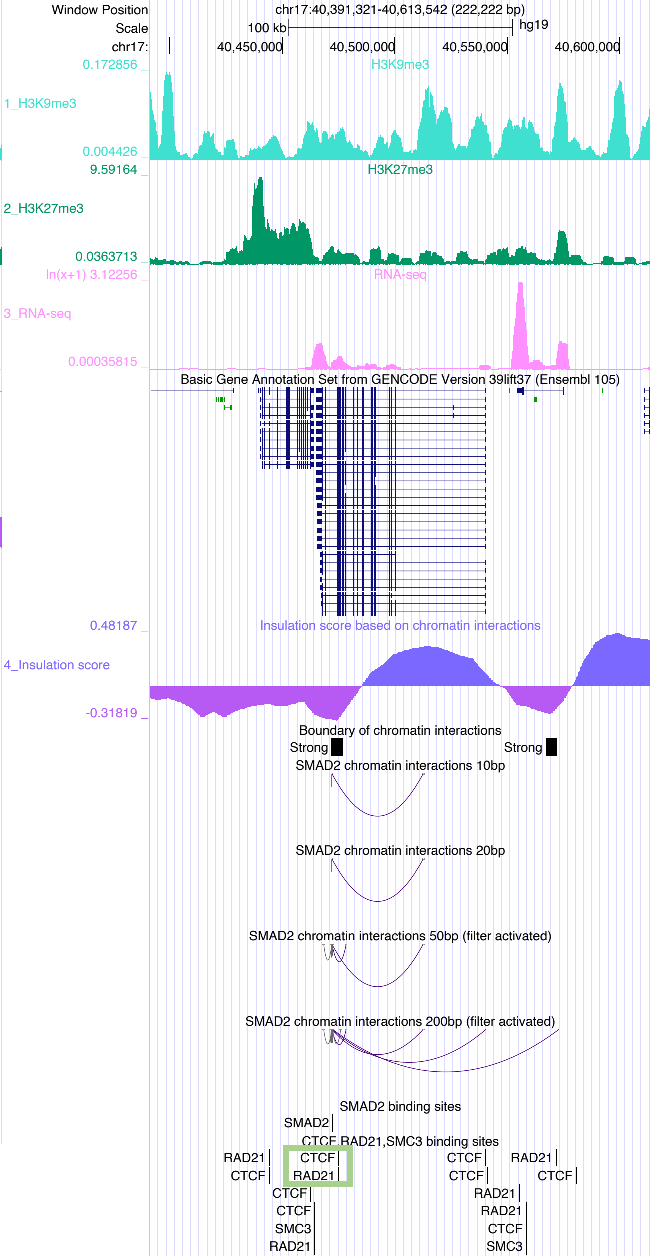

Window Position chr17:40,391,321-40,613,542 (222,222 bp)
Scale 100 kb hg19
chr17: 40,450,000 40,500,000 40,550,000 40,600,000
1_H3K9me3
2_H3K27me3
3_RNA-seq
Basic Gene Annotation Set from GENCODE Version 39lift37 (Ensembl 105)
Insulation score based on chromatin interactions
4_Insulation score
Boundary of chromatin interactions
Strong
Strong
SMAD2 chromatin interactions 10bp
SMAD2 chromatin interactions 20bp
SMAD2 chromatin interactions 50bp (filter activated)
SMAD2 chromatin interactions 200bp (filter activated)
SMAD2 binding sites
SMAD2
CTCF,RAD21,SMC3 binding sites
RAD21
CTCF
CTCF
RAD21
CTCF
CTCF
SMC3
RAD21
CTCF
CTCF
RAD21
RAD21
RAD21
CTCF
SMC3
CTCF

### SMAD3

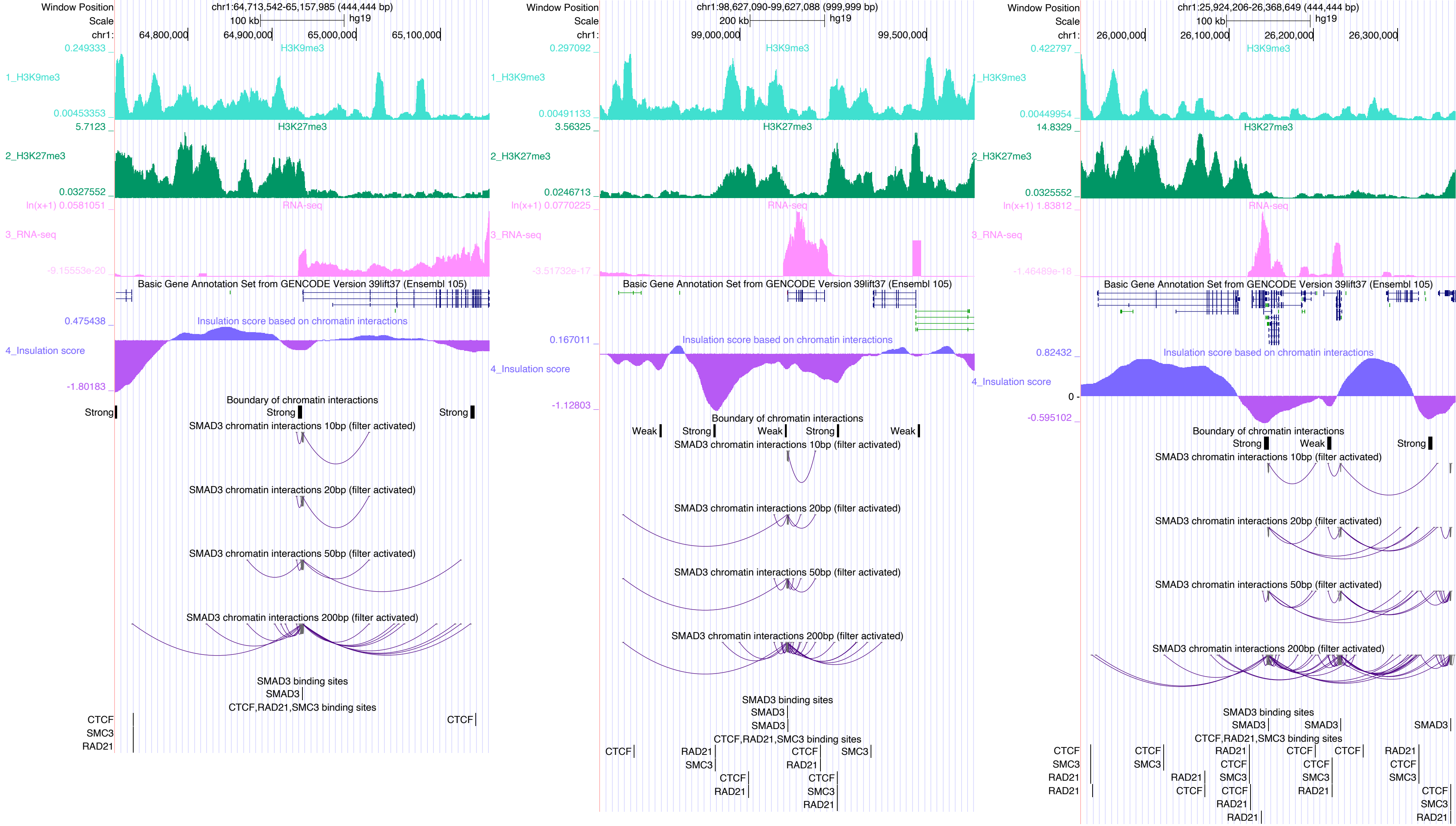

Window Position
Scale
chr1:
chr1:64,713,542-65,157,985 (444,444 bp)
100 kb
hg19
64,800,000
64,900,000
65,000,000
65,100,000
H3K9me3
1_H3K9me3
0.249333
0.00453353
H3K27me3
2_H3K27me3
5.7123
0.0327552
RNA-seq
3_RNA-seq
ln(x+1) 0.0581051
-9.15553e-20
Basic Gene Annotation Set from GENCODE Version 39lift37 (Ensembl 105)
Insulation score based on chromatin interactions
4_Insulation score
0.475438
-1.80183
Boundary of chromatin interactions
Strong
Strong
Strong
SMAD3 chromatin interactions 10bp (filter activated)
SMAD3 chromatin interactions 20bp (filter activated)
SMAD3 chromatin interactions 50bp (filter activated)
SMAD3 chromatin interactions 200bp (filter activated)
SMAD3 binding sites
SMAD3
CTCF,RAD21,SMC3 binding sites
CTCF
SMC3
RAD21
CTCF
chr1:98,627,090-99,627,088 (999,999 bp)
200 kb
hg19
99,000,000
99,500,000
0.297092
0.00491133
3.56325
0.0246713
ln(x+1) 0.0770225
-3.51732e-17
0.167011
-1.12803
Weak
Strong
Weak
Strong
Weak
chr1:25,924,206-26,368,649 (444,444 bp)
26,000,000
26,100,000
26,200,000
26,300,000
0.422797
0.00449954
14.8329
0.0325552
ln(x+1) 1.83812
-1.46489e-18
0.82432
0
-0.595102
Strong
Weak
Strong

# SP1

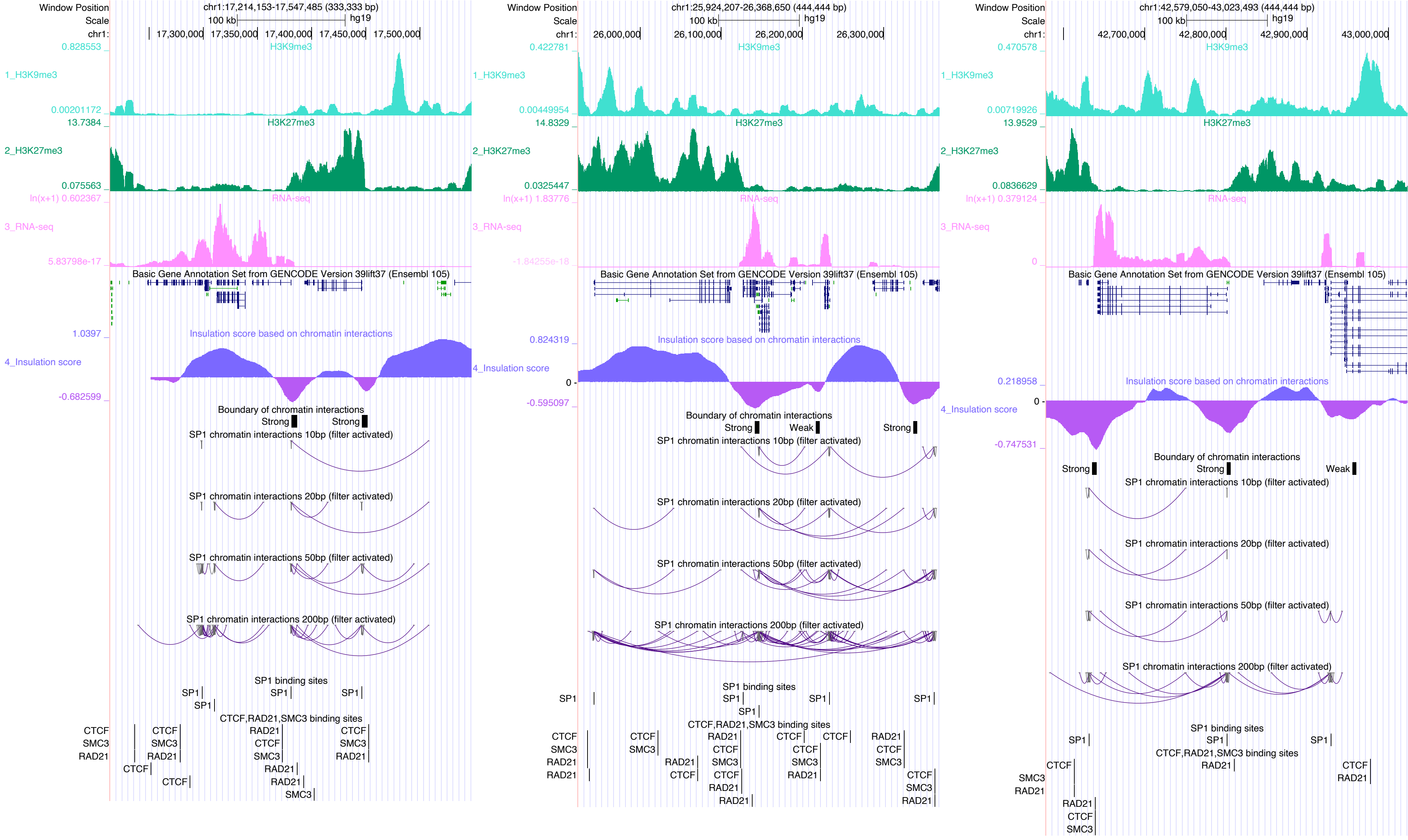

Window Position
Scale
chr1:
chr1:17,214,153-17,547,485 (333,333 bp)
100 kb
hg19
17,300,000
17,350,000
17,400,000
17,450,000
17,500,000
1_H3K9me3
2_H3K27me3
3_RNA-seq
4_Insulation score
Basic Gene Annotation Set from GENCODE Version 39lift37 (Ensembl 105)
Insulation score based on chromatin interactions
Boundary of chromatin interactions
Strong
Weak
SP1 chromatin interactions 10bp (filter activated)
SP1 chromatin interactions 20bp (filter activated)
SP1 chromatin interactions 50bp (filter activated)
SP1 chromatin interactions 200bp (filter activated)
SP1 binding sites
CTCF,RAD21,SMC3 binding sites
chr1:25,924,207-26,368,650 (444,444 bp)
26,000,000
26,100,000
26,200,000
26,300,000
chr1:42,579,050-43,023,493 (444,444 bp)
42,700,000
42,800,000
42,900,000
43,000,000

SP2

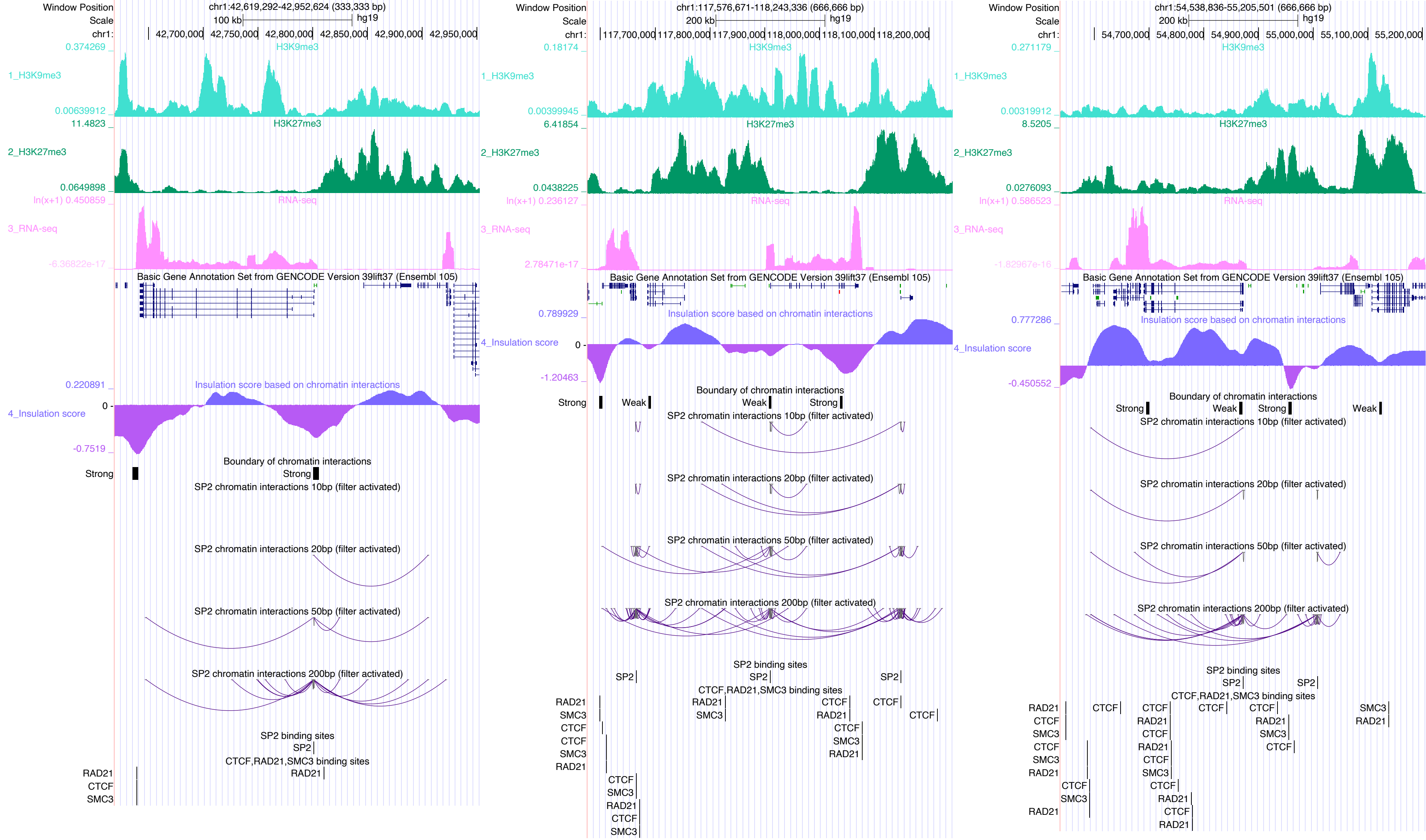
Window Position
Scale
chr1:
chr1:42,619,292-42,952,624 (333,333 bp)
100 kb
hg19
42,700,000 42,750,000 42,800,000 42,850,000 42,900,000 42,950,000
1_H3K9me3
H3K9me3
2_H3K27me3
H3K27me3
3_RNA-seq
RNA-seq
Basic Gene Annotation Set from GENCODE Version 39lift37 (Ensembl 105)
4_Insulation score
Insulation score based on chromatin interactions
Boundary of chromatin interactions
Strong
Weak
SP2 chromatin interactions 10bp (filter activated)
SP2 chromatin interactions 20bp (filter activated)
SP2 chromatin interactions 50bp (filter activated)
SP2 chromatin interactions 200bp (filter activated)
SP2 binding sites
CTCF,RAD21,SMC3 binding sites
chr1:117,576,671-118,243,336 (666,666 bp)
200 kb
117,700,000 117,800,000 117,900,000 118,000,000 118,100,000 118,200,000
chr1:54,538,836-55,205,501 (666,666 bp)
54,700,000 54,800,000 54,900,000 55,000,000 55,100,000 55,200,000

# SP5

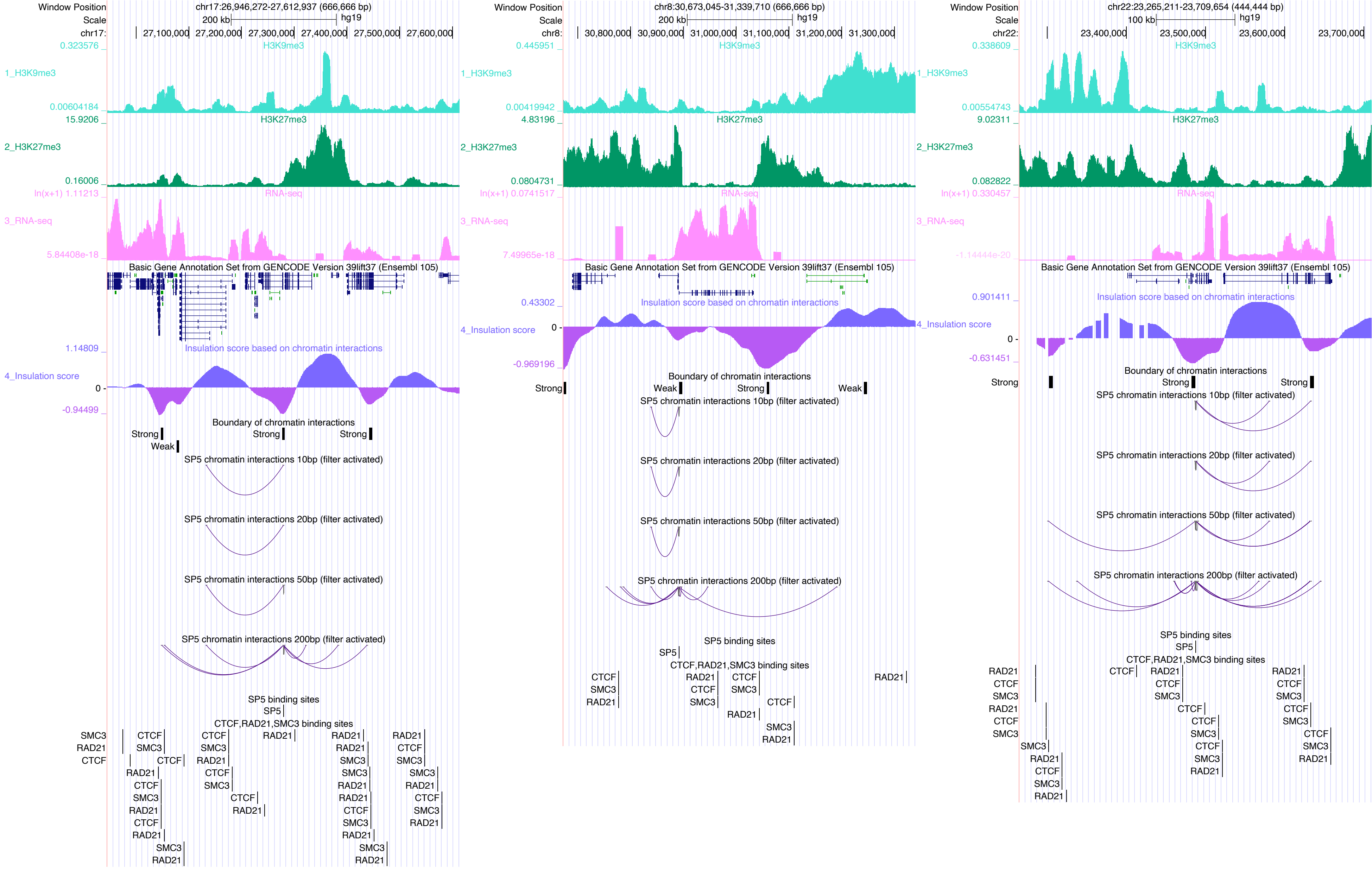

Window Position
chr17:26,946,272-27,612,937 (666,666 bp)
Scale
200 kb
hg19
chr17:
27,100,000
27,200,000
27,300,000
27,400,000
27,500,000
27,600,000
H3K9me3
1_H3K9me3
H3K27me3
2_H3K27me3
RNA-seq
3_RNA-seq
Basic Gene Annotation Set from GENCODE Version 39lift37 (Ensembl 105)
Insulation score based on chromatin interactions
4_Insulation score
Boundary of chromatin interactions
Strong
Weak
SP5 chromatin interactions 10bp (filter activated)
SP5 chromatin interactions 20bp (filter activated)
SP5 chromatin interactions 50bp (filter activated)
SP5 chromatin interactions 200bp (filter activated)
SP5 binding sites
SP5
CTCF,RAD21,SMC3 binding sites
CTCF
RAD21
SMC3
chr8:30,673,045-31,339,710 (666,666 bp)
chr8:
30,800,000
30,900,000
31,000,000
31,100,000
31,200,000
31,300,000
chr22:23,265,211-23,709,654 (444,444 bp)
100 kb
chr22:
23,400,000
23,500,000
23,600,000
23,700,000

SPI1

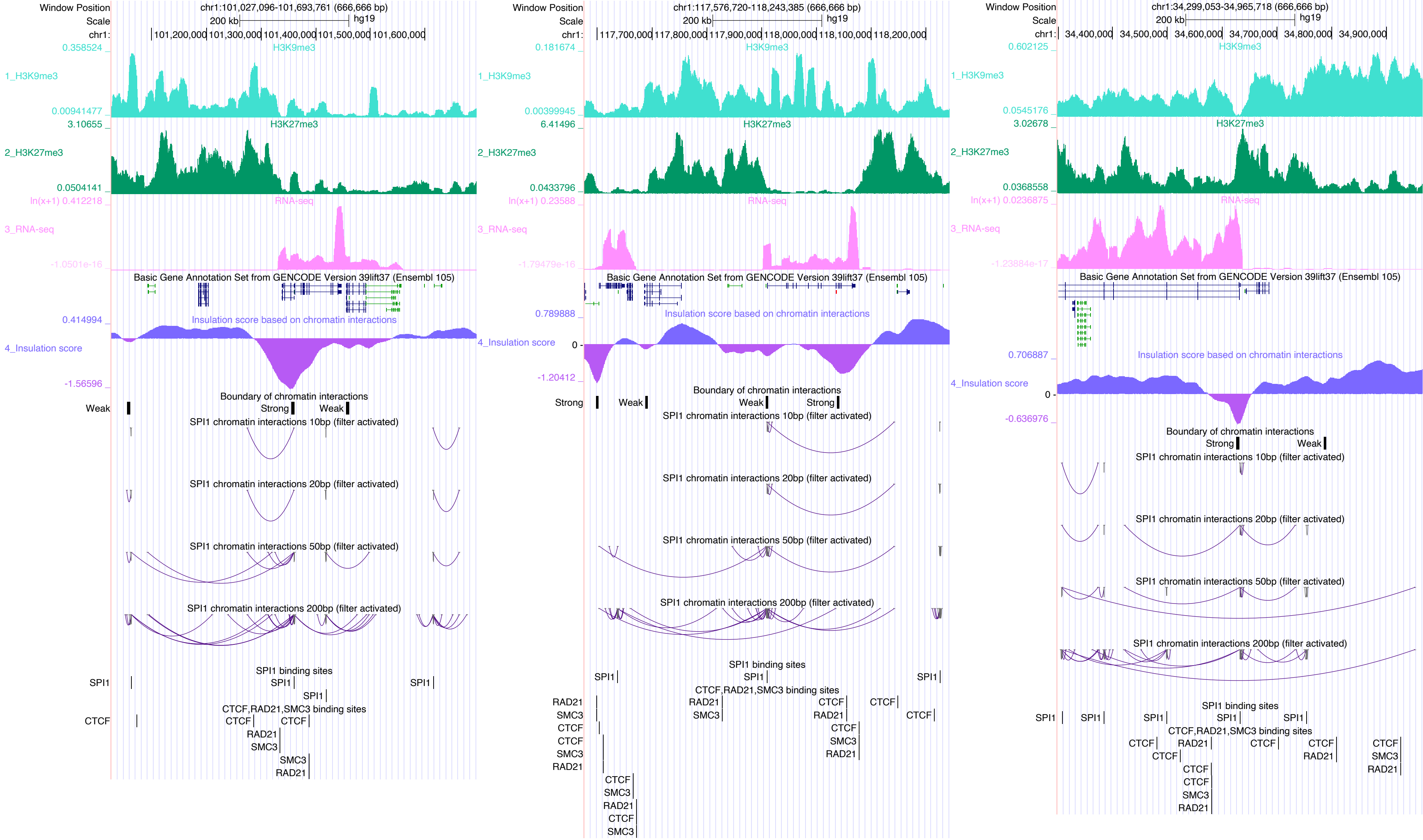
Window Position
chr1:101,027,096-101,693,761 (666,666 bp)
Scale
200 kb
hg19
chr1:
101,200,000
101,300,000
101,400,000
101,500,000
101,600,000
0.358524
H3K9me3
1_H3K9me3
0.00941477
3.10655
H3K27me3
2_H3K27me3
0.0504141
ln(x+1) 0.412218
RNA-seq
3_RNA-seq
-1.0501e-16
Basic Gene Annotation Set from GENCODE Version 39lift37 (Ensembl 105)
0.414994
Insulation score based on chromatin interactions
4_Insulation score
-1.56596
Boundary of chromatin interactions
Weak
Strong
Weak
SPI1 chromatin interactions 10bp (filter activated)
SPI1 chromatin interactions 20bp (filter activated)
SPI1 chromatin interactions 50bp (filter activated)
SPI1 chromatin interactions 200bp (filter activated)
SPI1 binding sites
SPI1
SPI1
SPI1
SPI1
CTCF,RAD21,SMC3 binding sites
CTCF
CTCF
CTCF
RAD21
SMC3
SMC3
RAD21
Window Position
chr1:117,576,720-118,243,385 (666,666 bp)
Scale
200 kb
hg19
chr1:
117,700,000
117,800,000
117,900,000
118,000,000
118,100,000
118,200,000
0.181674
H3K9me3
1_H3K9me3
0.00399945
6.41496
H3K27me3
2_H3K27me3
0.0433796
ln(x+1) 0.23588
RNA-seq
3_RNA-seq
-1.79479e-16
Basic Gene Annotation Set from GENCODE Version 39lift37 (Ensembl 105)
0.789888
Insulation score based on chromatin interactions
4_Insulation score
0
-1.20412
Boundary of chromatin interactions
Strong
Weak
Weak
Strong
SPI1 chromatin interactions 10bp (filter activated)
SPI1 chromatin interactions 20bp (filter activated)
SPI1 chromatin interactions 50bp (filter activated)
SPI1 chromatin interactions 200bp (filter activated)
SPI1 binding sites
SPI1
SPI1
SPI1
CTCF,RAD21,SMC3 binding sites
RAD21
SMC3
CTCF
CTCF
SMC3
RAD21
RAD21
SMC3
CTCF
RAD21
CTCF
CTCF
SMC3
RAD21
CTCF
SMC3
RAD21
CTCF
SMC3
Window Position
chr1:34,299,053-34,965,718 (666,666 bp)
Scale
200 kb
hg19
chr1:
34,400,000
34,500,000
34,600,000
34,700,000
34,800,000
34,900,000
0.602125
H3K9me3
1_H3K9me3
0.0545176
3.02678
H3K27me3
2_H3K27me3
0.0368558
ln(x+1) 0.0236875
RNA-seq
3_RNA-seq
-1.23884e-17
Basic Gene Annotation Set from GENCODE Version 39lift37 (Ensembl 105)
0.706887
Insulation score based on chromatin interactions
4_Insulation score
0
-0.636976
Boundary of chromatin interactions
Strong
Weak
SPI1 chromatin interactions 10bp (filter activated)
SPI1 chromatin interactions 20bp (filter activated)
SPI1 chromatin interactions 50bp (filter activated)
SPI1 chromatin interactions 200bp (filter activated)
SPI1 binding sites
SPI1
SPI1
SPI1
SPI1
SPI1
CTCF,RAD21,SMC3 binding sites
CTCF
RAD21
CTCF
CTCF
RAD21
CTCF
SMC3
RAD21
CTCF
CTCF
CTCF
SMC3
RAD21

### SPIB

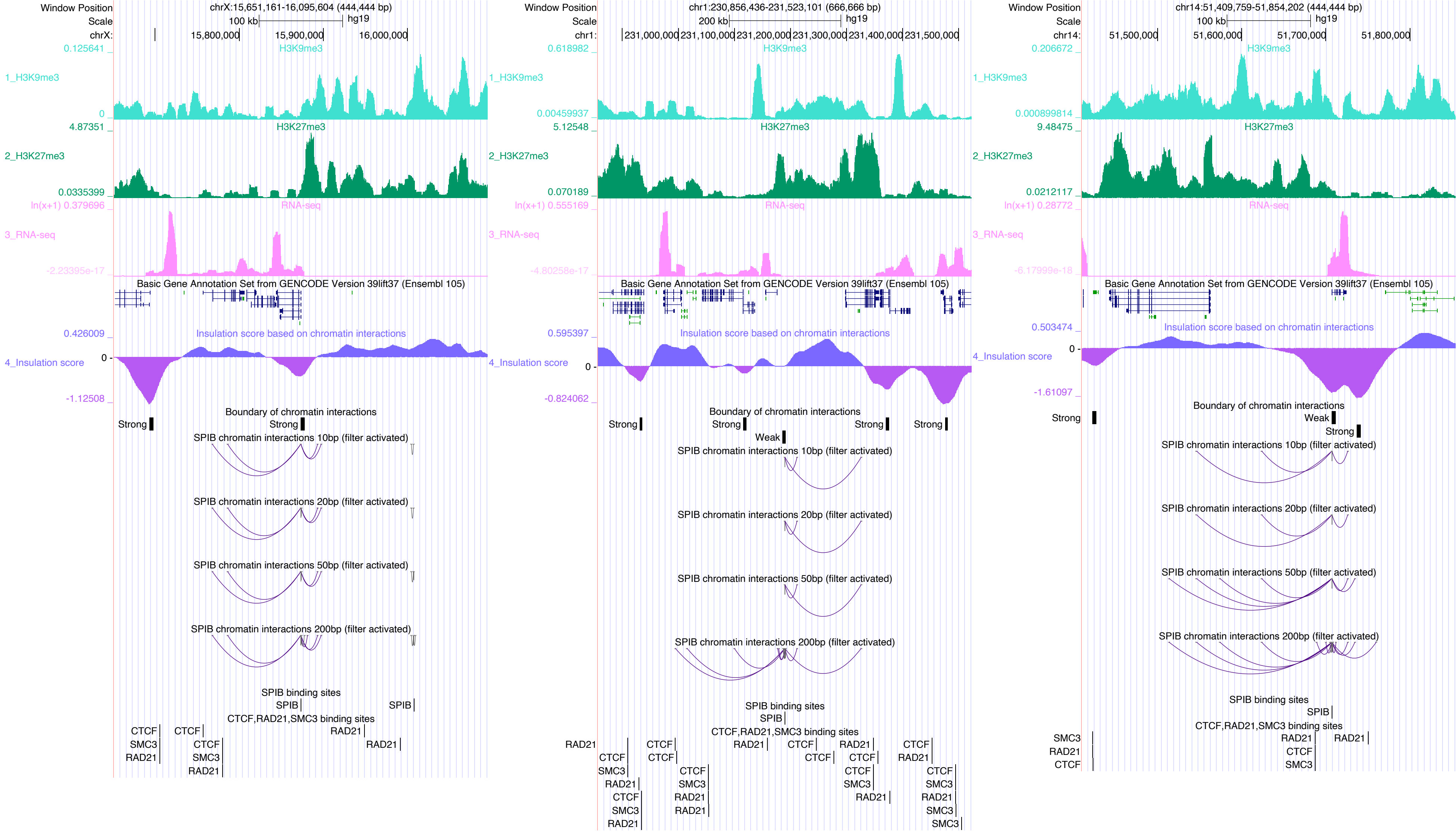

Window Position
chrX:15,651,161-16,095,604 (444,444 bp)
Scale
100 kb
hg19
chrX:
15,800,000
15,900,000
16,000,000
0.125641
H3K9me3
1_H3K9me3
0
4.87351
H3K27me3
2_H3K27me3
0.0335399
ln(x+1) 0.379696
RNA-seq
3_RNA-seq
-2.23395e-17
Basic Gene Annotation Set from GENCODE Version 39lift37 (Ensembl 105)
0.426009
Insulation score based on chromatin interactions
4_Insulation score
0
-1.12508
Boundary of chromatin interactions
Strong
Strong
SPIB chromatin interactions 10bp (filter activated)
SPIB chromatin interactions 20bp (filter activated)
SPIB chromatin interactions 50bp (filter activated)
SPIB chromatin interactions 200bp (filter activated)
SPIB binding sites
SPIB
SPIB
CTCF,RAD21,SMC3 binding sites
CTCF
SMC3
RAD21
CTCF
CTCF
SMC3
RAD21
RAD21
RAD21
Window Position
chr1:230,856,436-231,523,101 (666,666 bp)
Scale
200 kb
hg19
chr1:
231,000,000
231,100,000
231,200,000
231,300,000
231,400,000
231,500,000
0.618982
H3K9me3
1_H3K9me3
0.00459937
5.12548
H3K27me3
2_H3K27me3
0.070189
ln(x+1) 0.555169
RNA-seq
3_RNA-seq
-4.80258e-17
Basic Gene Annotation Set from GENCODE Version 39lift37 (Ensembl 105)
0.595397
Insulation score based on chromatin interactions
4_Insulation score
0
-0.824062
Boundary of chromatin interactions
Strong
Strong
Strong
Strong
Weak
SPIB chromatin interactions 10bp (filter activated)
SPIB chromatin interactions 20bp (filter activated)
SPIB chromatin interactions 50bp (filter activated)
SPIB chromatin interactions 200bp (filter activated)
SPIB binding sites
SPIB
CTCF,RAD21,SMC3 binding sites
RAD21
CTCF
CTCF
SMC3
RAD21
CTCF
SMC3
RAD21
CTCF
CTCF
CTCF
SMC3
RAD21
RAD21
RAD21
CTCF
CTCF
CTCF
CTCF
SMC3
RAD21
CTCF
RAD21
CTCF
SMC3
RAD21
SMC3
SMC3
Window Position
chr14:51,409,759-51,854,202 (444,444 bp)
Scale
100 kb
hg19
chr14:
51,500,000
51,600,000
51,700,000
51,800,000
0.206672
H3K9me3
1_H3K9me3
0.000899814
9.48475
H3K27me3
2_H3K27me3
0.0212117
ln(x+1) 0.28772
RNA-seq
3_RNA-seq
-6.17999e-18
Basic Gene Annotation Set from GENCODE Version 39lift37 (Ensembl 105)
0.503474
Insulation score based on chromatin interactions
4_Insulation score
0
-1.61097
Boundary of chromatin interactions
Strong
Weak
Strong
SPIB chromatin interactions 10bp (filter activated)
SPIB chromatin interactions 20bp (filter activated)
SPIB chromatin interactions 50bp (filter activated)
SPIB chromatin interactions 200bp (filter activated)
SPIB binding sites
SPIB
CTCF,RAD21,SMC3 binding sites
SMC3
RAD21
CTCF
RAD21
CTCF
SMC3
RAD21

### SREBF2

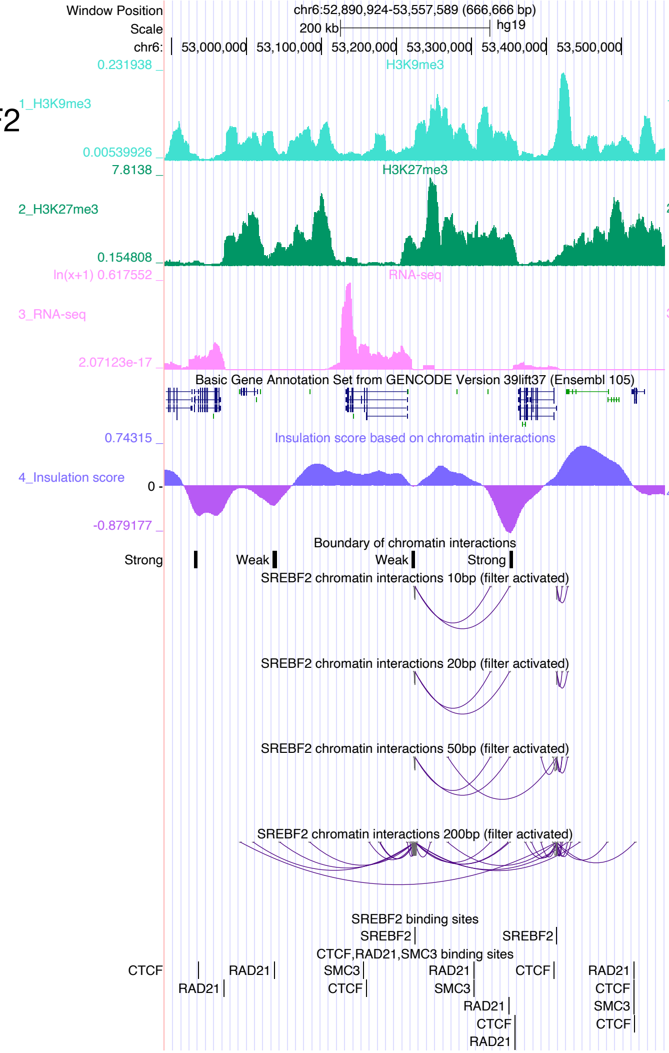

Window Position
chr6:52,890,924-53,557,589 (666,666 bp)
Scale 200 kb hg19
chr6: 53,000,000 53,100,000 53,200,000 53,300,000 53,400,000 53,500,000
1_H3K9me3
2_H3K27me3
3_RNA-seq
Basic Gene Annotation Set from GENCODE Version 39lift37 (Ensembl 105)
4_Insulation score
Insulation score based on chromatin interactions
Boundary of chromatin interactions
SREBF2 chromatin interactions 10bp (filter activated)
SREBF2 chromatin interactions 20bp (filter activated)
SREBF2 chromatin interactions 50bp (filter activated)
SREBF2 chromatin interactions 200bp (filter activated)
SREBF2 binding sites
CTCF,RAD21,SMC3 binding sites


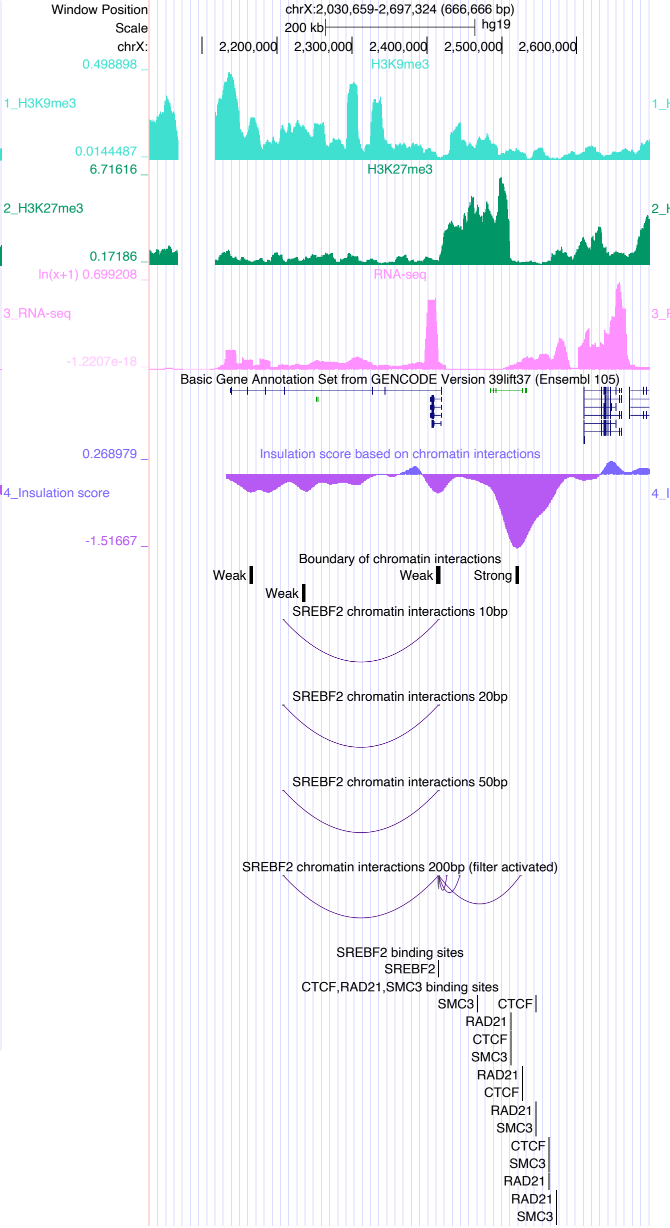

Window Position
chrX:2,030,659-2,697,324 (666,666 bp)
Scale 200 kb hg19
chrX: 2,200,000 2,300,000 2,400,000 2,500,000 2,600,000
1_H3K9me3
2_H3K27me3
3_RNA-seq
Basic Gene Annotation Set from GENCODE Version 39lift37 (Ensembl 105)
4_Insulation score
Insulation score based on chromatin interactions
Boundary of chromatin interactions
SREBF2 chromatin interactions 10bp
SREBF2 chromatin interactions 20bp
SREBF2 chromatin interactions 50bp
SREBF2 chromatin interactions 200bp (filter activated)
SREBF2 binding sites
CTCF,RAD21,SMC3 binding sites


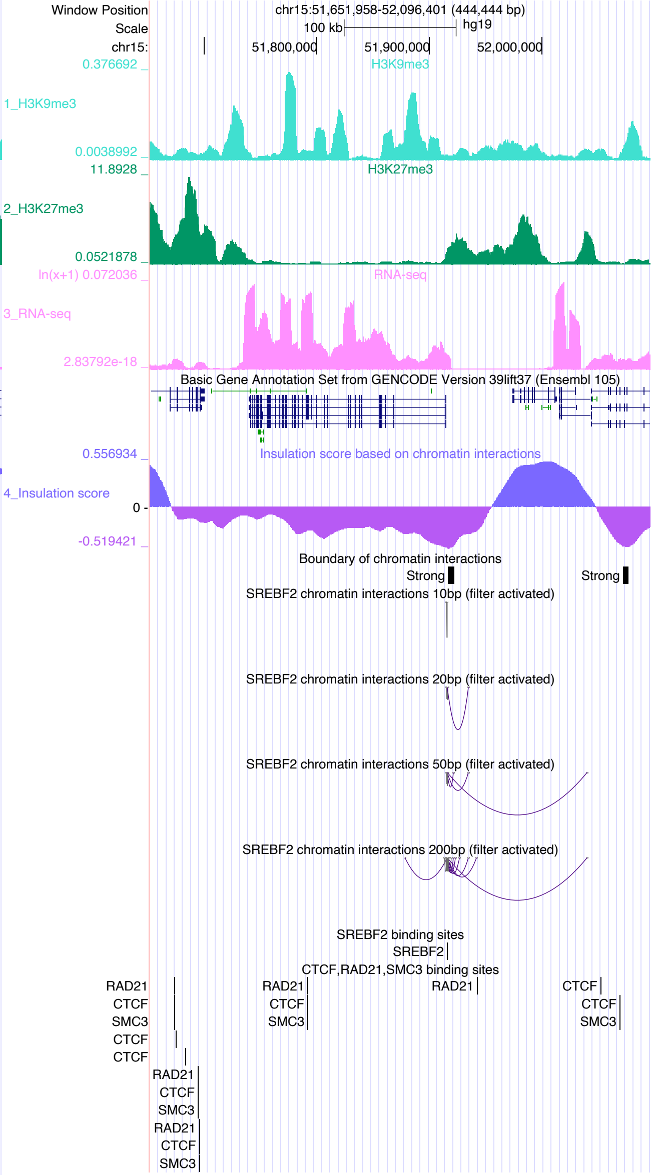

Window Position
chr15:51,651,958-52,096,401 (444,444 bp)
Scale 100 kb hg19
chr15: 51,800,000 51,900,000 52,000,000
1_H3K9me3
2_H3K27me3
3_RNA-seq
Basic Gene Annotation Set from GENCODE Version 39lift37 (Ensembl 105)
4_Insulation score
Insulation score based on chromatin interactions
Boundary of chromatin interactions
SREBF2 chromatin interactions 10bp (filter activated)
SREBF2 chromatin interactions 20bp (filter activated)
SREBF2 chromatin interactions 50bp (filter activated)
SREBF2 chromatin interactions 200bp (filter activated)
SREBF2 binding sites
CTCF,RAD21,SMC3 binding sites

### STAT1

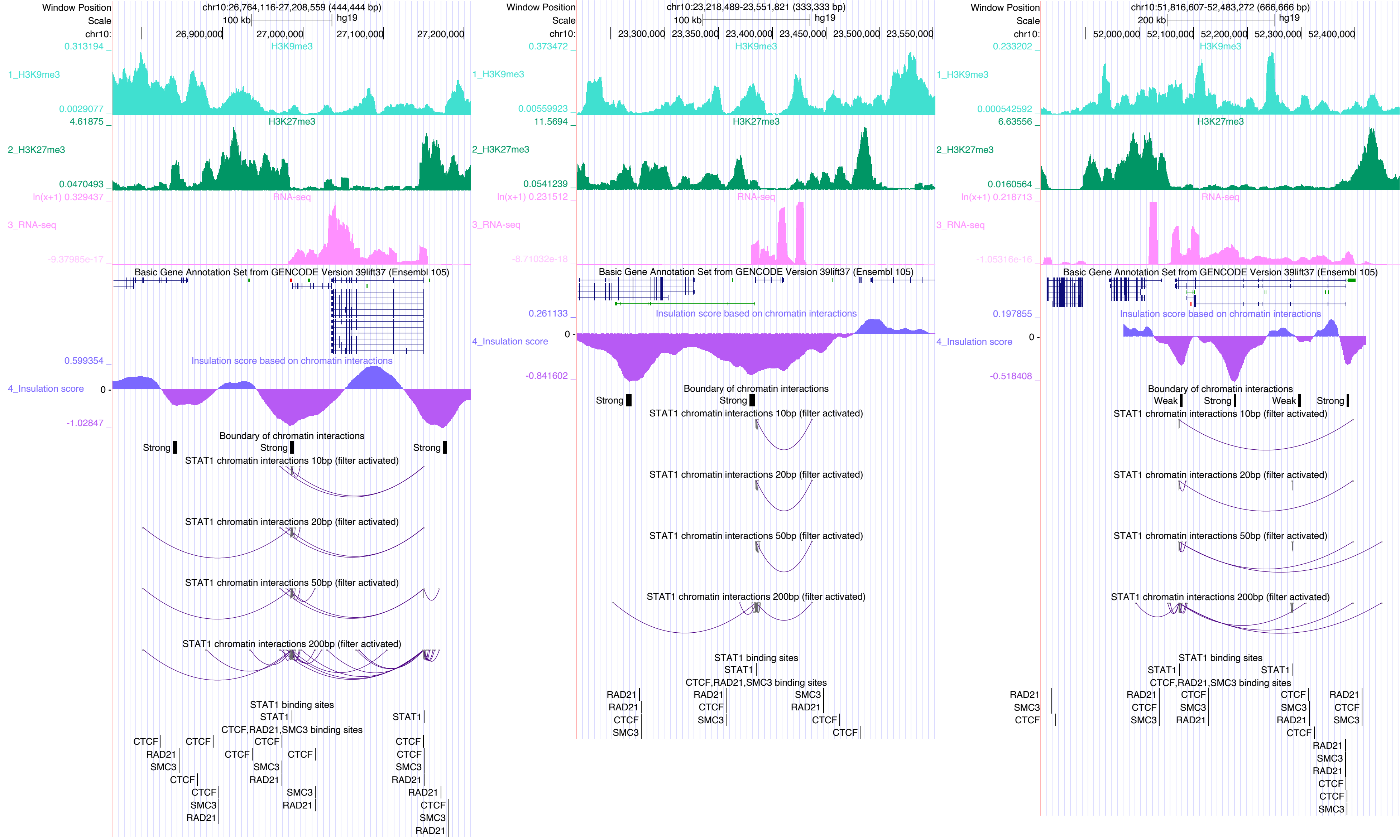

Window Position
Scale
chr10:
chr10:26,764,116-27,208,559 (444,444 bp)
100 kb
hg19
26,900,000
27,000,000
27,100,000
27,200,000
H3K9me3
1_H3K9me3
H3K27me3
2_H3K27me3
RNA-seq
3_RNA-seq
Basic Gene Annotation Set from GENCODE Version 39lift37 (Ensembl 105)
Insulation score based on chromatin interactions
4_Insulation score
Boundary of chromatin interactions
Strong
Weak
STAT1 chromatin interactions 10bp (filter activated)
STAT1 chromatin interactions 20bp (filter activated)
STAT1 chromatin interactions 50bp (filter activated)
STAT1 chromatin interactions 200bp (filter activated)
STAT1 binding sites
CTCF,RAD21,SMC3 binding sites
chr10:23,218,489-23,551,821 (333,333 bp)
23,300,000
23,350,000
23,400,000
23,450,000
23,500,000
23,550,000
chr10:51,816,607-52,483,272 (666,666 bp)
200 kb
52,000,000
52,100,000
52,200,000
52,300,000
52,400,000

#### STAT3

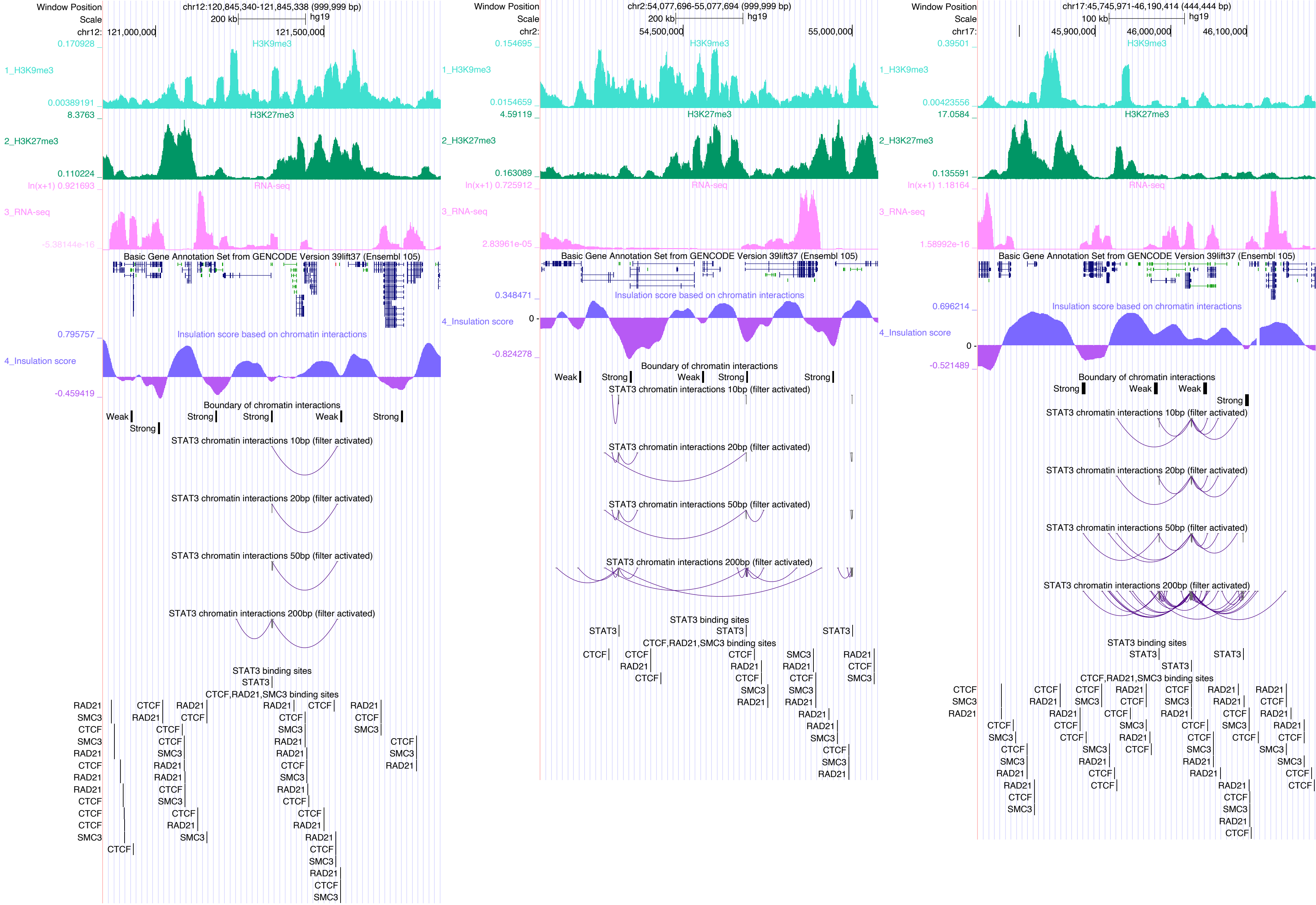

Window Position
chr12:120,845,340-121,845,338 (999,999 bp)
Scale
200 kb
hg19
chr12:
121,000,000
121,500,000
1_H3K9me3
H3K9me3
2_H3K27me3
H3K27me3
3_RNA-seq
RNA-seq
Basic Gene Annotation Set from GENCODE Version 39lift37 (Ensembl 105)
4_Insulation score
Insulation score based on chromatin interactions
Boundary of chromatin interactions
STAT3 chromatin interactions 10bp (filter activated)
STAT3 chromatin interactions 20bp (filter activated)
STAT3 chromatin interactions 50bp (filter activated)
STAT3 chromatin interactions 200bp (filter activated)
STAT3 binding sites
CTCF,RAD21,SMC3 binding sites
Window Position
chr2:54,077,696-55,077,694 (999,999 bp)
chr2:
54,500,000
55,000,000
Window Position
chr17:45,745,971-46,190,414 (444,444 bp)
100 kb
chr17:
45,900,000
46,000,000
46,100,000

### TAF1

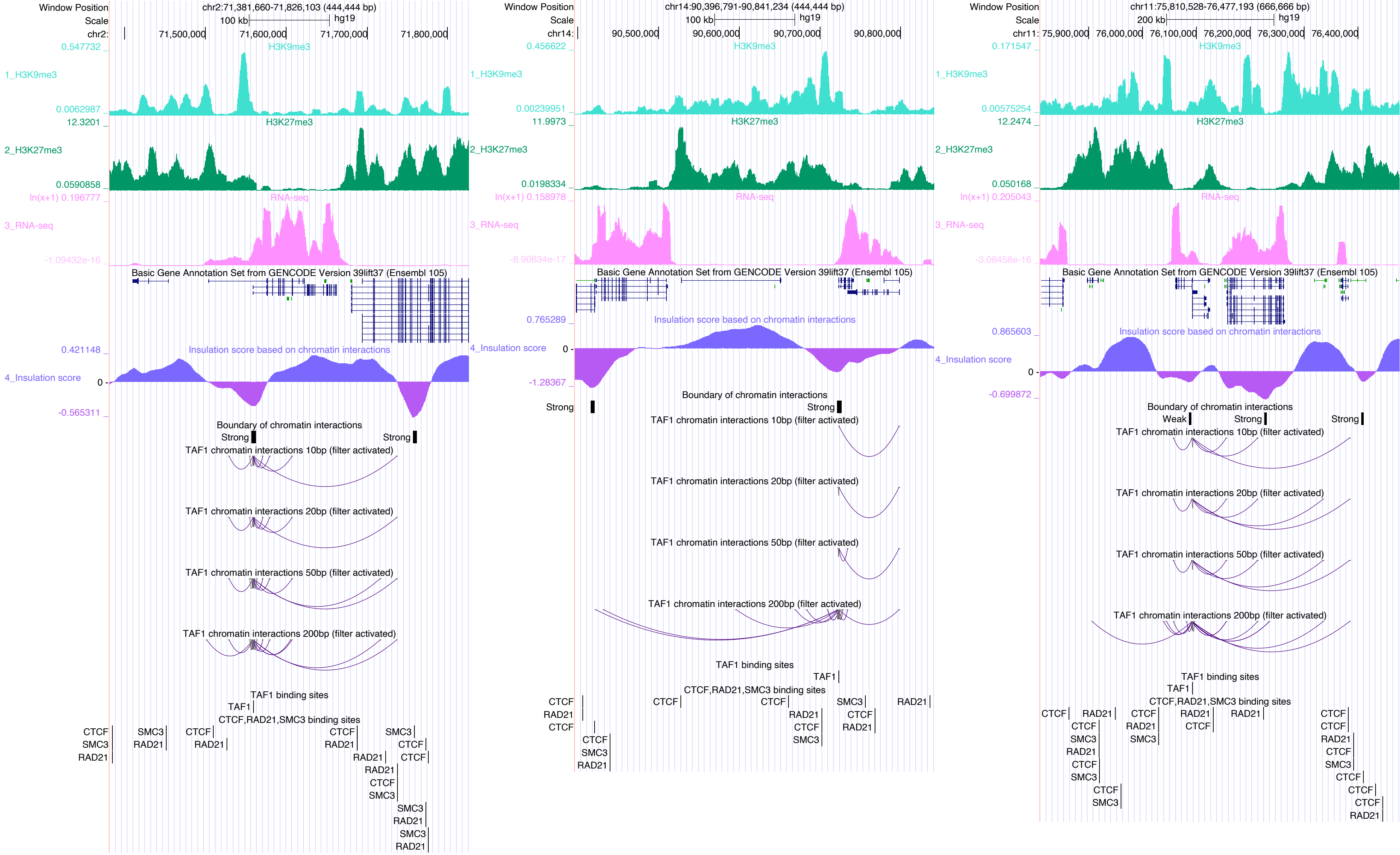

Window Position chr2:71,381,660-71,826,103 (444,444 bp)
Scale 100 kb hg19
chr2: 71,500,000 71,600,000 71,700,000 71,800,000
1_H3K9me3
H3K9me3
2_H3K27me3
H3K27me3
3_RNA-seq
RNA-seq
Basic Gene Annotation Set from GENCODE Version 39lift37 (Ensembl 105)
4_Insulation score
Insulation score based on chromatin interactions
Boundary of chromatin interactions
Strong
TAF1 chromatin interactions 10bp (filter activated)
TAF1 chromatin interactions 20bp (filter activated)
TAF1 chromatin interactions 50bp (filter activated)
TAF1 chromatin interactions 200bp (filter activated)
TAF1 binding sites
TAF1
CTCF,RAD21,SMC3 binding sites
CTCF
SMC3
RAD21
Window Position chr14:90,396,791-90,841,234 (444,444 bp)
chr14: 90,500,000 90,600,000 90,700,000 90,800,000
Window Position chr11:75,810,528-76,477,193 (666,666 bp)
Scale 200 kb hg19
chr11: 75,900,000 76,000,000 76,100,000 76,200,000 76,300,000 76,400,000
Weak

### TBX21

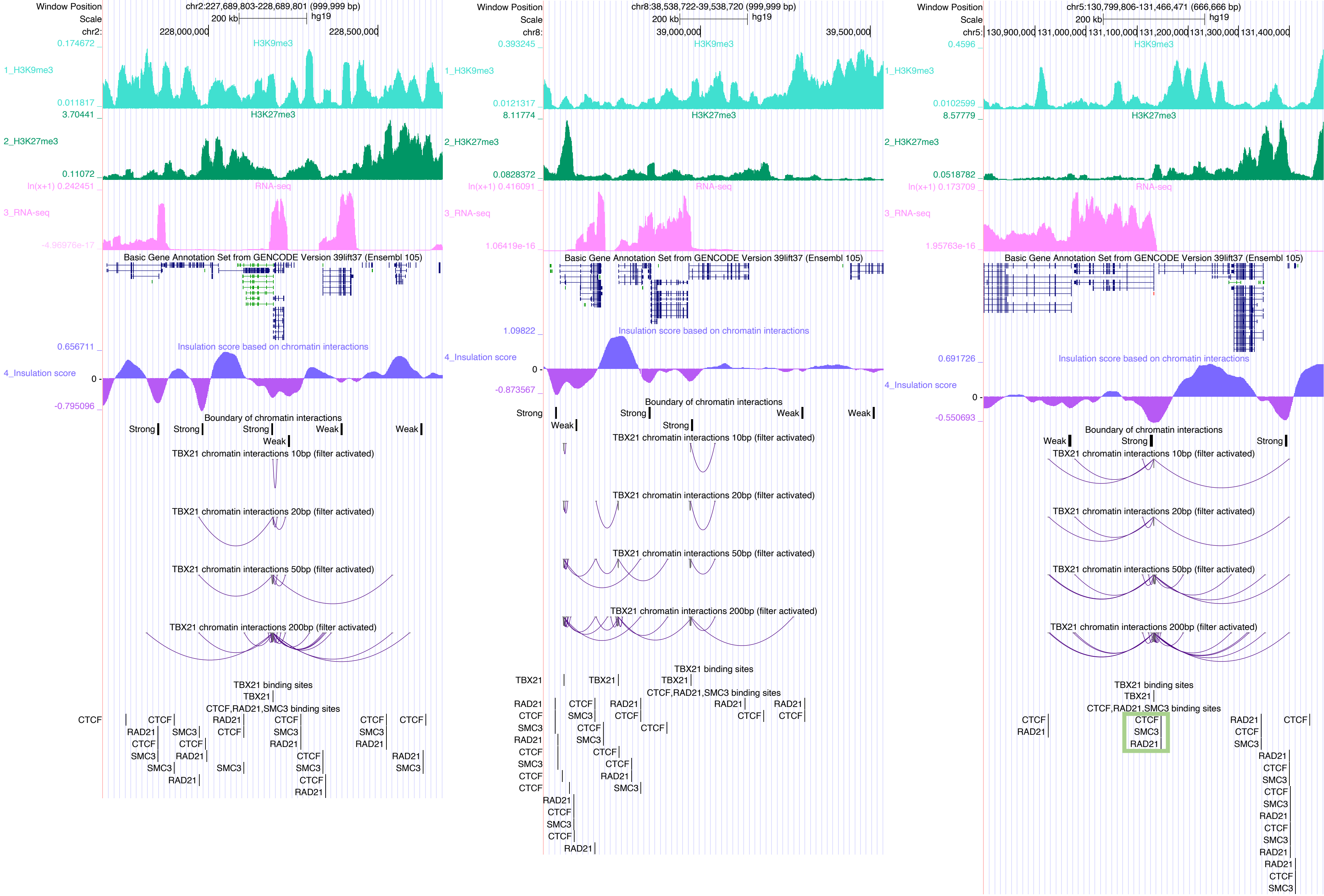

Window Position
Scale
chr2:227,689,803-228,689,801 (999,999 bp)
200 kb
hg19
chr2:
228,000,000
228,500,000
H3K9me3
1_H3K9me3
0.174672
0.011817
H3K27me3
2_H3K27me3
3.70441
0.11072
RNA-seq
3_RNA-seq
ln(x+1) 0.242451
-4.96976e-17
Basic Gene Annotation Set from GENCODE Version 39lift37 (Ensembl 105)
Insulation score based on chromatin interactions
4_Insulation score
0.656711
0
-0.795096
Boundary of chromatin interactions
Strong
Weak
TBX21 chromatin interactions 10bp (filter activated)
TBX21 chromatin interactions 20bp (filter activated)
TBX21 chromatin interactions 50bp (filter activated)
TBX21 chromatin interactions 200bp (filter activated)
TBX21 binding sites
TBX21
CTCF,RAD21,SMC3 binding sites
CTCF
RAD21
SMC3
chr8:38,538,722-39,538,720 (999,999 bp)
chr8:
39,000,000
39,500,000
0.393245
0.0121317
8.11774
0.0828372
ln(x+1) 0.416091
1.06419e-16
1.09822
-0.873567
chr5:130,799,806-131,466,471 (666,666 bp)
chr5:
130,900,000
131,000,000
131,100,000
131,200,000
131,300,000
131,400,000
0.4596
0.0102599
8.57779
0.0518782
ln(x+1) 0.173709
1.95763e-16
0.691726
-0.550693

### TCF3

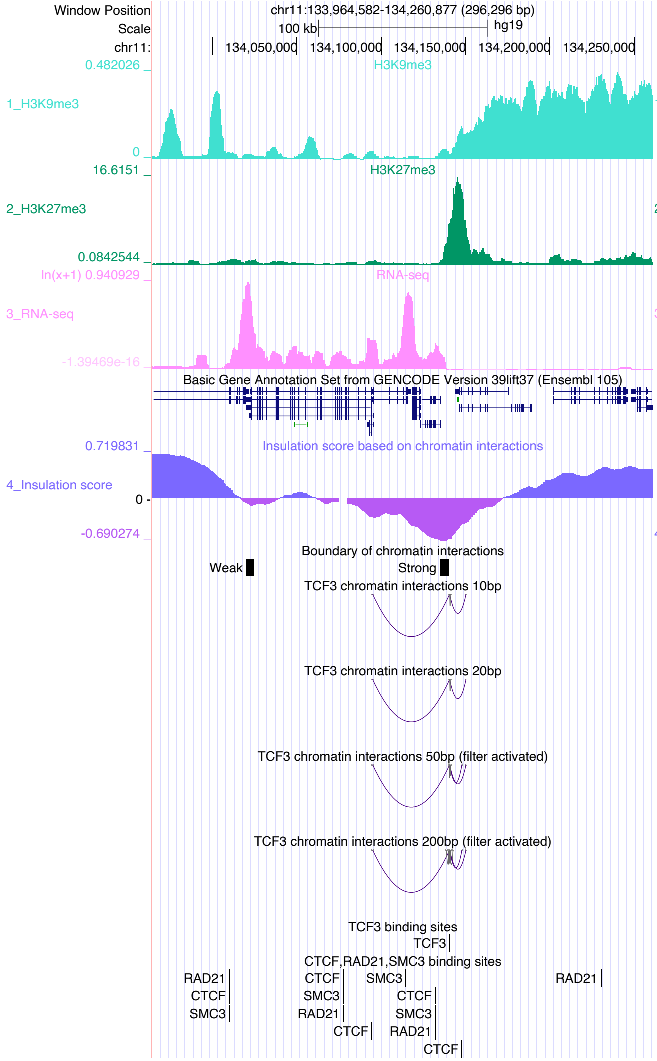

Window Position chr11:133,964,582-134,260,877 (296,296 bp)
Scale 100 kb hg19
H3K9me3
1_H3K9me3
H3K27me3
2_H3K27me3
RNA-seq
3_RNA-seq
Basic Gene Annotation Set from GENCODE Version 39lift37 (Ensembl 105)
Insulation score based on chromatin interactions
4_Insulation score
Boundary of chromatin interactions
TCF3 chromatin interactions 10bp
TCF3 chromatin interactions 20bp
TCF3 chromatin interactions 50bp (filter activated)
TCF3 chromatin interactions 200bp (filter activated)
TCF3 binding sites
CTCF,RAD21,SMC3 binding sites


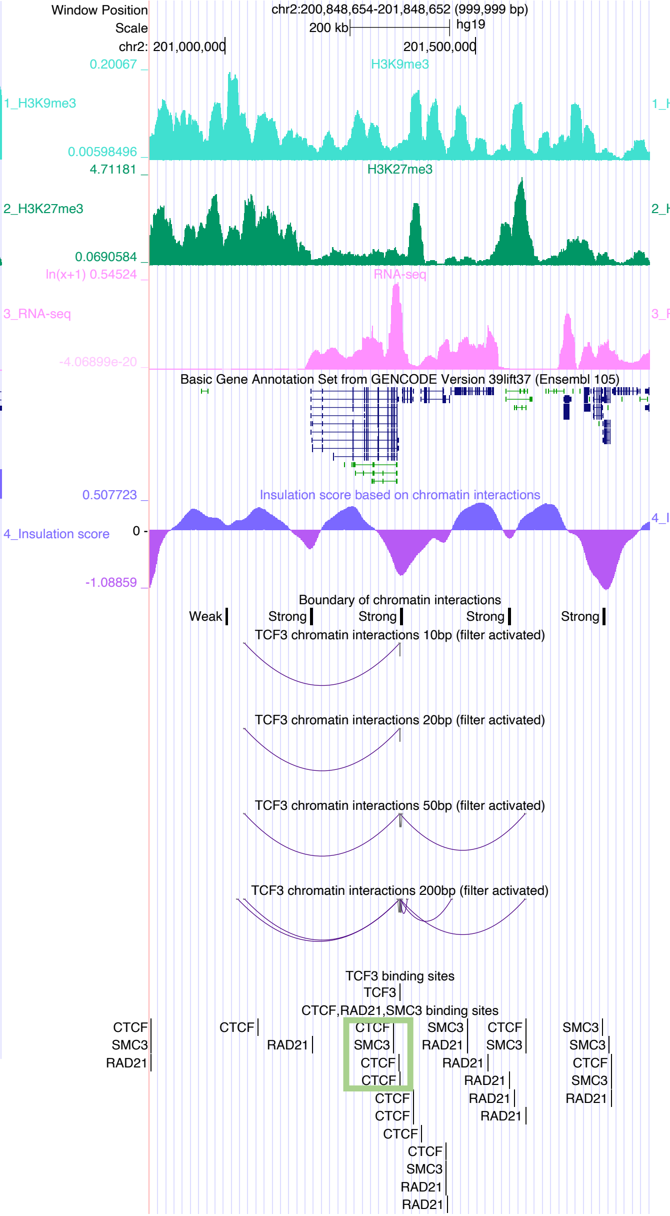

Window Position chr2:200,848,654-201,848,652 (999,999 bp)
Scale 200 kb hg19
H3K9me3
1_H3K9me3
H3K27me3
2_H3K27me3
RNA-seq
3_RNA-seq
Basic Gene Annotation Set from GENCODE Version 39lift37 (Ensembl 105)
Insulation score based on chromatin interactions
4_Insulation score
Boundary of chromatin interactions
TCF3 chromatin interactions 10bp (filter activated)
TCF3 chromatin interactions 20bp (filter activated)
TCF3 chromatin interactions 50bp (filter activated)
TCF3 chromatin interactions 200bp (filter activated)
TCF3 binding sites
CTCF,RAD21,SMC3 binding sites


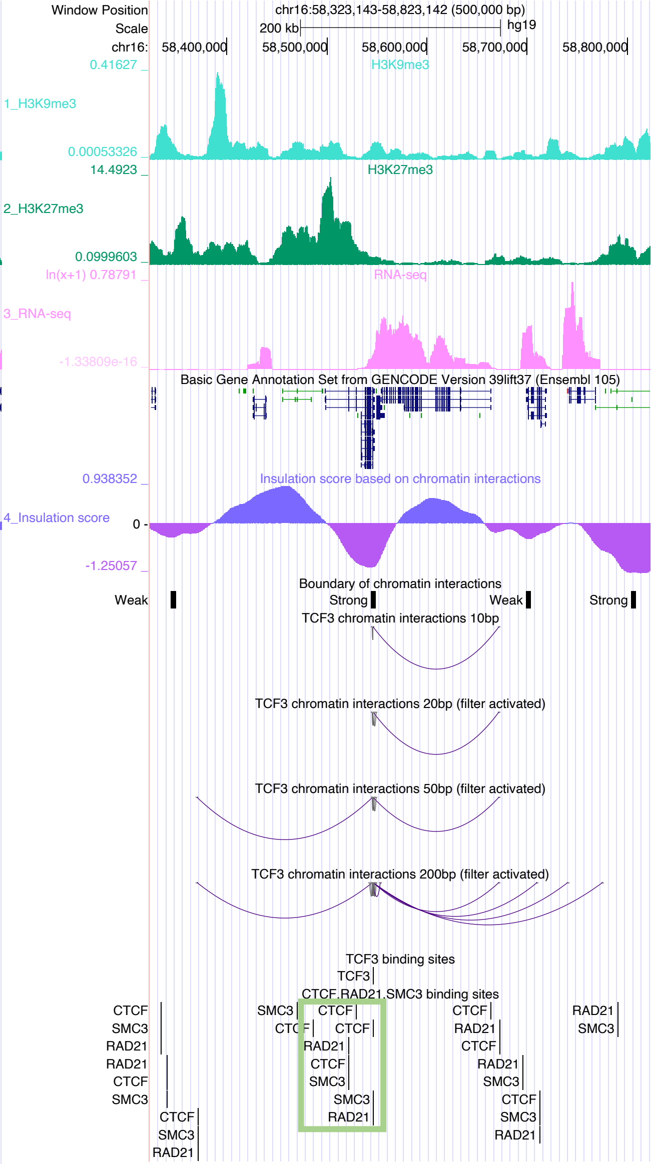

Window Position chr16:58,323,143-58,823,142 (500,000 bp)
Scale 200 kb hg19
H3K9me3
1_H3K9me3
H3K27me3
2_H3K27me3
RNA-seq
3_RNA-seq
Basic Gene Annotation Set from GENCODE Version 39lift37 (Ensembl 105)
Insulation score based on chromatin interactions
4_Insulation score
Boundary of chromatin interactions
TCF3 chromatin interactions 10bp
TCF3 chromatin interactions 20bp (filter activated)
TCF3 chromatin interactions 50bp (filter activated)
TCF3 chromatin interactions 200bp (filter activated)
TCF3 binding sites
CTCF,RAD21,SMC3 binding sites

### TCF4

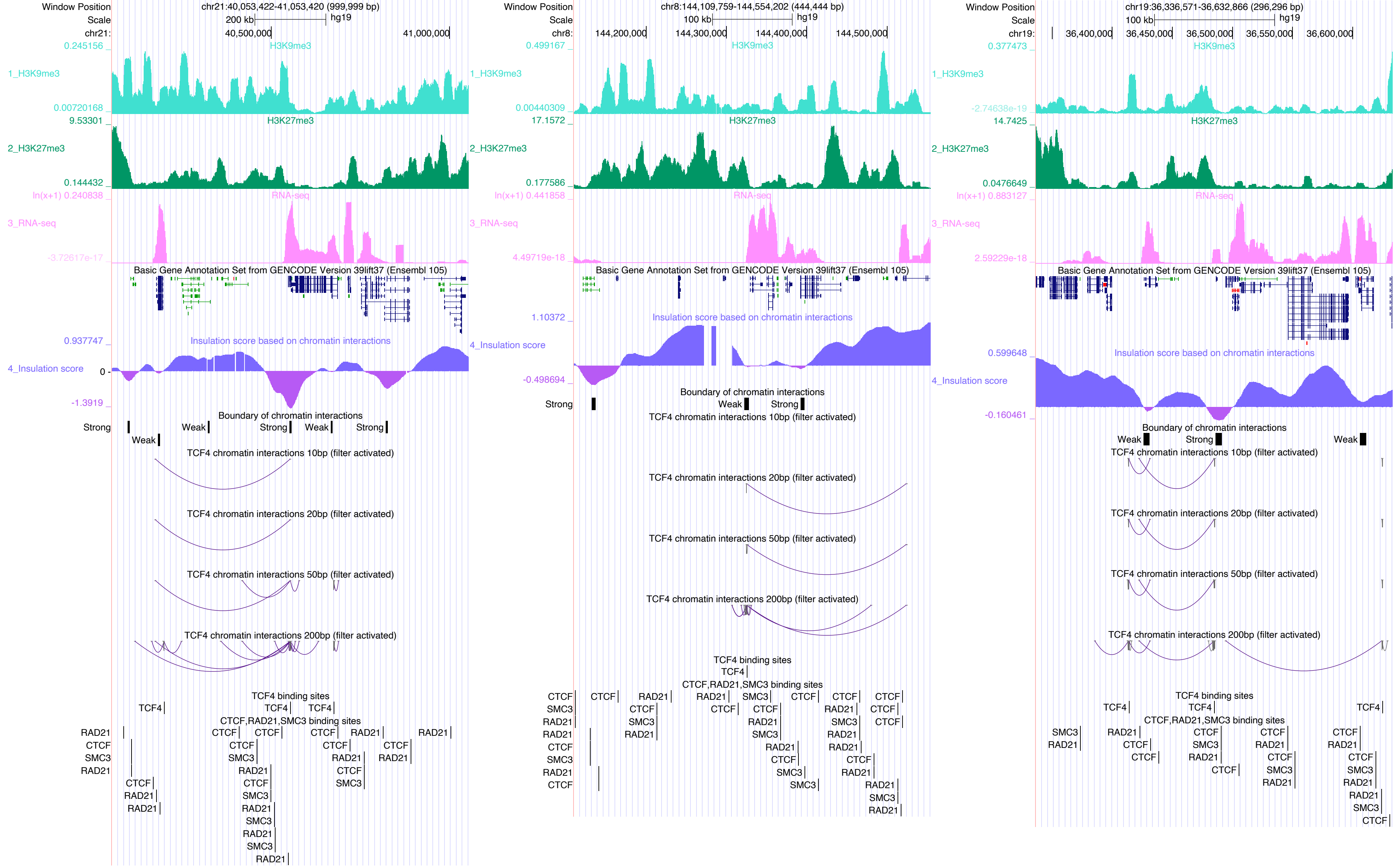
Window Position
chr21:40,053,422-41,053,420 (999,999 bp)
Scale
200 kb
hg19
chr21:
40,500,000
41,000,000
H3K9me3
1_H3K9me3
H3K27me3
2_H3K27me3
RNA-seq
3_RNA-seq
Basic Gene Annotation Set from GENCODE Version 39lift37 (Ensembl 105)
Insulation score based on chromatin interactions
4_Insulation score
Boundary of chromatin interactions
Strong
Weak
TCF4 chromatin interactions 10bp (filter activated)
TCF4 chromatin interactions 20bp (filter activated)
TCF4 chromatin interactions 50bp (filter activated)
TCF4 chromatin interactions 200bp (filter activated)
TCF4 binding sites
CTCF,RAD21,SMC3 binding sites
chr8:144,109,759-144,554,202 (444,444 bp)
100 kb
144,200,000
144,300,000
144,400,000
144,500,000
chr19:36,336,571-36,632,866 (296,296 bp)
36,400,000
36,450,000
36,500,000
36,550,000
36,600,000

### TCF12

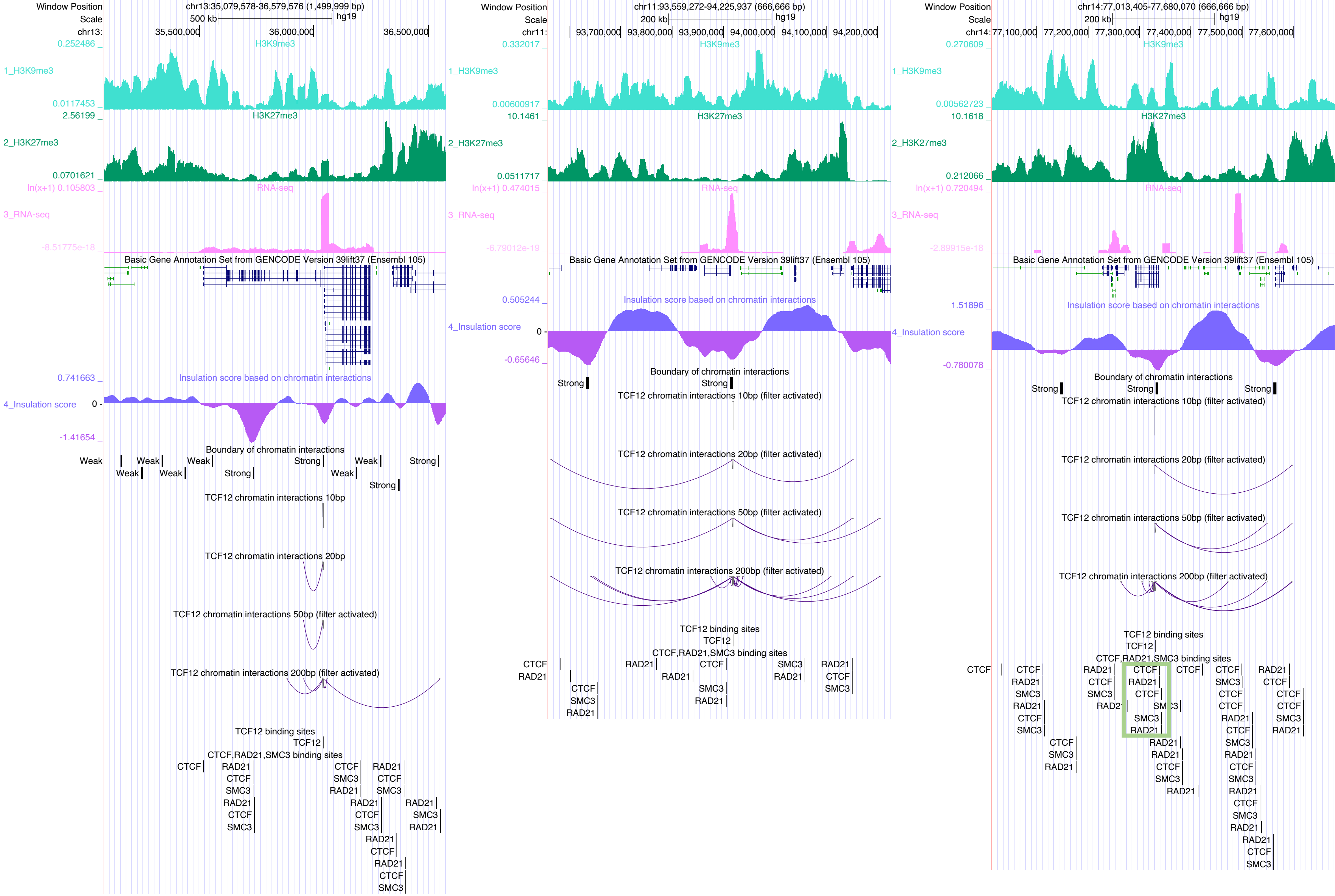

Window Position
Scale
chr13:35,079,578-36,579,576 (1,499,999 bp)
500 kb
hg19
35,500,000
36,000,000
36,500,000
1_H3K9me3
H3K9me3
2_H3K27me3
H3K27me3
3_RNA-seq
RNA-seq
Basic Gene Annotation Set from GENCODE Version 39lift37 (Ensembl 105)
4_Insulation score
Insulation score based on chromatin interactions
Boundary of chromatin interactions
Weak
Strong
TCF12 chromatin interactions 10bp
TCF12 chromatin interactions 20bp
TCF12 chromatin interactions 50bp (filter activated)
TCF12 chromatin interactions 200bp (filter activated)
TCF12 binding sites
TCF12
CTCF,RAD21,SMC3 binding sites
CTCF
RAD21
SMC3
chr11:93,559,272-94,225,937 (666,666 bp)
200 kb
93,700,000
93,800,000
93,900,000
94,000,000
94,100,000
94,200,000
TCF12 chromatin interactions 10bp (filter activated)
TCF12 chromatin interactions 20bp (filter activated)
chr14:77,013,405-77,680,070 (666,666 bp)
77,100,000
77,200,000
77,300,000
77,400,000
77,500,000
77,600,000

### TEAD3

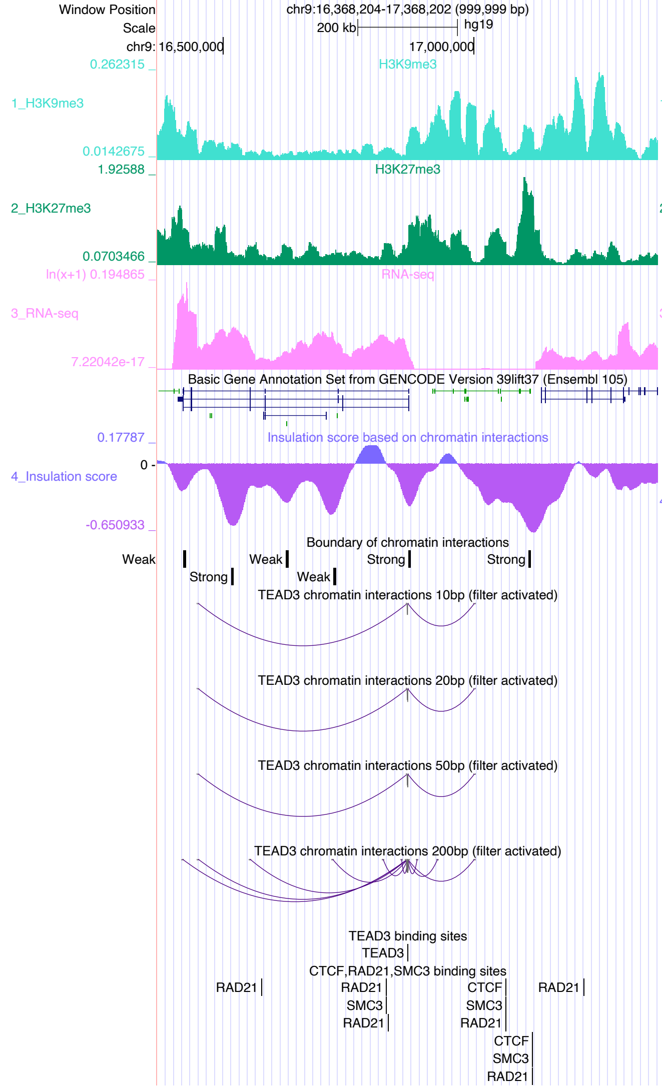

Window Position chr9:16,368,204-17,368,202 (999,999 bp)
Scale 200 kb hg19
chr9: 16,500,000 17,000,000
1_H3K9me3 H3K9me3 0.262315 0.0142675
2_H3K27me3 H3K27me3 1.92588 0.0703466
3_RNA-seq RNA-seq ln(x+1) 0.194865 7.22042e-17
Basic Gene Annotation Set from GENCODE Version 39lift37 (Ensembl 105)
4_Insulation score Insulation score based on chromatin interactions 0.17787 0 -0.650933
Boundary of chromatin interactions
Weak Strong Weak Weak Strong Strong
TEAD3 chromatin interactions 10bp (filter activated)
TEAD3 chromatin interactions 20bp (filter activated)
TEAD3 chromatin interactions 50bp (filter activated)
TEAD3 chromatin interactions 200bp (filter activated)
TEAD3 binding sites
TEAD3
CTCF,RAD21,SMC3 binding sites
RAD21 RAD21 SMC3 RAD21 CTCF SMC3 RAD21 RAD21 CTCF SMC3 RAD21


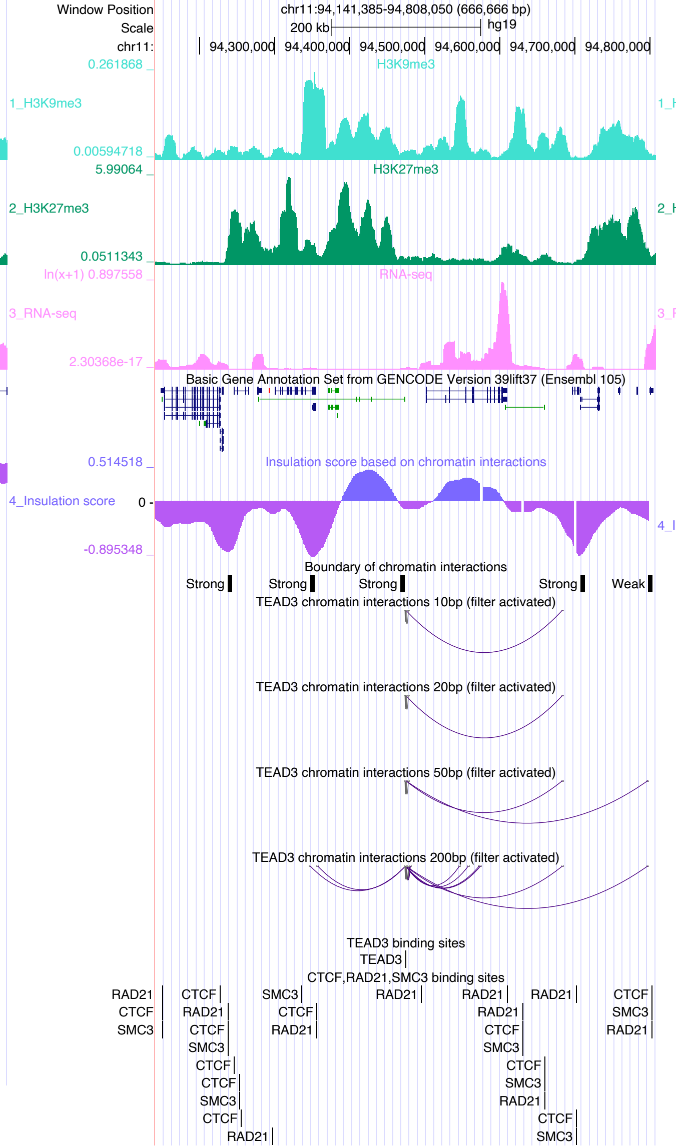

Window Position chr11:94,141,385-94,808,050 (666,666 bp)
Scale 200 kb hg19
chr11: 94,300,000 94,400,000 94,500,000 94,600,000 94,700,000 94,800,000
1_H3K9me3 H3K9me3 0.261868 0.00594718
2_H3K27me3 H3K27me3 5.99064 0.0511343
3_RNA-seq RNA-seq ln(x+1) 0.897558 2.30368e-17
Basic Gene Annotation Set from GENCODE Version 39lift37 (Ensembl 105)
4_Insulation score Insulation score based on chromatin interactions 0.514518 0 -0.895348
Boundary of chromatin interactions
Strong Strong Strong Strong Weak
TEAD3 chromatin interactions 10bp (filter activated)
TEAD3 chromatin interactions 20bp (filter activated)
TEAD3 chromatin interactions 50bp (filter activated)
TEAD3 chromatin interactions 200bp (filter activated)
TEAD3 binding sites
TEAD3
CTCF,RAD21,SMC3 binding sites
RAD21 CTCF SMC3 CTCF RAD21 CTCF SMC3 CTCF CTCF SMC3 CTCF RAD21 SMC3 CTCF RAD21 RAD21 RAD21 RAD21 CTCF SMC3 CTCF SMC3 RAD21 CTCF SMC3 RAD21 CTCF SMC3 RAD21


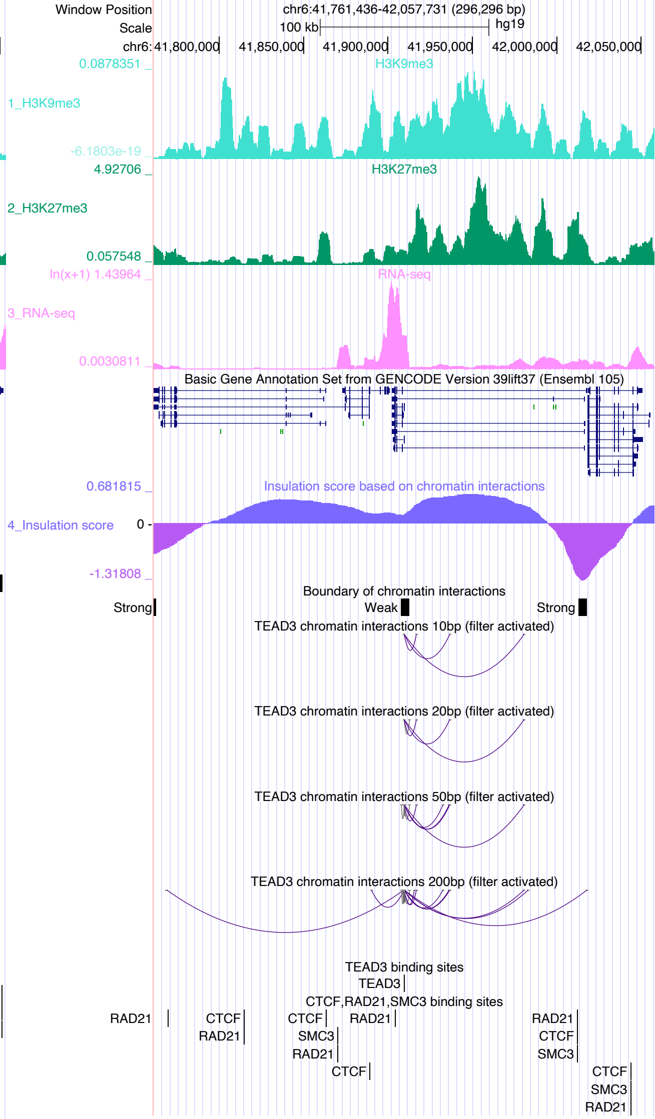

Window Position chr6:41,761,436-42,057,731 (296,296 bp)
Scale 100 kb hg19
chr6: 41,800,000 41,850,000 41,900,000 41,950,000 42,000,000 42,050,000
1_H3K9me3 H3K9me3 0.0878351 -6.1803e-19
2_H3K27me3 H3K27me3 4.92706 0.057548
3_RNA-seq RNA-seq ln(x+1) 1.43964 0.0030811
Basic Gene Annotation Set from GENCODE Version 39lift37 (Ensembl 105)
4_Insulation score Insulation score based on chromatin interactions 0.681815 0 -1.31808
Boundary of chromatin interactions
Strong Weak Strong
TEAD3 chromatin interactions 10bp (filter activated)
TEAD3 chromatin interactions 20bp (filter activated)
TEAD3 chromatin interactions 50bp (filter activated)
TEAD3 chromatin interactions 200bp (filter activated)
TEAD3 binding sites
TEAD3
CTCF,RAD21,SMC3 binding sites
RAD21 CTCF RAD21 CTCF SMC3 RAD21 CTCF RAD21 RAD21 CTCF SMC3 CTCF SMC3 RAD21

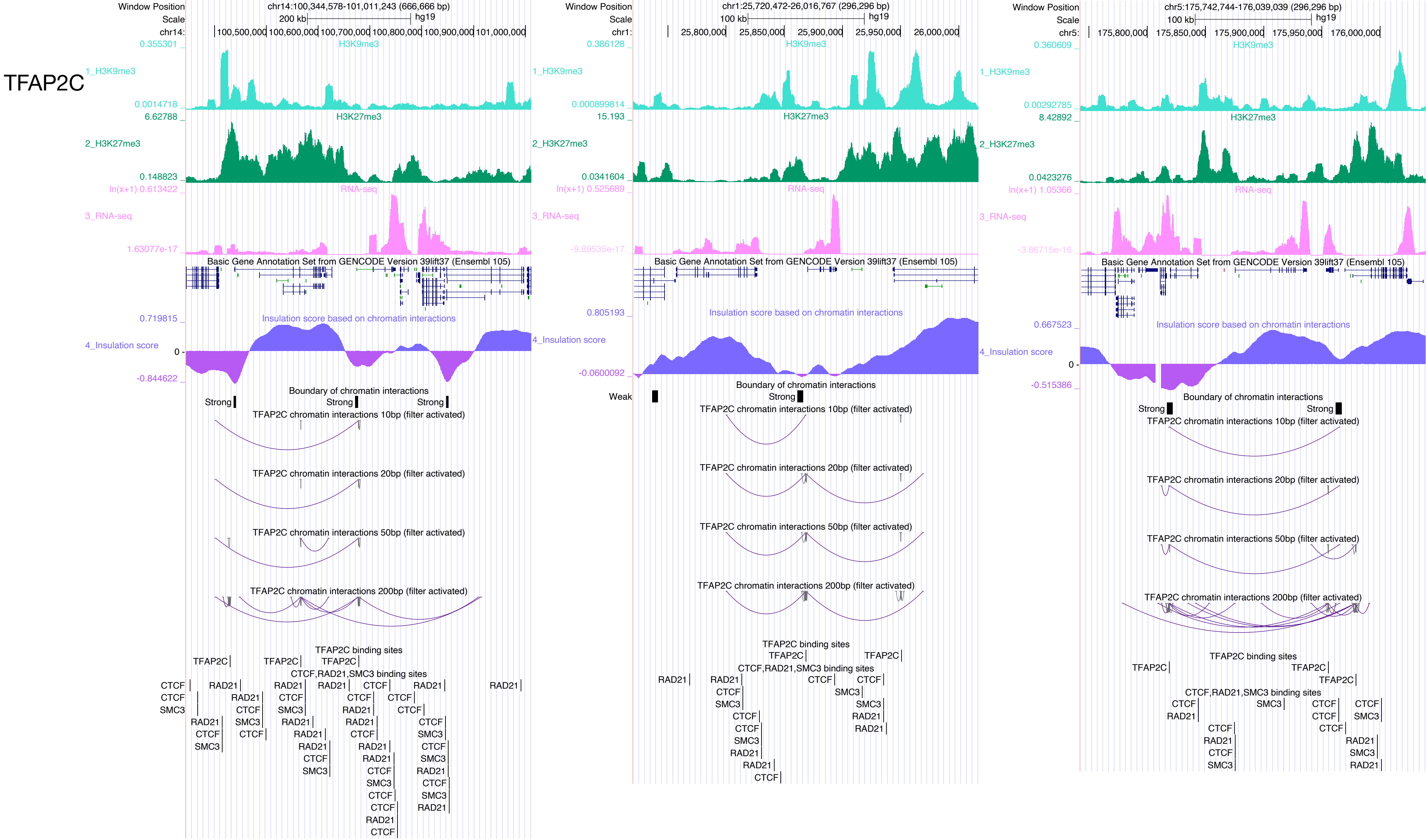

TFAP2C
Window Position
chr14:100,344,578-101,011,243 (666,666 bp)
Scale
200 kb
hg19
chr14:
100,500,000
100,600,000
100,700,000
100,800,000
100,900,000
101,000,000
H3K9me3
1_H3K9me3
H3K27me3
2_H3K27me3
RNA-seq
3_RNA-seq
Basic Gene Annotation Set from GENCODE Version 39lift37 (Ensembl 105)
Insulation score based on chromatin interactions
4_Insulation score
Boundary of chromatin interactions
Strong
Weak
TFAP2C chromatin interactions 10bp (filter activated)
TFAP2C chromatin interactions 20bp (filter activated)
TFAP2C chromatin interactions 50bp (filter activated)
TFAP2C chromatin interactions 200bp (filter activated)
TFAP2C binding sites
CTCF,RAD21,SMC3 binding sites
chr1:25,720,472-26,016,767 (296,296 bp)
100 kb
chr1:
25,800,000
25,850,000
25,900,000
25,950,000
26,000,000
chr5:175,742,744-176,039,039 (296,296 bp)
chr5:
175,800,000
175,850,000
175,900,000
175,950,000
176,000,000

### TFAP4

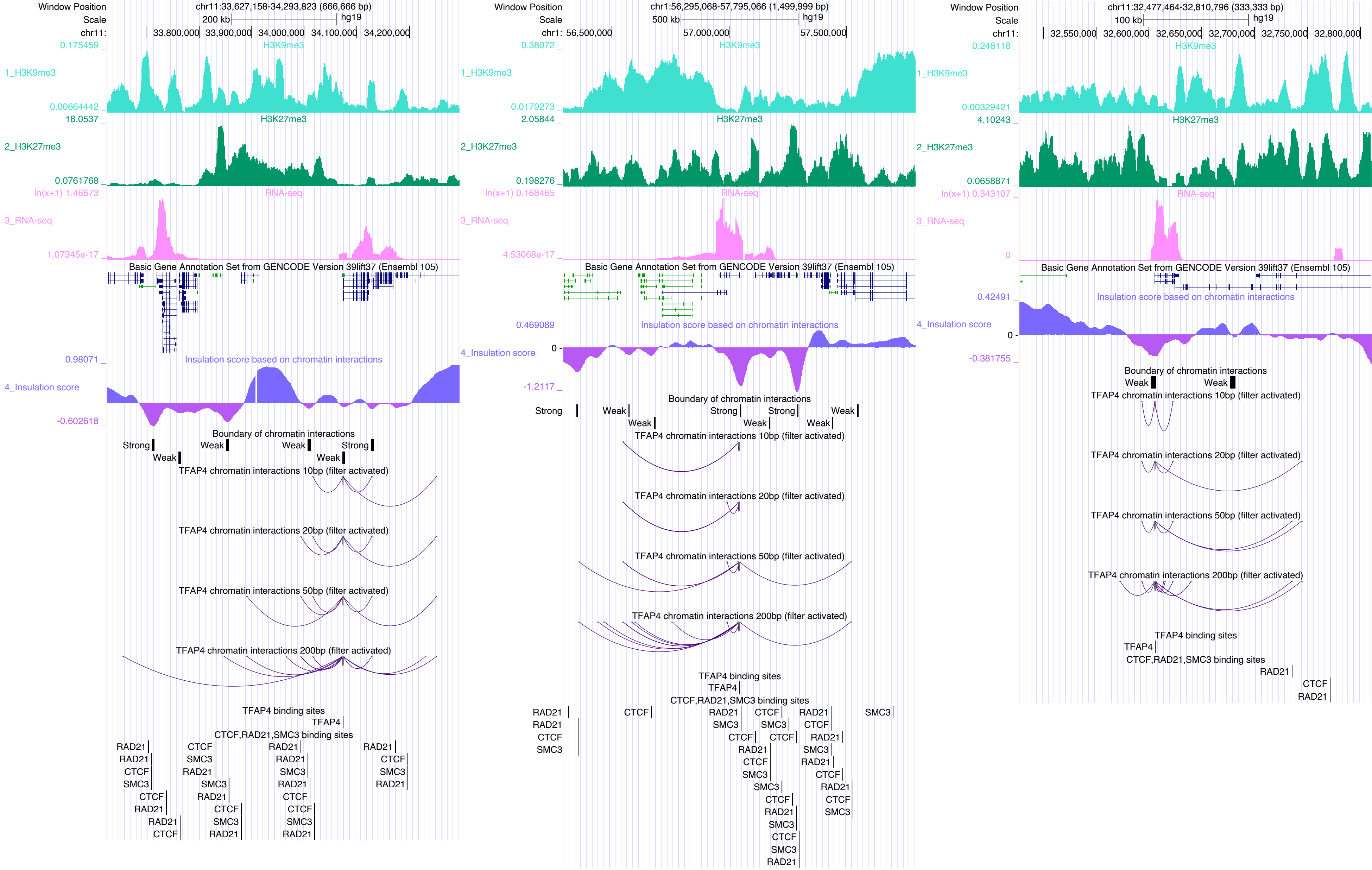

Window Position chr11:33,627,158-34,293,823 (666,666 bp)
Scale 200 kb hg19
chr11: 33,800,000 33,900,000 34,000,000 34,100,000 34,200,000
1_H3K9me3
2_H3K27me3
3_RNA-seq
Basic Gene Annotation Set from GENCODE Version 39lift37 (Ensembl 105)
4_Insulation score
Insulation score based on chromatin interactions
Boundary of chromatin interactions
TFAP4 chromatin interactions 10bp (filter activated)
TFAP4 chromatin interactions 20bp (filter activated)
TFAP4 chromatin interactions 50bp (filter activated)
TFAP4 chromatin interactions 200bp (filter activated)
TFAP4 binding sites
CTCF,RAD21,SMC3 binding sites
Window Position chr1:56,295,068-57,795,066 (1,499,999 bp)
Scale 500 kb hg19
chr1: 56,500,000 57,000,000 57,500,000
Window Position chr11:32,477,464-32,810,796 (333,333 bp)
Scale 100 kb hg19
chr11: 32,550,000 32,600,000 32,650,000 32,700,000 32,750,000 32,800,000

# TP53

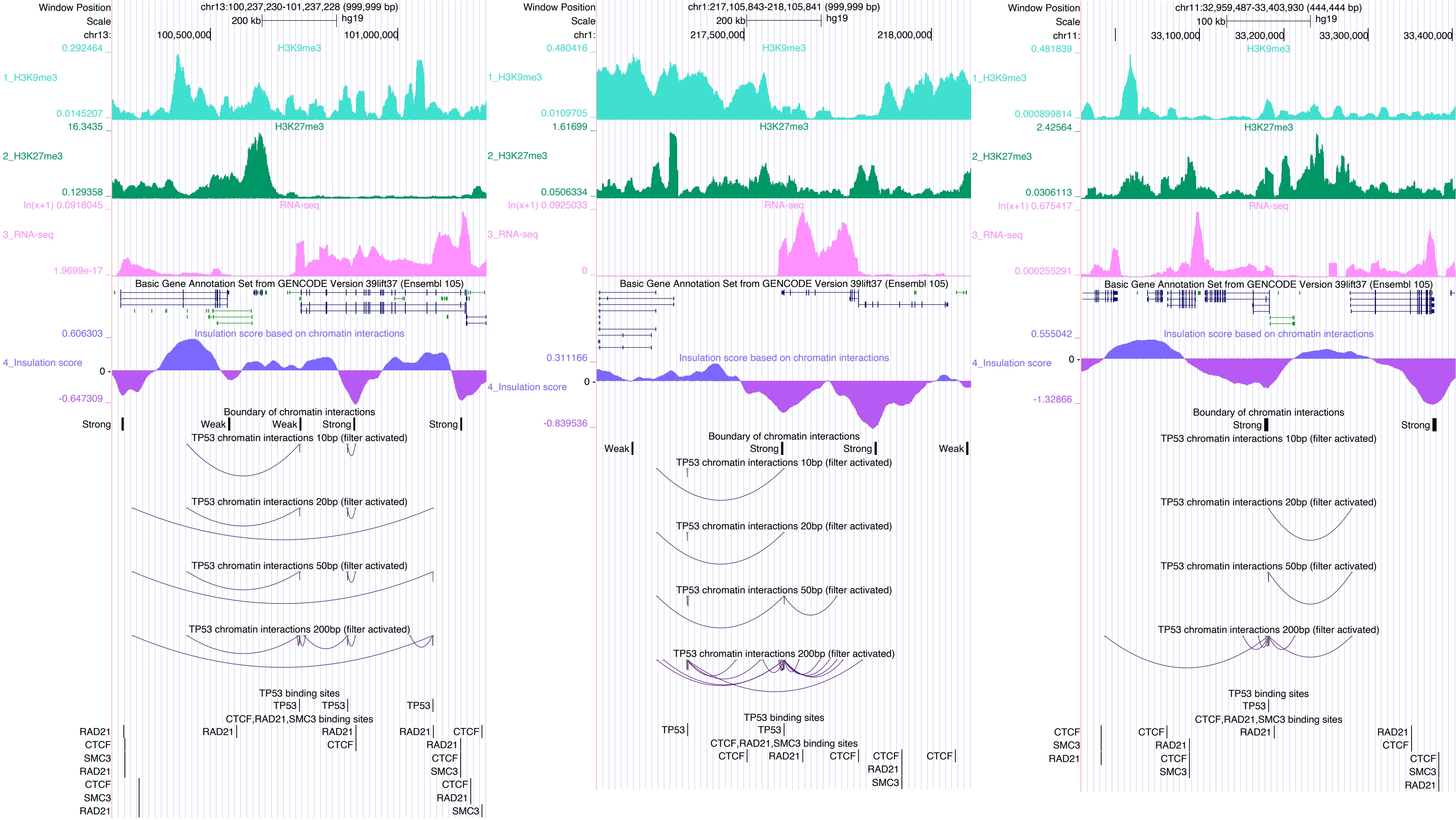
Window Position
chr13:100,237,230-101,237,228 (999,999 bp)
Scale
200 kb
hg19
chr13:
100,500,000
101,000,000
H3K9me3
1_H3K9me3
H3K27me3
2_H3K27me3
RNA-seq
3_RNA-seq
Basic Gene Annotation Set from GENCODE Version 39lift37 (Ensembl 105)
Insulation score based on chromatin interactions
4_Insulation score
Boundary of chromatin interactions
Strong
Weak
TP53 chromatin interactions 10bp (filter activated)
TP53 chromatin interactions 20bp (filter activated)
TP53 chromatin interactions 50bp (filter activated)
TP53 chromatin interactions 200bp (filter activated)
TP53 binding sites
TP53
CTCF,RAD21,SMC3 binding sites
RAD21
CTCF
SMC3
Window Position
chr1:217,105,843-218,105,841 (999,999 bp)
Scale
200 kb
hg19
chr1:
217,500,000
218,000,000
Window Position
chr11:32,959,487-33,403,930 (444,444 bp)
Scale
100 kb
hg19
chr11:
33,100,000
33,200,000
33,300,000
33,400,000

### TRIM28

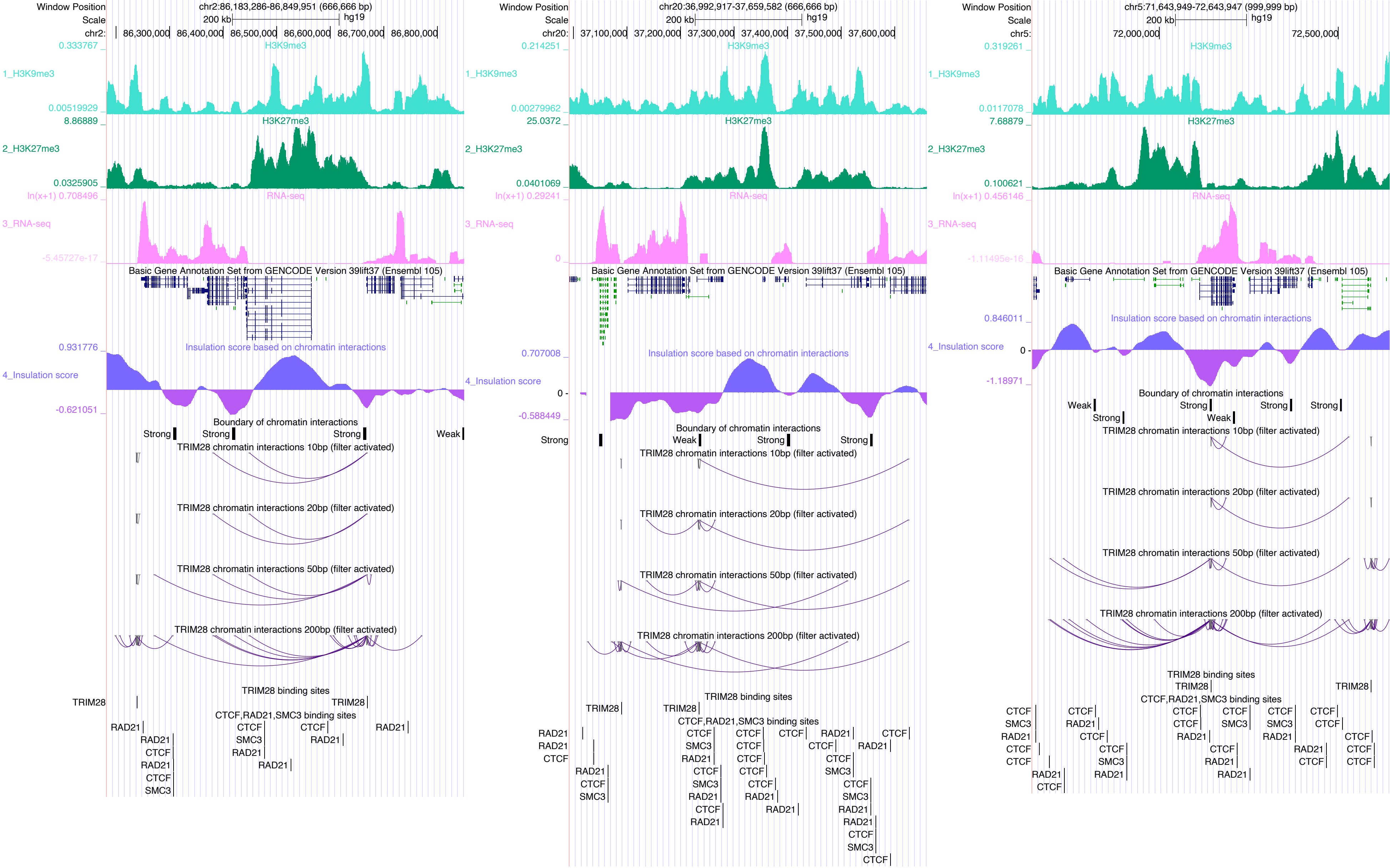

Window Position
Scale
chr2:86,183,286-86,849,951 (666,666 bp)
200 kb
hg19
chr2:
86,300,000
86,400,000
86,500,000
86,600,000
86,700,000
86,800,000
0.333767
H3K9me3
1_H3K9me3
0.00519929
8.86889
H3K27me3
2_H3K27me3
0.0325905
ln(x+1) 0.708496
RNA-seq
3_RNA-seq
-5.45727e-17
Basic Gene Annotation Set from GENCODE Version 39lift37 (Ensembl 105)
0.931776
Insulation score based on chromatin interactions
4_Insulation score
-0.621051
Boundary of chromatin interactions
Strong
Strong
Strong
Weak
TRIM28 chromatin interactions 10bp (filter activated)
TRIM28 chromatin interactions 20bp (filter activated)
TRIM28 chromatin interactions 50bp (filter activated)
TRIM28 chromatin interactions 200bp (filter activated)
TRIM28 binding sites
TRIM28
TRIM28
CTCF,RAD21,SMC3 binding sites
RAD21
RAD21
CTCF
RAD21
CTCF
SMC3
CTCF
SMC3
RAD21
RAD21
CTCF
RAD21
RAD21
chr20:36,992,917-37,659,582 (666,666 bp)
200 kb
hg19
chr20:
37,100,000
37,200,000
37,300,000
37,400,000
37,500,000
37,600,000
0.214251
H3K9me3
1_H3K9me3
0.00279962
25.0372
H3K27me3
2_H3K27me3
0.0401069
ln(x+1) 0.29241
RNA-seq
3_RNA-seq
0
Basic Gene Annotation Set from GENCODE Version 39lift37 (Ensembl 105)
0.707008
Insulation score based on chromatin interactions
4_Insulation score
0
-0.588449
Boundary of chromatin interactions
Strong
Weak
Strong
Strong
TRIM28 chromatin interactions 10bp (filter activated)
TRIM28 chromatin interactions 20bp (filter activated)
TRIM28 chromatin interactions 50bp (filter activated)
TRIM28 chromatin interactions 200bp (filter activated)
TRIM28 binding sites
TRIM28
TRIM28
CTCF,RAD21,SMC3 binding sites
RAD21
RAD21
CTCF
RAD21
CTCF
SMC3
chr5:71,643,949-72,643,947 (999,999 bp)
200 kb
hg19
chr5:
72,000,000
72,500,000
0.319261
H3K9me3
1_H3K9me3
0.0117078
7.68879
H3K27me3
2_H3K27me3
0.100621
ln(x+1) 0.456146
RNA-seq
3_RNA-seq
-1.11495e-16
Basic Gene Annotation Set from GENCODE Version 39lift37 (Ensembl 105)
0.846011
Insulation score based on chromatin interactions
4_Insulation score
0
-1.18971
Boundary of chromatin interactions
Weak
Strong
Strong
Strong
Strong
Weak
TRIM28 chromatin interactions 10bp (filter activated)
TRIM28 chromatin interactions 20bp (filter activated)
TRIM28 chromatin interactions 50bp (filter activated)
TRIM28 chromatin interactions 200bp (filter activated)
TRIM28 binding sites
TRIM28
TRIM28
CTCF,RAD21,SMC3 binding sites
CTCF
SMC3
RAD21
CTCF
CTCF

### USF1

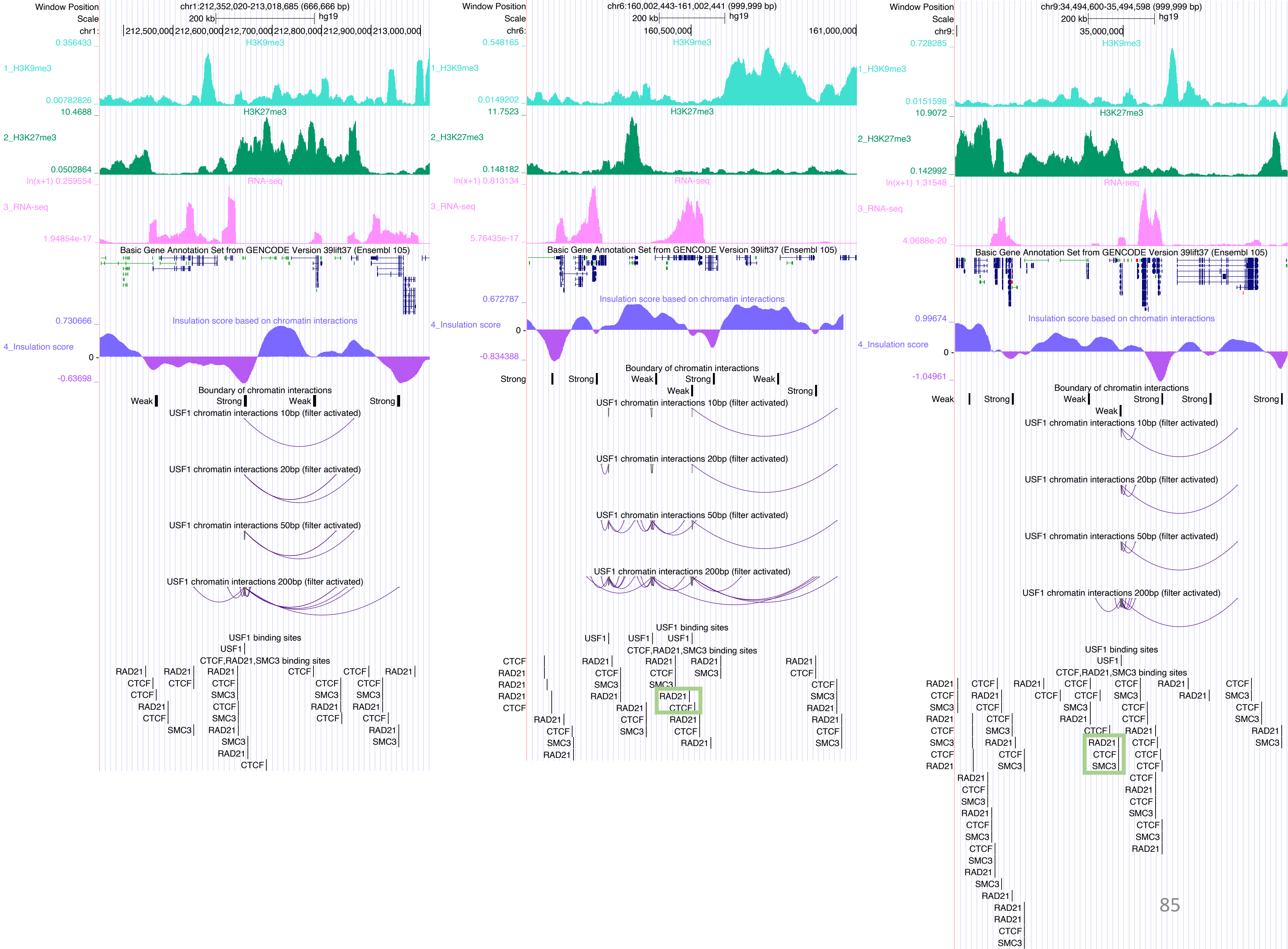
Window Position
chr1:212,352,020-213,018,685 (666,666 bp)
Scale
200 kb
hg19
chr1:
212,500,000
212,600,000
212,700,000
212,800,000
212,900,000
213,000,000
H3K9me3
1_H3K9me3
0.356433
0.00782826
H3K27me3
2_H3K27me3
10.4688
0.0502864
RNA-seq
3_RNA-seq
ln(x+1) 0.259554
1.94854e-17
Basic Gene Annotation Set from GENCODE Version 39lift37 (Ensembl 105)
Insulation score based on chromatin interactions
4_Insulation score
0.730666
-0.63698
Boundary of chromatin interactions
Weak
Strong
USF1 chromatin interactions 10bp (filter activated)
USF1 chromatin interactions 20bp (filter activated)
USF1 chromatin interactions 50bp (filter activated)
USF1 chromatin interactions 200bp (filter activated)
USF1 binding sites
CTCF,RAD21,SMC3 binding sites
Window Position
chr6:160,002,443-161,002,441 (999,999 bp)
chr6:
160,500,000
161,000,000
0.548165
0.0149202
11.7523
0.148182
ln(x+1) 0.813134
5.76435e-17
0.672787
-0.834388
Window Position
chr9:34,494,600-35,494,598 (999,999 bp)
chr9:
35,000,000
0.728285
0.0151598
10.9072
0.142992
ln(x+1) 1.31548
4.0688e-20
0.99674
-1.04961
RAD21
CTCF
SMC3
USF1

### VDR

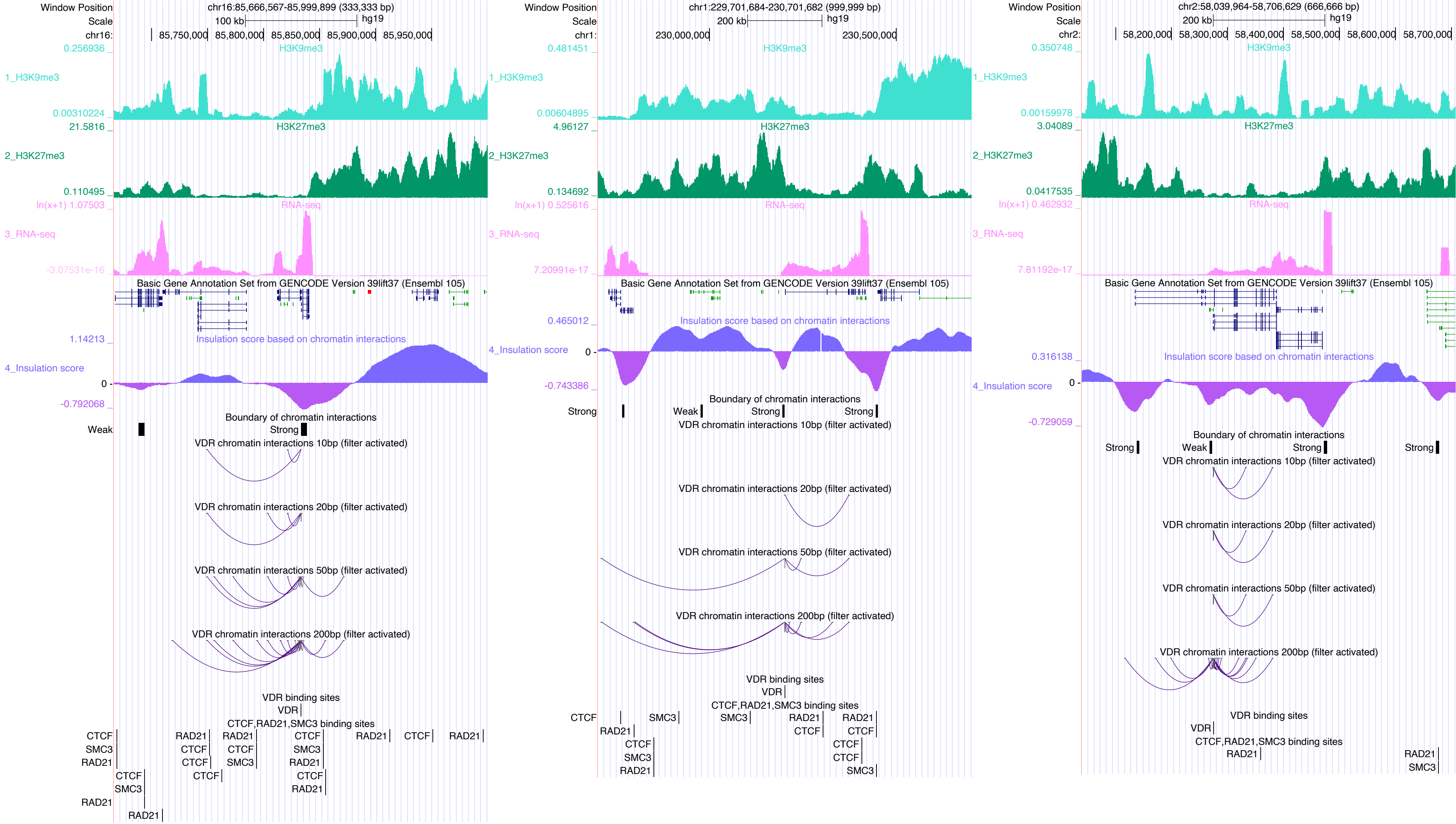

Window Position
chr16:85,666,567-85,999,899 (333,333 bp)
Scale
100 kb
hg19
chr16:
85,750,000
85,800,000
85,850,000
85,900,000
85,950,000
H3K9me3
1_H3K9me3
H3K27me3
2_H3K27me3
RNA-seq
3_RNA-seq
Basic Gene Annotation Set from GENCODE Version 39lift37 (Ensembl 105)
Insulation score based on chromatin interactions
4_Insulation score
Boundary of chromatin interactions
Weak
Strong
VDR chromatin interactions 10bp (filter activated)
VDR chromatin interactions 20bp (filter activated)
VDR chromatin interactions 50bp (filter activated)
VDR chromatin interactions 200bp (filter activated)
VDR binding sites
VDR
CTCF,RAD21,SMC3 binding sites
CTCF
SMC3
RAD21
Window Position
chr1:229,701,684-230,701,682 (999,999 bp)
200 kb
chr1:
230,000,000
230,500,000
Window Position
chr2:58,039,964-58,706,629 (666,666 bp)
chr2:
58,200,000
58,300,000
58,400,000
58,500,000
58,600,000
58,700,000

### ZBTB7A

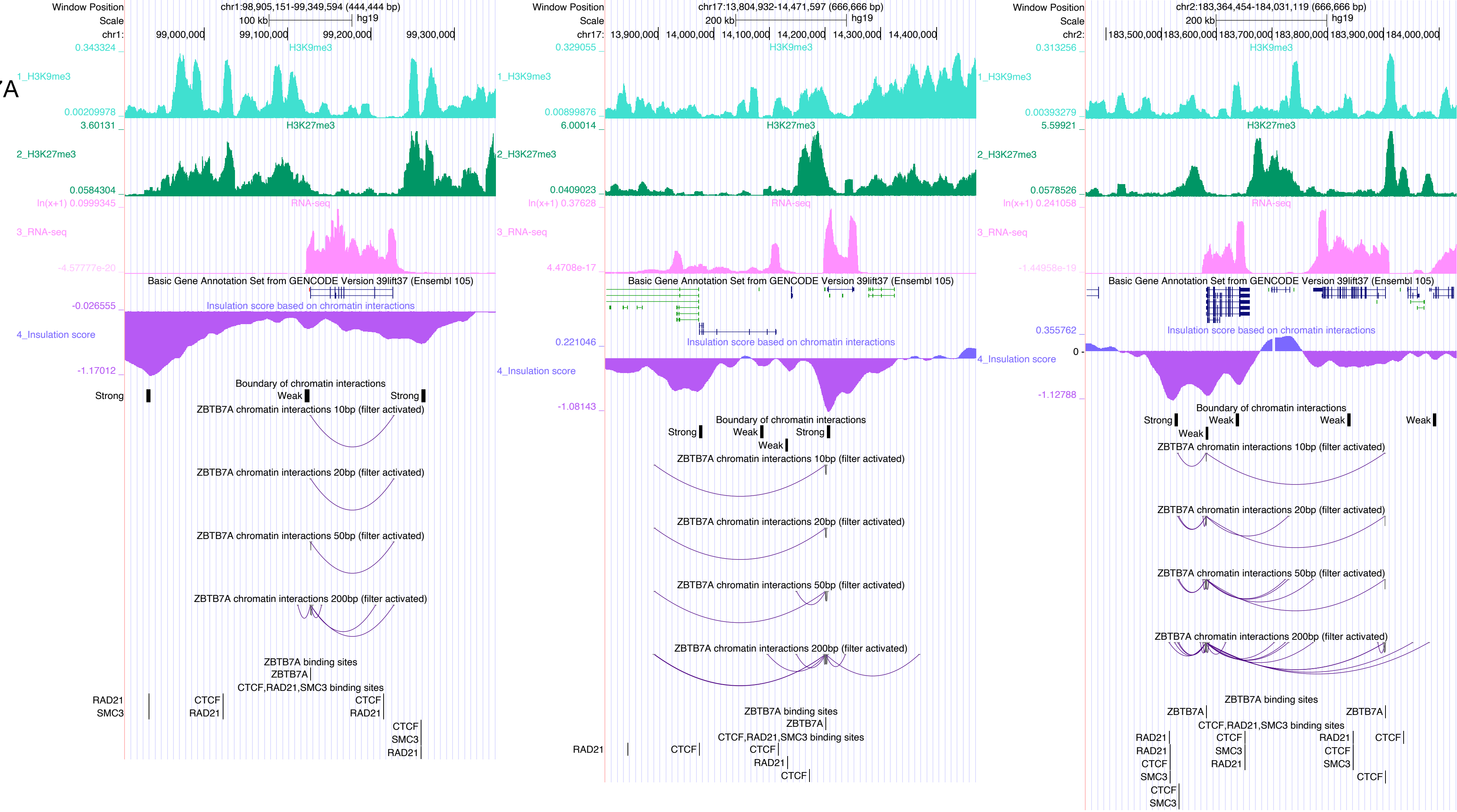

Window Position chr1:98,905,151-99,349,594 (444,444 bp)
Scale 100 kb hg19
chr1: 99,000,000 99,100,000 99,200,000 99,300,000
1_H3K9me3
2_H3K27me3
3_RNA-seq
Basic Gene Annotation Set from GENCODE Version 39lift37 (Ensembl 105)
4_Insulation score
Insulation score based on chromatin interactions
Boundary of chromatin interactions
Strong
Weak
ZBTB7A chromatin interactions 10bp (filter activated)
ZBTB7A chromatin interactions 20bp (filter activated)
ZBTB7A chromatin interactions 50bp (filter activated)
ZBTB7A chromatin interactions 200bp (filter activated)
ZBTB7A binding sites
CTCF,RAD21,SMC3 binding sites
Window Position chr17:13,804,932-14,471,597 (666,666 bp)
Scale 200 kb hg19
chr17: 13,900,000 14,000,000 14,100,000 14,200,000 14,300,000 14,400,000
Window Position chr2:183,364,454-184,031,119 (666,666 bp)
Scale 200 kb hg19
chr2: 183,500,000 183,600,000 183,700,000 183,800,000 183,900,000 184,000,000

### ZEB1

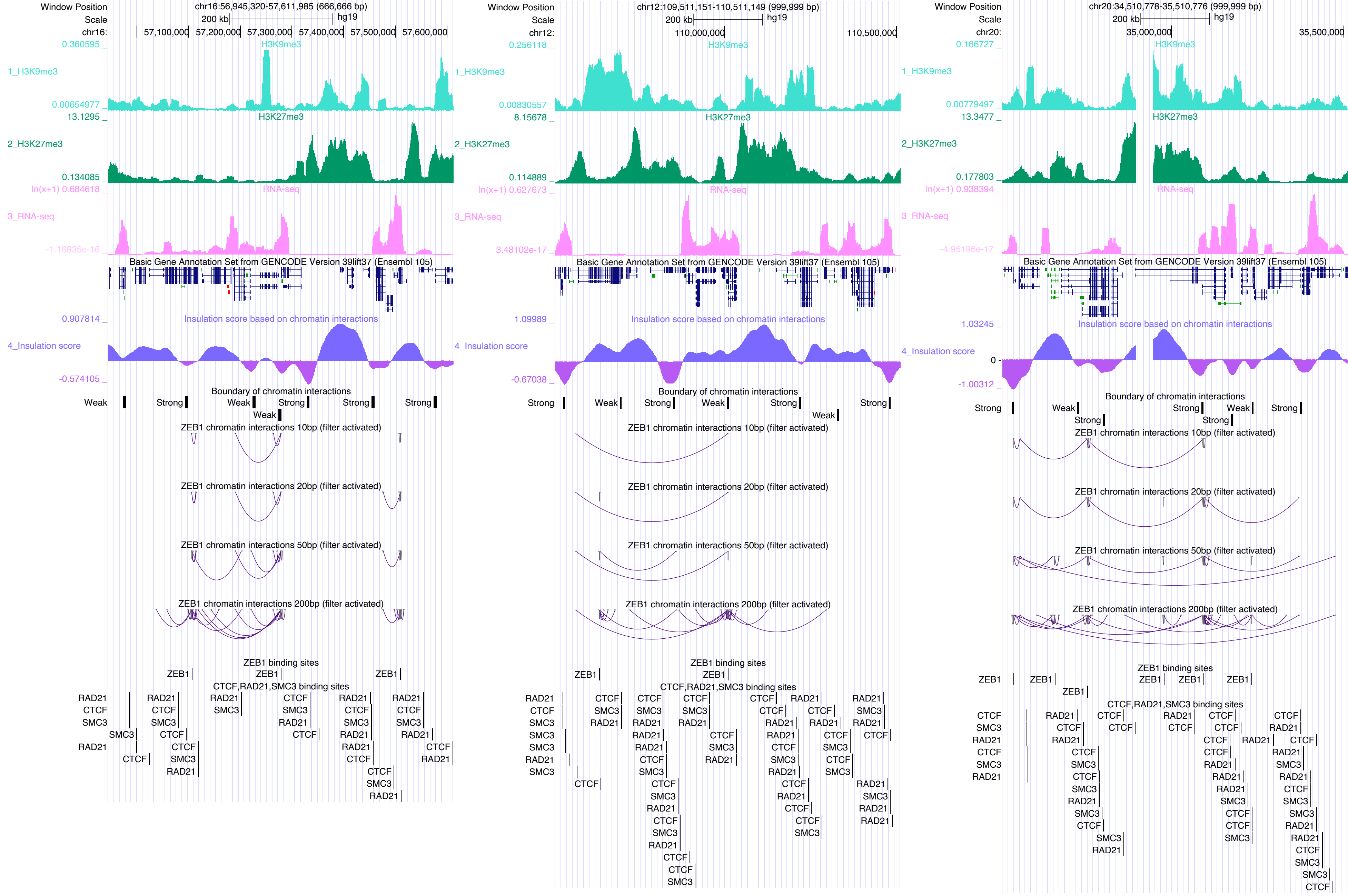

Window Position chr16:56,945,320-57,611,985 (666,666 bp)
Scale 200 kb hg19
H3K9me3
1_H3K9me3
H3K27me3
2_H3K27me3
RNA-seq
3_RNA-seq
Basic Gene Annotation Set from GENCODE Version 39lift37 (Ensembl 105)
Insulation score based on chromatin interactions
4_Insulation score
Boundary of chromatin interactions
ZEB1 chromatin interactions 10bp (filter activated)
ZEB1 chromatin interactions 20bp (filter activated)
ZEB1 chromatin interactions 50bp (filter activated)
ZEB1 chromatin interactions 200bp (filter activated)
ZEB1 binding sites
CTCF,RAD21,SMC3 binding sites
Window Position chr12:109,511,151-110,511,149 (999,999 bp)
Window Position chr20:34,510,778-35,510,776 (999,999 bp)

#### ZFP3

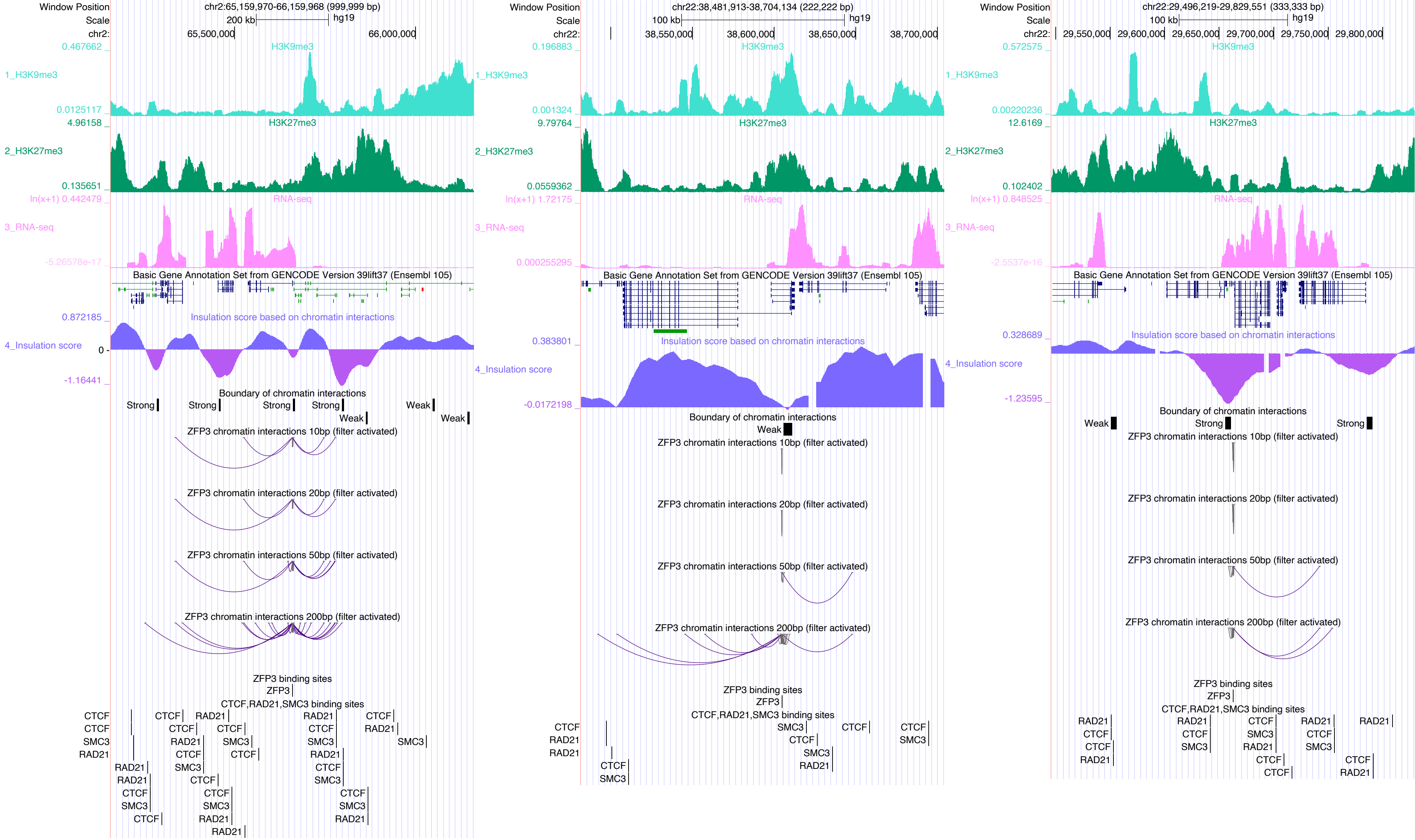

Window Position
chr2:65,159,970-66,159,968 (999,999 bp)
Scale
200 kb
hg19
chr2:
65,500,000
66,000,000
0.467662
H3K9me3
1_H3K9me3
0.0125117
4.96158
H3K27me3
2_H3K27me3
0.135651
ln(x+1) 0.442479
RNA-seq
3_RNA-seq
-5.26578e-17
Basic Gene Annotation Set from GENCODE Version 39lift37 (Ensembl 105)
0.872185
Insulation score based on chromatin interactions
4_Insulation score
0
-1.16441
Boundary of chromatin interactions
Strong
Weak
ZFP3 chromatin interactions 10bp (filter activated)
ZFP3 chromatin interactions 20bp (filter activated)
ZFP3 chromatin interactions 50bp (filter activated)
ZFP3 chromatin interactions 200bp (filter activated)
ZFP3 binding sites
ZFP3
CTCF,RAD21,SMC3 binding sites
CTCF
SMC3
RAD21
Window Position
chr22:38,481,913-38,704,134 (222,222 bp)
Scale
100 kb
hg19
chr22:
38,550,000
38,600,000
38,650,000
38,700,000
0.196883
H3K9me3
1_H3K9me3
0.001324
9.79764
H3K27me3
2_H3K27me3
0.0559362
ln(x+1) 1.72175
RNA-seq
3_RNA-seq
0.000255295
Basic Gene Annotation Set from GENCODE Version 39lift37 (Ensembl 105)
0.383801
Insulation score based on chromatin interactions
4_Insulation score
-0.0172198
Boundary of chromatin interactions
Weak
ZFP3 chromatin interactions 10bp (filter activated)
ZFP3 chromatin interactions 20bp (filter activated)
ZFP3 chromatin interactions 50bp (filter activated)
ZFP3 chromatin interactions 200bp (filter activated)
ZFP3 binding sites
ZFP3
CTCF,RAD21,SMC3 binding sites
CTCF
RAD21
SMC3
Window Position
chr22:29,496,219-29,829,551 (333,333 bp)
Scale
100 kb
hg19
chr22:
29,550,000
29,600,000
29,650,000
29,700,000
29,750,000
29,800,000
0.572575
H3K9me3
1_H3K9me3
0.00220236
12.6169
H3K27me3
2_H3K27me3
0.102402
ln(x+1) 0.848525
RNA-seq
3_RNA-seq
-2.5537e-16
Basic Gene Annotation Set from GENCODE Version 39lift37 (Ensembl 105)
0.328689
Insulation score based on chromatin interactions
4_Insulation score
-1.23595
Boundary of chromatin interactions
Weak
Strong
ZFP3 chromatin interactions 10bp (filter activated)
ZFP3 chromatin interactions 20bp (filter activated)
ZFP3 chromatin interactions 50bp (filter activated)
ZFP3 chromatin interactions 200bp (filter activated)
ZFP3 binding sites
ZFP3
CTCF,RAD21,SMC3 binding sites
RAD21
CTCF
SMC3

ZIC2

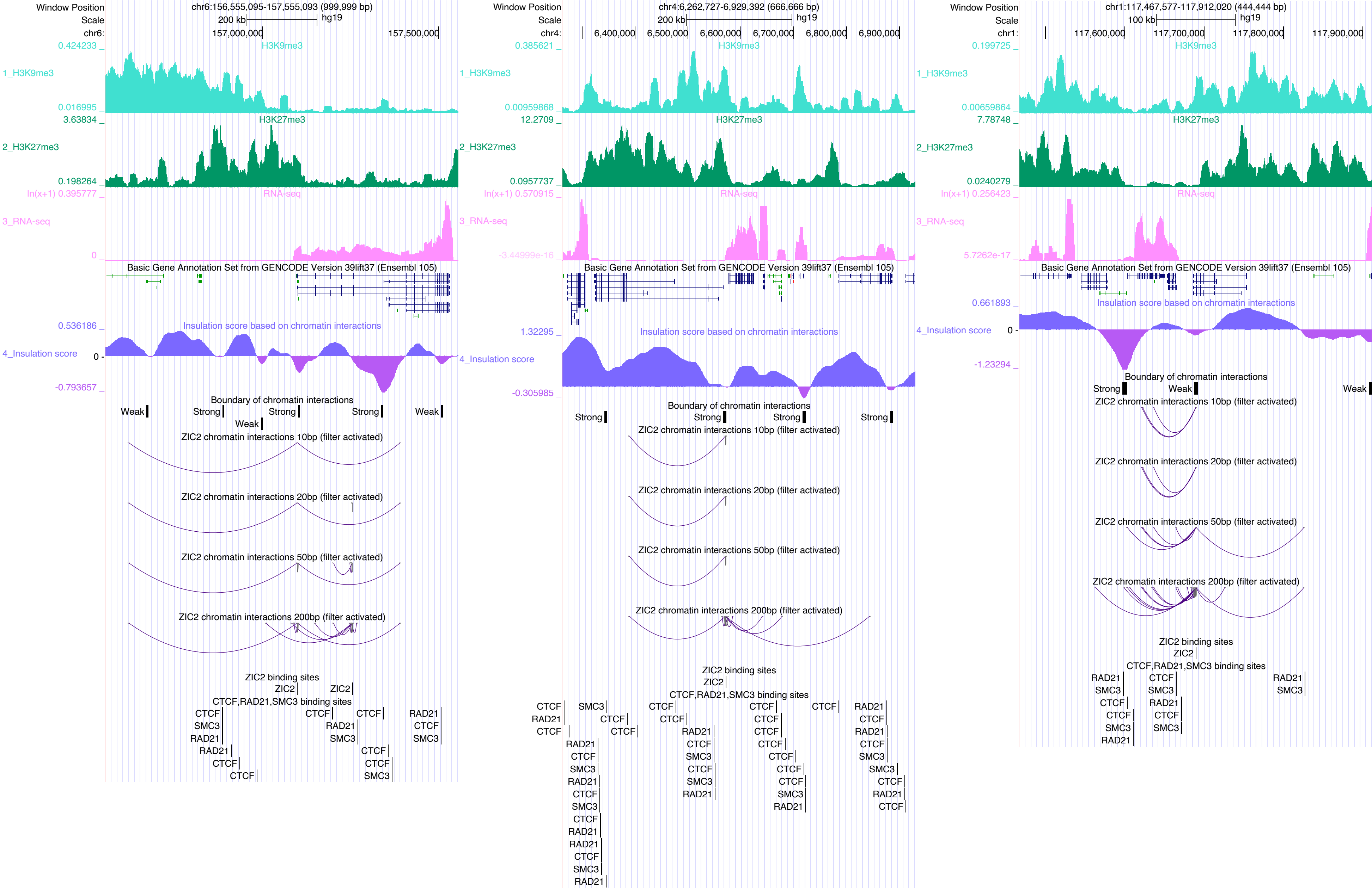

Window Position
chr6:156,555,095-157,555,093 (999,999 bp)
Scale
200 kb
hg19
chr6:
157,000,000
157,500,000
0.424233
H3K9me3
1_H3K9me3
0.016995
3.63834
H3K27me3
2_H3K27me3
0.198264
ln(x+1) 0.395777
RNA-seq
3_RNA-seq
0
Basic Gene Annotation Set from GENCODE Version 39lift37 (Ensembl 105)
0.536186
Insulation score based on chromatin interactions
4_Insulation score
0
-0.793657
Boundary of chromatin interactions
Weak
Strong
Strong
Strong
Weak
Weak
ZIC2 chromatin interactions 10bp (filter activated)
ZIC2 chromatin interactions 20bp (filter activated)
ZIC2 chromatin interactions 50bp (filter activated)
ZIC2 chromatin interactions 200bp (filter activated)
ZIC2 binding sites
ZIC2
ZIC2
CTCF,RAD21,SMC3 binding sites
CTCF
SMC3
RAD21
RAD21
CTCF
CTCF
CTCF
RAD21
SMC3
CTCF
CTCF
SMC3
RAD21
CTCF
SMC3
Window Position
chr4:6,262,727-6,929,392 (666,666 bp)
Scale
200 kb
hg19
chr4:
6,400,000
6,500,000
6,600,000
6,700,000
6,800,000
6,900,000
0.385621
H3K9me3
1_H3K9me3
0.00959868
12.2709
H3K27me3
2_H3K27me3
0.0957737
ln(x+1) 0.570915
RNA-seq
3_RNA-seq
-3.44999e-16
Basic Gene Annotation Set from GENCODE Version 39lift37 (Ensembl 105)
1.32295
Insulation score based on chromatin interactions
4_Insulation score
-0.305985
Boundary of chromatin interactions
Strong
Strong
Strong
Strong
ZIC2 chromatin interactions 10bp (filter activated)
ZIC2 chromatin interactions 20bp (filter activated)
ZIC2 chromatin interactions 50bp (filter activated)
ZIC2 chromatin interactions 200bp (filter activated)
ZIC2 binding sites
ZIC2
CTCF,RAD21,SMC3 binding sites
CTCF
RAD21
CTCF
SMC3
CTCF
CTCF
CTCF
CTCF
CTCF
CTCF
RAD21
CTCF
SMC3
RAD21
CTCF
SMC3
RAD21
CTCF
SMC3
CTCF
RAD21
RAD21
CTCF
SMC3
CTCF
SMC3
RAD21
CTCF
CTCF
SMC3
RAD21
CTCF
CTCF
RAD21
CTCF
CTCF
CTCF
SMC3
CTCF
SMC3
CTCF
SMC3
CTCF
RAD21
RAD21
RAD21
SMC3
CTCF
RAD21
CTCF
SMC3
RAD21
Window Position
chr1:117,467,577-117,912,020 (444,444 bp)
Scale
100 kb
hg19
chr1:
117,600,000
117,700,000
117,800,000
117,900,000
0.199725
H3K9me3
1_H3K9me3
0.00659864
7.78748
H3K27me3
2_H3K27me3
0.0240279
ln(x+1) 0.256423
RNA-seq
3_RNA-seq
5.7262e-17
Basic Gene Annotation Set from GENCODE Version 39lift37 (Ensembl 105)
0.661893
Insulation score based on chromatin interactions
4_Insulation score
0
-1.23294
Boundary of chromatin interactions
Strong
Weak
Weak
ZIC2 chromatin interactions 10bp (filter activated)
ZIC2 chromatin interactions 20bp (filter activated)
ZIC2 chromatin interactions 50bp (filter activated)
ZIC2 chromatin interactions 200bp (filter activated)
ZIC2 binding sites
ZIC2
CTCF,RAD21,SMC3 binding sites
RAD21
SMC3
CTCF
CTCF
SMC3
RAD21
CTCF
SMC3
RAD21
CTCF
SMC3
RAD21
SMC3

### ZNF18

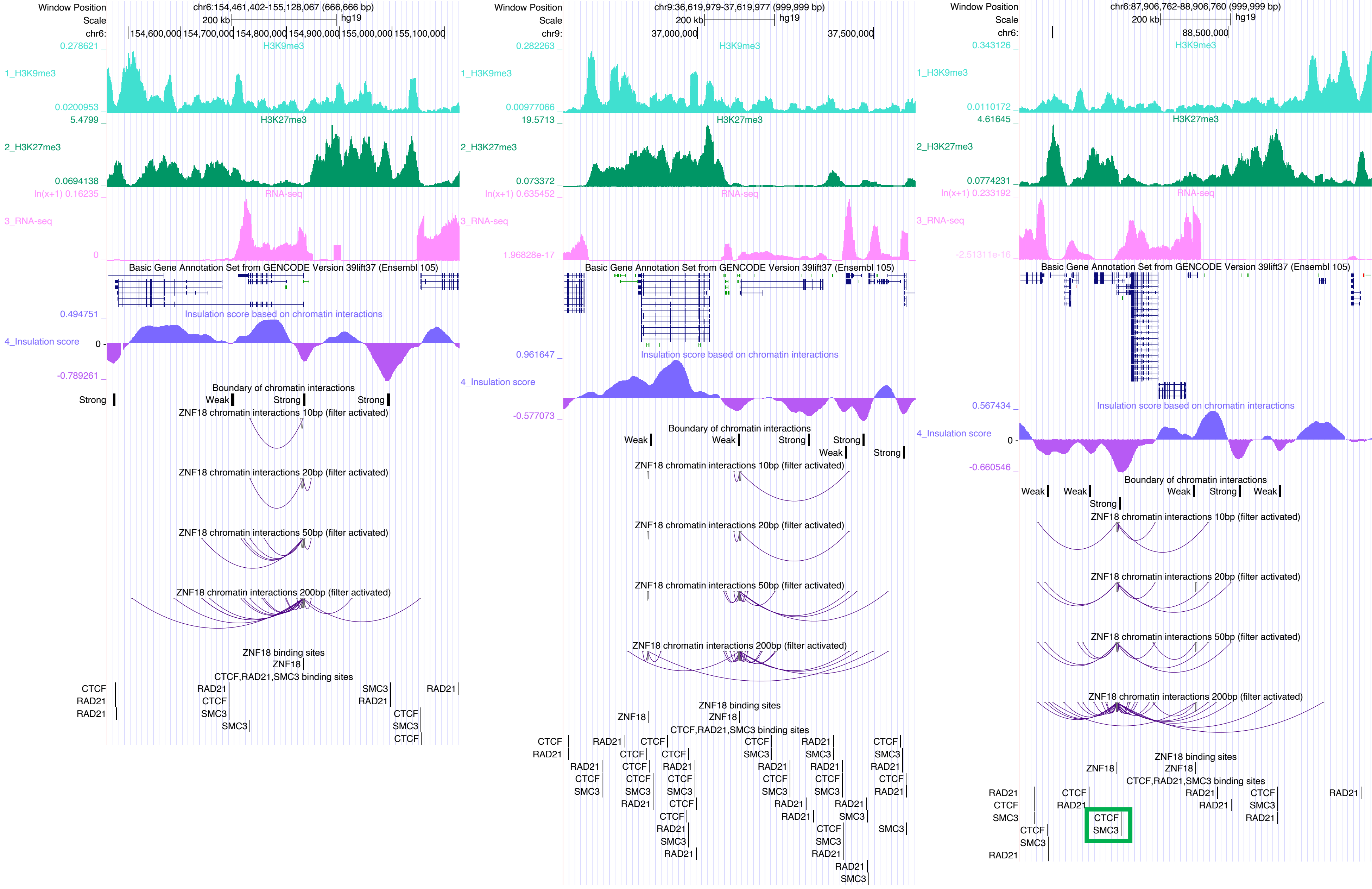
Window Position
chr6:154,461,402-155,128,067 (666,666 bp)
Scale
200 kb
hg19
1_H3K9me3
2_H3K27me3
3_RNA-seq
4_Insulation score
Basic Gene Annotation Set from GENCODE Version 39lift37 (Ensembl 105)
Insulation score based on chromatin interactions
Boundary of chromatin interactions
ZNF18 chromatin interactions 10bp (filter activated)
ZNF18 chromatin interactions 20bp (filter activated)
ZNF18 chromatin interactions 50bp (filter activated)
ZNF18 chromatin interactions 200bp (filter activated)
ZNF18 binding sites
CTCF,RAD21,SMC3 binding sites
chr9:36,619,979-37,619,977 (999,999 bp)
chr6:87,906,762-88,906,760 (999,999 bp)

ZNF35

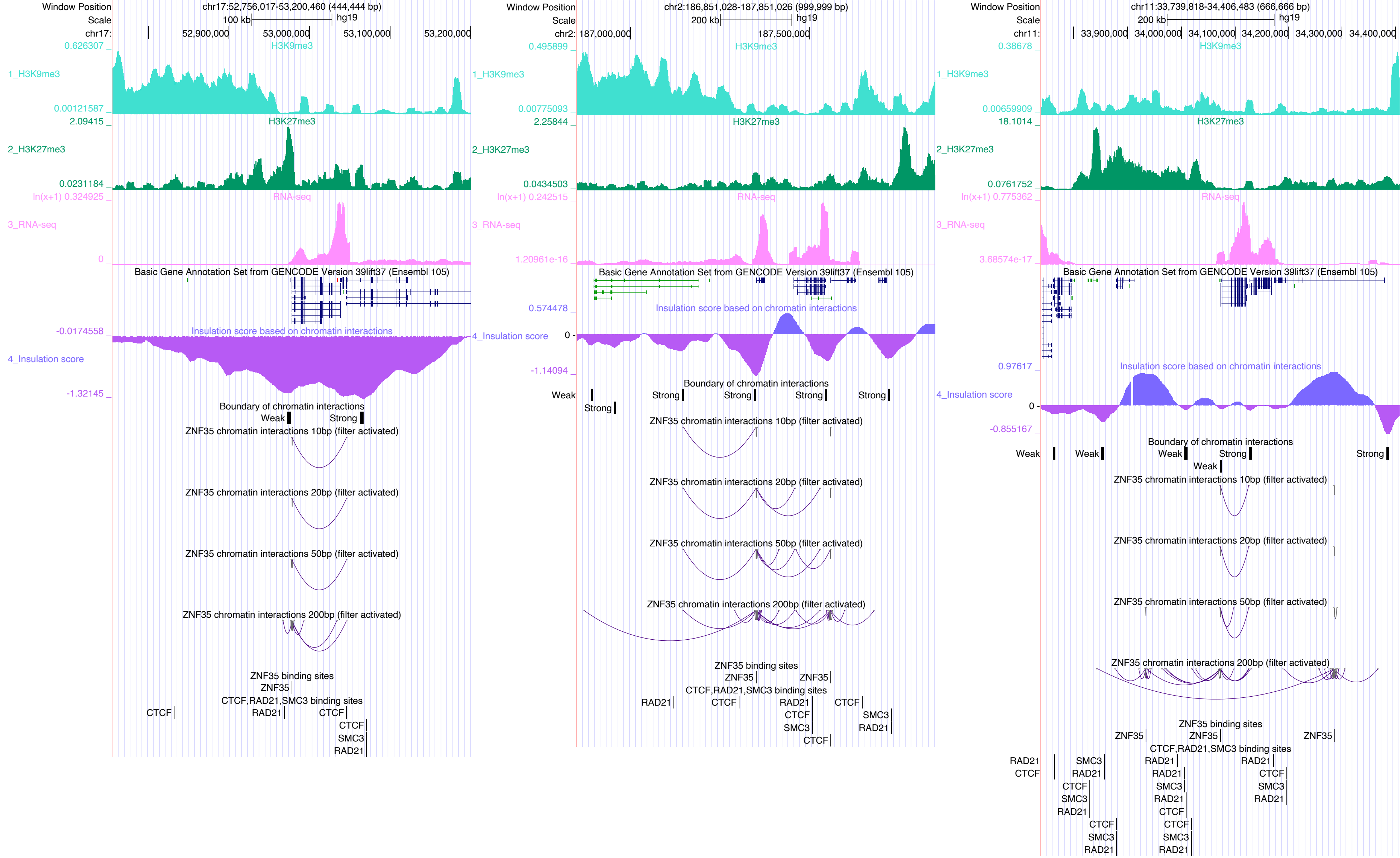

Window Position
chr17:52,756,017-53,200,460 (444,444 bp)
Scale
100 kb
hg19
chr17:
52,900,000
53,000,000
53,100,000
53,200,000
H3K9me3
1_H3K9me3
0.626307
0.00121587
H3K27me3
2_H3K27me3
2.09415
0.0231184
RNA-seq
3_RNA-seq
ln(x+1) 0.324925
0
Basic Gene Annotation Set from GENCODE Version 39lift37 (Ensembl 105)
Insulation score based on chromatin interactions
4_Insulation score
-0.0174558
-1.32145
Boundary of chromatin interactions
Weak
Strong
ZNF35 chromatin interactions 10bp (filter activated)
ZNF35 chromatin interactions 20bp (filter activated)
ZNF35 chromatin interactions 50bp (filter activated)
ZNF35 chromatin interactions 200bp (filter activated)
ZNF35 binding sites
ZNF35
CTCF,RAD21,SMC3 binding sites
CTCF
RAD21
CTCF
CTCF
SMC3
RAD21
Window Position
chr2:186,851,028-187,851,026 (999,999 bp)
Scale
200 kb
hg19
chr2:
187,000,000
187,500,000
H3K9me3
1_H3K9me3
0.495899
0.00775093
H3K27me3
2_H3K27me3
2.25844
0.0434503
RNA-seq
3_RNA-seq
ln(x+1) 0.242515
1.20961e-16
Basic Gene Annotation Set from GENCODE Version 39lift37 (Ensembl 105)
Insulation score based on chromatin interactions
4_Insulation score
0.574478
0
-1.14094
Boundary of chromatin interactions
Weak
Strong
Strong
Strong
Strong
Strong
ZNF35 chromatin interactions 10bp (filter activated)
ZNF35 chromatin interactions 20bp (filter activated)
ZNF35 chromatin interactions 50bp (filter activated)
ZNF35 chromatin interactions 200bp (filter activated)
ZNF35 binding sites
ZNF35
ZNF35
CTCF,RAD21,SMC3 binding sites
RAD21
CTCF
RAD21
CTCF
SMC3
CTCF
CTCF
SMC3
RAD21
Window Position
chr11:33,739,818-34,406,483 (666,666 bp)
Scale
200 kb
hg19
chr11:
33,900,000
34,000,000
34,100,000
34,200,000
34,300,000
34,400,000
H3K9me3
1_H3K9me3
0.38678
0.00659909
H3K27me3
2_H3K27me3
18.1014
0.0761752
RNA-seq
3_RNA-seq
ln(x+1) 0.775362
3.68574e-17
Basic Gene Annotation Set from GENCODE Version 39lift37 (Ensembl 105)
Insulation score based on chromatin interactions
4_Insulation score
0.97617
0
-0.855167
Boundary of chromatin interactions
Weak
Weak
Weak
Strong
Weak
Strong
ZNF35 chromatin interactions 10bp (filter activated)
ZNF35 chromatin interactions 20bp (filter activated)
ZNF35 chromatin interactions 50bp (filter activated)
ZNF35 chromatin interactions 200bp (filter activated)
ZNF35 binding sites
ZNF35
ZNF35
ZNF35
CTCF,RAD21,SMC3 binding sites
RAD21
CTCF
SMC3
RAD21
CTCF
SMC3
RAD21
CTCF
SMC3
RAD21
RAD21
RAD21
SMC3
RAD21
CTCF
CTCF
SMC3
RAD21
RAD21
CTCF
SMC3
RAD21

### ZNF143

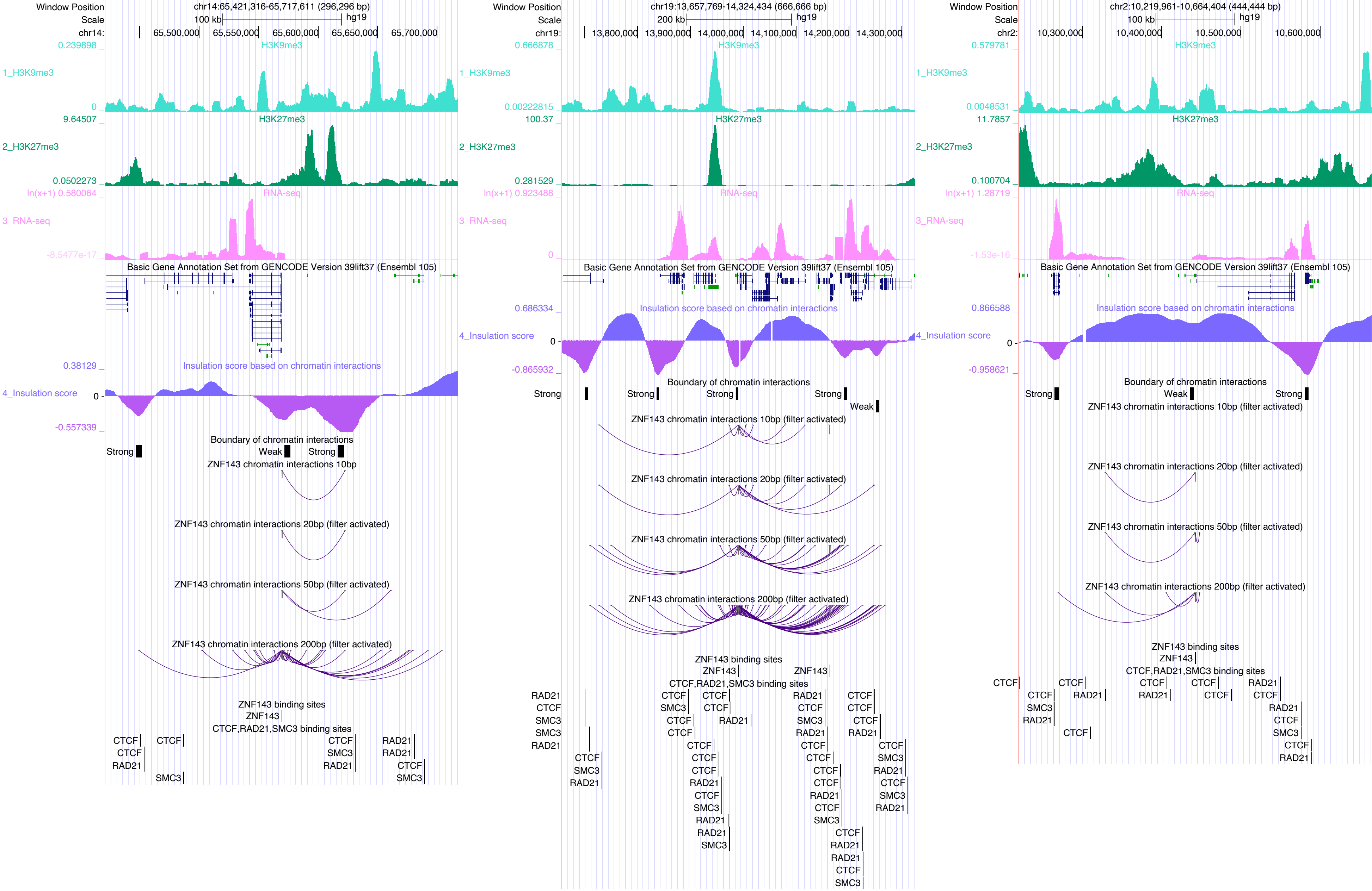

Window Position
Scale
chr14:65,421,316-65,717,611 (296,296 bp)
100 kb
hg19
chr14:
65,500,000
65,550,000
65,600,000
65,650,000
65,700,000
0.239898
H3K9me3
1_H3K9me3
0
9.64507
H3K27me3
2_H3K27me3
0.0502273
ln(x+1) 0.580064
RNA-seq
3_RNA-seq
-8.5477e-17
Basic Gene Annotation Set from GENCODE Version 39lift37 (Ensembl 105)
0.38129
Insulation score based on chromatin interactions
4_Insulation score
0
-0.557339
Boundary of chromatin interactions
Strong
Weak
Strong
ZNF143 chromatin interactions 10bp
ZNF143 chromatin interactions 20bp (filter activated)
ZNF143 chromatin interactions 50bp (filter activated)
ZNF143 chromatin interactions 200bp (filter activated)
ZNF143 binding sites
ZNF143
CTCF,RAD21,SMC3 binding sites
CTCF
CTCF
RAD21
CTCF
SMC3
CTCF
SMC3
RAD21
RAD21
RAD21
CTCF
SMC3
chr19:13,657,769-14,324,434 (666,666 bp)
200 kb
hg19
chr19:
13,800,000
13,900,000
14,000,000
14,100,000
14,200,000
14,300,000
0.666878
H3K9me3
1_H3K9me3
0.00222815
100.37
H3K27me3
2_H3K27me3
0.281529
ln(x+1) 0.923488
RNA-seq
3_RNA-seq
0
Basic Gene Annotation Set from GENCODE Version 39lift37 (Ensembl 105)
0.686334
Insulation score based on chromatin interactions
4_Insulation score
0
-0.865932
Boundary of chromatin interactions
Strong
Strong
Strong
Strong
Weak
ZNF143 chromatin interactions 10bp (filter activated)
ZNF143 chromatin interactions 20bp (filter activated)
ZNF143 chromatin interactions 50bp (filter activated)
ZNF143 chromatin interactions 200bp (filter activated)
ZNF143 binding sites
ZNF143
ZNF143
CTCF,RAD21,SMC3 binding sites
RAD21
CTCF
SMC3
SMC3
RAD21
CTCF
SMC3
RAD21
chr2:10,219,961-10,664,404 (444,444 bp)
100 kb
hg19
chr2:
10,300,000
10,400,000
10,500,000
10,600,000
0.579781
H3K9me3
1_H3K9me3
0.0048531
11.7857
H3K27me3
2_H3K27me3
0.100704
ln(x+1) 1.28719
RNA-seq
3_RNA-seq
-1.53e-16
Basic Gene Annotation Set from GENCODE Version 39lift37 (Ensembl 105)
0.866588
Insulation score based on chromatin interactions
4_Insulation score
0
-0.958621
Boundary of chromatin interactions
Strong
Weak
Strong
ZNF143 chromatin interactions 10bp (filter activated)
ZNF143 chromatin interactions 20bp (filter activated)
ZNF143 chromatin interactions 50bp (filter activated)
ZNF143 chromatin interactions 200bp (filter activated)
ZNF143 binding sites
ZNF143
CTCF,RAD21,SMC3 binding sites

### ZNF320

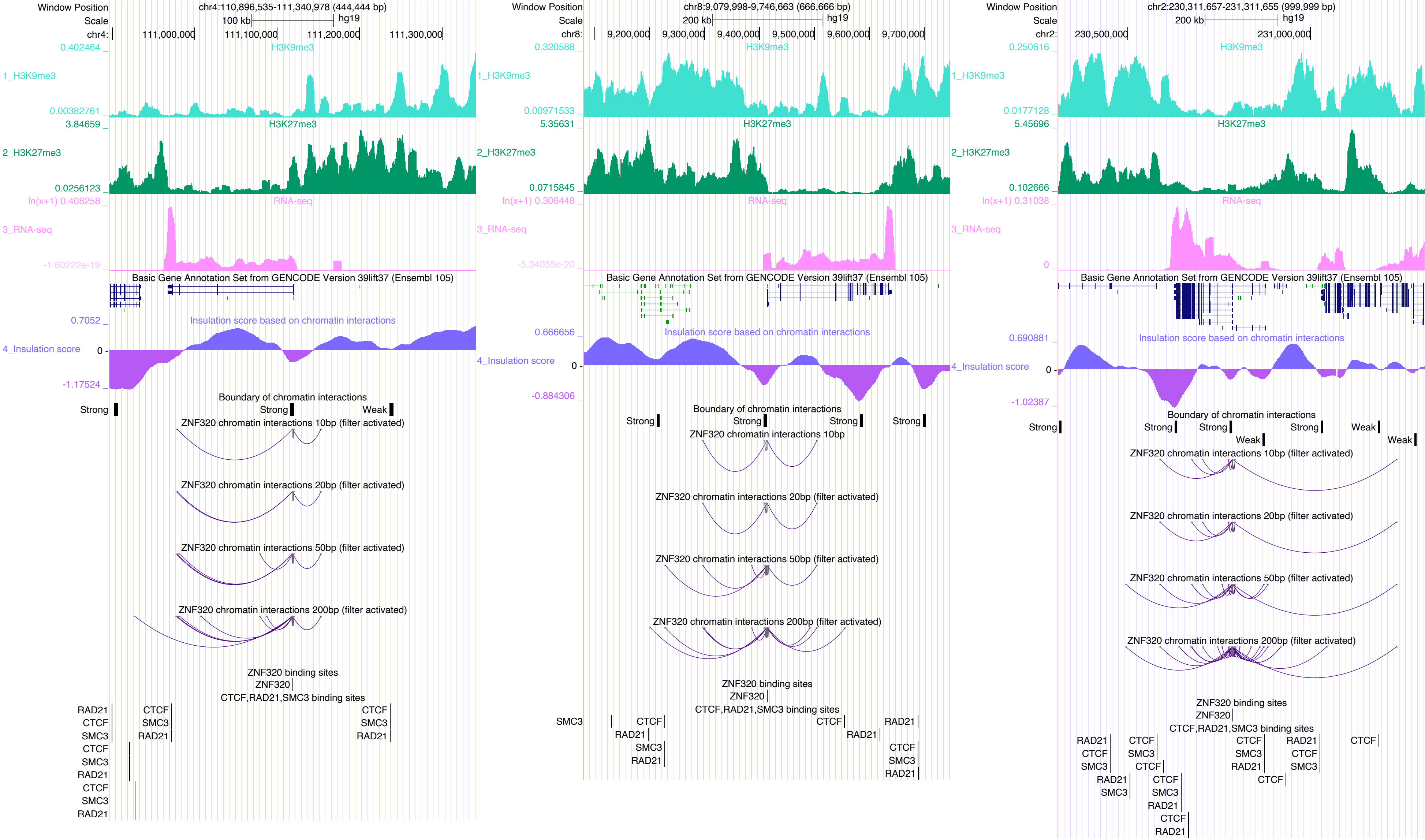

Window Position
Scale
chr4:110,896,535-111,340,978 (444,444 bp)
100 kb
hg19
111,000,000
111,100,000
111,200,000
111,300,000
1_H3K9me3
2_H3K27me3
3_RNA-seq
4_Insulation score
H3K9me3
H3K27me3
RNA-seq
Basic Gene Annotation Set from GENCODE Version 39lift37 (Ensembl 105)
Insulation score based on chromatin interactions
Boundary of chromatin interactions
Strong
Weak
ZNF320 chromatin interactions 10bp (filter activated)
ZNF320 chromatin interactions 20bp (filter activated)
ZNF320 chromatin interactions 50bp (filter activated)
ZNF320 chromatin interactions 200bp (filter activated)
ZNF320 binding sites
CTCF,RAD21,SMC3 binding sites
chr8:9,079,998-9,746,663 (666,666 bp)
200 kb
9,200,000
9,300,000
9,400,000
9,500,000
9,600,000
9,700,000
ZNF320 chromatin interactions 10bp
chr2:230,311,657-231,311,655 (999,999 bp)
230,500,000
231,000,000
CTCF
RAD21
SMC3

ZNF329

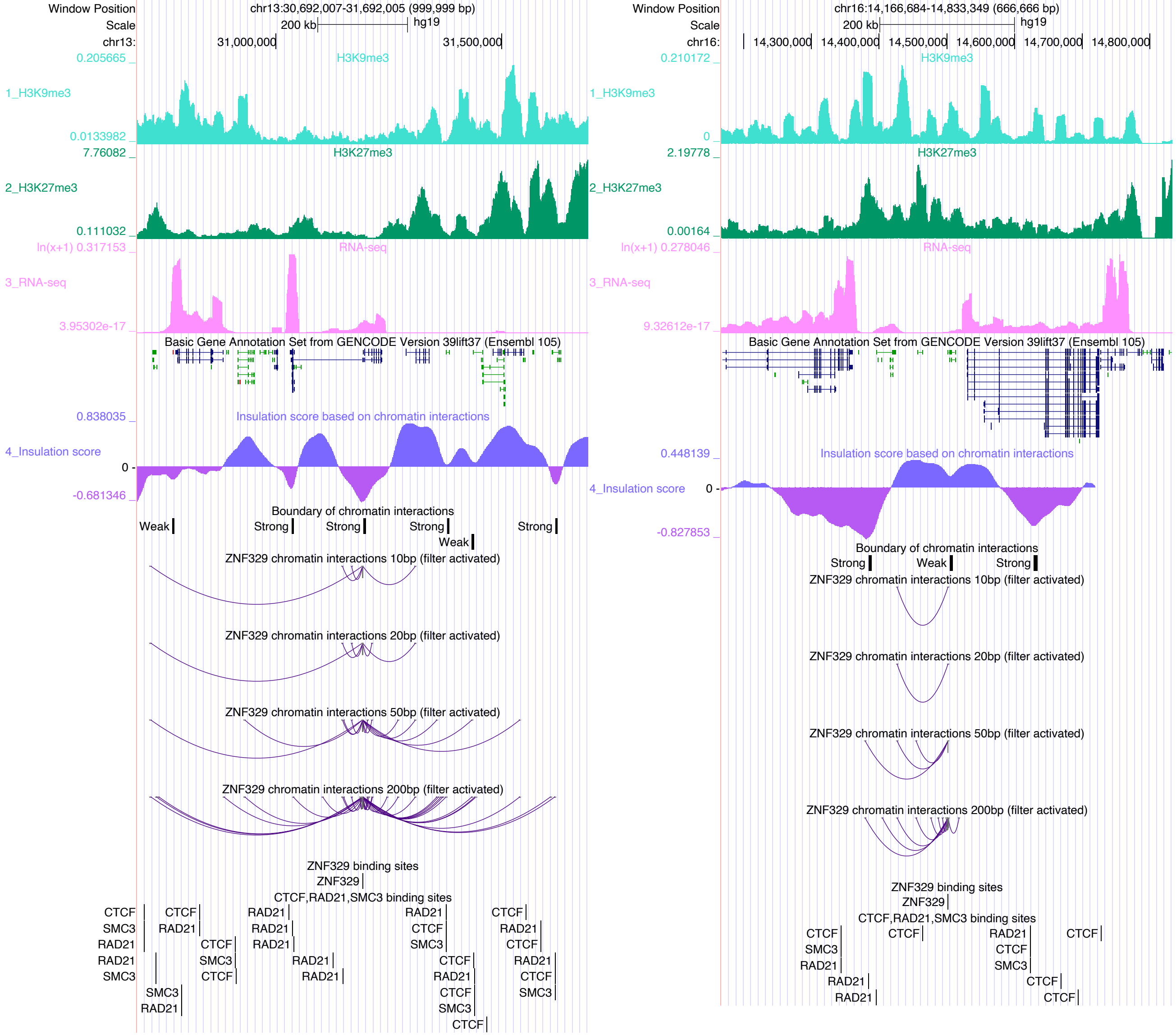

Window Position
chr13:30,692,007-31,692,005 (999,999 bp)
Scale
200 kb
hg19
chr13:
31,000,000
31,500,000
0.205665
H3K9me3
1_H3K9me3
0.0133982
7.76082
H3K27me3
2_H3K27me3
0.111032
ln(x+1) 0.317153
RNA-seq
3_RNA-seq
3.95302e-17
Basic Gene Annotation Set from GENCODE Version 39lift37 (Ensembl 105)
0.838035
Insulation score based on chromatin interactions
4_Insulation score
0
-0.681346
Boundary of chromatin interactions
Weak
Strong
Strong
Strong
Weak
Strong
ZNF329 chromatin interactions 10bp (filter activated)
ZNF329 chromatin interactions 20bp (filter activated)
ZNF329 chromatin interactions 50bp (filter activated)
ZNF329 chromatin interactions 200bp (filter activated)
ZNF329 binding sites
ZNF329
CTCF,RAD21,SMC3 binding sites
Window Position
chr16:14,166,684-14,833,349 (666,666 bp)
Scale
200 kb
hg19
chr16:
14,300,000
14,400,000
14,500,000
14,600,000
14,700,000
14,800,000
0.210172
H3K9me3
1_H3K9me3
0
2.19778
H3K27me3
2_H3K27me3
0.00164
ln(x+1) 0.278046
RNA-seq
3_RNA-seq
9.32612e-17
Basic Gene Annotation Set from GENCODE Version 39lift37 (Ensembl 105)
0.448139
Insulation score based on chromatin interactions
4_Insulation score
0
-0.827853
Boundary of chromatin interactions
Strong
Weak
Strong
ZNF329 chromatin interactions 10bp (filter activated)
ZNF329 chromatin interactions 20bp (filter activated)
ZNF329 chromatin interactions 50bp (filter activated)
ZNF329 chromatin interactions 200bp (filter activated)
ZNF329 binding sites
ZNF329
CTCF,RAD21,SMC3 binding sites

### ZNF449

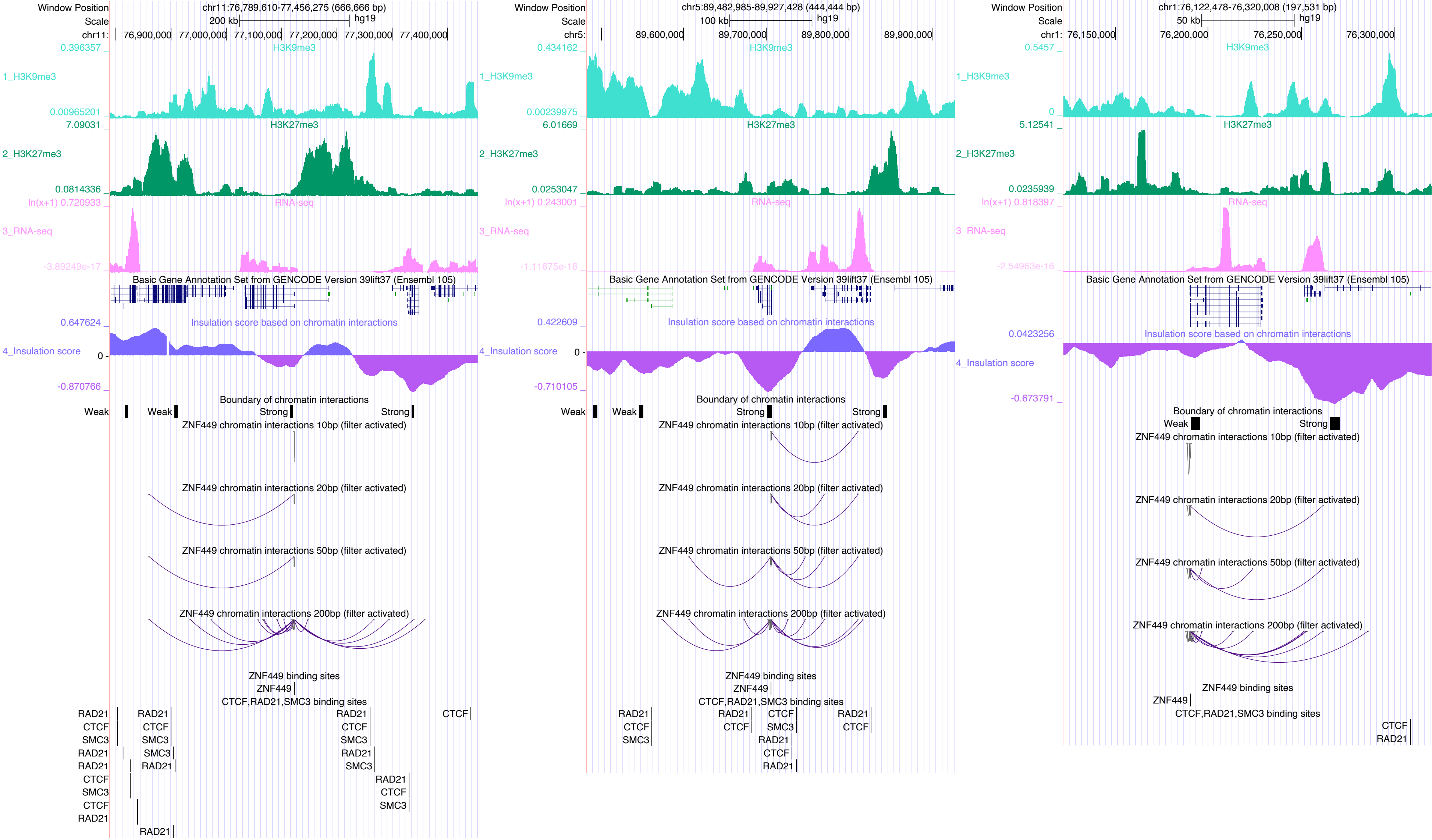

Window Position
chr11:76,789,610-77,456,275 (666,666 bp)
Scale
200 kb
hg19
chr11:
76,900,000
77,000,000
77,100,000
77,200,000
77,300,000
77,400,000
0.396357
H3K9me3
1_H3K9me3
0.00965201
7.09031
H3K27me3
2_H3K27me3
0.0814336
ln(x+1) 0.720933
RNA-seq
3_RNA-seq
-3.89249e-17
Basic Gene Annotation Set from GENCODE Version 39lift37 (Ensembl 105)
0.647624
Insulation score based on chromatin interactions
4_Insulation score
0
-0.870766
Boundary of chromatin interactions
Weak
Weak
Strong
Strong
ZNF449 chromatin interactions 10bp (filter activated)
ZNF449 chromatin interactions 20bp (filter activated)
ZNF449 chromatin interactions 50bp (filter activated)
ZNF449 chromatin interactions 200bp (filter activated)
ZNF449 binding sites
ZNF449
CTCF,RAD21,SMC3 binding sites
RAD21
CTCF
SMC3
RAD21
RAD21
CTCF
SMC3
CTCF
RAD21
RAD21
CTCF
SMC3
SMC3
RAD21
RAD21
RAD21
CTCF
SMC3
RAD21
SMC3
RAD21
CTCF
SMC3
CTCF
Window Position
chr5:89,482,985-89,927,428 (444,444 bp)
Scale
100 kb
hg19
chr5:
89,600,000
89,700,000
89,800,000
89,900,000
0.434162
H3K9me3
1_H3K9me3
0.00239975
6.01669
H3K27me3
2_H3K27me3
0.0253047
ln(x+1) 0.243001
RNA-seq
3_RNA-seq
-1.11675e-16
Basic Gene Annotation Set from GENCODE Version 39lift37 (Ensembl 105)
0.422609
Insulation score based on chromatin interactions
4_Insulation score
0
-0.710105
Boundary of chromatin interactions
Weak
Weak
Strong
Strong
ZNF449 chromatin interactions 10bp (filter activated)
ZNF449 chromatin interactions 20bp (filter activated)
ZNF449 chromatin interactions 50bp (filter activated)
ZNF449 chromatin interactions 200bp (filter activated)
ZNF449 binding sites
ZNF449
CTCF,RAD21,SMC3 binding sites
RAD21
CTCF
SMC3
RAD21
CTCF
CTCF
SMC3
RAD21
CTCF
RAD21
RAD21
CTCF
Window Position
chr1:76,122,478-76,320,008 (197,531 bp)
Scale
50 kb
hg19
chr1:
76,150,000
76,200,000
76,250,000
76,300,000
0.5457
H3K9me3
1_H3K9me3
0
5.12541
H3K27me3
2_H3K27me3
0.0235939
ln(x+1) 0.818397
RNA-seq
3_RNA-seq
-2.54963e-16
Basic Gene Annotation Set from GENCODE Version 39lift37 (Ensembl 105)
0.0423256
Insulation score based on chromatin interactions
4_Insulation score
-0.673791
Boundary of chromatin interactions
Weak
Strong
ZNF449 chromatin interactions 10bp (filter activated)
ZNF449 chromatin interactions 20bp (filter activated)
ZNF449 chromatin interactions 50bp (filter activated)
ZNF449 chromatin interactions 200bp (filter activated)
ZNF449 binding sites
ZNF449
CTCF,RAD21,SMC3 binding sites
CTCF
RAD21

ZZZ3

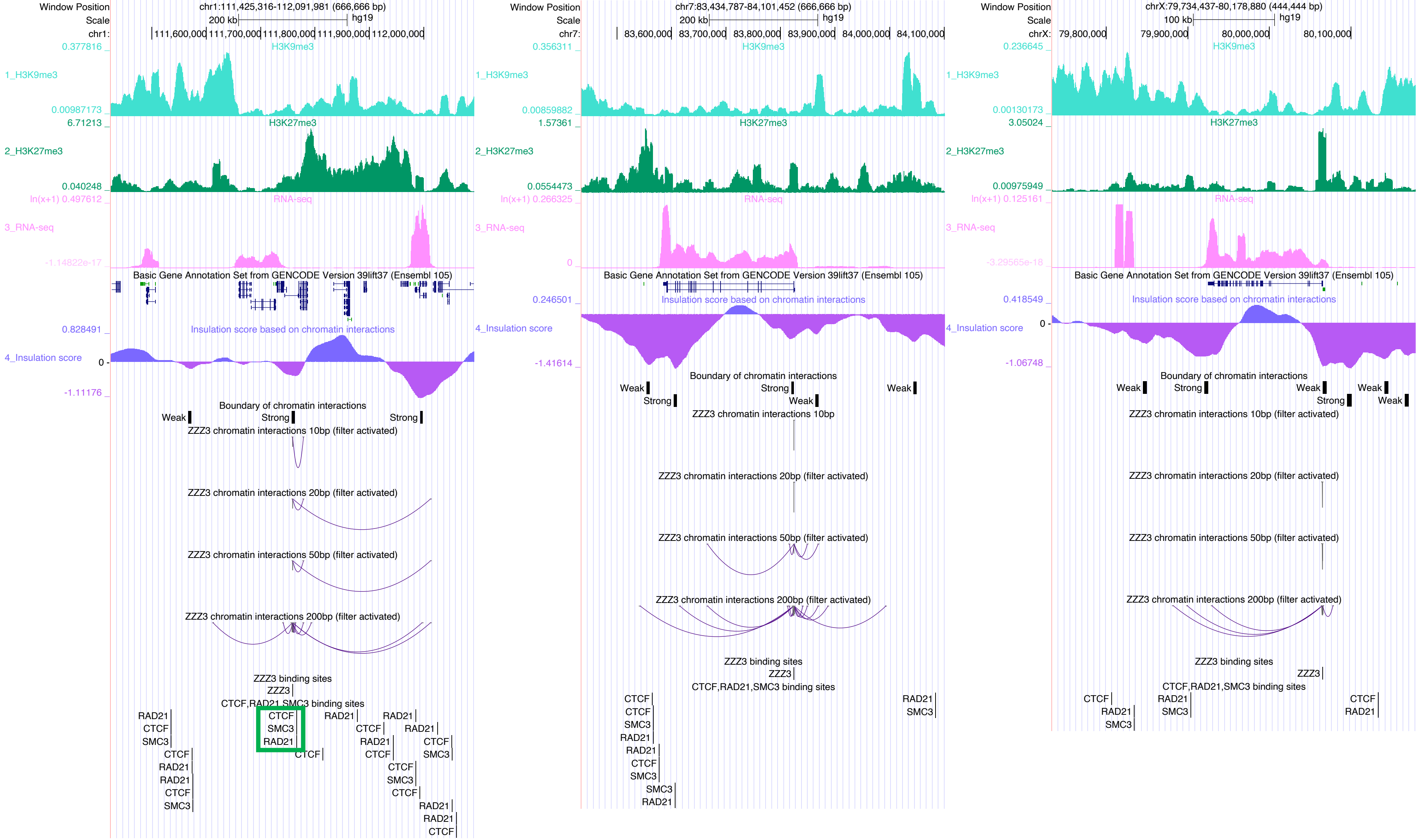

Window Position chr1:111,425,316-112,091,981 (666,666 bp)
Scale 200 kb hg19
H3K9me3
1_H3K9me3
H3K27me3
2_H3K27me3
RNA-seq
3_RNA-seq
Basic Gene Annotation Set from GENCODE Version 39lift37 (Ensembl 105)
Insulation score based on chromatin interactions
4_Insulation score
Boundary of chromatin interactions
ZZZ3 chromatin interactions 10bp (filter activated)
ZZZ3 chromatin interactions 20bp (filter activated)
ZZZ3 chromatin interactions 50bp (filter activated)
ZZZ3 chromatin interactions 200bp (filter activated)
ZZZ3 binding sites
CTCF,RAD21,SMC3 binding sites
Window Position chr7:83,434,787-84,101,452 (666,666 bp)
ZZZ3 chromatin interactions 10bp
Window Position chrX:79,734,437-80,178,880 (444,444 bp)
Scale 100 kb hg19

b

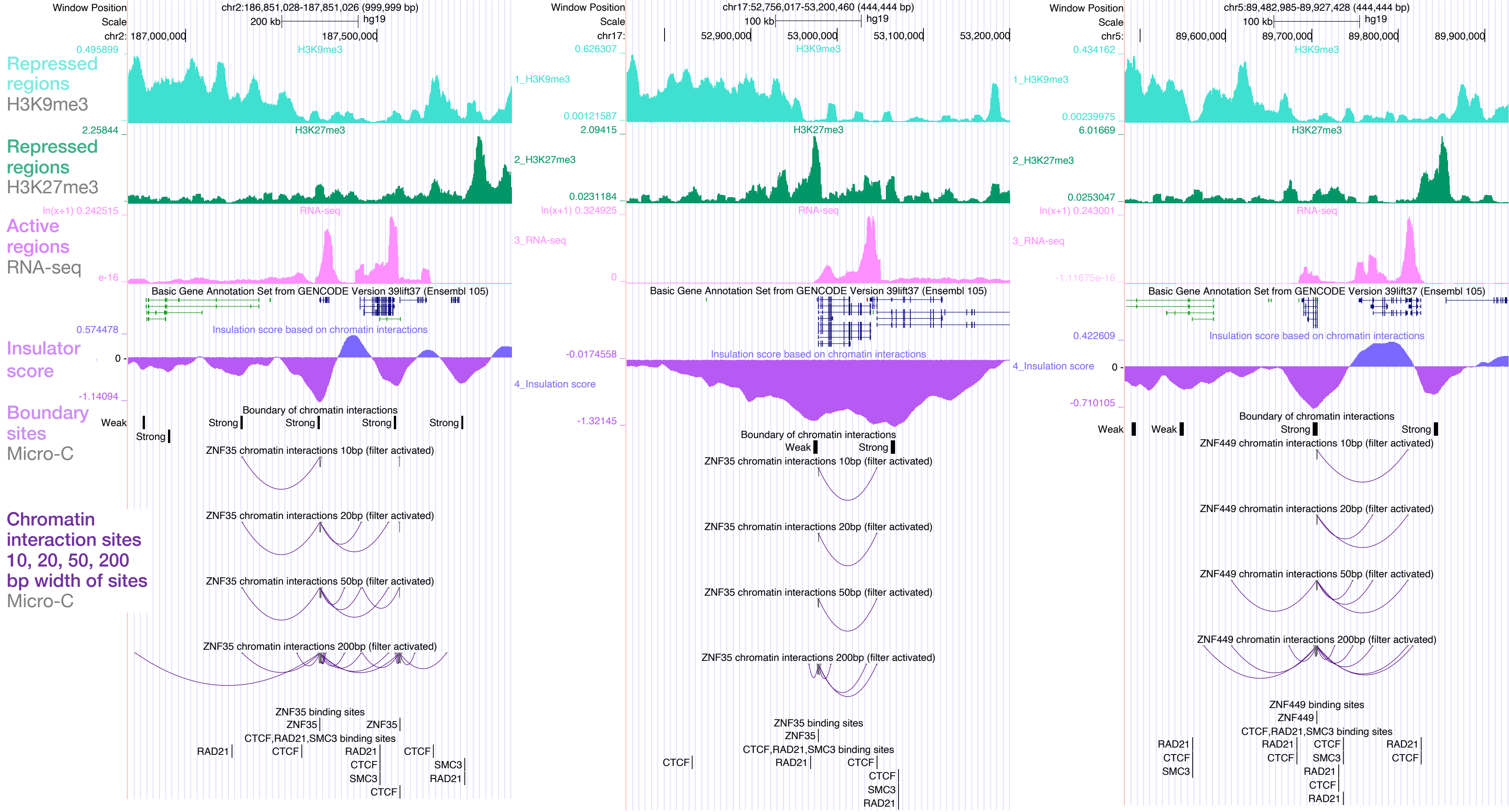


**b** Insulator-associated DNA-binding sites where H3K9me3 marks (top track) differ between the upstream and downstream regions of the site.

b

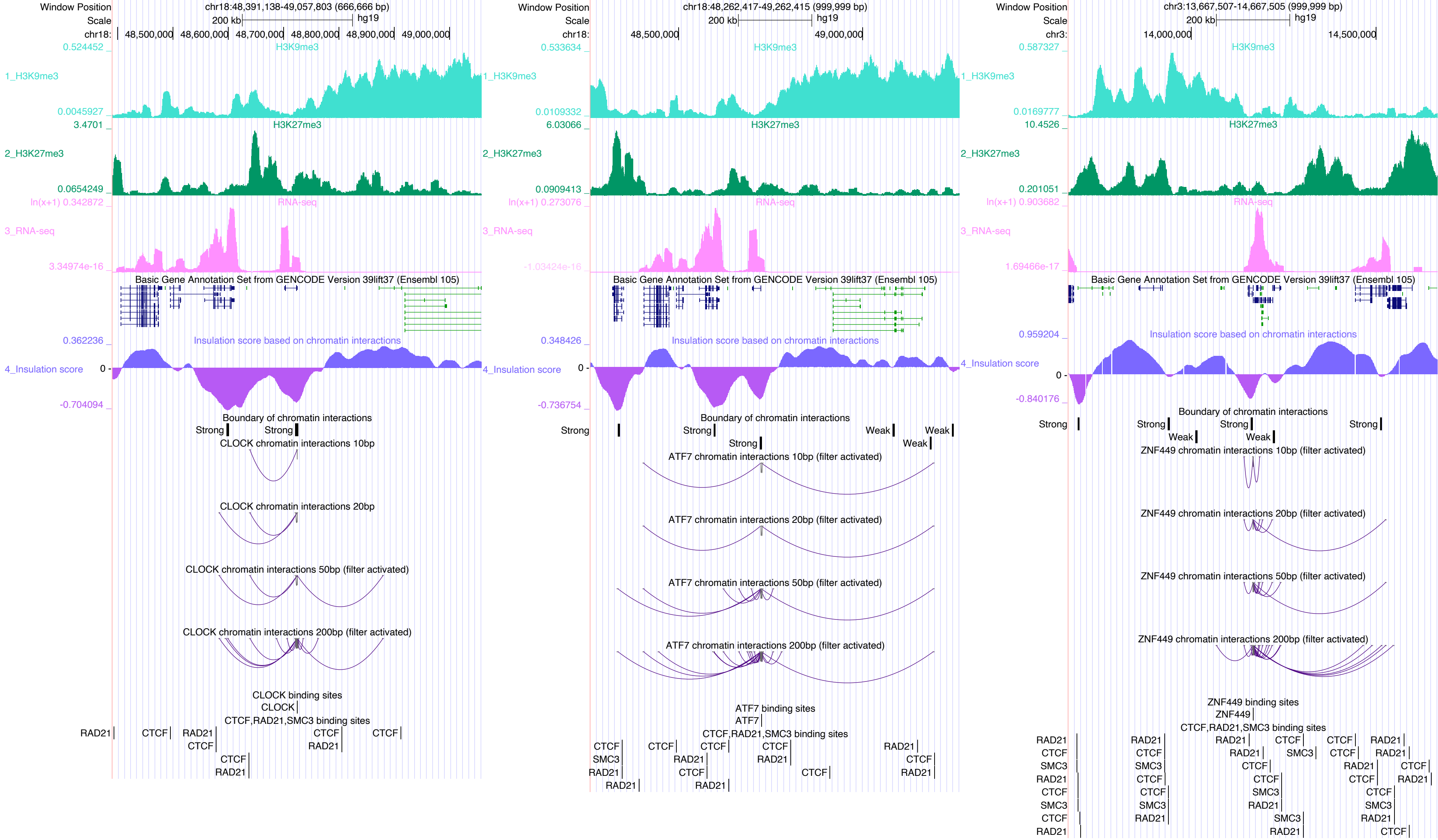
Window Position chr18:48,391,138-49,057,803 (666,666 bp)
Scale 200 kb hg19
chr18: 48,500,000 48,600,000 48,700,000 48,800,000 48,900,000 49,000,000
H3K9me3
1_H3K9me3
H3K27me3
2_H3K27me3
RNA-seq
3_RNA-seq
Basic Gene Annotation Set from GENCODE Version 39lift37 (Ensembl 105)
Insulation score based on chromatin interactions
4_Insulation score
Boundary of chromatin interactions
Strong
Weak
CLOCK chromatin interactions 10bp
CLOCK chromatin interactions 20bp
CLOCK chromatin interactions 50bp (filter activated)
CLOCK chromatin interactions 200bp (filter activated)
CLOCK binding sites
CLOCK
CTCF,RAD21,SMC3 binding sites
Window Position chr18:48,262,417-49,262,415 (999,999 bp)
chr18: 48,500,000 49,000,000
ATF7 chromatin interactions 10bp (filter activated)
ATF7 chromatin interactions 20bp (filter activated)
ATF7 chromatin interactions 50bp (filter activated)
ATF7 chromatin interactions 200bp (filter activated)
ATF7 binding sites
ATF7
Window Position chr3:13,667,507-14,667,505 (999,999 bp)
chr3: 14,000,000 14,500,000
ZNF449 chromatin interactions 10bp (filter activated)
ZNF449 chromatin interactions 20bp (filter activated)
ZNF449 chromatin interactions 50bp (filter activated)
ZNF449 chromatin interactions 200bp (filter activated)
ZNF449 binding sites
ZNF449
CTCF
RAD21
SMC3

b

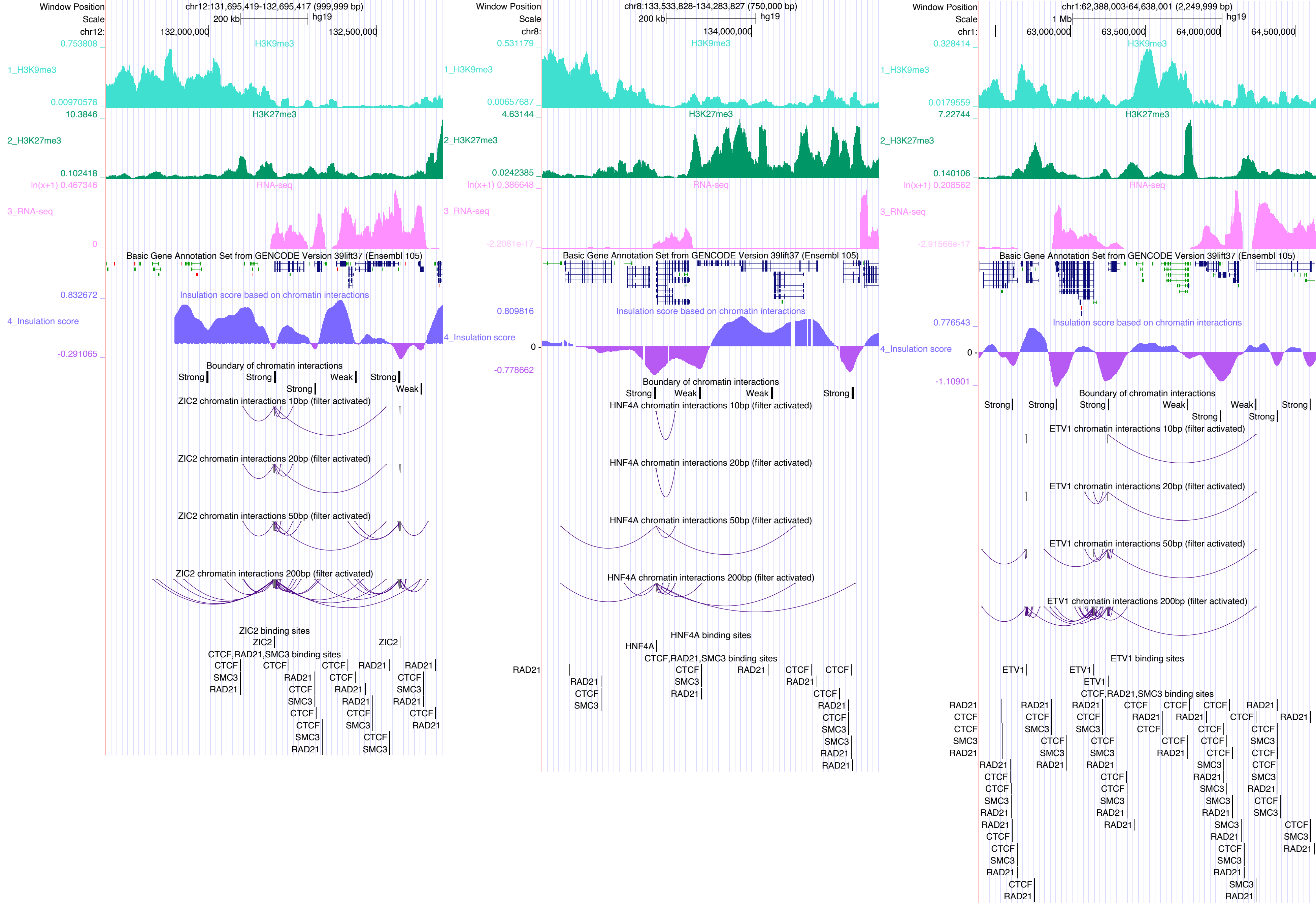

Window Position
chr12:131,695,419-132,695,417 (999,999 bp)
Scale
200 kb
hg19
chr12:
132,000,000
132,500,000
0.753808
H3K9me3
1_H3K9me3
0.00970578
10.3846
H3K27me3
2_H3K27me3
0.102418
ln(x+1) 0.467346
RNA-seq
3_RNA-seq
0
Basic Gene Annotation Set from GENCODE Version 39lift37 (Ensembl 105)
0.832672
Insulation score based on chromatin interactions
4_Insulation score
-0.291065
Boundary of chromatin interactions
ZIC2 chromatin interactions 10bp (filter activated)
ZIC2 chromatin interactions 20bp (filter activated)
ZIC2 chromatin interactions 50bp (filter activated)
ZIC2 chromatin interactions 200bp (filter activated)
ZIC2 binding sites
CTCF,RAD21,SMC3 binding sites
Window Position
chr8:133,533,828-134,283,827 (750,000 bp)
200 kb
chr8:
134,000,000
0.531179
0.00657687
4.63144
0.0242385
ln(x+1) 0.386648
-2.2081e-17
0.809816
0
-0.778662
HNF4A chromatin interactions 10bp (filter activated)
HNF4A chromatin interactions 20bp (filter activated)
HNF4A chromatin interactions 50bp (filter activated)
HNF4A chromatin interactions 200bp (filter activated)
HNF4A binding sites
Window Position
chr1:62,388,003-64,638,001 (2,249,999 bp)
1 Mb
chr1:
63,000,000
63,500,000
64,000,000
64,500,000
0.328414
0.0179559
7.22744
0.140106
ln(x+1) 0.208562
-2.91566e-17
0.776543
-1.10901
ETV1 chromatin interactions 10bp (filter activated)
ETV1 chromatin interactions 20bp (filter activated)
ETV1 chromatin interactions 50bp (filter activated)
ETV1 chromatin interactions 200bp (filter activated)
ETV1 binding sites

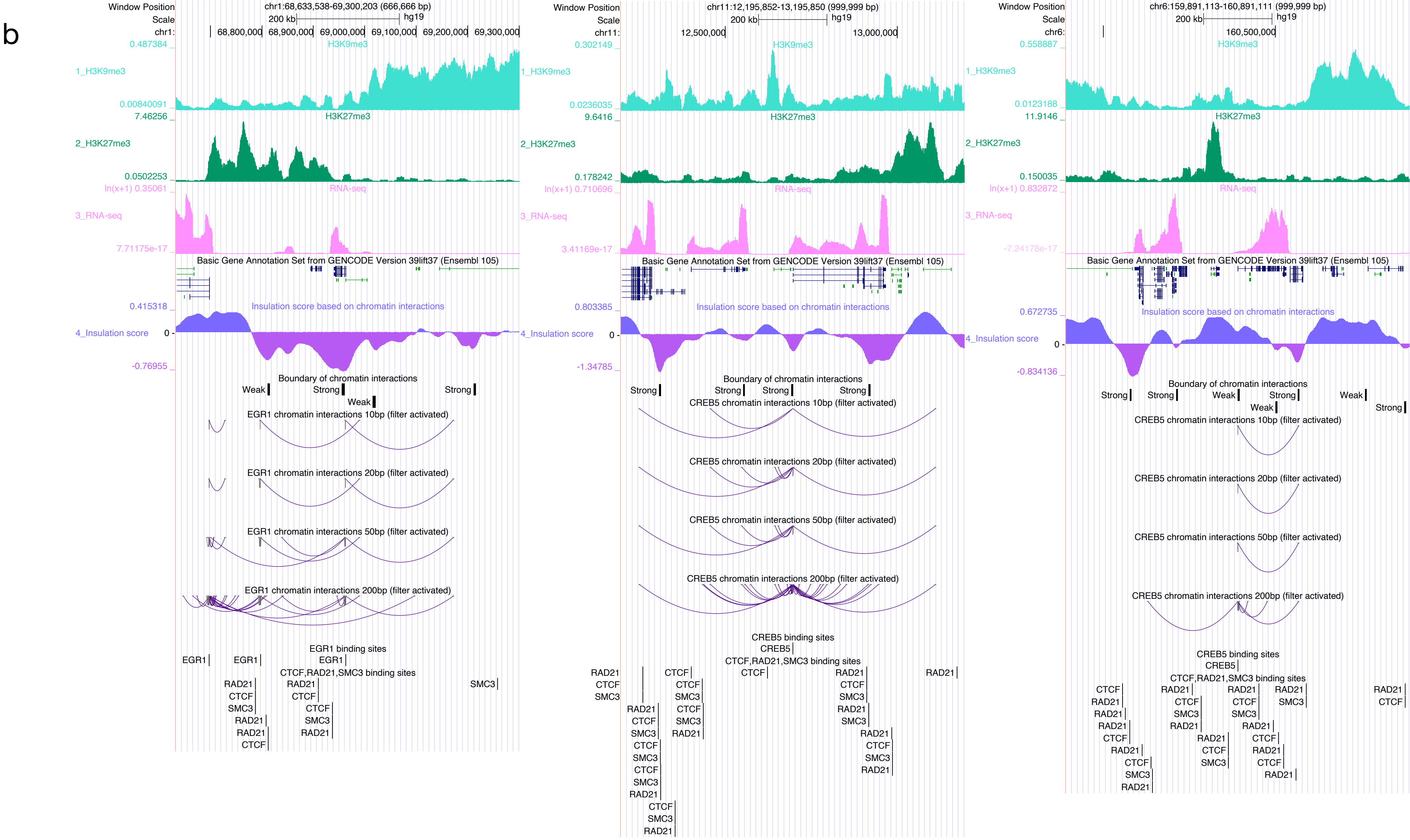
b
Window Position
Scale
chr1:68,633,538-69,300,203 (666,666 bp)
200 kb
hg19
68,800,000
68,900,000
69,000,000
69,100,000
69,200,000
69,300,000
H3K9me3
1_H3K9me3
H3K27me3
2_H3K27me3
RNA-seq
3_RNA-seq
Basic Gene Annotation Set from GENCODE Version 39lift37 (Ensembl 105)
Insulation score based on chromatin interactions
4_Insulation score
Boundary of chromatin interactions
Weak
Strong
EGR1 chromatin interactions 10bp (filter activated)
EGR1 chromatin interactions 20bp (filter activated)
EGR1 chromatin interactions 50bp (filter activated)
EGR1 chromatin interactions 200bp (filter activated)
EGR1 binding sites
CTCF,RAD21,SMC3 binding sites
chr11:12,195,852-13,195,850 (999,999 bp)
12,500,000
13,000,000
CREB5 chromatin interactions 10bp (filter activated)
CREB5 chromatin interactions 20bp (filter activated)
CREB5 chromatin interactions 50bp (filter activated)
CREB5 chromatin interactions 200bp (filter activated)
CREB5 binding sites
chr6:159,891,113-160,891,111 (999,999 bp)
160,500,000
EGR1
CREB5
CTCF
RAD21
SMC3

b

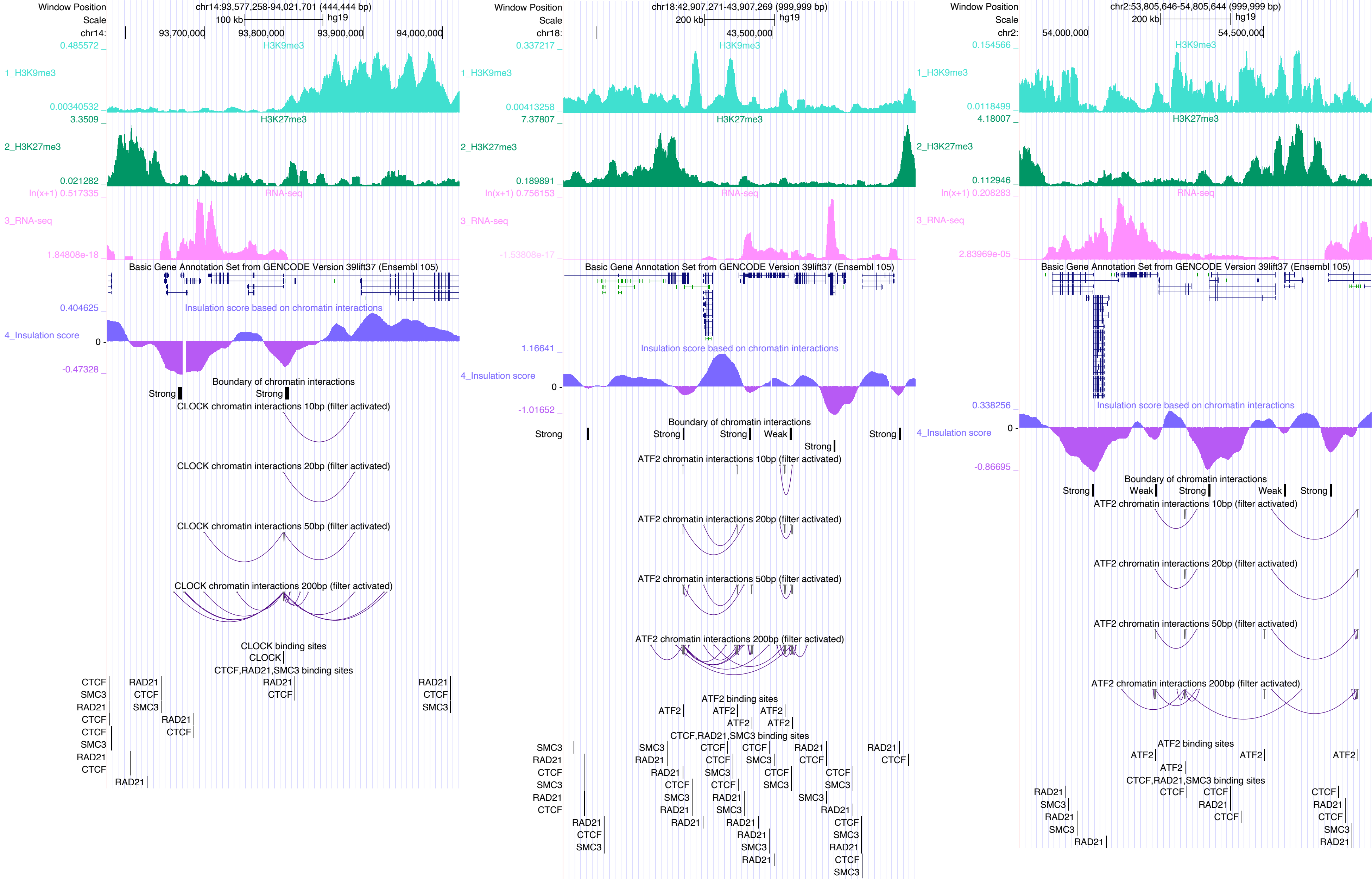
Window Position
Scale
chr14:93,577,258-94,021,701 (444,444 bp)
100 kb
hg19
chr14:
93,700,000
93,800,000
93,900,000
94,000,000
0.485572
H3K9me3
1_H3K9me3
0.00340532
3.3509
H3K27me3
2_H3K27me3
0.021282
ln(x+1) 0.517335
RNA-seq
3_RNA-seq
1.84808e-18
Basic Gene Annotation Set from GENCODE Version 39lift37 (Ensembl 105)
0.404625
Insulation score based on chromatin interactions
4_Insulation score
0
-0.47328
Boundary of chromatin interactions
Strong
Strong
CLOCK chromatin interactions 10bp (filter activated)
CLOCK chromatin interactions 20bp (filter activated)
CLOCK chromatin interactions 50bp (filter activated)
CLOCK chromatin interactions 200bp (filter activated)
CLOCK binding sites
CLOCK
CTCF,RAD21,SMC3 binding sites
CTCF
SMC3
RAD21
RAD21
CTCF
SMC3
Window Position
Scale
chr18:42,907,271-43,907,269 (999,999 bp)
200 kb
hg19
chr18:
43,500,000
0.337217
H3K9me3
1_H3K9me3
0.00413258
7.37807
H3K27me3
2_H3K27me3
0.189891
ln(x+1) 0.756153
RNA-seq
3_RNA-seq
-1.53808e-17
Basic Gene Annotation Set from GENCODE Version 39lift37 (Ensembl 105)
1.16641
Insulation score based on chromatin interactions
4_Insulation score
0
-1.01652
Boundary of chromatin interactions
Strong
Strong
Strong
Weak
Strong
Strong
ATF2 chromatin interactions 10bp (filter activated)
ATF2 chromatin interactions 20bp (filter activated)
ATF2 chromatin interactions 50bp (filter activated)
ATF2 chromatin interactions 200bp (filter activated)
ATF2 binding sites
ATF2
CTCF,RAD21,SMC3 binding sites
SMC3
RAD21
CTCF
Window Position
Scale
chr2:53,805,646-54,805,644 (999,999 bp)
200 kb
hg19
chr2:
54,000,000
54,500,000
0.154566
H3K9me3
1_H3K9me3
0.0118499
4.18007
H3K27me3
2_H3K27me3
0.112946
ln(x+1) 0.208283
RNA-seq
3_RNA-seq
2.83969e-05
Basic Gene Annotation Set from GENCODE Version 39lift37 (Ensembl 105)
0.338256
Insulation score based on chromatin interactions
4_Insulation score
0
-0.86695
Boundary of chromatin interactions
Strong
Weak
Strong
Weak
Strong
ATF2 chromatin interactions 10bp (filter activated)
ATF2 chromatin interactions 20bp (filter activated)
ATF2 chromatin interactions 50bp (filter activated)
ATF2 chromatin interactions 200bp (filter activated)
ATF2 binding sites
ATF2
CTCF,RAD21,SMC3 binding sites
RAD21
SMC3
CTCF

c

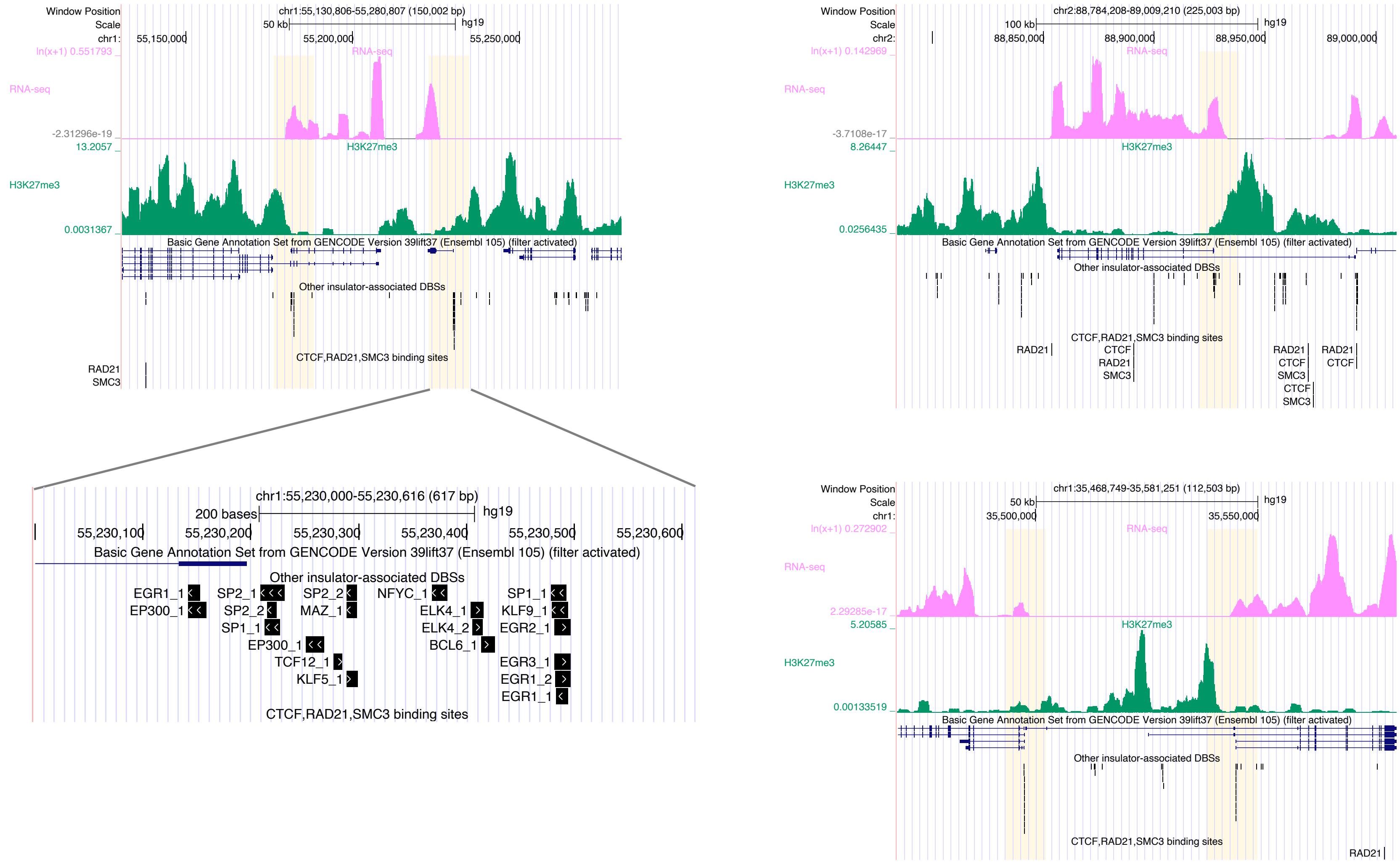


**c** Cluster of DNA-binding sites of insulator-associated DBPs. Clusters of DNA-binding sites of predicted insulator-associated DBPs are highlighted in yellow within insulator sites. Each dot represents the DNA-binding site of a predicted insulator-associated DBP.

d

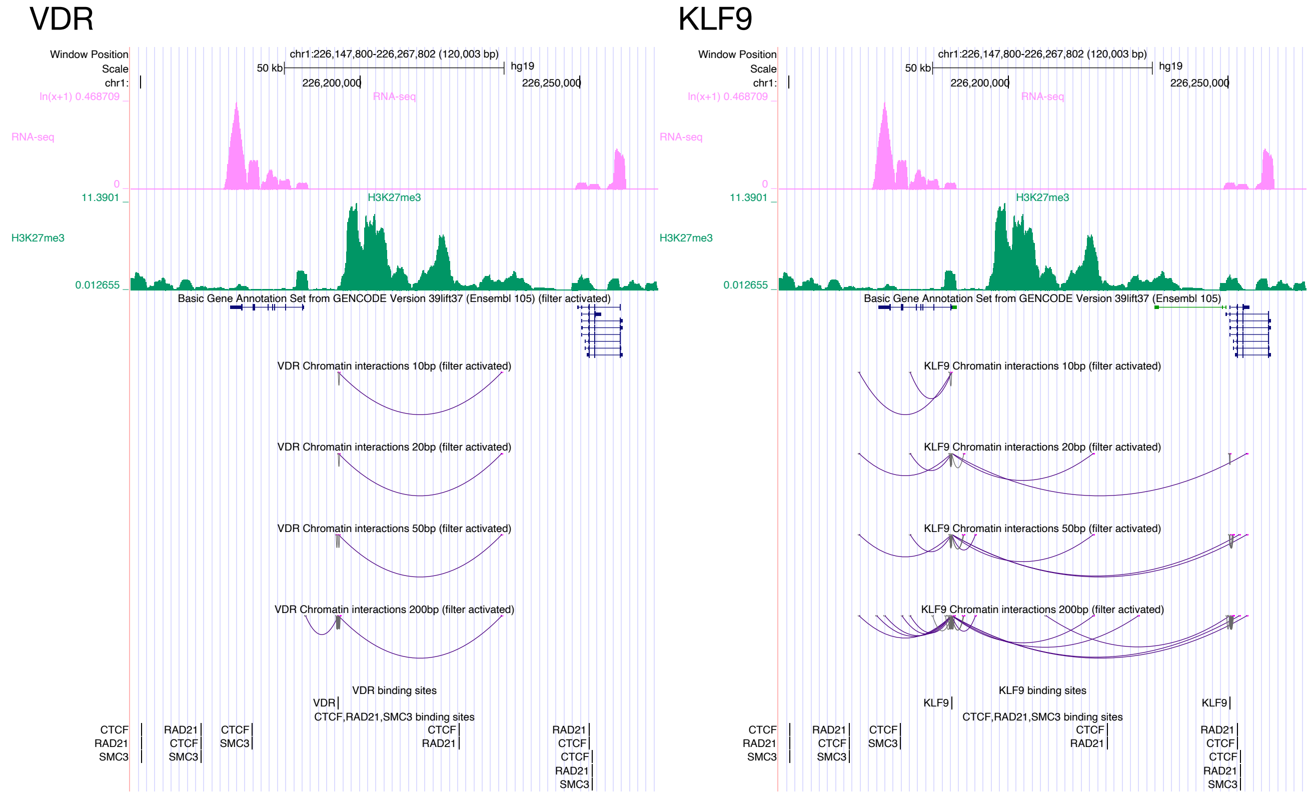


**d** Differences in chromatin interactions between DBPs at the same loci.

d

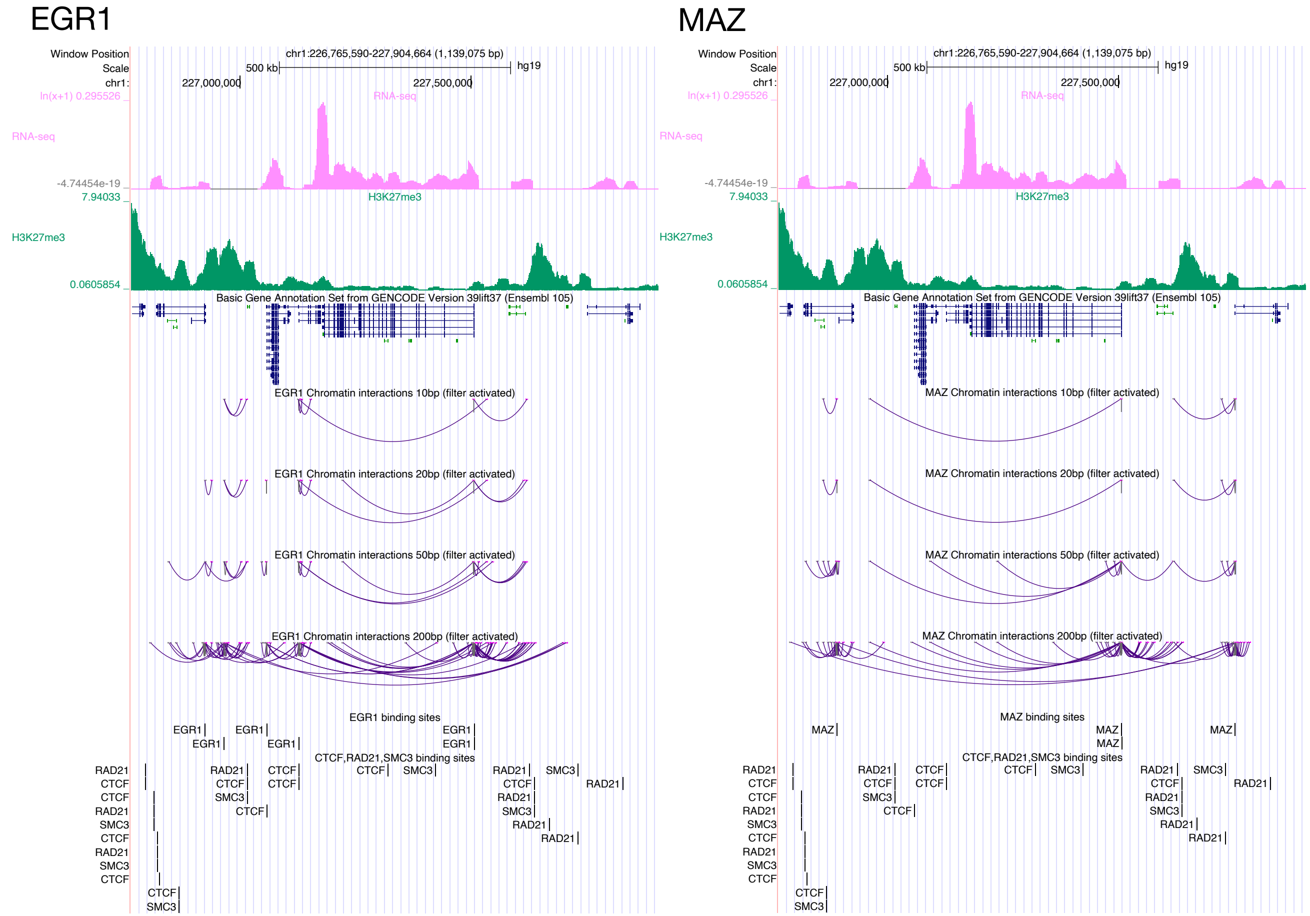


**d** Differences in chromatin interactions between DBPs at the same loci.

e

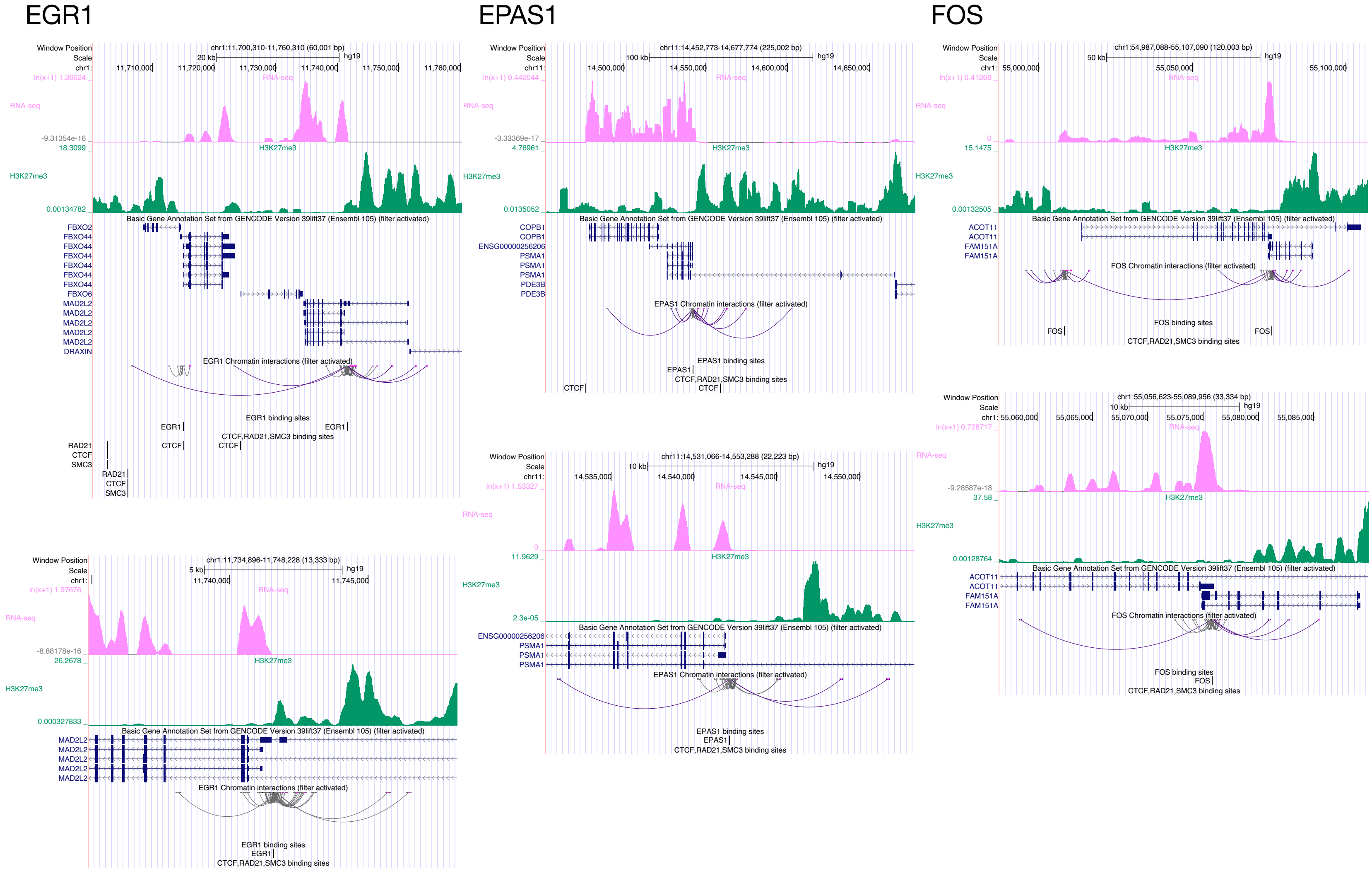


**e** Potential regulation of alternative transcription.

e

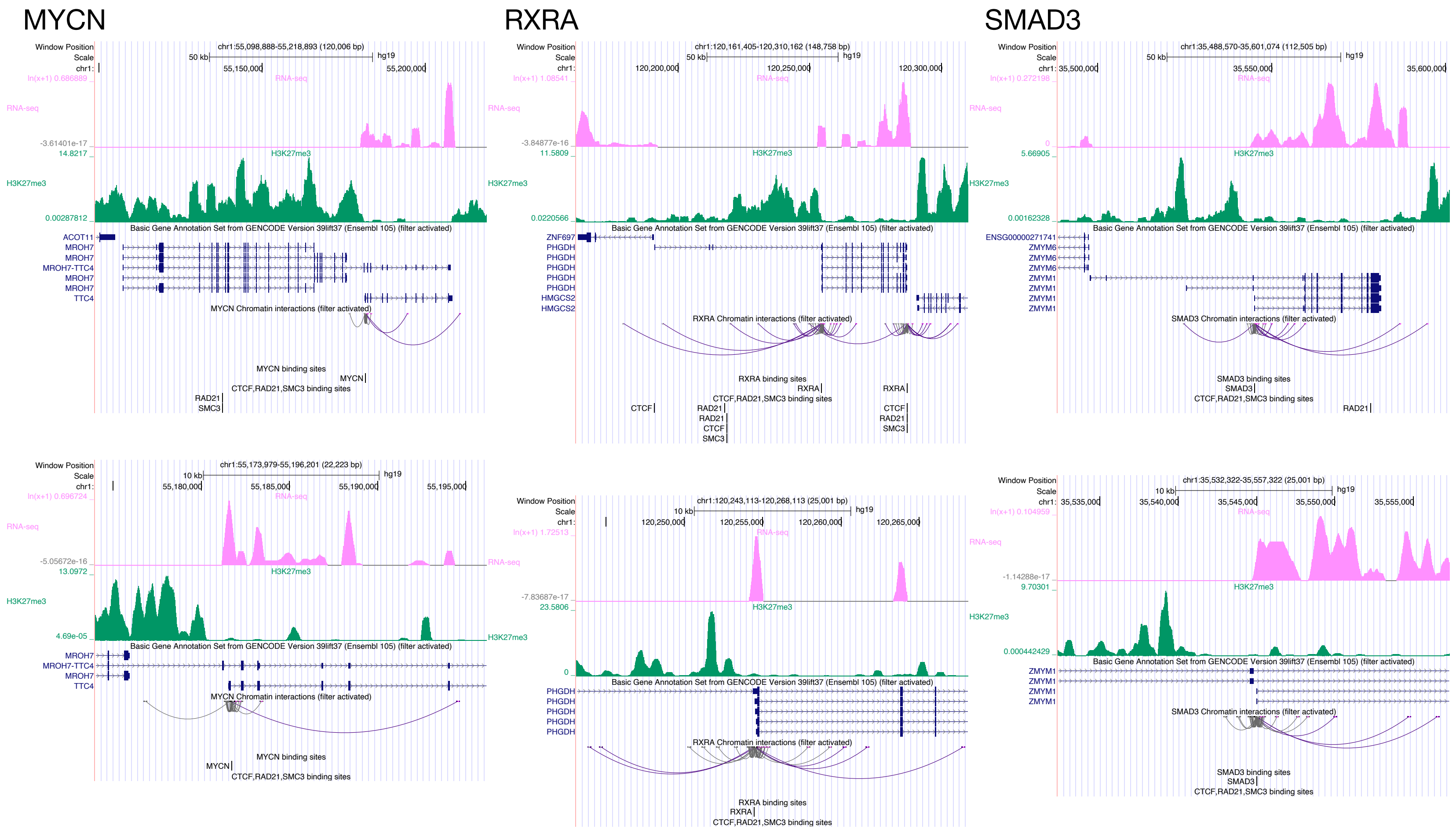


**e** Potential regulation of alternative transcription.

f

AHR

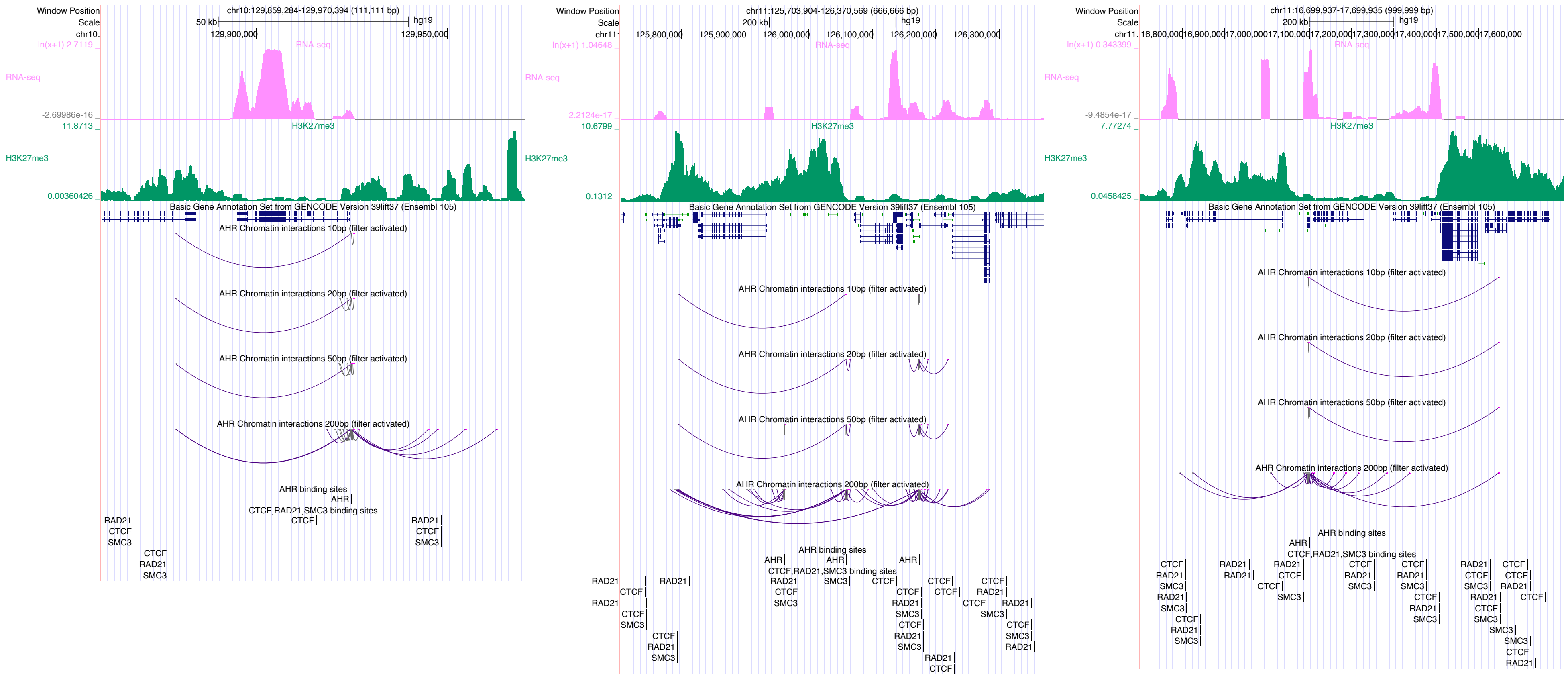


**f** Insulator sites identified as boundaries between transcriptionally repressed regions (based on H3K27me3 marks) and transcribed regions (based on RNA-seq data). Screenshots from the UCSC Genome Browser of regions surrounding insulator-associated DNA-binding sites.

### ASCL1

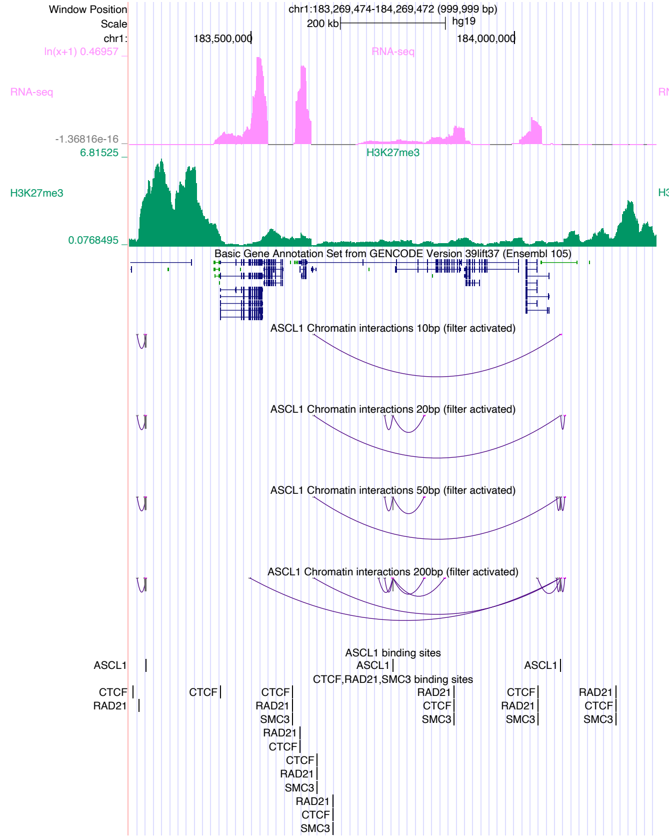

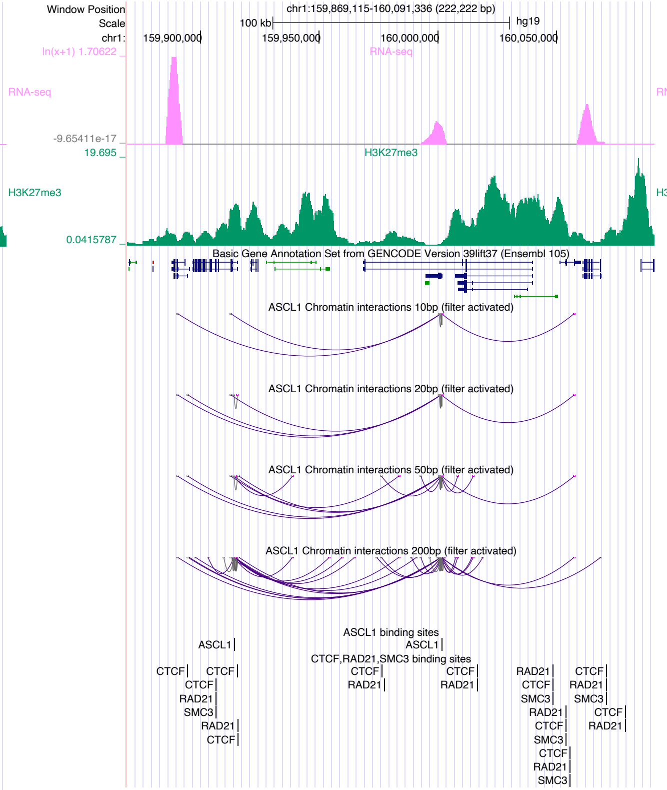

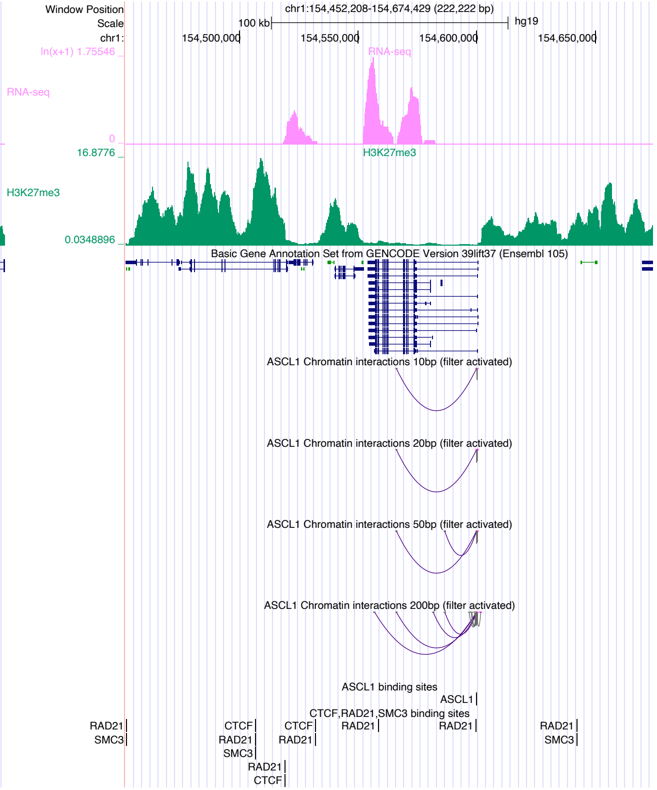

### ATF2

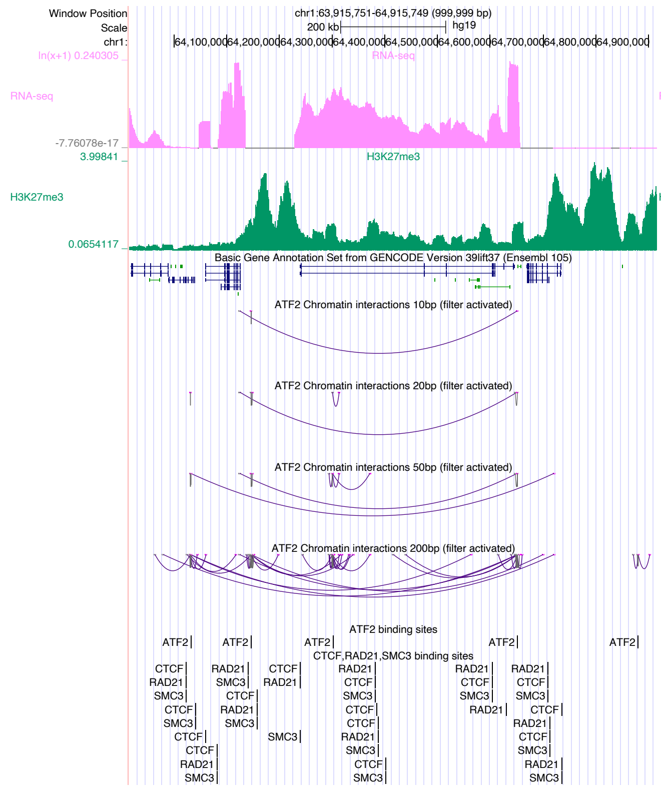

Window Position
chr1:63,915,751-64,915,749 (999,999 bp)
Scale
200 kb
hg19
RNA-seq
H3K27me3
Basic Gene Annotation Set from GENCODE Version 39lift37 (Ensembl 105)
ATF2 Chromatin interactions 10bp (filter activated)
ATF2 Chromatin interactions 20bp (filter activated)
ATF2 Chromatin interactions 50bp (filter activated)
ATF2 Chromatin interactions 200bp (filter activated)
ATF2 binding sites
CTCF,RAD21,SMC3 binding sites


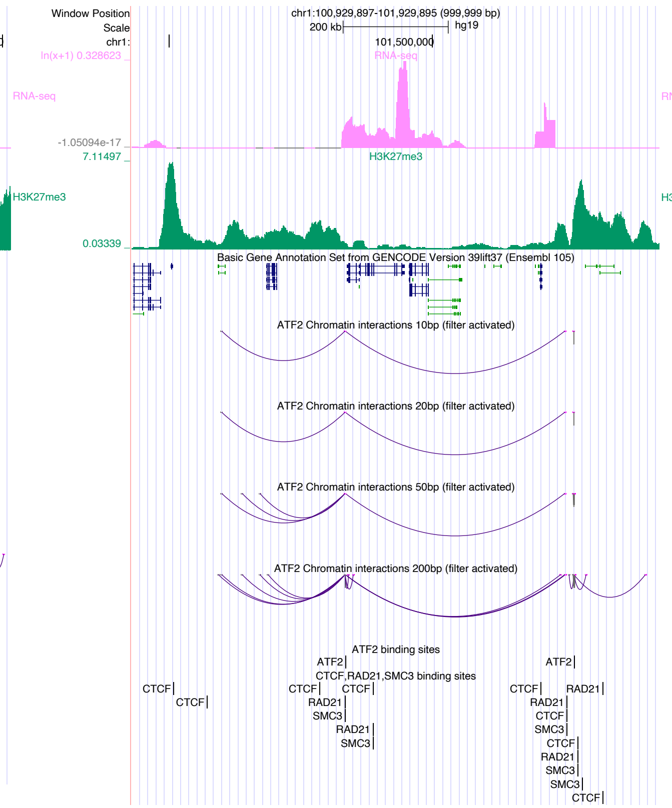

Window Position
chr1:100,929,897-101,929,895 (999,999 bp)
Scale
200 kb
hg19
101,500,000
RNA-seq
H3K27me3
Basic Gene Annotation Set from GENCODE Version 39lift37 (Ensembl 105)
ATF2 Chromatin interactions 10bp (filter activated)
ATF2 Chromatin interactions 20bp (filter activated)
ATF2 Chromatin interactions 50bp (filter activated)
ATF2 Chromatin interactions 200bp (filter activated)
ATF2 binding sites
CTCF,RAD21,SMC3 binding sites


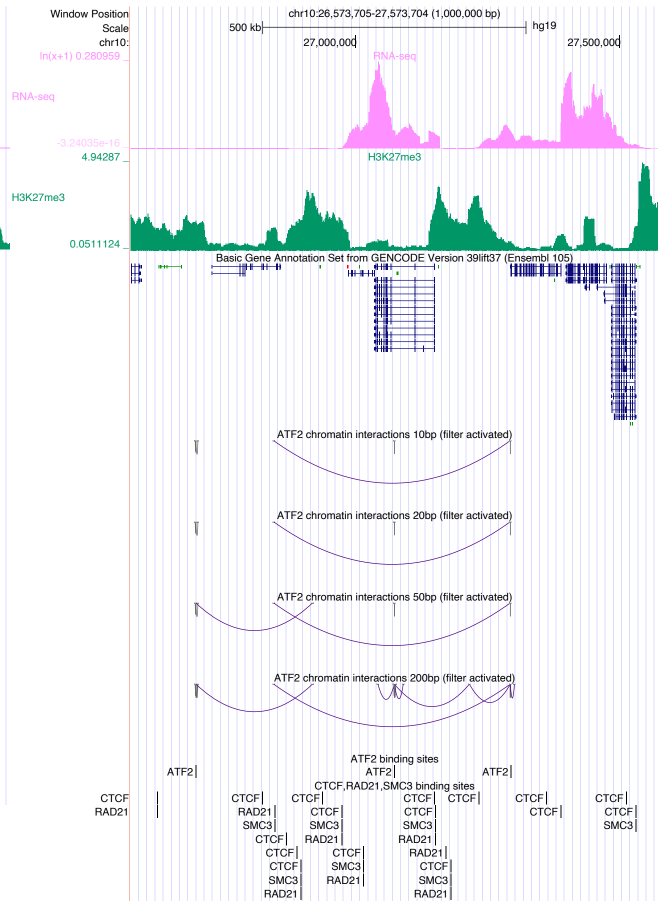

Window Position
chr10:26,573,705-27,573,704 (1,000,000 bp)
Scale
500 kb
hg19
27,000,000
27,500,000
RNA-seq
H3K27me3
Basic Gene Annotation Set from GENCODE Version 39lift37 (Ensembl 105)
ATF2 chromatin interactions 10bp (filter activated)
ATF2 chromatin interactions 20bp (filter activated)
ATF2 chromatin interactions 50bp (filter activated)
ATF2 chromatin interactions 200bp (filter activated)
ATF2 binding sites
CTCF,RAD21,SMC3 binding sites

### ATF7

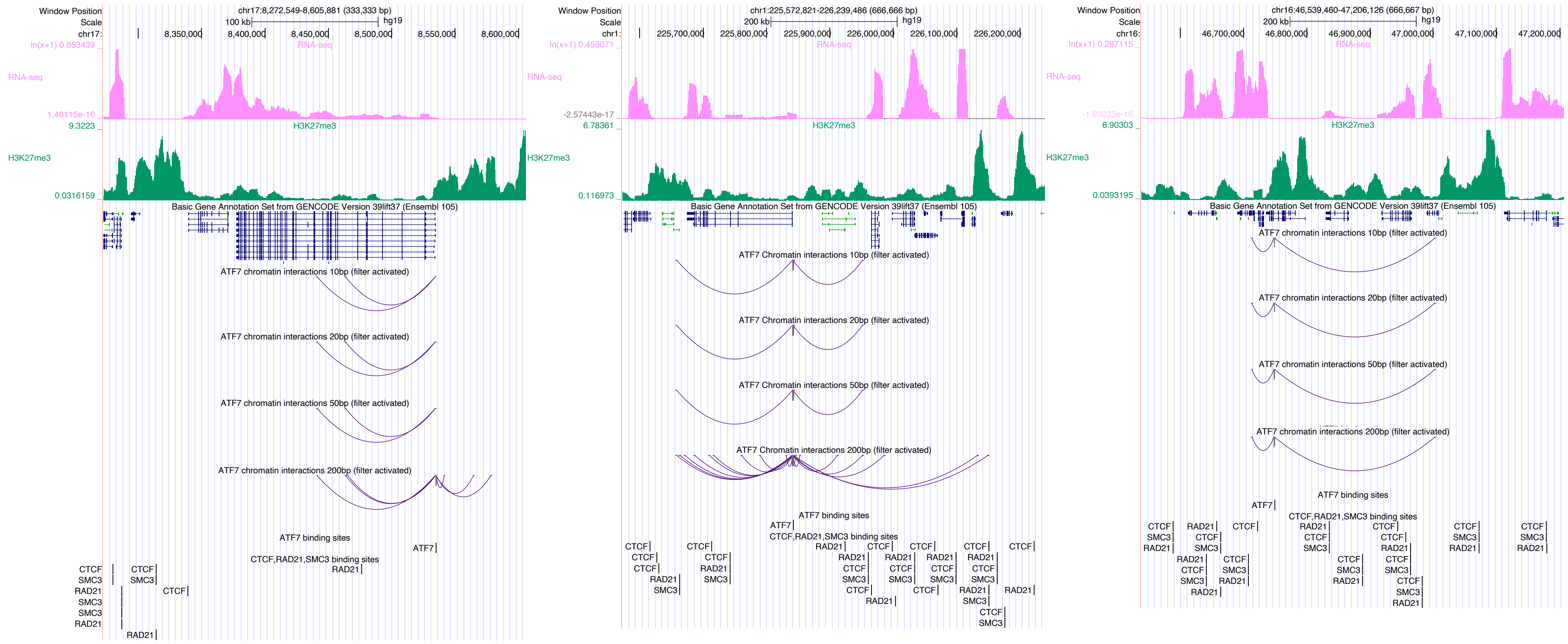

Window Position
chr17:8,272,549-8,605,881 (333,333 bp)
chr1:225,572,821-226,239,486 (666,666 bp)
chr16:46,539,460-47,206,126 (666,667 bp)
RNA-seq
H3K27me3
Basic Gene Annotation Set from GENCODE Version 39lift37 (Ensembl 105)
ATF7 chromatin interactions 10bp (filter activated)
ATF7 chromatin interactions 20bp (filter activated)
ATF7 chromatin interactions 50bp (filter activated)
ATF7 chromatin interactions 200bp (filter activated)
ATF7 binding sites
CTCF,RAD21,SMC3 binding sites

### BATF

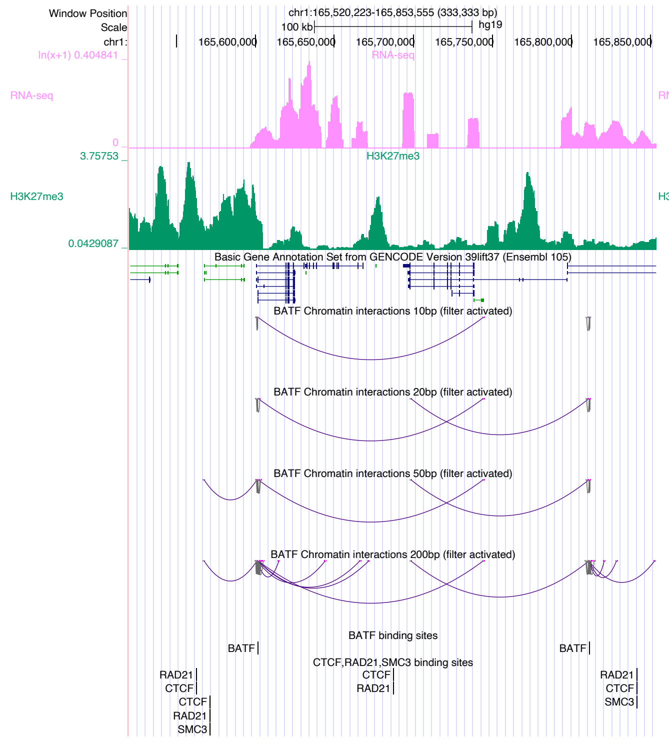
Window Position
Scale
chr1:
chr1:165,520,223-165,853,555 (333,333 bp)
100 kb
hg19
165,600,000
165,650,000
165,700,000
165,750,000
165,800,000
165,850,000
ln(x+1) 0.404841
RNA-seq
0
3.75753
H3K27me3
0.0429087
Basic Gene Annotation Set from GENCODE Version 39lift37 (Ensembl 105)
BATF Chromatin interactions 10bp (filter activated)
BATF Chromatin interactions 20bp (filter activated)
BATF Chromatin interactions 50bp (filter activated)
BATF Chromatin interactions 200bp (filter activated)
BATF binding sites
BATF
CTCF,RAD21,SMC3 binding sites
RAD21
CTCF
SMC3

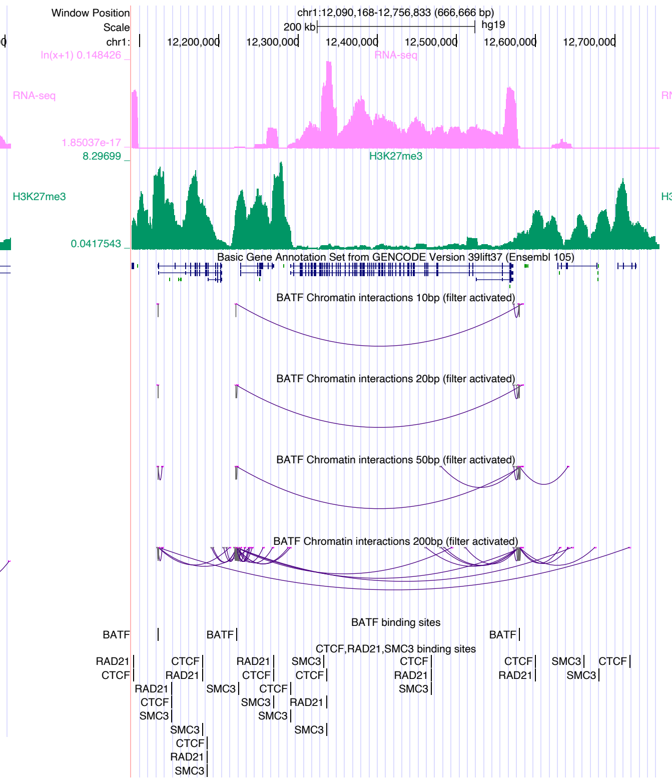
Window Position
Scale
chr1:
chr1:12,090,168-12,756,833 (666,666 bp)
200 kb
hg19
12,200,000
12,300,000
12,400,000
12,500,000
12,600,000
12,700,000
ln(x+1) 0.148426
RNA-seq
1.85037e-17
8.29699
H3K27me3
0.0417543
Basic Gene Annotation Set from GENCODE Version 39lift37 (Ensembl 105)
BATF Chromatin interactions 10bp (filter activated)
BATF Chromatin interactions 20bp (filter activated)
BATF Chromatin interactions 50bp (filter activated)
BATF Chromatin interactions 200bp (filter activated)
BATF binding sites
BATF
CTCF,RAD21,SMC3 binding sites
RAD21
CTCF
SMC3

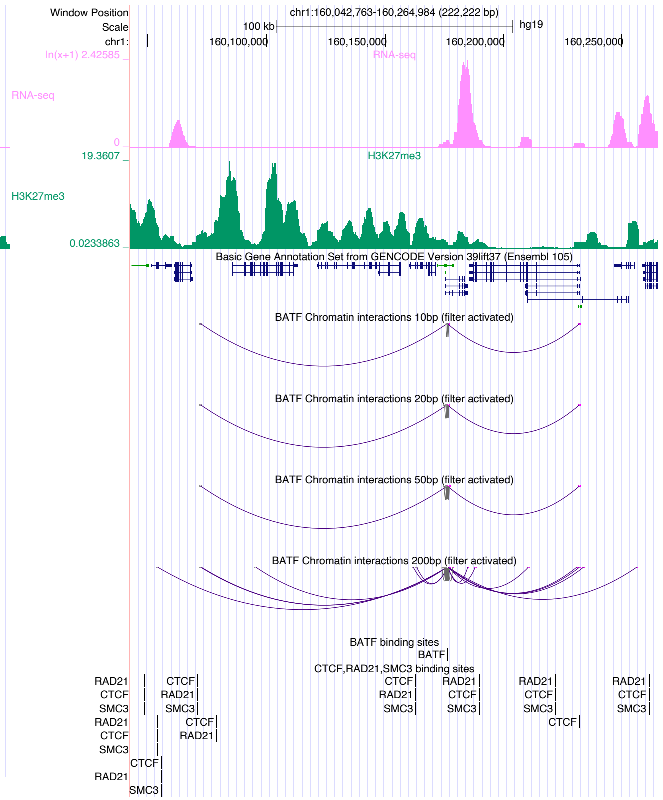
Window Position
Scale
chr1:
chr1:160,042,763-160,264,984 (222,222 bp)
100 kb
hg19
160,100,000
160,150,000
160,200,000
160,250,000
ln(x+1) 2.42585
RNA-seq
0
19.3607
H3K27me3
0.0233863
Basic Gene Annotation Set from GENCODE Version 39lift37 (Ensembl 105)
BATF Chromatin interactions 10bp (filter activated)
BATF Chromatin interactions 20bp (filter activated)
BATF Chromatin interactions 50bp (filter activated)
BATF Chromatin interactions 200bp (filter activated)
BATF binding sites
BATF
CTCF,RAD21,SMC3 binding sites
RAD21
CTCF
SMC3

### BATF3

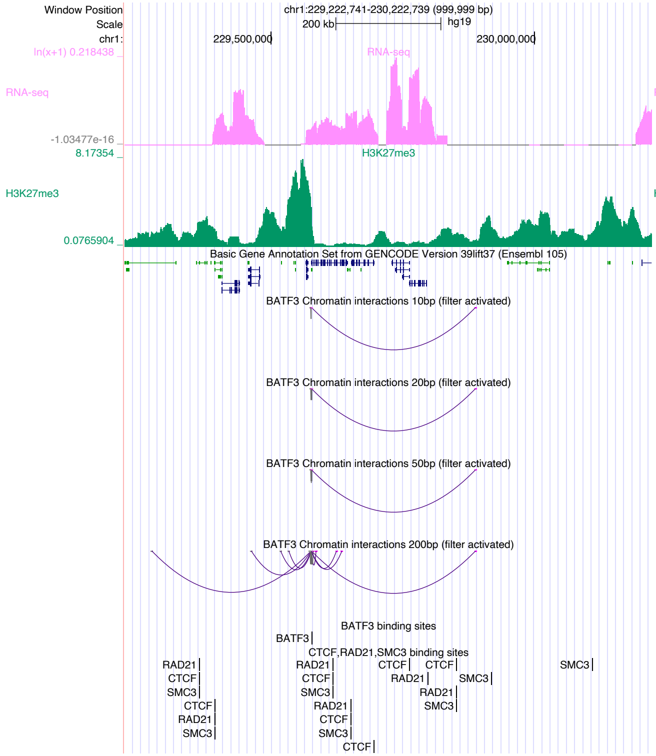

Window Position chr1:229,222,741-230,222,739 (999,999 bp)
Scale 200 kb hg19
chr1: 229,500,000 230,000,000
RNA-seq
H3K27me3
Basic Gene Annotation Set from GENCODE Version 39lift37 (Ensembl 105)
BATF3 Chromatin interactions 10bp (filter activated)
BATF3 Chromatin interactions 20bp (filter activated)
BATF3 Chromatin interactions 50bp (filter activated)
BATF3 Chromatin interactions 200bp (filter activated)
BATF3 binding sites
CTCF,RAD21,SMC3 binding sites


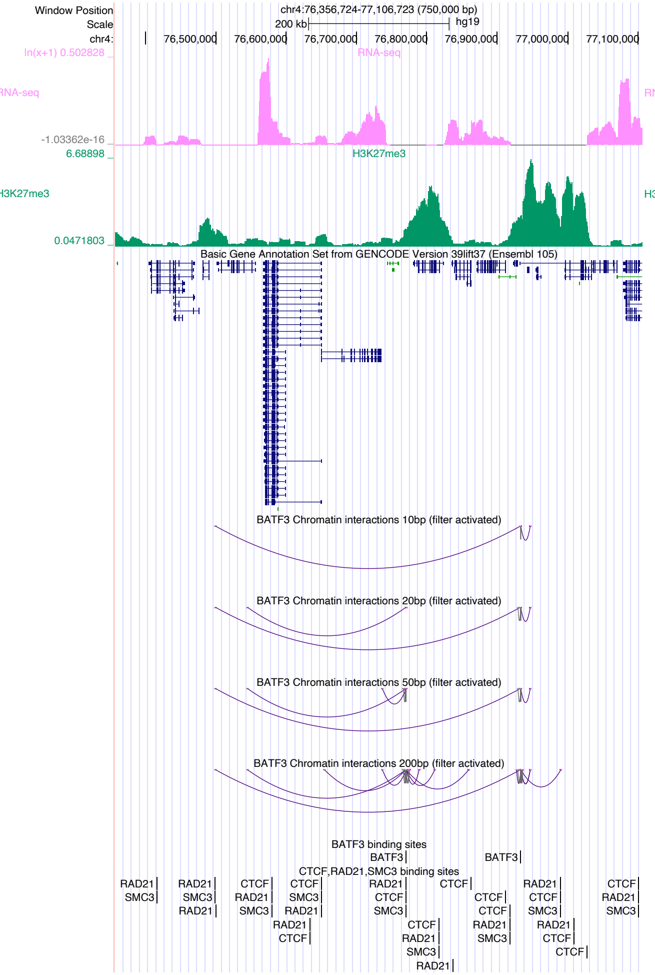

Window Position chr4:76,356,724-77,106,723 (750,000 bp)
Scale 200 kb hg19
chr4: 76,500,000 76,600,000 76,700,000 76,800,000 76,900,000 77,000,000 77,100,000
RNA-seq
H3K27me3
Basic Gene Annotation Set from GENCODE Version 39lift37 (Ensembl 105)
BATF3 Chromatin interactions 10bp (filter activated)
BATF3 Chromatin interactions 20bp (filter activated)
BATF3 Chromatin interactions 50bp (filter activated)
BATF3 Chromatin interactions 200bp (filter activated)
BATF3 binding sites
CTCF,RAD21,SMC3 binding sites


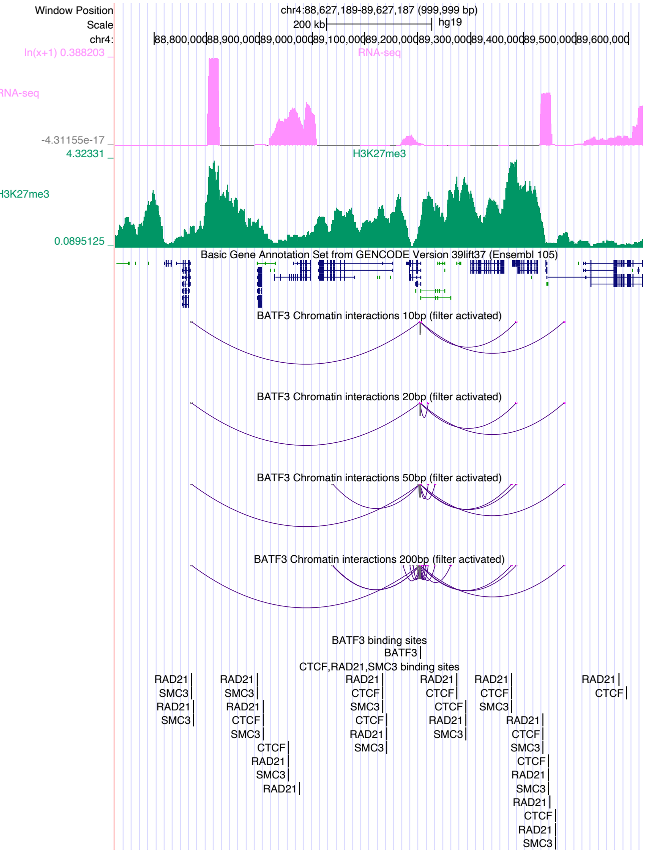

Window Position chr4:88,627,189-89,627,187 (999,999 bp)
Scale 200 kb hg19
chr4: 88,800,000 88,900,000 89,000,000 89,100,000 89,200,000 89,300,000 89,400,000 89,500,000 89,600,000
RNA-seq
H3K27me3
Basic Gene Annotation Set from GENCODE Version 39lift37 (Ensembl 105)
BATF3 Chromatin interactions 10bp (filter activated)
BATF3 Chromatin interactions 20bp (filter activated)
BATF3 Chromatin interactions 50bp (filter activated)
BATF3 Chromatin interactions 200bp (filter activated)
BATF3 binding sites
CTCF,RAD21,SMC3 binding sites

### BCL6

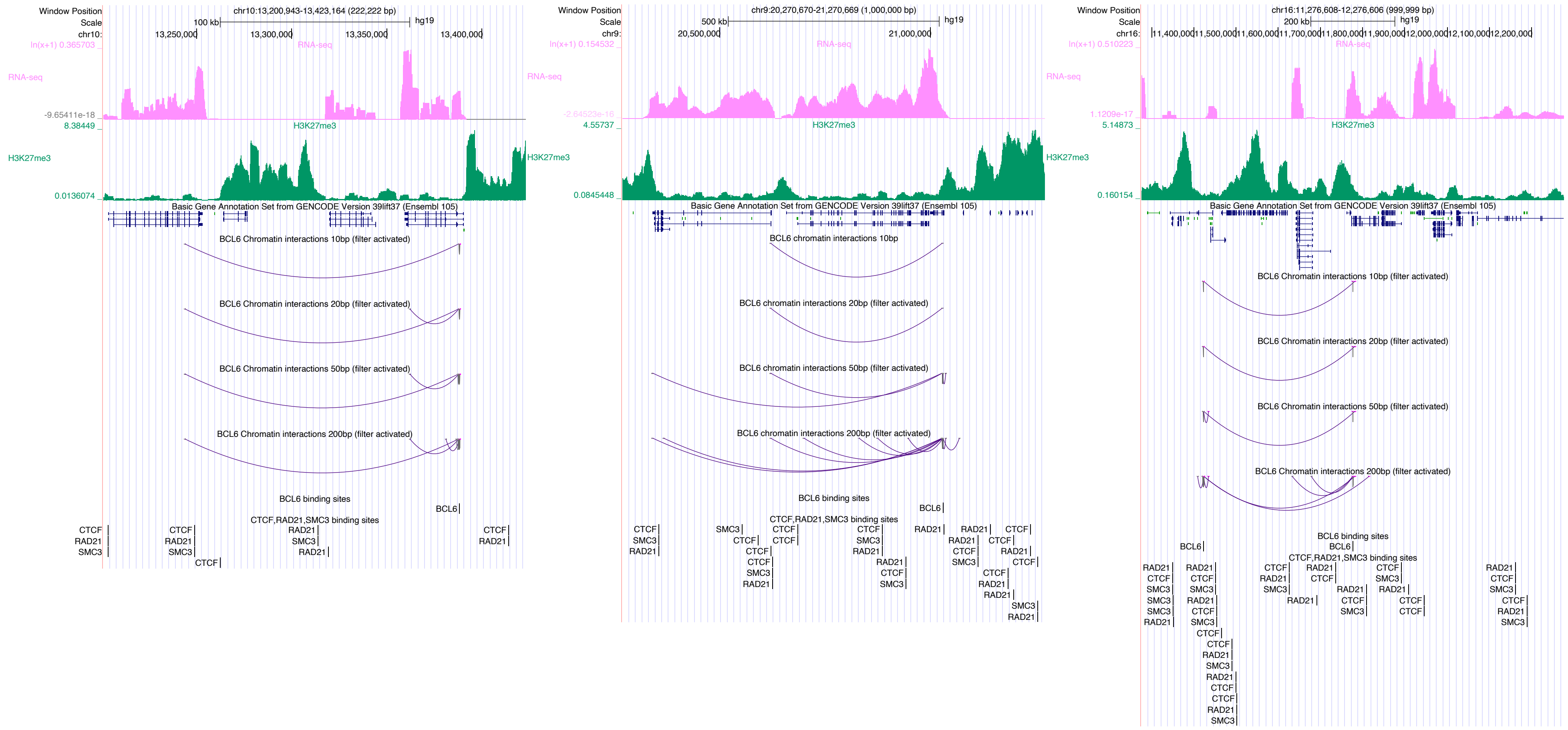

Window Position
Scale
chr10:13,200,943-13,423,164 (222,222 bp)
100 kb
hg19
13,250,000
13,300,000
13,350,000
13,400,000
RNA-seq
H3K27me3
Basic Gene Annotation Set from GENCODE Version 39lift37 (Ensembl 105)
BCL6 Chromatin interactions 10bp (filter activated)
BCL6 Chromatin interactions 20bp (filter activated)
BCL6 Chromatin interactions 50bp (filter activated)
BCL6 Chromatin interactions 200bp (filter activated)
BCL6 binding sites
CTCF,RAD21,SMC3 binding sites
chr9:20,270,670-21,270,669 (1,000,000 bp)
500 kb
20,500,000
21,000,000
BCL6 chromatin interactions 10bp
BCL6 chromatin interactions 20bp (filter activated)
BCL6 chromatin interactions 50bp (filter activated)
BCL6 chromatin interactions 200bp (filter activated)
chr16:11,276,608-12,276,606 (999,999 bp)
200 kb

### CDX2

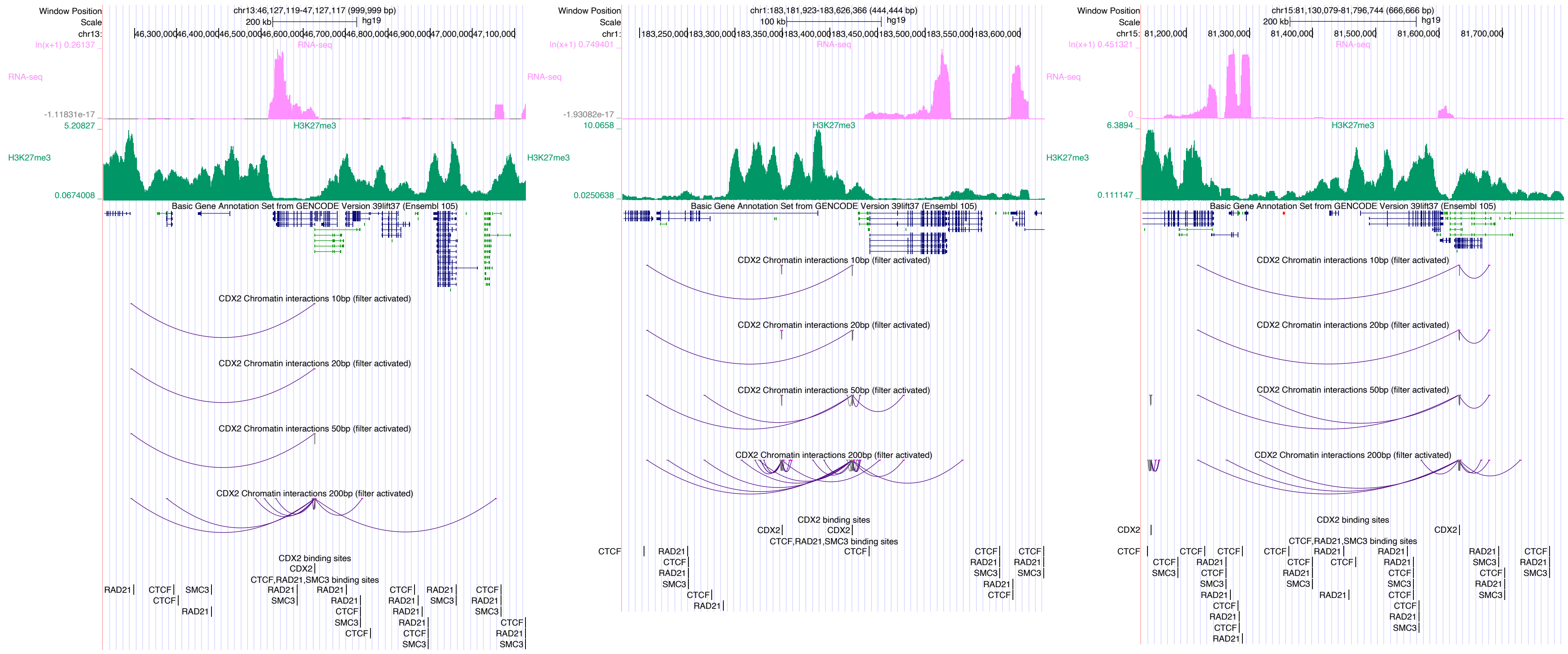
Window Position
Scale
chr13:46,127,119-47,127,117 (999,999 bp)
200 kb
hg19
RNA-seq
H3K27me3
Basic Gene Annotation Set from GENCODE Version 39lift37 (Ensembl 105)
CDX2 Chromatin interactions 10bp (filter activated)
CDX2 Chromatin interactions 20bp (filter activated)
CDX2 Chromatin interactions 50bp (filter activated)
CDX2 Chromatin interactions 200bp (filter activated)
CDX2 binding sites
CTCF,RAD21,SMC3 binding sites
chr1:183,181,923-183,626,366 (444,444 bp)
100 kb
chr15:81,130,079-81,796,744 (666,666 bp)
200 kb

### CEBPA

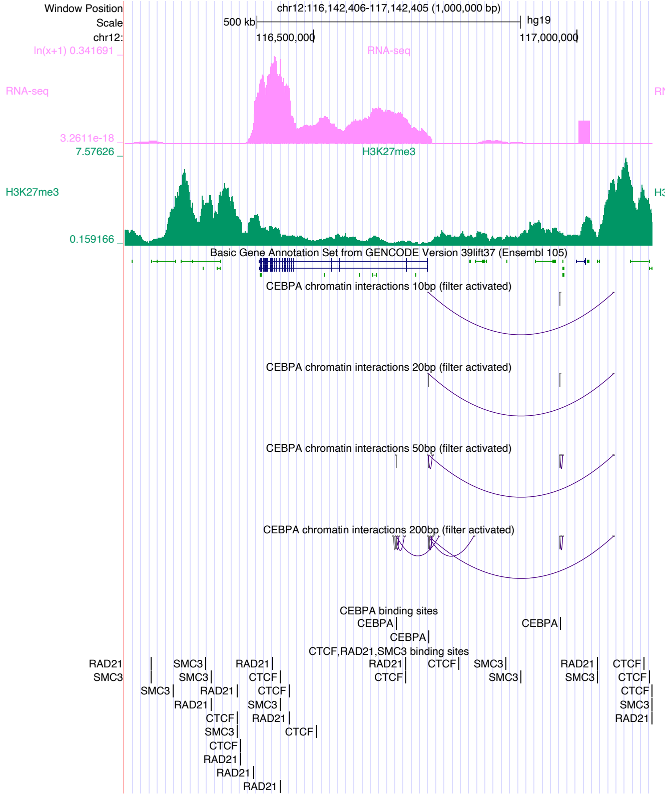

Window Position
chr12:116,142,406-117,142,405 (1,000,000 bp)
Scale
500 kb
hg19
chr12:
116,500,000
117,000,000
RNA-seq
H3K27me3
Basic Gene Annotation Set from GENCODE Version 39lift37 (Ensembl 105)
CEBPA chromatin interactions 10bp (filter activated)
CEBPA chromatin interactions 20bp (filter activated)
CEBPA chromatin interactions 50bp (filter activated)
CEBPA chromatin interactions 200bp (filter activated)
CEBPA binding sites
CTCF,RAD21,SMC3 binding sites


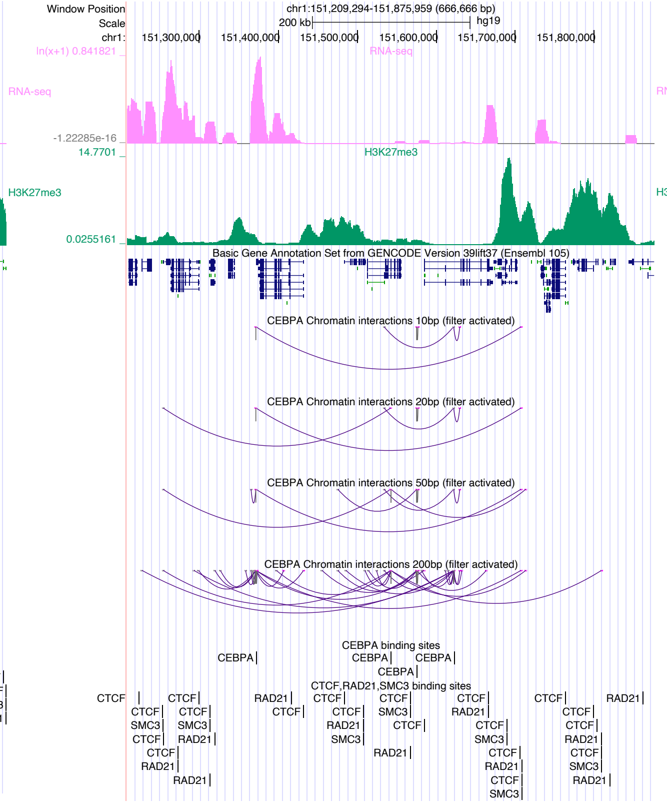

Window Position
chr1:151,209,294-151,875,959 (666,666 bp)
Scale
200 kb
hg19
chr1:
151,300,000
151,400,000
151,500,000
151,600,000
151,700,000
151,800,000
RNA-seq
H3K27me3
Basic Gene Annotation Set from GENCODE Version 39lift37 (Ensembl 105)
CEBPA Chromatin interactions 10bp (filter activated)
CEBPA Chromatin interactions 20bp (filter activated)
CEBPA Chromatin interactions 50bp (filter activated)
CEBPA Chromatin interactions 200bp (filter activated)
CEBPA binding sites
CTCF,RAD21,SMC3 binding sites


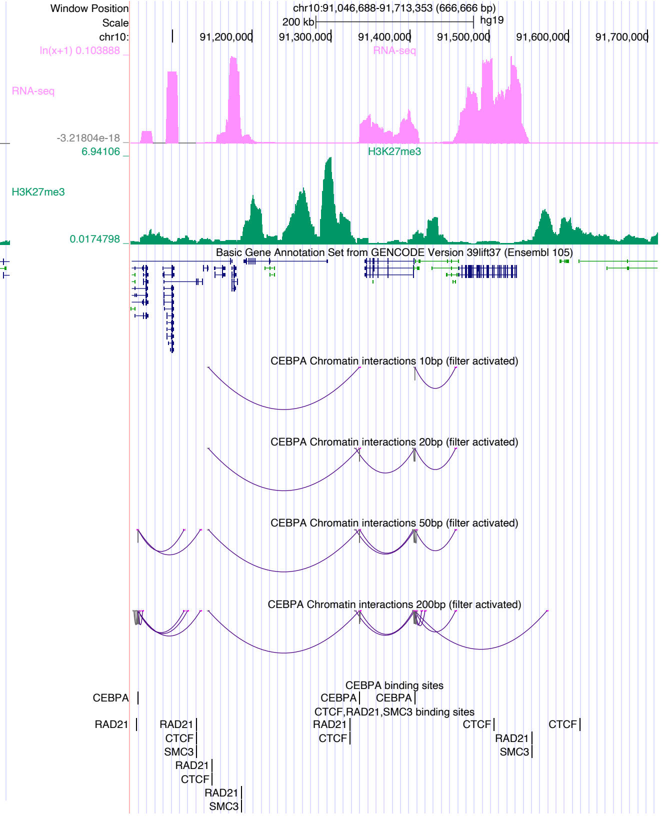

Window Position
chr10:91,046,688-91,713,353 (666,666 bp)
Scale
200 kb
hg19
chr10:
91,200,000
91,300,000
91,400,000
91,500,000
91,600,000
91,700,000
RNA-seq
H3K27me3
Basic Gene Annotation Set from GENCODE Version 39lift37 (Ensembl 105)
CEBPA Chromatin interactions 10bp (filter activated)
CEBPA Chromatin interactions 20bp (filter activated)
CEBPA Chromatin interactions 50bp (filter activated)
CEBPA Chromatin interactions 200bp (filter activated)
CEBPA binding sites
CTCF,RAD21,SMC3 binding sites

### CLOCK

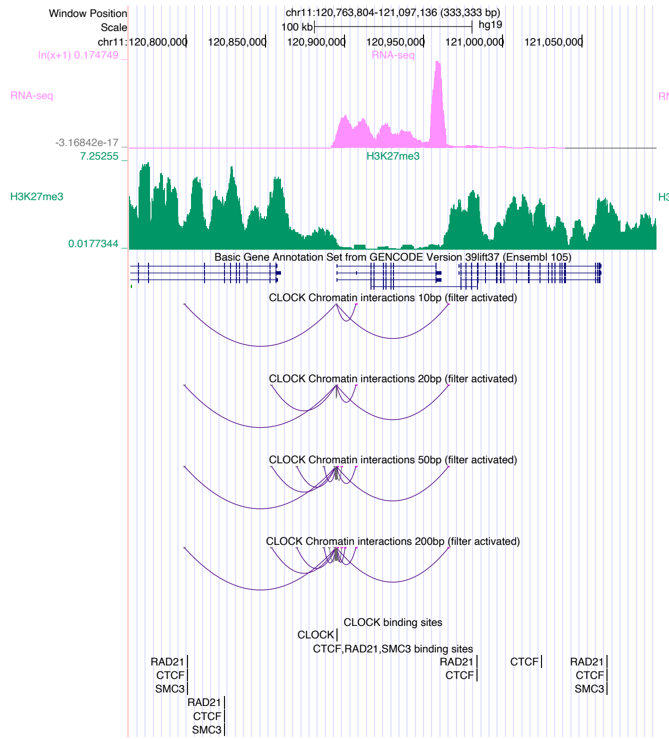

Window Position
Scale
chr11:120,763,804-121,097,136 (333,333 bp)
100 kb
hg19
chr11: 120,800,000 120,850,000 120,900,000 120,950,000 121,000,000 121,050,000
ln(x+1) 0.174749
RNA-seq
-3.16842e-17
7.25255
H3K27me3
0.0177344
Basic Gene Annotation Set from GENCODE Version 39lift37 (Ensembl 105)
CLOCK Chromatin interactions 10bp (filter activated)
CLOCK Chromatin interactions 20bp (filter activated)
CLOCK Chromatin interactions 50bp (filter activated)
CLOCK Chromatin interactions 200bp (filter activated)
CLOCK binding sites
CLOCK
CTCF,RAD21,SMC3 binding sites
RAD21
CTCF
SMC3


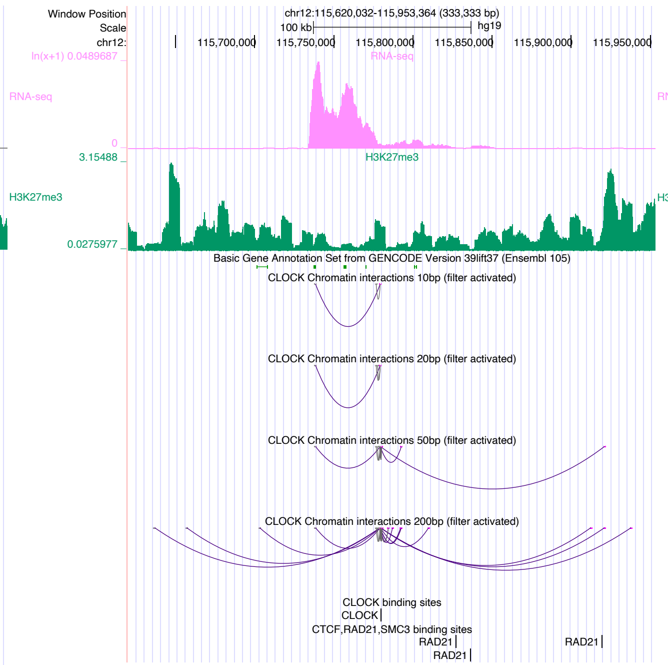

Window Position
Scale
chr12:115,620,032-115,953,364 (333,333 bp)
100 kb
hg19
chr12: 115,700,000 115,750,000 115,800,000 115,850,000 115,900,000 115,950,000
ln(x+1) 0.0489687
RNA-seq
0
3.15488
H3K27me3
0.0275977
Basic Gene Annotation Set from GENCODE Version 39lift37 (Ensembl 105)
CLOCK Chromatin interactions 10bp (filter activated)
CLOCK Chromatin interactions 20bp (filter activated)
CLOCK Chromatin interactions 50bp (filter activated)
CLOCK Chromatin interactions 200bp (filter activated)
CLOCK binding sites
CLOCK
CTCF,RAD21,SMC3 binding sites
RAD21


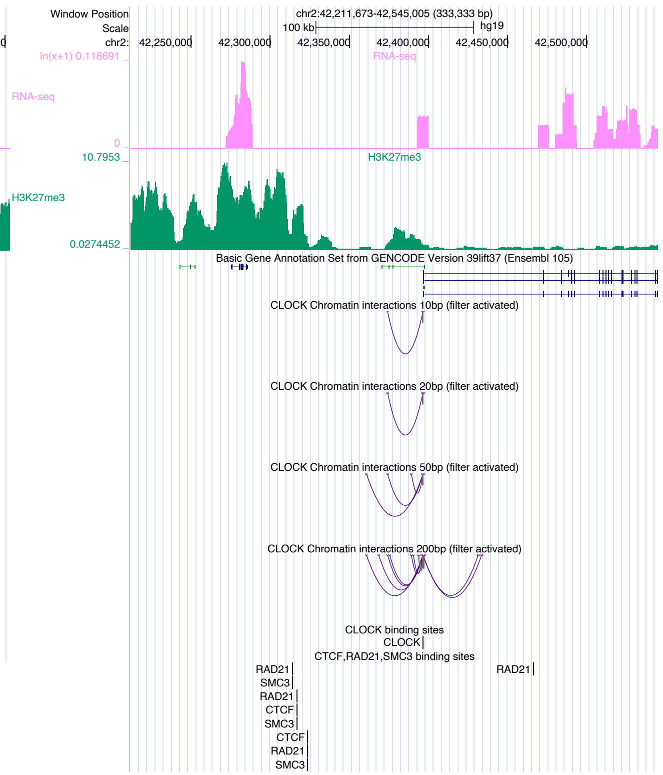

Window Position
Scale
chr2:42,211,673-42,545,005 (333,333 bp)
100 kb
hg19
chr2: 42,250,000 42,300,000 42,350,000 42,400,000 42,450,000 42,500,000
ln(x+1) 0.118691
RNA-seq
0
10.7953
H3K27me3
0.0274452
Basic Gene Annotation Set from GENCODE Version 39lift37 (Ensembl 105)
CLOCK Chromatin interactions 10bp (filter activated)
CLOCK Chromatin interactions 20bp (filter activated)
CLOCK Chromatin interactions 50bp (filter activated)
CLOCK Chromatin interactions 200bp (filter activated)
CLOCK binding sites
CLOCK
CTCF,RAD21,SMC3 binding sites
RAD21
SMC3
CTCF

### CREB5

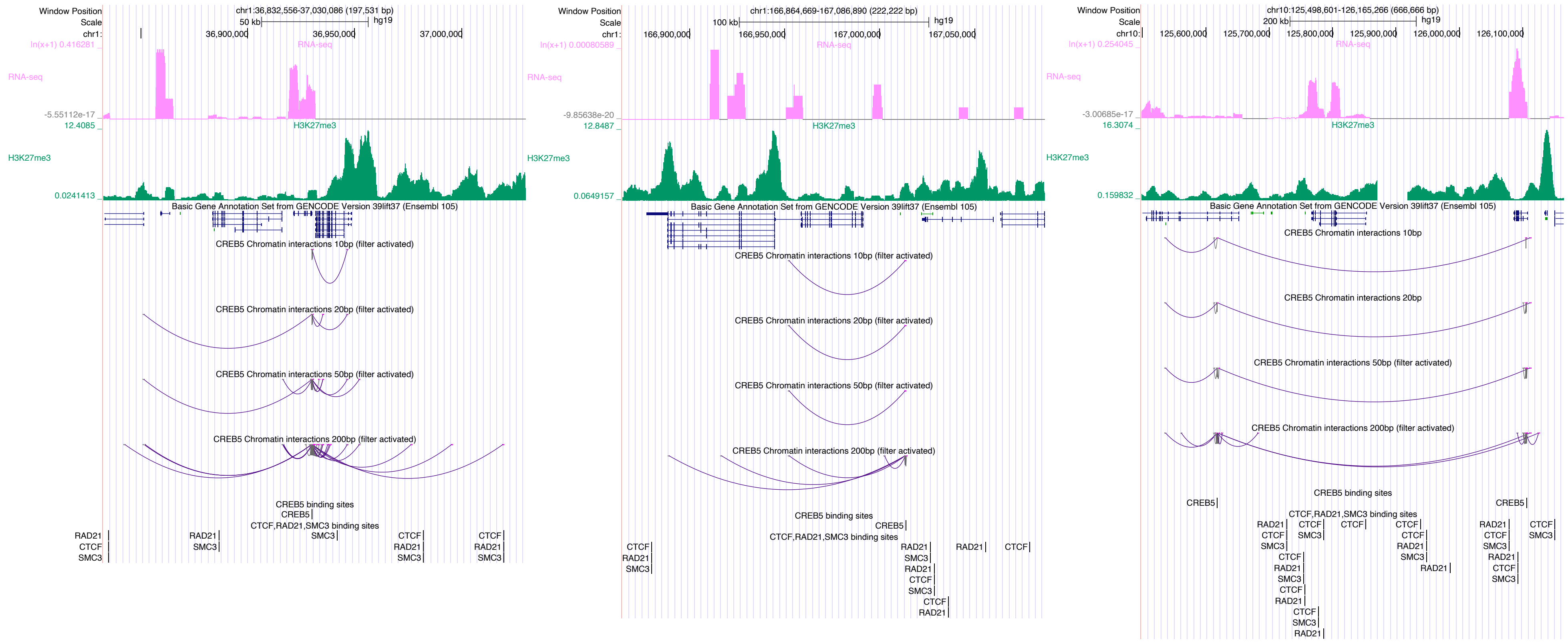
Window Position
chr1:36,832,556-37,030,086 (197,531 bp)
Scale
50 kb
hg19
RNA-seq
H3K27me3
Basic Gene Annotation Set from GENCODE Version 39lift37 (Ensembl 105)
CREB5 Chromatin interactions 10bp (filter activated)
CREB5 Chromatin interactions 20bp (filter activated)
CREB5 Chromatin interactions 50bp (filter activated)
CREB5 Chromatin interactions 200bp (filter activated)
CREB5 binding sites
CTCF,RAD21,SMC3 binding sites
chr1:166,864,669-167,086,890 (222,222 bp)
100 kb
chr10:125,498,601-126,165,266 (666,666 bp)
200 kb
CREB5 Chromatin interactions 20bp
CREB5 Chromatin interactions 50bp (filter activated)
CREB5 Chromatin interactions 200bp (filter activated)

### CTCFL

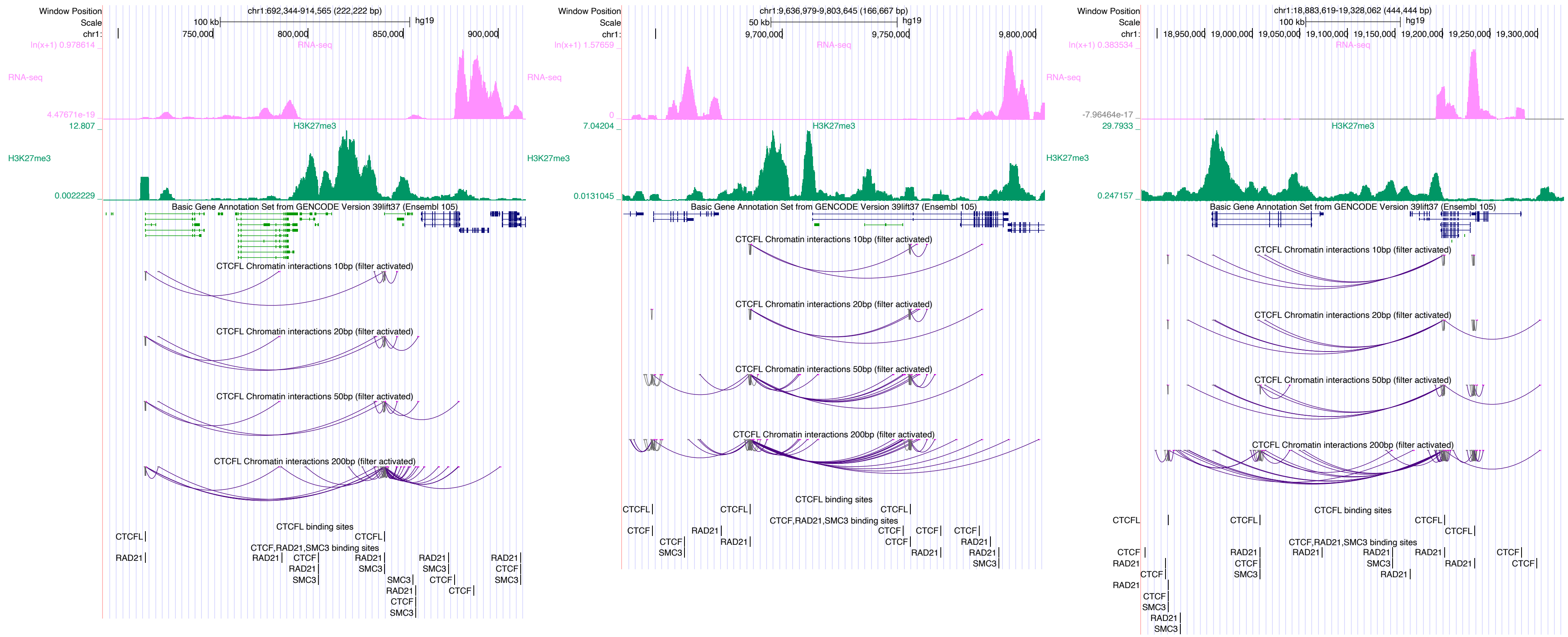

Window Position
Scale
chr1:
chr1:692,344-914,565 (222,222 bp)
100 kb
hg19
750,000
800,000
850,000
900,000
RNA-seq
H3K27me3
Basic Gene Annotation Set from GENCODE Version 39lift37 (Ensembl 105)
CTCFL Chromatin interactions 10bp (filter activated)
CTCFL Chromatin interactions 20bp (filter activated)
CTCFL Chromatin interactions 50bp (filter activated)
CTCFL Chromatin interactions 200bp (filter activated)
CTCFL binding sites
CTCF,RAD21,SMC3 binding sites
chr1:9,636,979-9,803,645 (166,667 bp)
50 kb
9,700,000
9,750,000
9,800,000
chr1:18,883,619-19,328,062 (444,444 bp)
18,950,000
19,000,000
19,050,000
19,100,000
19,150,000
19,200,000
19,250,000
19,300,000

### DUX4

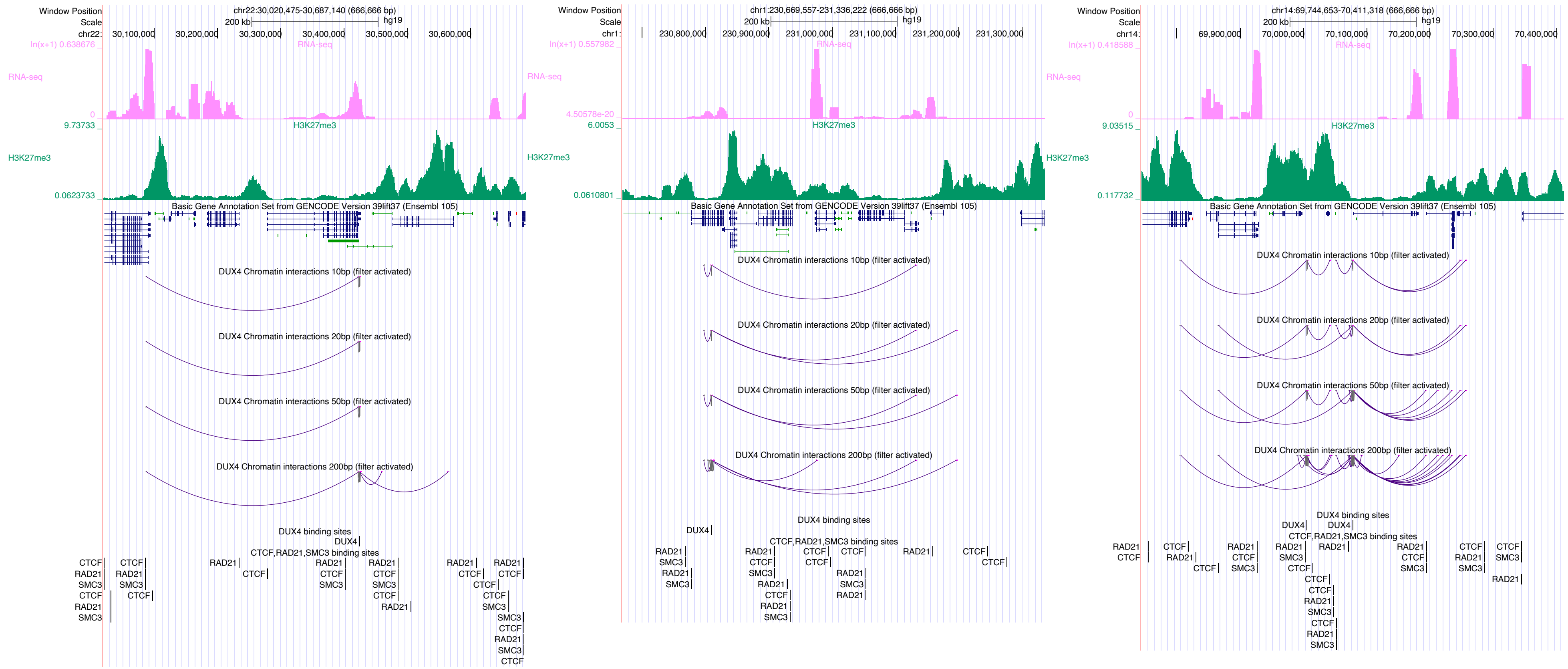

Window Position
Scale
chr22:30,020,475-30,687,140 (666,666 bp)
200 kb
hg19
RNA-seq
H3K27me3
Basic Gene Annotation Set from GENCODE Version 39lift37 (Ensembl 105)
DUX4 Chromatin interactions 10bp (filter activated)
DUX4 Chromatin interactions 20bp (filter activated)
DUX4 Chromatin interactions 50bp (filter activated)
DUX4 Chromatin interactions 200bp (filter activated)
DUX4 binding sites
CTCF,RAD21,SMC3 binding sites
chr1:230,669,557-231,336,222 (666,666 bp)
chr14:69,744,653-70,411,318 (666,666 bp)

### EBF1

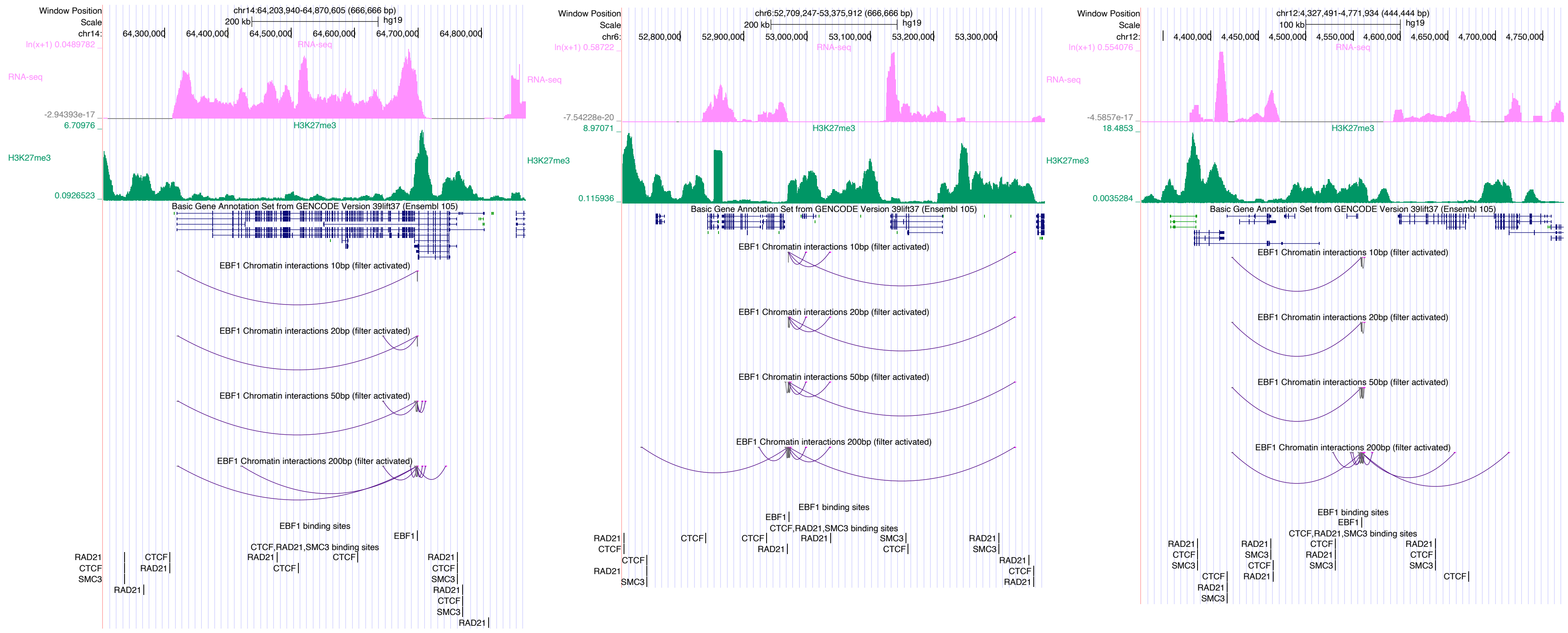

### EGR1

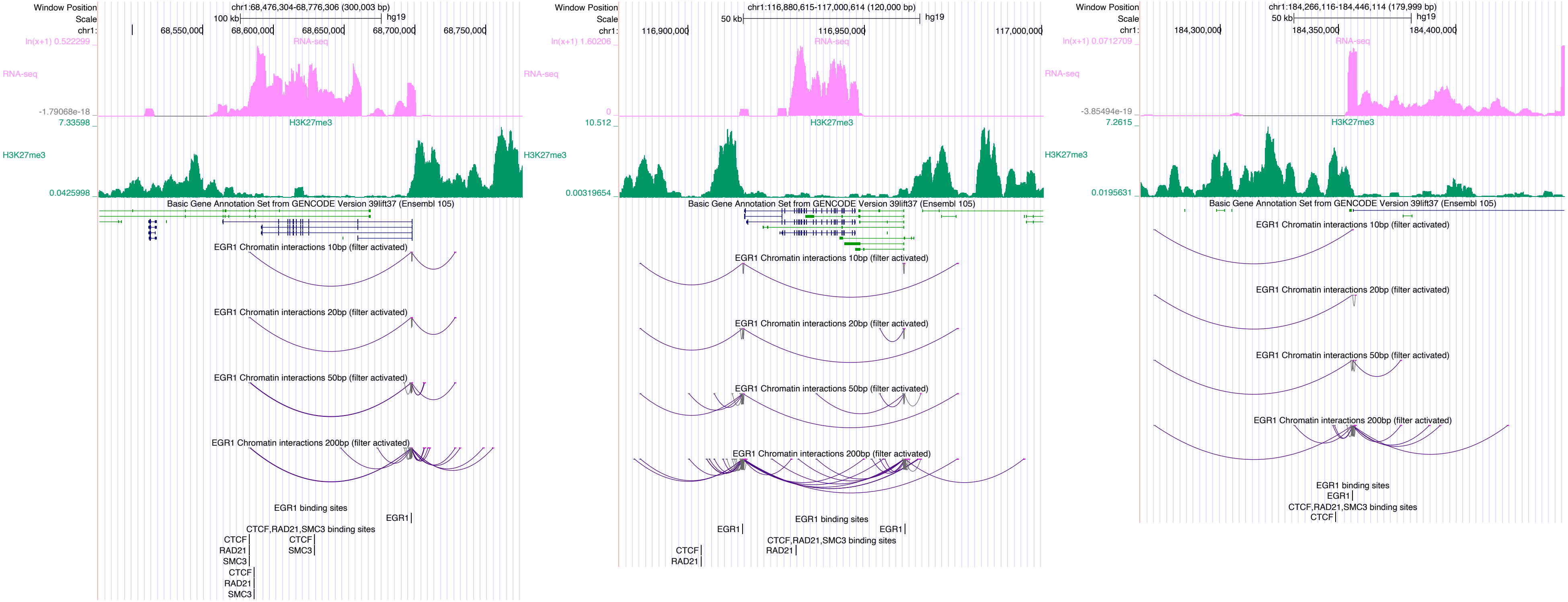

Window Position
Scale
chr1:
chr1:68,476,304-68,776,306 (300,003 bp)
100 kb
hg19
68,550,000
68,600,000
68,650,000
68,700,000
68,750,000
ln(x+1) 0.522299
RNA-seq
-1.79068e-18
7.33598
H3K27me3
0.0425998
Basic Gene Annotation Set from GENCODE Version 39lift37 (Ensembl 105)
EGR1 Chromatin interactions 10bp (filter activated)
EGR1 Chromatin interactions 20bp (filter activated)
EGR1 Chromatin interactions 50bp (filter activated)
EGR1 Chromatin interactions 200bp (filter activated)
EGR1 binding sites
EGR1
CTCF,RAD21,SMC3 binding sites
CTCF
RAD21
SMC3
chr1:116,880,615-117,000,614 (120,000 bp)
50 kb
116,900,000
116,950,000
117,000,000
ln(x+1) 1.60206
0
10.512
0.00319654
chr1:184,266,116-184,446,114 (179,999 bp)
184,300,000
184,350,000
184,400,000
ln(x+1) 0.0712709
-3.85494e-19
7.2615
0.0195631

### EGR2

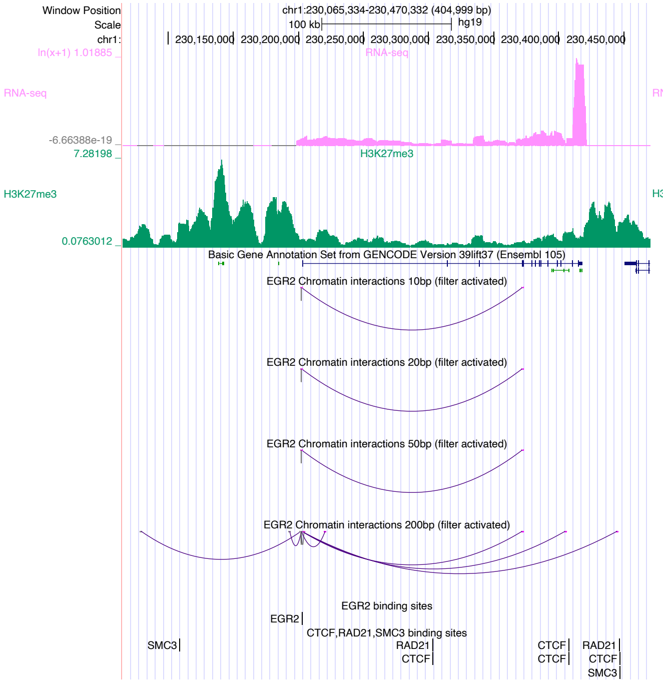
Window Position
chr1:230,065,334-230,470,332 (404,999 bp)
Scale
100 kb
hg19
chr1:
230,150,000
230,200,000
230,250,000
230,300,000
230,350,000
230,400,000
230,450,000
ln(x+1) 1.01885
RNA-seq
-6.66388e-19
7.28198
H3K27me3
0.0763012
Basic Gene Annotation Set from GENCODE Version 39lift37 (Ensembl 105)
EGR2 Chromatin interactions 10bp (filter activated)
EGR2 Chromatin interactions 20bp (filter activated)
EGR2 Chromatin interactions 50bp (filter activated)
EGR2 Chromatin interactions 200bp (filter activated)
EGR2 binding sites
EGR2
CTCF,RAD21,SMC3 binding sites
SMC3
RAD21
CTCF
CTCF
CTCF
RAD21
CTCF
SMC3

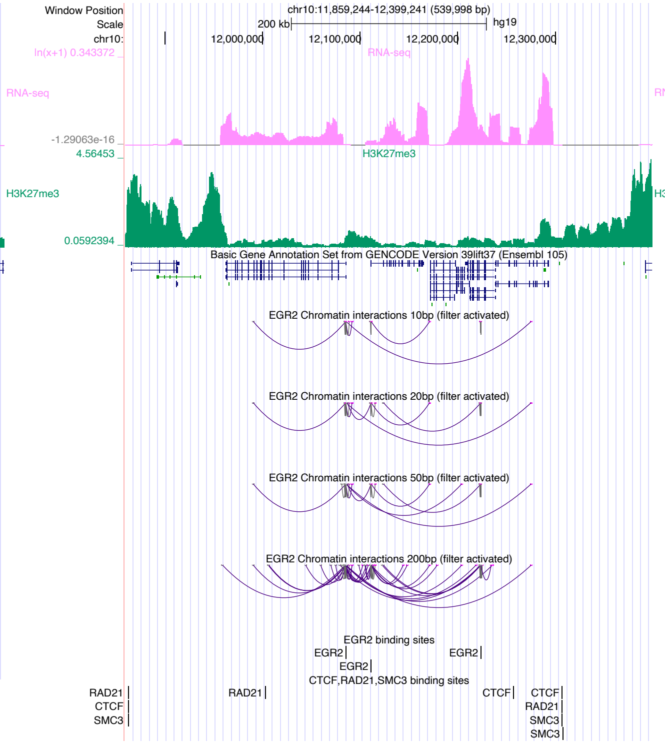
Window Position
chr10:11,859,244-12,399,241 (539,998 bp)
Scale
200 kb
hg19
chr10:
12,000,000
12,100,000
12,200,000
12,300,000
ln(x+1) 0.343372
RNA-seq
-1.29063e-16
4.56453
H3K27me3
0.0592394
Basic Gene Annotation Set from GENCODE Version 39lift37 (Ensembl 105)
EGR2 Chromatin interactions 10bp (filter activated)
EGR2 Chromatin interactions 20bp (filter activated)
EGR2 Chromatin interactions 50bp (filter activated)
EGR2 Chromatin interactions 200bp (filter activated)
EGR2 binding sites
EGR2
EGR2
EGR2
CTCF,RAD21,SMC3 binding sites
RAD21
CTCF
SMC3
RAD21
CTCF
CTCF
RAD21
SMC3
SMC3

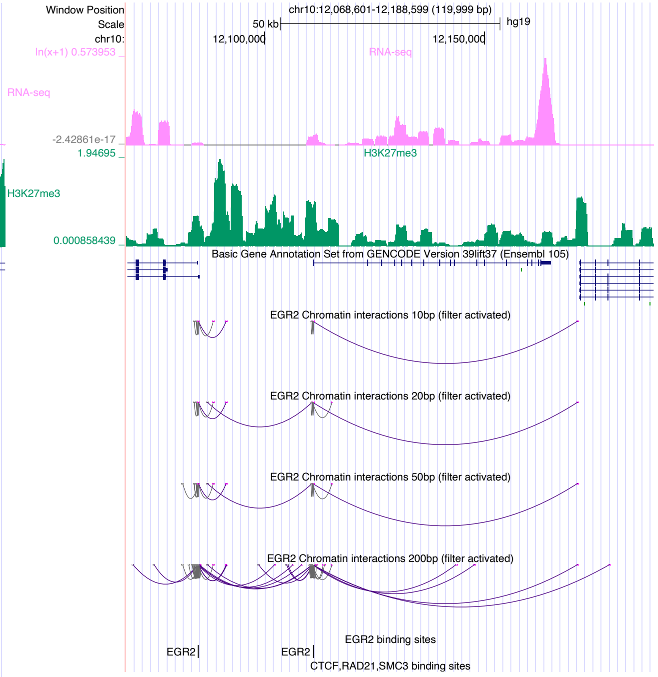
Window Position
chr10:12,068,601-12,188,599 (119,999 bp)
Scale
50 kb
hg19
chr10:
12,100,000
12,150,000
ln(x+1) 0.573953
RNA-seq
-2.42861e-17
1.94695
H3K27me3
0.000858439
Basic Gene Annotation Set from GENCODE Version 39lift37 (Ensembl 105)
EGR2 Chromatin interactions 10bp (filter activated)
EGR2 Chromatin interactions 20bp (filter activated)
EGR2 Chromatin interactions 50bp (filter activated)
EGR2 Chromatin interactions 200bp (filter activated)
EGR2 binding sites
EGR2
EGR2
CTCF,RAD21,SMC3 binding sites

### EGR3

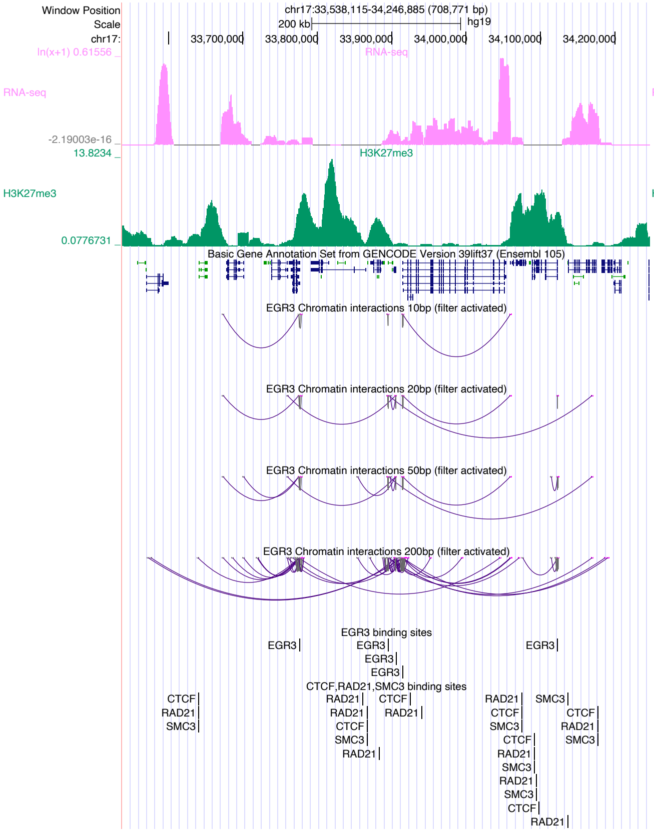

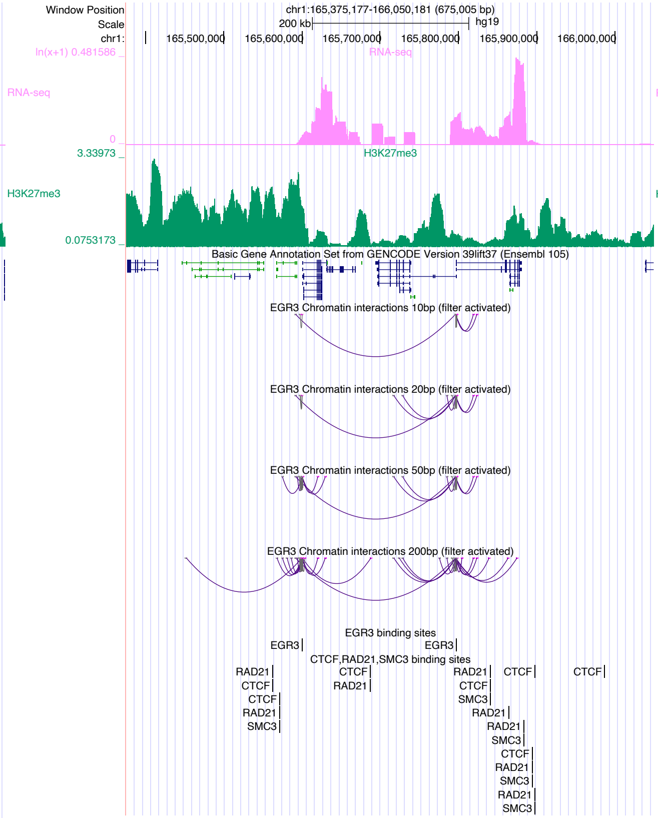

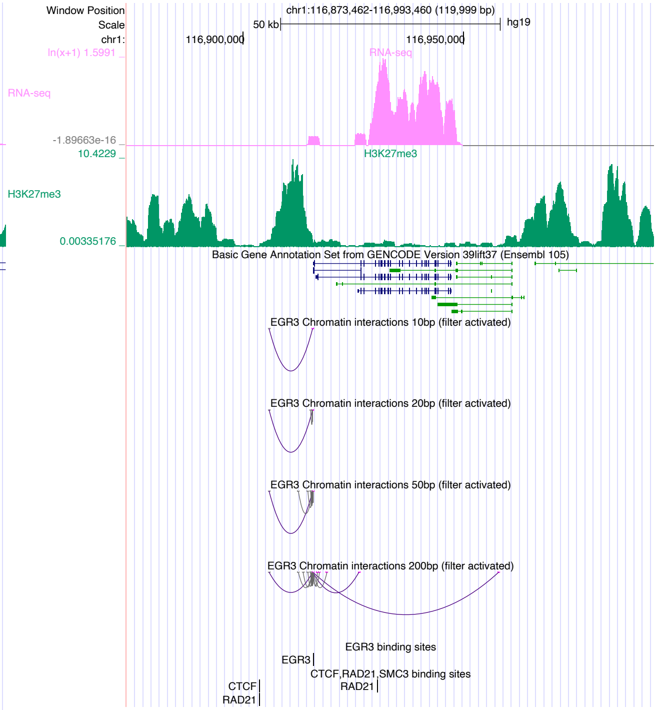

### ELK4

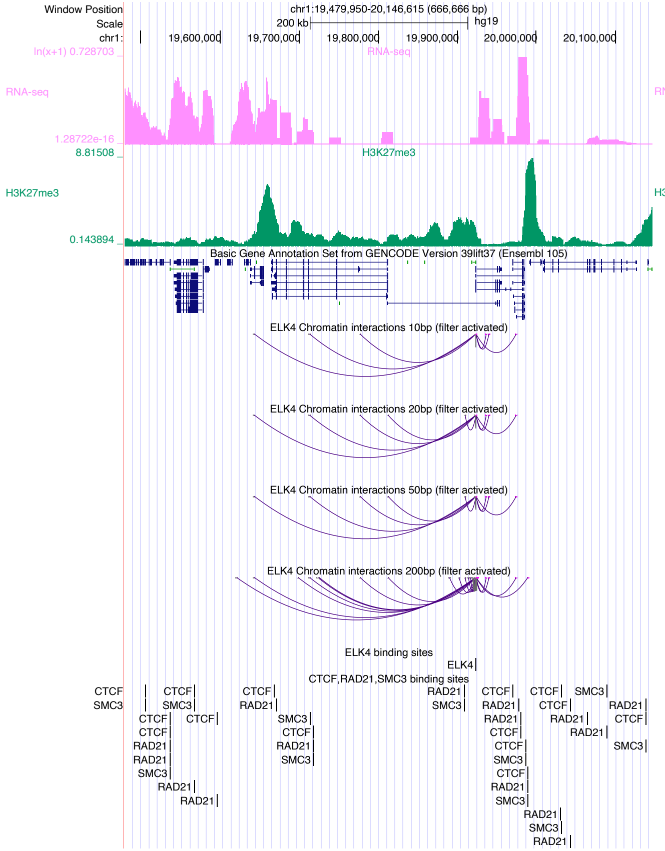

Window Position
chr1:19,479,950-20,146,615 (666,666 bp)
Scale
200 kb
hg19
chr1:
19,600,000
19,700,000
19,800,000
19,900,000
20,000,000
20,100,000
RNA-seq
H3K27me3
Basic Gene Annotation Set from GENCODE Version 39lift37 (Ensembl 105)
ELK4 Chromatin interactions 10bp (filter activated)
ELK4 Chromatin interactions 20bp (filter activated)
ELK4 Chromatin interactions 50bp (filter activated)
ELK4 Chromatin interactions 200bp (filter activated)
ELK4 binding sites
CTCF,RAD21,SMC3 binding sites


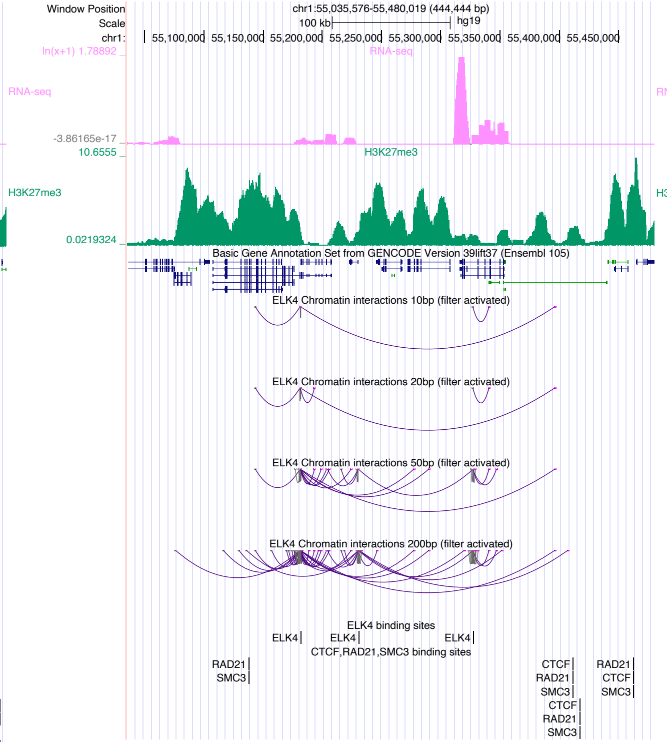

Window Position
chr1:55,035,576-55,480,019 (444,444 bp)
Scale
100 kb
hg19
chr1:
55,100,000
55,150,000
55,200,000
55,250,000
55,300,000
55,350,000
55,400,000
55,450,000
RNA-seq
H3K27me3
Basic Gene Annotation Set from GENCODE Version 39lift37 (Ensembl 105)
ELK4 Chromatin interactions 10bp (filter activated)
ELK4 Chromatin interactions 20bp (filter activated)
ELK4 Chromatin interactions 50bp (filter activated)
ELK4 Chromatin interactions 200bp (filter activated)
ELK4 binding sites
CTCF,RAD21,SMC3 binding sites


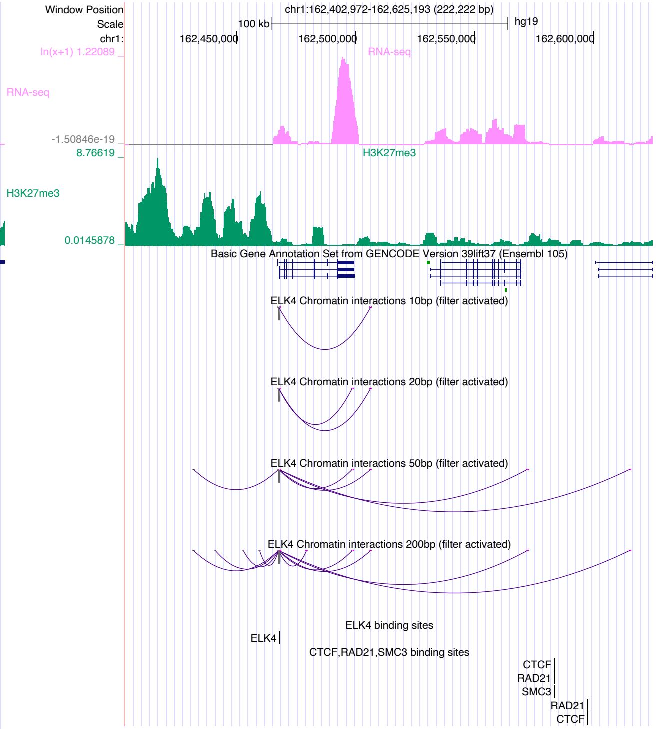

Window Position
chr1:162,402,972-162,625,193 (222,222 bp)
Scale
100 kb
hg19
chr1:
162,450,000
162,500,000
162,550,000
162,600,000
RNA-seq
H3K27me3
Basic Gene Annotation Set from GENCODE Version 39lift37 (Ensembl 105)
ELK4 Chromatin interactions 10bp (filter activated)
ELK4 Chromatin interactions 20bp (filter activated)
ELK4 Chromatin interactions 50bp (filter activated)
ELK4 Chromatin interactions 200bp (filter activated)
ELK4 binding sites
CTCF,RAD21,SMC3 binding sites

# EP300

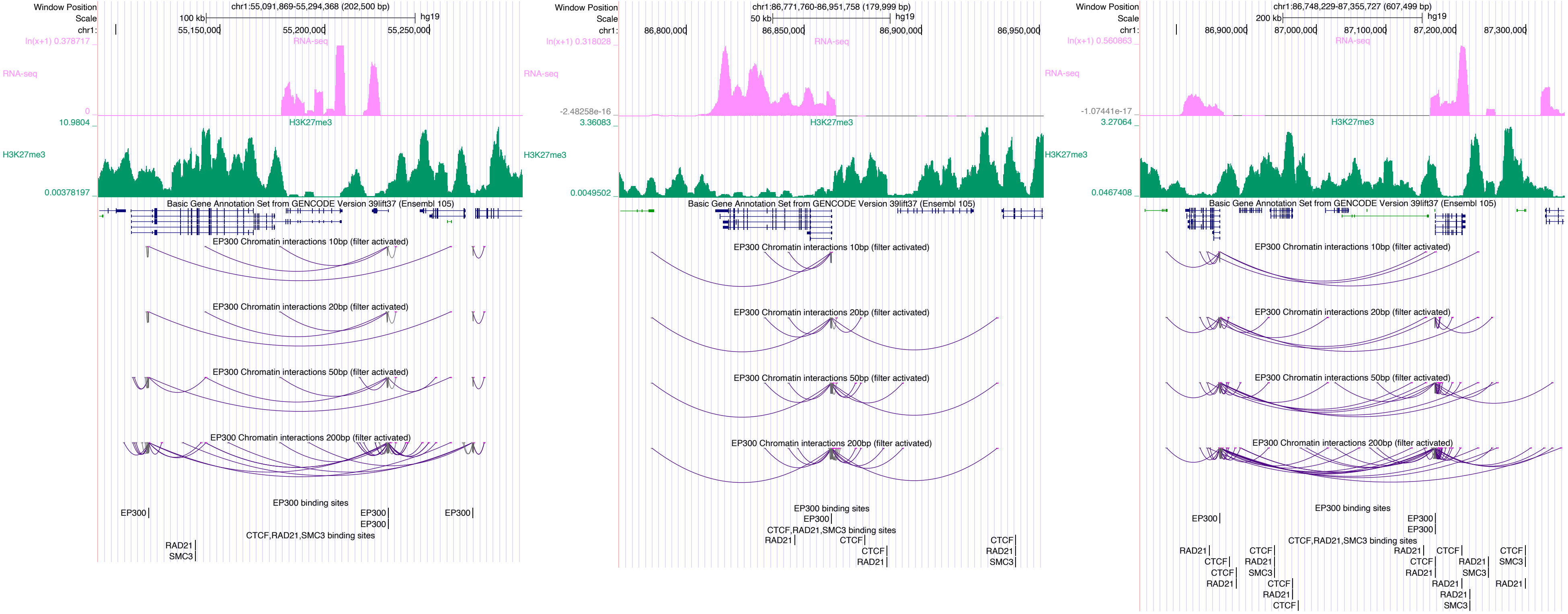

Window Position
Scale
chr1:
chr1:55,091,869-55,294,368 (202,500 bp)
100 kb
hg19
55,150,000
55,200,000
55,250,000
ln(x+1) 0.378717
RNA-seq
0
10.9804
H3K27me3
0.00378197
Basic Gene Annotation Set from GENCODE Version 39lift37 (Ensembl 105)
EP300 Chromatin interactions 10bp (filter activated)
EP300 Chromatin interactions 20bp (filter activated)
EP300 Chromatin interactions 50bp (filter activated)
EP300 Chromatin interactions 200bp (filter activated)
EP300 binding sites
EP300
CTCF,RAD21,SMC3 binding sites
RAD21
SMC3
chr1:86,771,760-86,951,758 (179,999 bp)
50 kb
86,800,000
86,850,000
86,900,000
86,950,000
ln(x+1) 0.318028
-2.48258e-16
3.36083
0.0049502
CTCF
chr1:86,748,229-87,355,727 (607,499 bp)
200 kb
86,900,000
87,000,000
87,100,000
87,200,000
87,300,000
ln(x+1) 0.560863
-1.07441e-17
3.27064
0.0467408

### EPAS1

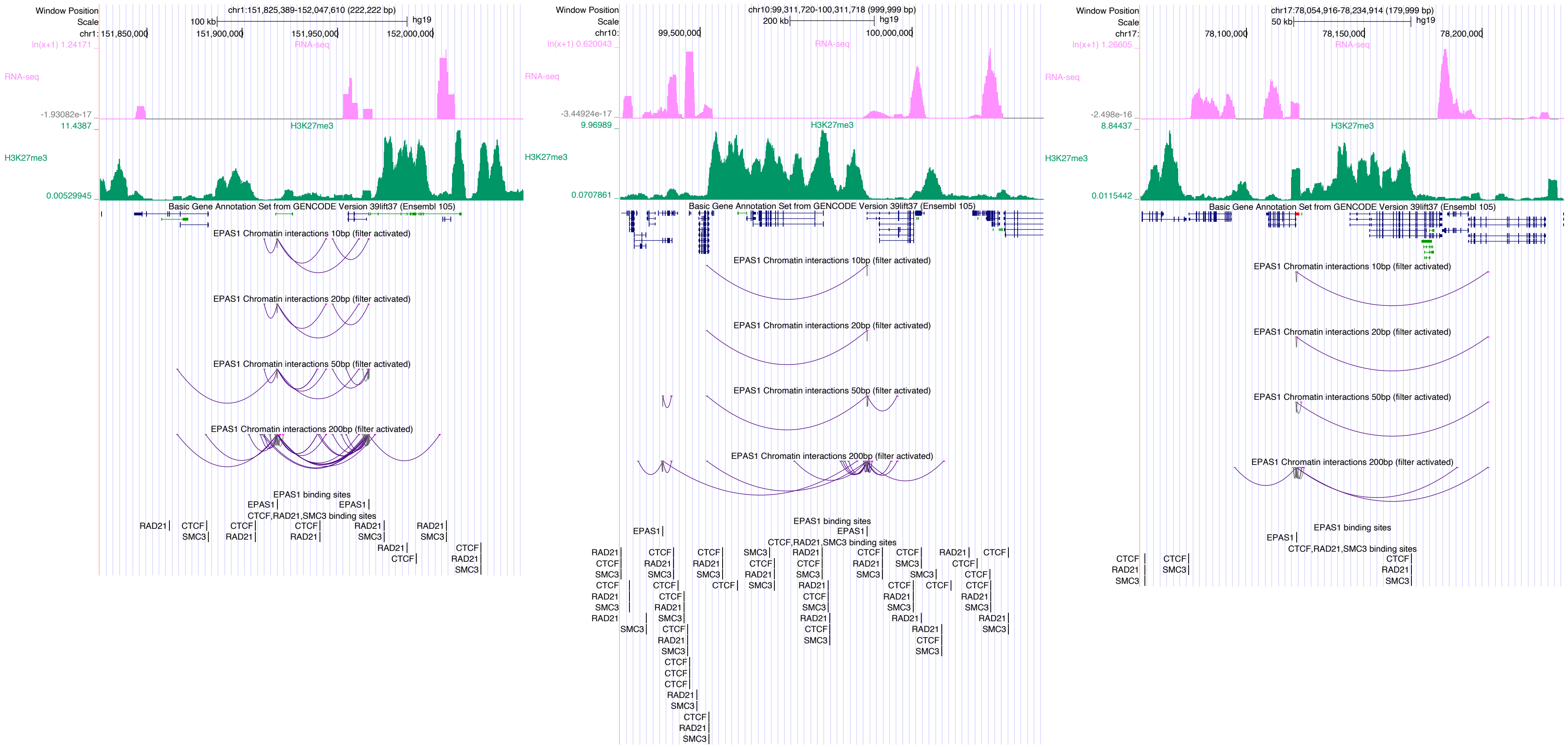

chr1:151,825,389-152,047,610 (222,222 bp)
chr10:99,311,720-100,311,718 (999,999 bp)
chr17:78,054,916-78,234,914 (179,999 bp)
RNA-seq
H3K27me3
Basic Gene Annotation Set from GENCODE Version 39lift37 (Ensembl 105)
EPAS1 Chromatin interactions 10bp (filter activated)
EPAS1 Chromatin interactions 20bp (filter activated)
EPAS1 Chromatin interactions 50bp (filter activated)
EPAS1 Chromatin interactions 200bp (filter activated)
EPAS1 binding sites
CTCF,RAD21,SMC3 binding sites

### ETV1

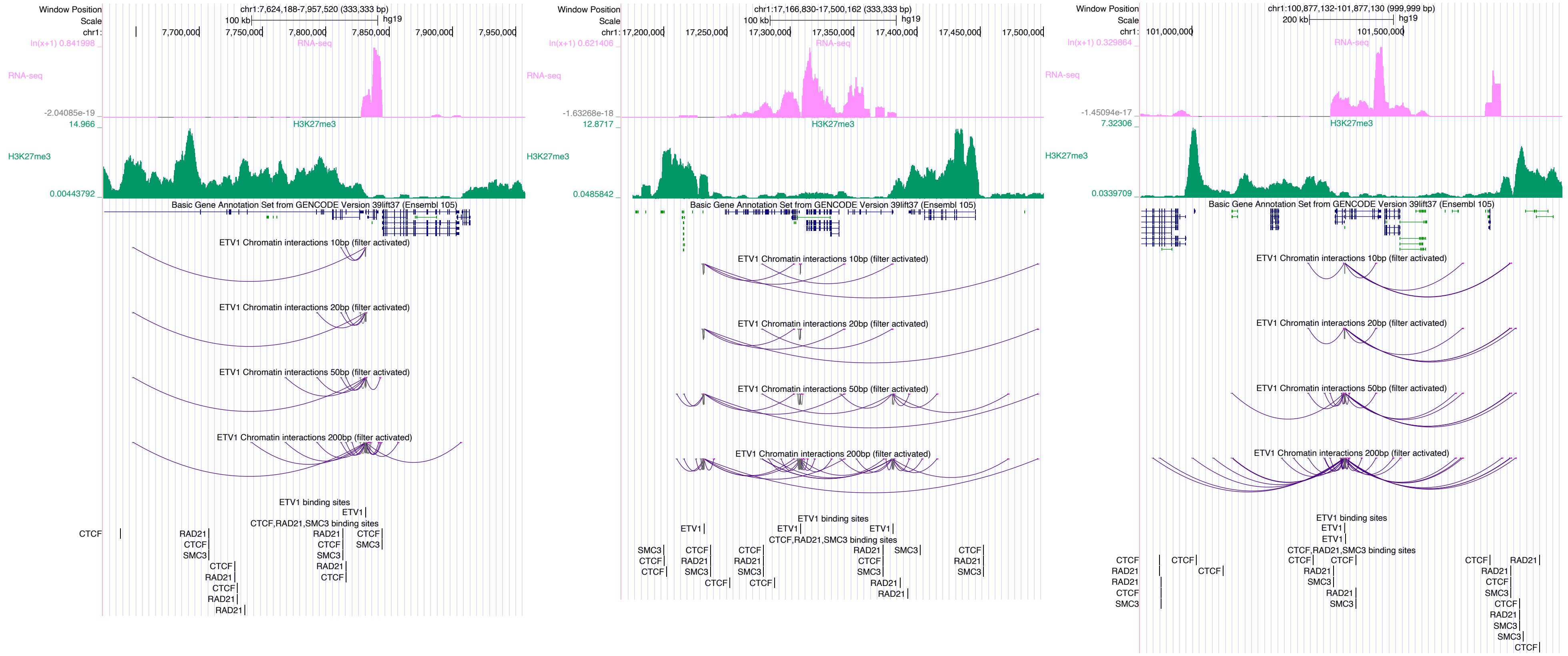
Window Position
Scale
chr1:7,624,188-7,957,520 (333,333 bp)
100 kb
hg19
7,700,000
7,750,000
7,800,000
7,850,000
7,900,000
7,950,000
RNA-seq
ln(x+1) 0.841998
-2.04085e-19
H3K27me3
14.966
0.00443792
Basic Gene Annotation Set from GENCODE Version 39lift37 (Ensembl 105)
ETV1 Chromatin interactions 10bp (filter activated)
ETV1 Chromatin interactions 20bp (filter activated)
ETV1 Chromatin interactions 50bp (filter activated)
ETV1 Chromatin interactions 200bp (filter activated)
ETV1 binding sites
CTCF,RAD21,SMC3 binding sites
chr1:17,166,830-17,500,162 (333,333 bp)
17,200,000
17,250,000
17,300,000
17,350,000
17,400,000
17,450,000
17,500,000
ln(x+1) 0.621406
-1.63268e-18
12.8717
0.0485842
chr1:100,877,132-101,877,130 (999,999 bp)
200 kb
101,000,000
101,500,000
ln(x+1) 0.329864
-1.45094e-17
7.32306
0.0339709
ETV1
CTCF
RAD21
SMC3

### FOS

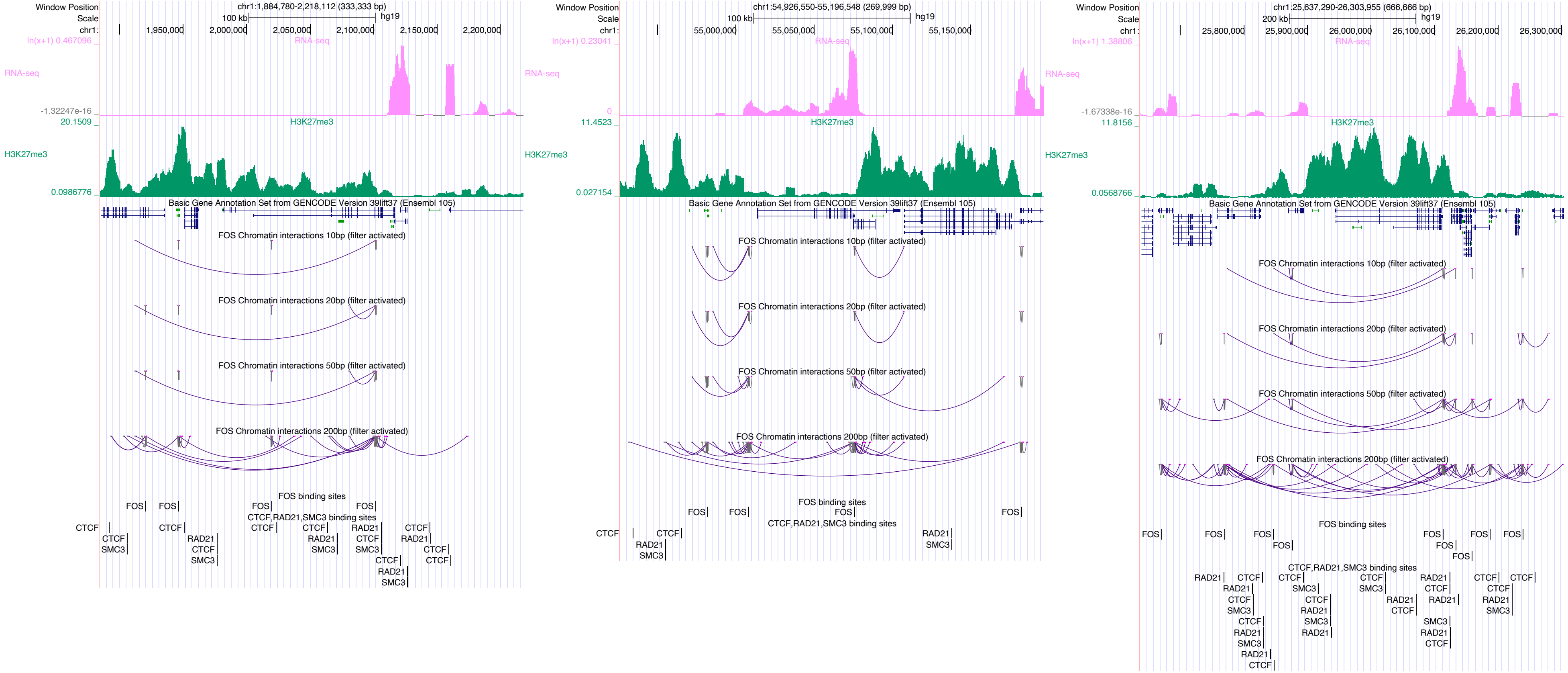
Window Position
chr1:1,884,780-2,218,112 (333,333 bp)
chr1:54,926,550-55,196,548 (269,999 bp)
chr1:25,637,290-26,303,955 (666,666 bp)
RNA-seq
H3K27me3
Basic Gene Annotation Set from GENCODE Version 39lift37 (Ensembl 105)
FOS Chromatin interactions 10bp (filter activated)
FOS Chromatin interactions 20bp (filter activated)
FOS Chromatin interactions 50bp (filter activated)
FOS Chromatin interactions 200bp (filter activated)
FOS binding sites
CTCF,RAD21,SMC3 binding sites

### FOSL2

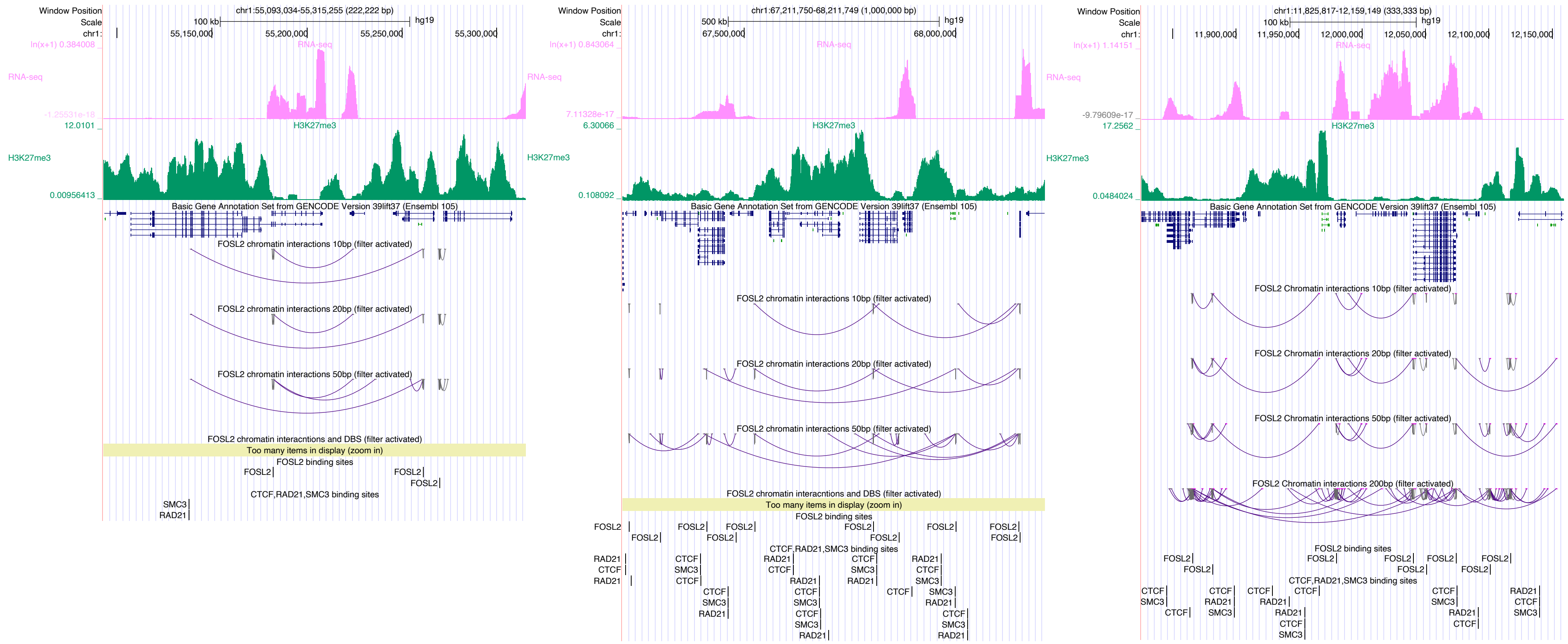

Window Position
chr1:55,093,034-55,315,255 (222,222 bp)
Scale
100 kb
hg19
chr1:
55,150,000
55,200,000
55,250,000
55,300,000
RNA-seq
H3K27me3
Basic Gene Annotation Set from GENCODE Version 39lift37 (Ensembl 105)
FOSL2 chromatin interactions 10bp (filter activated)
FOSL2 chromatin interactions 20bp (filter activated)
FOSL2 chromatin interactions 50bp (filter activated)
FOSL2 chromatin interacntions and DBS (filter activated)
Too many items in display (zoom in)
FOSL2 binding sites
CTCF,RAD21,SMC3 binding sites
Window Position
chr1:67,211,750-68,211,749 (1,000,000 bp)
500 kb
67,500,000
68,000,000
FOSL2 chromatin interactions 10bp (filter activated)
FOSL2 chromatin interactions 20bp (filter activated)
FOSL2 chromatin interactions 50bp (filter activated)
FOSL2 chromatin interacntions and DBS (filter activated)
Too many items in display (zoom in)
FOSL2 binding sites
CTCF,RAD21,SMC3 binding sites
Window Position
chr1:11,825,817-12,159,149 (333,333 bp)
100 kb
11,900,000
11,950,000
12,000,000
12,050,000
12,100,000
12,150,000
FOSL2 Chromatin interactions 10bp (filter activated)
FOSL2 Chromatin interactions 20bp (filter activated)
FOSL2 Chromatin interactions 50bp (filter activated)
FOSL2 Chromatin interactions 200bp (filter activated)
FOSL2 binding sites
CTCF,RAD21,SMC3 binding sites

### FOXA3

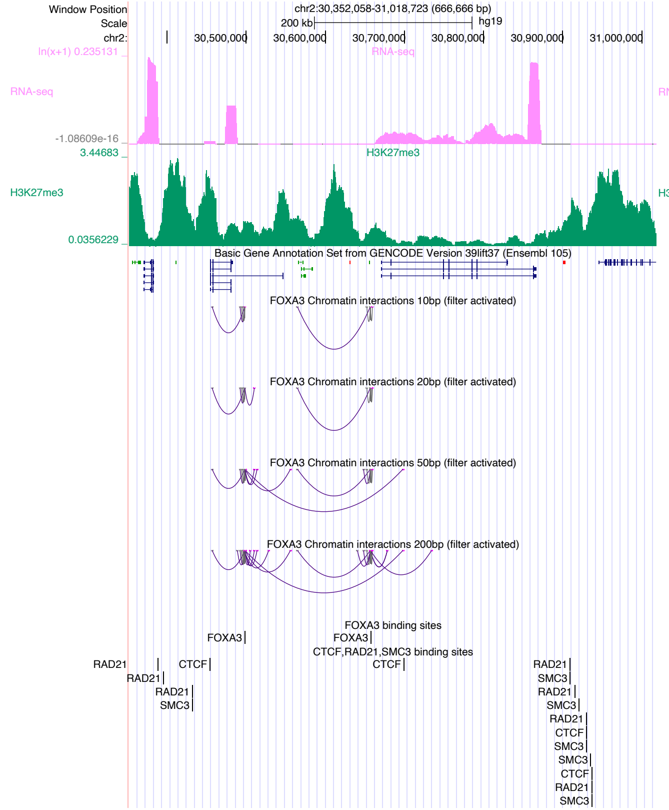

Window Position
chr2:30,352,058-31,018,723 (666,666 bp)
Scale
200 kb
hg19
chr2:
30,500,000
30,600,000
30,700,000
30,800,000
30,900,000
31,000,000
RNA-seq
H3K27me3
Basic Gene Annotation Set from GENCODE Version 39lift37 (Ensembl 105)
FOXA3 Chromatin interactions 10bp (filter activated)
FOXA3 Chromatin interactions 20bp (filter activated)
FOXA3 Chromatin interactions 50bp (filter activated)
FOXA3 Chromatin interactions 200bp (filter activated)
FOXA3 binding sites
CTCF,RAD21,SMC3 binding sites


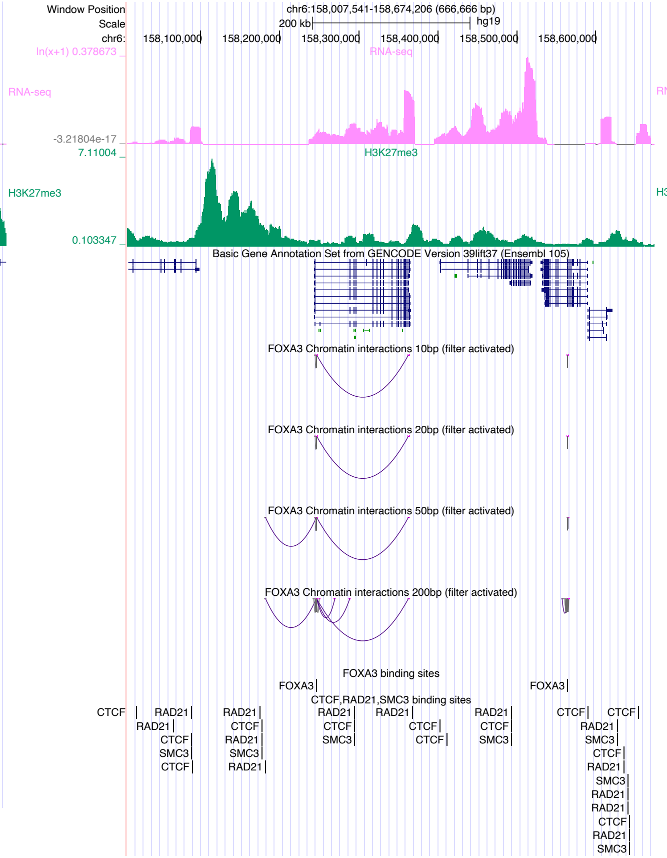

Window Position
chr6:158,007,541-158,674,206 (666,666 bp)
Scale
200 kb
hg19
chr6:
158,100,000
158,200,000
158,300,000
158,400,000
158,500,000
158,600,000
RNA-seq
H3K27me3
Basic Gene Annotation Set from GENCODE Version 39lift37 (Ensembl 105)
FOXA3 Chromatin interactions 10bp (filter activated)
FOXA3 Chromatin interactions 20bp (filter activated)
FOXA3 Chromatin interactions 50bp (filter activated)
FOXA3 Chromatin interactions 200bp (filter activated)
FOXA3 binding sites
CTCF,RAD21,SMC3 binding sites


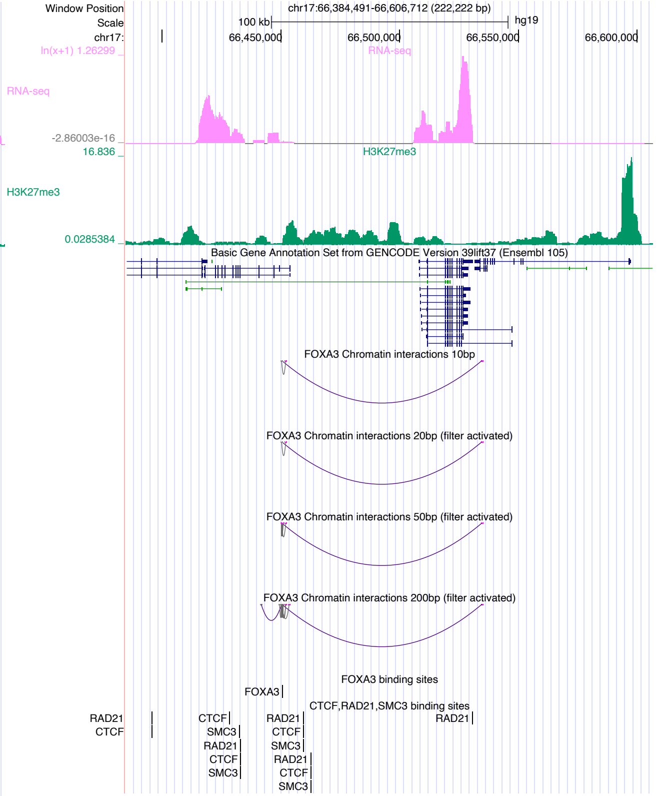

Window Position
chr17:66,384,491-66,606,712 (222,222 bp)
Scale
100 kb
hg19
chr17:
66,450,000
66,500,000
66,550,000
66,600,000
RNA-seq
H3K27me3
Basic Gene Annotation Set from GENCODE Version 39lift37 (Ensembl 105)
FOXA3 Chromatin interactions 10bp
FOXA3 Chromatin interactions 20bp (filter activated)
FOXA3 Chromatin interactions 50bp (filter activated)
FOXA3 Chromatin interactions 200bp (filter activated)
FOXA3 binding sites
CTCF,RAD21,SMC3 binding sites

### FOXH1

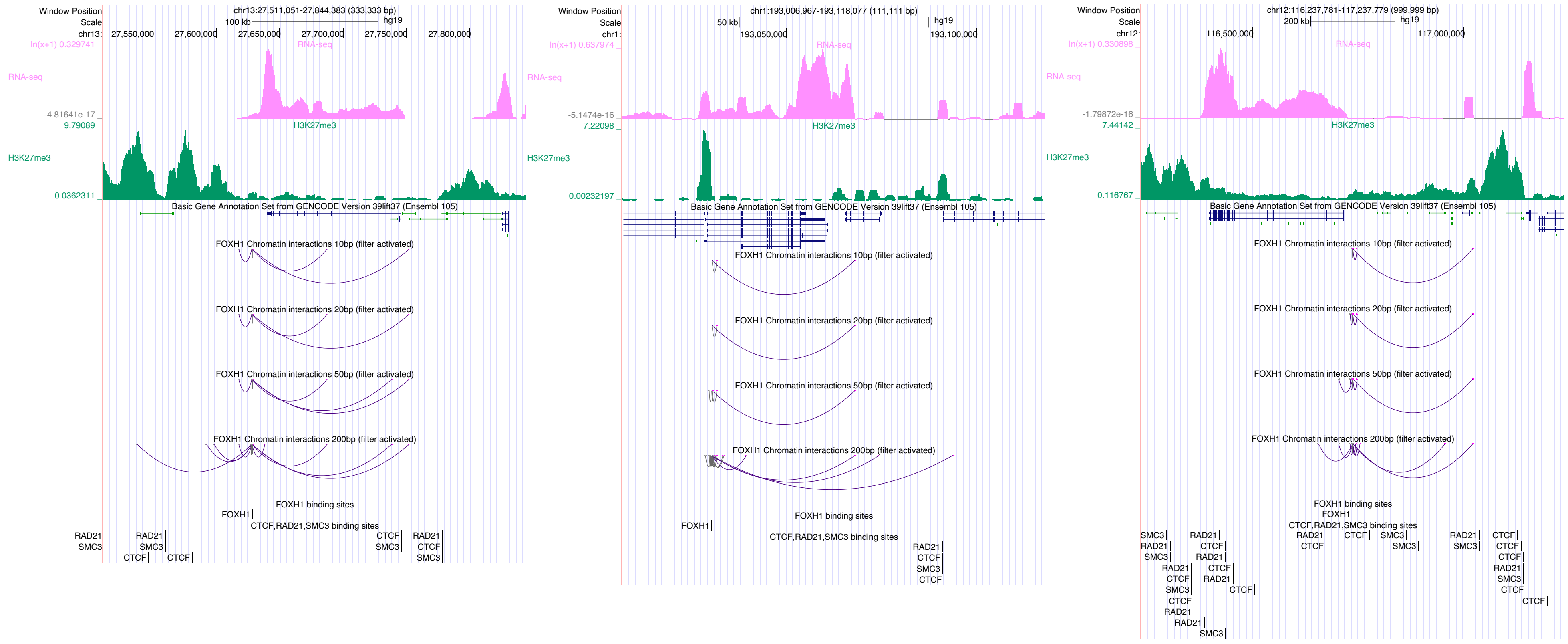

Window Position
chr13:27,511,051-27,844,383 (333,333 bp)
Scale
100 kb
hg19
chr13:
27,550,000
27,600,000
27,650,000
27,700,000
27,750,000
27,800,000
RNA-seq
H3K27me3
Basic Gene Annotation Set from GENCODE Version 39lift37 (Ensembl 105)
FOXH1 Chromatin interactions 10bp (filter activated)
FOXH1 Chromatin interactions 20bp (filter activated)
FOXH1 Chromatin interactions 50bp (filter activated)
FOXH1 Chromatin interactions 200bp (filter activated)
FOXH1 binding sites
FOXH1
CTCF,RAD21,SMC3 binding sites
Window Position
chr1:193,006,967-193,118,077 (111,111 bp)
Scale
50 kb
hg19
chr1:
193,050,000
193,100,000
Window Position
chr12:116,237,781-117,237,779 (999,999 bp)
Scale
200 kb
hg19
chr12:
116,500,000
117,000,000

### FOXK1

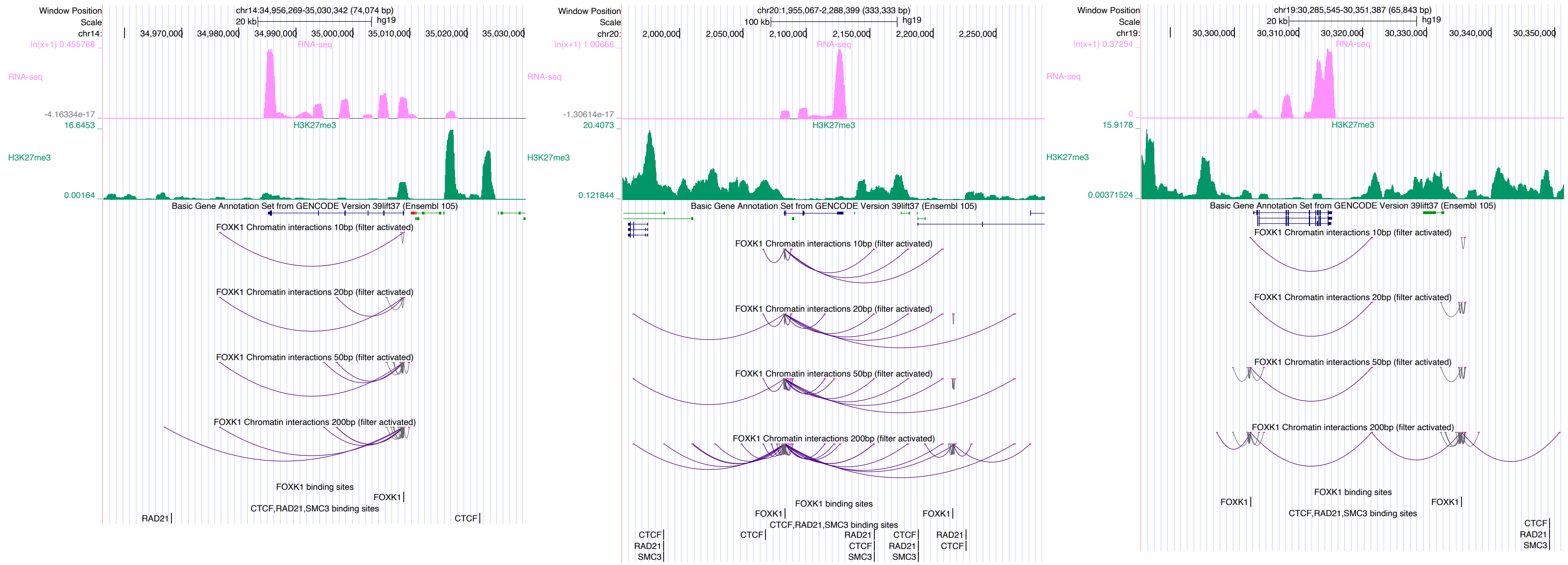
Window Position
chr14:34,956,269-35,030,342 (74,074 bp)
Scale
20 kb
hg19
chr14:
34,970,000
34,980,000
34,990,000
35,000,000
35,010,000
35,020,000
35,030,000
ln(x+1) 0.455768
RNA-seq
-4.16334e-17
16.6453
H3K27me3
0.00164
Basic Gene Annotation Set from GENCODE Version 39lift37 (Ensembl 105)
FOXK1 Chromatin interactions 10bp (filter activated)
FOXK1 Chromatin interactions 20bp (filter activated)
FOXK1 Chromatin interactions 50bp (filter activated)
FOXK1 Chromatin interactions 200bp (filter activated)
FOXK1 binding sites
FOXK1
CTCF,RAD21,SMC3 binding sites
RAD21
CTCF
Window Position
chr20:1,955,067-2,288,399 (333,333 bp)
Scale
100 kb
hg19
chr20:
2,000,000
2,050,000
2,100,000
2,150,000
2,200,000
2,250,000
ln(x+1) 1.00668
RNA-seq
-1.30614e-17
20.4073
H3K27me3
0.121844
Basic Gene Annotation Set from GENCODE Version 39lift37 (Ensembl 105)
FOXK1 Chromatin interactions 10bp (filter activated)
FOXK1 Chromatin interactions 20bp (filter activated)
FOXK1 Chromatin interactions 50bp (filter activated)
FOXK1 Chromatin interactions 200bp (filter activated)
FOXK1 binding sites
FOXK1
FOXK1
CTCF,RAD21,SMC3 binding sites
CTCF
RAD21
SMC3
CTCF
RAD21
CTCF
SMC3
CTCF
RAD21
SMC3
RAD21
CTCF
Window Position
chr19:30,285,545-30,351,387 (65,843 bp)
Scale
20 kb
hg19
chr19:
30,300,000
30,310,000
30,320,000
30,330,000
30,340,000
30,350,000
ln(x+1) 0.37254
RNA-seq
0
15.9178
H3K27me3
0.00371524
Basic Gene Annotation Set from GENCODE Version 39lift37 (Ensembl 105)
FOXK1 Chromatin interactions 10bp (filter activated)
FOXK1 Chromatin interactions 20bp (filter activated)
FOXK1 Chromatin interactions 50bp (filter activated)
FOXK1 Chromatin interactions 200bp (filter activated)
FOXK1 binding sites
FOXK1
FOXK1
CTCF,RAD21,SMC3 binding sites
CTCF
RAD21
SMC3

### GATA2

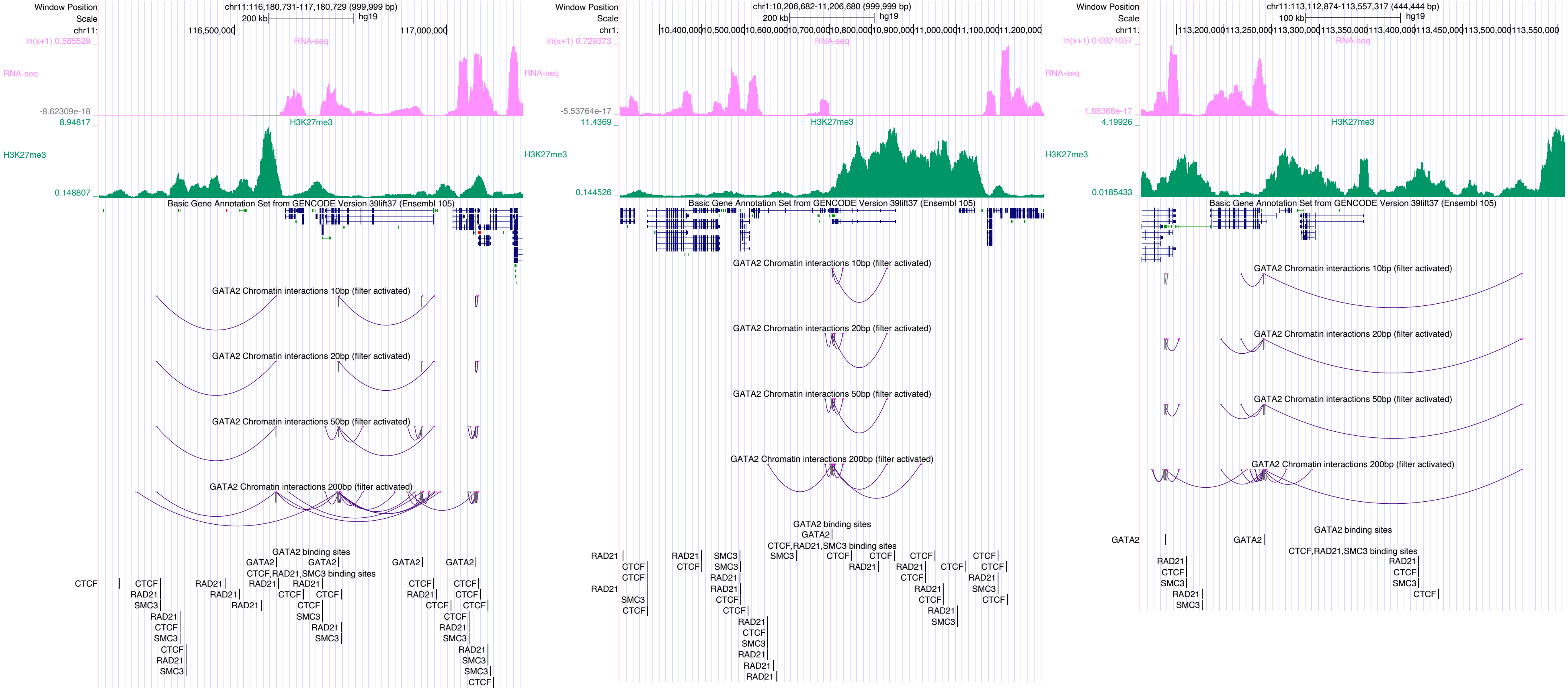

Window Position
Scale
chr11:116,180,731-117,180,729 (999,999 bp)
200 kb
hg19
RNA-seq
H3K27me3
Basic Gene Annotation Set from GENCODE Version 39lift37 (Ensembl 105)
GATA2 Chromatin interactions 10bp (filter activated)
GATA2 Chromatin interactions 20bp (filter activated)
GATA2 Chromatin interactions 50bp (filter activated)
GATA2 Chromatin interactions 200bp (filter activated)
GATA2 binding sites
CTCF,RAD21,SMC3 binding sites
chr1:10,206,682-11,206,680 (999,999 bp)
chr11:113,112,874-113,557,317 (444,444 bp)
100 kb

### GRHL1

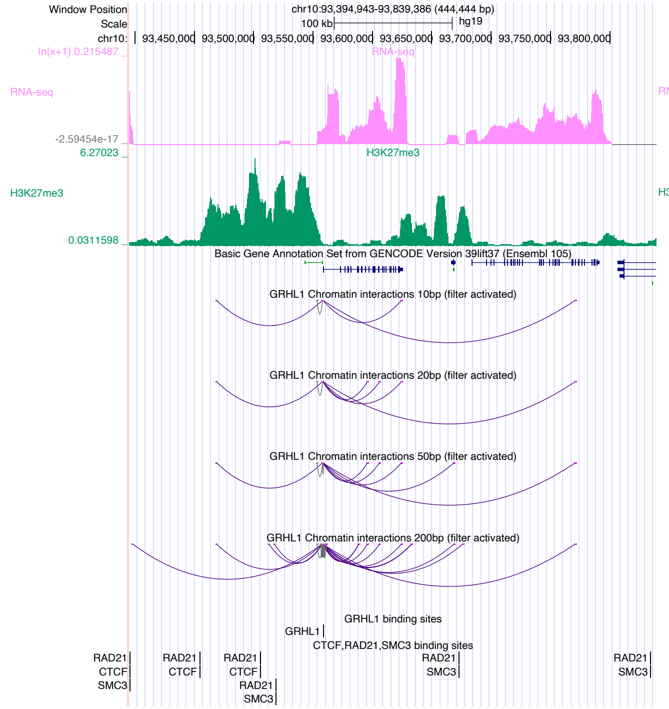

Window Position chr10:93,394,943-93,839,386 (444,444 bp)
Scale 100 kb hg19
chr10: 93,450,000 93,500,000 93,550,000 93,600,000 93,650,000 93,700,000 93,750,000 93,800,000
RNA-seq
H3K27me3
Basic Gene Annotation Set from GENCODE Version 39lift37 (Ensembl 105)
GRHL1 Chromatin interactions 10bp (filter activated)
GRHL1 Chromatin interactions 20bp (filter activated)
GRHL1 Chromatin interactions 50bp (filter activated)
GRHL1 Chromatin interactions 200bp (filter activated)
GRHL1 binding sites
CTCF,RAD21,SMC3 binding sites


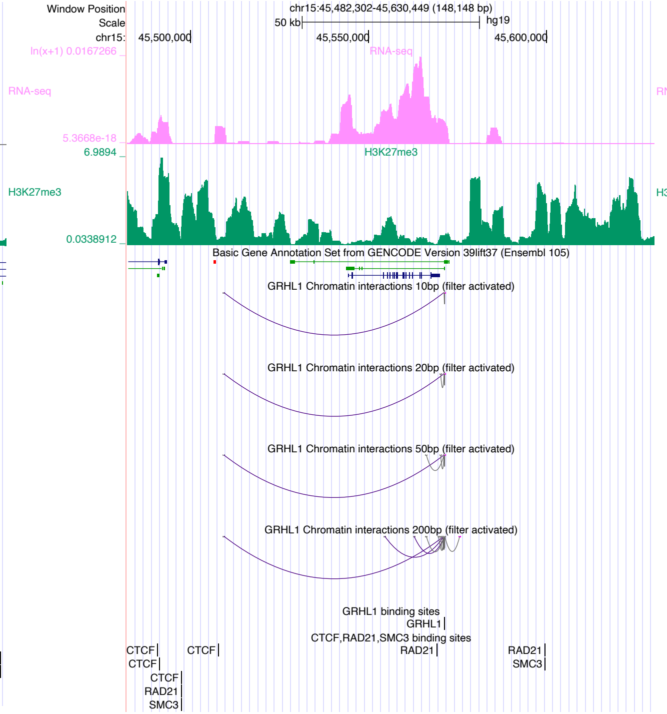

Window Position chr15:45,482,302-45,630,449 (148,148 bp)
Scale 50 kb hg19
chr15: 45,500,000 45,550,000 45,600,000
RNA-seq
H3K27me3
Basic Gene Annotation Set from GENCODE Version 39lift37 (Ensembl 105)
GRHL1 Chromatin interactions 10bp (filter activated)
GRHL1 Chromatin interactions 20bp (filter activated)
GRHL1 Chromatin interactions 50bp (filter activated)
GRHL1 Chromatin interactions 200bp (filter activated)
GRHL1 binding sites
CTCF,RAD21,SMC3 binding sites


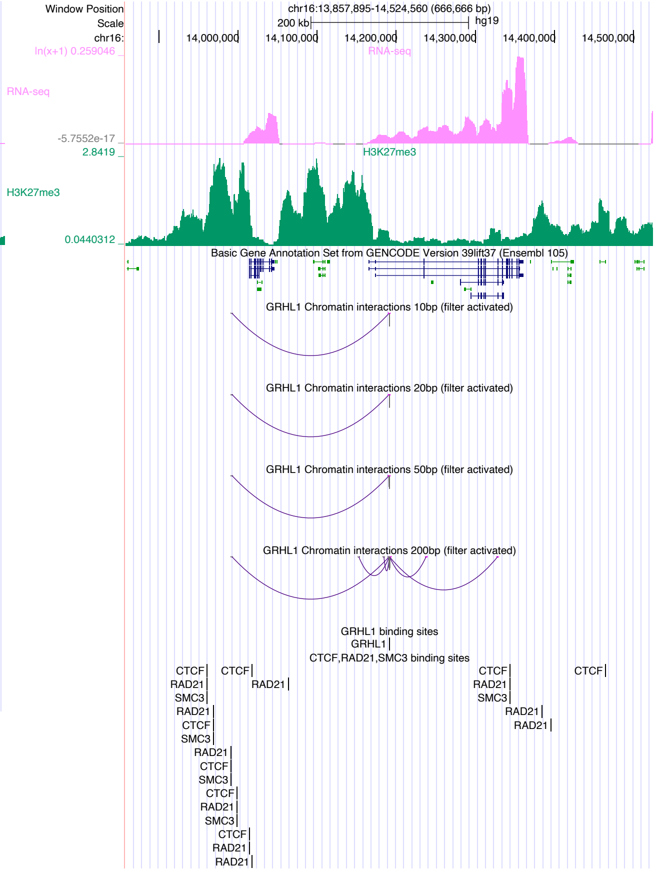

Window Position chr16:13,857,895-14,524,560 (666,666 bp)
Scale 200 kb hg19
chr16: 14,000,000 14,100,000 14,200,000 14,300,000 14,400,000 14,500,000
RNA-seq
H3K27me3
Basic Gene Annotation Set from GENCODE Version 39lift37 (Ensembl 105)
GRHL1 Chromatin interactions 10bp (filter activated)
GRHL1 Chromatin interactions 20bp (filter activated)
GRHL1 Chromatin interactions 50bp (filter activated)
GRHL1 Chromatin interactions 200bp (filter activated)
GRHL1 binding sites
CTCF,RAD21,SMC3 binding sites

### GRHL2

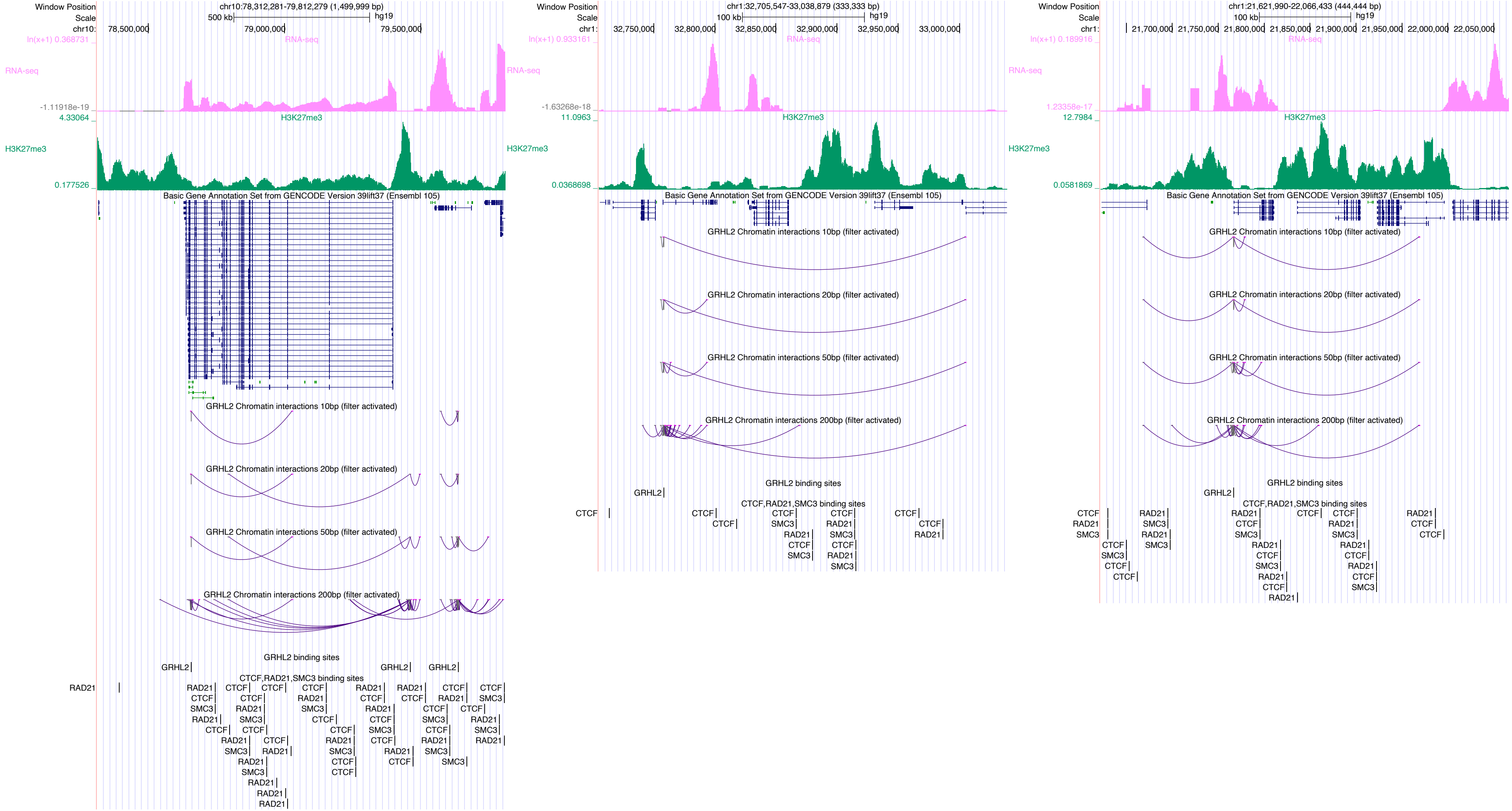
Window Position
chr10:78,312,281-79,812,279 (1,499,999 bp)
Scale
500 kb
hg19
chr10:
78,500,000
79,000,000
79,500,000
ln(x+1) 0.368731
RNA-seq
-1.11918e-19
4.33064
H3K27me3
0.177526
Basic Gene Annotation Set from GENCODE Version 39lift37 (Ensembl 105)
GRHL2 Chromatin interactions 10bp (filter activated)
GRHL2 Chromatin interactions 20bp (filter activated)
GRHL2 Chromatin interactions 50bp (filter activated)
GRHL2 Chromatin interactions 200bp (filter activated)
GRHL2 binding sites
CTCF,RAD21,SMC3 binding sites
Window Position
chr1:32,705,547-33,038,879 (333,333 bp)
Scale
100 kb
hg19
chr1:
32,750,000
32,800,000
32,850,000
32,900,000
32,950,000
33,000,000
ln(x+1) 0.933161
RNA-seq
-1.63268e-18
11.0963
H3K27me3
0.0368698
Basic Gene Annotation Set from GENCODE Version 39lift37 (Ensembl 105)
GRHL2 Chromatin interactions 10bp (filter activated)
GRHL2 Chromatin interactions 20bp (filter activated)
GRHL2 Chromatin interactions 50bp (filter activated)
GRHL2 Chromatin interactions 200bp (filter activated)
GRHL2 binding sites
CTCF,RAD21,SMC3 binding sites
Window Position
chr1:21,621,990-22,066,433 (444,444 bp)
Scale
100 kb
hg19
chr1:
21,700,000
21,750,000
21,800,000
21,850,000
21,900,000
21,950,000
22,000,000
22,050,000
ln(x+1) 0.189916
RNA-seq
1.23358e-17
12.7984
H3K27me3
0.0581869
Basic Gene Annotation Set from GENCODE Version 39lift37 (Ensembl 105)
GRHL2 Chromatin interactions 10bp (filter activated)
GRHL2 Chromatin interactions 20bp (filter activated)
GRHL2 Chromatin interactions 50bp (filter activated)
GRHL2 Chromatin interactions 200bp (filter activated)
GRHL2 binding sites
CTCF,RAD21,SMC3 binding sites

### HNF4A

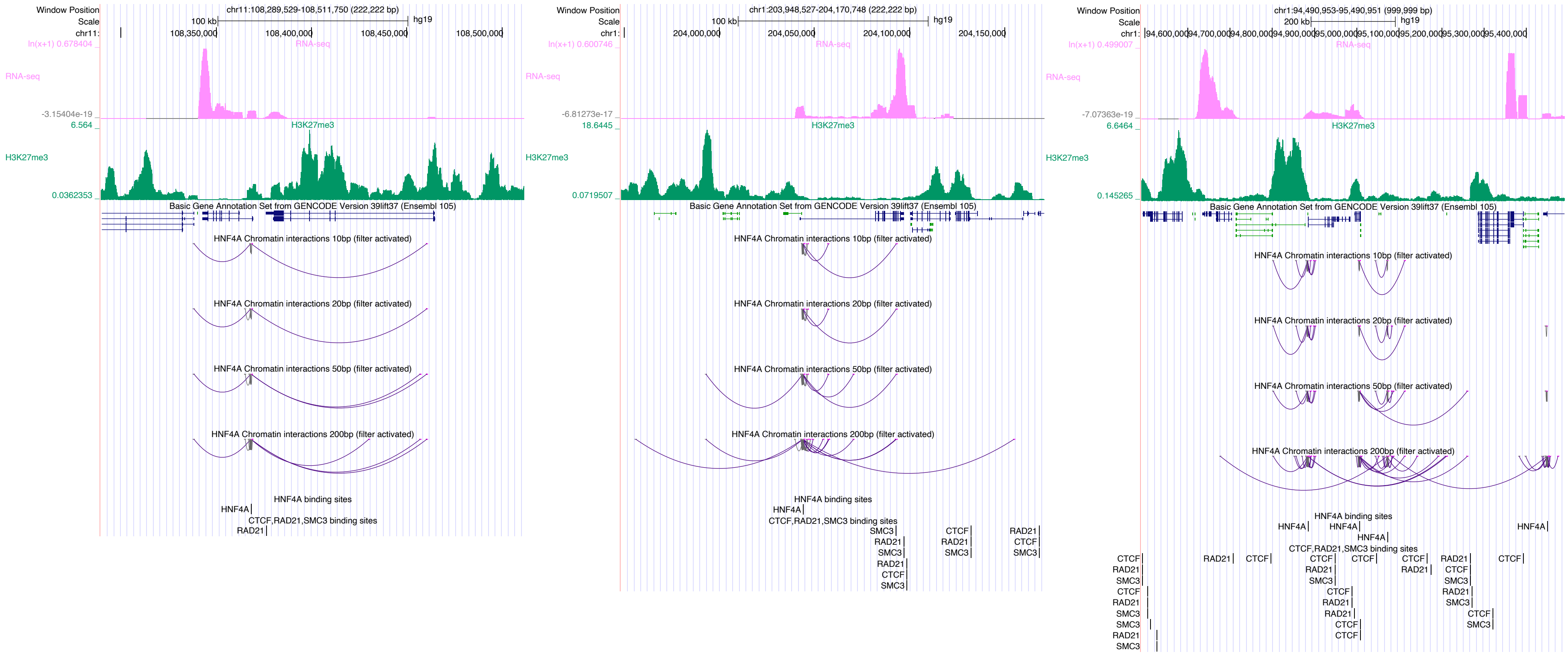

Window Position
Scale
chr11:108,289,529-108,511,750 (222,222 bp)
100 kb
hg19
108,350,000
108,400,000
108,450,000
108,500,000
RNA-seq
H3K27me3
Basic Gene Annotation Set from GENCODE Version 39lift37 (Ensembl 105)
HNF4A Chromatin interactions 10bp (filter activated)
HNF4A Chromatin interactions 20bp (filter activated)
HNF4A Chromatin interactions 50bp (filter activated)
HNF4A Chromatin interactions 200bp (filter activated)
HNF4A binding sites
CTCF,RAD21,SMC3 binding sites
chr1:203,948,527-204,170,748 (222,222 bp)
204,000,000
204,050,000
204,100,000
204,150,000
chr1:94,490,953-95,490,951 (999,999 bp)
200 kb
94,600,000
94,700,000
94,800,000
94,900,000
95,000,000
95,100,000
95,200,000
95,300,000
95,400,000

### JUN

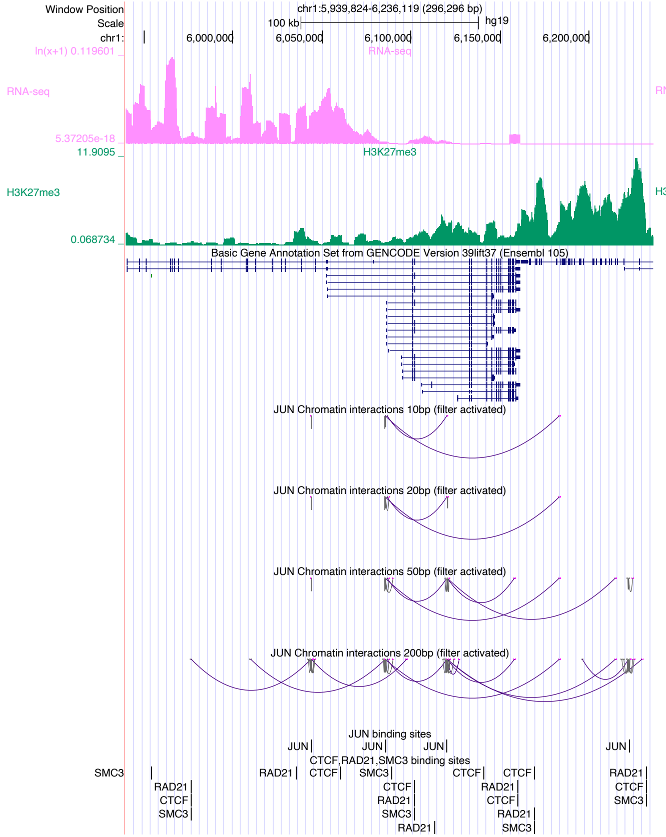

Window Position
chr1:5,939,824-6,236,119 (296,296 bp)
Scale
100 kb
hg19
chr1:
6,000,000
6,050,000
6,100,000
6,150,000
6,200,000
ln(x+1) 0.119601
RNA-seq
5.37205e-18
11.9095
H3K27me3
0.068734
Basic Gene Annotation Set from GENCODE Version 39lift37 (Ensembl 105)
JUN Chromatin interactions 10bp (filter activated)
JUN Chromatin interactions 20bp (filter activated)
JUN Chromatin interactions 50bp (filter activated)
JUN Chromatin interactions 200bp (filter activated)
JUN binding sites
CTCF,RAD21,SMC3 binding sites


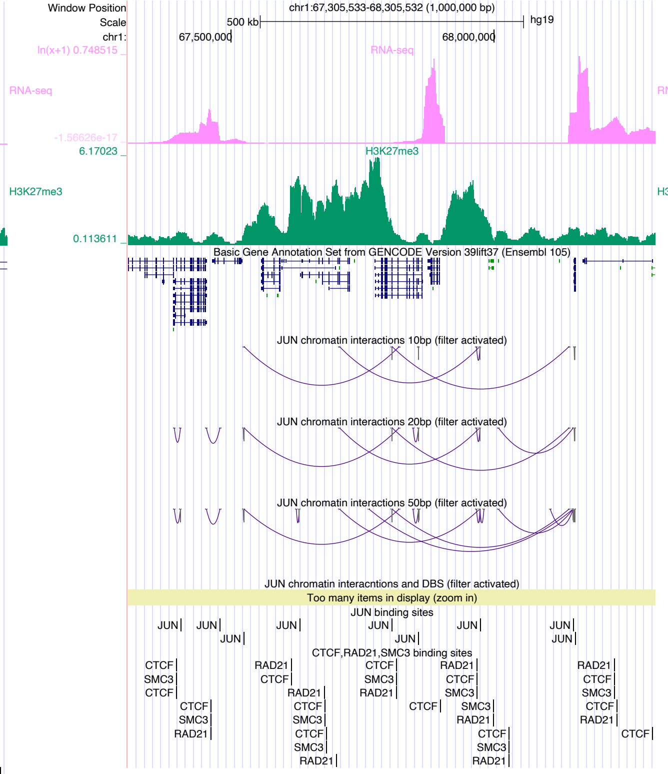

Window Position
chr1:67,305,533-68,305,532 (1,000,000 bp)
Scale
500 kb
hg19
chr1:
67,500,000
68,000,000
ln(x+1) 0.748515
RNA-seq
-1.56626e-17
6.17023
H3K27me3
0.113611
Basic Gene Annotation Set from GENCODE Version 39lift37 (Ensembl 105)
JUN chromatin interactions 10bp (filter activated)
JUN chromatin interactions 20bp (filter activated)
JUN chromatin interactions 50bp (filter activated)
JUN chromatin interacntions and DBS (filter activated)
Too many items in display (zoom in)
JUN binding sites
CTCF,RAD21,SMC3 binding sites


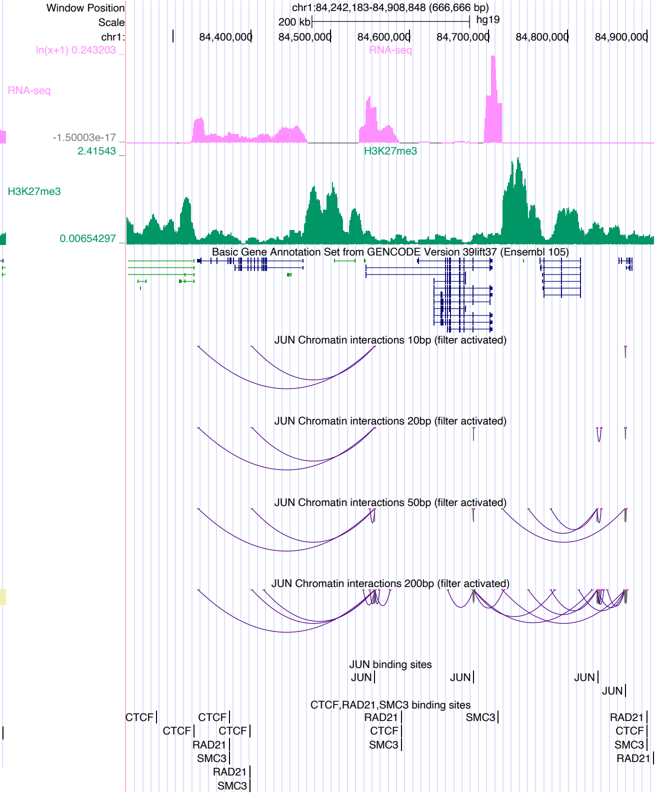

Window Position
chr1:84,242,183-84,908,848 (666,666 bp)
Scale
200 kb
hg19
chr1:
84,400,000
84,500,000
84,600,000
84,700,000
84,800,000
84,900,000
ln(x+1) 0.243203
RNA-seq
-1.50003e-17
2.41543
H3K27me3
0.00654297
Basic Gene Annotation Set from GENCODE Version 39lift37 (Ensembl 105)
JUN Chromatin interactions 10bp (filter activated)
JUN Chromatin interactions 20bp (filter activated)
JUN Chromatin interactions 50bp (filter activated)
JUN Chromatin interactions 200bp (filter activated)
JUN binding sites
CTCF,RAD21,SMC3 binding sites

### JUNB

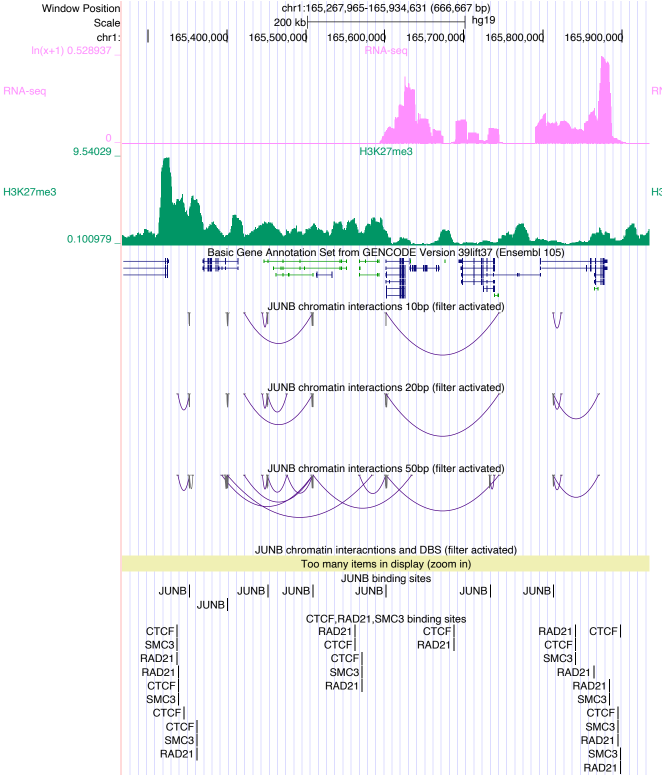
Window Position
chr1:165,267,965-165,934,631 (666,667 bp)
Scale
200 kb
hg19
chr1:
165,400,000
165,500,000
165,600,000
165,700,000
165,800,000
165,900,000
ln(x+1) 0.528937
RNA-seq
0
9.54029
H3K27me3
0.100979
Basic Gene Annotation Set from GENCODE Version 39lift37 (Ensembl 105)
JUNB chromatin interactions 10bp (filter activated)
JUNB chromatin interactions 20bp (filter activated)
JUNB chromatin interactions 50bp (filter activated)
JUNB chromatin interacntions and DBS (filter activated)
Too many items in display (zoom in)
JUNB binding sites
CTCF,RAD21,SMC3 binding sites

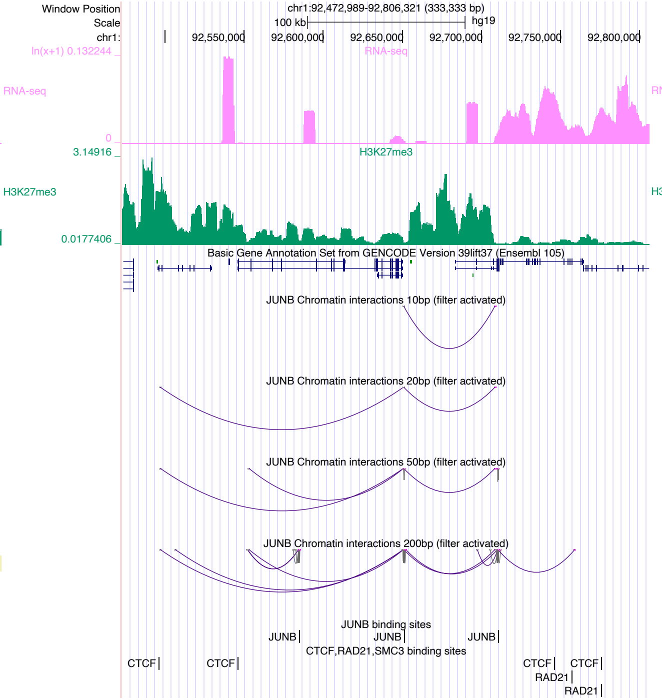
Window Position
chr1:92,472,989-92,806,321 (333,333 bp)
Scale
100 kb
hg19
chr1:
92,550,000
92,600,000
92,650,000
92,700,000
92,750,000
92,800,000
ln(x+1) 0.132244
RNA-seq
0
3.14916
H3K27me3
0.0177406
Basic Gene Annotation Set from GENCODE Version 39lift37 (Ensembl 105)
JUNB Chromatin interactions 10bp (filter activated)
JUNB Chromatin interactions 20bp (filter activated)
JUNB Chromatin interactions 50bp (filter activated)
JUNB Chromatin interactions 200bp (filter activated)
JUNB binding sites
CTCF,RAD21,SMC3 binding sites

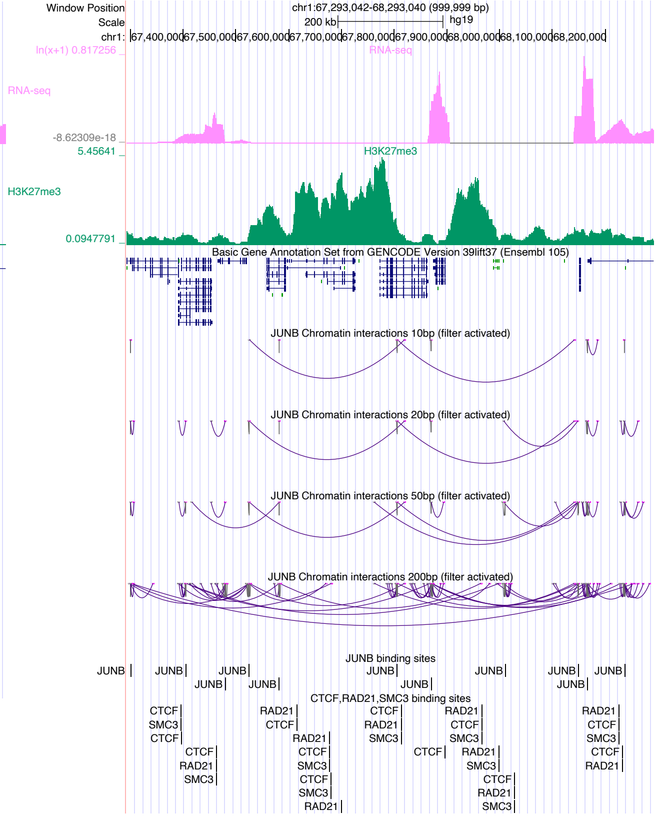
Window Position
chr1:67,293,042-68,293,040 (999,999 bp)
Scale
200 kb
hg19
chr1:
67,400,000
67,500,000
67,600,000
67,700,000
67,800,000
67,900,000
68,000,000
68,100,000
68,200,000
ln(x+1) 0.817256
RNA-seq
-8.62309e-18
5.45641
H3K27me3
0.0947791
Basic Gene Annotation Set from GENCODE Version 39lift37 (Ensembl 105)
JUNB Chromatin interactions 10bp (filter activated)
JUNB Chromatin interactions 20bp (filter activated)
JUNB Chromatin interactions 50bp (filter activated)
JUNB Chromatin interactions 200bp (filter activated)
JUNB binding sites
CTCF,RAD21,SMC3 binding sites

### JUND

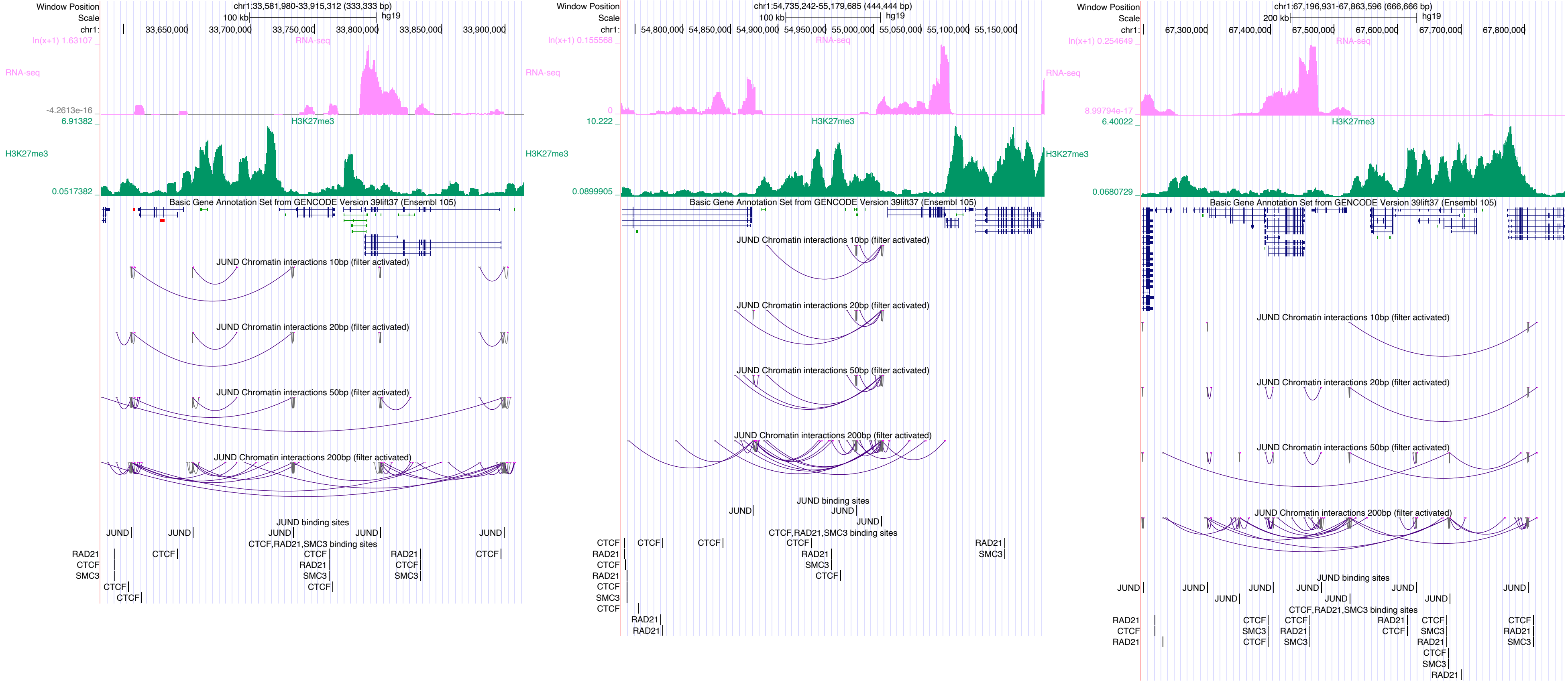

Window Position
Scale
chr1:
chr1:33,581,980-33,915,312 (333,333 bp)
100 kb
hg19
33,650,000
33,700,000
33,750,000
33,800,000
33,850,000
33,900,000
RNA-seq
ln(x+1) 1.63107
-4.2613e-16
H3K27me3
6.91382
0.0517382
Basic Gene Annotation Set from GENCODE Version 39lift37 (Ensembl 105)
JUND Chromatin interactions 10bp (filter activated)
JUND Chromatin interactions 20bp (filter activated)
JUND Chromatin interactions 50bp (filter activated)
JUND Chromatin interactions 200bp (filter activated)
JUND binding sites
CTCF,RAD21,SMC3 binding sites
JUND
CTCF
RAD21
SMC3
chr1:54,735,242-55,179,685 (444,444 bp)
54,800,000
54,850,000
54,900,000
54,950,000
55,000,000
55,050,000
55,100,000
55,150,000
ln(x+1) 0.155568
0
10.222
0.0899905
chr1:67,196,931-67,863,596 (666,666 bp)
200 kb
67,300,000
67,400,000
67,500,000
67,600,000
67,700,000
67,800,000
ln(x+1) 0.254649
8.99794e-17
6.40022
0.0680729

### KLF1

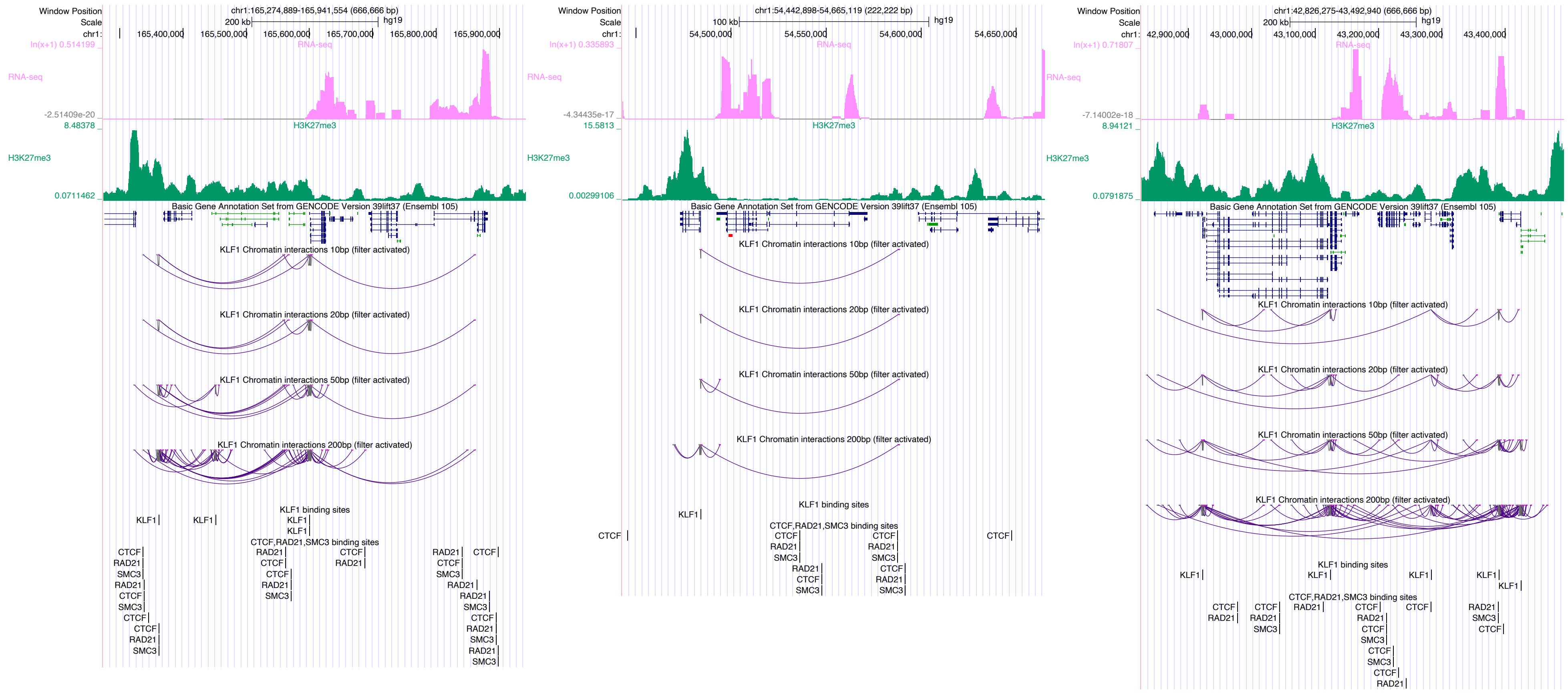
Window Position
Scale
chr1:
chr1:165,274,889-165,941,554 (666,666 bp)
200 kb
hg19
165,400,000
165,500,000
165,600,000
165,700,000
165,800,000
165,900,000
ln(x+1) 0.514199
RNA-seq
-2.51409e-20
8.48378
H3K27me3
0.0711462
Basic Gene Annotation Set from GENCODE Version 39lift37 (Ensembl 105)
KLF1 Chromatin interactions 10bp (filter activated)
KLF1 Chromatin interactions 20bp (filter activated)
KLF1 Chromatin interactions 50bp (filter activated)
KLF1 Chromatin interactions 200bp (filter activated)
KLF1 binding sites
CTCF,RAD21,SMC3 binding sites
chr1:54,442,898-54,665,119 (222,222 bp)
100 kb
54,500,000
54,550,000
54,600,000
54,650,000
ln(x+1) 0.335893
-4.34435e-17
15.5813
0.00299106
chr1:42,826,275-43,492,940 (666,666 bp)
42,900,000
43,000,000
43,100,000
43,200,000
43,300,000
43,400,000
ln(x+1) 0.71807
-7.14002e-18
8.94121
0.0791875
KLF1
CTCF
RAD21
SMC3

### KLF5

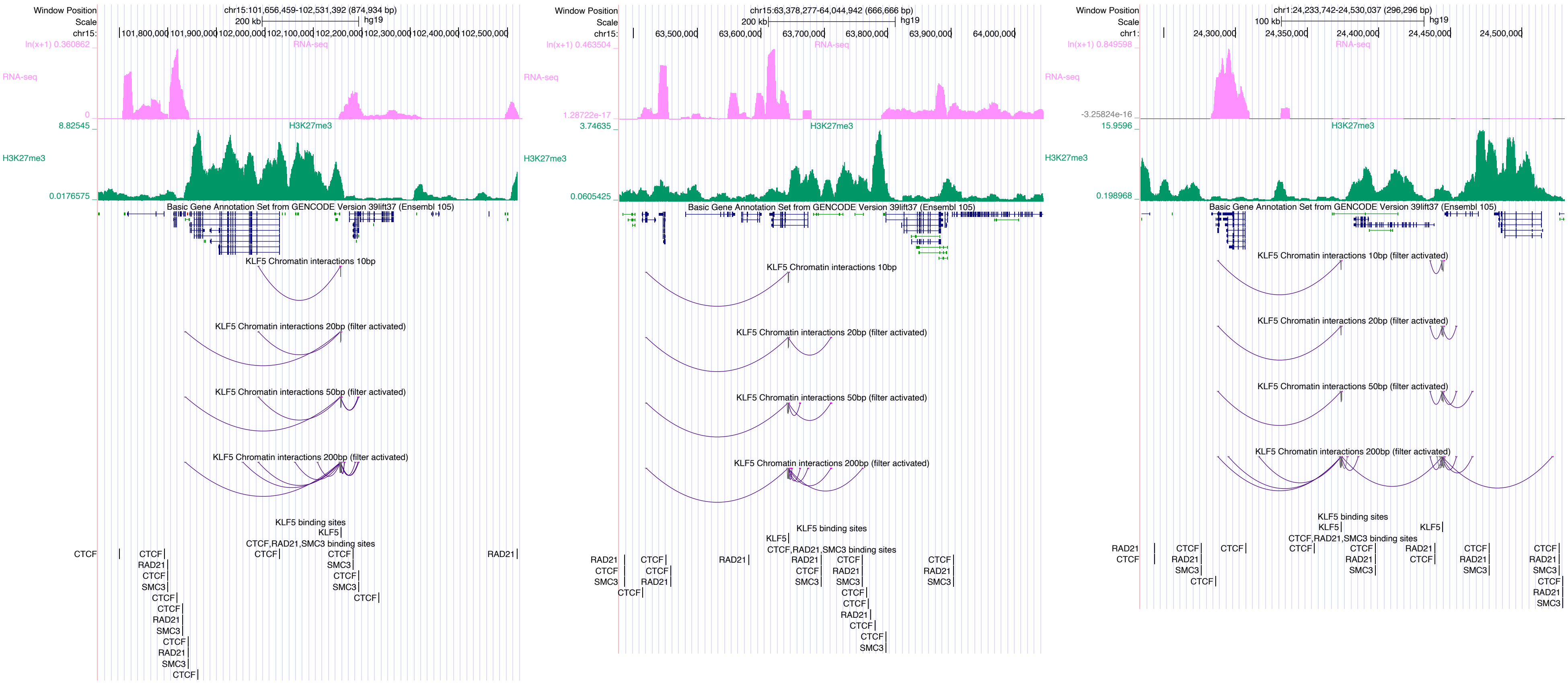

Window Position
Scale
chr15:101,656,459-102,531,392 (874,934 bp)
200 kb
hg19
chr15:
101,800,000
101,900,000
102,000,000
102,100,000
102,200,000
102,300,000
102,400,000
102,500,000
ln(x+1) 0.360862
RNA-seq
0
8.82545
H3K27me3
0.0176575
Basic Gene Annotation Set from GENCODE Version 39lift37 (Ensembl 105)
KLF5 Chromatin interactions 10bp
KLF5 Chromatin interactions 20bp (filter activated)
KLF5 Chromatin interactions 50bp (filter activated)
KLF5 Chromatin interactions 200bp (filter activated)
KLF5 binding sites
KLF5
CTCF,RAD21,SMC3 binding sites
CTCF
RAD21
SMC3
chr15:63,378,277-64,044,942 (666,666 bp)
63,500,000
63,600,000
63,700,000
63,800,000
63,900,000
64,000,000
ln(x+1) 0.463504
1.28722e-17
3.74635
0.0605425
chr1:24,233,742-24,530,037 (296,296 bp)
100 kb
chr1:
24,300,000
24,350,000
24,400,000
24,450,000
24,500,000
ln(x+1) 0.849598
-3.25824e-16
15.9596
0.198968

### KLF8

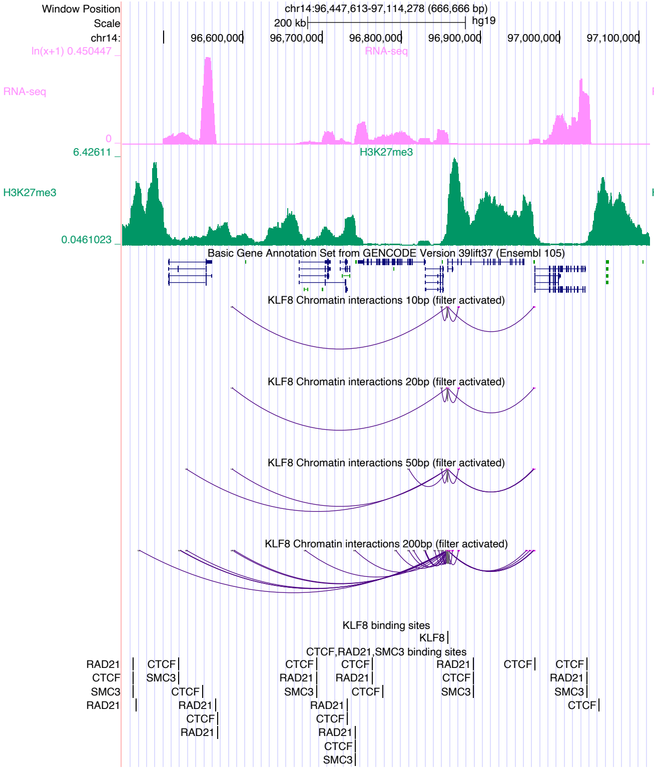

Window Position
chr14:96,447,613-97,114,278 (666,666 bp)
Scale
200 kb
hg19
chr14:
96,600,000
96,700,000
96,800,000
96,900,000
97,000,000
97,100,000
RNA-seq
H3K27me3
Basic Gene Annotation Set from GENCODE Version 39lift37 (Ensembl 105)
KLF8 Chromatin interactions 10bp (filter activated)
KLF8 Chromatin interactions 20bp (filter activated)
KLF8 Chromatin interactions 50bp (filter activated)
KLF8 Chromatin interactions 200bp (filter activated)
KLF8 binding sites
CTCF,RAD21,SMC3 binding sites


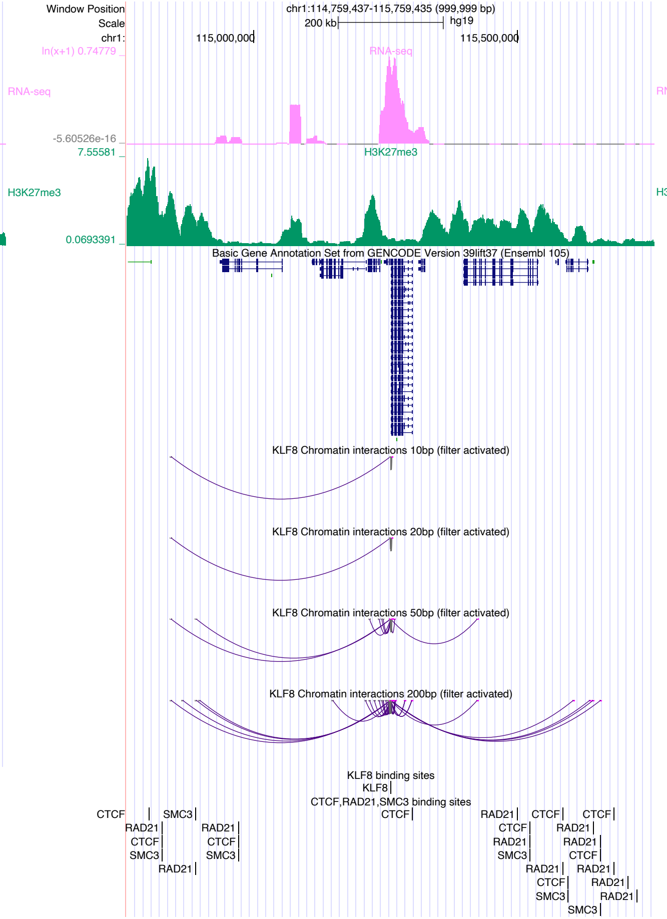

Window Position
chr1:114,759,437-115,759,435 (999,999 bp)
Scale
200 kb
hg19
chr1:
115,000,000
115,500,000
RNA-seq
H3K27me3
Basic Gene Annotation Set from GENCODE Version 39lift37 (Ensembl 105)
KLF8 Chromatin interactions 10bp (filter activated)
KLF8 Chromatin interactions 20bp (filter activated)
KLF8 Chromatin interactions 50bp (filter activated)
KLF8 Chromatin interactions 200bp (filter activated)
KLF8 binding sites
CTCF,RAD21,SMC3 binding sites


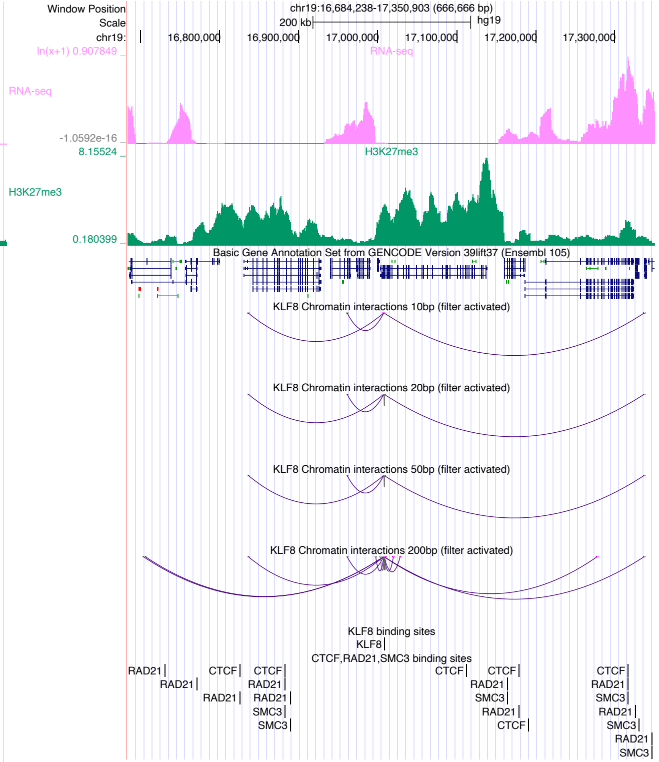

Window Position
chr19:16,684,238-17,350,903 (666,666 bp)
Scale
200 kb
hg19
chr19:
16,800,000
16,900,000
17,000,000
17,100,000
17,200,000
17,300,000
RNA-seq
H3K27me3
Basic Gene Annotation Set from GENCODE Version 39lift37 (Ensembl 105)
KLF8 Chromatin interactions 10bp (filter activated)
KLF8 Chromatin interactions 20bp (filter activated)
KLF8 Chromatin interactions 50bp (filter activated)
KLF8 Chromatin interactions 200bp (filter activated)
KLF8 binding sites
CTCF,RAD21,SMC3 binding sites

### KLF9

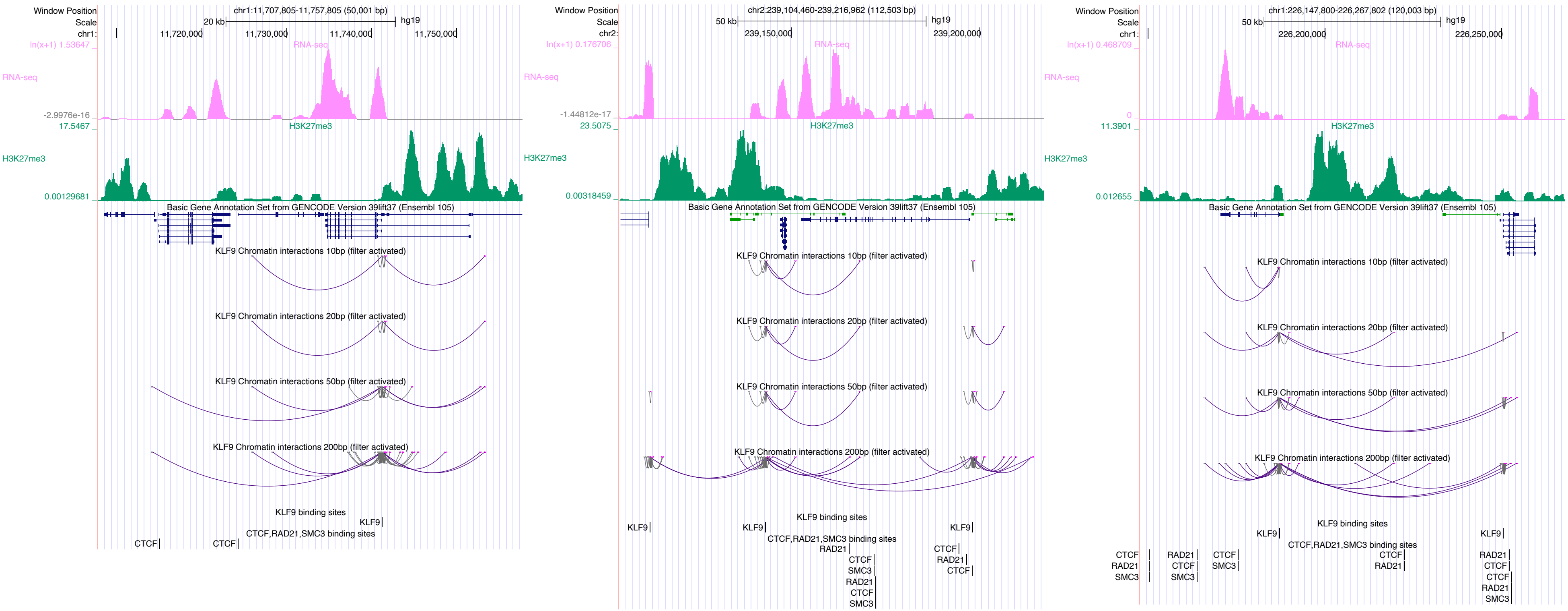
Window Position
Scale
chr1:11,707,805-11,757,805 (50,001 bp)
20 kb
hg19
11,720,000
11,730,000
11,740,000
11,750,000
RNA-seq
H3K27me3
Basic Gene Annotation Set from GENCODE Version 39lift37 (Ensembl 105)
KLF9 Chromatin interactions 10bp (filter activated)
KLF9 Chromatin interactions 20bp (filter activated)
KLF9 Chromatin interactions 50bp (filter activated)
KLF9 Chromatin interactions 200bp (filter activated)
KLF9 binding sites
CTCF,RAD21,SMC3 binding sites
chr2:239,104,460-239,216,962 (112,503 bp)
50 kb
239,150,000
239,200,000
chr1:226,147,800-226,267,802 (120,003 bp)
226,200,000
226,250,000
KLF9
CTCF
RAD21
SMC3

### KLF16

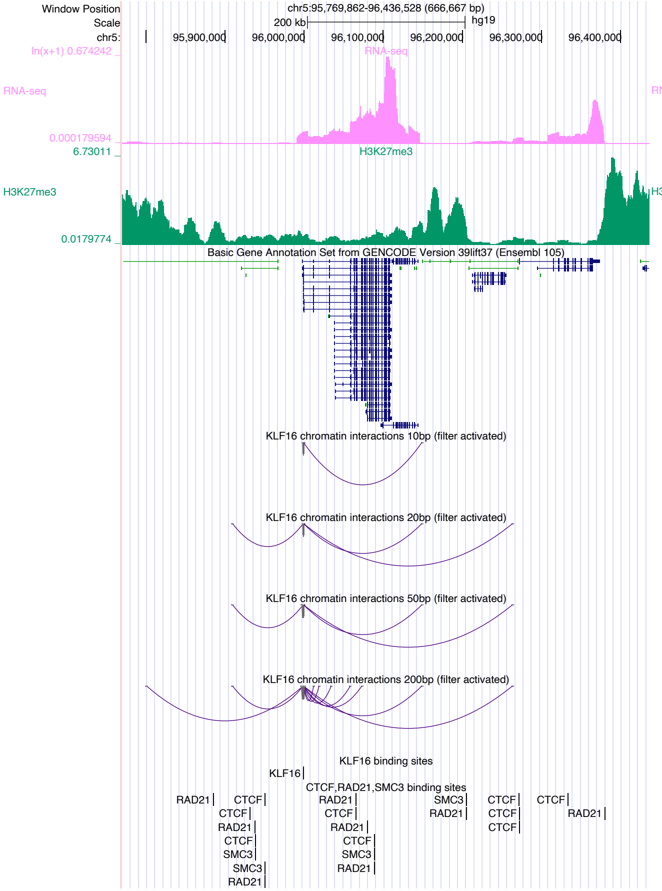

chr5:95,769,862-96,436,528 (666,667 bp)
RNA-seq
H3K27me3
Basic Gene Annotation Set from GENCODE Version 39lift37 (Ensembl 105)
KLF16 chromatin interactions 10bp (filter activated)
KLF16 chromatin interactions 20bp (filter activated)
KLF16 chromatin interactions 50bp (filter activated)
KLF16 chromatin interactions 200bp (filter activated)
KLF16 binding sites
CTCF,RAD21,SMC3 binding sites


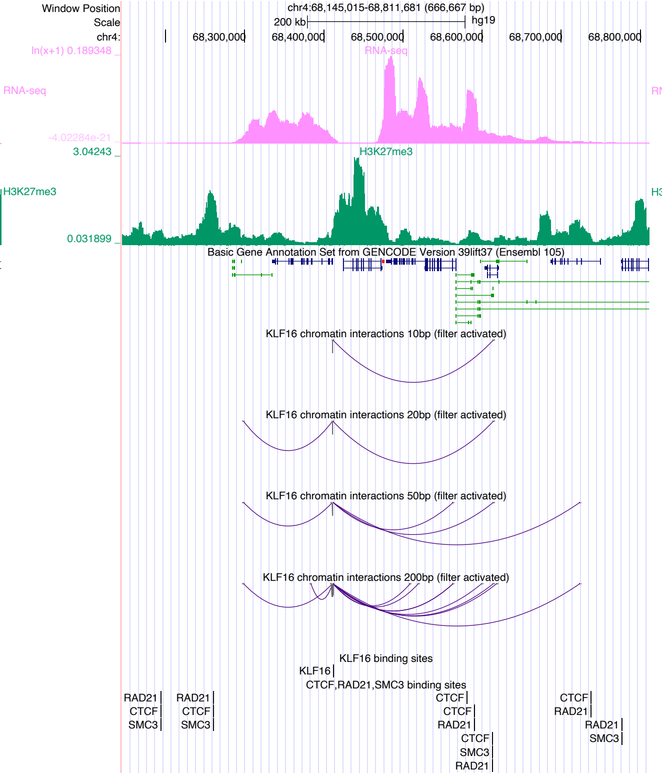

chr4:68,145,015-68,811,681 (666,667 bp)
RNA-seq
H3K27me3
Basic Gene Annotation Set from GENCODE Version 39lift37 (Ensembl 105)
KLF16 chromatin interactions 10bp (filter activated)
KLF16 chromatin interactions 20bp (filter activated)
KLF16 chromatin interactions 50bp (filter activated)
KLF16 chromatin interactions 200bp (filter activated)
KLF16 binding sites
CTCF,RAD21,SMC3 binding sites


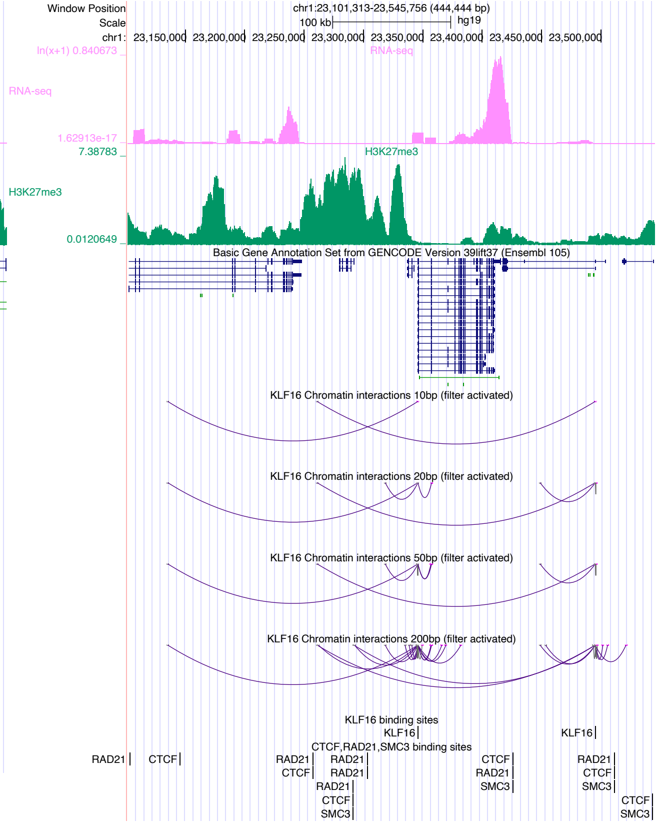

chr1:23,101,313-23,545,756 (444,444 bp)
RNA-seq
H3K27me3
Basic Gene Annotation Set from GENCODE Version 39lift37 (Ensembl 105)
KLF16 Chromatin interactions 10bp (filter activated)
KLF16 Chromatin interactions 20bp (filter activated)
KLF16 Chromatin interactions 50bp (filter activated)
KLF16 Chromatin interactions 200bp (filter activated)
KLF16 binding sites
CTCF,RAD21,SMC3 binding sites

### MAFG

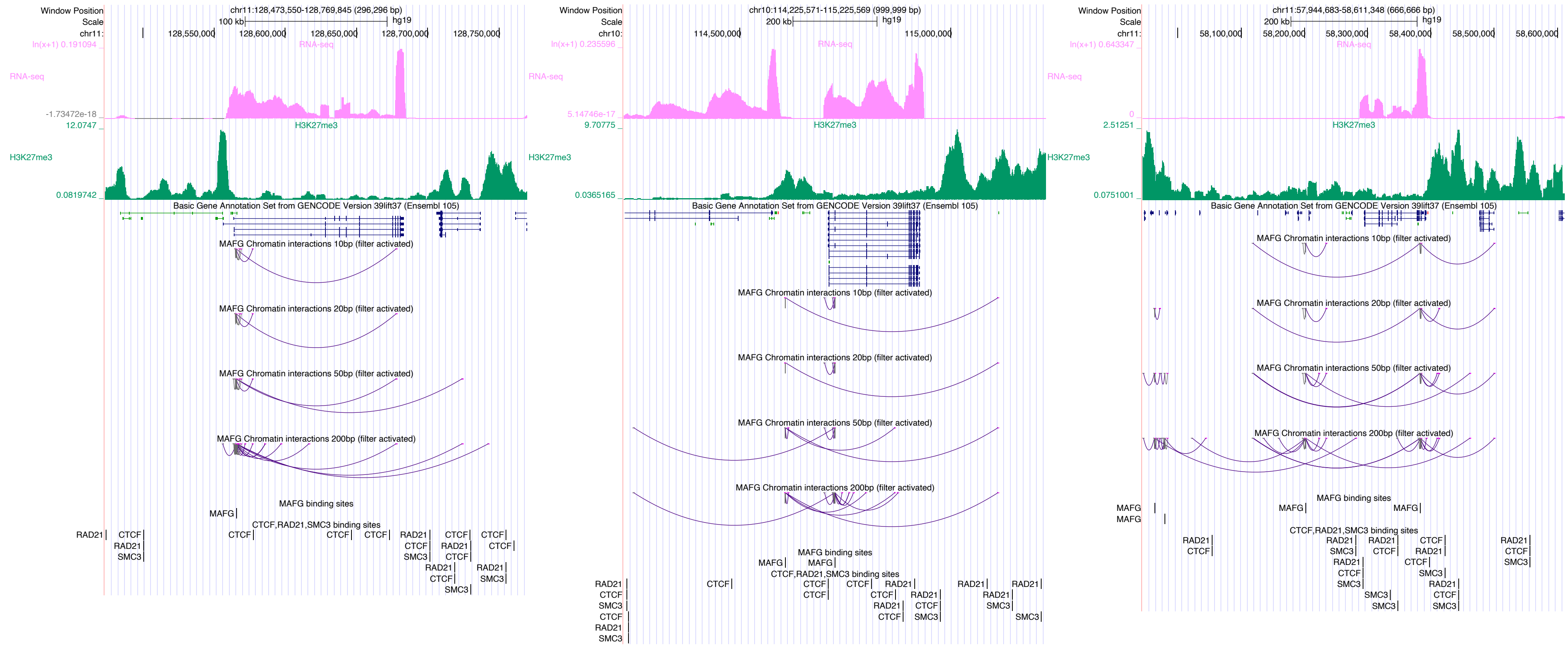

Window Position
Scale
chr11:128,473,550-128,769,845 (296,296 bp)
100 kb
hg19
128,550,000
128,600,000
128,650,000
128,700,000
128,750,000
RNA-seq
H3K27me3
Basic Gene Annotation Set from GENCODE Version 39lift37 (Ensembl 105)
MAFG Chromatin interactions 10bp (filter activated)
MAFG Chromatin interactions 20bp (filter activated)
MAFG Chromatin interactions 50bp (filter activated)
MAFG Chromatin interactions 200bp (filter activated)
MAFG binding sites
CTCF,RAD21,SMC3 binding sites
chr10:114,225,571-115,225,569 (999,999 bp)
200 kb
114,500,000
115,000,000
chr11:57,944,683-58,611,348 (666,666 bp)
58,100,000
58,200,000
58,300,000
58,400,000
58,500,000
58,600,000

### MAZ

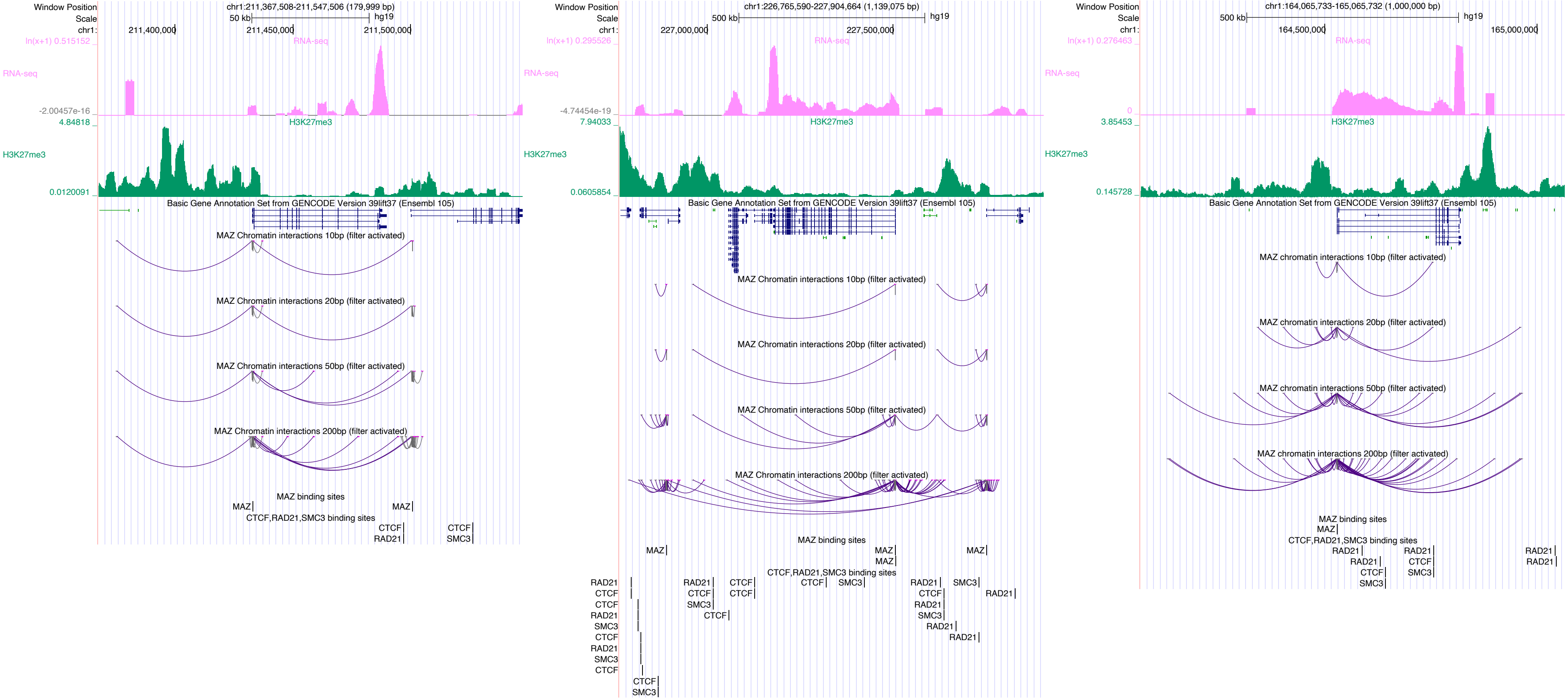

Window Position
Scale
chr1:
chr1:211,367,508-211,547,506 (179,999 bp)
50 kb
hg19
211,400,000
211,450,000
211,500,000
ln(x+1) 0.515152
RNA-seq
-2.00457e-16
4.84818
H3K27me3
0.0120091
Basic Gene Annotation Set from GENCODE Version 39lift37 (Ensembl 105)
MAZ Chromatin interactions 10bp (filter activated)
MAZ Chromatin interactions 20bp (filter activated)
MAZ Chromatin interactions 50bp (filter activated)
MAZ Chromatin interactions 200bp (filter activated)
MAZ binding sites
MAZ
CTCF,RAD21,SMC3 binding sites
CTCF
RAD21
SMC3
chr1:226,765,590-227,904,664 (1,139,075 bp)
500 kb
227,000,000
227,500,000
ln(x+1) 0.295526
-4.74454e-19
7.94033
0.0605854
chr1:164,065,733-165,065,732 (1,000,000 bp)
164,500,000
165,000,000
ln(x+1) 0.276463
0
3.85453
0.145728
MAZ chromatin interactions 10bp (filter activated)
MAZ chromatin interactions 20bp (filter activated)
MAZ chromatin interactions 50bp (filter activated)
MAZ chromatin interactions 200bp (filter activated)

### MECP2

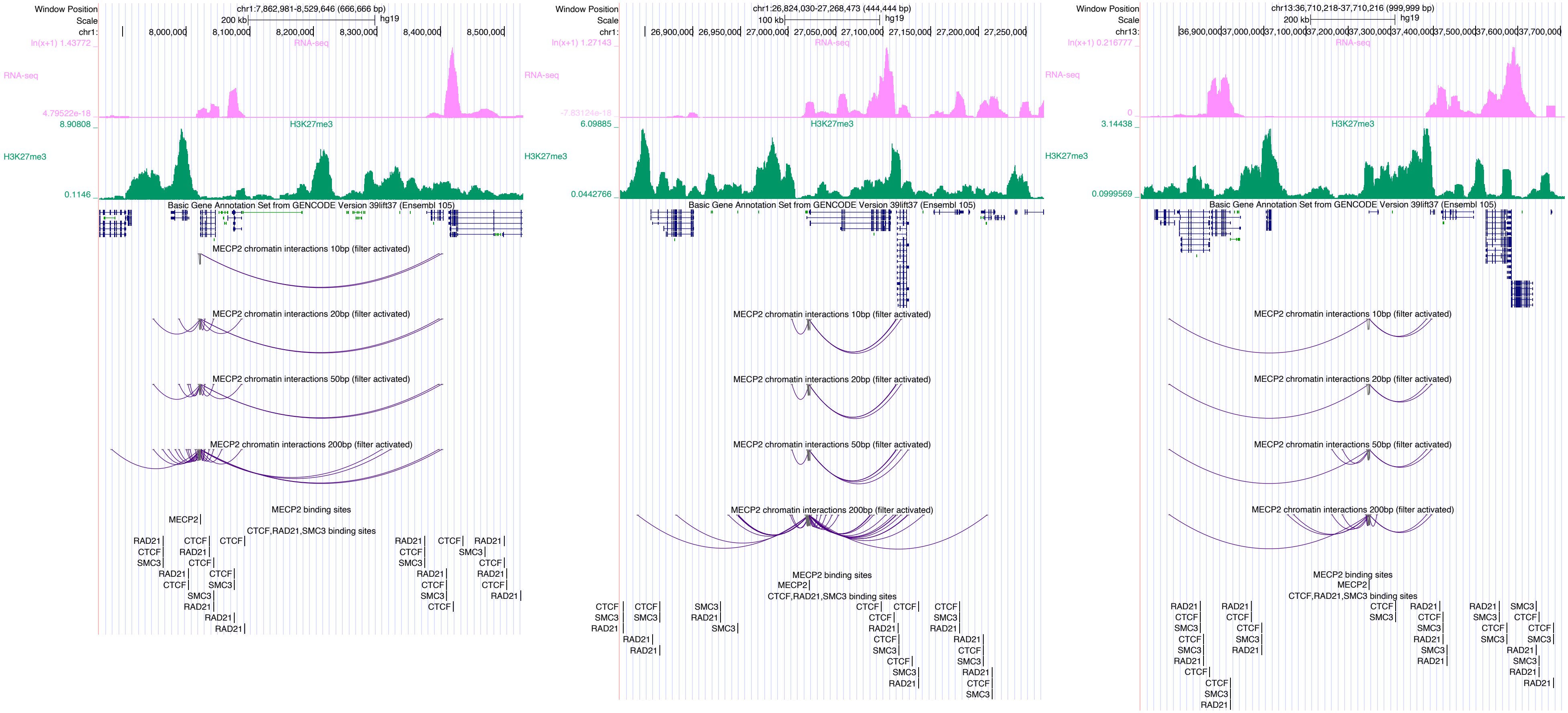

Window Position
chr1:7,862,981-8,529,646 (666,666 bp)
Scale
200 kb
hg19
RNA-seq
H3K27me3
Basic Gene Annotation Set from GENCODE Version 39lift37 (Ensembl 105)
MECP2 chromatin interactions 10bp (filter activated)
MECP2 chromatin interactions 20bp (filter activated)
MECP2 chromatin interactions 50bp (filter activated)
MECP2 chromatin interactions 200bp (filter activated)
MECP2 binding sites
CTCF,RAD21,SMC3 binding sites
chr1:26,824,030-27,268,473 (444,444 bp)
100 kb
chr13:36,710,218-37,710,216 (999,999 bp)

### MEF2B

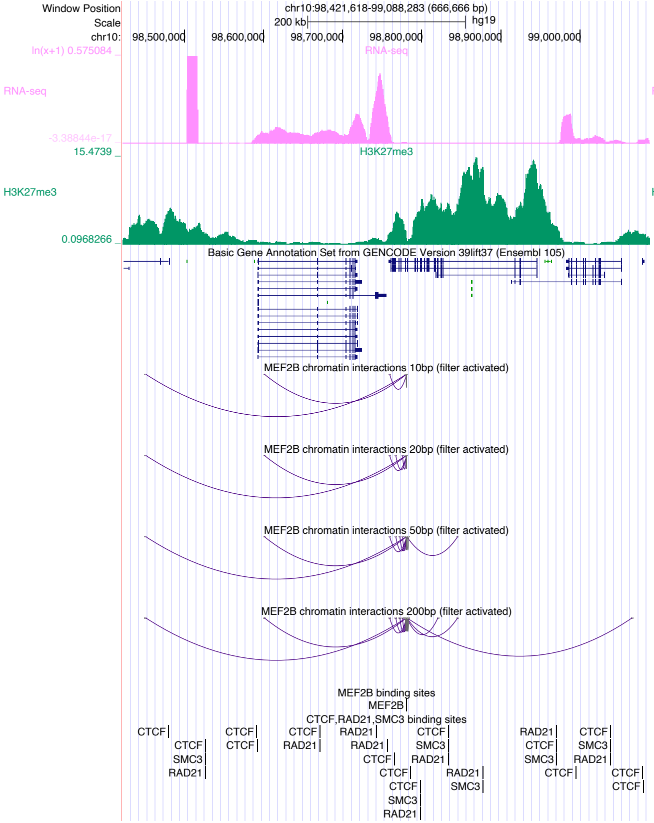

Window Position chr10:98,421,618-99,088,283 (666,666 bp)
Scale 200 kb hg19
chr10: 98,500,000 98,600,000 98,700,000 98,800,000 98,900,000 99,000,000
RNA-seq
H3K27me3
Basic Gene Annotation Set from GENCODE Version 39lift37 (Ensembl 105)
MEF2B chromatin interactions 10bp (filter activated)
MEF2B chromatin interactions 20bp (filter activated)
MEF2B chromatin interactions 50bp (filter activated)
MEF2B chromatin interactions 200bp (filter activated)
MEF2B binding sites
CTCF,RAD21,SMC3 binding sites


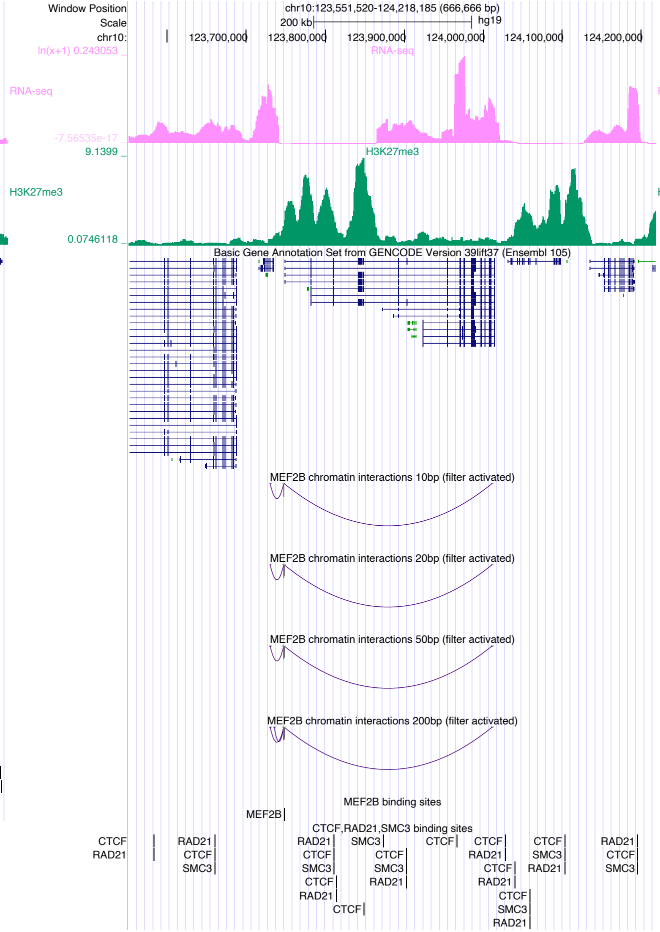

Window Position chr10:123,551,520-124,218,185 (666,666 bp)
Scale 200 kb hg19
chr10: 123,700,000 123,800,000 123,900,000 124,000,000 124,100,000 124,200,000
RNA-seq
H3K27me3
Basic Gene Annotation Set from GENCODE Version 39lift37 (Ensembl 105)
MEF2B chromatin interactions 10bp (filter activated)
MEF2B chromatin interactions 20bp (filter activated)
MEF2B chromatin interactions 50bp (filter activated)
MEF2B chromatin interactions 200bp (filter activated)
MEF2B binding sites
CTCF,RAD21,SMC3 binding sites


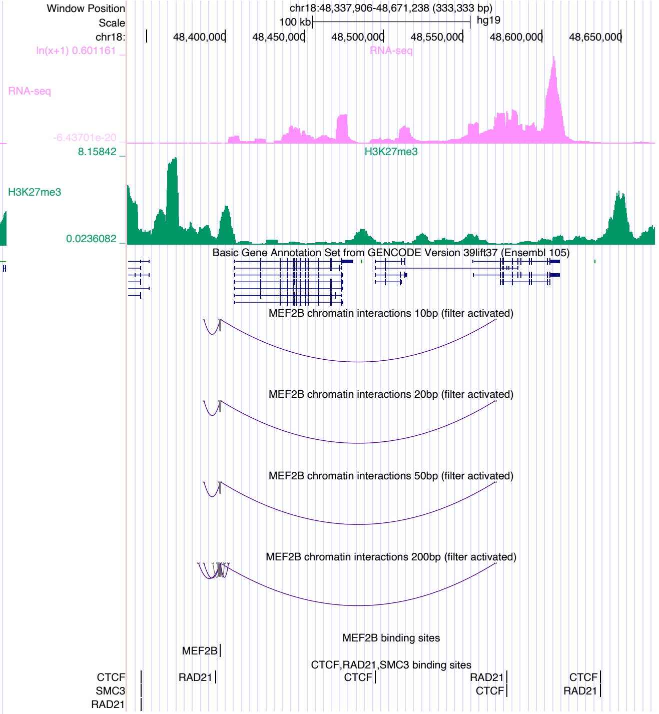

Window Position chr18:48,337,906-48,671,238 (333,333 bp)
Scale 100 kb hg19
chr18: 48,400,000 48,450,000 48,500,000 48,550,000 48,600,000 48,650,000
RNA-seq
H3K27me3
Basic Gene Annotation Set from GENCODE Version 39lift37 (Ensembl 105)
MEF2B chromatin interactions 10bp (filter activated)
MEF2B chromatin interactions 20bp (filter activated)
MEF2B chromatin interactions 50bp (filter activated)
MEF2B chromatin interactions 200bp (filter activated)
MEF2B binding sites
CTCF,RAD21,SMC3 binding sites

### MEIS2

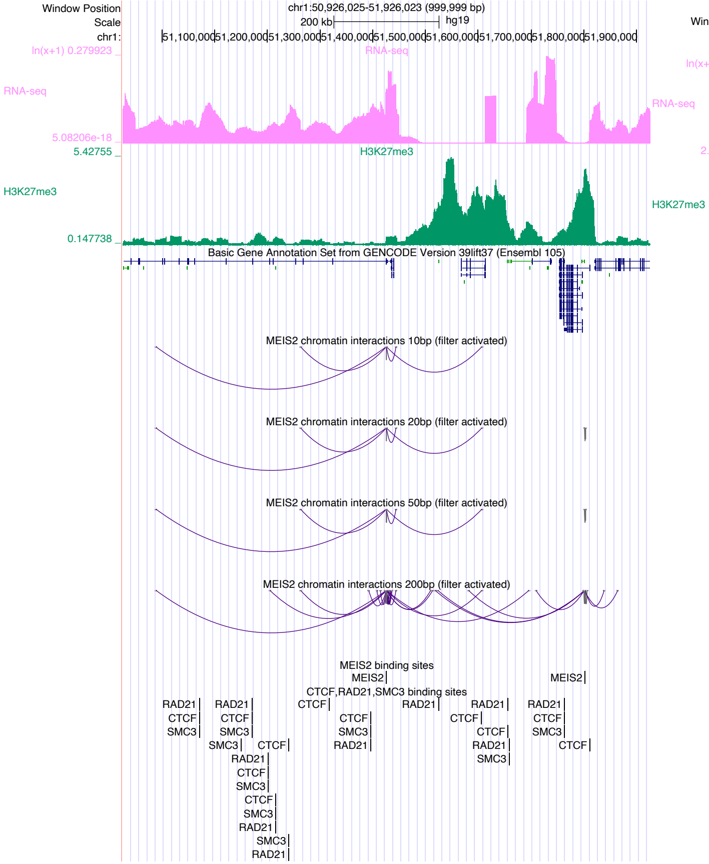

chr1:50,926,025-51,926,023 (999,999 bp)
RNA-seq
H3K27me3
Basic Gene Annotation Set from GENCODE Version 39lift37 (Ensembl 105)
MEIS2 chromatin interactions 10bp (filter activated)
MEIS2 chromatin interactions 20bp (filter activated)
MEIS2 chromatin interactions 50bp (filter activated)
MEIS2 chromatin interactions 200bp (filter activated)
MEIS2 binding sites
CTCF,RAD21,SMC3 binding sites


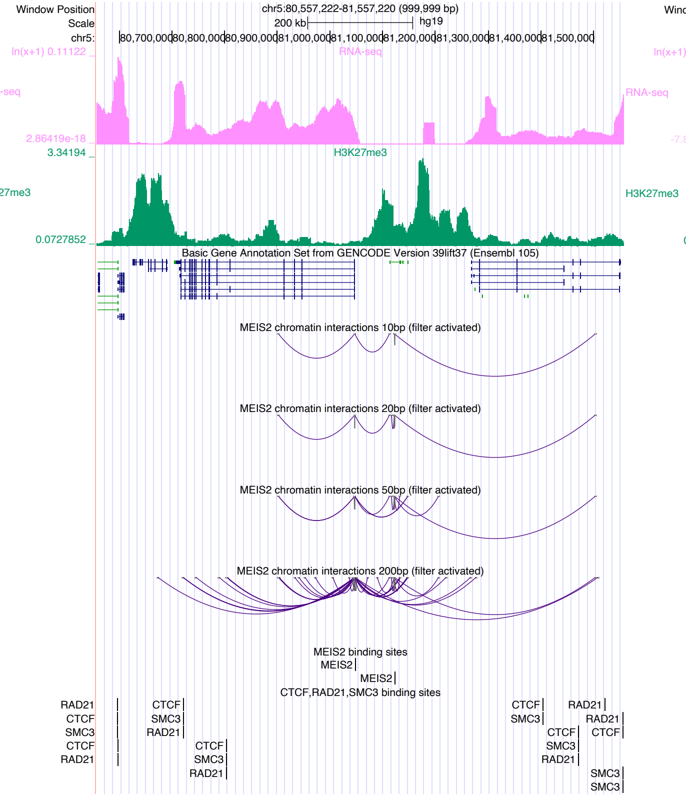

chr5:80,557,222-81,557,220 (999,999 bp)
RNA-seq
H3K27me3
Basic Gene Annotation Set from GENCODE Version 39lift37 (Ensembl 105)
MEIS2 chromatin interactions 10bp (filter activated)
MEIS2 chromatin interactions 20bp (filter activated)
MEIS2 chromatin interactions 50bp (filter activated)
MEIS2 chromatin interactions 200bp (filter activated)
MEIS2 binding sites
CTCF,RAD21,SMC3 binding sites


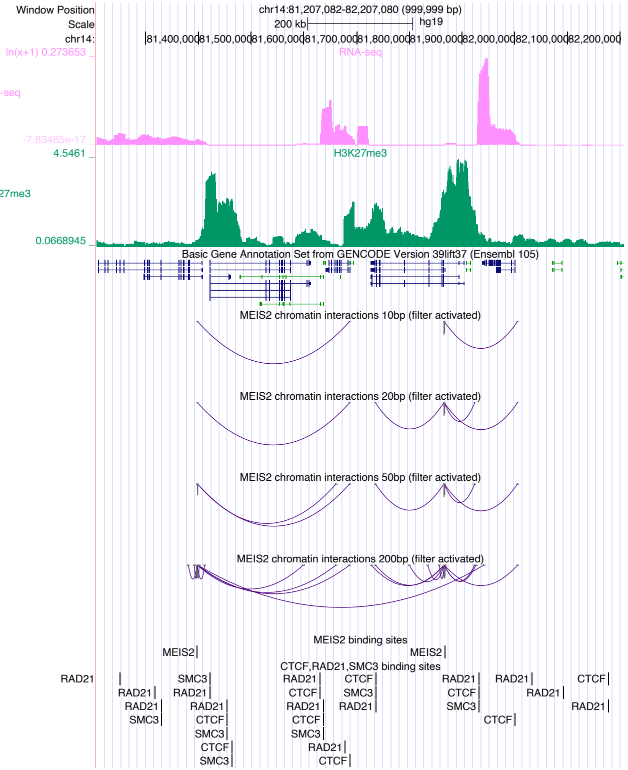

chr14:81,207,082-82,207,080 (999,999 bp)
RNA-seq
H3K27me3
Basic Gene Annotation Set from GENCODE Version 39lift37 (Ensembl 105)
MEIS2 chromatin interactions 10bp (filter activated)
MEIS2 chromatin interactions 20bp (filter activated)
MEIS2 chromatin interactions 50bp (filter activated)
MEIS2 chromatin interactions 200bp (filter activated)
MEIS2 binding sites
CTCF,RAD21,SMC3 binding sites

### MYB

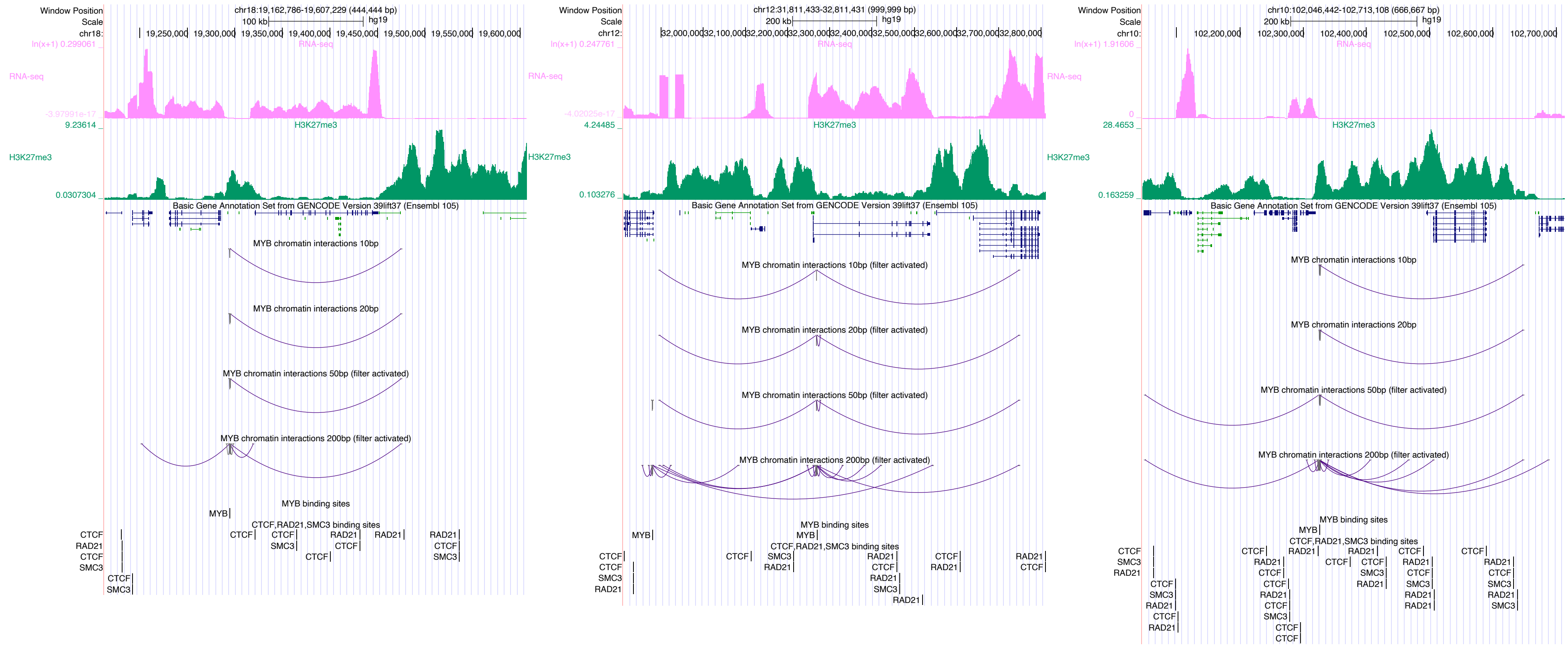

Window Position
Scale
chr18:
chr18:19,162,786-19,607,229 (444,444 bp)
100 kb
hg19
19,250,000
19,300,000
19,350,000
19,400,000
19,450,000
19,500,000
19,550,000
19,600,000
ln(x+1) 0.299061
RNA-seq
-3.97991e-17
9.23614
H3K27me3
0.0307304
Basic Gene Annotation Set from GENCODE Version 39lift37 (Ensembl 105)
MYB chromatin interactions 10bp
MYB chromatin interactions 20bp
MYB chromatin interactions 50bp (filter activated)
MYB chromatin interactions 200bp (filter activated)
MYB binding sites
MYB
CTCF,RAD21,SMC3 binding sites
CTCF
RAD21
SMC3
chr12:31,811,433-32,811,431 (999,999 bp)
200 kb
chr12:
32,000,000
32,100,000
32,200,000
32,300,000
32,400,000
32,500,000
32,600,000
32,700,000
32,800,000
ln(x+1) 0.247761
-4.02025e-17
4.24485
0.103276
MYB chromatin interactions 10bp (filter activated)
MYB chromatin interactions 20bp (filter activated)
chr10:102,046,442-102,713,108 (666,667 bp)
chr10:
102,200,000
102,300,000
102,400,000
102,500,000
102,600,000
102,700,000
ln(x+1) 1.91606
0
28.4653
0.163259

### MYCN

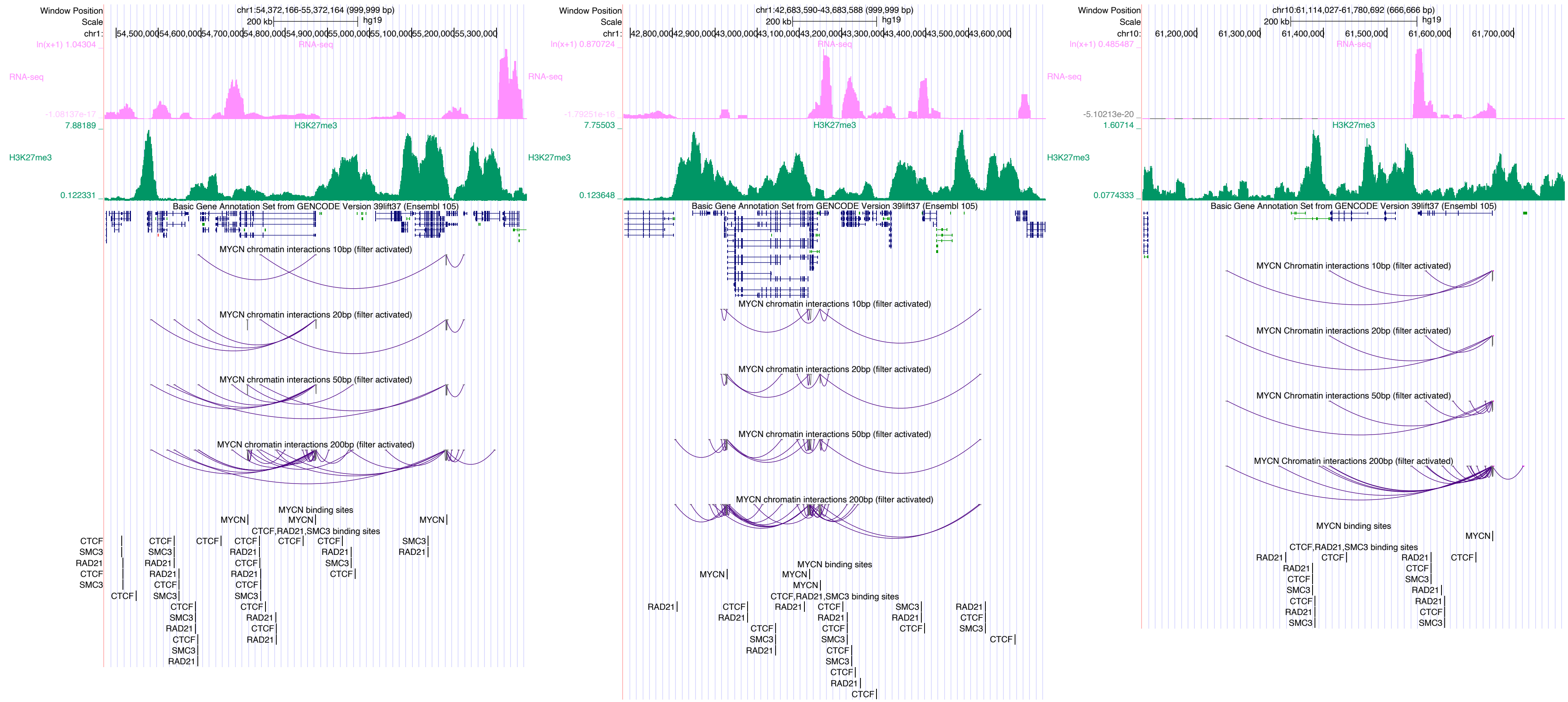
Window Position
Scale
chr1:54,372,166-55,372,164 (999,999 bp)
200 kb
hg19
chr1:
54,500,000
54,600,000
54,700,000
54,800,000
54,900,000
55,000,000
55,100,000
55,200,000
55,300,000
ln(x+1) 1.04304
RNA-seq
-1.08137e-17
7.88189
H3K27me3
0.122331
Basic Gene Annotation Set from GENCODE Version 39lift37 (Ensembl 105)
MYCN chromatin interactions 10bp (filter activated)
MYCN chromatin interactions 20bp (filter activated)
MYCN chromatin interactions 50bp (filter activated)
MYCN chromatin interactions 200bp (filter activated)
MYCN binding sites
CTCF,RAD21,SMC3 binding sites
chr1:42,683,590-43,683,588 (999,999 bp)
42,800,000
42,900,000
43,000,000
43,100,000
43,200,000
43,300,000
43,400,000
43,500,000
43,600,000
ln(x+1) 0.870724
-1.79251e-16
7.75503
0.123648
chr10:61,114,027-61,780,692 (666,666 bp)
chr10:
61,200,000
61,300,000
61,400,000
61,500,000
61,600,000
61,700,000
ln(x+1) 0.485487
-5.10213e-20
1.60714
0.0774333
MYCN Chromatin interactions 10bp (filter activated)
MYCN Chromatin interactions 20bp (filter activated)
MYCN Chromatin interactions 50bp (filter activated)
MYCN Chromatin interactions 200bp (filter activated)
MYCN
CTCF
SMC3
RAD21

### MYOD1

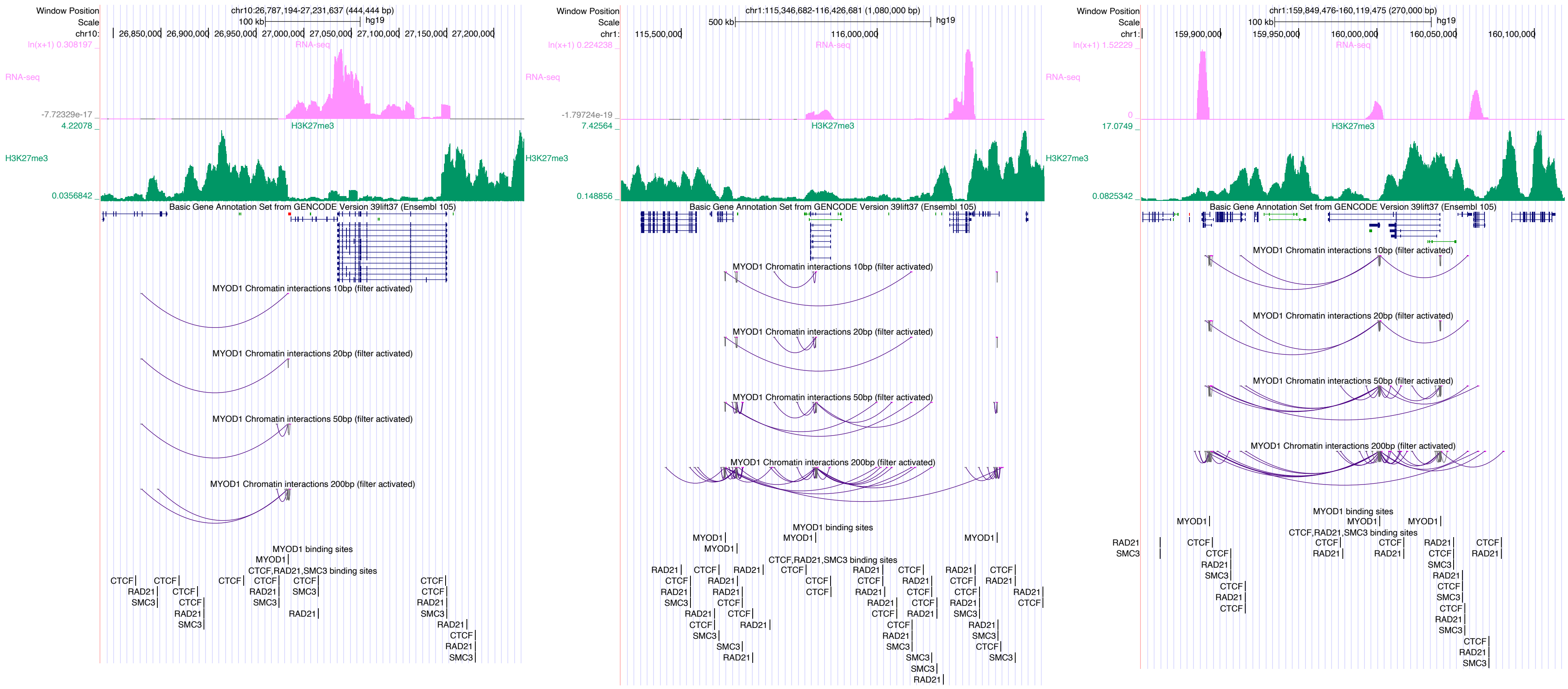

Window Position
chr10:26,787,194-27,231,637 (444,444 bp)
Scale
100 kb
hg19
chr10:
26,850,000
26,900,000
26,950,000
27,000,000
27,050,000
27,100,000
27,150,000
27,200,000
RNA-seq
H3K27me3
Basic Gene Annotation Set from GENCODE Version 39lift37 (Ensembl 105)
MYOD1 Chromatin interactions 10bp (filter activated)
MYOD1 Chromatin interactions 20bp (filter activated)
MYOD1 Chromatin interactions 50bp (filter activated)
MYOD1 Chromatin interactions 200bp (filter activated)
MYOD1 binding sites
CTCF,RAD21,SMC3 binding sites
chr1:115,346,682-116,426,681 (1,080,000 bp)
500 kb
115,500,000
116,000,000
chr1:159,849,476-160,119,475 (270,000 bp)
159,900,000
159,950,000
160,000,000
160,050,000
160,100,000

### MYOG

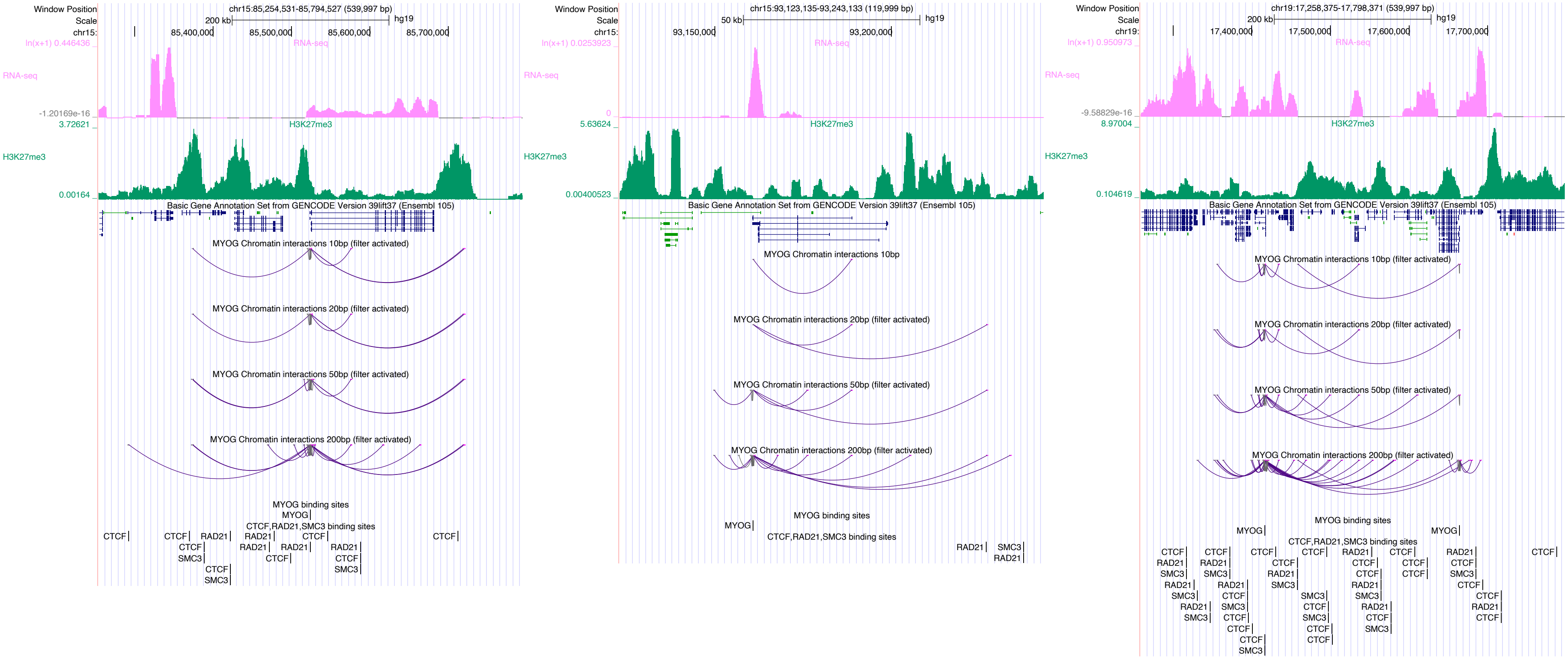

Window Position
Scale
chr15:85,254,531-85,794,527 (539,997 bp)
200 kb
hg19
RNA-seq
H3K27me3
Basic Gene Annotation Set from GENCODE Version 39lift37 (Ensembl 105)
MYOG Chromatin interactions 10bp (filter activated)
MYOG Chromatin interactions 20bp (filter activated)
MYOG Chromatin interactions 50bp (filter activated)
MYOG Chromatin interactions 200bp (filter activated)
MYOG binding sites
CTCF,RAD21,SMC3 binding sites
chr15:93,123,135-93,243,133 (119,999 bp)
50 kb
MYOG Chromatin interactions 10bp
chr19:17,258,375-17,798,371 (539,997 bp)

### NFYC

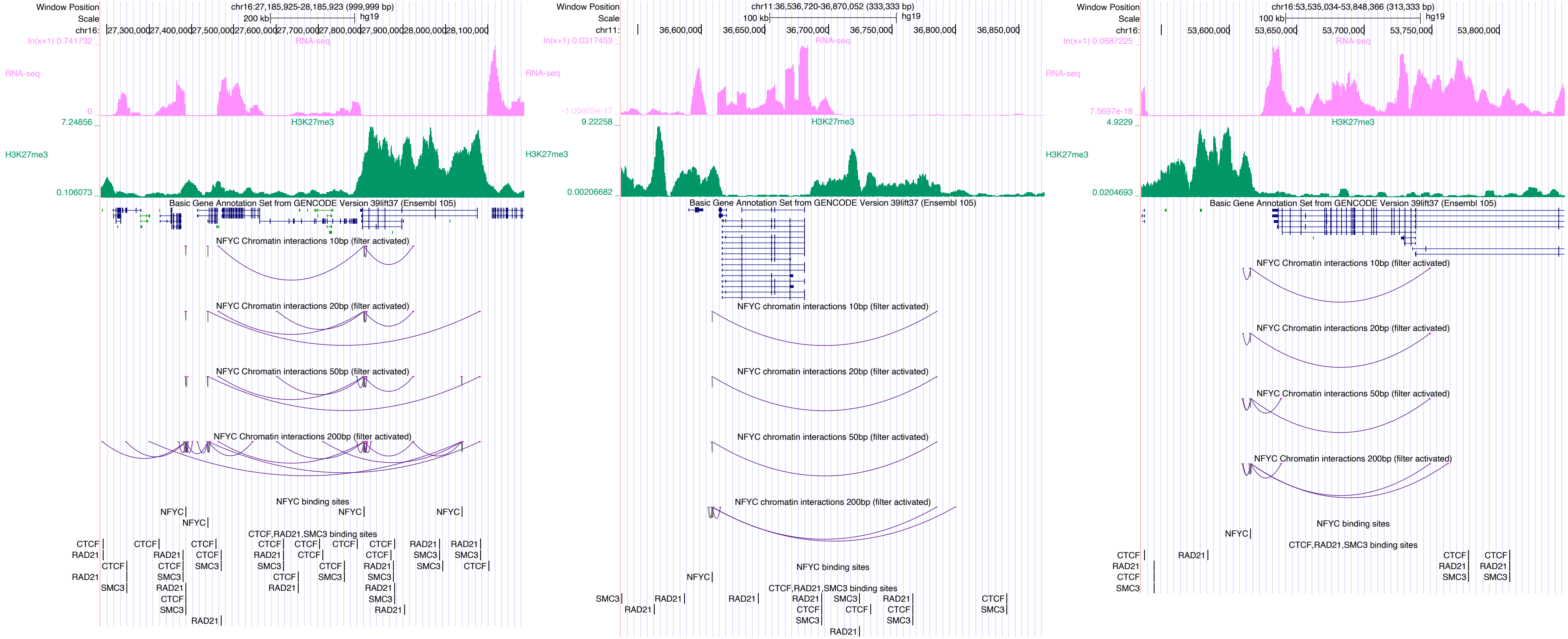

Window Position
Scale
chr16:27,185,925-28,185,923 (999,999 bp)
200 kb
hg19
RNA-seq
H3K27me3
Basic Gene Annotation Set from GENCODE Version 39lift37 (Ensembl 105)
NFYC Chromatin interactions 10bp (filter activated)
NFYC Chromatin interactions 20bp (filter activated)
NFYC Chromatin interactions 50bp (filter activated)
NFYC Chromatin interactions 200bp (filter activated)
NFYC binding sites
CTCF,RAD21,SMC3 binding sites
chr11:36,536,720-36,870,052 (333,333 bp)
100 kb
chr16:53,535,034-53,848,366 (313,333 bp)

# NR2C2

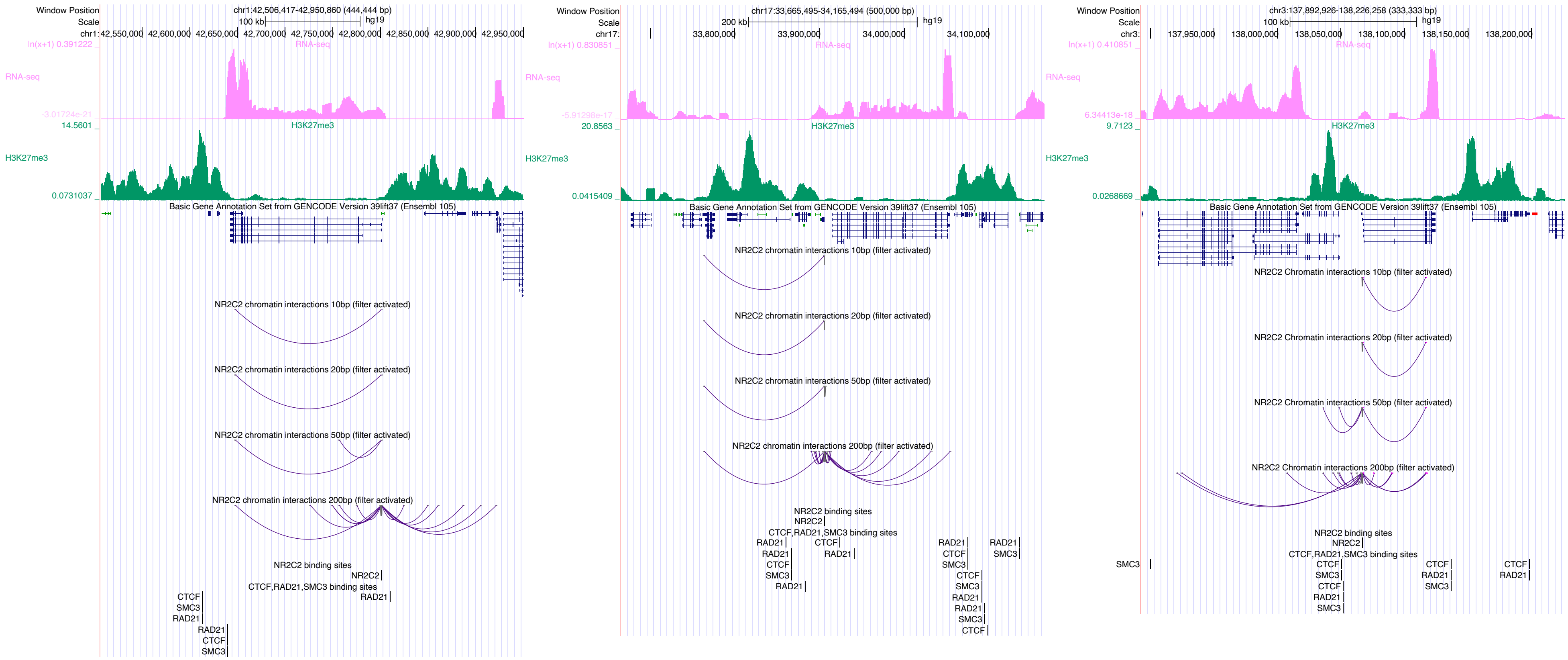

Window Position
chr1:42,506,417-42,950,860 (444,444 bp)
Scale
100 kb
hg19
RNA-seq
H3K27me3
Basic Gene Annotation Set from GENCODE Version 39lift37 (Ensembl 105)
NR2C2 chromatin interactions 10bp (filter activated)
NR2C2 chromatin interactions 20bp (filter activated)
NR2C2 chromatin interactions 50bp (filter activated)
NR2C2 chromatin interactions 200bp (filter activated)
NR2C2 binding sites
CTCF,RAD21,SMC3 binding sites
chr17:33,665,495-34,165,494 (500,000 bp)
200 kb
chr3:137,892,926-138,226,258 (333,333 bp)
NR2C2 Chromatin interactions 10bp (filter activated)
NR2C2 Chromatin interactions 20bp (filter activated)
NR2C2 Chromatin interactions 50bp (filter activated)
NR2C2 Chromatin interactions 200bp (filter activated)

# NR2F2

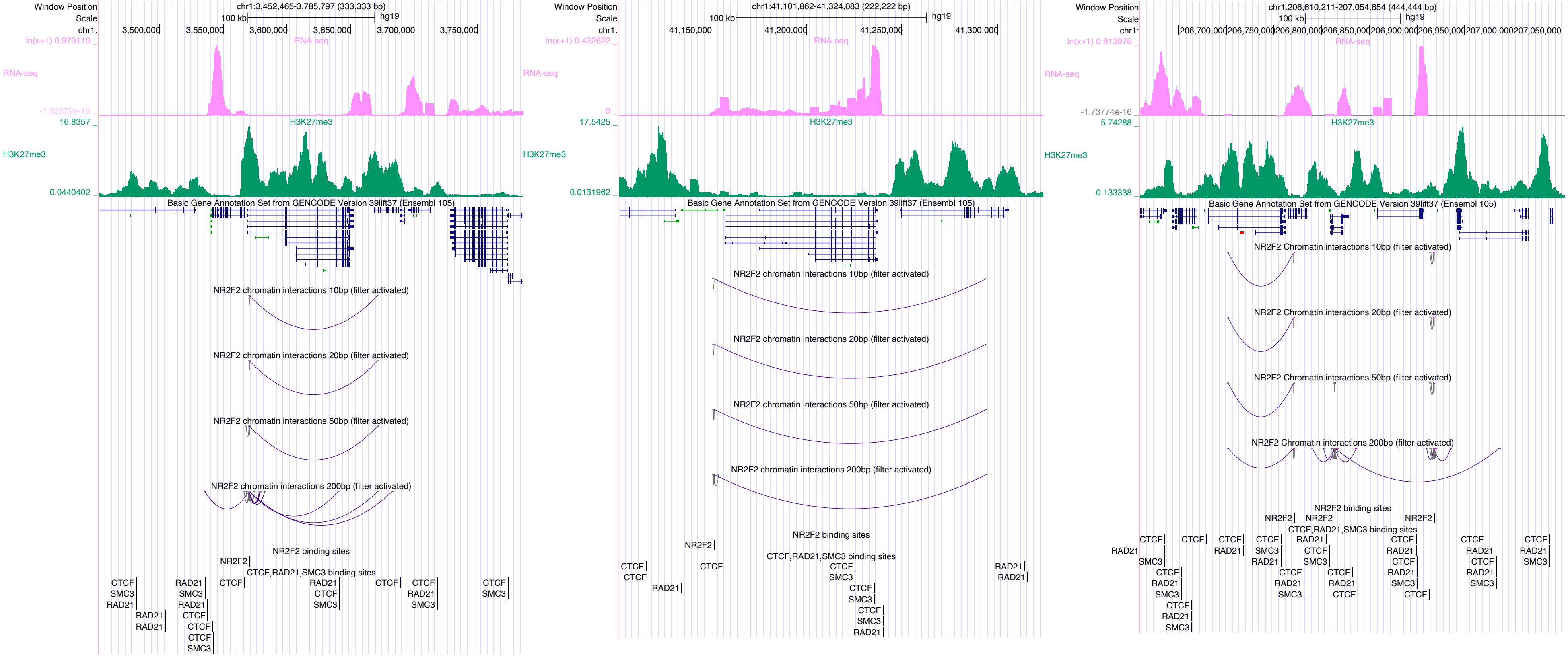

Window Position
Scale
chr1:
chr1:3,452,465-3,785,797 (333,333 bp)
100 kb
hg19
3,500,000
3,550,000
3,600,000
3,650,000
3,700,000
3,750,000
ln(x+1) 0.979119
RNA-seq
-1.52879e-19
16.8357
H3K27me3
0.0440402
Basic Gene Annotation Set from GENCODE Version 39lift37 (Ensembl 105)
NR2F2 chromatin interactions 10bp (filter activated)
NR2F2 chromatin interactions 20bp (filter activated)
NR2F2 chromatin interactions 50bp (filter activated)
NR2F2 chromatin interactions 200bp (filter activated)
NR2F2 binding sites
CTCF,RAD21,SMC3 binding sites
chr1:41,101,862-41,324,083 (222,222 bp)
41,150,000
41,200,000
41,250,000
41,300,000
ln(x+1) 0.432622
0
17.5425
0.0131962
chr1:206,610,211-207,054,654 (444,444 bp)
206,700,000
206,750,000
206,800,000
206,850,000
206,900,000
206,950,000
207,000,000
207,050,000
ln(x+1) 0.813976
-1.73774e-16
5.74288
0.133338
NR2F2 Chromatin interactions 10bp (filter activated)
NR2F2 Chromatin interactions 20bp (filter activated)
NR2F2 Chromatin interactions 50bp (filter activated)
NR2F2 Chromatin interactions 200bp (filter activated)
NR2F2
CTCF
RAD21
SMC3

# NR3C1

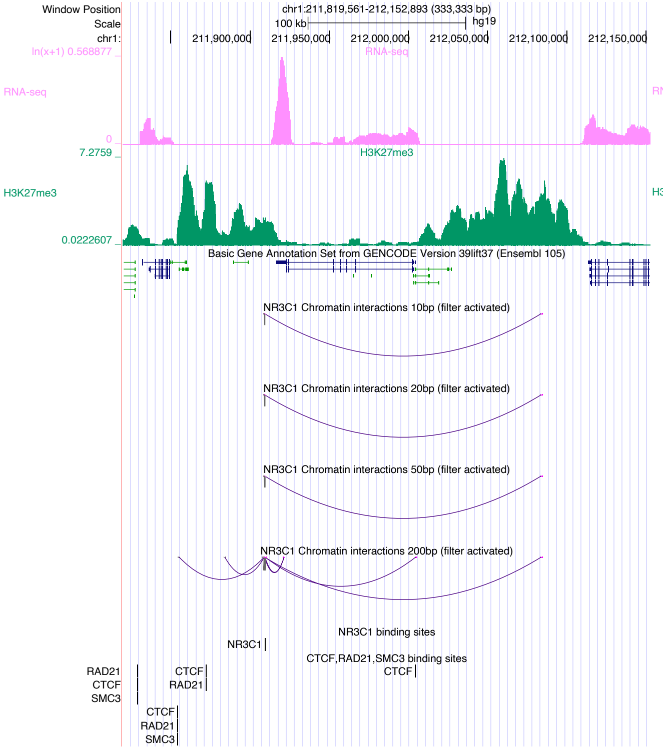

Window Position chr1:211,819,561-212,152,893 (333,333 bp)
Scale 100 kb hg19
chr1: 211,900,000 211,950,000 212,000,000 212,050,000 212,100,000 212,150,000
RNA-seq
H3K27me3
Basic Gene Annotation Set from GENCODE Version 39lift37 (Ensembl 105)
NR3C1 Chromatin interactions 10bp (filter activated)
NR3C1 Chromatin interactions 20bp (filter activated)
NR3C1 Chromatin interactions 50bp (filter activated)
NR3C1 Chromatin interactions 200bp (filter activated)
NR3C1 binding sites
CTCF,RAD21,SMC3 binding sites


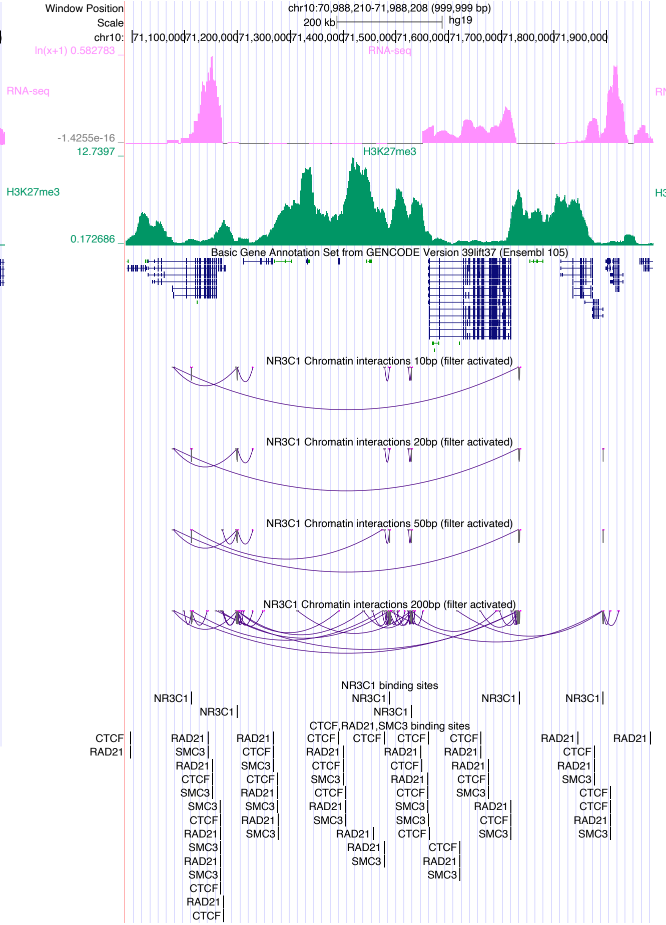

Window Position chr10:70,988,210-71,988,208 (999,999 bp)
Scale 200 kb hg19
chr10: 71,100,000 71,200,000 71,300,000 71,400,000 71,500,000 71,600,000 71,700,000 71,800,000 71,900,000
RNA-seq
H3K27me3
Basic Gene Annotation Set from GENCODE Version 39lift37 (Ensembl 105)
NR3C1 Chromatin interactions 10bp (filter activated)
NR3C1 Chromatin interactions 20bp (filter activated)
NR3C1 Chromatin interactions 50bp (filter activated)
NR3C1 Chromatin interactions 200bp (filter activated)
NR3C1 binding sites
CTCF,RAD21,SMC3 binding sites


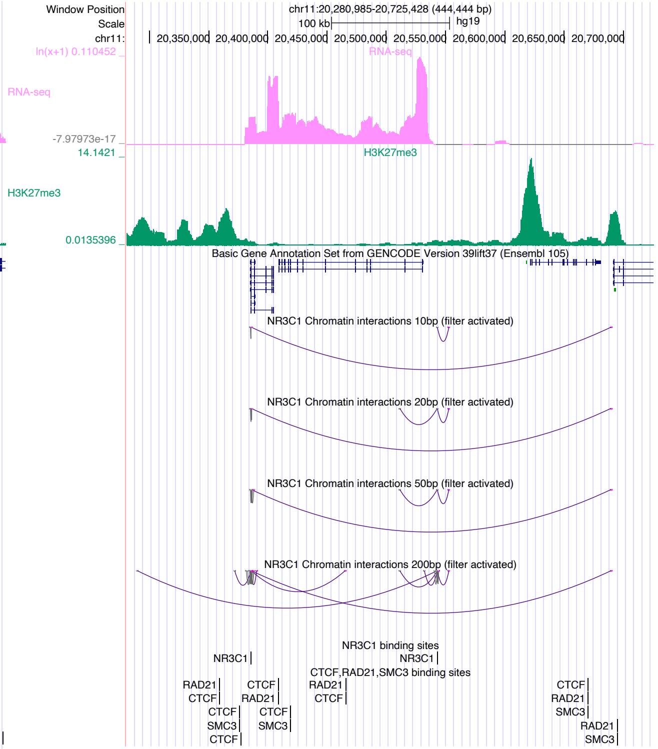

Window Position chr11:20,280,985-20,725,428 (444,444 bp)
Scale 100 kb hg19
chr11: 20,350,000 20,400,000 20,450,000 20,500,000 20,550,000 20,600,000 20,650,000 20,700,000
RNA-seq
H3K27me3
Basic Gene Annotation Set from GENCODE Version 39lift37 (Ensembl 105)
NR3C1 Chromatin interactions 10bp (filter activated)
NR3C1 Chromatin interactions 20bp (filter activated)
NR3C1 Chromatin interactions 50bp (filter activated)
NR3C1 Chromatin interactions 200bp (filter activated)
NR3C1 binding sites
CTCF,RAD21,SMC3 binding sites

# NR5A2

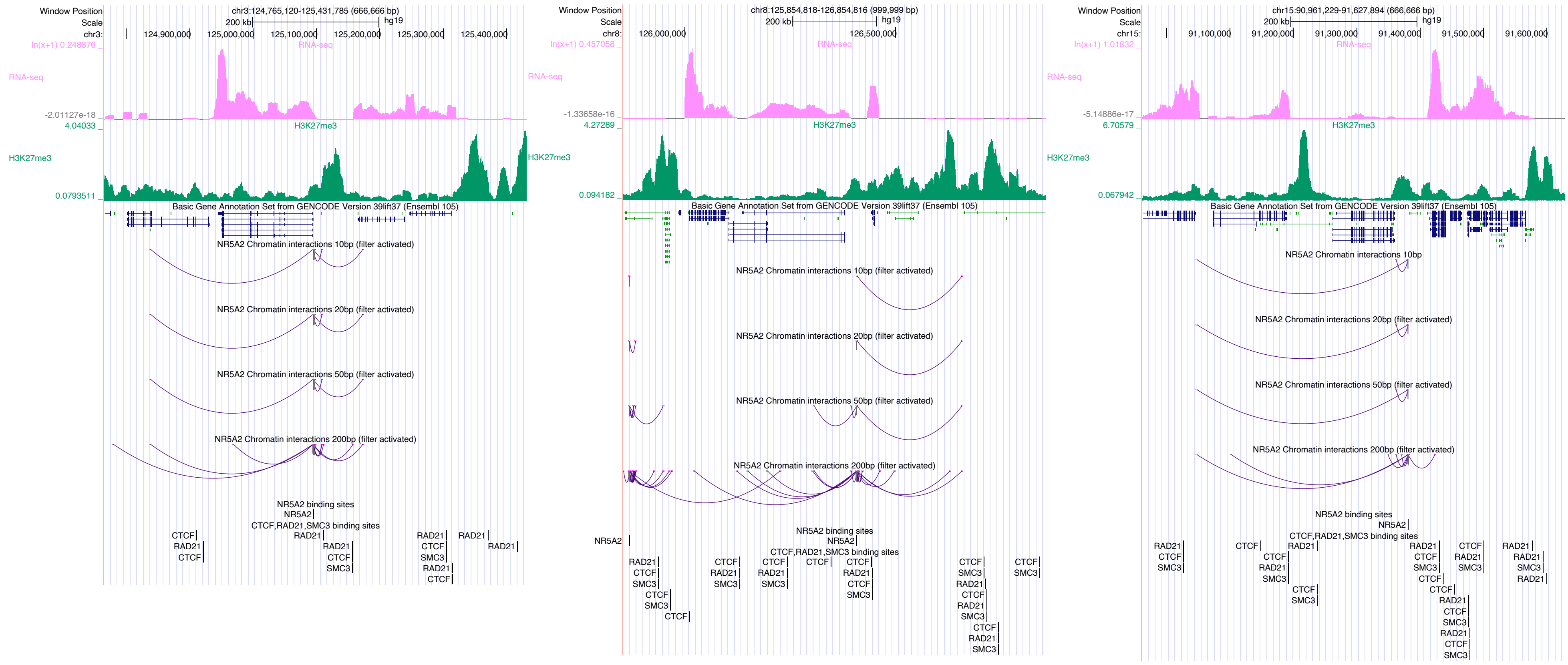

Window Position
chr3:124,765,120-125,431,785 (666,666 bp)
Scale
200 kb
hg19
RNA-seq
H3K27me3
Basic Gene Annotation Set from GENCODE Version 39lift37 (Ensembl 105)
NR5A2 Chromatin interactions 10bp (filter activated)
NR5A2 Chromatin interactions 20bp (filter activated)
NR5A2 Chromatin interactions 50bp (filter activated)
NR5A2 Chromatin interactions 200bp (filter activated)
NR5A2 binding sites
CTCF,RAD21,SMC3 binding sites
chr8:125,854,818-126,854,816 (999,999 bp)
chr15:90,961,229-91,627,894 (666,666 bp)
NR5A2 Chromatin interactions 10bp

### PAX5

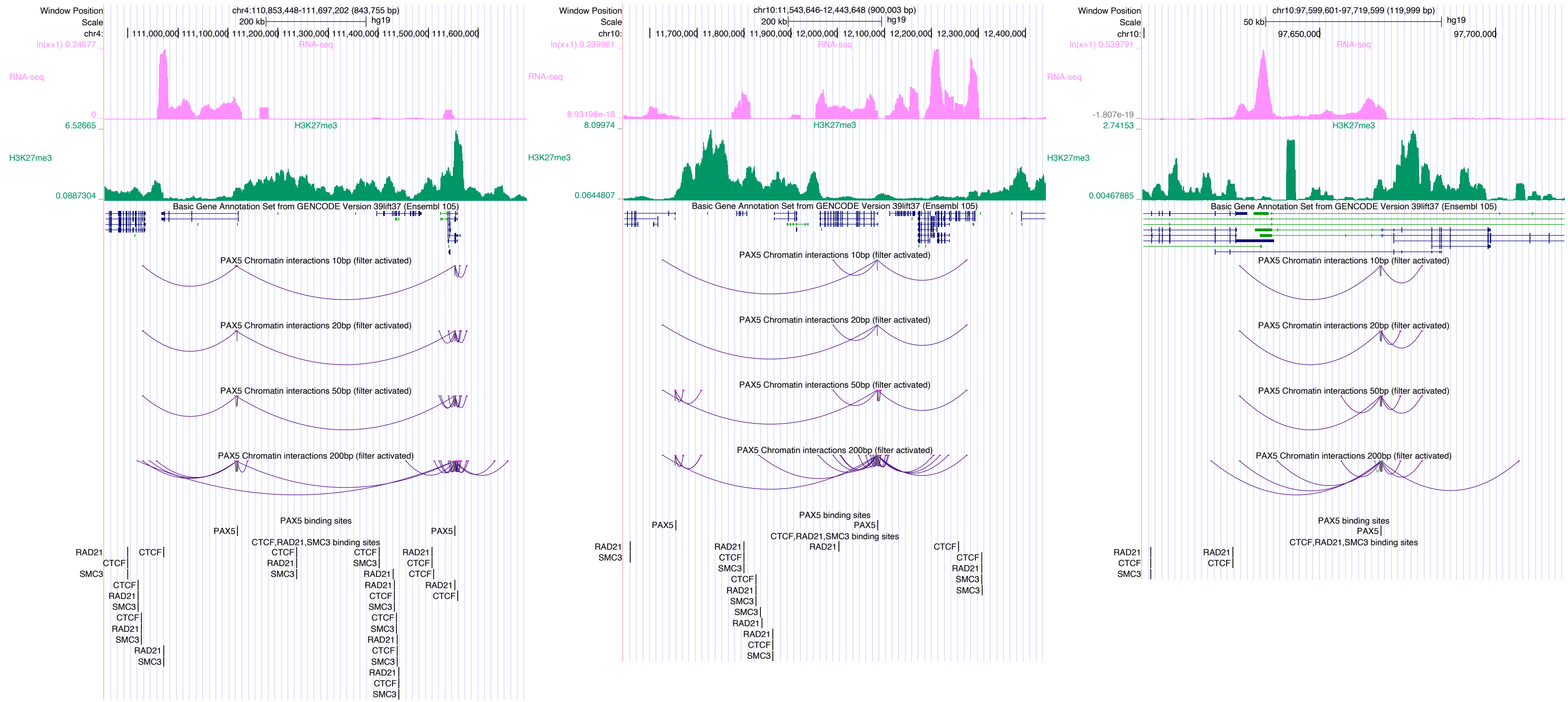

Window Position
chr4:110,853,448-111,697,202 (843,755 bp)
Scale
200 kb
hg19
chr4:
111,000,000 111,100,000 111,200,000 111,300,000 111,400,000 111,500,000 111,600,000
ln(x+1) 0.24677
RNA-seq
0
6.52665
H3K27me3
0.0887304
Basic Gene Annotation Set from GENCODE Version 39lift37 (Ensembl 105)
PAX5 Chromatin interactions 10bp (filter activated)
PAX5 Chromatin interactions 20bp (filter activated)
PAX5 Chromatin interactions 50bp (filter activated)
PAX5 Chromatin interactions 200bp (filter activated)
PAX5 binding sites
PAX5
CTCF,RAD21,SMC3 binding sites
RAD21
CTCF
SMC3
Window Position
chr10:11,543,646-12,443,648 (900,003 bp)
200 kb
chr10:
11,700,000 11,800,000 11,900,000 12,000,000 12,100,000 12,200,000 12,300,000 12,400,000
ln(x+1) 0.239961
8.93198e-18
8.09974
0.0644807
Window Position
chr10:97,599,601-97,719,599 (119,999 bp)
50 kb
97,650,000
97,700,000
ln(x+1) 0.539791
-1.807e-19
2.74153
0.00467885

### PBX2

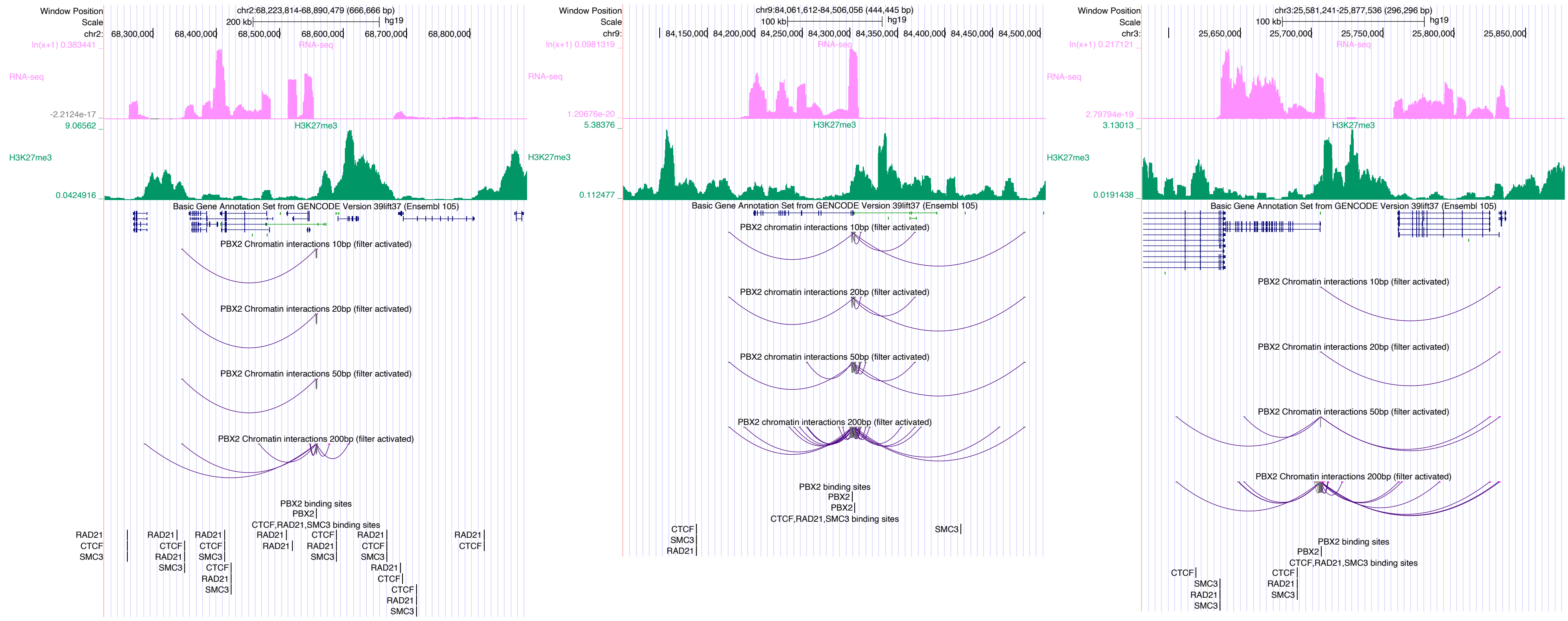
Window Position chr2:68,223,814-68,890,479 (666,666 bp)
Scale 200 kb hg19
chr2: 68,300,000 68,400,000 68,500,000 68,600,000 68,700,000 68,800,000
RNA-seq
H3K27me3
Basic Gene Annotation Set from GENCODE Version 39lift37 (Ensembl 105)
PBX2 Chromatin interactions 10bp (filter activated)
PBX2 Chromatin interactions 20bp (filter activated)
PBX2 Chromatin interactions 50bp (filter activated)
PBX2 Chromatin interactions 200bp (filter activated)
PBX2 binding sites
CTCF,RAD21,SMC3 binding sites
Window Position chr9:84,061,612-84,506,056 (444,445 bp)
Scale 100 kb hg19
chr9: 84,150,000 84,200,000 84,250,000 84,300,000 84,350,000 84,400,000 84,450,000 84,500,000
PBX2 chromatin interactions 10bp (filter activated)
PBX2 chromatin interactions 20bp (filter activated)
PBX2 chromatin interactions 50bp (filter activated)
PBX2 chromatin interactions 200bp (filter activated)
Window Position chr3:25,581,241-25,877,536 (296,296 bp)
Scale 100 kb hg19
chr3: 25,650,000 25,700,000 25,750,000 25,800,000 25,850,000

### PRDM9

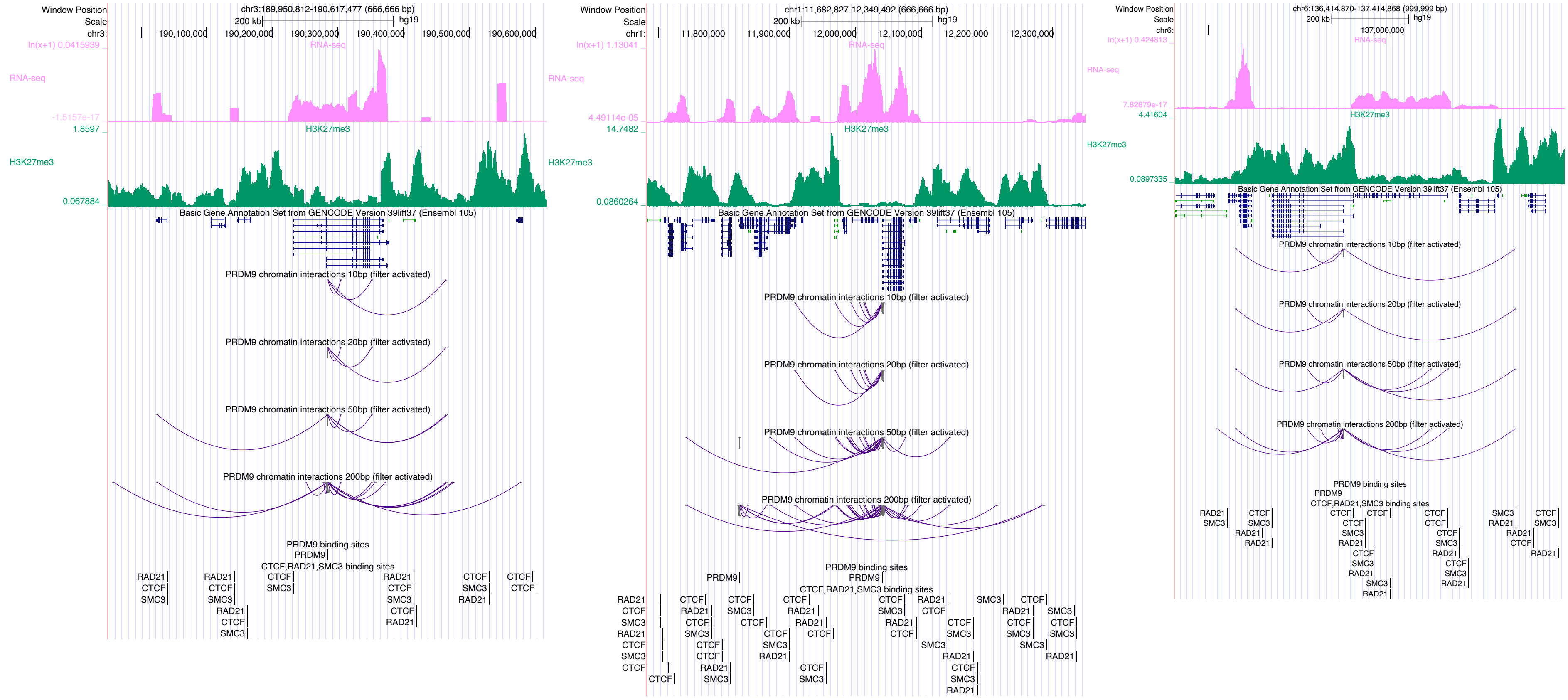

Window Position
Scale
chr3:189,950,812-190,617,477 (666,666 bp)
200 kb
hg19
190,100,000
190,200,000
190,300,000
190,400,000
190,500,000
190,600,000
RNA-seq
H3K27me3
Basic Gene Annotation Set from GENCODE Version 39lift37 (Ensembl 105)
PRDM9 chromatin interactions 10bp (filter activated)
PRDM9 chromatin interactions 20bp (filter activated)
PRDM9 chromatin interactions 50bp (filter activated)
PRDM9 chromatin interactions 200bp (filter activated)
PRDM9 binding sites
CTCF,RAD21,SMC3 binding sites
chr1:11,682,827-12,349,492 (666,666 bp)
11,800,000
11,900,000
12,000,000
12,100,000
12,200,000
12,300,000
chr6:136,414,870-137,414,868 (999,999 bp)
137,000,000

### RBPJ

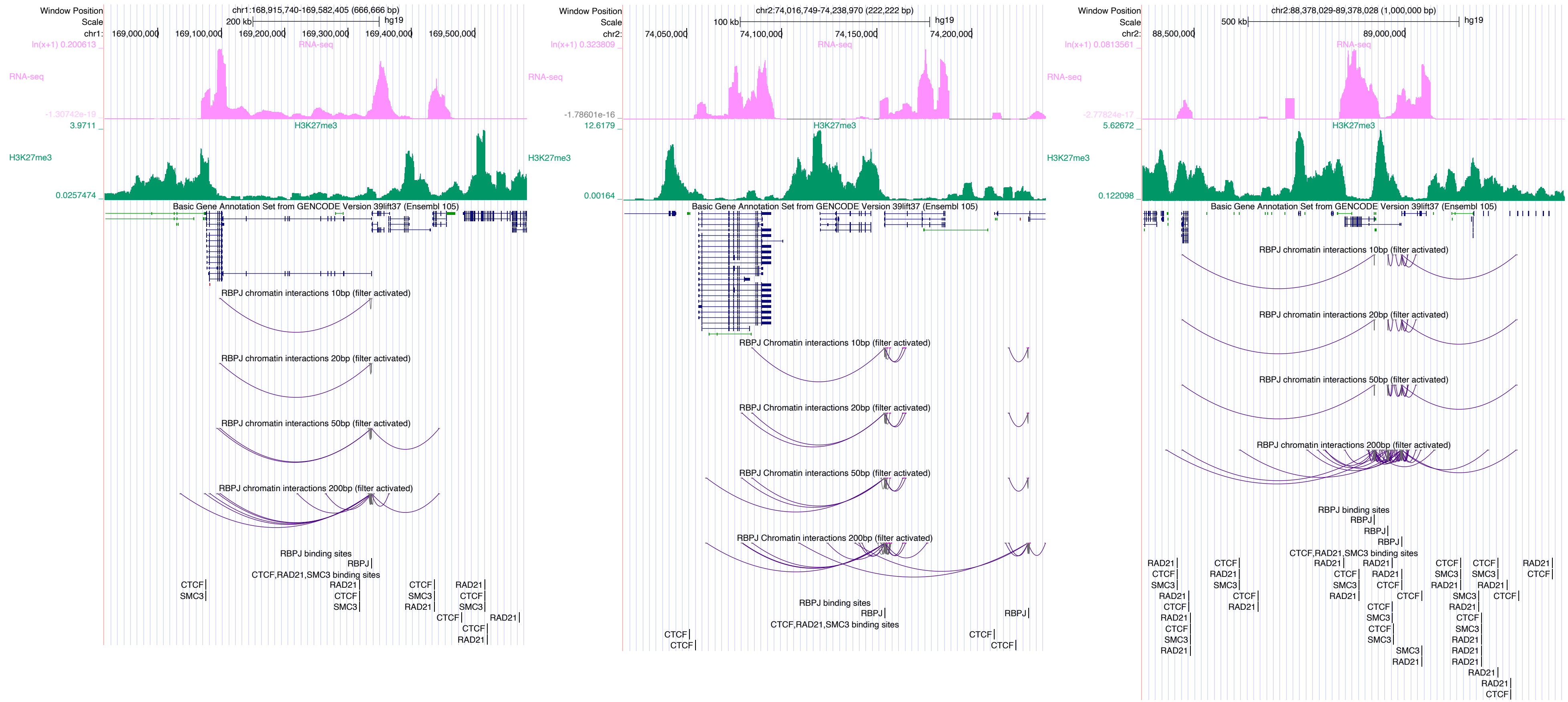

Window Position
Scale
chr1:168,915,740-169,582,405 (666,666 bp)
200 kb
hg19
chr1: 169,000,000 169,100,000 169,200,000 169,300,000 169,400,000 169,500,000
RNA-seq
H3K27me3
Basic Gene Annotation Set from GENCODE Version 39lift37 (Ensembl 105)
RBPJ chromatin interactions 10bp (filter activated)
RBPJ chromatin interactions 20bp (filter activated)
RBPJ chromatin interactions 50bp (filter activated)
RBPJ chromatin interactions 200bp (filter activated)
RBPJ binding sites
RBPJ
CTCF,RAD21,SMC3 binding sites
CTCF
SMC3
RAD21
chr2:74,016,749-74,238,970 (222,222 bp)
100 kb
chr2: 74,050,000 74,100,000 74,150,000 74,200,000
RBPJ Chromatin interactions 10bp (filter activated)
RBPJ Chromatin interactions 20bp (filter activated)
RBPJ Chromatin interactions 50bp (filter activated)
RBPJ Chromatin interactions 200bp (filter activated)
chr2:88,378,029-89,378,028 (1,000,000 bp)
500 kb
chr2: 88,500,000 89,000,000

### REST

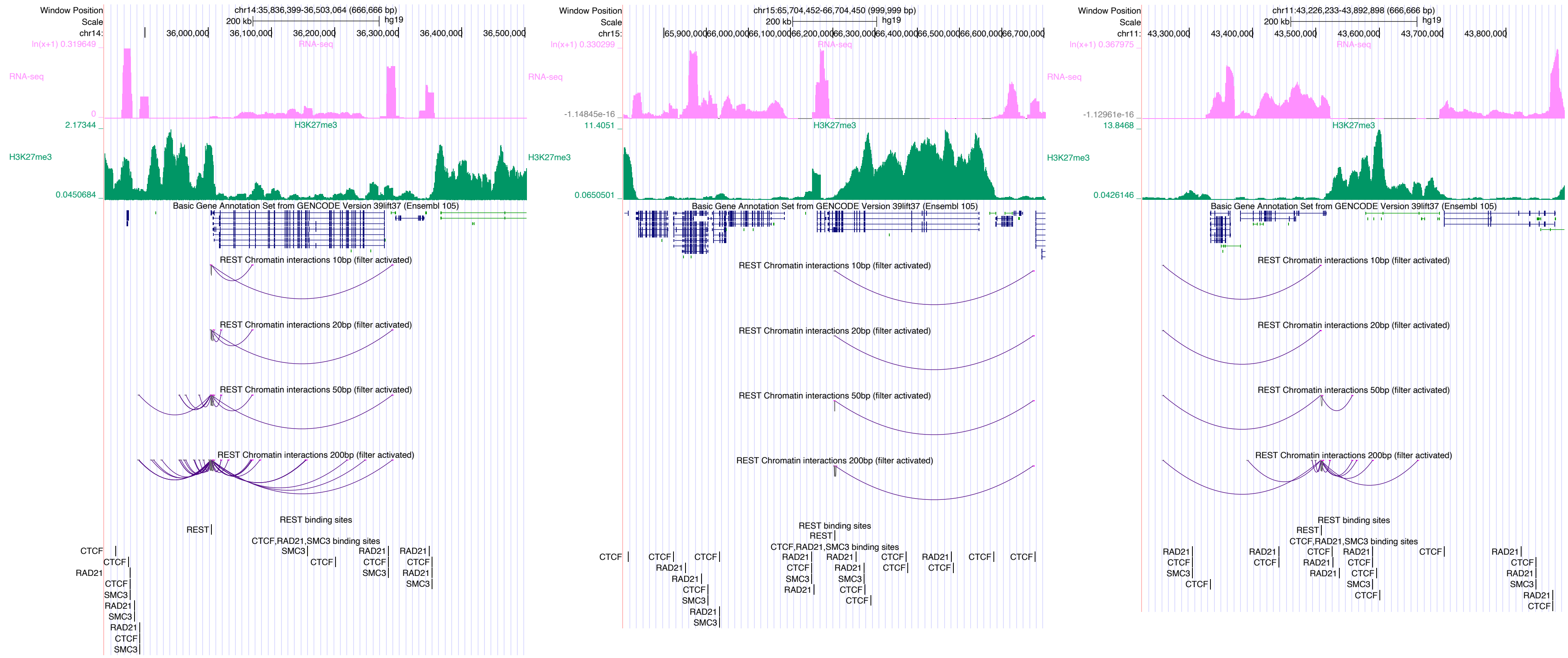
chr14:35,836,399-36,503,064 (666,666 bp)
chr15:65,704,452-66,704,450 (999,999 bp)
chr11:43,226,233-43,892,898 (666,666 bp)
RNA-seq
H3K27me3
Basic Gene Annotation Set from GENCODE Version 39lift37 (Ensembl 105)
REST Chromatin interactions 10bp (filter activated)
REST Chromatin interactions 20bp (filter activated)
REST Chromatin interactions 50bp (filter activated)
REST Chromatin interactions 200bp (filter activated)
REST binding sites
CTCF,RAD21,SMC3 binding sites

### RUNX1

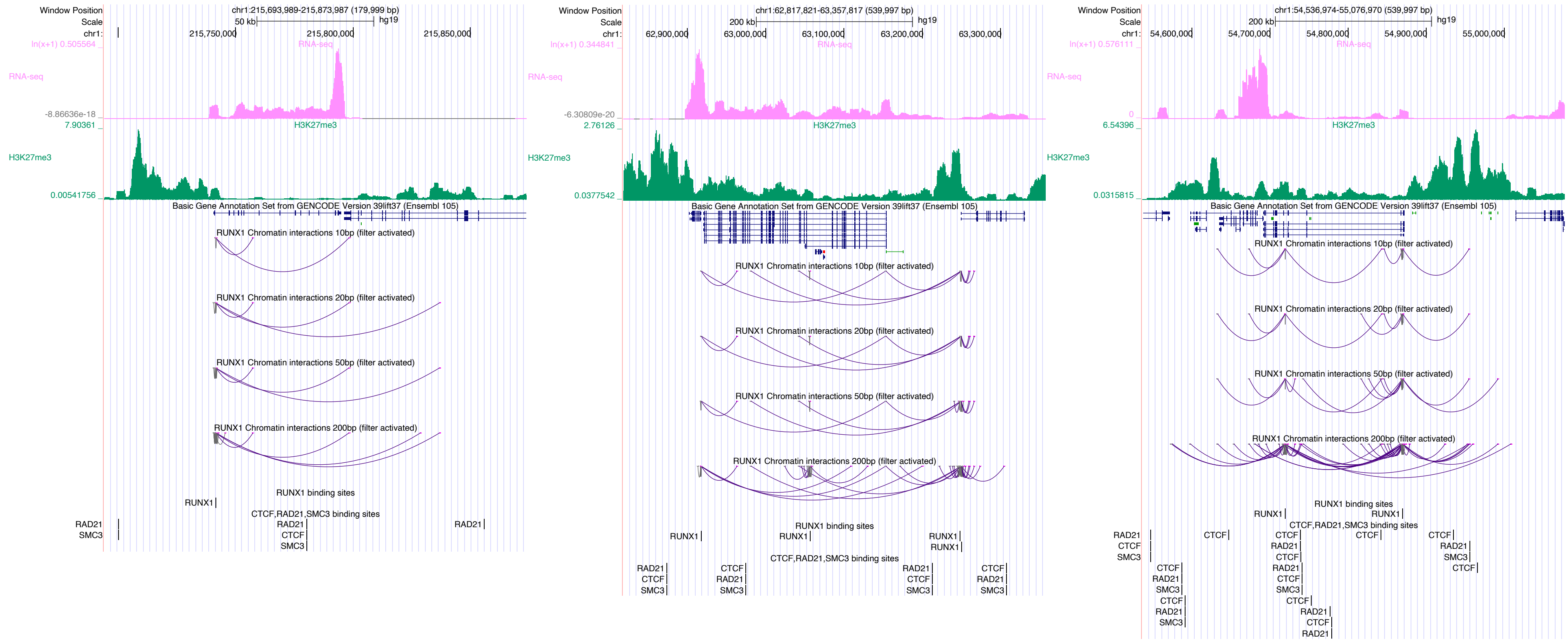
Window Position chr1:215,693,989-215,873,987 (179,999 bp)
Scale 50 kb hg19
chr1: 215,750,000 215,800,000 215,850,000
RNA-seq
H3K27me3
Basic Gene Annotation Set from GENCODE Version 39lift37 (Ensembl 105)
RUNX1 Chromatin interactions 10bp (filter activated)
RUNX1 Chromatin interactions 20bp (filter activated)
RUNX1 Chromatin interactions 50bp (filter activated)
RUNX1 Chromatin interactions 200bp (filter activated)
RUNX1 binding sites
CTCF,RAD21,SMC3 binding sites
Window Position chr1:62,817,821-63,357,817 (539,997 bp)
Scale 200 kb hg19
chr1: 62,900,000 63,000,000 63,100,000 63,200,000 63,300,000
Window Position chr1:54,536,974-55,076,970 (539,997 bp)
Scale 200 kb hg19
chr1: 54,600,000 54,700,000 54,800,000 54,900,000 55,000,000

### RUNX3

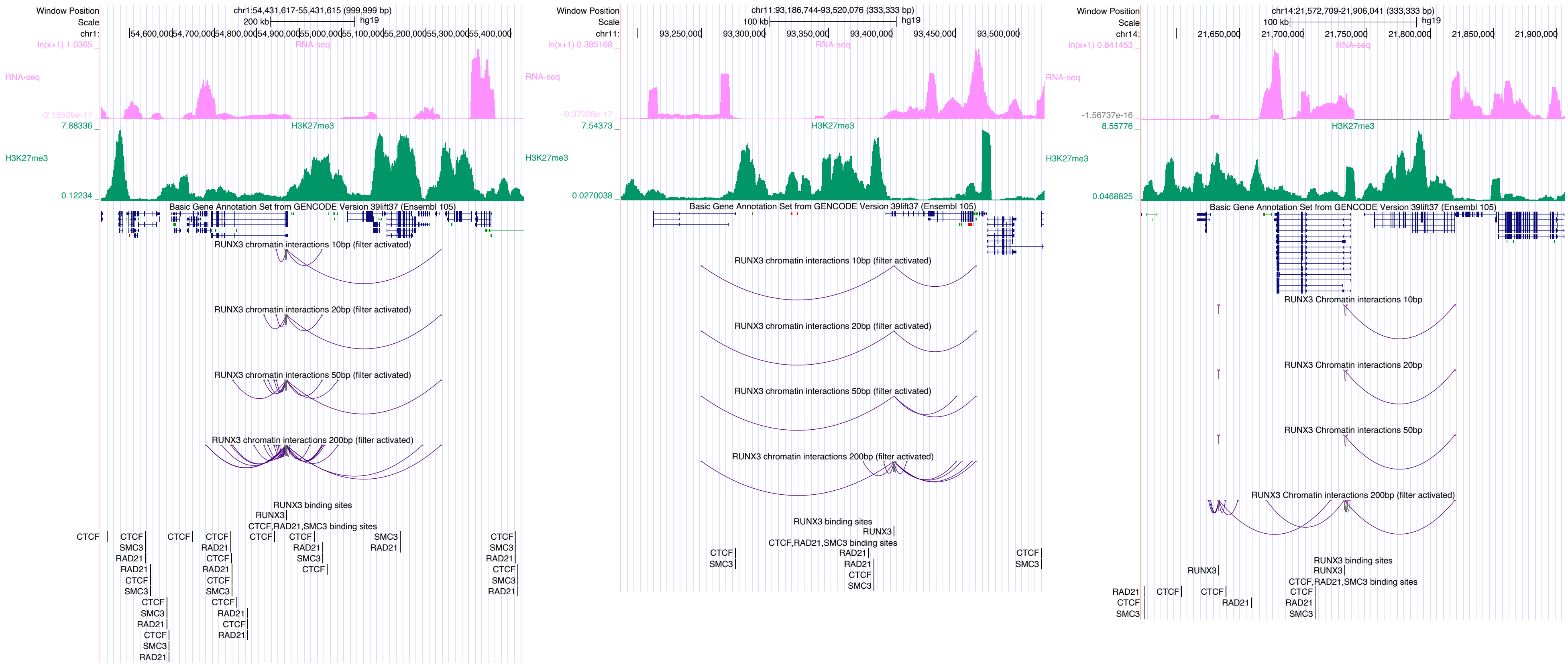
Window Position
chr1:54,431,617-55,431,615 (999,999 bp)
RNA-seq
H3K27me3
Basic Gene Annotation Set from GENCODE Version 39lift37 (Ensembl 105)
RUNX3 chromatin interactions 10bp (filter activated)
RUNX3 chromatin interactions 20bp (filter activated)
RUNX3 chromatin interactions 50bp (filter activated)
RUNX3 chromatin interactions 200bp (filter activated)
RUNX3 binding sites
CTCF,RAD21,SMC3 binding sites
chr11:93,186,744-93,520,076 (333,333 bp)
chr14:21,572,709-21,906,041 (333,333 bp)
RUNX3 Chromatin interactions 10bp
RUNX3 Chromatin interactions 20bp
RUNX3 Chromatin interactions 50bp
RUNX3 Chromatin interactions 200bp (filter activated)

### RXRA

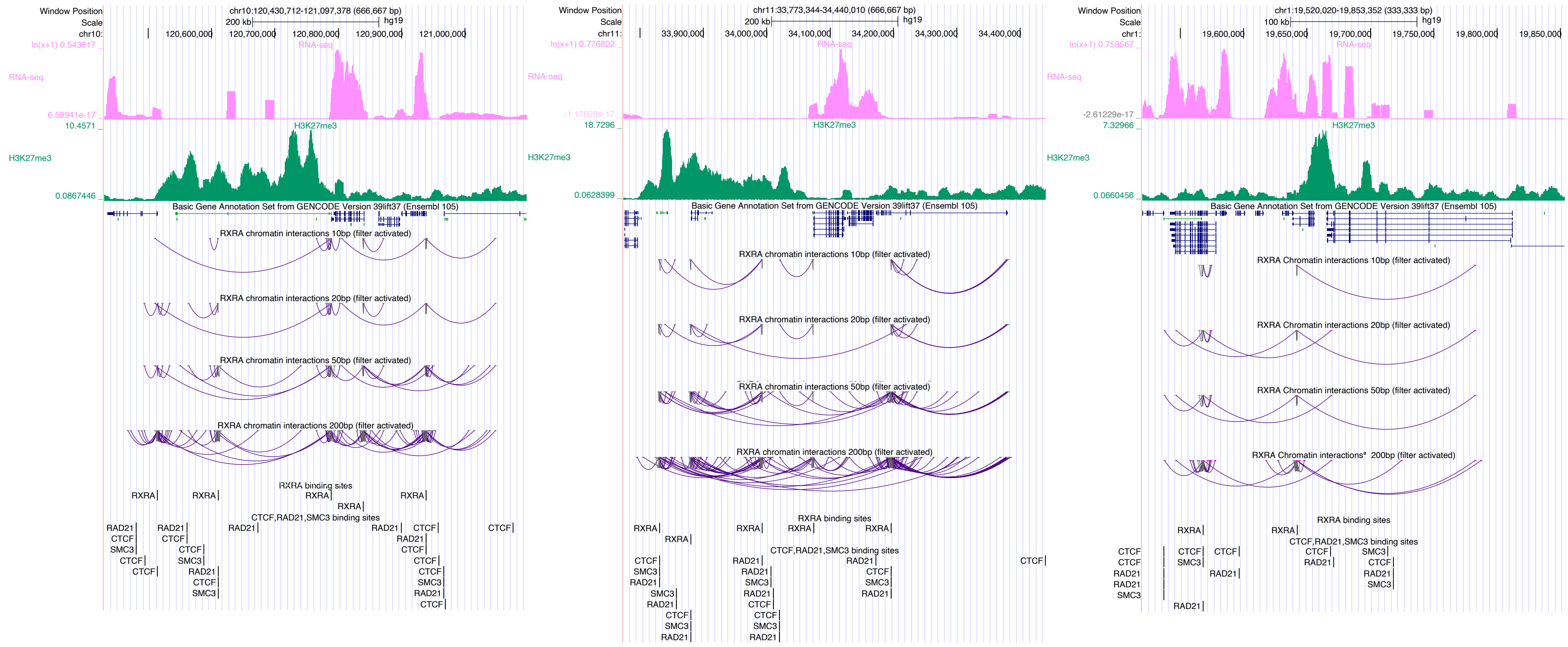
Window Position
chr10:120,430,712-121,097,378 (666,667 bp)
Scale
200 kb
hg19
RNA-seq
H3K27me3
Basic Gene Annotation Set from GENCODE Version 39lift37 (Ensembl 105)
RXRA chromatin interactions 10bp (filter activated)
RXRA chromatin interactions 20bp (filter activated)
RXRA chromatin interactions 50bp (filter activated)
RXRA chromatin interactions 200bp (filter activated)
RXRA binding sites
CTCF,RAD21,SMC3 binding sites
Window Position
chr11:33,773,344-34,440,010 (666,667 bp)
Window Position
chr1:19,520,020-19,853,352 (333,333 bp)
100 kb
RXRA Chromatin interactions 10bp (filter activated)
RXRA Chromatin interactions 20bp (filter activated)
RXRA Chromatin interactions 50bp (filter activated)

### SCRT1

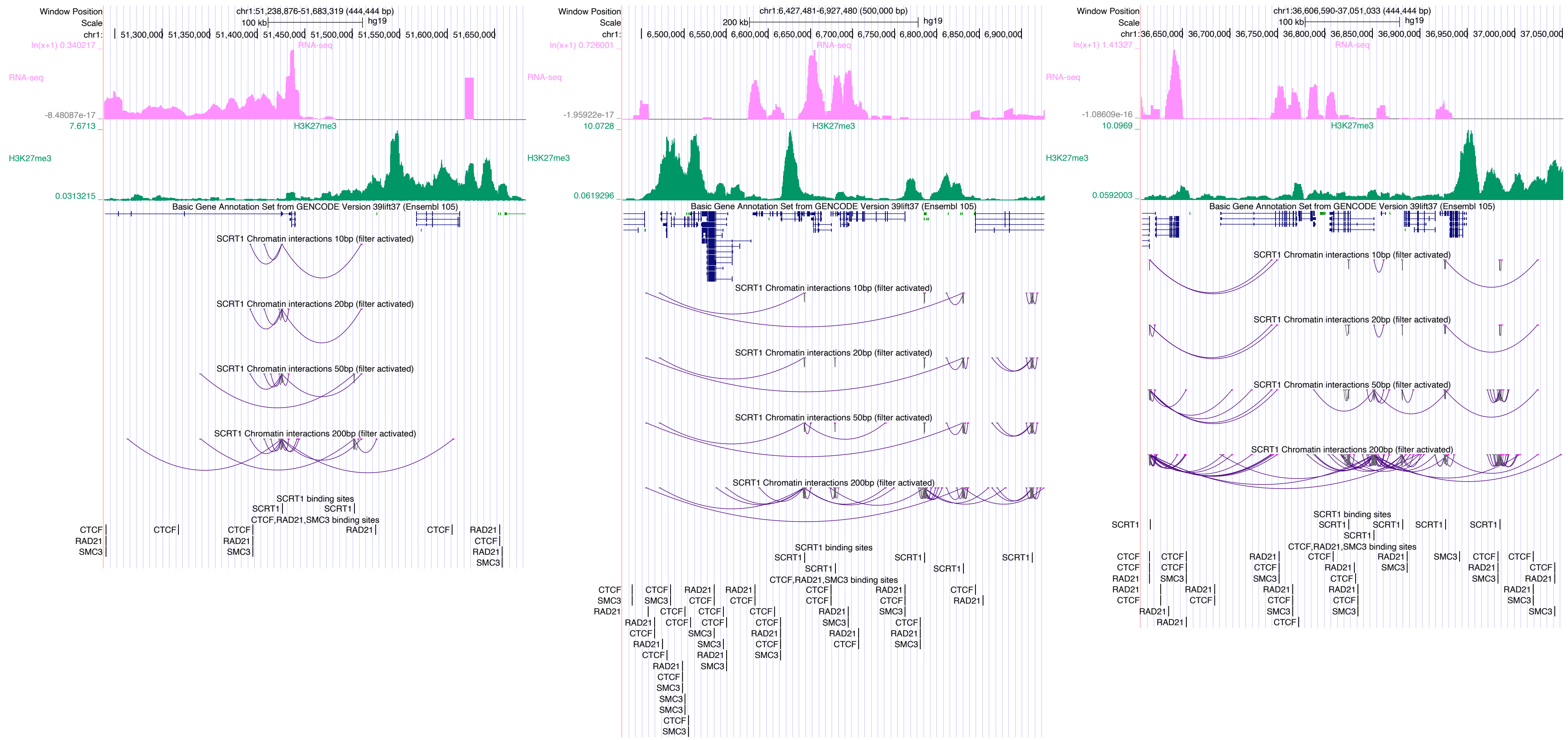
Window Position
chr1:51,238,876-51,683,319 (444,444 bp)
Scale
100 kb
hg19
chr1:
51,300,000
51,350,000
51,400,000
51,450,000
51,500,000
51,550,000
51,600,000
51,650,000
ln(x+1) 0.340217
RNA-seq
-8.48087e-17
7.6713
H3K27me3
0.0313215
Basic Gene Annotation Set from GENCODE Version 39lift37 (Ensembl 105)
SCRT1 Chromatin interactions 10bp (filter activated)
SCRT1 Chromatin interactions 20bp (filter activated)
SCRT1 Chromatin interactions 50bp (filter activated)
SCRT1 Chromatin interactions 200bp (filter activated)
SCRT1 binding sites
CTCF,RAD21,SMC3 binding sites
chr1:6,427,481-6,927,480 (500,000 bp)
200 kb
6,500,000
6,550,000
6,600,000
6,650,000
6,700,000
6,750,000
6,800,000
6,850,000
6,900,000
ln(x+1) 0.726001
-1.95922e-17
10.0728
0.0619296
chr1:36,606,590-37,051,033 (444,444 bp)
36,650,000
36,700,000
36,750,000
36,800,000
36,850,000
36,900,000
36,950,000
37,000,000
37,050,000
ln(x+1) 1.41327
-1.08609e-16
10.0969
0.0592003
SCRT1
CTCF
RAD21
SMC3

### SIN3A

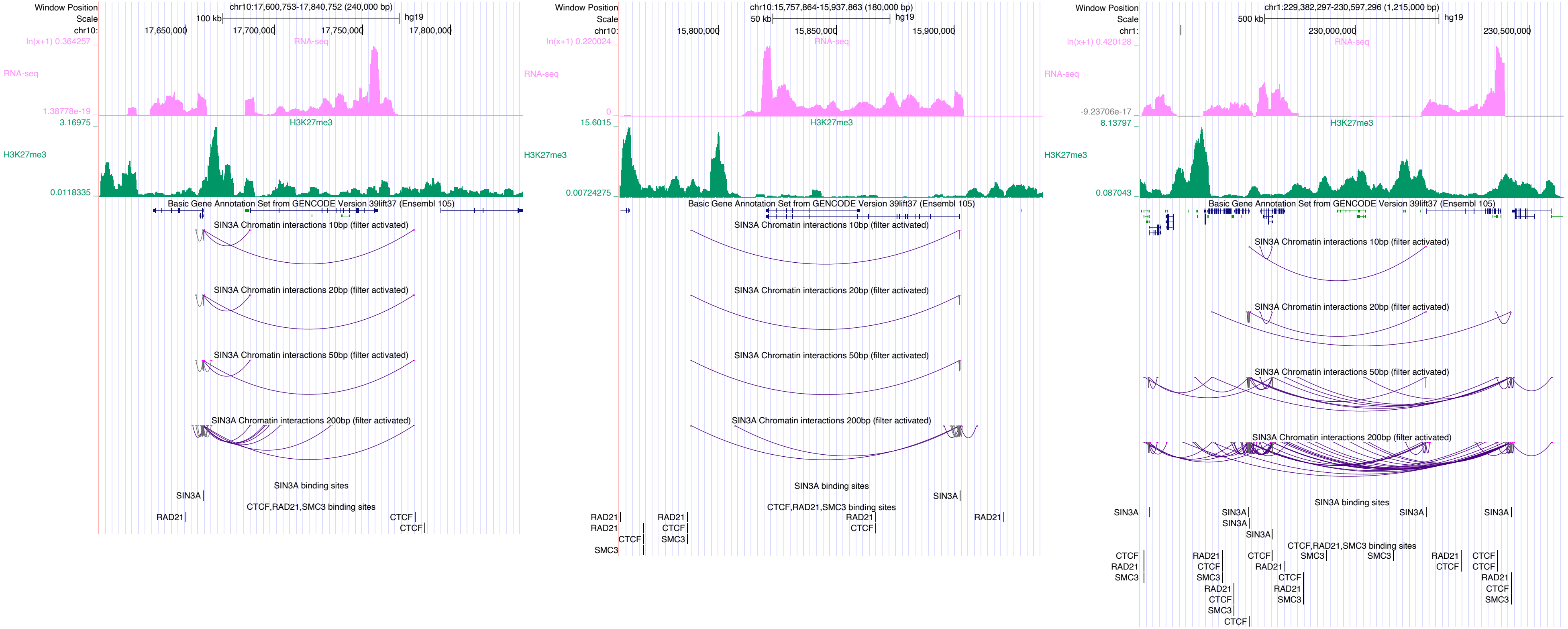

Window Position
Scale
chr10:
chr10:17,600,753-17,840,752 (240,000 bp)
100 kb
hg19
17,650,000
17,700,000
17,750,000
17,800,000
ln(x+1) 0.364257
RNA-seq
1.38778e-19
3.16975
H3K27me3
0.0118335
Basic Gene Annotation Set from GENCODE Version 39lift37 (Ensembl 105)
SIN3A Chromatin interactions 10bp (filter activated)
SIN3A Chromatin interactions 20bp (filter activated)
SIN3A Chromatin interactions 50bp (filter activated)
SIN3A Chromatin interactions 200bp (filter activated)
SIN3A binding sites
SIN3A
CTCF,RAD21,SMC3 binding sites
RAD21
CTCF
CTCF
Window Position
Scale
chr10:
chr10:15,757,864-15,937,863 (180,000 bp)
50 kb
hg19
15,800,000
15,850,000
15,900,000
ln(x+1) 0.220024
RNA-seq
0
15.6015
H3K27me3
0.00724275
Basic Gene Annotation Set from GENCODE Version 39lift37 (Ensembl 105)
SIN3A Chromatin interactions 10bp (filter activated)
SIN3A Chromatin interactions 20bp (filter activated)
SIN3A Chromatin interactions 50bp (filter activated)
SIN3A Chromatin interactions 200bp (filter activated)
SIN3A binding sites
SIN3A
CTCF,RAD21,SMC3 binding sites
RAD21
RAD21
CTCF
SMC3
RAD21
CTCF
SMC3
RAD21
CTCF
RAD21
Window Position
Scale
chr1:
chr1:229,382,297-230,597,296 (1,215,000 bp)
500 kb
hg19
230,000,000
230,500,000
ln(x+1) 0.420128
RNA-seq
-9.23706e-17
8.13797
H3K27me3
0.087043
Basic Gene Annotation Set from GENCODE Version 39lift37 (Ensembl 105)
SIN3A Chromatin interactions 10bp (filter activated)
SIN3A Chromatin interactions 20bp (filter activated)
SIN3A Chromatin interactions 50bp (filter activated)
SIN3A Chromatin interactions 200bp (filter activated)
SIN3A binding sites
SIN3A
SIN3A
SIN3A
SIN3A
SIN3A
SIN3A
CTCF,RAD21,SMC3 binding sites
CTCF
RAD21
SMC3
RAD21
CTCF
SMC3
RAD21
CTCF
SMC3
CTCF
CTCF
RAD21
CTCF
RAD21
SMC3
SMC3
SMC3
RAD21
CTCF
CTCF
CTCF
RAD21
CTCF
SMC3

### SMAD1

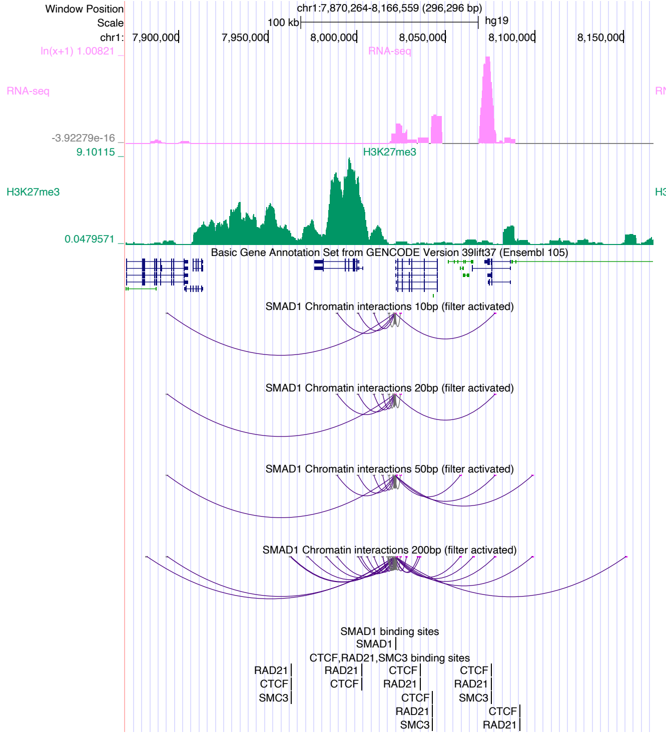

Window Position
chr1:7,870,264-8,166,559 (296,296 bp)
Scale
100 kb
hg19
RNA-seq
H3K27me3
Basic Gene Annotation Set from GENCODE Version 39lift37 (Ensembl 105)
SMAD1 Chromatin interactions 10bp (filter activated)
SMAD1 Chromatin interactions 20bp (filter activated)
SMAD1 Chromatin interactions 50bp (filter activated)
SMAD1 Chromatin interactions 200bp (filter activated)
SMAD1 binding sites
CTCF,RAD21,SMC3 binding sites


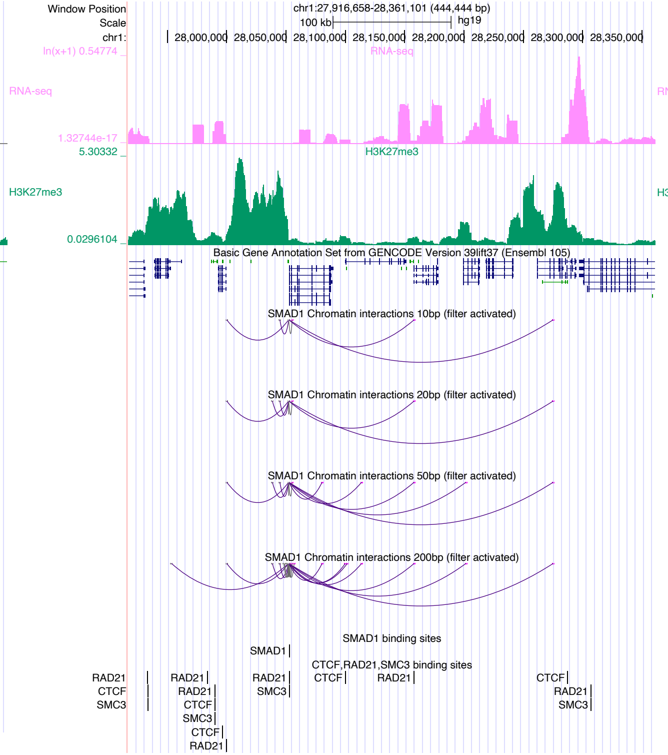

Window Position
chr1:27,916,658-28,361,101 (444,444 bp)
Scale
100 kb
hg19
RNA-seq
H3K27me3
Basic Gene Annotation Set from GENCODE Version 39lift37 (Ensembl 105)
SMAD1 Chromatin interactions 10bp (filter activated)
SMAD1 Chromatin interactions 20bp (filter activated)
SMAD1 Chromatin interactions 50bp (filter activated)
SMAD1 Chromatin interactions 200bp (filter activated)
SMAD1 binding sites
CTCF,RAD21,SMC3 binding sites


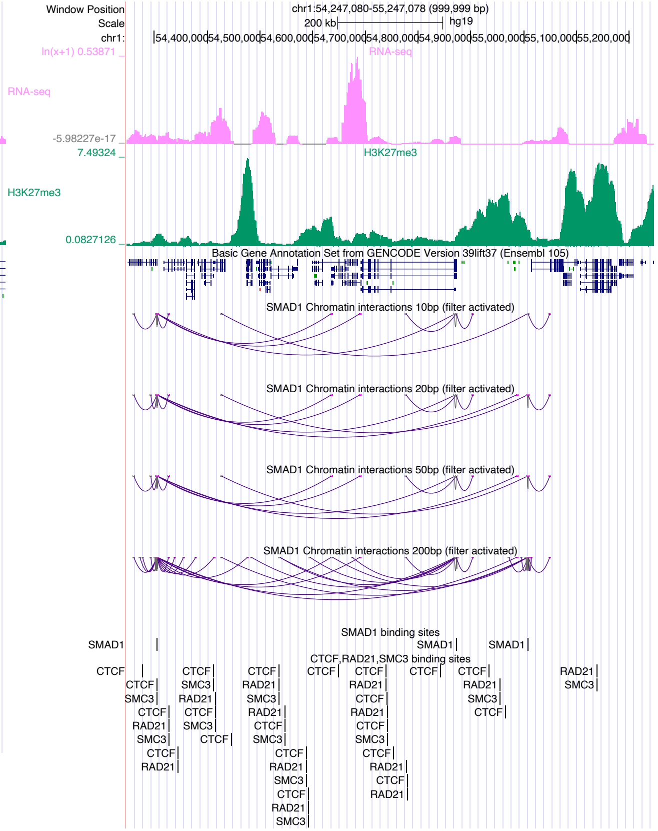

Window Position
chr1:54,247,080-55,247,078 (999,999 bp)
Scale
200 kb
hg19
RNA-seq
H3K27me3
Basic Gene Annotation Set from GENCODE Version 39lift37 (Ensembl 105)
SMAD1 Chromatin interactions 10bp (filter activated)
SMAD1 Chromatin interactions 20bp (filter activated)
SMAD1 Chromatin interactions 50bp (filter activated)
SMAD1 Chromatin interactions 200bp (filter activated)
SMAD1 binding sites
CTCF,RAD21,SMC3 binding sites

### SMAD2

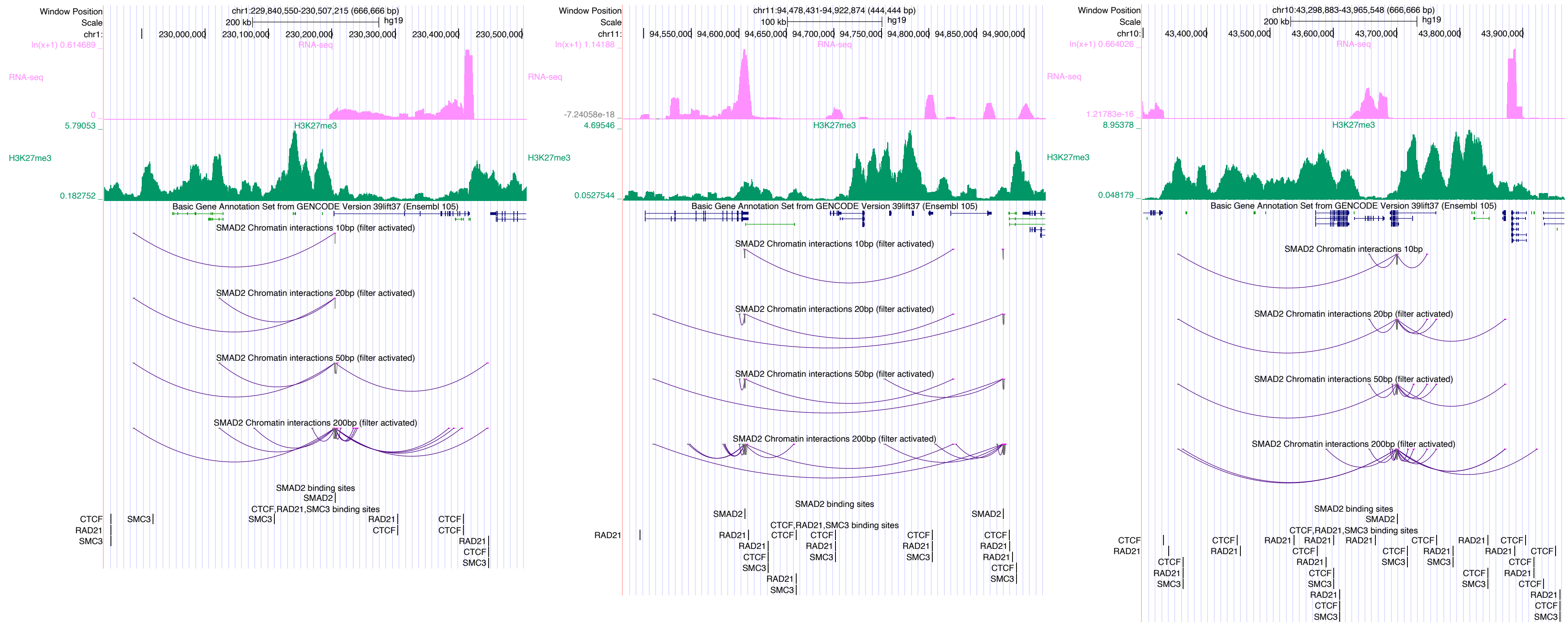

chr1:229,840,550-230,507,215 (666,666 bp)
chr11:94,478,431-94,922,874 (444,444 bp)
chr10:43,298,883-43,965,548 (666,666 bp)
RNA-seq
H3K27me3
Basic Gene Annotation Set from GENCODE Version 39lift37 (Ensembl 105)
SMAD2 Chromatin interactions 10bp (filter activated)
SMAD2 Chromatin interactions 20bp (filter activated)
SMAD2 Chromatin interactions 50bp (filter activated)
SMAD2 Chromatin interactions 200bp (filter activated)
SMAD2 binding sites
CTCF,RAD21,SMC3 binding sites

### SMAD3

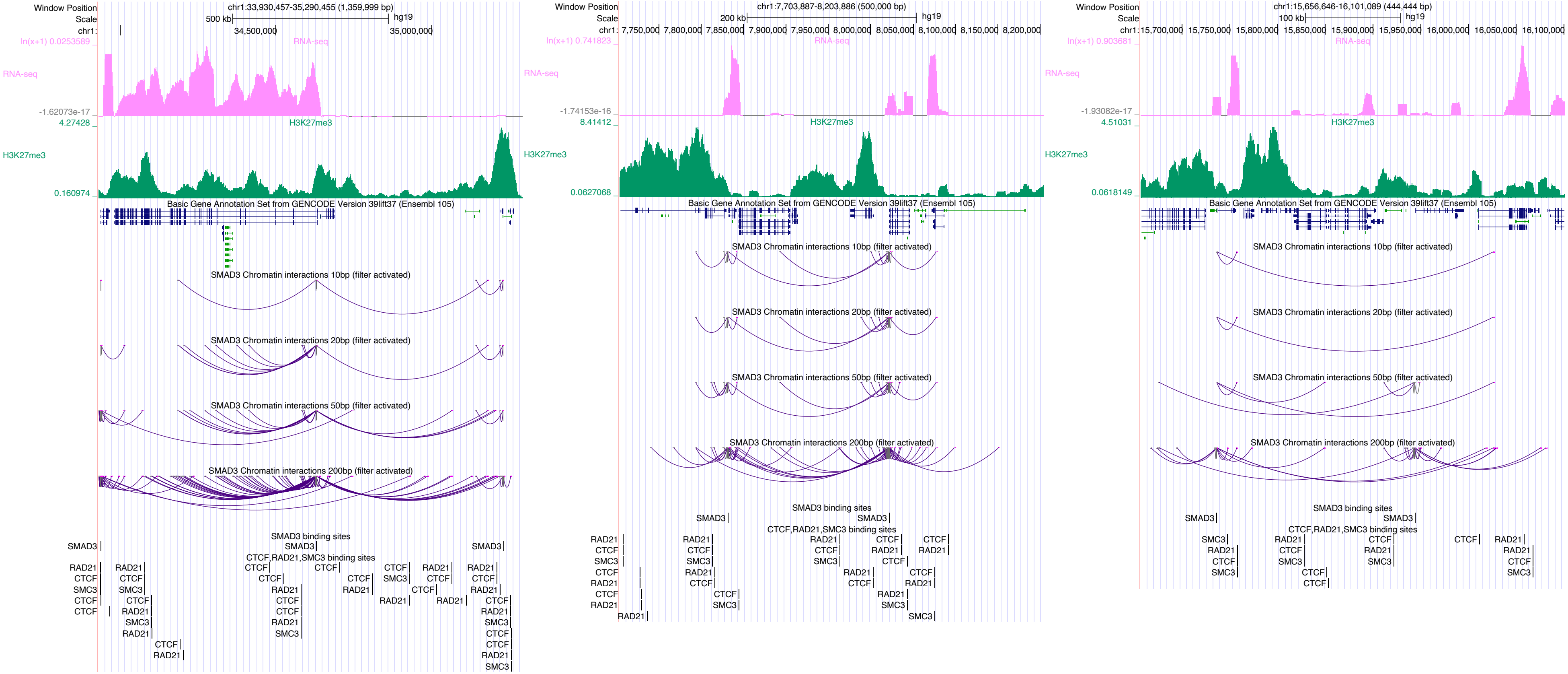

Window Position
Scale
chr1:33,930,457-35,290,455 (1,359,999 bp)
500 kb
hg19
34,500,000
35,000,000
RNA-seq
ln(x+1) 0.0253589
-1.62073e-17
H3K27me3
4.27428
0.160974
Basic Gene Annotation Set from GENCODE Version 39lift37 (Ensembl 105)
SMAD3 Chromatin interactions 10bp (filter activated)
SMAD3 Chromatin interactions 20bp (filter activated)
SMAD3 Chromatin interactions 50bp (filter activated)
SMAD3 Chromatin interactions 200bp (filter activated)
SMAD3 binding sites
CTCF,RAD21,SMC3 binding sites
chr1:7,703,887-8,203,886 (500,000 bp)
200 kb
7,750,000
7,800,000
7,850,000
7,900,000
7,950,000
8,000,000
8,050,000
8,100,000
8,150,000
8,200,000
ln(x+1) 0.741823
-1.74153e-16
8.41412
0.0627068
chr1:15,656,646-16,101,089 (444,444 bp)
100 kb
15,700,000
15,750,000
15,800,000
15,850,000
15,900,000
15,950,000
16,000,000
16,050,000
16,100,000
ln(x+1) 0.903681
-1.93082e-17
4.51031
0.0618149
SMAD3
CTCF
RAD21
SMC3

# SP1

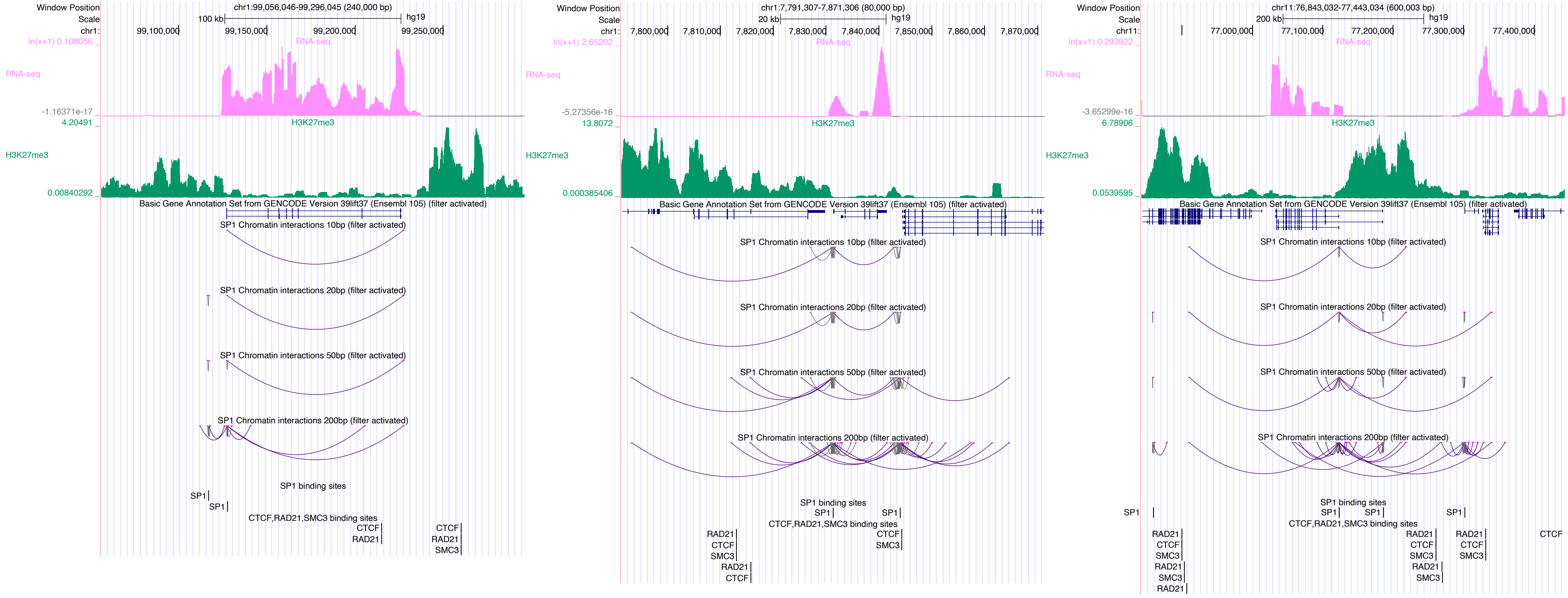

Window Position
Scale
chr1:99,056,046-99,296,045 (240,000 bp)
100 kb
hg19
99,100,000
99,150,000
99,200,000
99,250,000
RNA-seq
H3K27me3
Basic Gene Annotation Set from GENCODE Version 39lift37 (Ensembl 105) (filter activated)
SP1 Chromatin interactions 10bp (filter activated)
SP1 Chromatin interactions 20bp (filter activated)
SP1 Chromatin interactions 50bp (filter activated)
SP1 Chromatin interactions 200bp (filter activated)
SP1 binding sites
CTCF,RAD21,SMC3 binding sites
chr1:7,791,307-7,871,306 (80,000 bp)
20 kb
7,800,000
7,810,000
7,820,000
7,830,000
7,840,000
7,850,000
7,860,000
7,870,000
chr11:76,843,032-77,443,034 (600,003 bp)
200 kb
77,000,000
77,100,000
77,200,000
77,300,000
77,400,000

# SP2

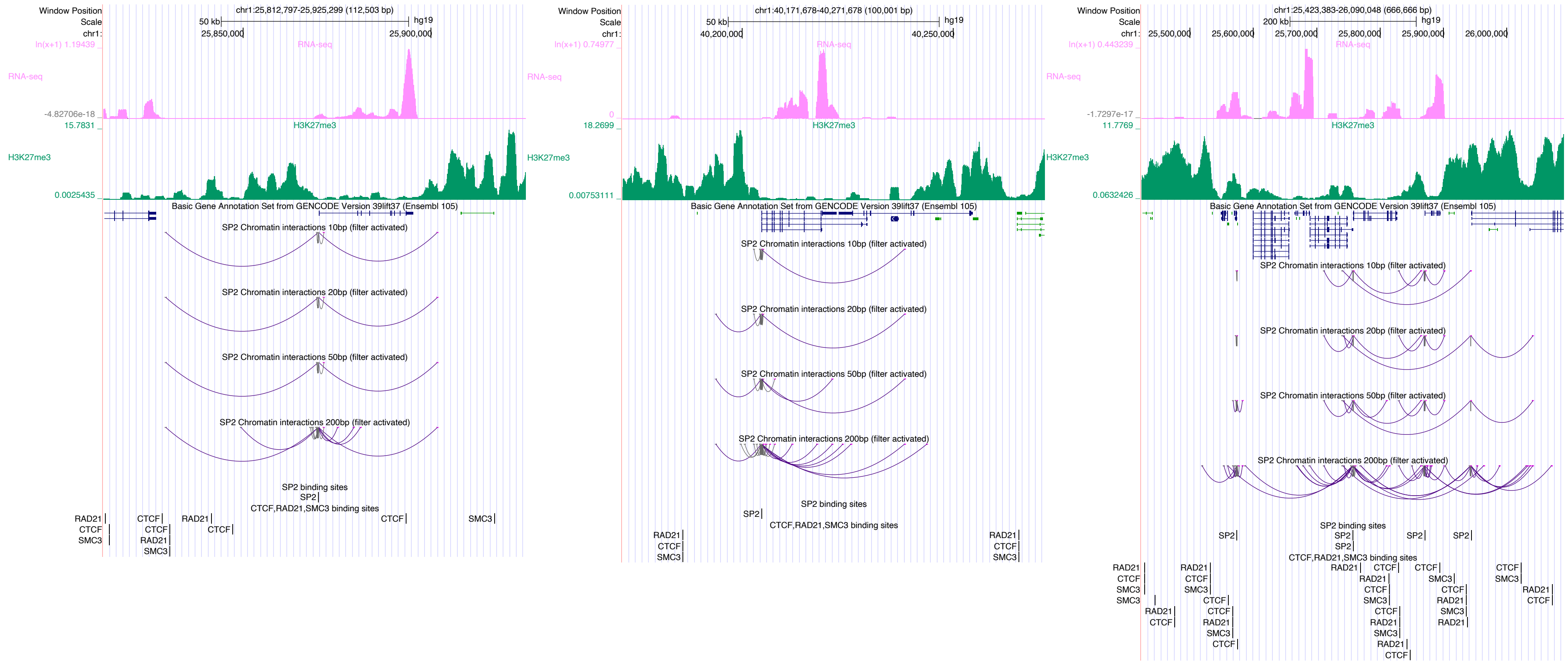
Window Position
Scale
chr1:
chr1:25,812,797-25,925,299 (112,503 bp)
50 kb
hg19
25,850,000
25,900,000
ln(x+1) 1.19439
RNA-seq
-4.82706e-18
15.7831
H3K27me3
0.0025435
Basic Gene Annotation Set from GENCODE Version 39lift37 (Ensembl 105)
SP2 Chromatin interactions 10bp (filter activated)
SP2 Chromatin interactions 20bp (filter activated)
SP2 Chromatin interactions 50bp (filter activated)
SP2 Chromatin interactions 200bp (filter activated)
SP2 binding sites
SP2
CTCF,RAD21,SMC3 binding sites
RAD21
CTCF
SMC3
chr1:40,171,678-40,271,678 (100,001 bp)
40,200,000
40,250,000
ln(x+1) 0.74977
0
18.2699
0.00753111
chr1:25,423,383-26,090,048 (666,666 bp)
200 kb
25,500,000
25,600,000
25,700,000
25,800,000
25,900,000
26,000,000
ln(x+1) 0.443239
-1.7297e-17
11.7769
0.0632426

# SP5

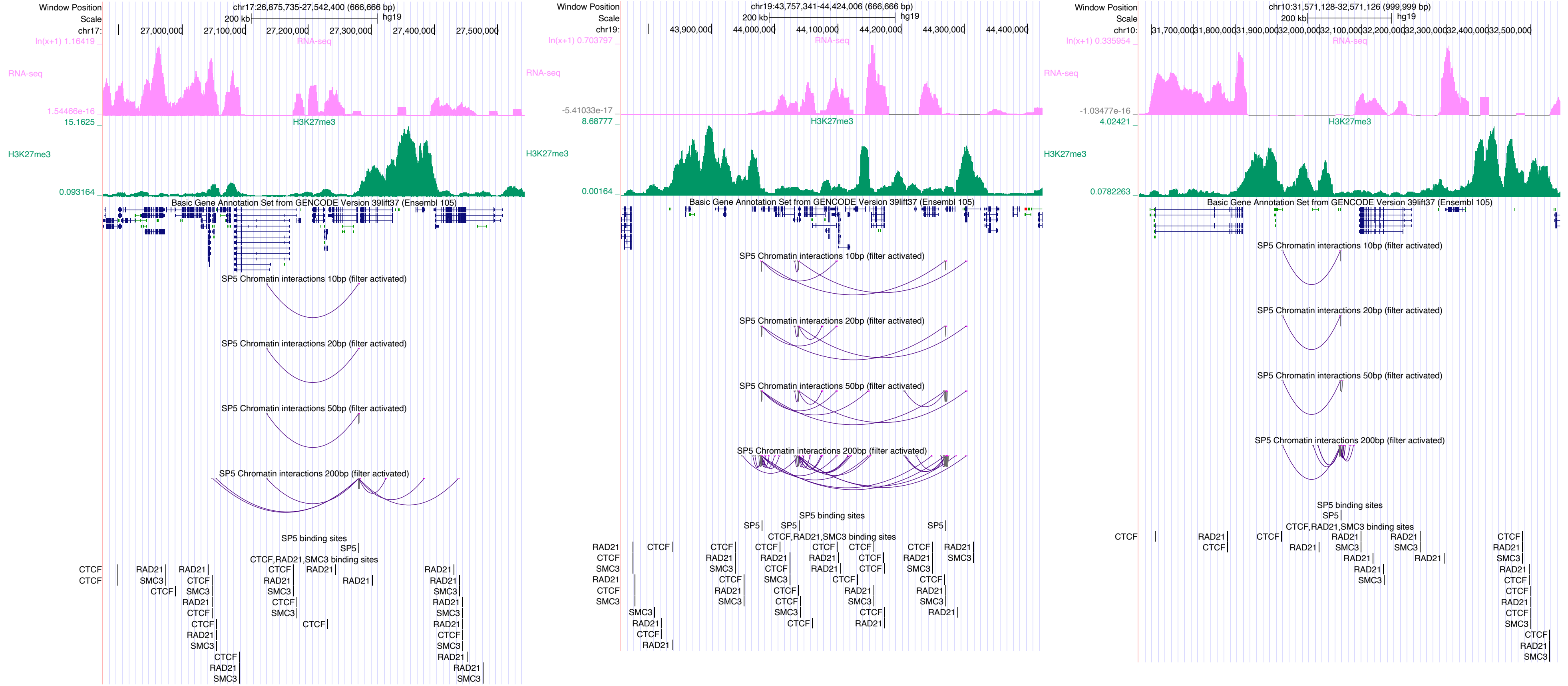
Window Position
Scale
chr17:26,875,735-27,542,400 (666,666 bp)
200 kb
hg19
chr17:
27,000,000
27,100,000
27,200,000
27,300,000
27,400,000
27,500,000
ln(x+1) 1.16419
RNA-seq
1.54466e-16
15.1625
H3K27me3
0.093164
Basic Gene Annotation Set from GENCODE Version 39lift37 (Ensembl 105)
SP5 Chromatin interactions 10bp (filter activated)
SP5 Chromatin interactions 20bp (filter activated)
SP5 Chromatin interactions 50bp (filter activated)
SP5 Chromatin interactions 200bp (filter activated)
SP5 binding sites
SP5
CTCF,RAD21,SMC3 binding sites
chr19:43,757,341-44,424,006 (666,666 bp)
chr19:
43,900,000
44,000,000
44,100,000
44,200,000
44,300,000
44,400,000
ln(x+1) 0.703797
-5.41033e-17
8.68777
0.00164
chr10:31,571,128-32,571,126 (999,999 bp)
chr10:
31,700,000
31,800,000
31,900,000
32,000,000
32,100,000
32,200,000
32,300,000
32,400,000
32,500,000
ln(x+1) 0.335954
-1.03477e-16
4.02421
0.0782263
CTCF
RAD21
SMC3

### SPI1

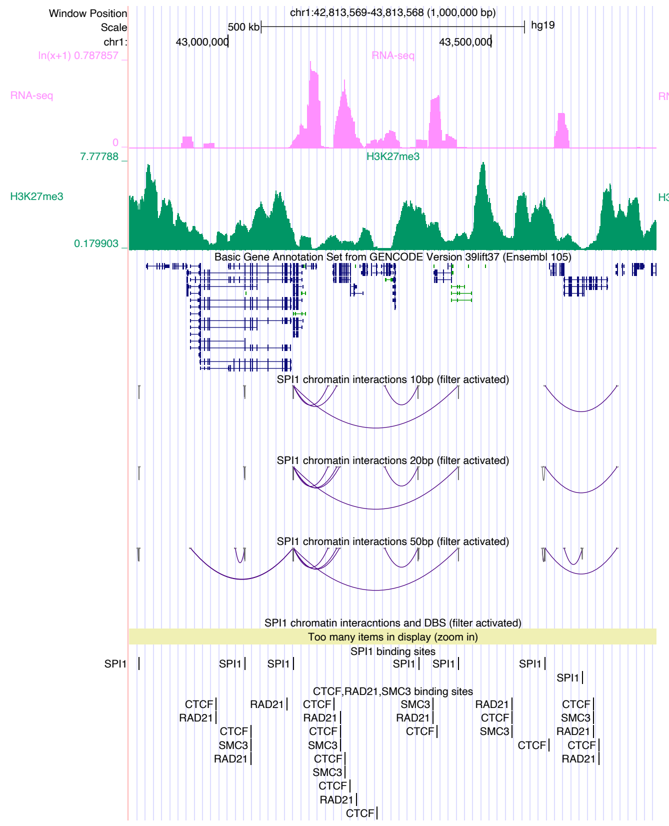

Window Position
Scale
chr1:
chr1:42,813,569-43,813,568 (1,000,000 bp)
500 kb
hg19
43,000,000
43,500,000
RNA-seq
H3K27me3
Basic Gene Annotation Set from GENCODE Version 39lift37 (Ensembl 105)
SPI1 chromatin interactions 10bp (filter activated)
SPI1 chromatin interactions 20bp (filter activated)
SPI1 chromatin interactions 50bp (filter activated)
SPI1 chromatin interacntions and DBS (filter activated)
Too many items in display (zoom in)
SPI1 binding sites
CTCF,RAD21,SMC3 binding sites


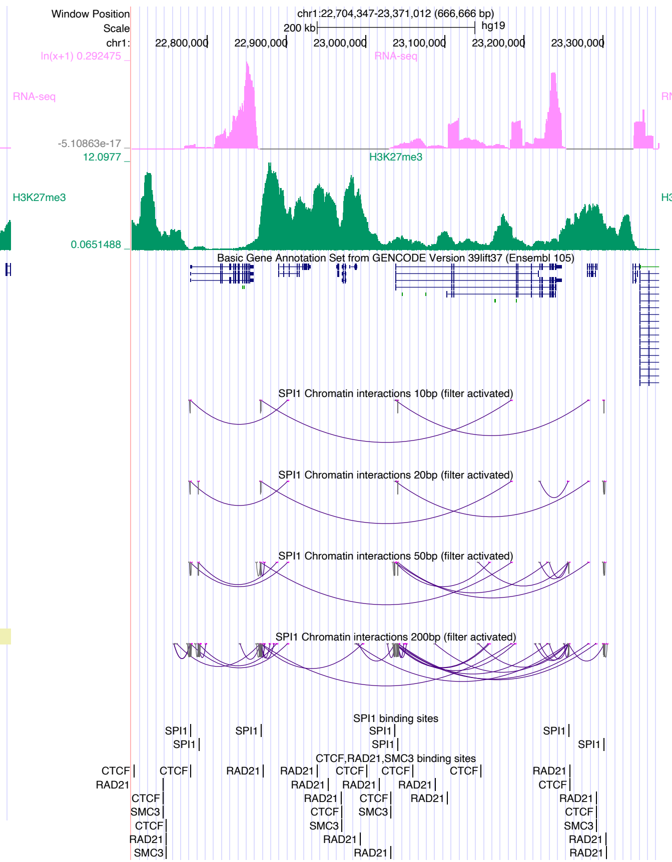

Window Position
Scale
chr1:
chr1:22,704,347-23,371,012 (666,666 bp)
200 kb
hg19
22,800,000
22,900,000
23,000,000
23,100,000
23,200,000
23,300,000
RNA-seq
H3K27me3
Basic Gene Annotation Set from GENCODE Version 39lift37 (Ensembl 105)
SPI1 Chromatin interactions 10bp (filter activated)
SPI1 Chromatin interactions 20bp (filter activated)
SPI1 Chromatin interactions 50bp (filter activated)
SPI1 Chromatin interactions 200bp (filter activated)
SPI1 binding sites
CTCF,RAD21,SMC3 binding sites


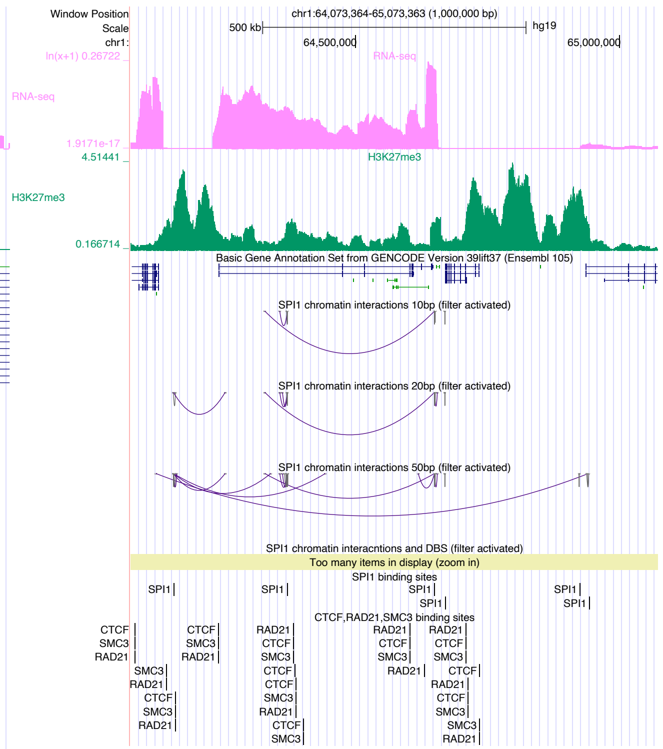

Window Position
Scale
chr1:
chr1:64,073,364-65,073,363 (1,000,000 bp)
500 kb
hg19
64,500,000
65,000,000
RNA-seq
H3K27me3
Basic Gene Annotation Set from GENCODE Version 39lift37 (Ensembl 105)
SPI1 chromatin interactions 10bp (filter activated)
SPI1 chromatin interactions 20bp (filter activated)
SPI1 chromatin interactions 50bp (filter activated)
SPI1 chromatin interacntions and DBS (filter activated)
Too many items in display (zoom in)
SPI1 binding sites
CTCF,RAD21,SMC3 binding sites

### SPIB

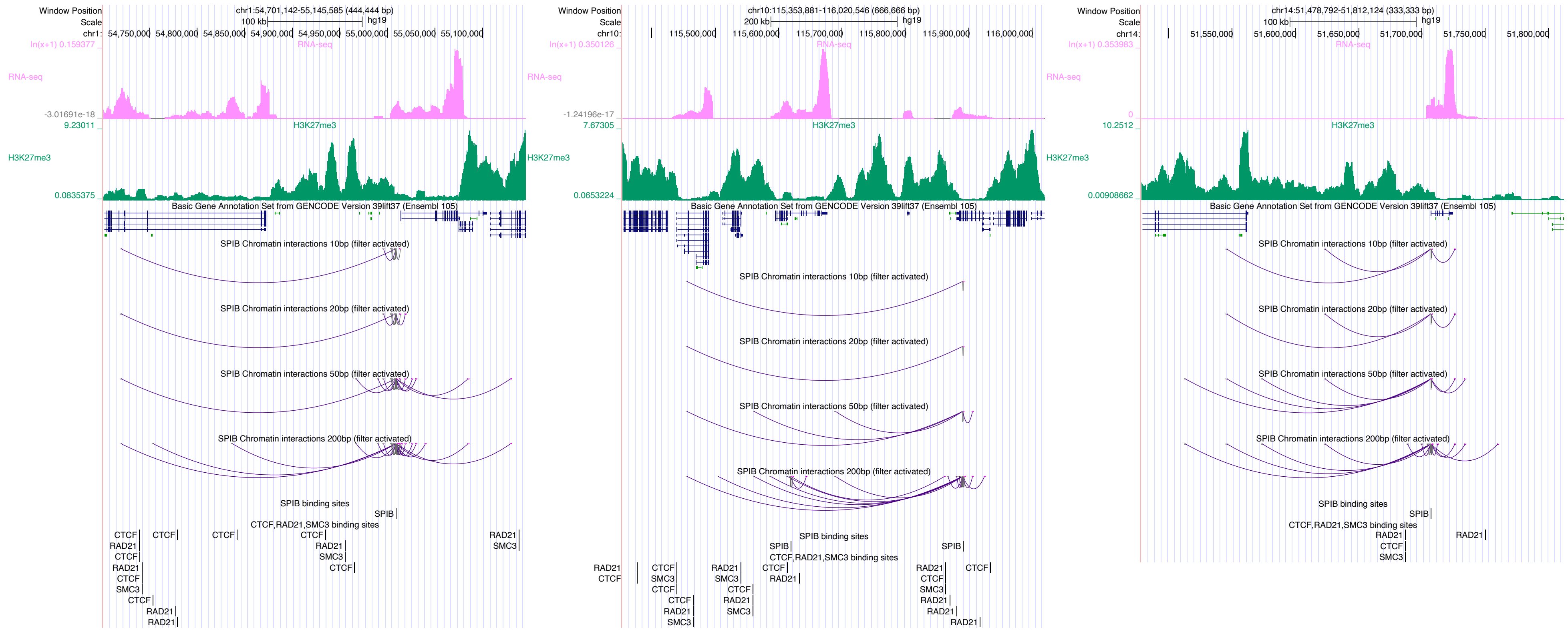

Window Position
chr1:54,701,142-55,145,585 (444,444 bp)
Scale
100 kb
hg19
RNA-seq
H3K27me3
Basic Gene Annotation Set from GENCODE Version 39lift37 (Ensembl 105)
SPIB Chromatin interactions 10bp (filter activated)
SPIB Chromatin interactions 20bp (filter activated)
SPIB Chromatin interactions 50bp (filter activated)
SPIB Chromatin interactions 200bp (filter activated)
SPIB binding sites
CTCF,RAD21,SMC3 binding sites
Window Position
chr10:115,353,881-116,020,546 (666,666 bp)
200 kb
Window Position
chr14:51,478,792-51,812,124 (333,333 bp)
100 kb

### SREBF2

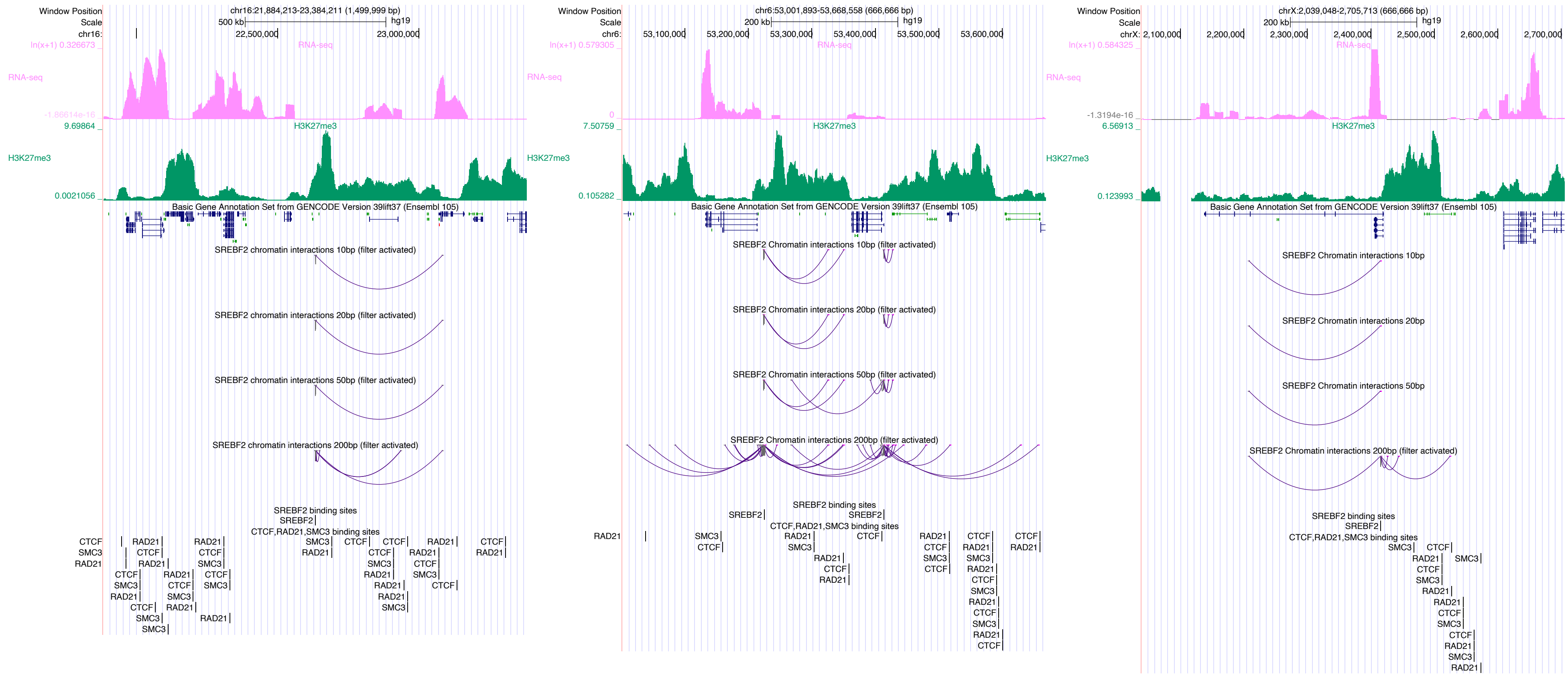
Window Position
chr16:21,884,213-23,384,211 (1,499,999 bp)
Scale
500 kb
hg19
22,500,000
23,000,000
RNA-seq
H3K27me3
Basic Gene Annotation Set from GENCODE Version 39lift37 (Ensembl 105)
SREBF2 chromatin interactions 10bp (filter activated)
SREBF2 chromatin interactions 20bp (filter activated)
SREBF2 chromatin interactions 50bp (filter activated)
SREBF2 chromatin interactions 200bp (filter activated)
SREBF2 binding sites
CTCF,RAD21,SMC3 binding sites
Window Position
chr6:53,001,893-53,668,558 (666,666 bp)
200 kb
53,100,000
53,200,000
53,300,000
53,400,000
53,500,000
53,600,000
SREBF2 Chromatin interactions 10bp (filter activated)
SREBF2 Chromatin interactions 20bp (filter activated)
SREBF2 Chromatin interactions 50bp (filter activated)
SREBF2 Chromatin interactions 200bp (filter activated)
Window Position
chrX:2,039,048-2,705,713 (666,666 bp)
2,100,000
2,200,000
2,300,000
2,400,000
2,500,000
2,600,000
2,700,000
SREBF2 Chromatin interactions 10bp
SREBF2 Chromatin interactions 20bp
SREBF2 Chromatin interactions 50bp
SREBF2 Chromatin interactions 200bp (filter activated)

### STAT1

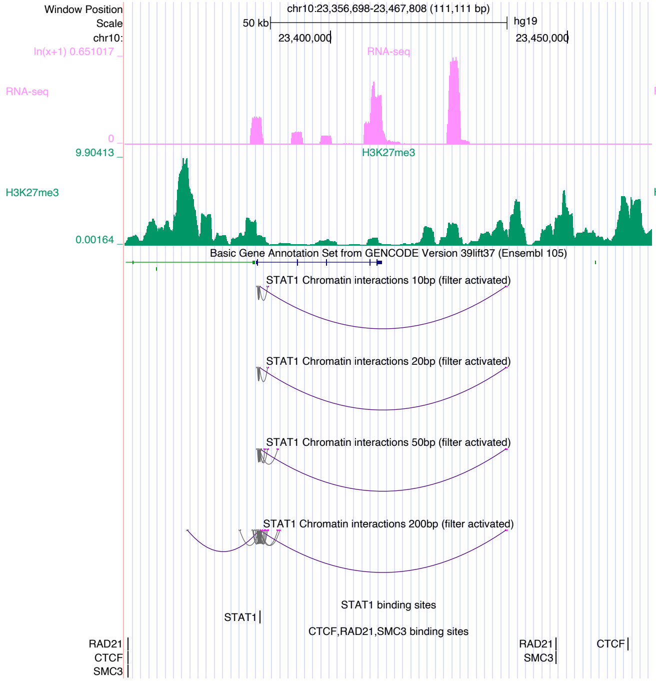
Window Position
Scale
chr10:
chr10:23,356,698-23,467,808 (111,111 bp)
50 kb
hg19
23,400,000
23,450,000
ln(x+1) 0.651017
RNA-seq
0
9.90413
H3K27me3
0.00164
Basic Gene Annotation Set from GENCODE Version 39lift37 (Ensembl 105)
STAT1 Chromatin interactions 10bp (filter activated)
STAT1 Chromatin interactions 20bp (filter activated)
STAT1 Chromatin interactions 50bp (filter activated)
STAT1 Chromatin interactions 200bp (filter activated)
STAT1 binding sites
STAT1
CTCF,RAD21,SMC3 binding sites
RAD21
CTCF
SMC3

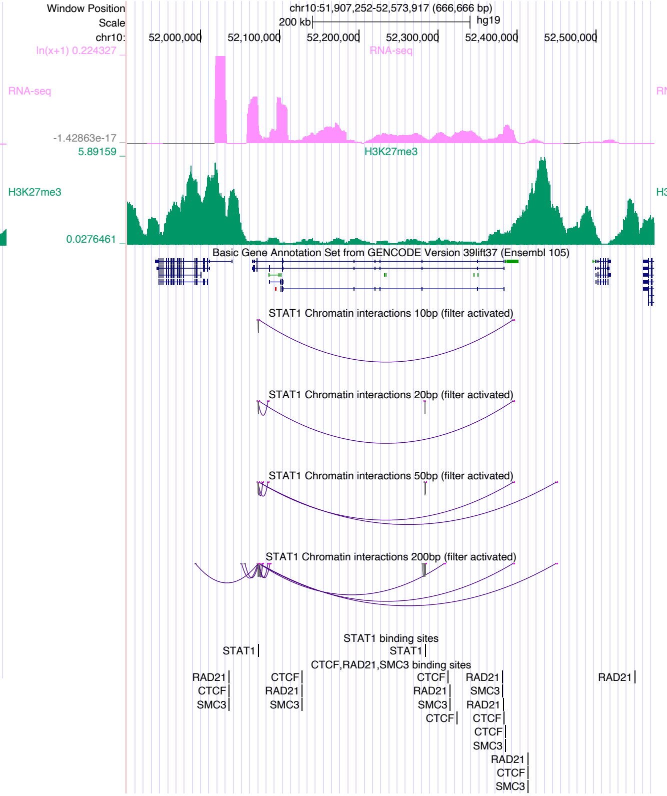
Window Position
Scale
chr10:
chr10:51,907,252-52,573,917 (666,666 bp)
200 kb
hg19
52,000,000
52,100,000
52,200,000
52,300,000
52,400,000
52,500,000
ln(x+1) 0.224327
RNA-seq
-1.42863e-17
5.89159
H3K27me3
0.0276461
Basic Gene Annotation Set from GENCODE Version 39lift37 (Ensembl 105)
STAT1 Chromatin interactions 10bp (filter activated)
STAT1 Chromatin interactions 20bp (filter activated)
STAT1 Chromatin interactions 50bp (filter activated)
STAT1 Chromatin interactions 200bp (filter activated)
STAT1 binding sites
STAT1
CTCF,RAD21,SMC3 binding sites
RAD21
CTCF
SMC3

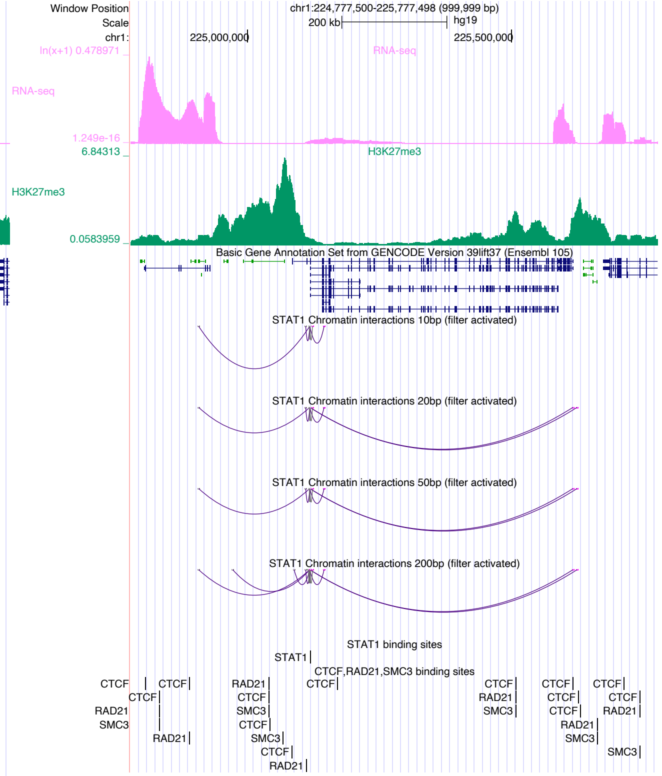
Window Position
Scale
chr1:
chr1:224,777,500-225,777,498 (999,999 bp)
200 kb
hg19
225,000,000
225,500,000
ln(x+1) 0.478971
RNA-seq
1.249e-16
6.84313
H3K27me3
0.0583959
Basic Gene Annotation Set from GENCODE Version 39lift37 (Ensembl 105)
STAT1 Chromatin interactions 10bp (filter activated)
STAT1 Chromatin interactions 20bp (filter activated)
STAT1 Chromatin interactions 50bp (filter activated)
STAT1 Chromatin interactions 200bp (filter activated)
STAT1 binding sites
STAT1
CTCF,RAD21,SMC3 binding sites
CTCF
RAD21
SMC3

### STAT3

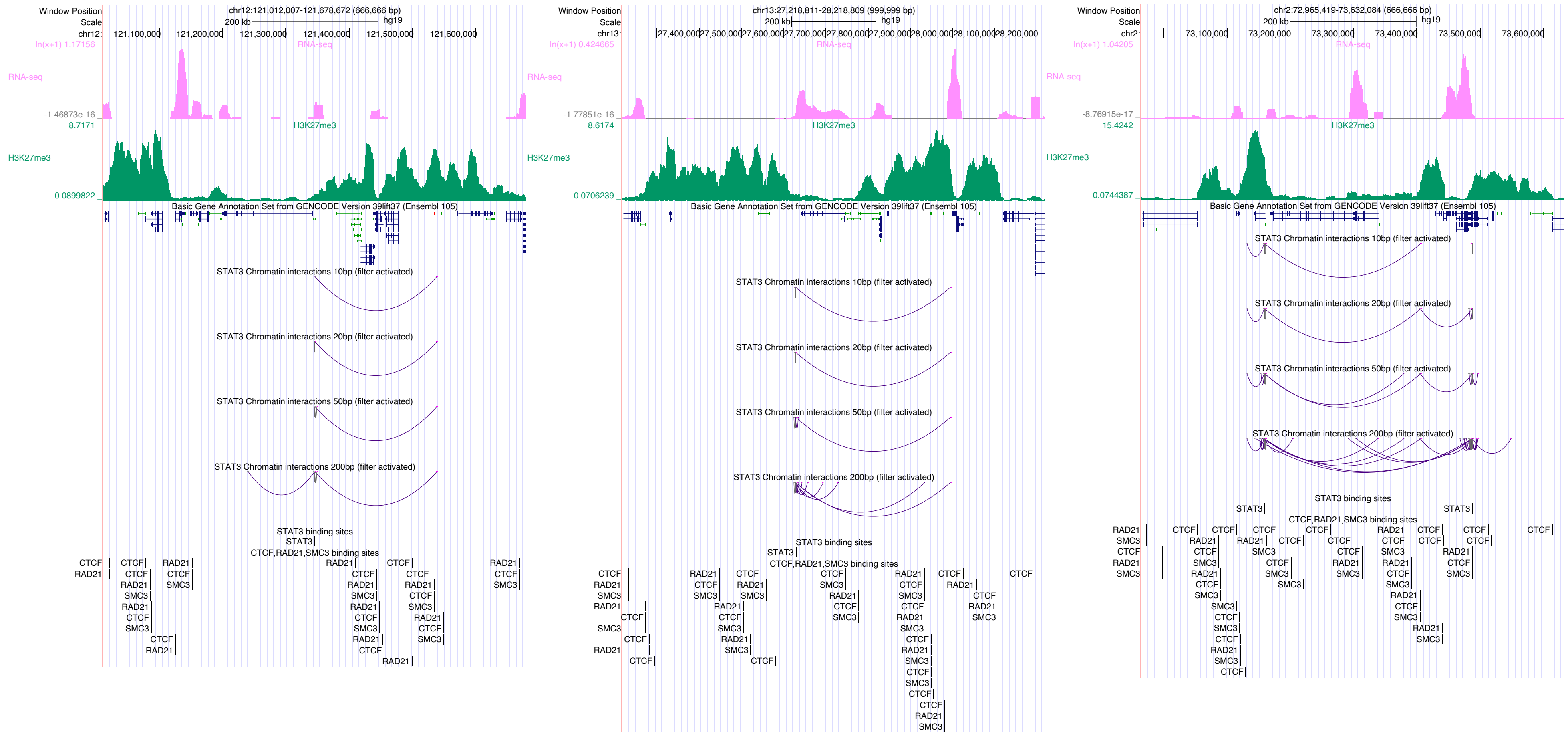
Window Position
chr12:121,012,007-121,678,672 (666,666 bp)
RNA-seq
H3K27me3
Basic Gene Annotation Set from GENCODE Version 39lift37 (Ensembl 105)
STAT3 Chromatin interactions 10bp (filter activated)
STAT3 Chromatin interactions 20bp (filter activated)
STAT3 Chromatin interactions 50bp (filter activated)
STAT3 Chromatin interactions 200bp (filter activated)
STAT3 binding sites
CTCF,RAD21,SMC3 binding sites
chr13:27,218,811-28,218,809 (999,999 bp)
chr2:72,965,419-73,632,084 (666,666 bp)

### TAF1

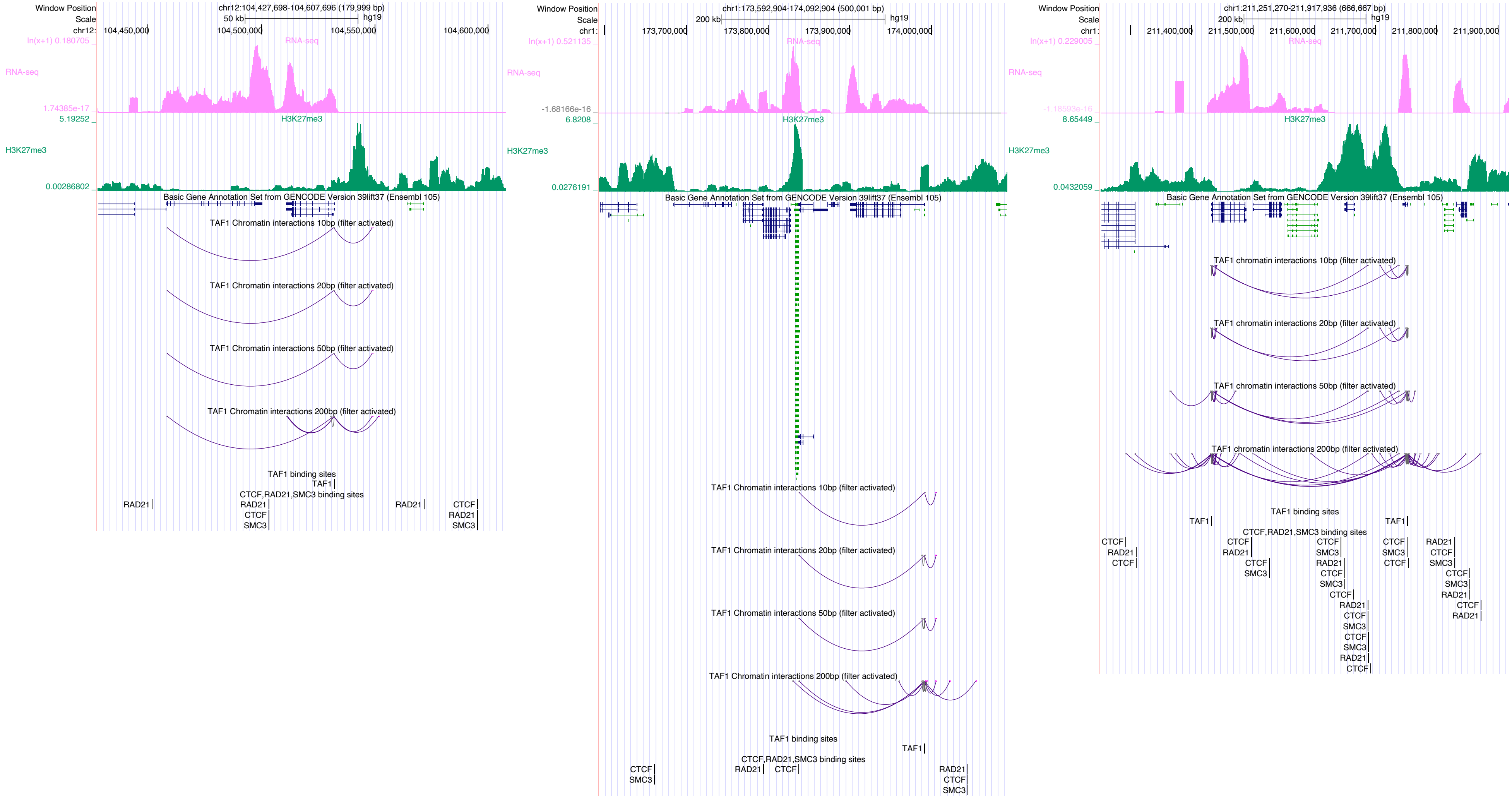

Window Position
Scale
chr12:
chr12:104,427,698-104,607,696 (179,999 bp)
50 kb
hg19
104,450,000
104,500,000
104,550,000
104,600,000
ln(x+1) 0.180705
RNA-seq
1.74385e-17
5.19252
H3K27me3
0.00286802
Basic Gene Annotation Set from GENCODE Version 39lift37 (Ensembl 105)
TAF1 Chromatin interactions 10bp (filter activated)
TAF1 Chromatin interactions 20bp (filter activated)
TAF1 Chromatin interactions 50bp (filter activated)
TAF1 Chromatin interactions 200bp (filter activated)
TAF1 binding sites
TAF1
CTCF,RAD21,SMC3 binding sites
RAD21
CTCF
SMC3
Window Position
Scale
chr1:
chr1:173,592,904-174,092,904 (500,001 bp)
200 kb
hg19
173,700,000
173,800,000
173,900,000
174,000,000
ln(x+1) 0.521135
RNA-seq
-1.68166e-16
6.8208
H3K27me3
0.0276191
Basic Gene Annotation Set from GENCODE Version 39lift37 (Ensembl 105)
TAF1 Chromatin interactions 10bp (filter activated)
TAF1 Chromatin interactions 20bp (filter activated)
TAF1 Chromatin interactions 50bp (filter activated)
TAF1 Chromatin interactions 200bp (filter activated)
TAF1 binding sites
TAF1
CTCF,RAD21,SMC3 binding sites
CTCF
SMC3
RAD21
Window Position
Scale
chr1:
chr1:211,251,270-211,917,936 (666,667 bp)
200 kb
hg19
211,400,000
211,500,000
211,600,000
211,700,000
211,800,000
211,900,000
ln(x+1) 0.229005
RNA-seq
-1.18593e-16
8.65449
H3K27me3
0.0432059
Basic Gene Annotation Set from GENCODE Version 39lift37 (Ensembl 105)
TAF1 chromatin interactions 10bp (filter activated)
TAF1 chromatin interactions 20bp (filter activated)
TAF1 chromatin interactions 50bp (filter activated)
TAF1 chromatin interactions 200bp (filter activated)
TAF1 binding sites
TAF1
CTCF,RAD21,SMC3 binding sites
CTCF
RAD21
SMC3

### TBX21

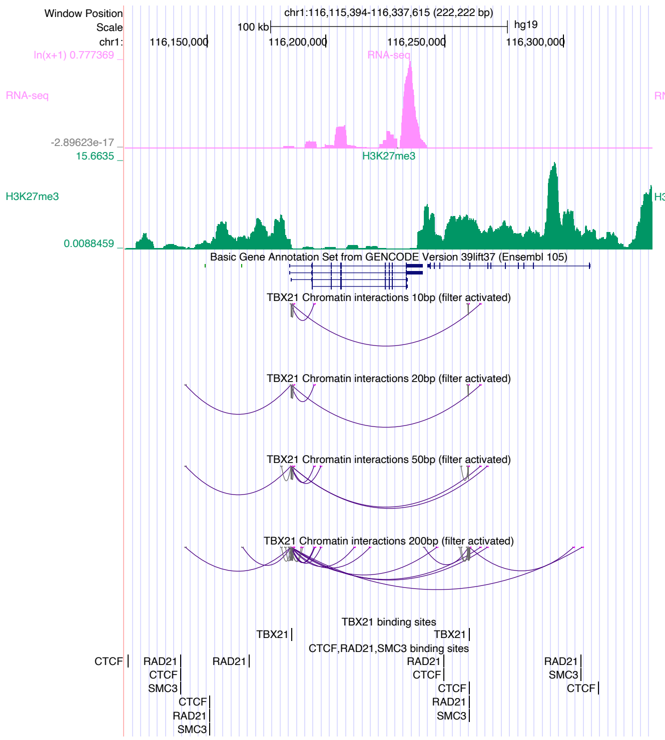

Window Position chr1:116,115,394-116,337,615 (222,222 bp)
Scale 100 kb hg19
RNA-seq
H3K27me3
Basic Gene Annotation Set from GENCODE Version 39lift37 (Ensembl 105)
TBX21 Chromatin interactions 10bp (filter activated)
TBX21 Chromatin interactions 20bp (filter activated)
TBX21 Chromatin interactions 50bp (filter activated)
TBX21 Chromatin interactions 200bp (filter activated)
TBX21 binding sites
CTCF,RAD21,SMC3 binding sites


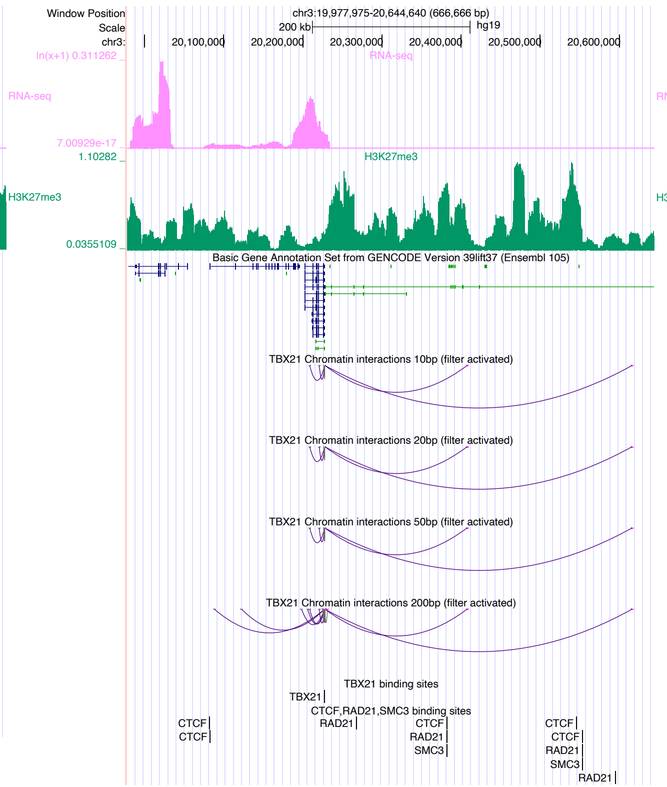

Window Position chr3:19,977,975-20,644,640 (666,666 bp)
Scale 200 kb hg19
RNA-seq
H3K27me3
Basic Gene Annotation Set from GENCODE Version 39lift37 (Ensembl 105)
TBX21 Chromatin interactions 10bp (filter activated)
TBX21 Chromatin interactions 20bp (filter activated)
TBX21 Chromatin interactions 50bp (filter activated)
TBX21 Chromatin interactions 200bp (filter activated)
TBX21 binding sites
CTCF,RAD21,SMC3 binding sites


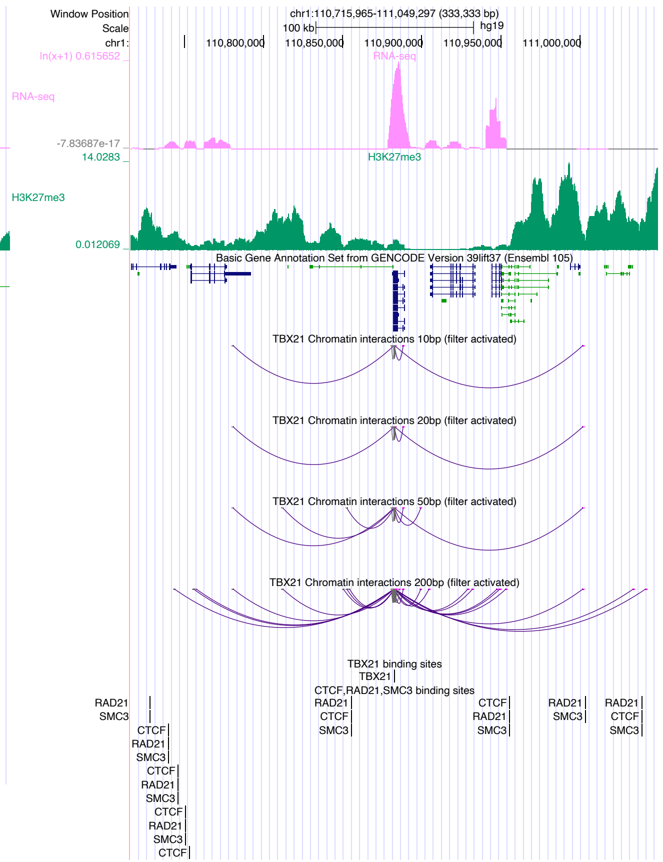

Window Position chr1:110,715,965-111,049,297 (333,333 bp)
Scale 100 kb hg19
RNA-seq
H3K27me3
Basic Gene Annotation Set from GENCODE Version 39lift37 (Ensembl 105)
TBX21 Chromatin interactions 10bp (filter activated)
TBX21 Chromatin interactions 20bp (filter activated)
TBX21 Chromatin interactions 50bp (filter activated)
TBX21 Chromatin interactions 200bp (filter activated)
TBX21 binding sites
CTCF,RAD21,SMC3 binding sites

### TCF3

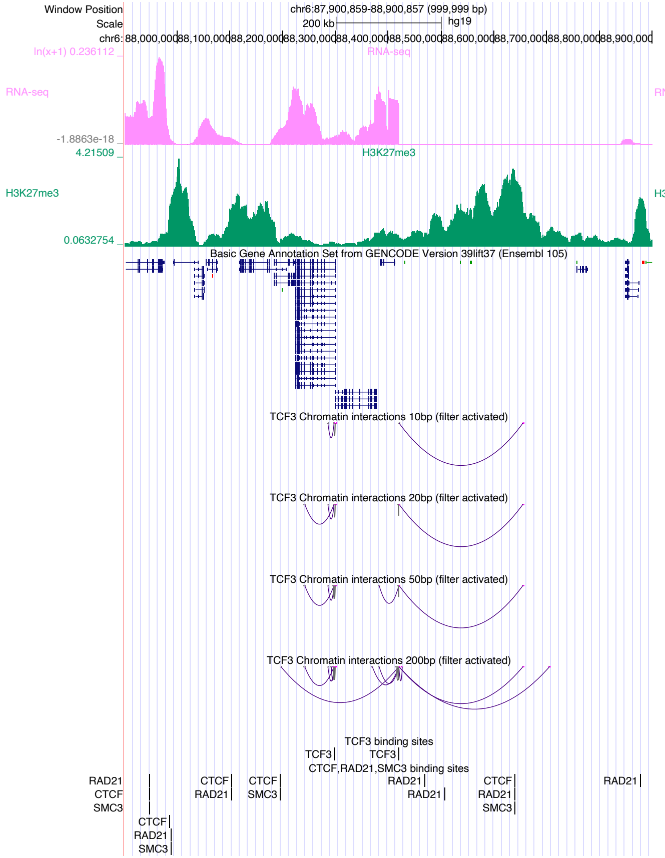

Window Position chr6:87,900,859-88,900,857 (999,999 bp)
Scale 200 kb hg19
RNA-seq
H3K27me3
Basic Gene Annotation Set from GENCODE Version 39lift37 (Ensembl 105)
TCF3 Chromatin interactions 10bp (filter activated)
TCF3 Chromatin interactions 20bp (filter activated)
TCF3 Chromatin interactions 50bp (filter activated)
TCF3 Chromatin interactions 200bp (filter activated)
TCF3 binding sites
CTCF,RAD21,SMC3 binding sites


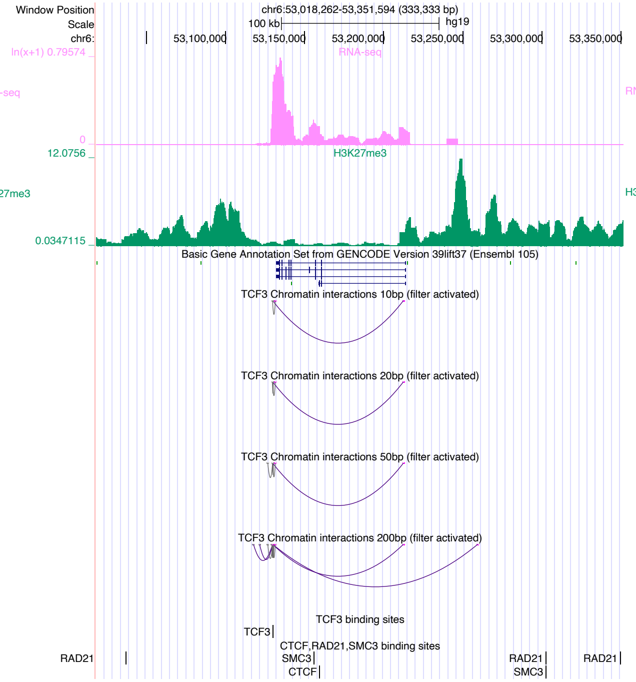

Window Position chr6:53,018,262-53,351,594 (333,333 bp)
Scale 100 kb hg19
RNA-seq
H3K27me3
Basic Gene Annotation Set from GENCODE Version 39lift37 (Ensembl 105)
TCF3 Chromatin interactions 10bp (filter activated)
TCF3 Chromatin interactions 20bp (filter activated)
TCF3 Chromatin interactions 50bp (filter activated)
TCF3 Chromatin interactions 200bp (filter activated)
TCF3 binding sites
CTCF,RAD21,SMC3 binding sites


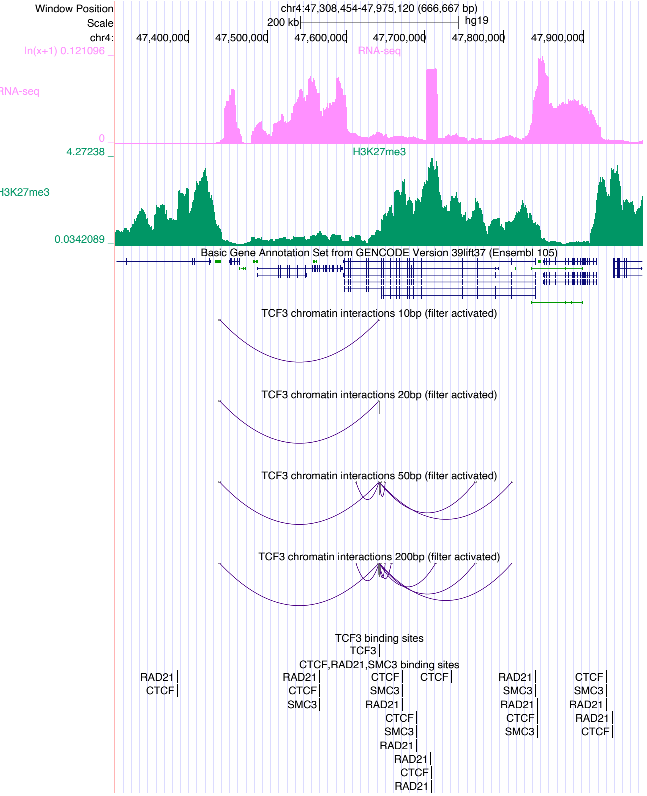

Window Position chr4:47,308,454-47,975,120 (666,667 bp)
Scale 200 kb hg19
RNA-seq
H3K27me3
Basic Gene Annotation Set from GENCODE Version 39lift37 (Ensembl 105)
TCF3 chromatin interactions 10bp (filter activated)
TCF3 chromatin interactions 20bp (filter activated)
TCF3 chromatin interactions 50bp (filter activated)
TCF3 chromatin interactions 200bp (filter activated)
TCF3 binding sites
CTCF,RAD21,SMC3 binding sites

### TCF4

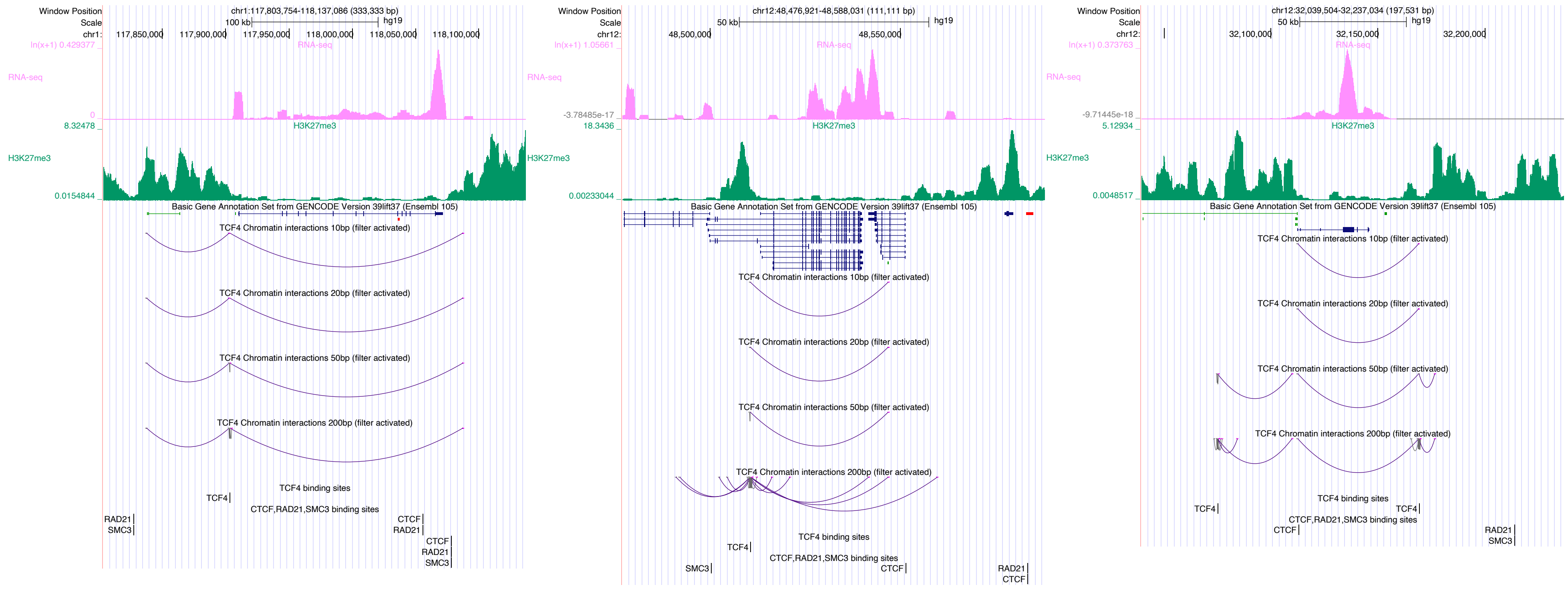
Window Position
chr1:117,803,754-118,137,086 (333,333 bp)
Scale
100 kb
hg19
chr1:
117,850,000
117,900,000
117,950,000
118,000,000
118,050,000
118,100,000
ln(x+1) 0.429377
RNA-seq
0
8.32478
H3K27me3
0.0154844
Basic Gene Annotation Set from GENCODE Version 39lift37 (Ensembl 105)
TCF4 Chromatin interactions 10bp (filter activated)
TCF4 Chromatin interactions 20bp (filter activated)
TCF4 Chromatin interactions 50bp (filter activated)
TCF4 Chromatin interactions 200bp (filter activated)
TCF4 binding sites
TCF4
CTCF,RAD21,SMC3 binding sites
RAD21
SMC3
CTCF
RAD21
CTCF
RAD21
SMC3
Window Position
chr12:48,476,921-48,588,031 (111,111 bp)
Scale
50 kb
hg19
chr12:
48,500,000
48,550,000
ln(x+1) 1.05661
RNA-seq
-3.78485e-17
18.3436
H3K27me3
0.00233044
Basic Gene Annotation Set from GENCODE Version 39lift37 (Ensembl 105)
TCF4 Chromatin interactions 10bp (filter activated)
TCF4 Chromatin interactions 20bp (filter activated)
TCF4 Chromatin interactions 50bp (filter activated)
TCF4 Chromatin interactions 200bp (filter activated)
TCF4 binding sites
TCF4
CTCF,RAD21,SMC3 binding sites
SMC3
CTCF
RAD21
CTCF
Window Position
chr12:32,039,504-32,237,034 (197,531 bp)
Scale
50 kb
hg19
chr12:
32,100,000
32,150,000
32,200,000
ln(x+1) 0.373763
RNA-seq
-9.71445e-18
5.12934
H3K27me3
0.0048517
Basic Gene Annotation Set from GENCODE Version 39lift37 (Ensembl 105)
TCF4 Chromatin interactions 10bp (filter activated)
TCF4 Chromatin interactions 20bp (filter activated)
TCF4 Chromatin interactions 50bp (filter activated)
TCF4 Chromatin interactions 200bp (filter activated)
TCF4 binding sites
TCF4
TCF4
CTCF,RAD21,SMC3 binding sites
CTCF
RAD21
SMC3

### TCF12

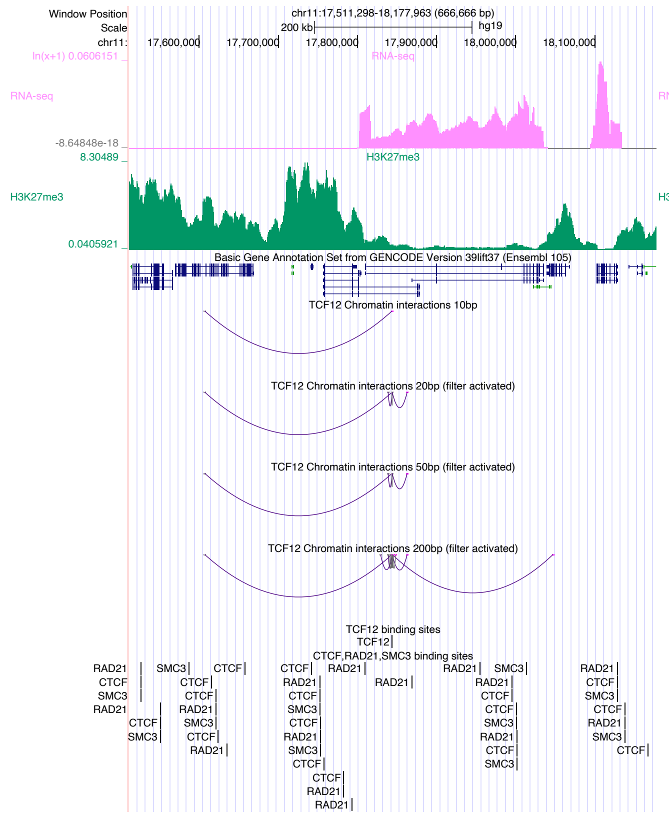

chr11:17,511,298-18,177,963 (666,666 bp)
RNA-seq
H3K27me3
Basic Gene Annotation Set from GENCODE Version 39lift37 (Ensembl 105)
TCF12 Chromatin interactions 10bp
TCF12 Chromatin interactions 20bp (filter activated)
TCF12 Chromatin interactions 50bp (filter activated)
TCF12 Chromatin interactions 200bp (filter activated)
TCF12 binding sites
CTCF,RAD21,SMC3 binding sites


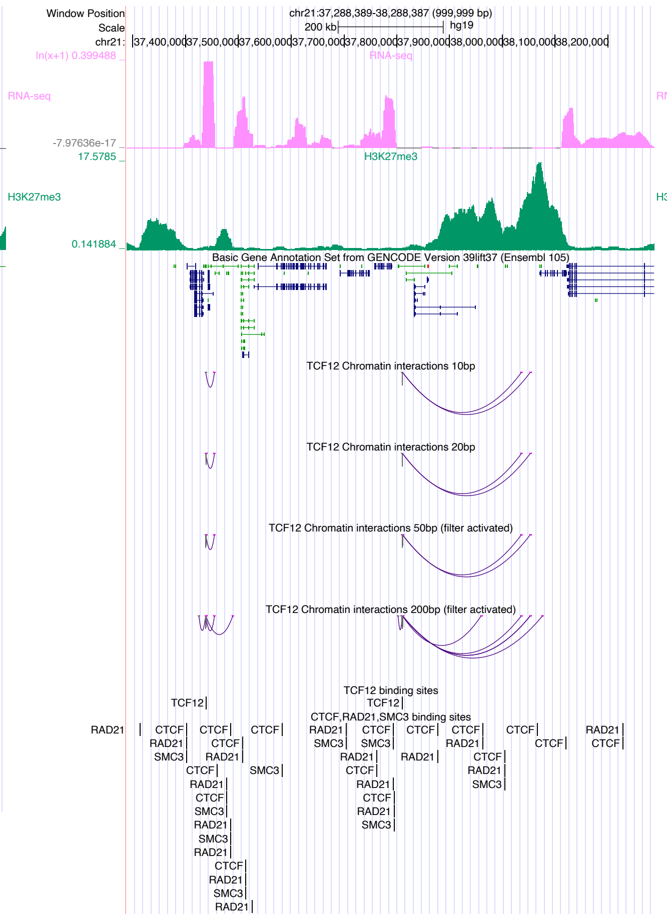

chr21:37,288,389-38,288,387 (999,999 bp)
RNA-seq
H3K27me3
Basic Gene Annotation Set from GENCODE Version 39lift37 (Ensembl 105)
TCF12 Chromatin interactions 10bp
TCF12 Chromatin interactions 20bp
TCF12 Chromatin interactions 50bp (filter activated)
TCF12 Chromatin interactions 200bp (filter activated)
TCF12 binding sites
CTCF,RAD21,SMC3 binding sites


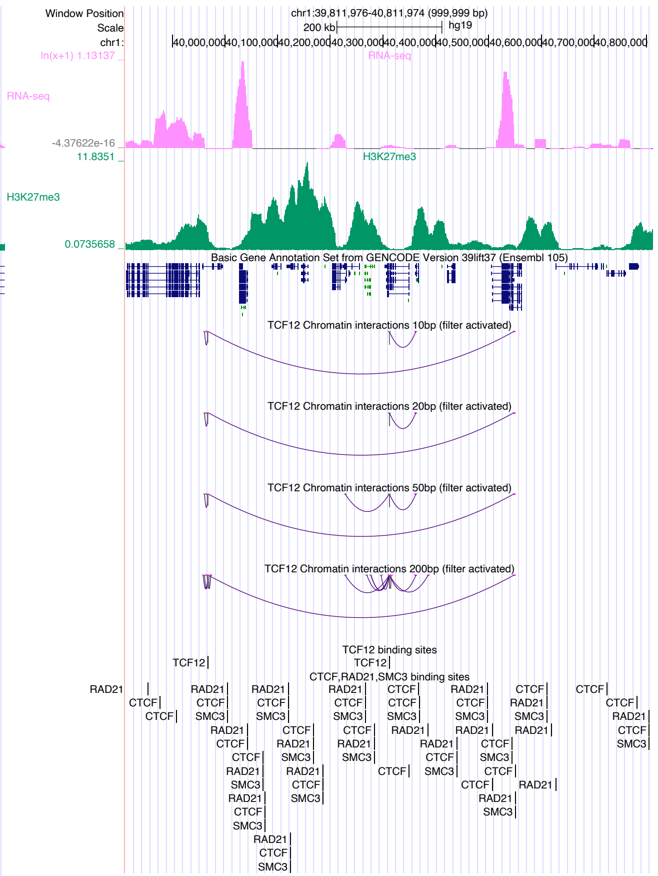

chr1:39,811,976-40,811,974 (999,999 bp)
RNA-seq
H3K27me3
Basic Gene Annotation Set from GENCODE Version 39lift37 (Ensembl 105)
TCF12 Chromatin interactions 10bp (filter activated)
TCF12 Chromatin interactions 20bp (filter activated)
TCF12 Chromatin interactions 50bp (filter activated)
TCF12 Chromatin interactions 200bp (filter activated)
TCF12 binding sites
CTCF,RAD21,SMC3 binding sites

### TEAD3

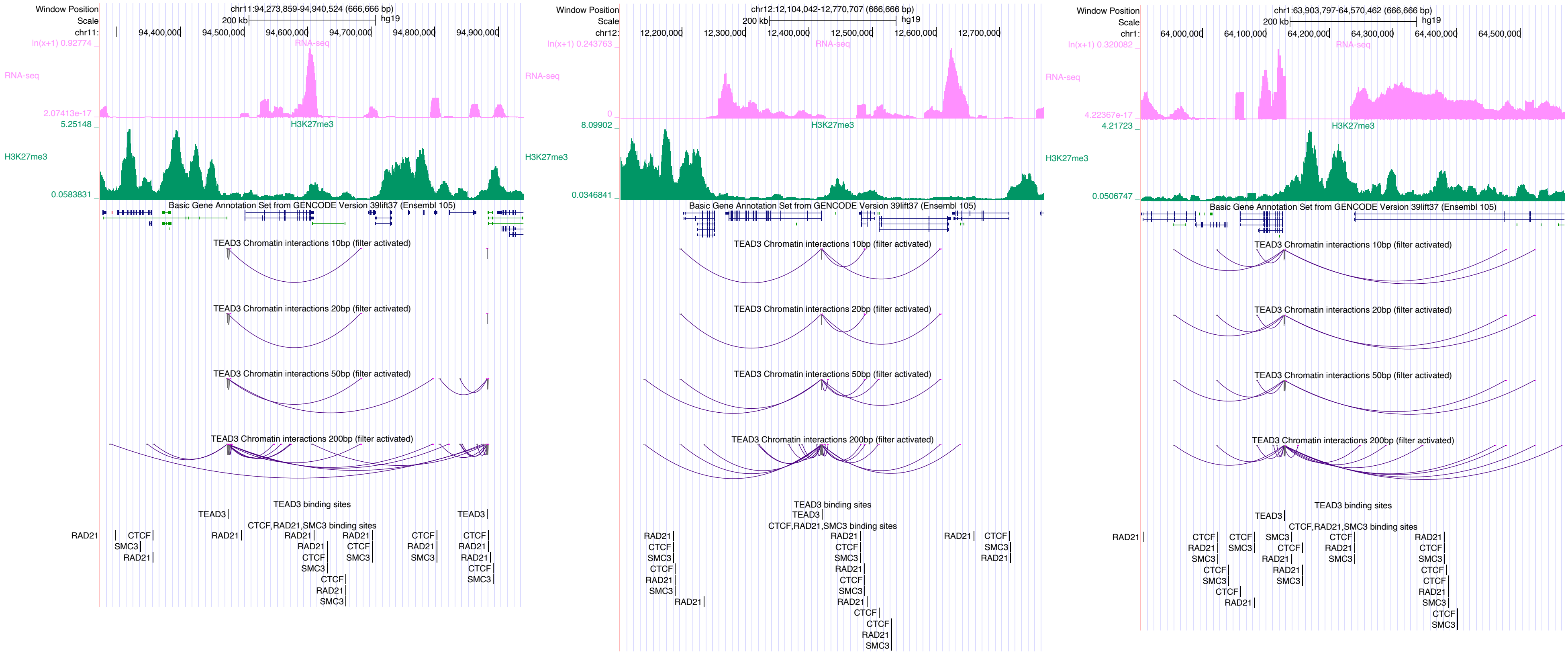

Window Position
Scale
chr11:94,273,859-94,940,524 (666,666 bp)
200 kb
hg19
chr11:
94,400,000
94,500,000
94,600,000
94,700,000
94,800,000
94,900,000
ln(x+1) 0.92774
RNA-seq
2.07413e-17
5.25148
H3K27me3
0.0583831
Basic Gene Annotation Set from GENCODE Version 39lift37 (Ensembl 105)
TEAD3 Chromatin interactions 10bp (filter activated)
TEAD3 Chromatin interactions 20bp (filter activated)
TEAD3 Chromatin interactions 50bp (filter activated)
TEAD3 Chromatin interactions 200bp (filter activated)
TEAD3 binding sites
CTCF,RAD21,SMC3 binding sites
chr12:12,104,042-12,770,707 (666,666 bp)
chr12:
12,200,000
12,300,000
12,400,000
12,500,000
12,600,000
12,700,000
ln(x+1) 0.243763
0
8.09902
0.0346841
chr1:63,903,797-64,570,462 (666,666 bp)
chr1:
64,000,000
64,100,000
64,200,000
64,300,000
64,400,000
64,500,000
ln(x+1) 0.320082
4.22367e-17
4.21723
0.0506747

### TFAP2C

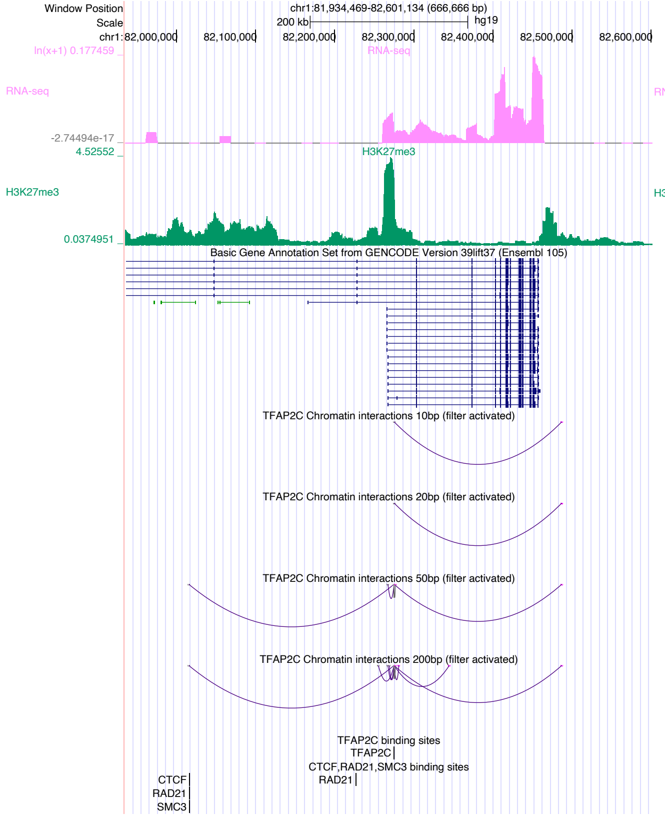

Window Position
chr1:81,934,469-82,601,134 (666,666 bp)
Scale
200 kb
hg19
RNA-seq
H3K27me3
Basic Gene Annotation Set from GENCODE Version 39lift37 (Ensembl 105)
TFAP2C Chromatin interactions 10bp (filter activated)
TFAP2C Chromatin interactions 20bp (filter activated)
TFAP2C Chromatin interactions 50bp (filter activated)
TFAP2C Chromatin interactions 200bp (filter activated)
TFAP2C binding sites
CTCF,RAD21,SMC3 binding sites


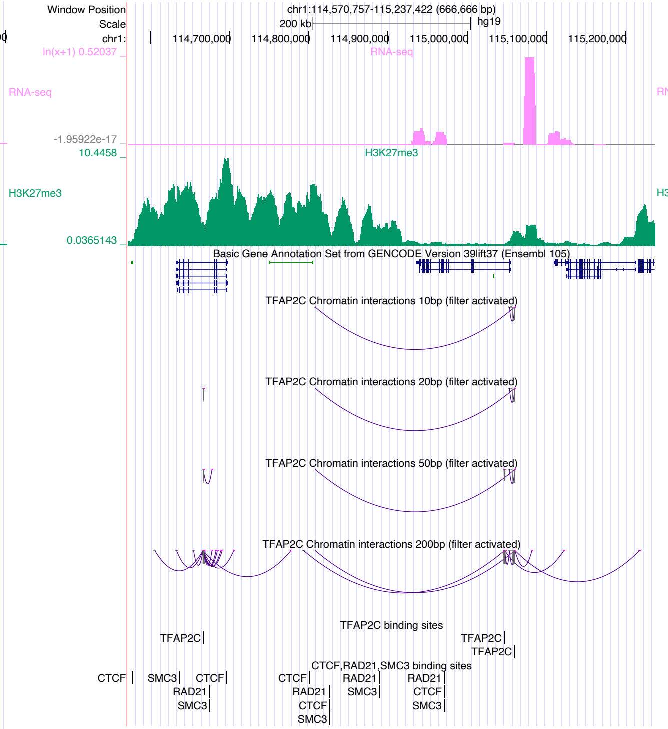

Window Position
chr1:114,570,757-115,237,422 (666,666 bp)
Scale
200 kb
hg19
RNA-seq
H3K27me3
Basic Gene Annotation Set from GENCODE Version 39lift37 (Ensembl 105)
TFAP2C Chromatin interactions 10bp (filter activated)
TFAP2C Chromatin interactions 20bp (filter activated)
TFAP2C Chromatin interactions 50bp (filter activated)
TFAP2C Chromatin interactions 200bp (filter activated)
TFAP2C binding sites
CTCF,RAD21,SMC3 binding sites


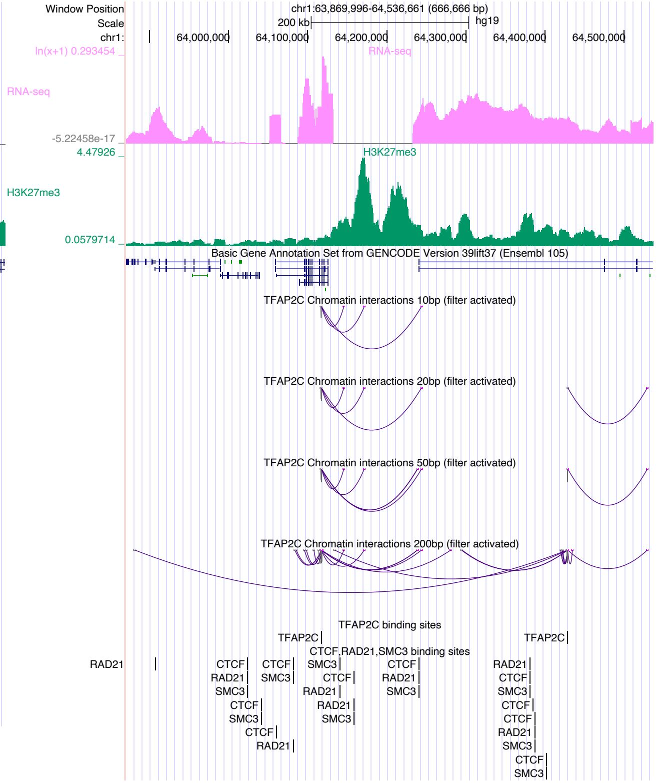

Window Position
chr1:63,869,996-64,536,661 (666,666 bp)
Scale
200 kb
hg19
RNA-seq
H3K27me3
Basic Gene Annotation Set from GENCODE Version 39lift37 (Ensembl 105)
TFAP2C Chromatin interactions 10bp (filter activated)
TFAP2C Chromatin interactions 20bp (filter activated)
TFAP2C Chromatin interactions 50bp (filter activated)
TFAP2C Chromatin interactions 200bp (filter activated)
TFAP2C binding sites
CTCF,RAD21,SMC3 binding sites

### TFAP4

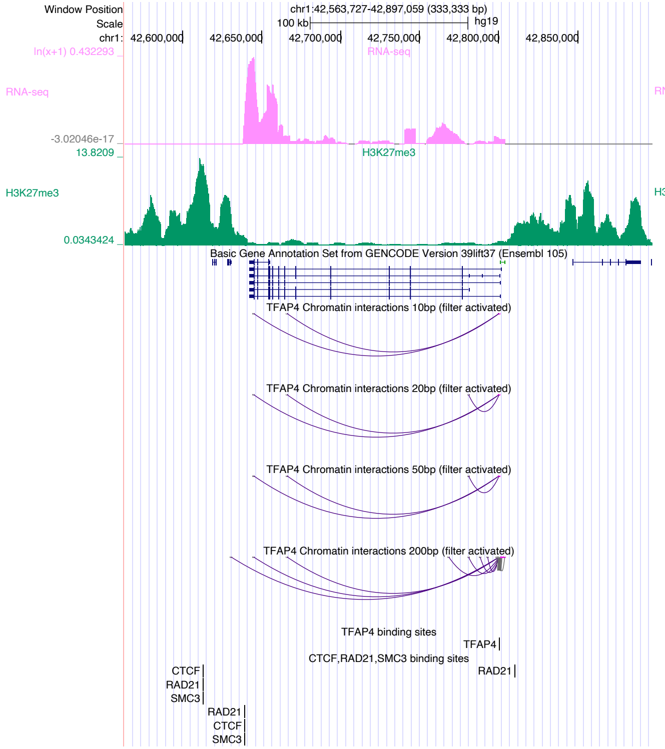

Window Position
chr1:42,563,727-42,897,059 (333,333 bp)
Scale
100 kb
hg19
chr1:
42,600,000
42,650,000
42,700,000
42,750,000
42,800,000
42,850,000
ln(x+1) 0.432293
RNA-seq
-3.02046e-17
13.8209
H3K27me3
0.0343424
Basic Gene Annotation Set from GENCODE Version 39lift37 (Ensembl 105)
TFAP4 Chromatin interactions 10bp (filter activated)
TFAP4 Chromatin interactions 20bp (filter activated)
TFAP4 Chromatin interactions 50bp (filter activated)
TFAP4 Chromatin interactions 200bp (filter activated)
TFAP4 binding sites
TFAP4
CTCF,RAD21,SMC3 binding sites


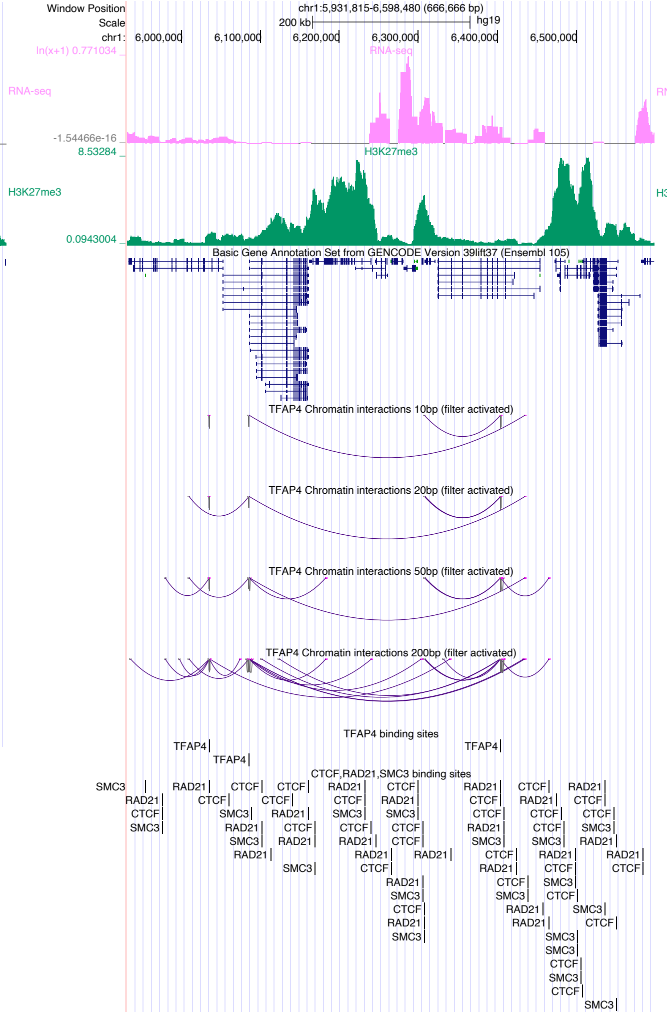

Window Position
chr1:5,931,815-6,598,480 (666,666 bp)
Scale
200 kb
hg19
chr1:
6,000,000
6,100,000
6,200,000
6,300,000
6,400,000
6,500,000
ln(x+1) 0.771034
RNA-seq
-1.54466e-16
8.53284
H3K27me3
0.0943004
Basic Gene Annotation Set from GENCODE Version 39lift37 (Ensembl 105)
TFAP4 Chromatin interactions 10bp (filter activated)
TFAP4 Chromatin interactions 20bp (filter activated)
TFAP4 Chromatin interactions 50bp (filter activated)
TFAP4 Chromatin interactions 200bp (filter activated)
TFAP4 binding sites
TFAP4
CTCF,RAD21,SMC3 binding sites


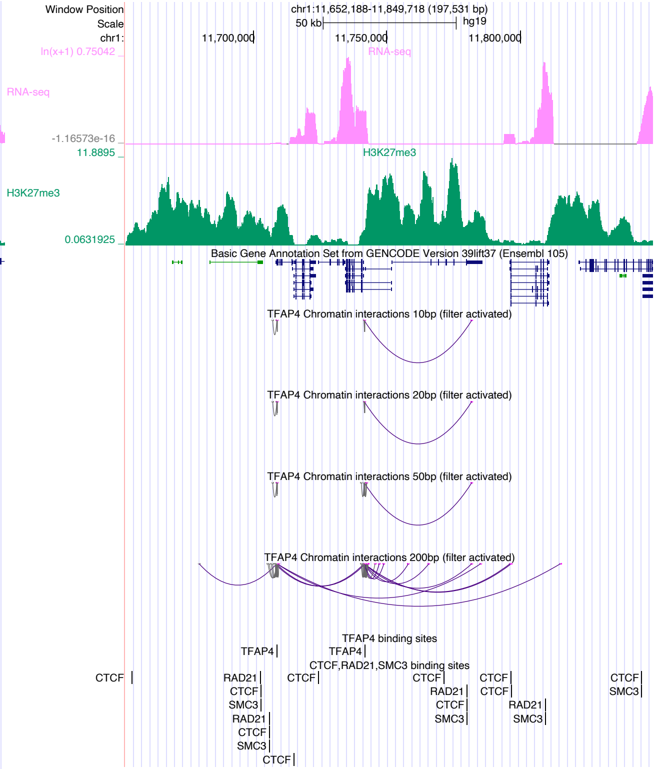

Window Position
chr1:11,652,188-11,849,718 (197,531 bp)
Scale
50 kb
hg19
chr1:
11,700,000
11,750,000
11,800,000
ln(x+1) 0.75042
RNA-seq
-1.16573e-16
11.8895
H3K27me3
0.0631925
Basic Gene Annotation Set from GENCODE Version 39lift37 (Ensembl 105)
TFAP4 Chromatin interactions 10bp (filter activated)
TFAP4 Chromatin interactions 20bp (filter activated)
TFAP4 Chromatin interactions 50bp (filter activated)
TFAP4 Chromatin interactions 200bp (filter activated)
TFAP4 binding sites
TFAP4
CTCF,RAD21,SMC3 binding sites

# TP53

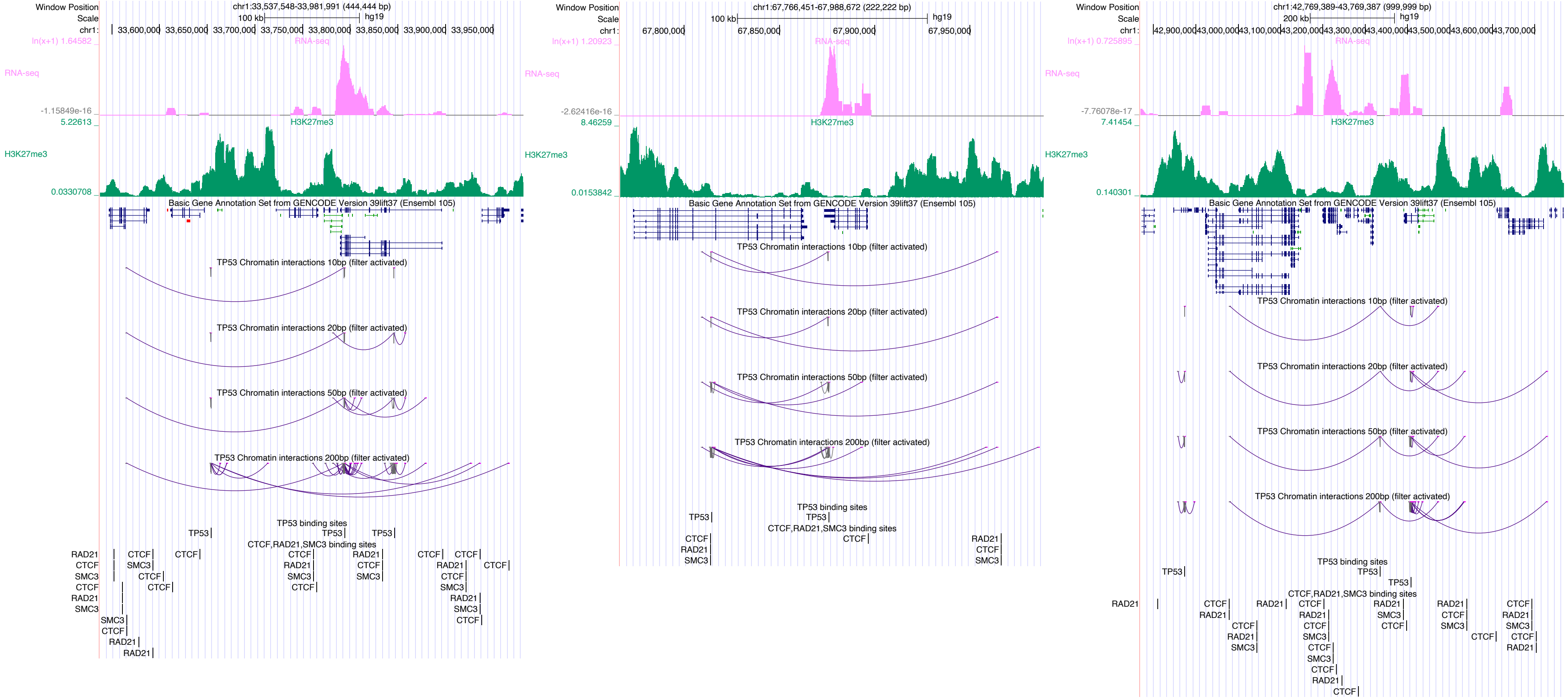

Window Position
Scale
chr1:
chr1:33,537,548-33,981,991 (444,444 bp)
100 kb
hg19
33,600,000
33,650,000
33,700,000
33,750,000
33,800,000
33,850,000
33,900,000
33,950,000
ln(x+1) 1.64582
RNA-seq
-1.15849e-16
5.22613
H3K27me3
0.0330708
Basic Gene Annotation Set from GENCODE Version 39lift37 (Ensembl 105)
TP53 Chromatin interactions 10bp (filter activated)
TP53 Chromatin interactions 20bp (filter activated)
TP53 Chromatin interactions 50bp (filter activated)
TP53 Chromatin interactions 200bp (filter activated)
TP53 binding sites
TP53
CTCF,RAD21,SMC3 binding sites
RAD21
CTCF
SMC3
chr1:67,766,451-67,988,672 (222,222 bp)
67,800,000
67,850,000
67,900,000
67,950,000
ln(x+1) 1.20923
-2.62416e-16
8.46259
0.0153842
chr1:42,769,389-43,769,387 (999,999 bp)
200 kb
42,900,000
43,000,000
43,100,000
43,200,000
43,300,000
43,400,000
43,500,000
43,600,000
43,700,000
ln(x+1) 0.725895
-7.76078e-17
7.41454
0.140301

### TRIM28

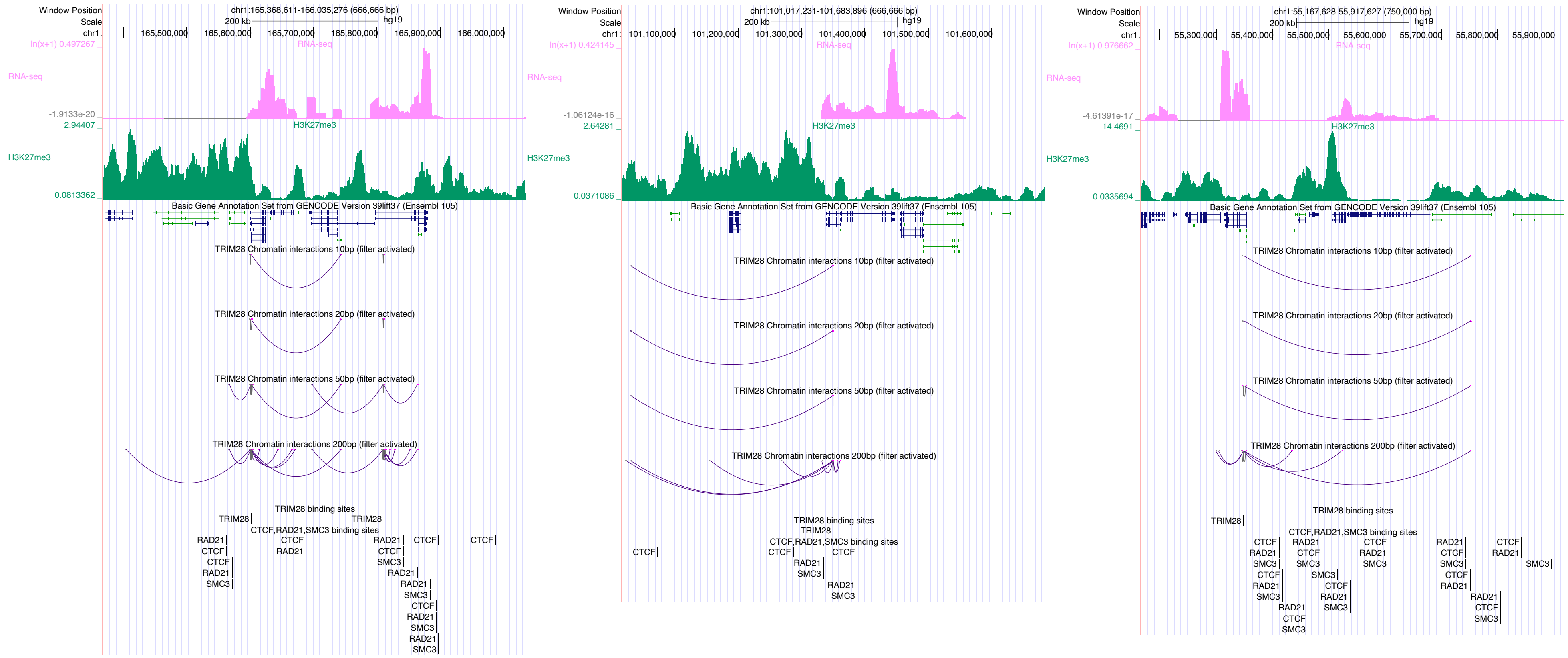

Window Position
Scale
chr1:165,368,611-166,035,276 (666,666 bp)
200 kb
hg19
RNA-seq
H3K27me3
Basic Gene Annotation Set from GENCODE Version 39lift37 (Ensembl 105)
TRIM28 Chromatin interactions 10bp (filter activated)
TRIM28 Chromatin interactions 20bp (filter activated)
TRIM28 Chromatin interactions 50bp (filter activated)
TRIM28 Chromatin interactions 200bp (filter activated)
TRIM28 binding sites
CTCF,RAD21,SMC3 binding sites
chr1:101,017,231-101,683,896 (666,666 bp)
chr1:55,167,628-55,917,627 (750,000 bp)

### USF1

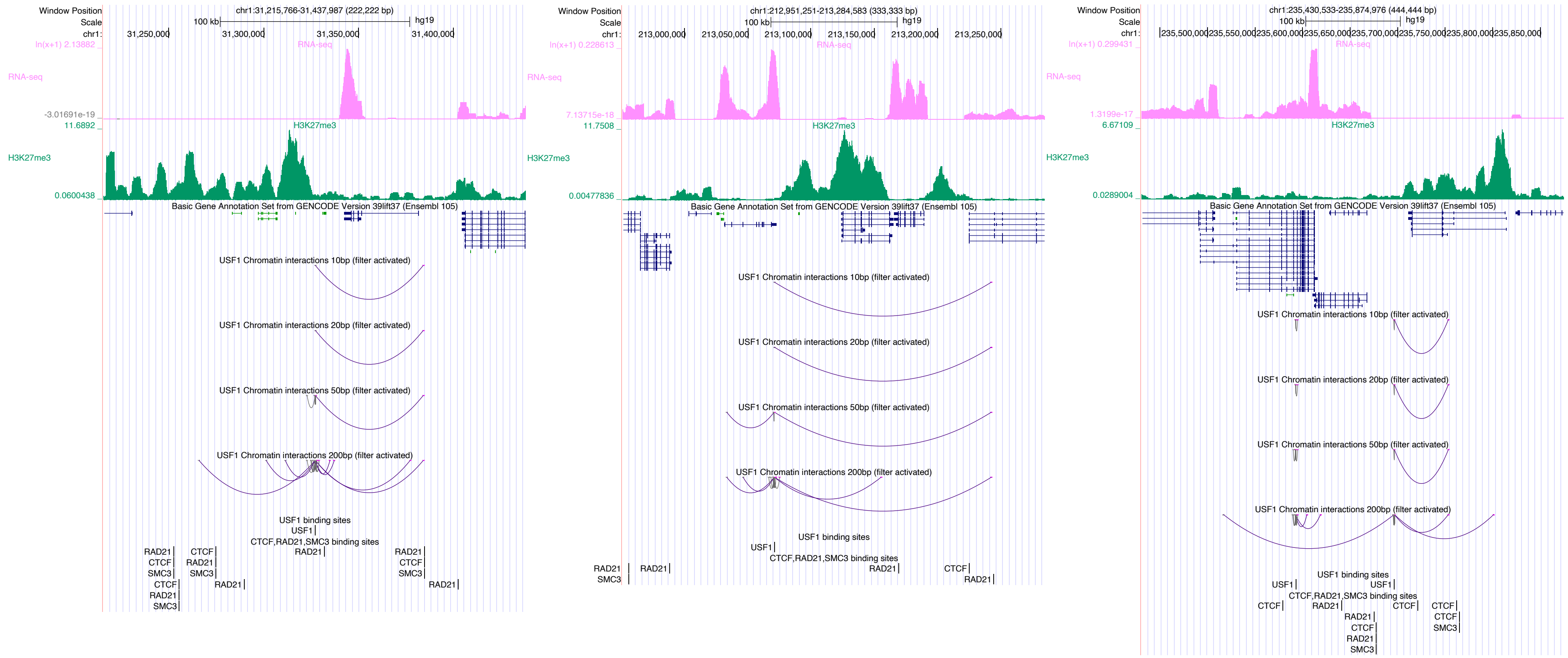

Window Position
Scale
chr1:
chr1:31,215,766-31,437,987 (222,222 bp)
100 kb
hg19
31,250,000
31,300,000
31,350,000
31,400,000
ln(x+1) 2.13882
RNA-seq
-3.01691e-19
11.6892
H3K27me3
0.0600438
Basic Gene Annotation Set from GENCODE Version 39lift37 (Ensembl 105)
USF1 Chromatin interactions 10bp (filter activated)
USF1 Chromatin interactions 20bp (filter activated)
USF1 Chromatin interactions 50bp (filter activated)
USF1 Chromatin interactions 200bp (filter activated)
USF1 binding sites
USF1
CTCF,RAD21,SMC3 binding sites
RAD21
CTCF
SMC3
chr1:212,951,251-213,284,583 (333,333 bp)
213,000,000
213,050,000
213,100,000
213,150,000
213,200,000
213,250,000
ln(x+1) 0.228613
7.13715e-18
11.7508
0.00477836
chr1:235,430,533-235,874,976 (444,444 bp)
235,500,000
235,550,000
235,600,000
235,650,000
235,700,000
235,750,000
235,800,000
235,850,000
ln(x+1) 0.299431
1.3199e-17
6.67109
0.0289004

### VDR

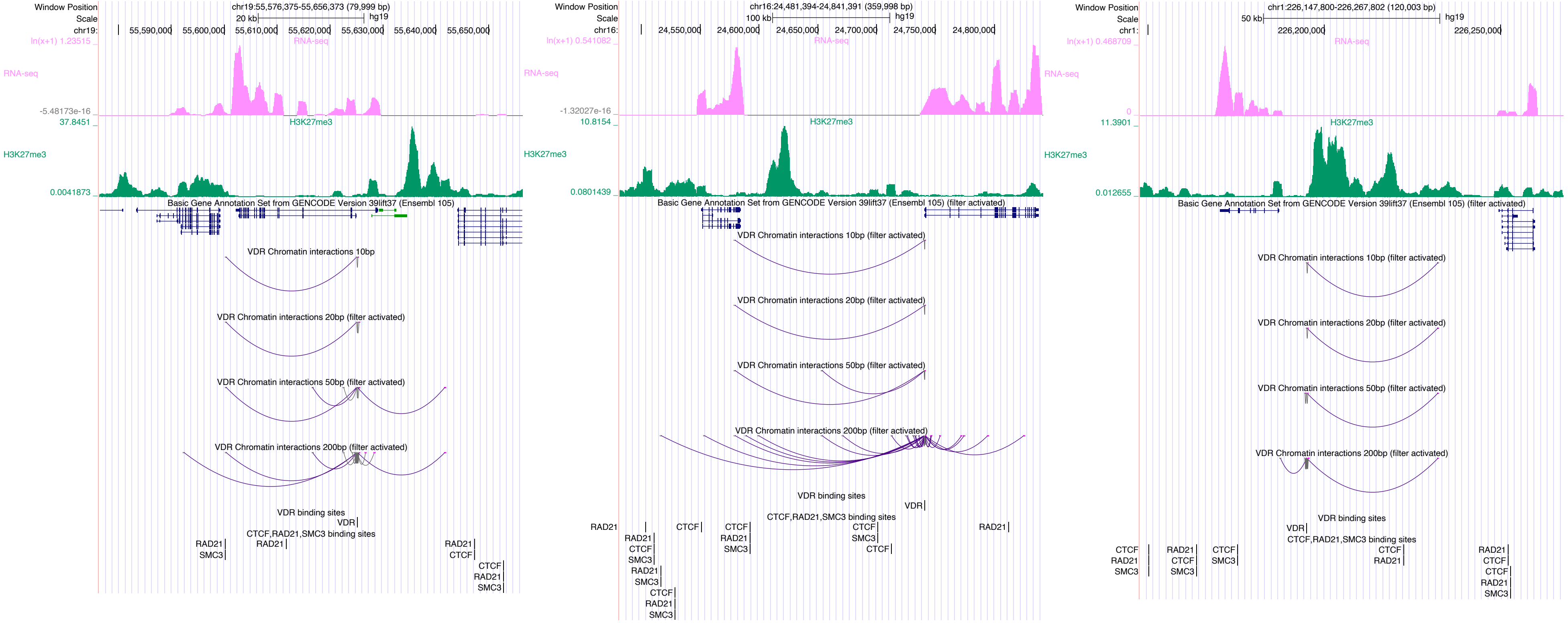

Window Position
Scale
chr19:
chr19:55,576,375-55,656,373 (79,999 bp)
20 kb
hg19
55,590,000
55,600,000
55,610,000
55,620,000
55,630,000
55,640,000
55,650,000
ln(x+1) 1.23515
RNA-seq
RNA-seq
-5.48173e-16
37.8451
H3K27me3
H3K27me3
0.0041873
Basic Gene Annotation Set from GENCODE Version 39lift37 (Ensembl 105)
VDR Chromatin interactions 10bp
VDR Chromatin interactions 20bp (filter activated)
VDR Chromatin interactions 50bp (filter activated)
VDR Chromatin interactions 200bp (filter activated)
VDR binding sites
VDR
CTCF,RAD21,SMC3 binding sites
RAD21
SMC3
RAD21
RAD21
CTCF
CTCF
RAD21
SMC3
Window Position
Scale
chr16:
chr16:24,481,394-24,841,391 (359,998 bp)
100 kb
hg19
24,550,000
24,600,000
24,650,000
24,700,000
24,750,000
24,800,000
ln(x+1) 0.541082
RNA-seq
RNA-seq
-1.32027e-16
10.8154
H3K27me3
H3K27me3
0.0801439
Basic Gene Annotation Set from GENCODE Version 39lift37 (Ensembl 105) (filter activated)
VDR Chromatin interactions 10bp (filter activated)
VDR Chromatin interactions 20bp (filter activated)
VDR Chromatin interactions 50bp (filter activated)
VDR Chromatin interactions 200bp (filter activated)
VDR binding sites
VDR
CTCF,RAD21,SMC3 binding sites
RAD21
CTCF
CTCF
RAD21
RAD21
CTCF
SMC3
CTCF
RAD21
SMC3
SMC3
CTCF
RAD21
SMC3
CTCF
RAD21
SMC3
Window Position
Scale
chr1:
chr1:226,147,800-226,267,802 (120,003 bp)
50 kb
hg19
226,200,000
226,250,000
ln(x+1) 0.468709
RNA-seq
RNA-seq
0
11.3901
H3K27me3
H3K27me3
0.012655
Basic Gene Annotation Set from GENCODE Version 39lift37 (Ensembl 105) (filter activated)
VDR Chromatin interactions 10bp (filter activated)
VDR Chromatin interactions 20bp (filter activated)
VDR Chromatin interactions 50bp (filter activated)
VDR Chromatin interactions 200bp (filter activated)
VDR binding sites
VDR
CTCF,RAD21,SMC3 binding sites
CTCF
RAD21
SMC3
RAD21
CTCF
SMC3
CTCF
SMC3
CTCF
RAD21
RAD21
CTCF
CTCF
RAD21
SMC3

### ZBTB7A

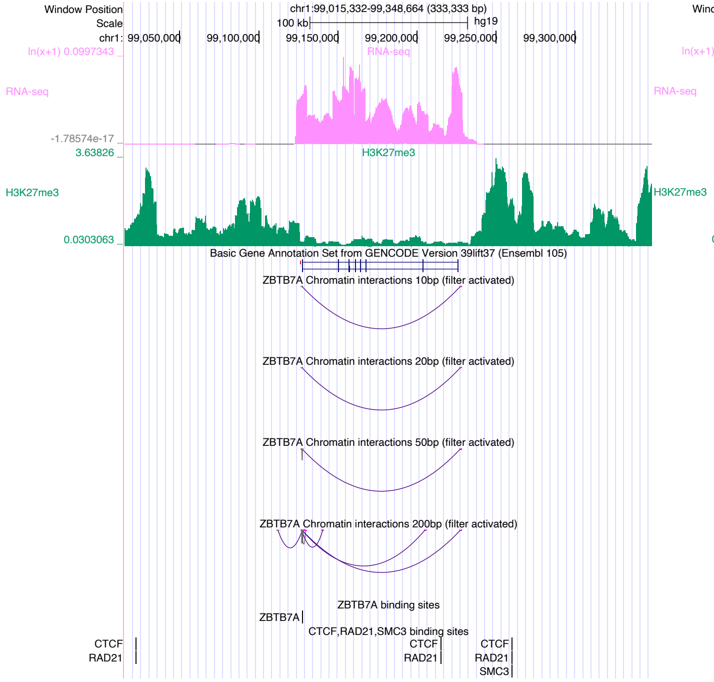

Window Position
chr1:99,015,332-99,348,664 (333,333 bp)
Scale
100 kb
hg19
chr1: 99,050,000 99,100,000 99,150,000 99,200,000 99,250,000 99,300,000
RNA-seq
H3K27me3
Basic Gene Annotation Set from GENCODE Version 39lift37 (Ensembl 105)
ZBTB7A Chromatin interactions 10bp (filter activated)
ZBTB7A Chromatin interactions 20bp (filter activated)
ZBTB7A Chromatin interactions 50bp (filter activated)
ZBTB7A Chromatin interactions 200bp (filter activated)
ZBTB7A binding sites
CTCF,RAD21,SMC3 binding sites


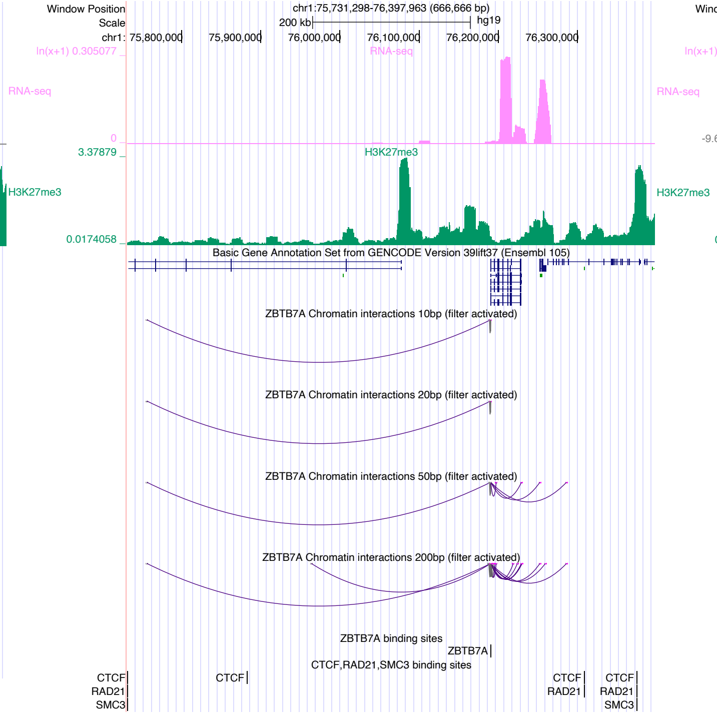

Window Position
chr1:75,731,298-76,397,963 (666,666 bp)
Scale
200 kb
hg19
chr1: 75,800,000 75,900,000 76,000,000 76,100,000 76,200,000 76,300,000
RNA-seq
H3K27me3
Basic Gene Annotation Set from GENCODE Version 39lift37 (Ensembl 105)
ZBTB7A Chromatin interactions 10bp (filter activated)
ZBTB7A Chromatin interactions 20bp (filter activated)
ZBTB7A Chromatin interactions 50bp (filter activated)
ZBTB7A Chromatin interactions 200bp (filter activated)
ZBTB7A binding sites
CTCF,RAD21,SMC3 binding sites


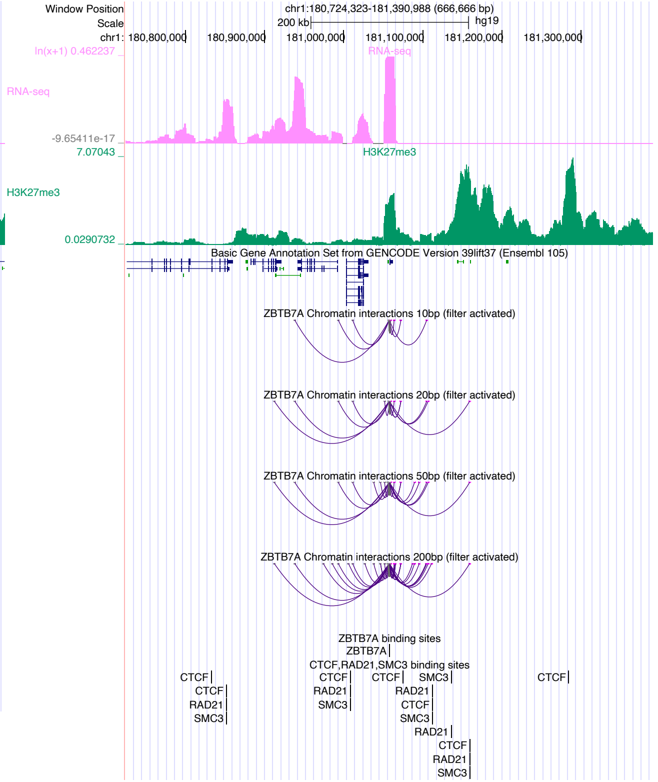

Window Position
chr1:180,724,323-181,390,988 (666,666 bp)
Scale
200 kb
hg19
chr1: 180,800,000 180,900,000 181,000,000 181,100,000 181,200,000 181,300,000
RNA-seq
H3K27me3
Basic Gene Annotation Set from GENCODE Version 39lift37 (Ensembl 105)
ZBTB7A Chromatin interactions 10bp (filter activated)
ZBTB7A Chromatin interactions 20bp (filter activated)
ZBTB7A Chromatin interactions 50bp (filter activated)
ZBTB7A Chromatin interactions 200bp (filter activated)
ZBTB7A binding sites
CTCF,RAD21,SMC3 binding sites

### ZEB1

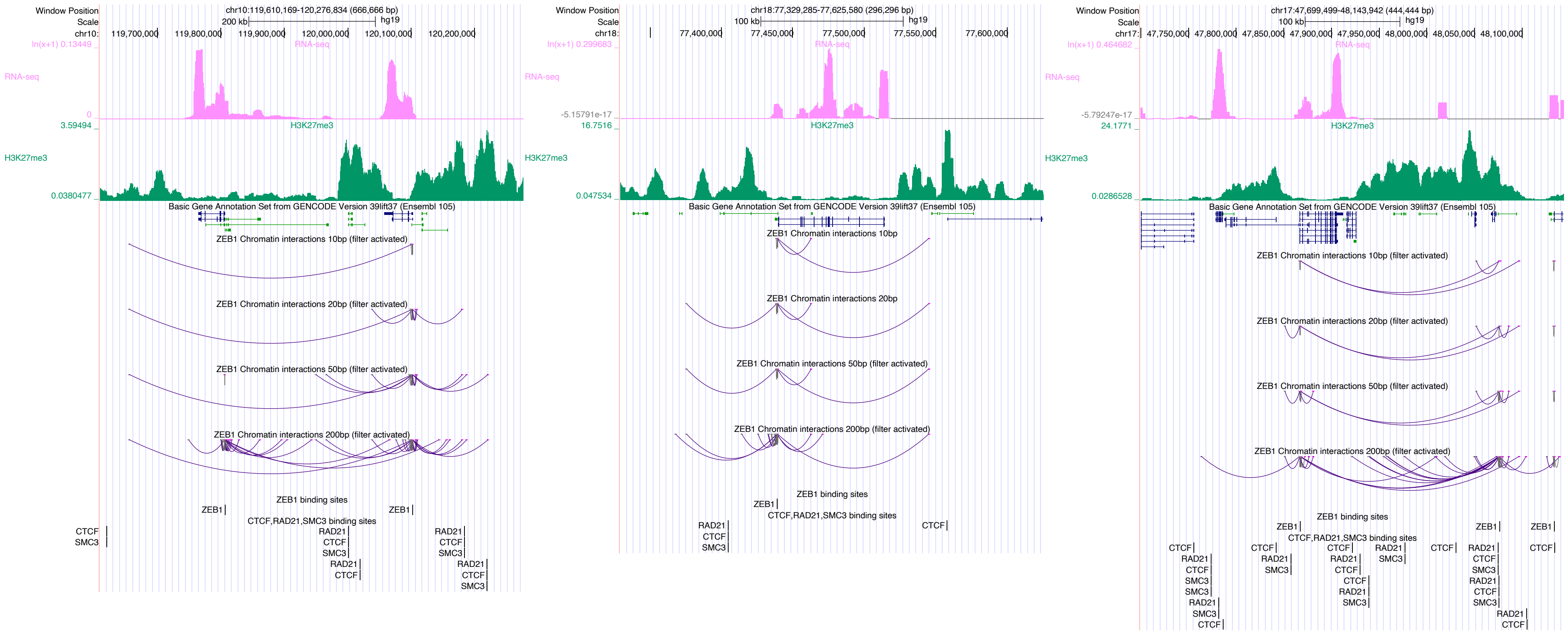
Window Position
Scale
chr10:119,610,169-120,276,834 (666,666 bp)
200 kb
hg19
chr10:
119,700,000
119,800,000
119,900,000
120,000,000
120,100,000
120,200,000
ln(x+1) 0.13449
RNA-seq
0
3.59494
H3K27me3
0.0380477
Basic Gene Annotation Set from GENCODE Version 39lift37 (Ensembl 105)
ZEB1 Chromatin interactions 10bp (filter activated)
ZEB1 Chromatin interactions 20bp (filter activated)
ZEB1 Chromatin interactions 50bp (filter activated)
ZEB1 Chromatin interactions 200bp (filter activated)
ZEB1 binding sites
ZEB1
CTCF,RAD21,SMC3 binding sites
CTCF
SMC3
RAD21
chr18:77,329,285-77,625,580 (296,296 bp)
100 kb
chr18:
77,400,000
77,450,000
77,500,000
77,550,000
77,600,000
ln(x+1) 0.299683
-5.15791e-17
16.7516
0.047534
ZEB1 Chromatin interactions 10bp
ZEB1 Chromatin interactions 20bp
chr17:47,699,499-48,143,942 (444,444 bp)
chr17:
47,750,000
47,800,000
47,850,000
47,900,000
47,950,000
48,000,000
48,050,000
48,100,000
ln(x+1) 0.464682
-5.79247e-17
24.1771
0.0286528

### ZFP3

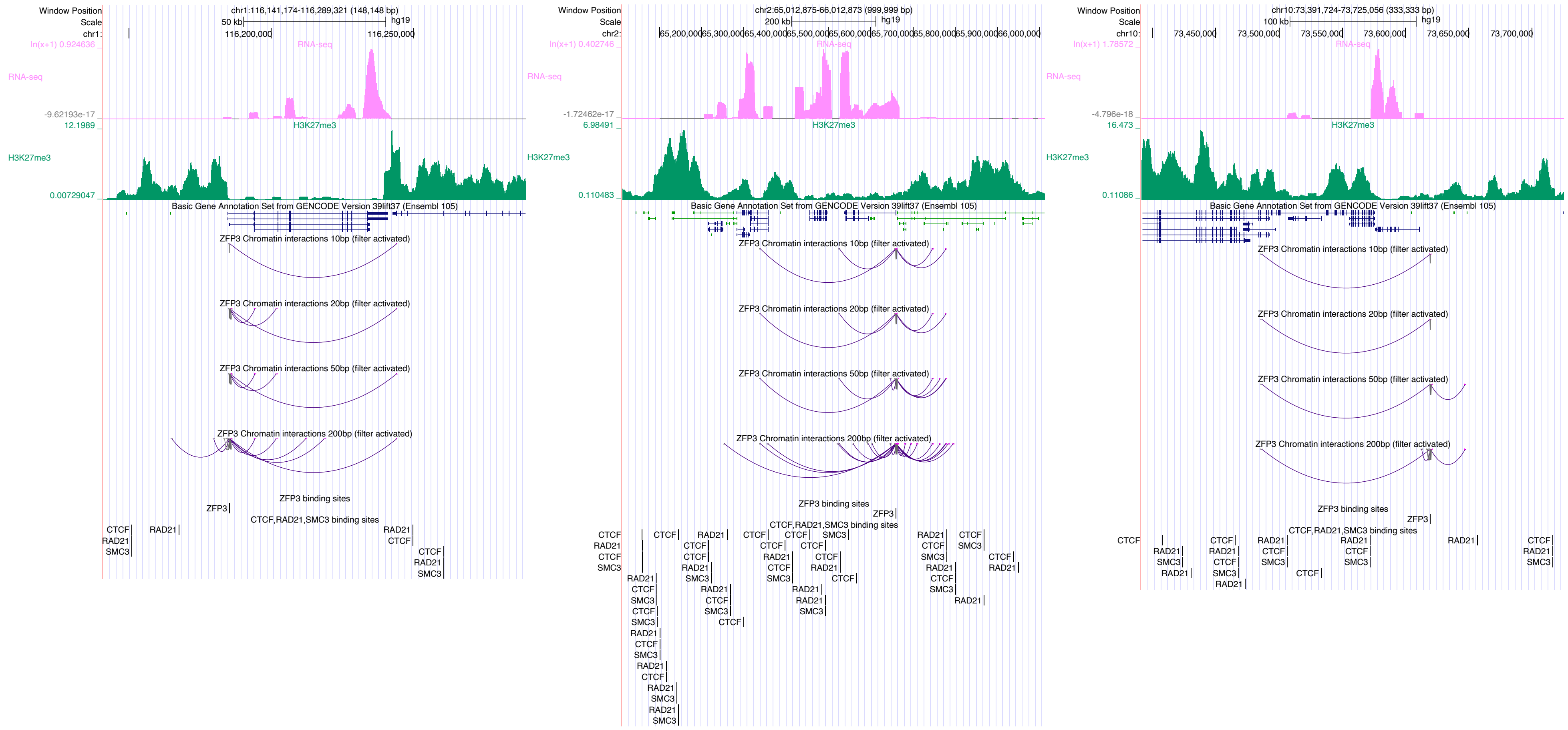

Window Position
chr1:116,141,174-116,289,321 (148,148 bp)
Scale
50 kb
hg19
116,200,000
116,250,000
RNA-seq
ln(x+1) 0.924636
-9.62193e-17
H3K27me3
12.1989
0.00729047
Basic Gene Annotation Set from GENCODE Version 39lift37 (Ensembl 105)
ZFP3 Chromatin interactions 10bp (filter activated)
ZFP3 Chromatin interactions 20bp (filter activated)
ZFP3 Chromatin interactions 50bp (filter activated)
ZFP3 Chromatin interactions 200bp (filter activated)
ZFP3 binding sites
ZFP3
CTCF,RAD21,SMC3 binding sites
CTCF
RAD21
SMC3
chr2:65,012,875-66,012,873 (999,999 bp)
200 kb
65,200,000
65,300,000
65,400,000
65,500,000
65,600,000
65,700,000
65,800,000
65,900,000
66,000,000
ln(x+1) 0.402746
-1.72462e-17
6.98491
0.110483
chr10:73,391,724-73,725,056 (333,333 bp)
100 kb
73,450,000
73,500,000
73,550,000
73,600,000
73,650,000
73,700,000
ln(x+1) 1.78572
-4.796e-18
16.473
0.11086

### ZIC2

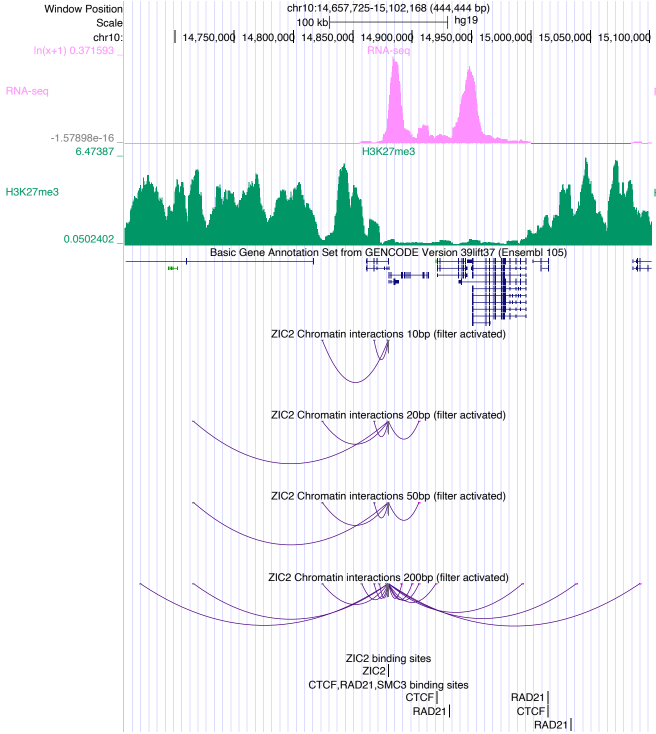

Window Position
Scale
chr10:
chr10:14,657,725-15,102,168 (444,444 bp)
100 kb
hg19
14,750,000
14,800,000
14,850,000
14,900,000
14,950,000
15,000,000
15,050,000
15,100,000
ln(x+1) 0.371593
RNA-seq
-1.57898e-16
6.47387
H3K27me3
0.0502402
Basic Gene Annotation Set from GENCODE Version 39lift37 (Ensembl 105)
ZIC2 Chromatin interactions 10bp (filter activated)
ZIC2 Chromatin interactions 20bp (filter activated)
ZIC2 Chromatin interactions 50bp (filter activated)
ZIC2 Chromatin interactions 200bp (filter activated)
ZIC2 binding sites
ZIC2
CTCF,RAD21,SMC3 binding sites
CTCF
RAD21
RAD21
CTCF
RAD21


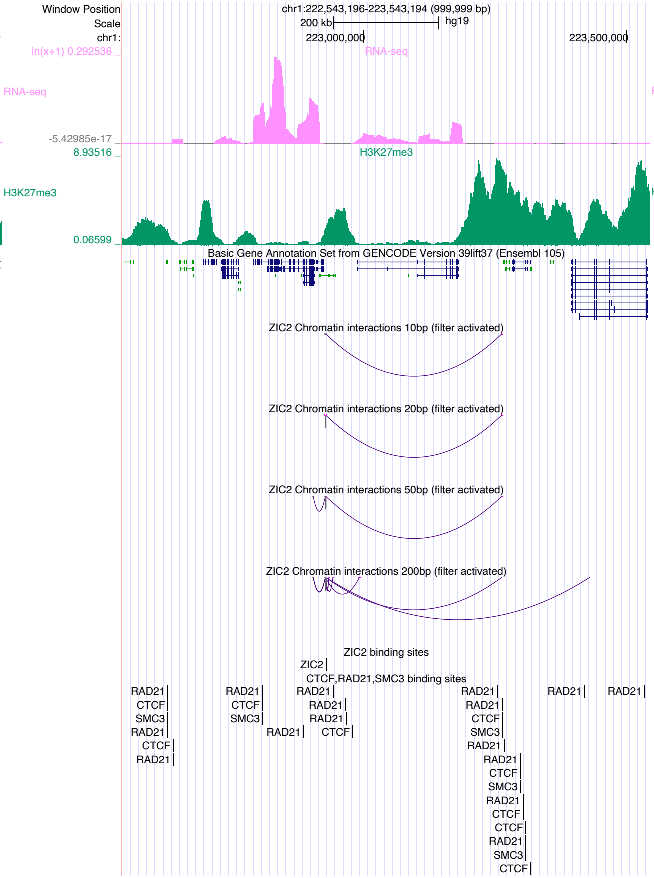

Window Position
Scale
chr1:
chr1:222,543,196-223,543,194 (999,999 bp)
200 kb
hg19
223,000,000
223,500,000
ln(x+1) 0.292536
RNA-seq
-5.42985e-17
8.93516
H3K27me3
0.06599
Basic Gene Annotation Set from GENCODE Version 39lift37 (Ensembl 105)
ZIC2 Chromatin interactions 10bp (filter activated)
ZIC2 Chromatin interactions 20bp (filter activated)
ZIC2 Chromatin interactions 50bp (filter activated)
ZIC2 Chromatin interactions 200bp (filter activated)
ZIC2 binding sites
ZIC2
CTCF,RAD21,SMC3 binding sites
RAD21
CTCF
SMC3
RAD21
CTCF
RAD21
RAD21
CTCF
SMC3
RAD21
RAD21
RAD21
RAD21
CTCF
RAD21
RAD21
CTCF
SMC3
RAD21
RAD21
CTCF
SMC3
RAD21
CTCF
CTCF
RAD21
SMC3
CTCF
RAD21
RAD21


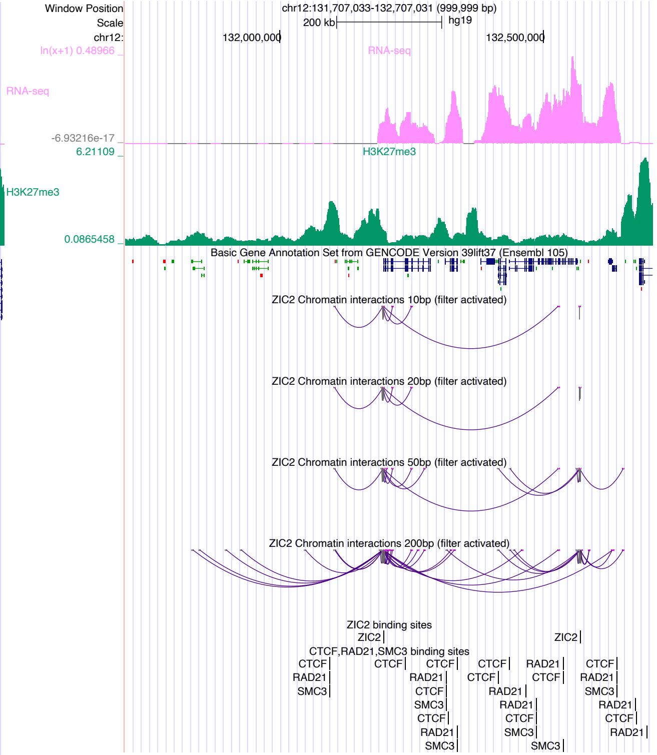

Window Position
Scale
chr12:
chr12:131,707,033-132,707,031 (999,999 bp)
200 kb
hg19
132,000,000
132,500,000
ln(x+1) 0.48966
RNA-seq
-6.93216e-17
6.21109
H3K27me3
0.0865458
Basic Gene Annotation Set from GENCODE Version 39lift37 (Ensembl 105)
ZIC2 Chromatin interactions 10bp (filter activated)
ZIC2 Chromatin interactions 20bp (filter activated)
ZIC2 Chromatin interactions 50bp (filter activated)
ZIC2 Chromatin interactions 200bp (filter activated)
ZIC2 binding sites
ZIC2
ZIC2
CTCF,RAD21,SMC3 binding sites
CTCF
RAD21
SMC3
CTCF
CTCF
RAD21
CTCF
SMC3
CTCF
RAD21
SMC3
CTCF
CTCF
RAD21
RAD21
CTCF
SMC3
RAD21
CTCF
SMC3
CTCF
RAD21
SMC3
RAD21
CTCF
RAD21

### ZNF18

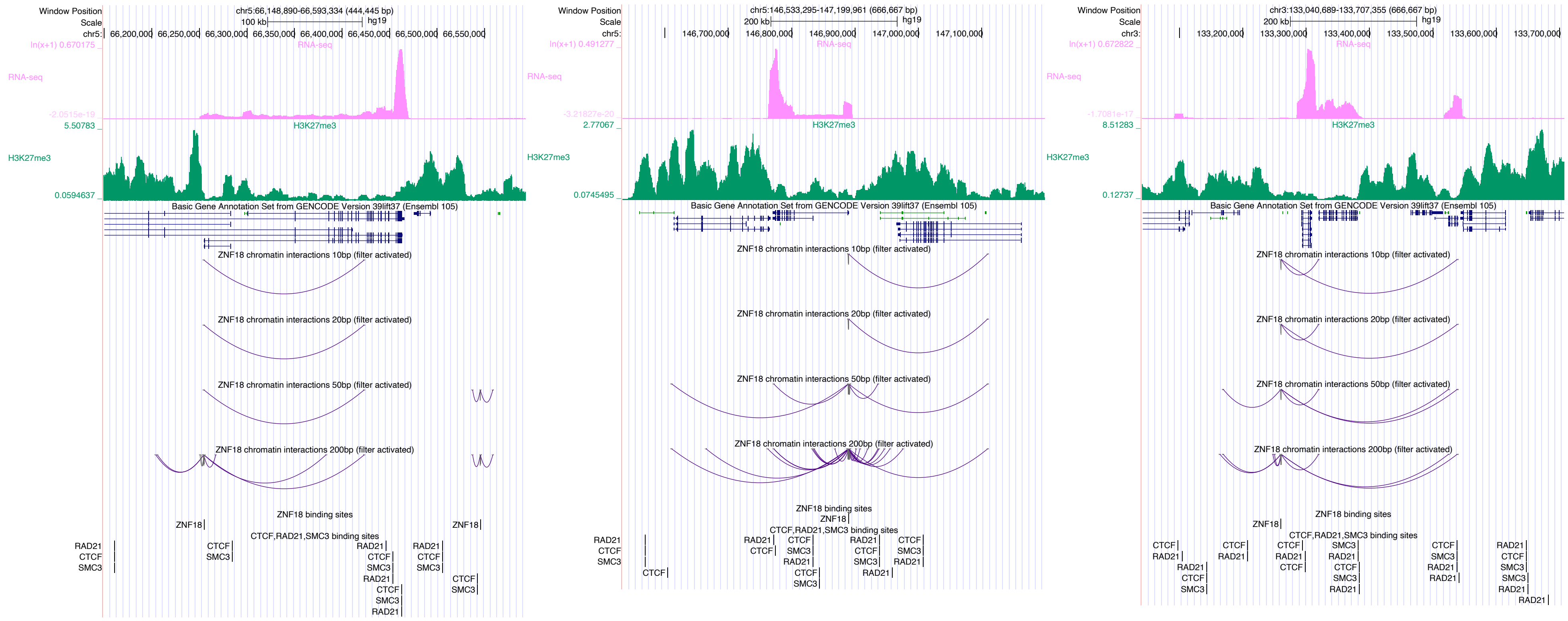
Window Position
Scale
chr5:66,148,890-66,593,334 (444,445 bp)
100 kb
hg19
RNA-seq
H3K27me3
Basic Gene Annotation Set from GENCODE Version 39lift37 (Ensembl 105)
ZNF18 chromatin interactions 10bp (filter activated)
ZNF18 chromatin interactions 20bp (filter activated)
ZNF18 chromatin interactions 50bp (filter activated)
ZNF18 chromatin interactions 200bp (filter activated)
ZNF18 binding sites
CTCF,RAD21,SMC3 binding sites
chr5:146,533,295-147,199,961 (666,667 bp)
200 kb
chr3:133,040,689-133,707,355 (666,667 bp)

### ZNF35

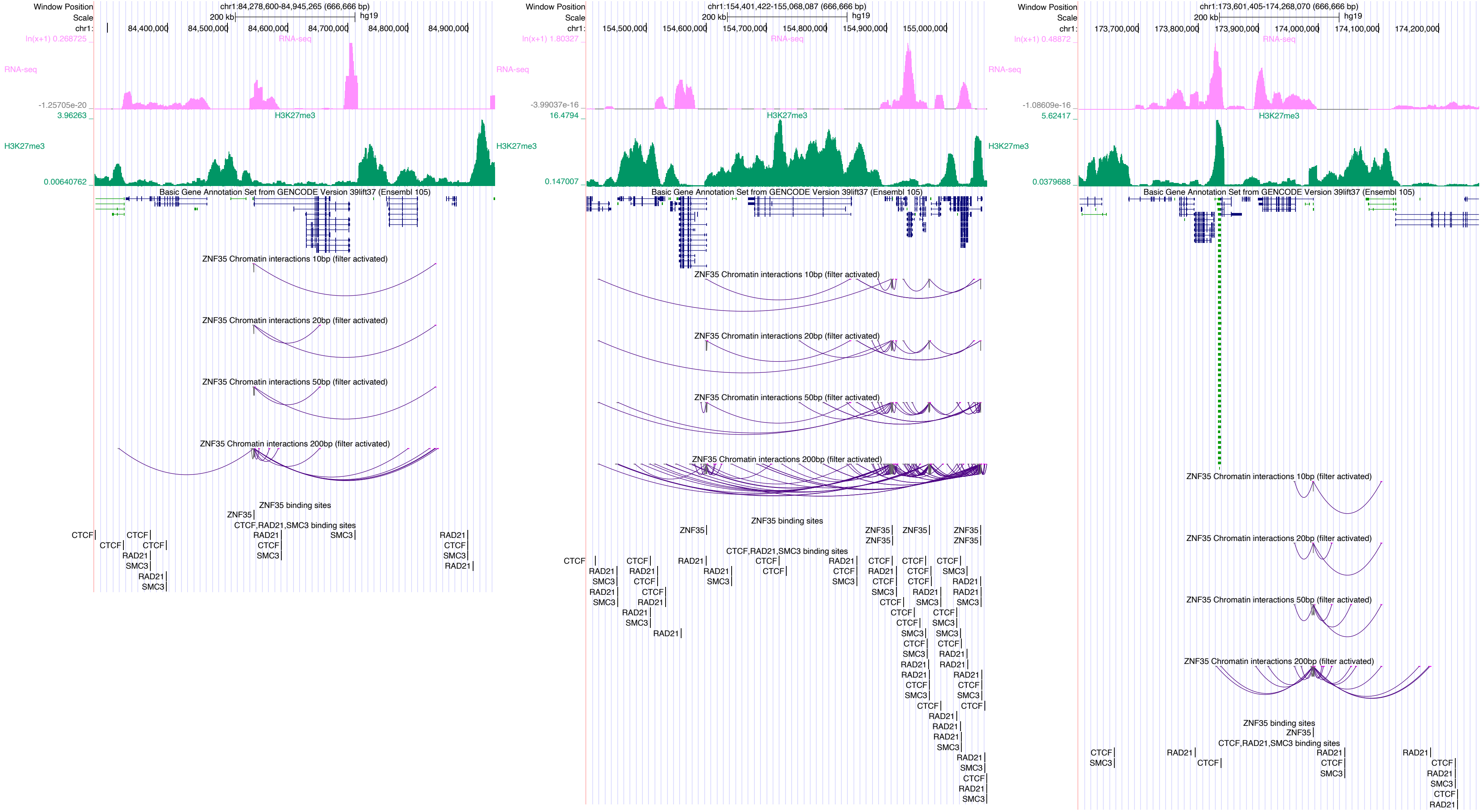

Window Position
Scale
chr1:
chr1:84,278,600-84,945,265 (666,666 bp)
200 kb
hg19
84,400,000
84,500,000
84,600,000
84,700,000
84,800,000
84,900,000
RNA-seq
H3K27me3
Basic Gene Annotation Set from GENCODE Version 39lift37 (Ensembl 105)
ZNF35 Chromatin interactions 10bp (filter activated)
ZNF35 Chromatin interactions 20bp (filter activated)
ZNF35 Chromatin interactions 50bp (filter activated)
ZNF35 Chromatin interactions 200bp (filter activated)
ZNF35 binding sites
CTCF,RAD21,SMC3 binding sites
chr1:154,401,422-155,068,087 (666,666 bp)
154,500,000
154,600,000
154,700,000
154,800,000
154,900,000
155,000,000
chr1:173,601,405-174,268,070 (666,666 bp)
173,700,000
173,800,000
173,900,000
174,000,000
174,100,000
174,200,000

### ZNF143

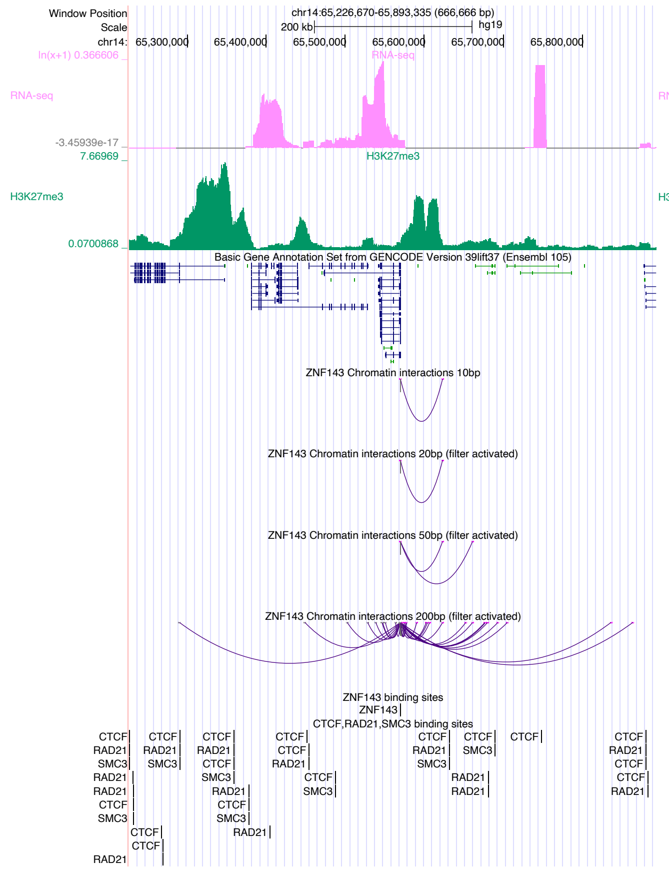
Window Position
chr14:65,226,670-65,893,335 (666,666 bp)
Scale
200 kb
hg19
chr14: 65,300,000 65,400,000 65,500,000 65,600,000 65,700,000 65,800,000
RNA-seq
H3K27me3
Basic Gene Annotation Set from GENCODE Version 39lift37 (Ensembl 105)
ZNF143 Chromatin interactions 10bp
ZNF143 Chromatin interactions 20bp (filter activated)
ZNF143 Chromatin interactions 50bp (filter activated)
ZNF143 Chromatin interactions 200bp (filter activated)
ZNF143 binding sites
CTCF,RAD21,SMC3 binding sites

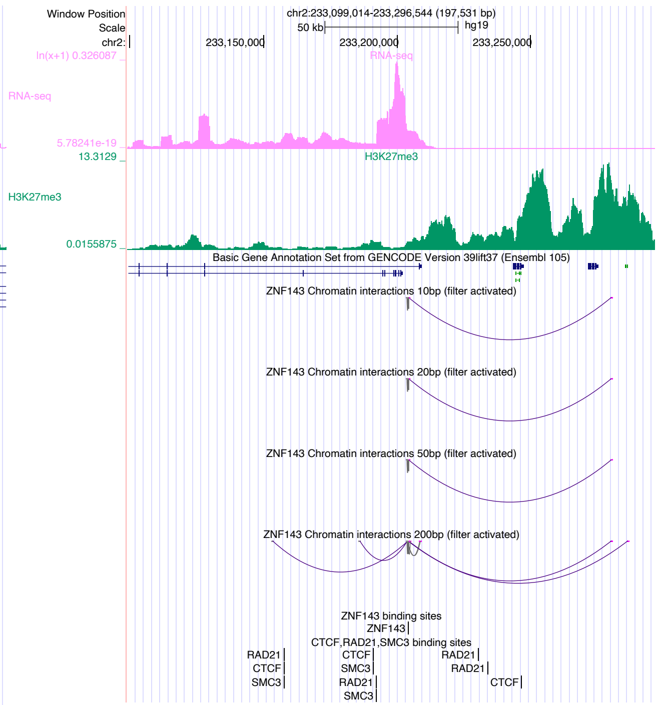
Window Position
chr2:233,099,014-233,296,544 (197,531 bp)
Scale
50 kb
hg19
chr2: 233,150,000 233,200,000 233,250,000
RNA-seq
H3K27me3
Basic Gene Annotation Set from GENCODE Version 39lift37 (Ensembl 105)
ZNF143 Chromatin interactions 10bp (filter activated)
ZNF143 Chromatin interactions 20bp (filter activated)
ZNF143 Chromatin interactions 50bp (filter activated)
ZNF143 Chromatin interactions 200bp (filter activated)
ZNF143 binding sites
CTCF,RAD21,SMC3 binding sites

### ZNF320

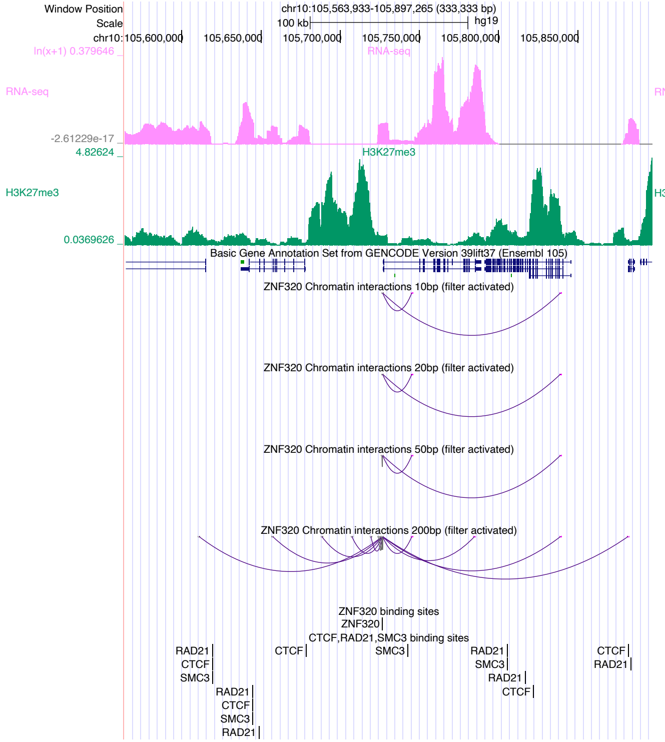
Window Position
chr10:105,563,933-105,897,265 (333,333 bp)
Scale
100 kb
hg19
chr10:105,600,000
105,650,000
105,700,000
105,750,000
105,800,000
105,850,000
ln(x+1) 0.379646
RNA-seq
-2.61229e-17
4.82624
H3K27me3
0.0369626
Basic Gene Annotation Set from GENCODE Version 39lift37 (Ensembl 105)
ZNF320 Chromatin interactions 10bp (filter activated)
ZNF320 Chromatin interactions 20bp (filter activated)
ZNF320 Chromatin interactions 50bp (filter activated)
ZNF320 Chromatin interactions 200bp (filter activated)
ZNF320 binding sites
ZNF320
CTCF,RAD21,SMC3 binding sites
RAD21
CTCF
SMC3

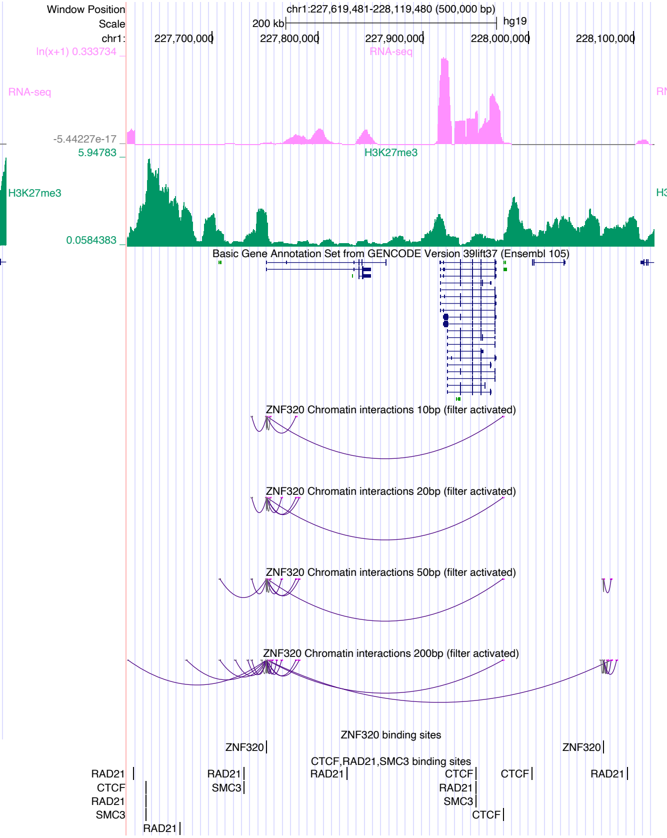
Window Position
chr1:227,619,481-228,119,480 (500,000 bp)
Scale
200 kb
hg19
chr1:
227,700,000
227,800,000
227,900,000
228,000,000
228,100,000
ln(x+1) 0.333734
RNA-seq
-5.44227e-17
5.94783
H3K27me3
0.0584383
Basic Gene Annotation Set from GENCODE Version 39lift37 (Ensembl 105)
ZNF320 Chromatin interactions 10bp (filter activated)
ZNF320 Chromatin interactions 20bp (filter activated)
ZNF320 Chromatin interactions 50bp (filter activated)
ZNF320 Chromatin interactions 200bp (filter activated)
ZNF320 binding sites
ZNF320
CTCF,RAD21,SMC3 binding sites
RAD21
CTCF
SMC3

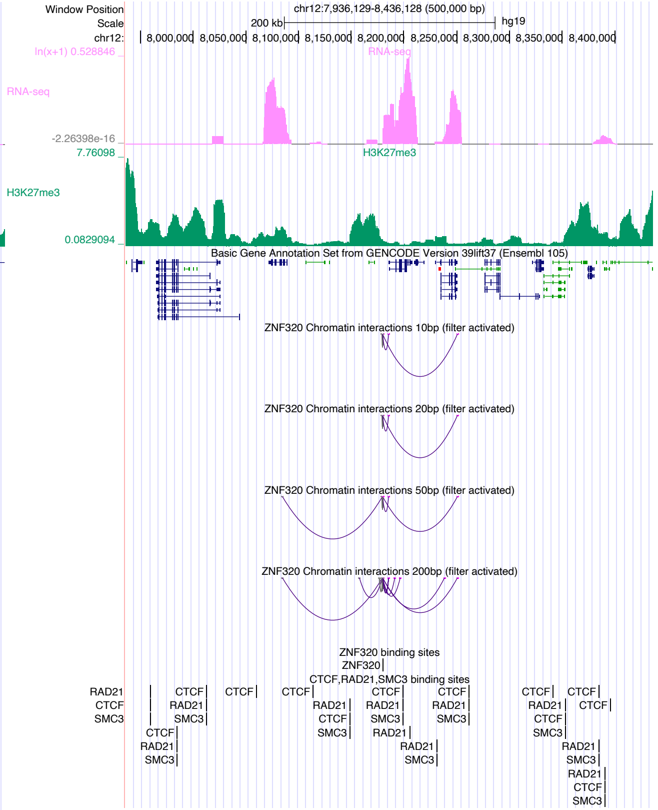
Window Position
chr12:7,936,129-8,436,128 (500,000 bp)
Scale
200 kb
hg19
chr12:
8,000,000
8,050,000
8,100,000
8,150,000
8,200,000
8,250,000
8,300,000
8,350,000
8,400,000
ln(x+1) 0.528846
RNA-seq
-2.26398e-16
7.76098
H3K27me3
0.0829094
Basic Gene Annotation Set from GENCODE Version 39lift37 (Ensembl 105)
ZNF320 Chromatin interactions 10bp (filter activated)
ZNF320 Chromatin interactions 20bp (filter activated)
ZNF320 Chromatin interactions 50bp (filter activated)
ZNF320 Chromatin interactions 200bp (filter activated)
ZNF320 binding sites
ZNF320
CTCF,RAD21,SMC3 binding sites
RAD21
CTCF
SMC3

### ZNF329

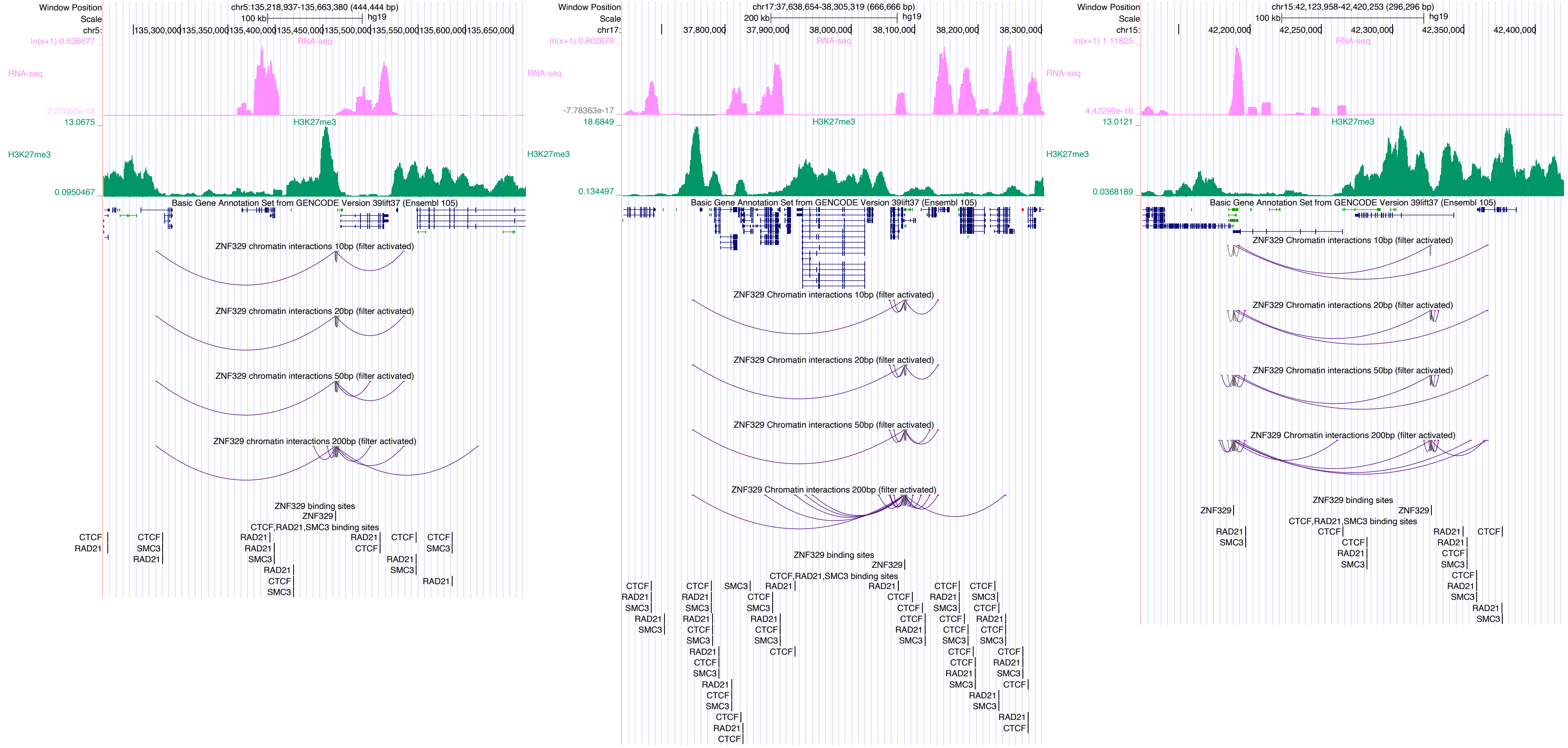

Window Position
chr5:135,218,937-135,663,380 (444,444 bp)
Scale
100 kb
hg19
chr5:
135,300,000
135,350,000
135,400,000
135,450,000
135,500,000
135,550,000
135,600,000
135,650,000
ln(x+1) 0.636677
RNA-seq
-2.27668e-16
13.0675
H3K27me3
0.0950467
Basic Gene Annotation Set from GENCODE Version 39lift37 (Ensembl 105)
ZNF329 chromatin interactions 10bp (filter activated)
ZNF329 chromatin interactions 20bp (filter activated)
ZNF329 chromatin interactions 50bp (filter activated)
ZNF329 chromatin interactions 200bp (filter activated)
ZNF329 binding sites
ZNF329
CTCF,RAD21,SMC3 binding sites
CTCF
RAD21
SMC3
Window Position
chr17:37,638,654-38,305,319 (666,666 bp)
200 kb
hg19
chr17:
37,800,000
37,900,000
38,000,000
38,100,000
38,200,000
38,300,000
ln(x+1) 0.802679
RNA-seq
-7.78363e-17
18.6849
H3K27me3
0.134497
Basic Gene Annotation Set from GENCODE Version 39lift37 (Ensembl 105)
ZNF329 Chromatin interactions 10bp (filter activated)
ZNF329 Chromatin interactions 20bp (filter activated)
ZNF329 Chromatin interactions 50bp (filter activated)
ZNF329 Chromatin interactions 200bp (filter activated)
ZNF329 binding sites
ZNF329
CTCF,RAD21,SMC3 binding sites
Window Position
chr15:42,123,958-42,420,253 (296,296 bp)
100 kb
hg19
chr15:
42,200,000
42,250,000
42,300,000
42,350,000
42,400,000
ln(x+1) 1.11825
RNA-seq
4.42299e-16
13.0121
H3K27me3
0.0368189
Basic Gene Annotation Set from GENCODE Version 39lift37 (Ensembl 105)
ZNF329 Chromatin interactions 10bp (filter activated)
ZNF329 Chromatin interactions 20bp (filter activated)
ZNF329 Chromatin interactions 50bp (filter activated)
ZNF329 Chromatin interactions 200bp (filter activated)
ZNF329 binding sites
ZNF329
CTCF,RAD21,SMC3 binding sites

### ZNF449

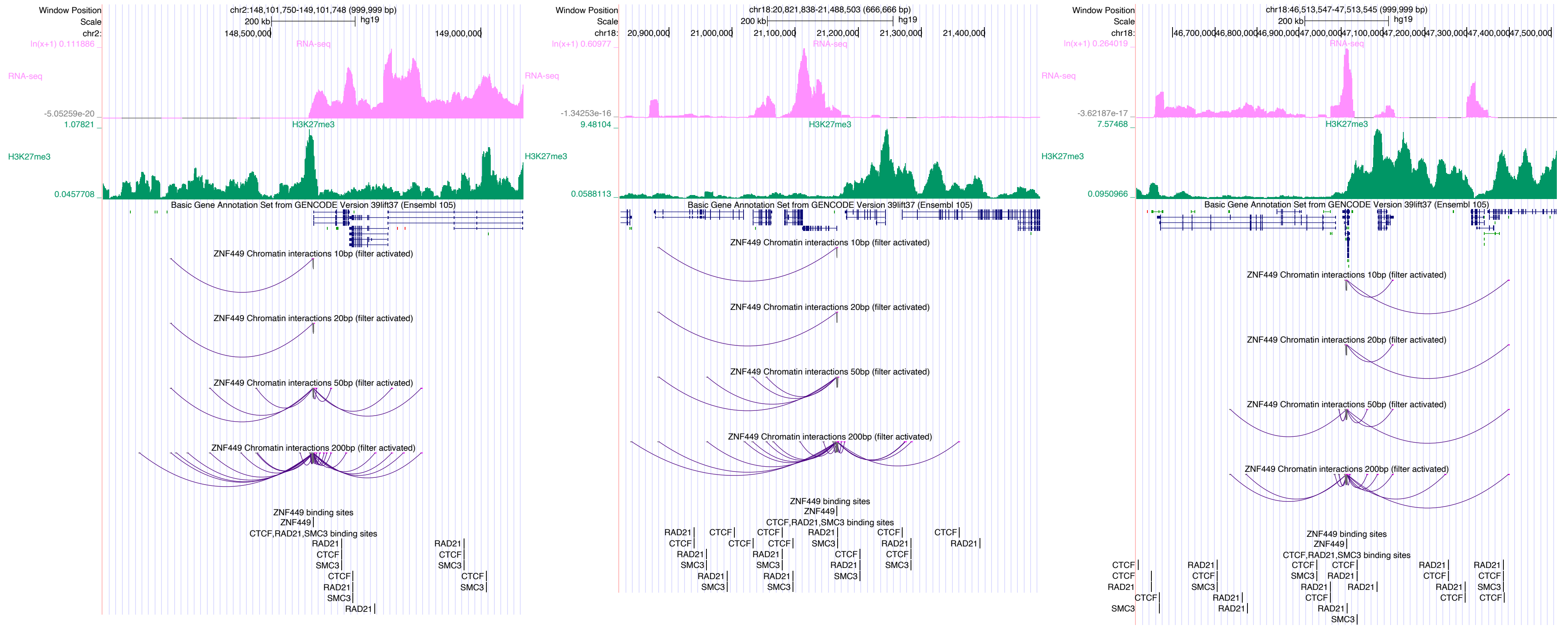

Window Position
chr2:148,101,750-149,101,748 (999,999 bp)
Scale
200 kb
hg19
chr2:
148,500,000
149,000,000
RNA-seq
H3K27me3
Basic Gene Annotation Set from GENCODE Version 39lift37 (Ensembl 105)
ZNF449 Chromatin interactions 10bp (filter activated)
ZNF449 Chromatin interactions 20bp (filter activated)
ZNF449 Chromatin interactions 50bp (filter activated)
ZNF449 Chromatin interactions 200bp (filter activated)
ZNF449 binding sites
CTCF,RAD21,SMC3 binding sites
chr18:20,821,838-21,488,503 (666,666 bp)
chr18:
20,900,000
21,000,000
21,100,000
21,200,000
21,300,000
21,400,000
chr18:46,513,547-47,513,545 (999,999 bp)
46,700,000
46,800,000
46,900,000
47,000,000
47,100,000
47,200,000
47,300,000
47,400,000
47,500,000

### ZZZ3

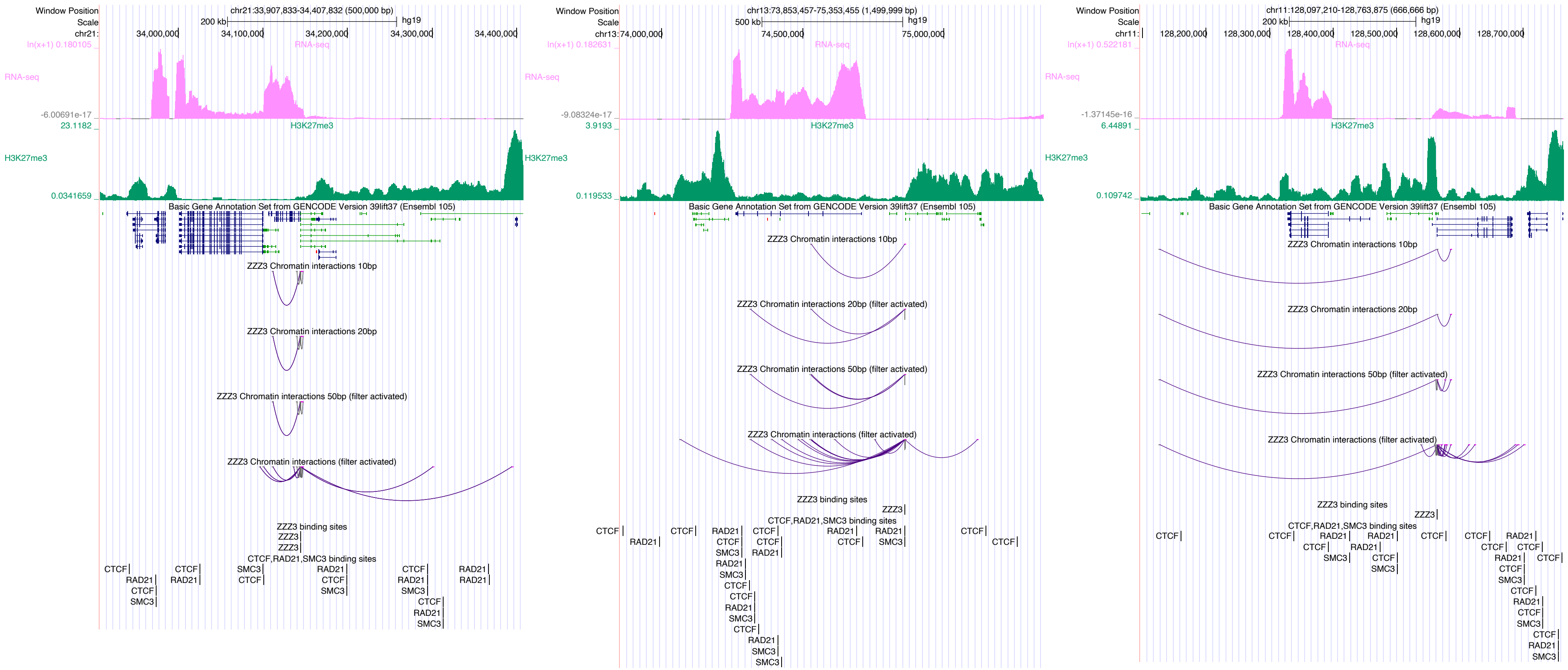
Window Position
Scale
chr21:33,907,833-34,407,832 (500,000 bp)
200 kb
hg19
chr21:
34,000,000
34,100,000
34,200,000
34,300,000
34,400,000
ln(x+1) 0.180105
RNA-seq
-6.00691e-17
23.1182
H3K27me3
0.0341659
Basic Gene Annotation Set from GENCODE Version 39lift37 (Ensembl 105)
ZZZ3 Chromatin interactions 10bp
ZZZ3 Chromatin interactions 20bp
ZZZ3 Chromatin interactions 50bp (filter activated)
ZZZ3 Chromatin interactions (filter activated)
ZZZ3 binding sites
ZZZ3
CTCF,RAD21,SMC3 binding sites
CTCF
RAD21
SMC3
chr13:73,853,457-75,353,455 (1,499,999 bp)
500 kb
chr13:74,000,000
74,500,000
75,000,000
ln(x+1) 0.182631
-9.08324e-17
3.9193
0.119533
ZZZ3 Chromatin interactions 20bp (filter activated)
chr11:128,097,210-128,763,875 (666,666 bp)
chr11:
128,200,000
128,300,000
128,400,000
128,500,000
128,600,000
128,700,000
ln(x+1) 0.522181
-1.37145e-16
6.44891
0.109742
