## Supplementary material for "Systematic discovery of directional regulatory motifs associated with human insulator sites": Supplementary_Fig4.pdf

a

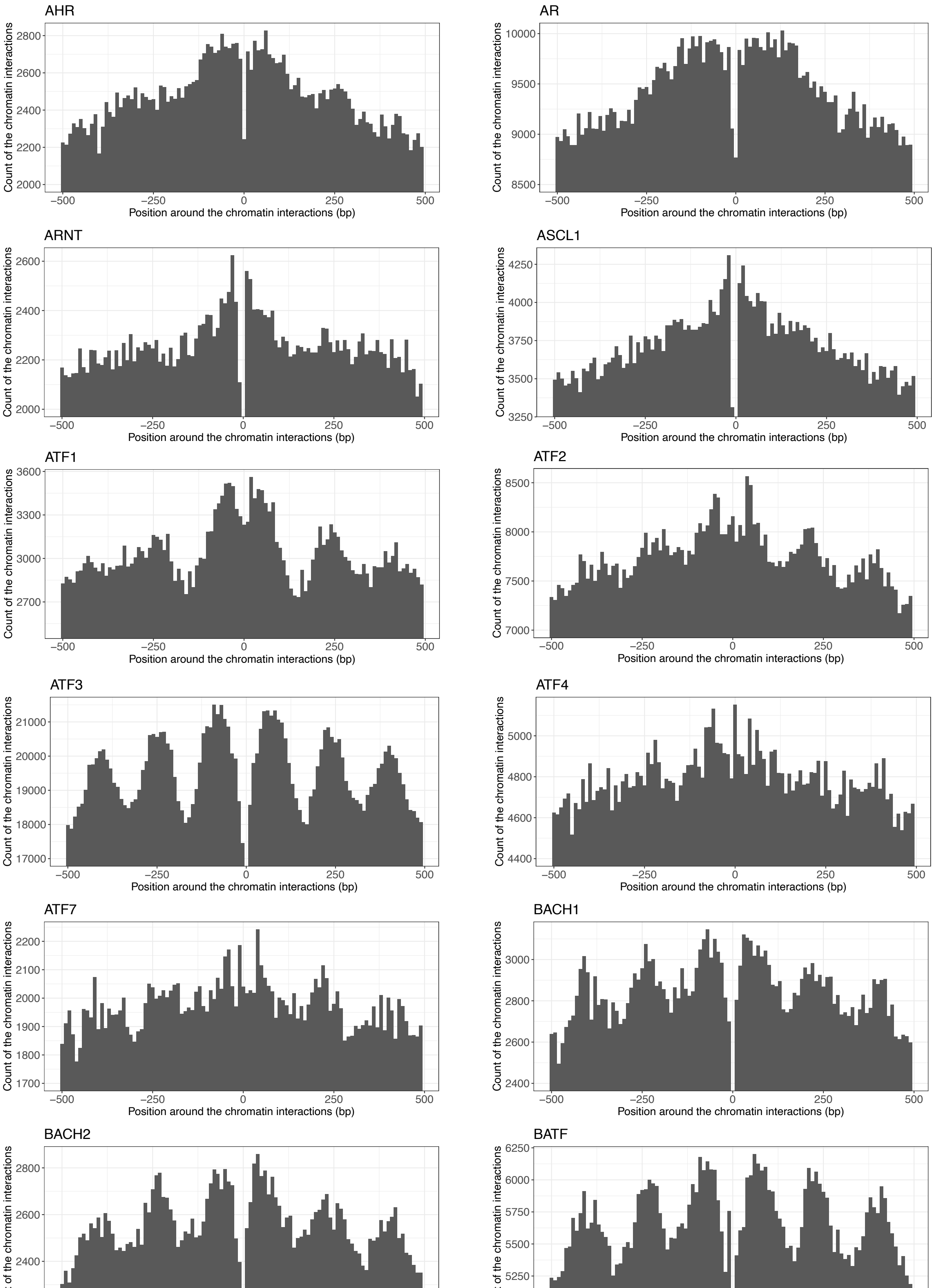


Supplementary Fig. 4. Distribution of DNA-binding sites not overlapping with CTCF, RAD21, or SMC3 around chromatin interaction sites. **a** 240 DBPs, each with at least 1,000 DNA-binding sites identified from open chromatin regions in HFF cells. **b** Two DBPs with fewer than 1,000 DNA-binding sites identified from open chromatin regions in HFF cells. **c** The distribution patterns of DNA-binding sites for "CTCF, RAD21, and SMC3" were similar to those for "BACH2, FOS, ATF3, NFE2, and MAFK".

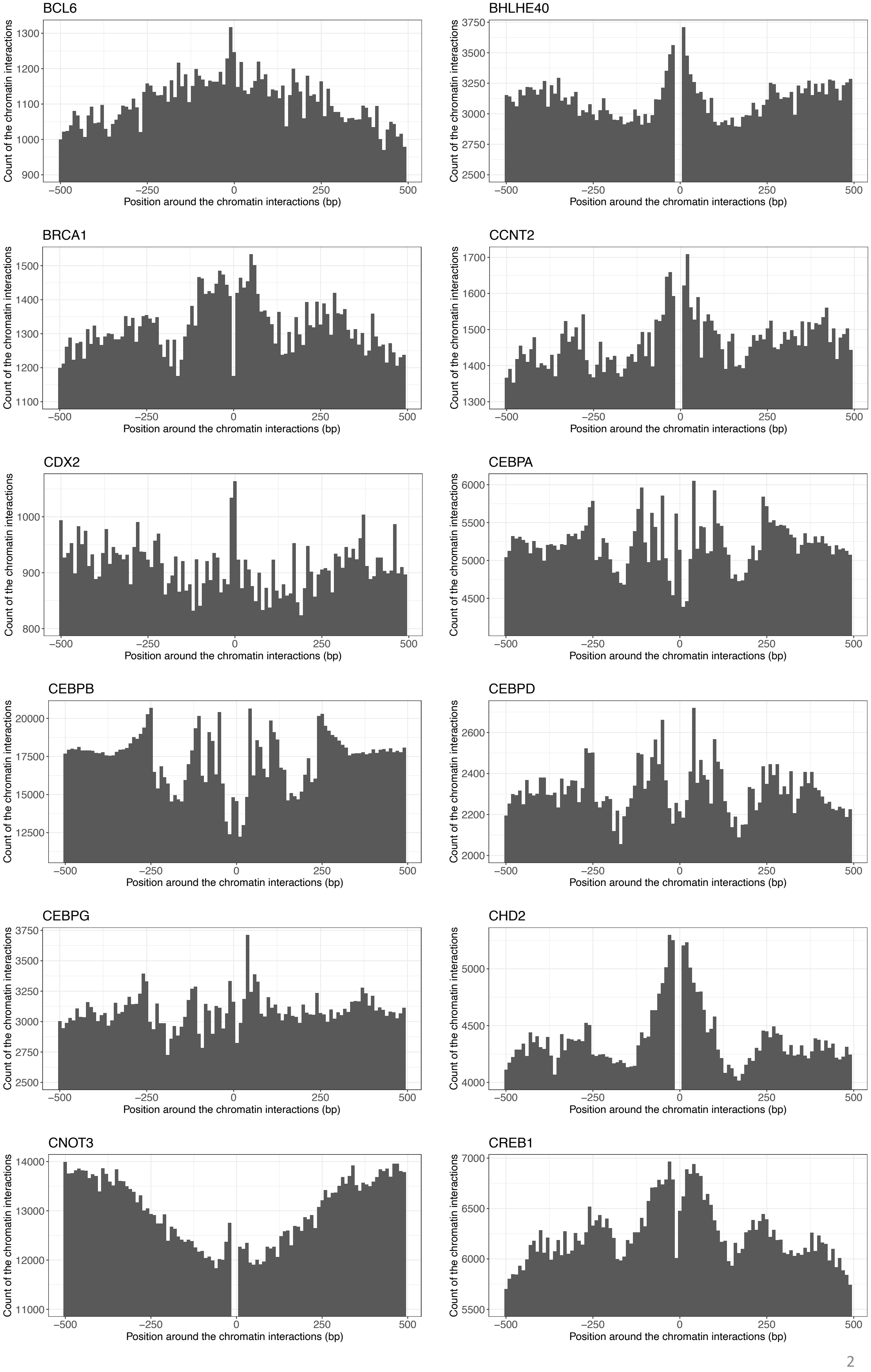
BCL6
Count of the chromatin interactions
900
1000
1100
1200
1300
−500
−250
0
250
500
Position around the chromatin interactions (bp)
BHLHE40
Count of the chromatin interactions
2500
2750
3000
3250
3500
3750
−500
−250
0
250
500
Position around the chromatin interactions (bp)
BRCA1
Count of the chromatin interactions
1100
1200
1300
1400
1500
−500
−250
0
250
500
Position around the chromatin interactions (bp)
CCNT2
Count of the chromatin interactions
1300
1400
1500
1600
1700
−500
−250
0
250
500
Position around the chromatin interactions (bp)
CDX2
Count of the chromatin interactions
800
900
1000
−500
−250
0
250
500
Position around the chromatin interactions (bp)
CEBPA
Count of the chromatin interactions
4500
5000
5500
6000
−500
−250
0
250
500
Position around the chromatin interactions (bp)
CEBPB
Count of the chromatin interactions
12500
15000
17500
20000
−500
−250
0
250
500
Position around the chromatin interactions (bp)
CEBPD
Count of the chromatin interactions
2000
2200
2400
2600
−500
−250
0
250
500
Position around the chromatin interactions (bp)
CEBPG
Count of the chromatin interactions
2500
2750
3000
3250
3500
3750
−500
−250
0
250
500
Position around the chromatin interactions (bp)
CHD2
Count of the chromatin interactions
4000
4500
5000
−500
−250
0
250
500
Position around the chromatin interactions (bp)
CNOT3
Count of the chromatin interactions
11000
12000
13000
14000
−500
−250
0
250
500
Position around the chromatin interactions (bp)
CREB1
Count of the chromatin interactions
5500
6000
6500
7000
−500
−250
0
250
500
Position around the chromatin interactions (bp)

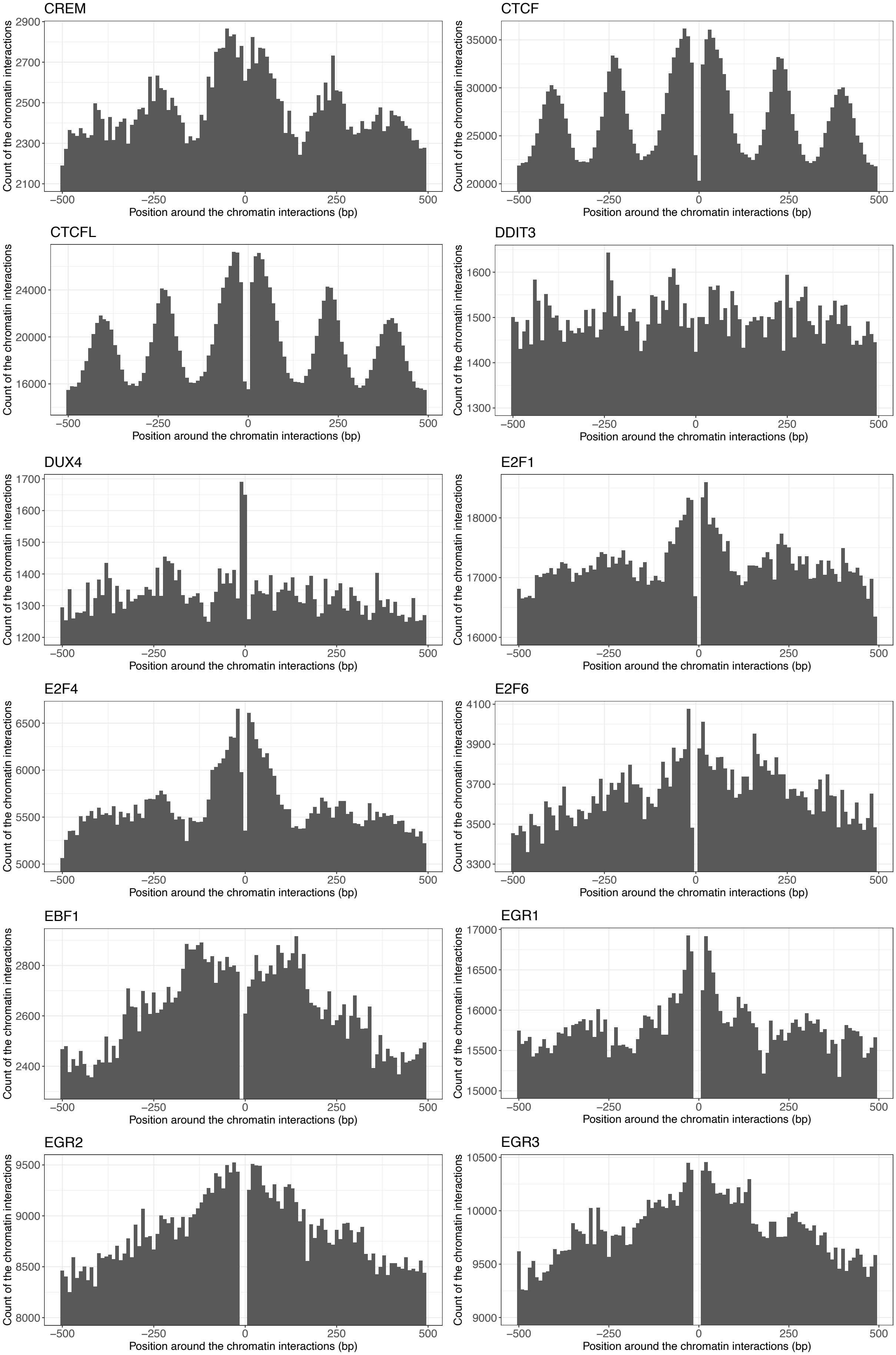
CREM
CTCF
CTCFL
DDIT3
DUX4
E2F1
E2F4
E2F6
EBF1
EGR1
EGR2
EGR3
Count of the chromatin interactions
Position around the chromatin interactions (bp)
−500
−250
0
250
500

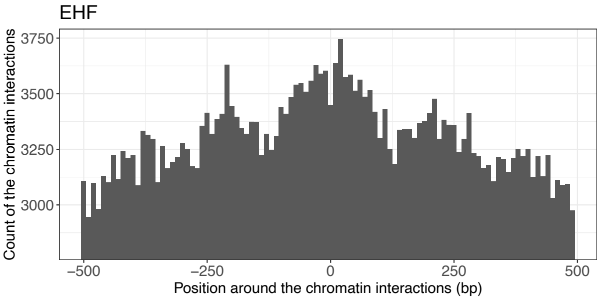
EHF
Count of the chromatin interactions
3750
3500
3250
3000
−500
−250
0
250
500
Position around the chromatin interactions (bp)

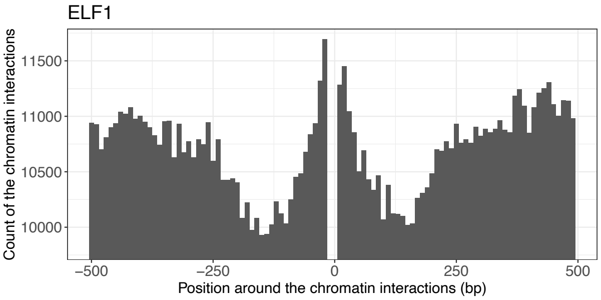
ELF1
Count of the chromatin interactions
11500
11000
10500
10000
−500
−250
0
250
500
Position around the chromatin interactions (bp)

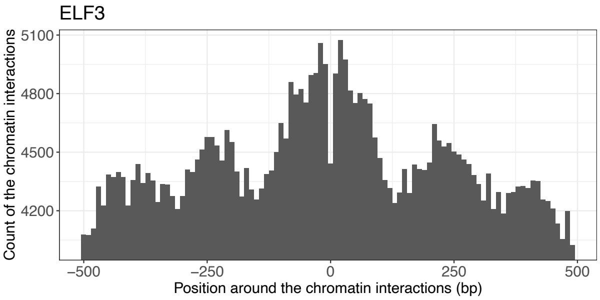
ELF3
Count of the chromatin interactions
5100
4800
4500
4200
−500
−250
0
250
500
Position around the chromatin interactions (bp)

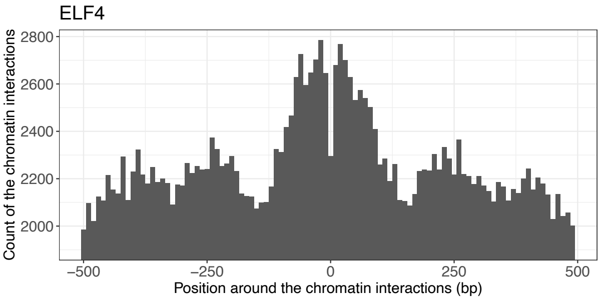
ELF4
Count of the chromatin interactions
2800
2600
2400
2200
2000
−500
−250
0
250
500
Position around the chromatin interactions (bp)

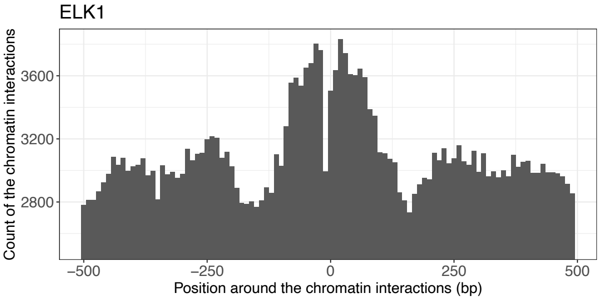
ELK1
Count of the chromatin interactions
3600
3200
2800
−500
−250
0
250
500
Position around the chromatin interactions (bp)

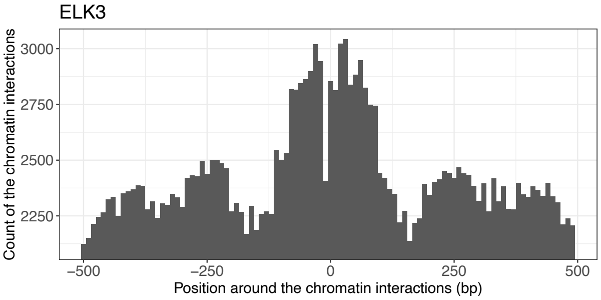
ELK3
Count of the chromatin interactions
3000
2750
2500
2250
−500
−250
0
250
500
Position around the chromatin interactions (bp)

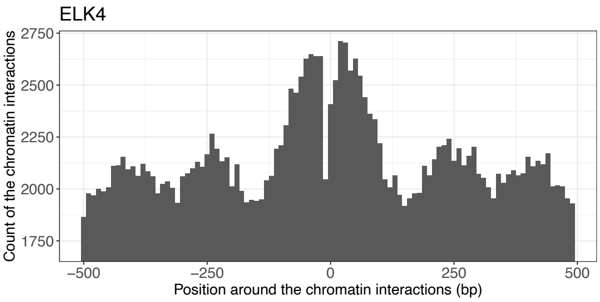
ELK4
Count of the chromatin interactions
2750
2500
2250
2000
1750
−500
−250
0
250
500
Position around the chromatin interactions (bp)

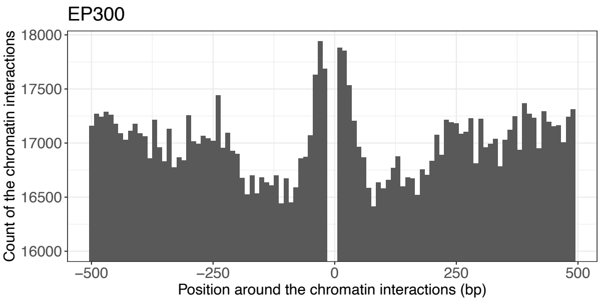
EP300
Count of the chromatin interactions
18000
17500
17000
16500
16000
−500
−250
0
250
500
Position around the chromatin interactions (bp)

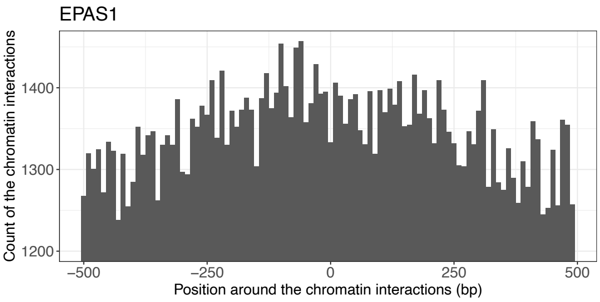
EPAS1
Count of the chromatin interactions
1400
1300
1200
−500
−250
0
250
500
Position around the chromatin interactions (bp)

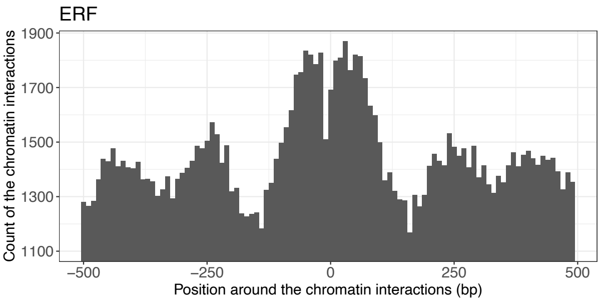
ERF
Count of the chromatin interactions
1900
1700
1500
1300
1100
−500
−250
0
250
500
Position around the chromatin interactions (bp)

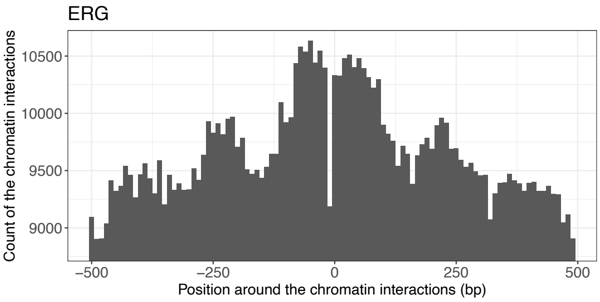
ERG
Count of the chromatin interactions
10500
10000
9500
9000
−500
−250
0
250
500
Position around the chromatin interactions (bp)

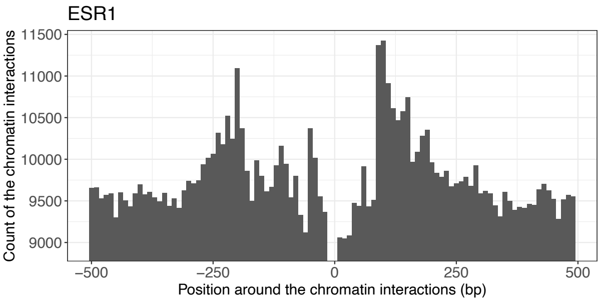
ESR1
Count of the chromatin interactions
11500
11000
10500
10000
9500
9000
−500
−250
0
250
500
Position around the chromatin interactions (bp)

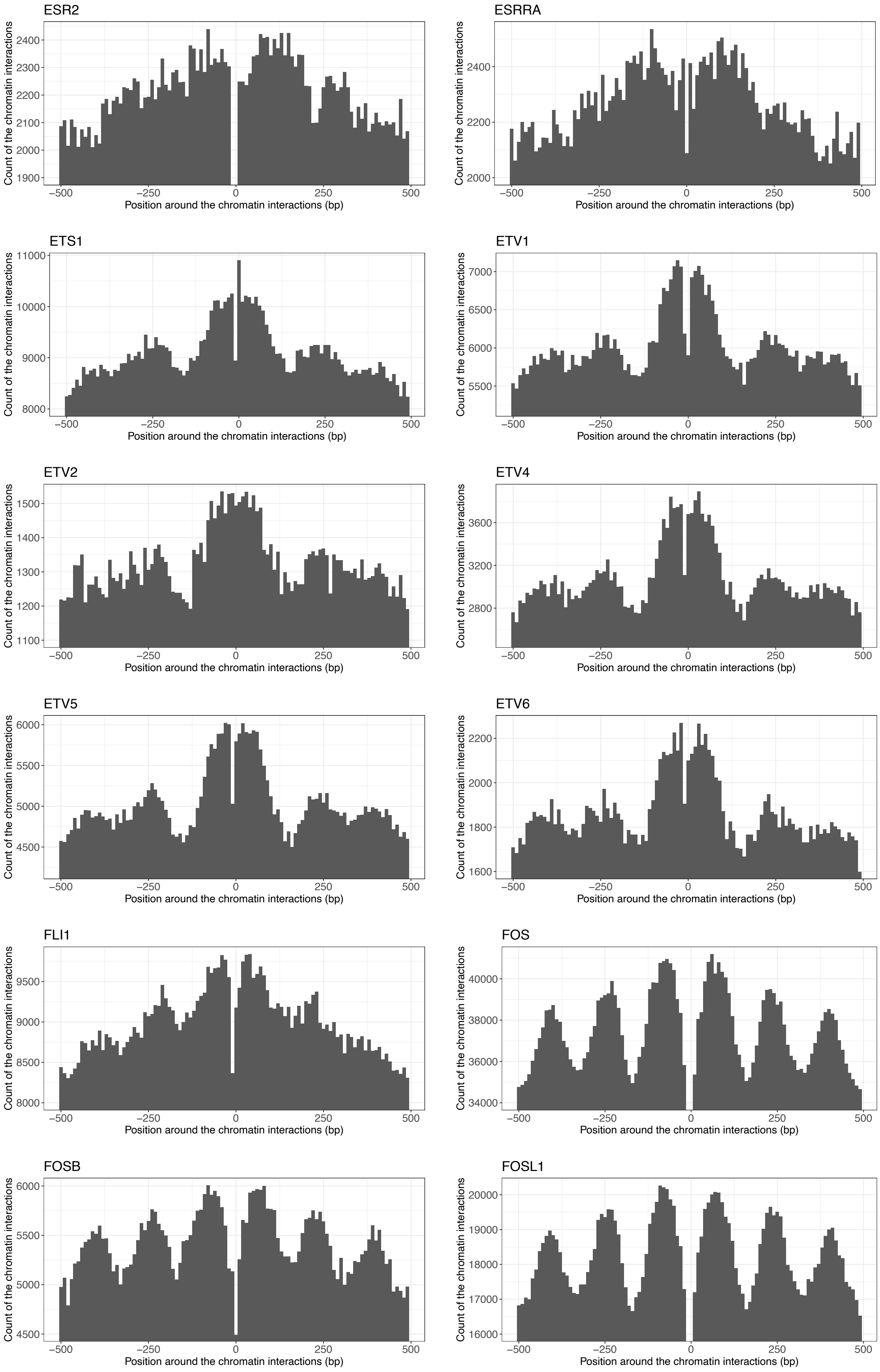
ESR2
ESRRA
ETS1
ETV1
ETV2
ETV4
ETV5
ETV6
FLI1
FOS
FOSB
FOSL1
Count of the chromatin interactions
Position around the chromatin interactions (bp)

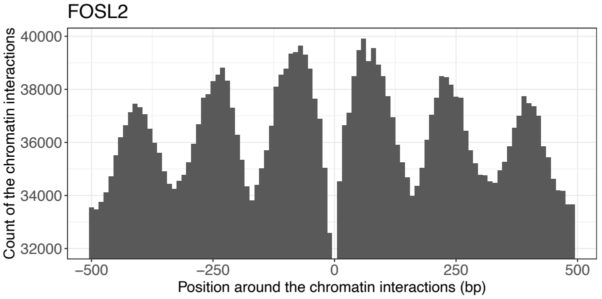
FOSL2
Count of the chromatin interactions
40000
38000
36000
34000
32000
−500
−250
0
250
500
Position around the chromatin interactions (bp)

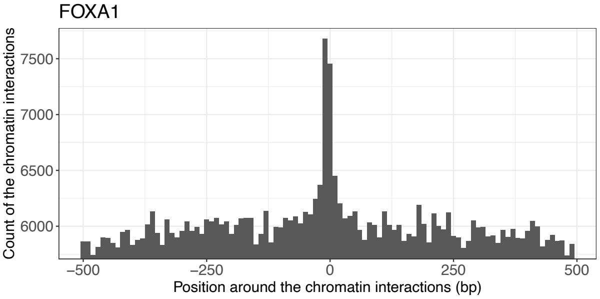
FOXA1
Count of the chromatin interactions
7500
7000
6500
6000
−500
−250
0
250
500
Position around the chromatin interactions (bp)

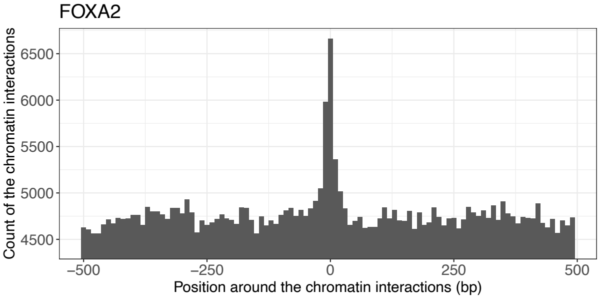
FOXA2
Count of the chromatin interactions
6500
6000
5500
5000
4500
−500
−250
0
250
500
Position around the chromatin interactions (bp)

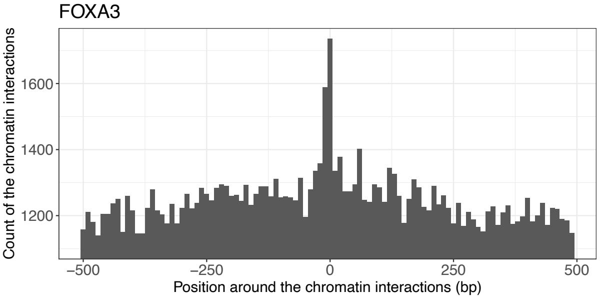
FOXA3
Count of the chromatin interactions
1600
1400
1200
−500
−250
0
250
500
Position around the chromatin interactions (bp)

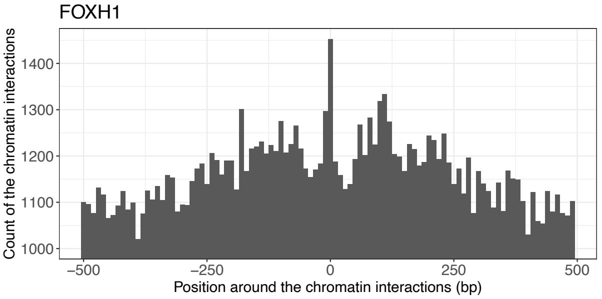
FOXH1
Count of the chromatin interactions
1400
1300
1200
1100
1000
−500
−250
0
250
500
Position around the chromatin interactions (bp)

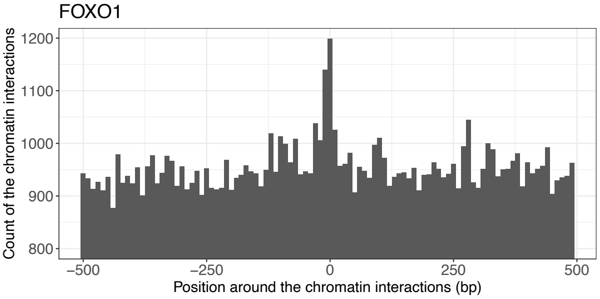
FOXO1
Count of the chromatin interactions
1200
1100
1000
900
800
−500
−250
0
250
500
Position around the chromatin interactions (bp)

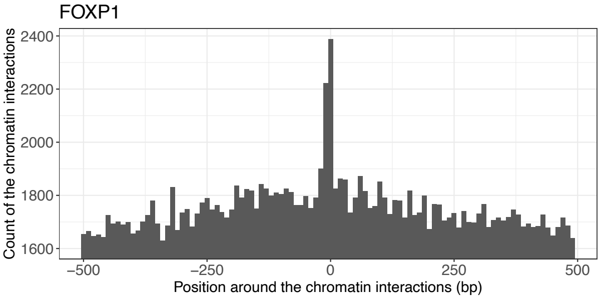
FOXP1
Count of the chromatin interactions
2400
2200
2000
1800
1600
−500
−250
0
250
500
Position around the chromatin interactions (bp)

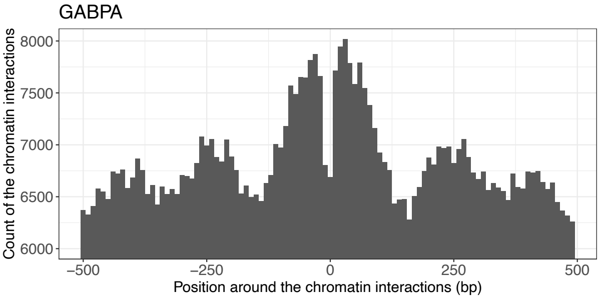
GABPA
Count of the chromatin interactions
8000
7500
7000
6500
6000
−500
−250
0
250
500
Position around the chromatin interactions (bp)

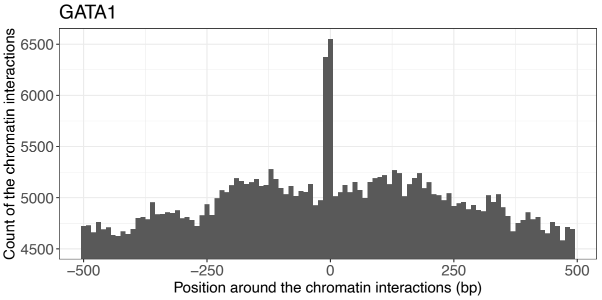
GATA1
Count of the chromatin interactions
6500
6000
5500
5000
4500
−500
−250
0
250
500
Position around the chromatin interactions (bp)

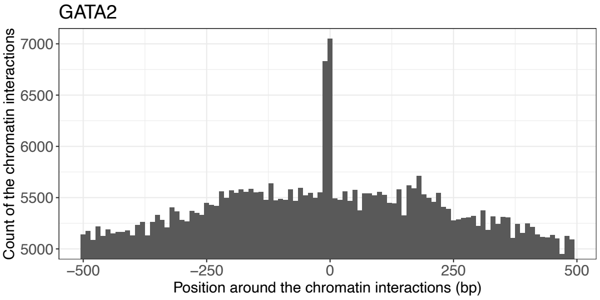
GATA2
Count of the chromatin interactions
7000
6500
6000
5500
5000
−500
−250
0
250
500
Position around the chromatin interactions (bp)

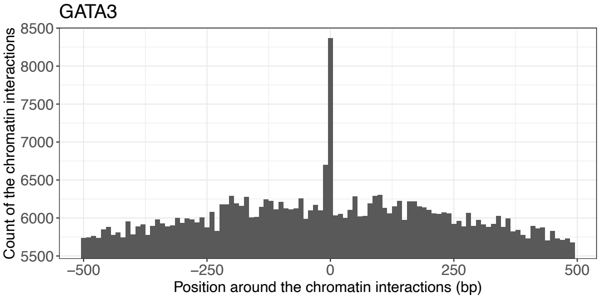
GATA3
Count of the chromatin interactions
8500
8000
7500
7000
6500
6000
5500
−500
−250
0
250
500
Position around the chromatin interactions (bp)

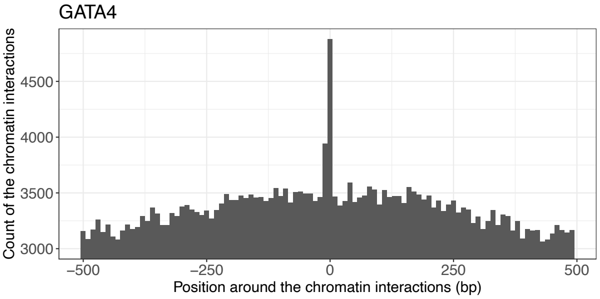
GATA4
Count of the chromatin interactions
4500
4000
3500
3000
−500
−250
0
250
500
Position around the chromatin interactions (bp)

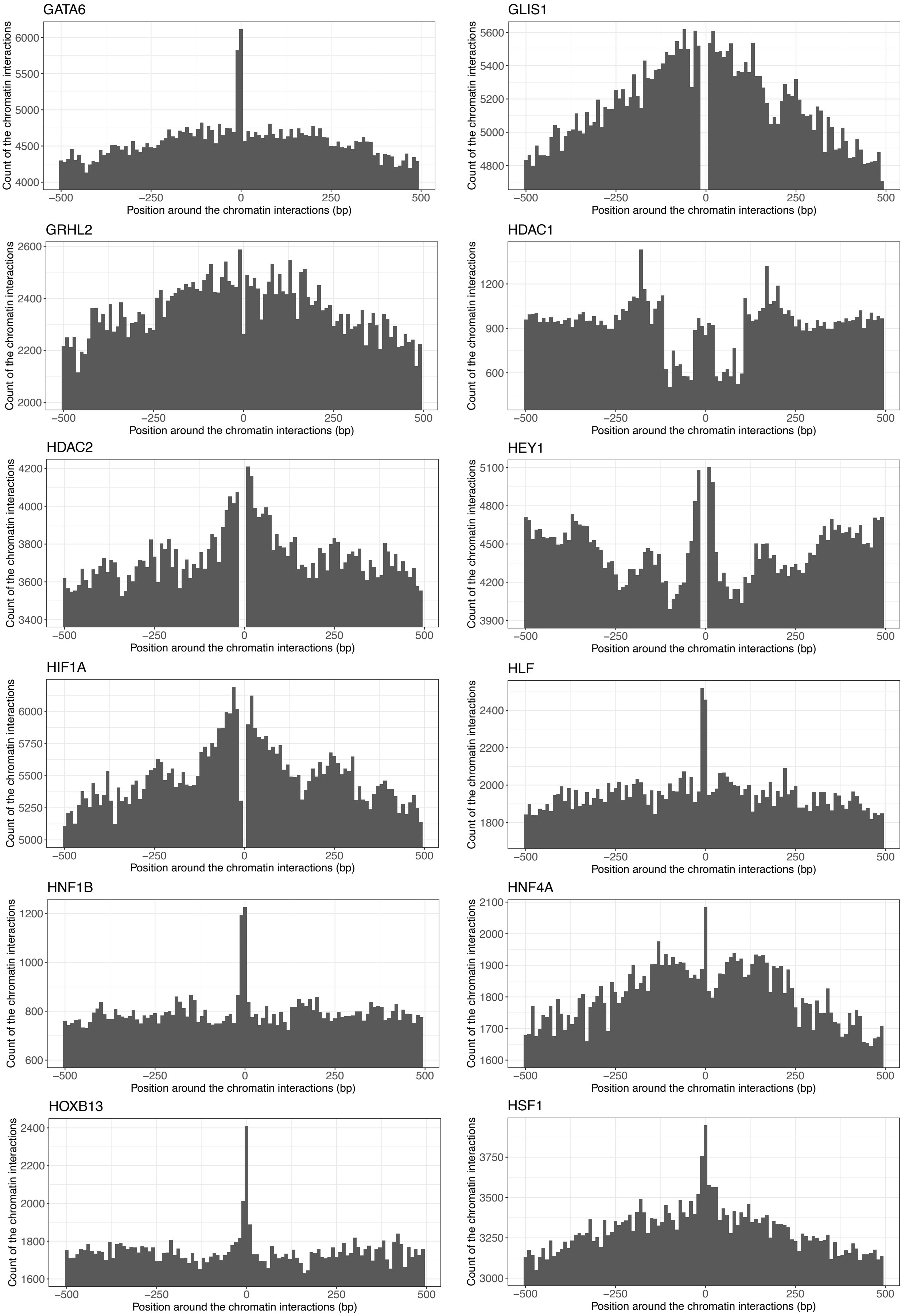
GATA6
GLIS1
GRHL2
HDAC1
HDAC2
HEY1
HIF1A
HLF
HNF1B
HNF4A
HOXB13
HSF1
Count of the chromatin interactions
Position around the chromatin interactions (bp)

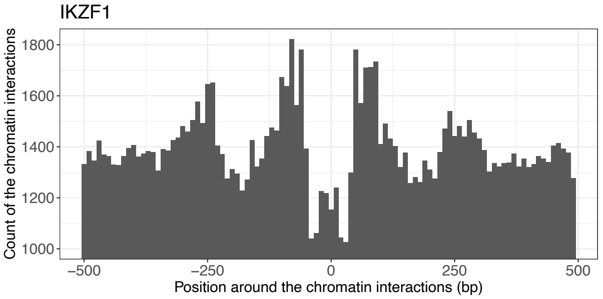
IKZF1
Count of the chromatin interactions
1800
1600
1400
1200
1000
−500
−250
0
250
500
Position around the chromatin interactions (bp)

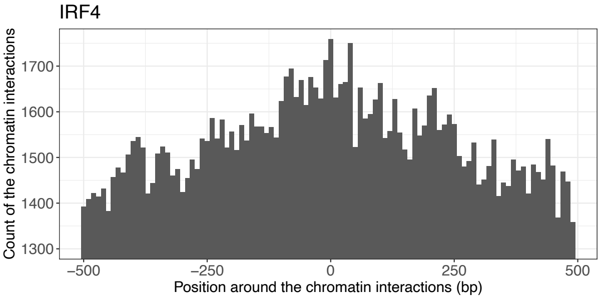
IRF4
Count of the chromatin interactions
1700
1600
1500
1400
1300
−500
−250
0
250
500
Position around the chromatin interactions (bp)

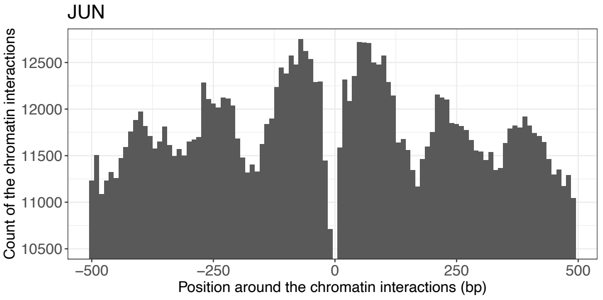
JUN
Count of the chromatin interactions
12500
12000
11500
11000
10500
−500
−250
0
250
500
Position around the chromatin interactions (bp)

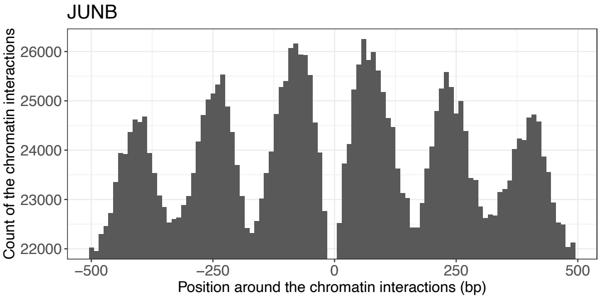
JUNB
Count of the chromatin interactions
26000
25000
24000
23000
22000
−500
−250
0
250
500
Position around the chromatin interactions (bp)

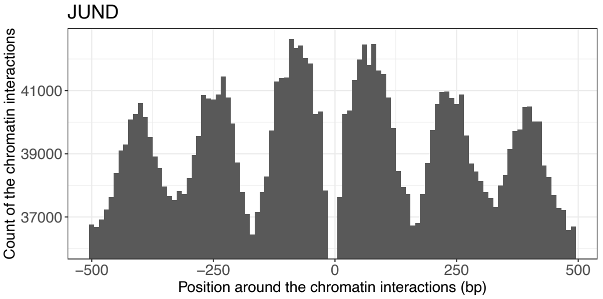
JUND
Count of the chromatin interactions
41000
39000
37000
−500
−250
0
250
500
Position around the chromatin interactions (bp)

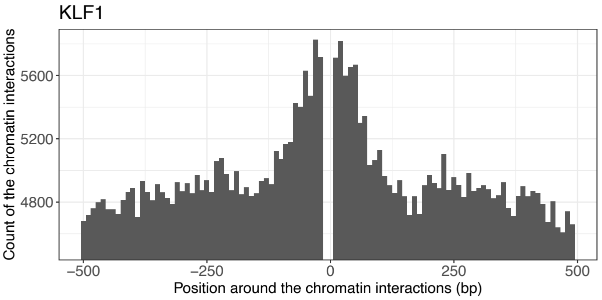
KLF1
Count of the chromatin interactions
5600
5200
4800
−500
−250
0
250
500
Position around the chromatin interactions (bp)

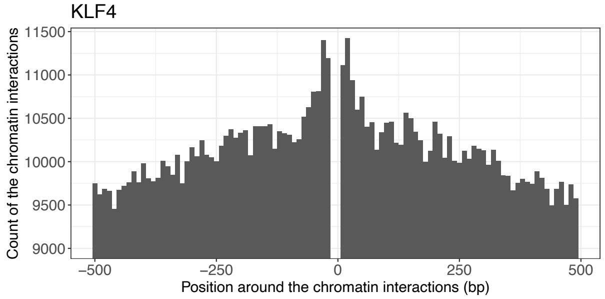
KLF4
Count of the chromatin interactions
11500
11000
10500
10000
9500
9000
−500
−250
0
250
500
Position around the chromatin interactions (bp)

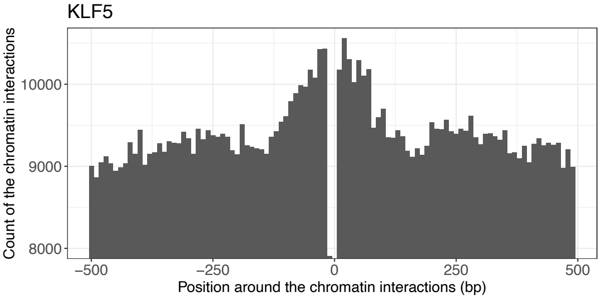
KLF5
Count of the chromatin interactions
10000
9000
8000
−500
−250
0
250
500
Position around the chromatin interactions (bp)

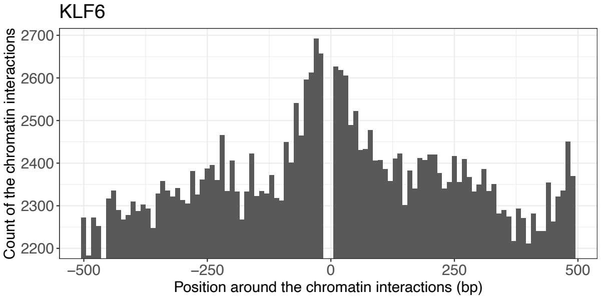
KLF6
Count of the chromatin interactions
2700
2600
2500
2400
2300
2200
−500
−250
0
250
500
Position around the chromatin interactions (bp)

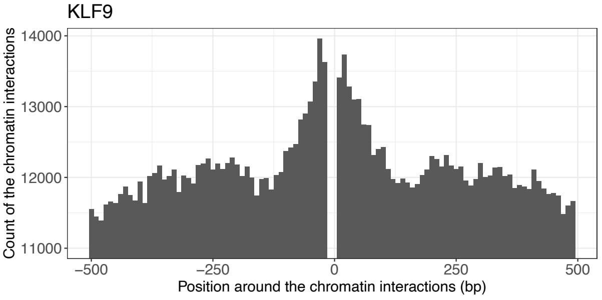
KLF9
Count of the chromatin interactions
14000
13000
12000
11000
−500
−250
0
250
500
Position around the chromatin interactions (bp)

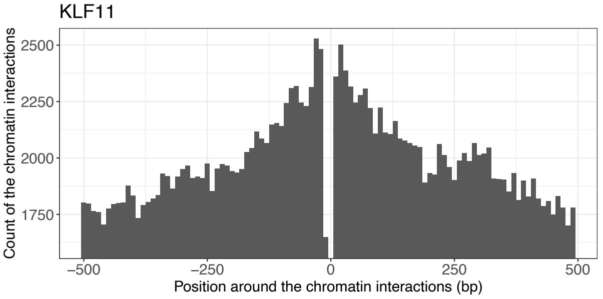
KLF11
Count of the chromatin interactions
2500
2250
2000
1750
−500
−250
0
250
500
Position around the chromatin interactions (bp)

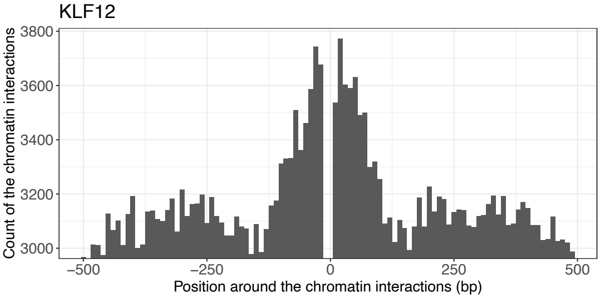
KLF12
Count of the chromatin interactions
3800
3600
3400
3200
3000
−500
−250
0
250
500
Position around the chromatin interactions (bp)

KLF15

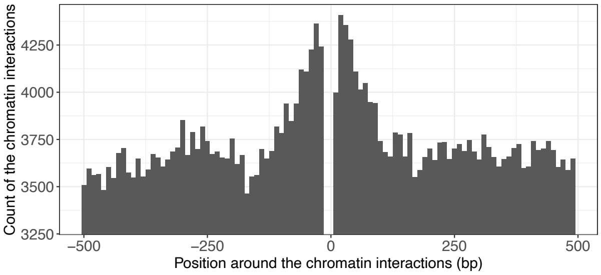


MAF

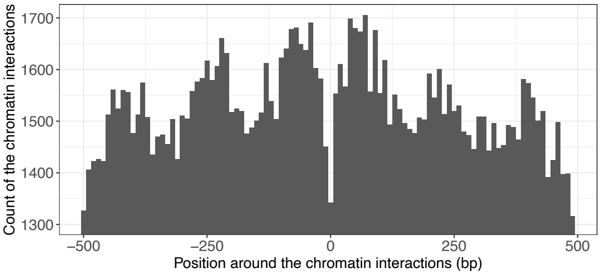


MAFF

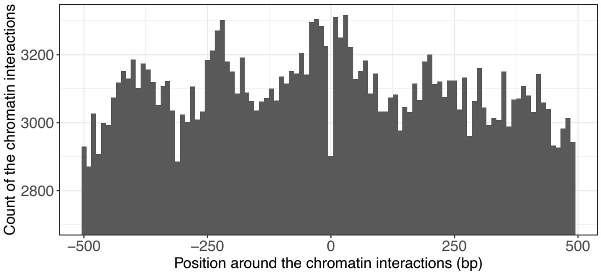


MAFG

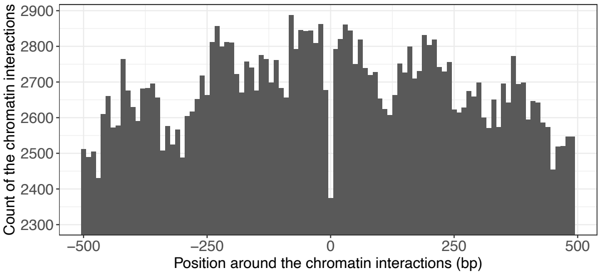


MAFK

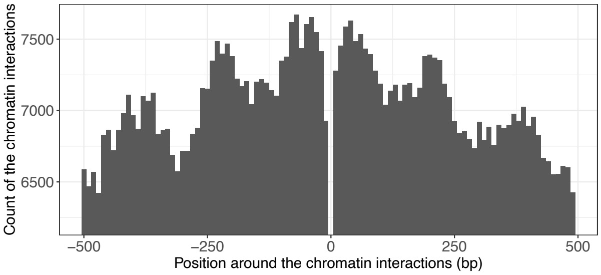


MAX

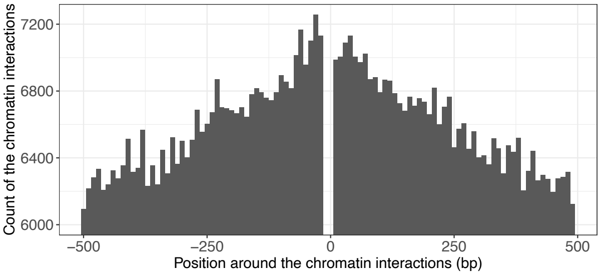


MAZ

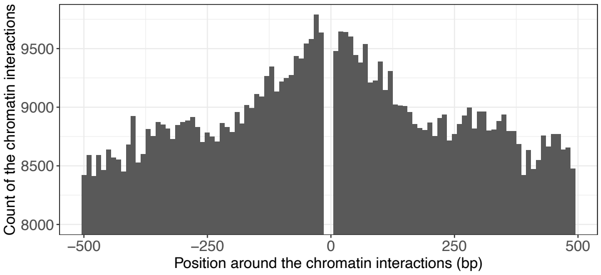


MBD2

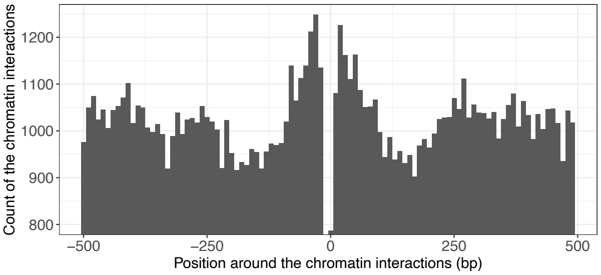


MITF

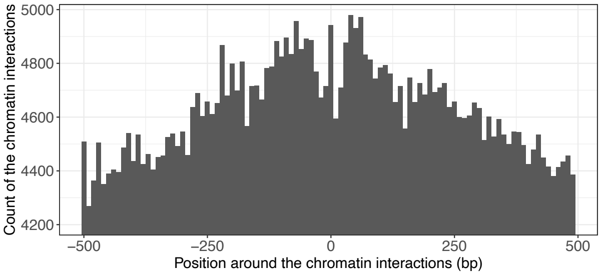


MNT

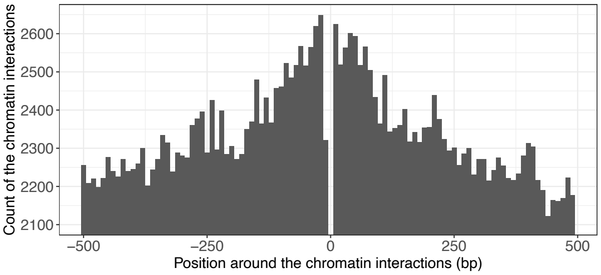


MXI1

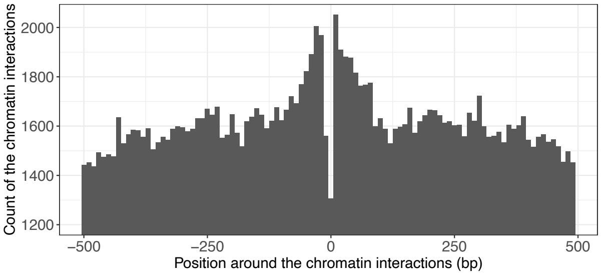


MYB

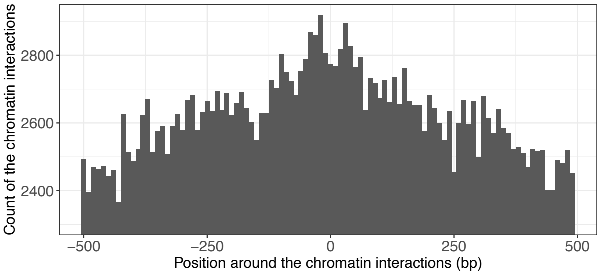

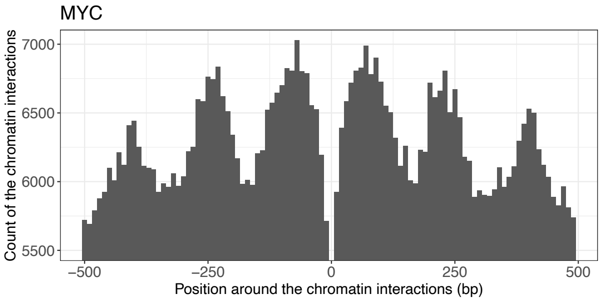
MYC
Count of the chromatin interactions
7000
6500
6000
5500
−500
−250
0
250
500
Position around the chromatin interactions (bp)

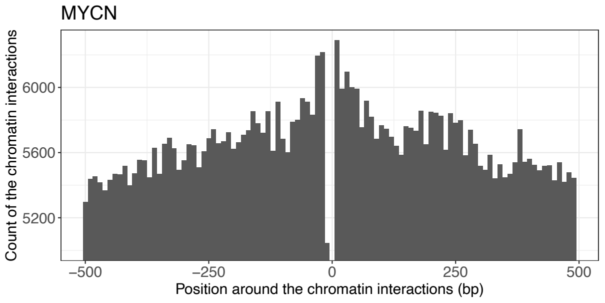
MYCN
Count of the chromatin interactions
6000
5600
5200
−500
−250
0
250
500
Position around the chromatin interactions (bp)

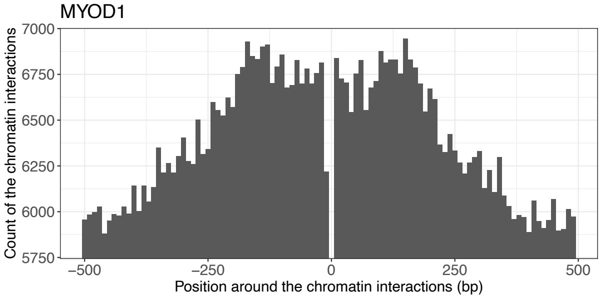
MYOD1
Count of the chromatin interactions
7000
6750
6500
6250
6000
5750
−500
−250
0
250
500
Position around the chromatin interactions (bp)

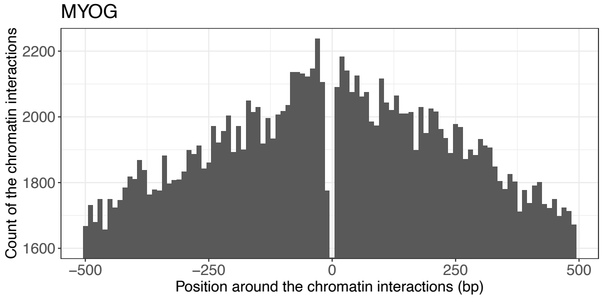
MYOG
Count of the chromatin interactions
2200
2000
1800
1600
−500
−250
0
250
500
Position around the chromatin interactions (bp)

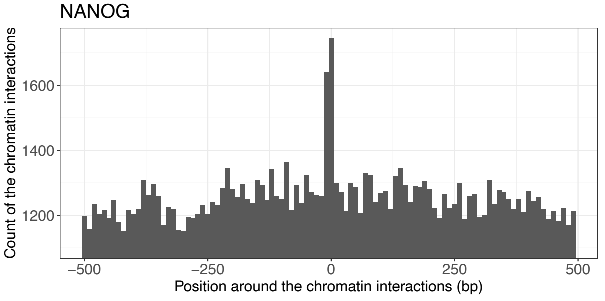
NANOG
Count of the chromatin interactions
1600
1400
1200
−500
−250
0
250
500
Position around the chromatin interactions (bp)

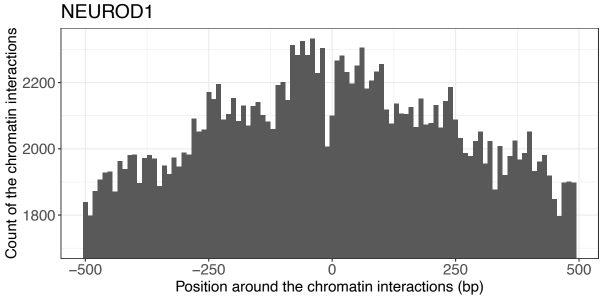
NEUROD1
Count of the chromatin interactions
2200
2000
1800
−500
−250
0
250
500
Position around the chromatin interactions (bp)

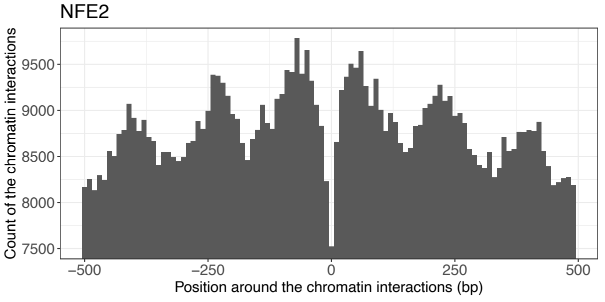
NFE2
Count of the chromatin interactions
9500
9000
8500
8000
7500
−500
−250
0
250
500
Position around the chromatin interactions (bp)

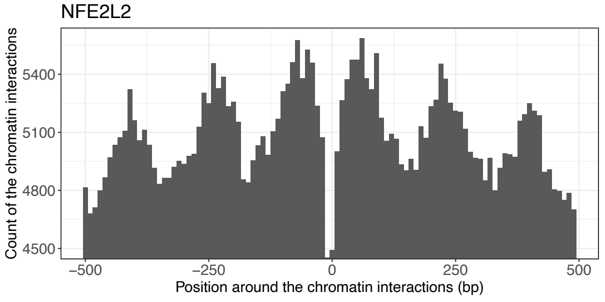
NFE2L2
Count of the chromatin interactions
5400
5100
4800
4500
−500
−250
0
250
500
Position around the chromatin interactions (bp)

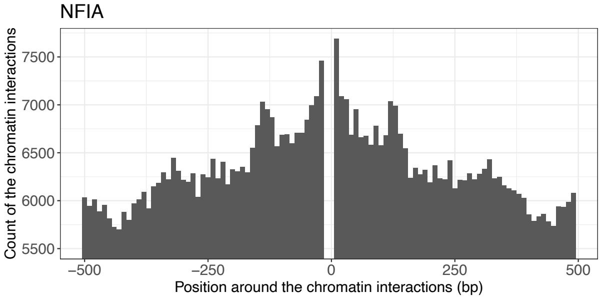
NFIA
Count of the chromatin interactions
7500
7000
6500
6000
5500
−500
−250
0
250
500
Position around the chromatin interactions (bp)

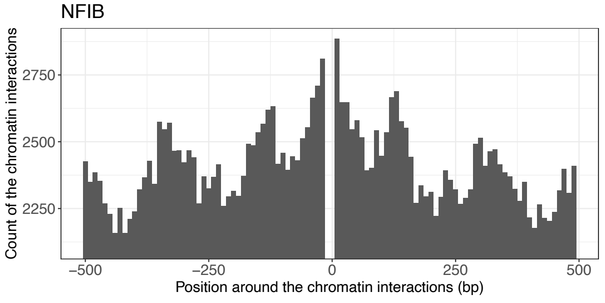
NFIB
Count of the chromatin interactions
2750
2500
2250
−500
−250
0
250
500
Position around the chromatin interactions (bp)

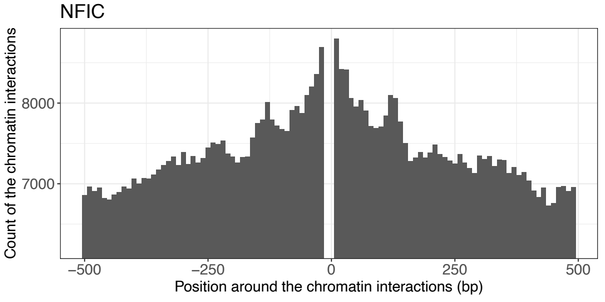
NFIC
Count of the chromatin interactions
8000
7000
−500
−250
0
250
500
Position around the chromatin interactions (bp)

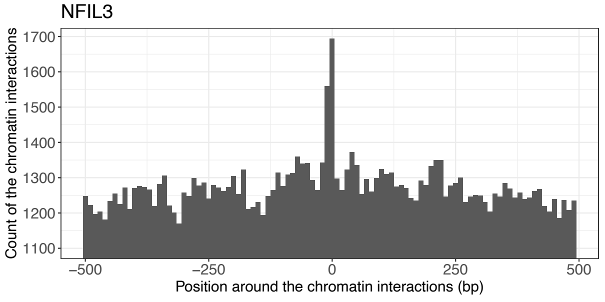
NFIL3
Count of the chromatin interactions
1700
1600
1500
1400
1300
1200
1100
−500
−250
0
250
500
Position around the chromatin interactions (bp)

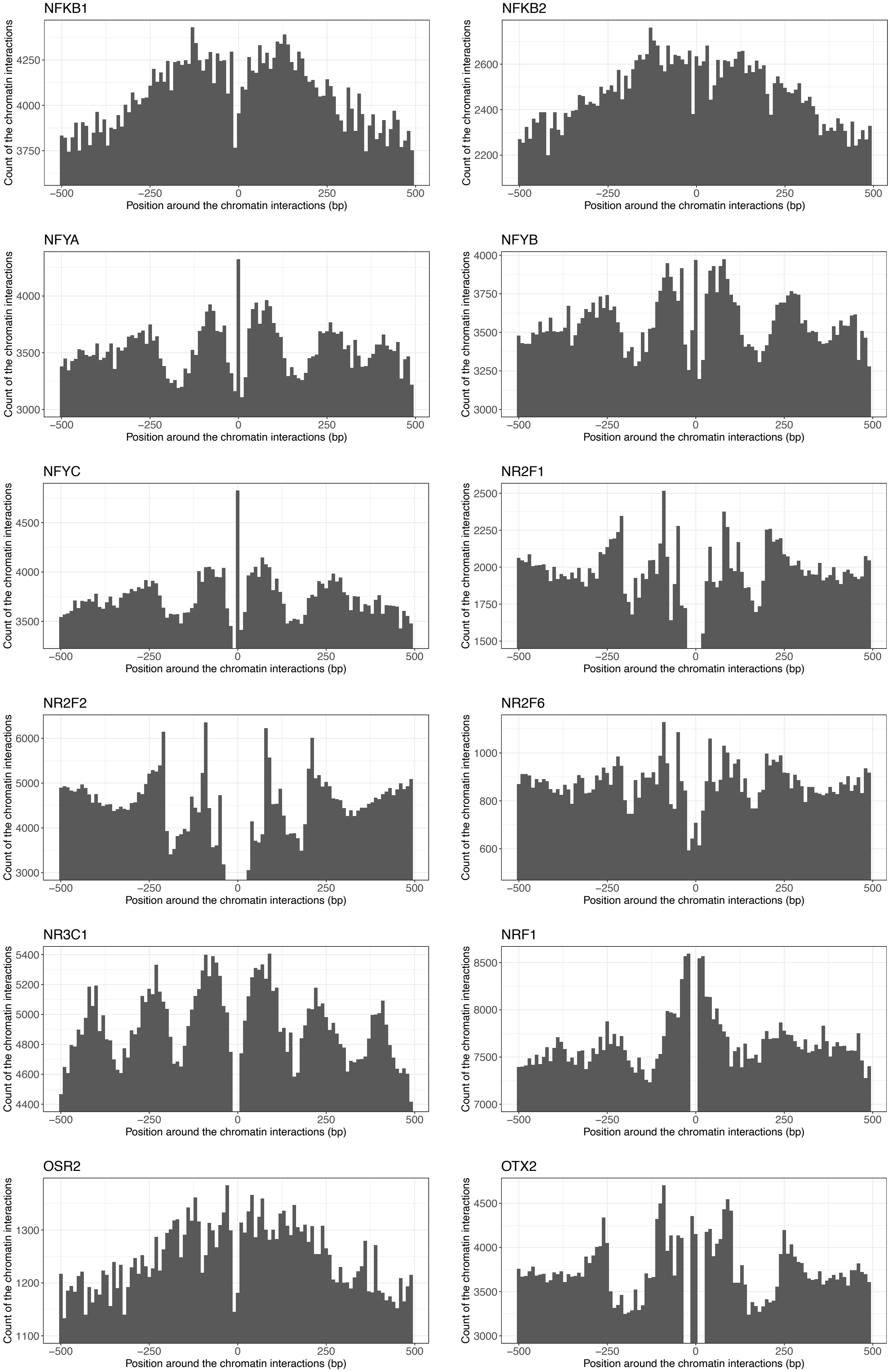
NFKB1
NFKB2
NFYA
NFYB
NFYC
NR2F1
NR2F2
NR2F6
NR3C1
NRF1
OSR2
OTX2
Count of the chromatin interactions
Position around the chromatin interactions (bp)
−500
−250
0
250
500

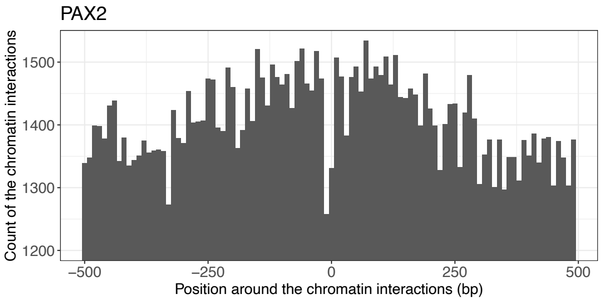
PAX2
Count of the chromatin interactions
1500
1400
1300
1200
−500
−250
0
250
500
Position around the chromatin interactions (bp)

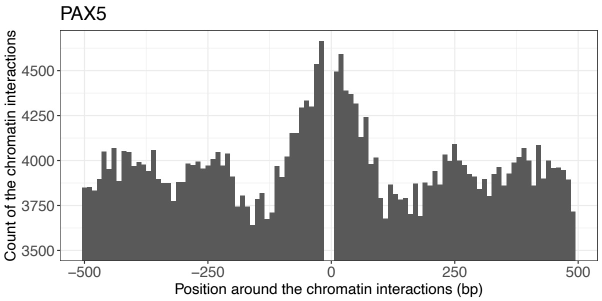
PAX5
Count of the chromatin interactions
4500
4250
4000
3750
3500
−500
−250
0
250
500
Position around the chromatin interactions (bp)

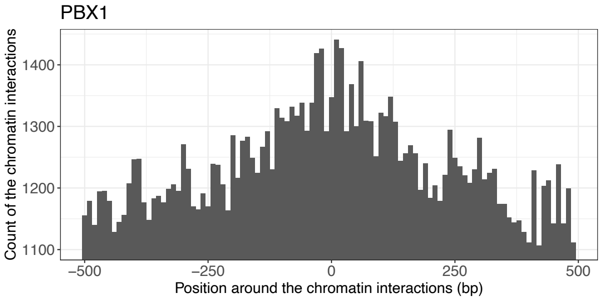
PBX1
Count of the chromatin interactions
1400
1300
1200
1100
−500
−250
0
250
500
Position around the chromatin interactions (bp)

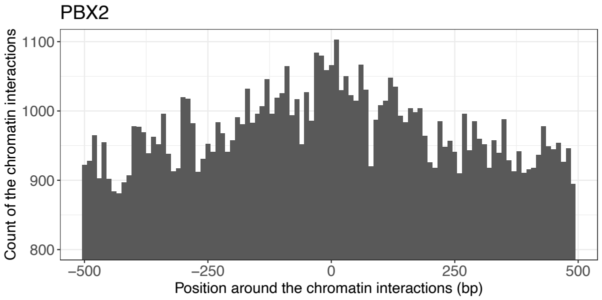
PBX2
Count of the chromatin interactions
1100
1000
900
800
−500
−250
0
250
500
Position around the chromatin interactions (bp)

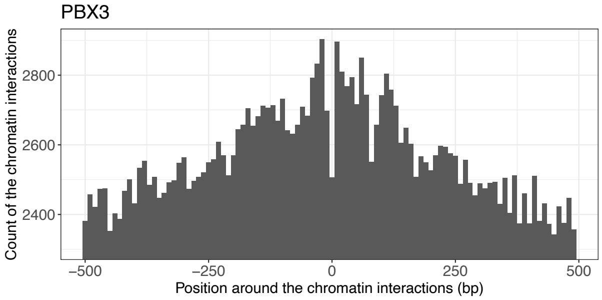
PBX3
Count of the chromatin interactions
2800
2600
2400
−500
−250
0
250
500
Position around the chromatin interactions (bp)

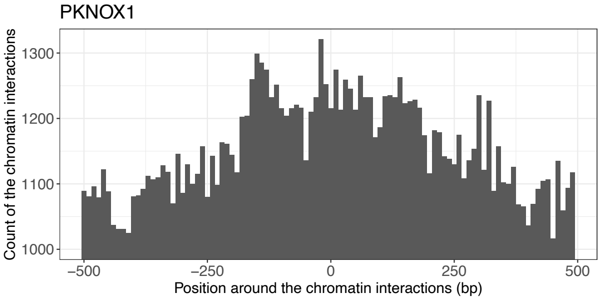
PKNOX1
Count of the chromatin interactions
1300
1200
1100
1000
−500
−250
0
250
500
Position around the chromatin interactions (bp)

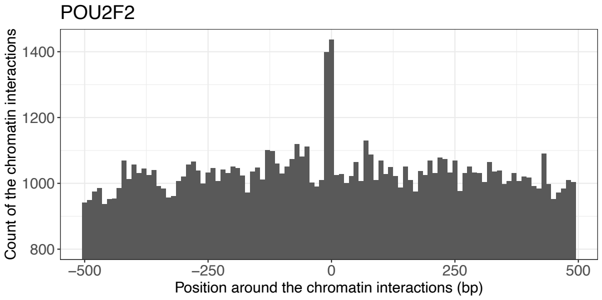
POU2F2
Count of the chromatin interactions
1400
1200
1000
800
−500
−250
0
250
500
Position around the chromatin interactions (bp)

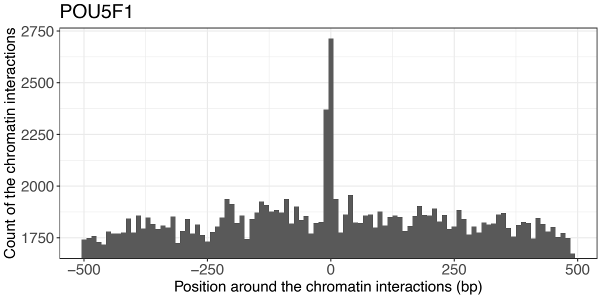
POU5F1
Count of the chromatin interactions
2750
2500
2250
2000
1750
−500
−250
0
250
500
Position around the chromatin interactions (bp)

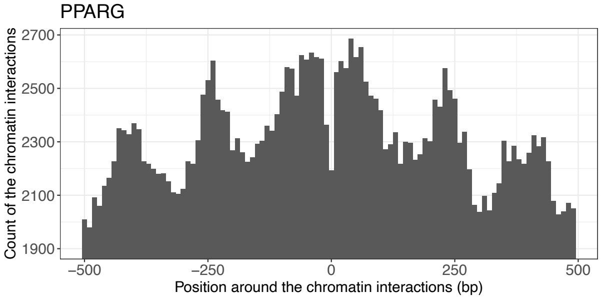
PPARG
Count of the chromatin interactions
2700
2500
2300
2100
1900
−500
−250
0
250
500
Position around the chromatin interactions (bp)

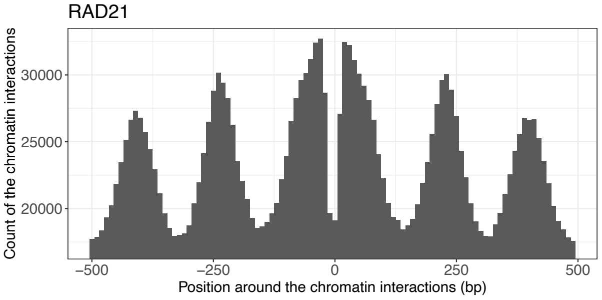
RAD21
Count of the chromatin interactions
30000
25000
20000
−500
−250
0
250
500
Position around the chromatin interactions (bp)

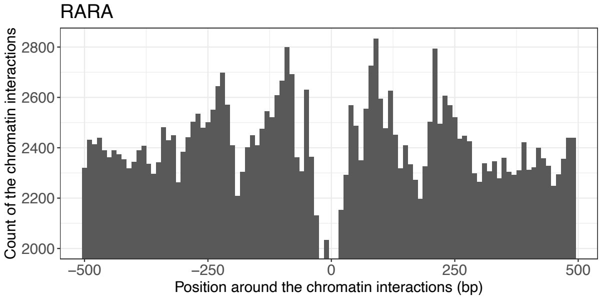
RARA
Count of the chromatin interactions
2800
2600
2400
2200
2000
−500
−250
0
250
500
Position around the chromatin interactions (bp)

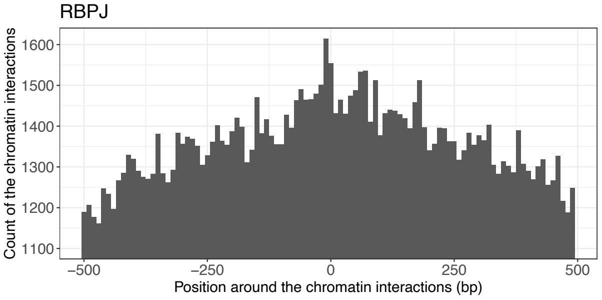
RBPJ
Count of the chromatin interactions
1600
1500
1400
1300
1200
1100
−500
−250
0
250
500
Position around the chromatin interactions (bp)

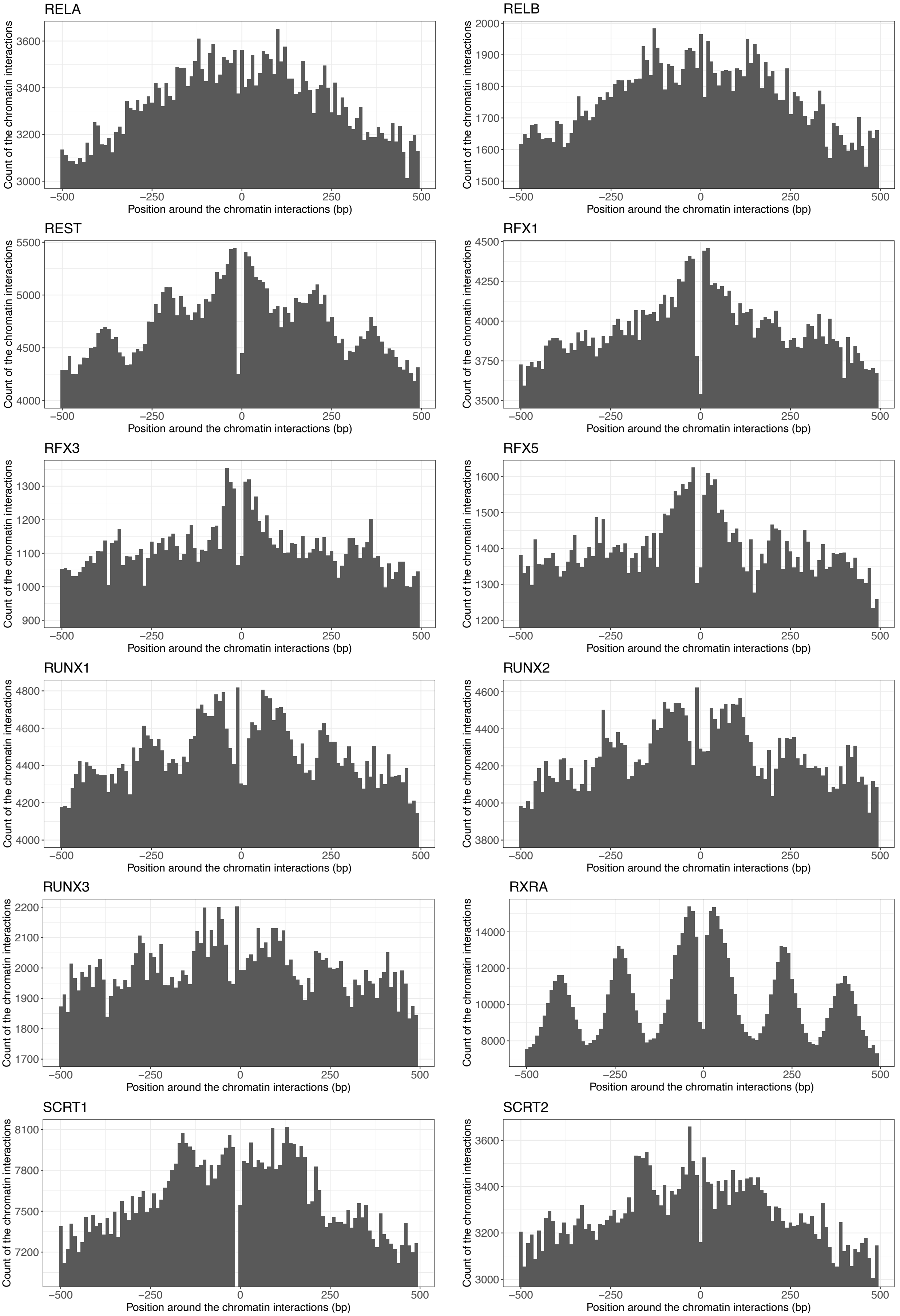
RELA
RELB
REST
RFX1
RFX3
RFX5
RUNX1
RUNX2
RUNX3
RXRA
SCRT1
SCRT2
Count of the chromatin interactions
Position around the chromatin interactions (bp)
−500
−250
0
250
500

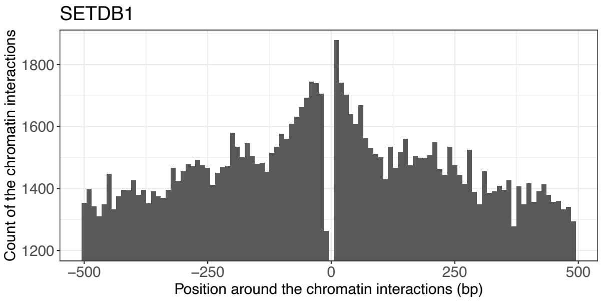
SETDB1
Count of the chromatin interactions
1200
1400
1600
1800
−500
−250
0
250
500
Position around the chromatin interactions (bp)

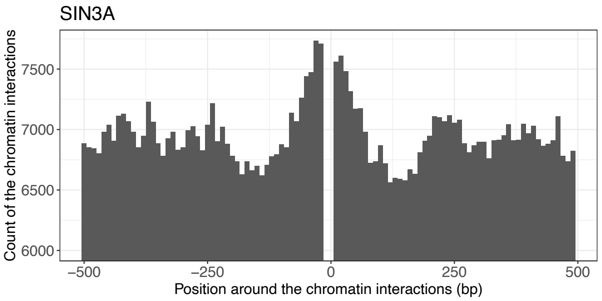
SIN3A
Count of the chromatin interactions
6000
6500
7000
7500
−500
−250
0
250
500
Position around the chromatin interactions (bp)

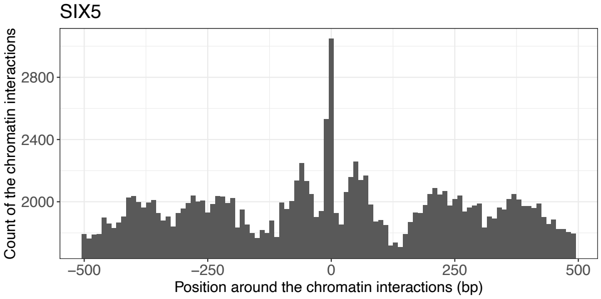
SIX5
Count of the chromatin interactions
2000
2400
2800
−500
−250
0
250
500
Position around the chromatin interactions (bp)

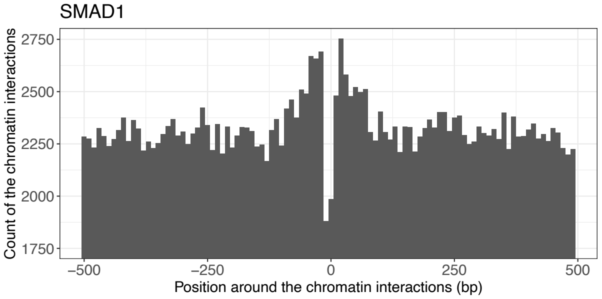
SMAD1
Count of the chromatin interactions
1750
2000
2250
2500
2750
−500
−250
0
250
500
Position around the chromatin interactions (bp)

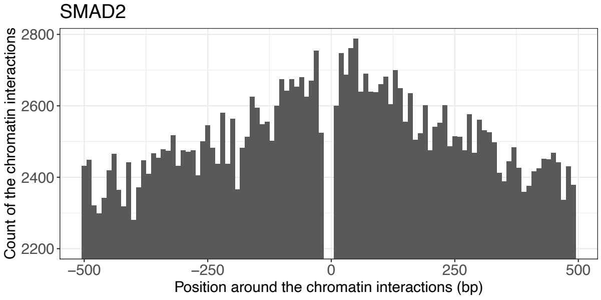
SMAD2
Count of the chromatin interactions
2200
2400
2600
2800
−500
−250
0
250
500
Position around the chromatin interactions (bp)

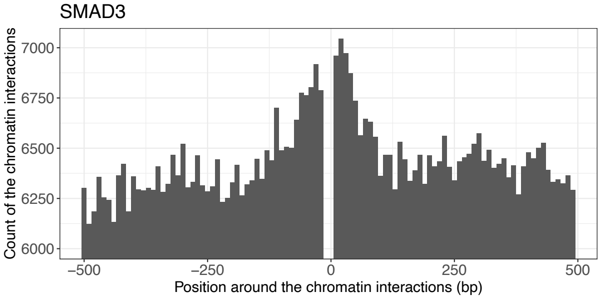
SMAD3
Count of the chromatin interactions
6000
6250
6500
6750
7000
−500
−250
0
250
500
Position around the chromatin interactions (bp)

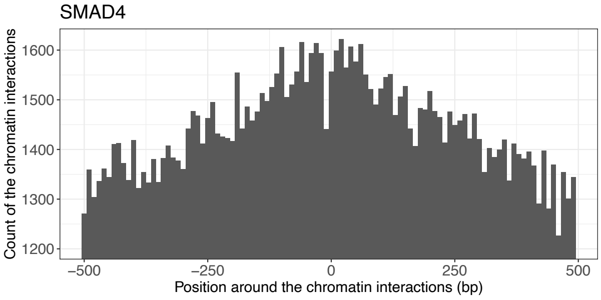
SMAD4
Count of the chromatin interactions
1200
1300
1400
1500
1600
−500
−250
0
250
500
Position around the chromatin interactions (bp)

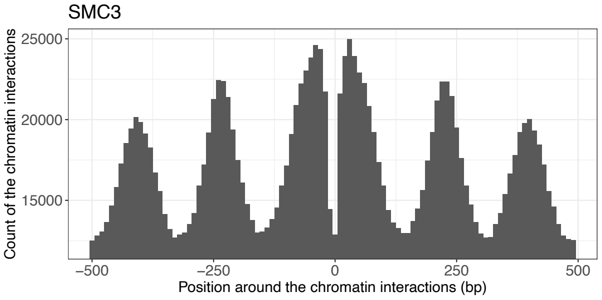
SMC3
Count of the chromatin interactions
15000
20000
25000
−500
−250
0
250
500
Position around the chromatin interactions (bp)

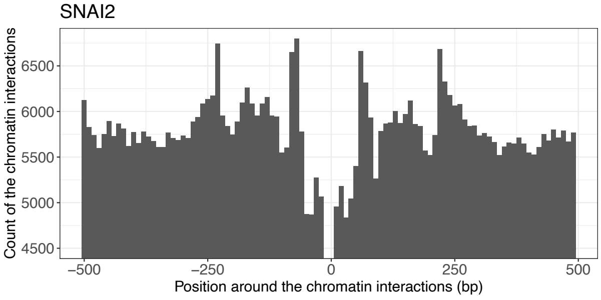
SNAI2
Count of the chromatin interactions
4500
5000
5500
6000
6500
−500
−250
0
250
500
Position around the chromatin interactions (bp)

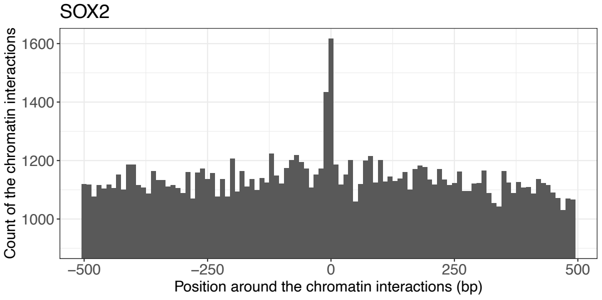
SOX2
Count of the chromatin interactions
1000
1200
1400
1600
−500
−250
0
250
500
Position around the chromatin interactions (bp)

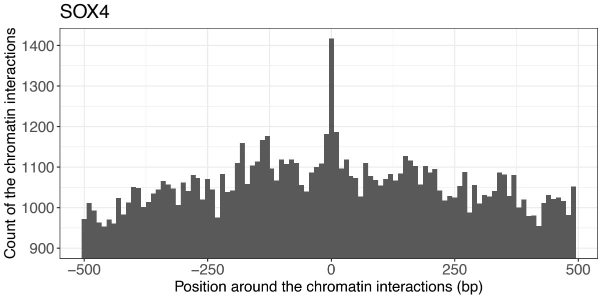
SOX4
Count of the chromatin interactions
900
1000
1100
1200
1300
1400
−500
−250
0
250
500
Position around the chromatin interactions (bp)

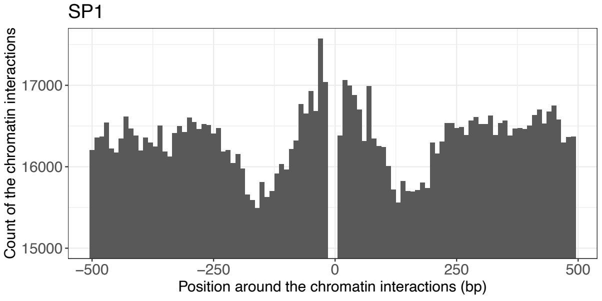
SP1
Count of the chromatin interactions
15000
16000
17000
−500
−250
0
250
500
Position around the chromatin interactions (bp)

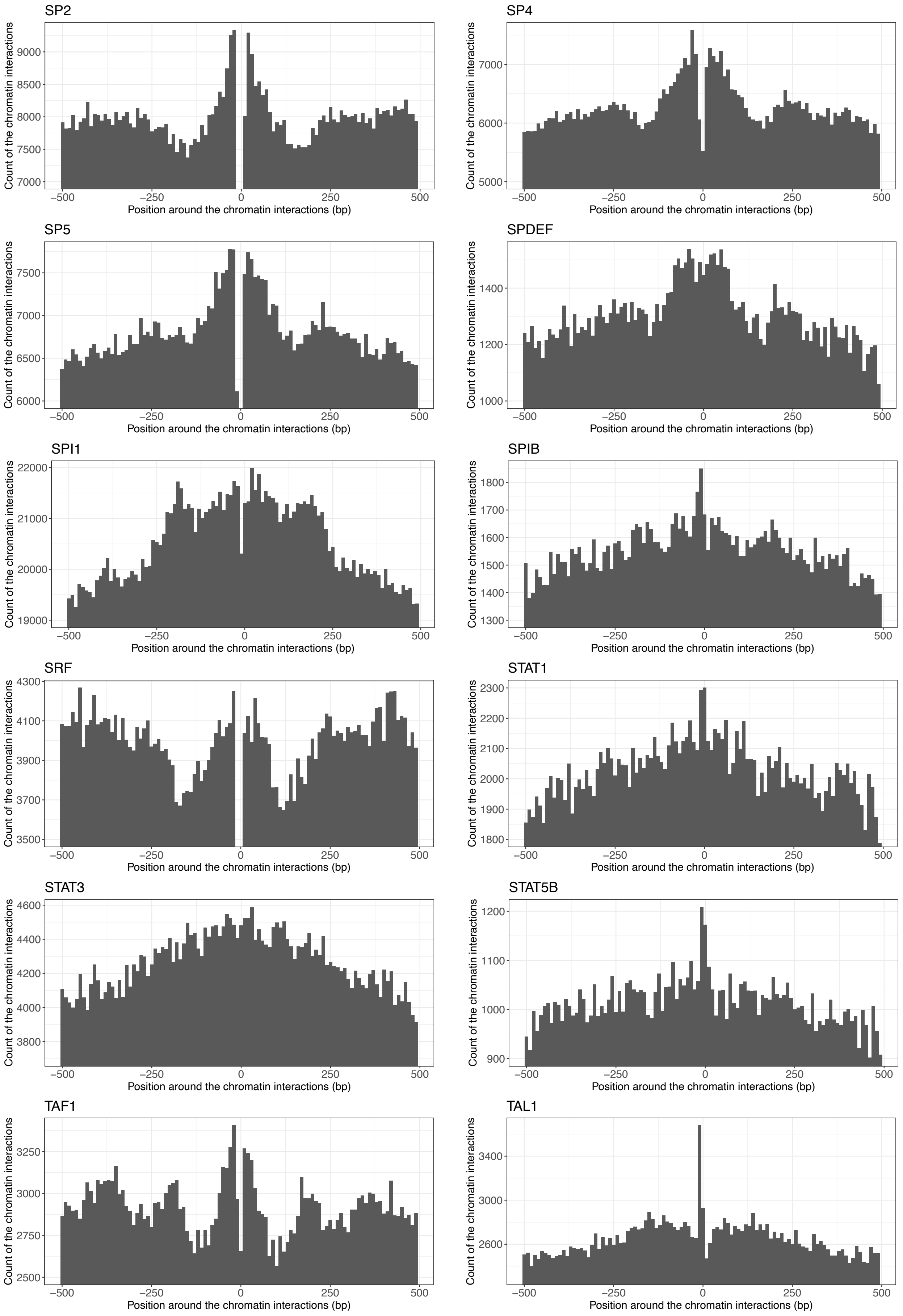
SP2
SP4
SP5
SPDEF
SPI1
SPIB
SRF
STAT1
STAT3
STAT5B
TAF1
TAL1
Count of the chromatin interactions
Position around the chromatin interactions (bp)

TCF3

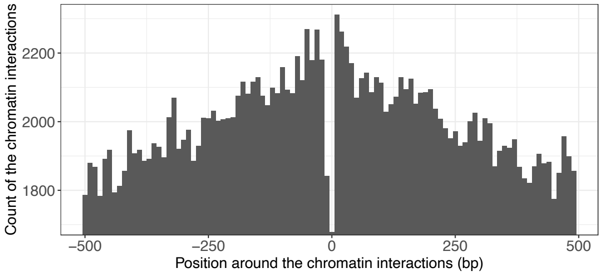


TCF4

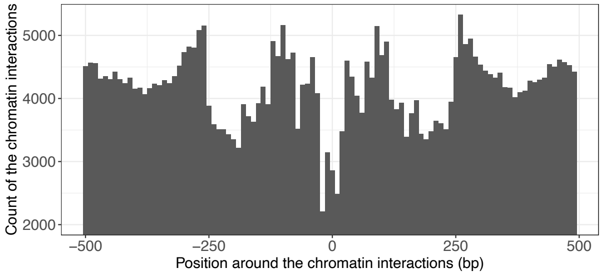


TCF7L2

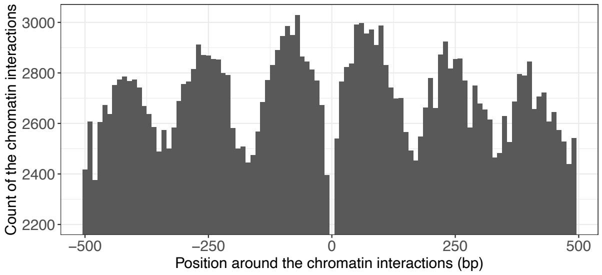


TCF12

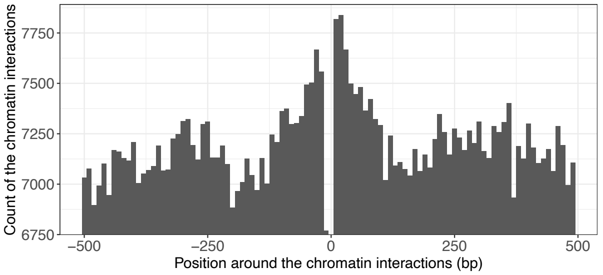


TCF21

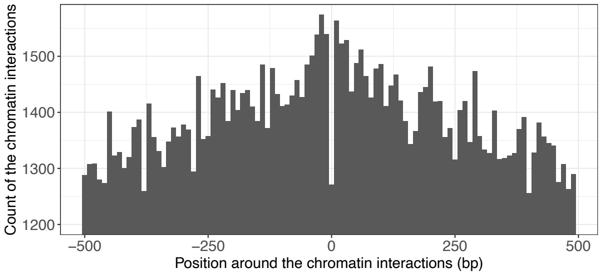


TEAD1

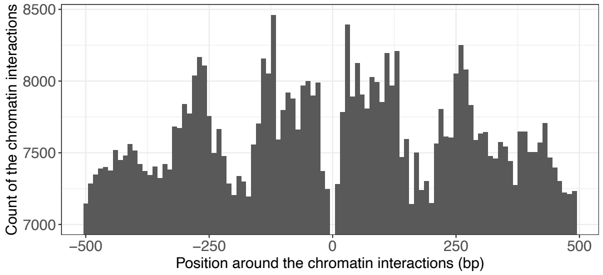


TEAD3

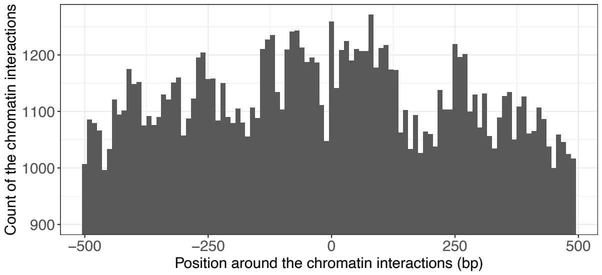


TEAD4

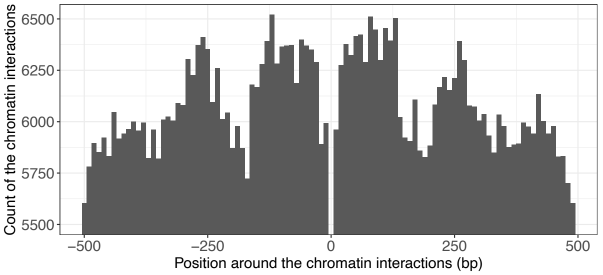


TFAP2A

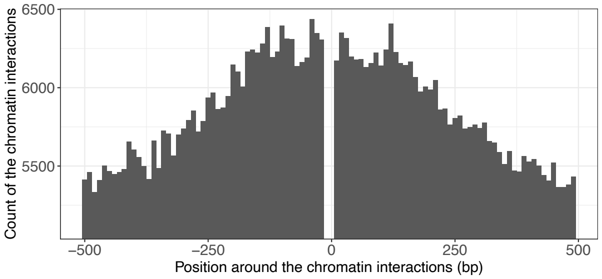


TFAP2C

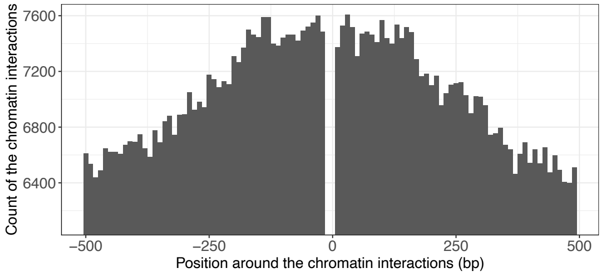


TFAP4

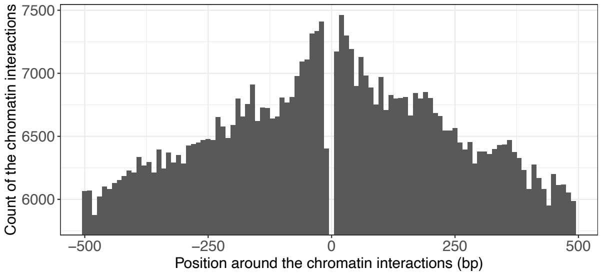


TFDP1

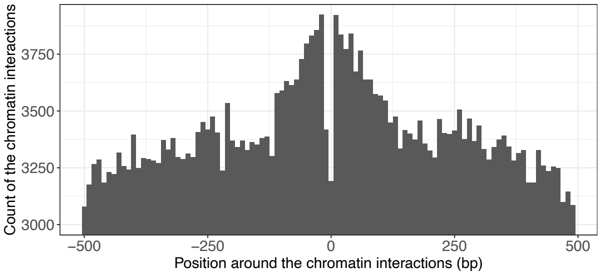

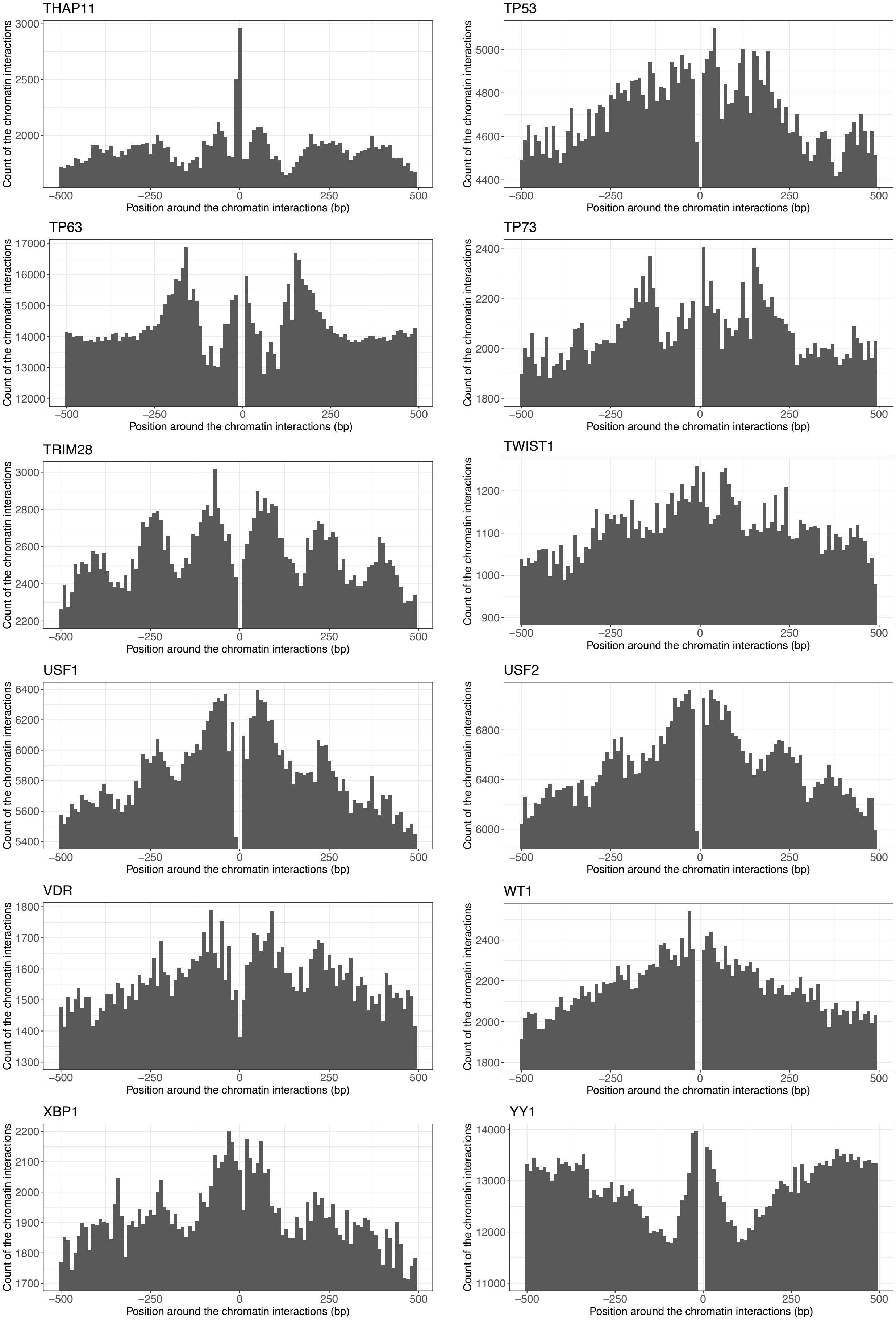
THAP11
3000
2500
2000
TP53
5000
4800
4600
4400
TP63
17000
16000
15000
14000
13000
12000
TP73
2400
2200
2000
1800
TRIM28
3000
2800
2600
2400
2200
TWIST1
1200
1100
1000
900
USF1
6400
6200
6000
5800
5600
5400
USF2
6800
6400
6000
VDR
1800
1700
1600
1500
1400
1300
WT1
2400
2200
2000
1800
XBP1
2200
2100
2000
1900
1800
1700
YY1
14000
13000
12000
11000
Count of the chromatin interactions
−500
−250
0
250
500
Position around the chromatin interactions (bp)

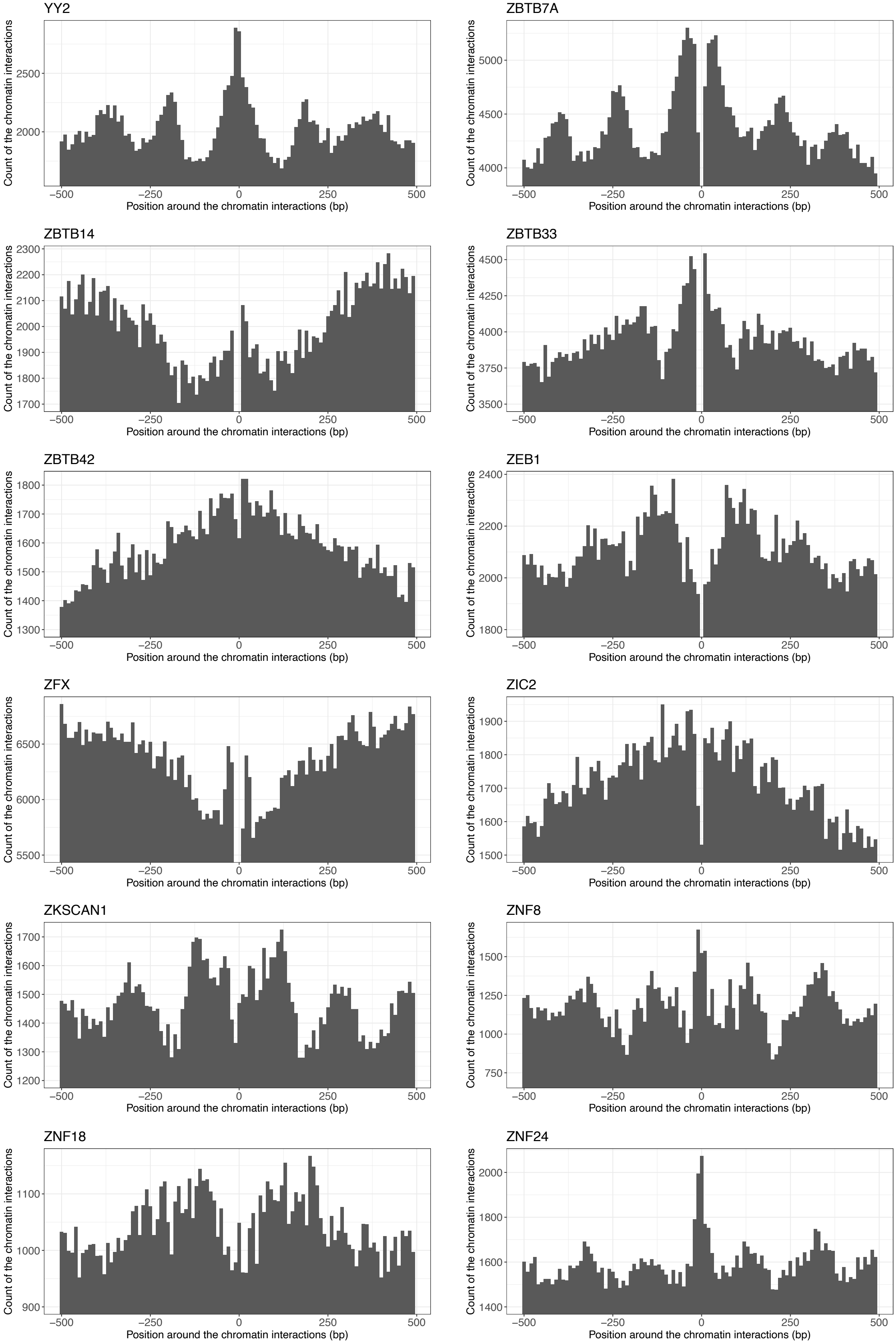
YY2
ZBTB7A
ZBTB14
ZBTB33
ZBTB42
ZEB1
ZFX
ZIC2
ZKSCAN1
ZNF8
ZNF18
ZNF24
Count of the chromatin interactions
Position around the chromatin interactions (bp)

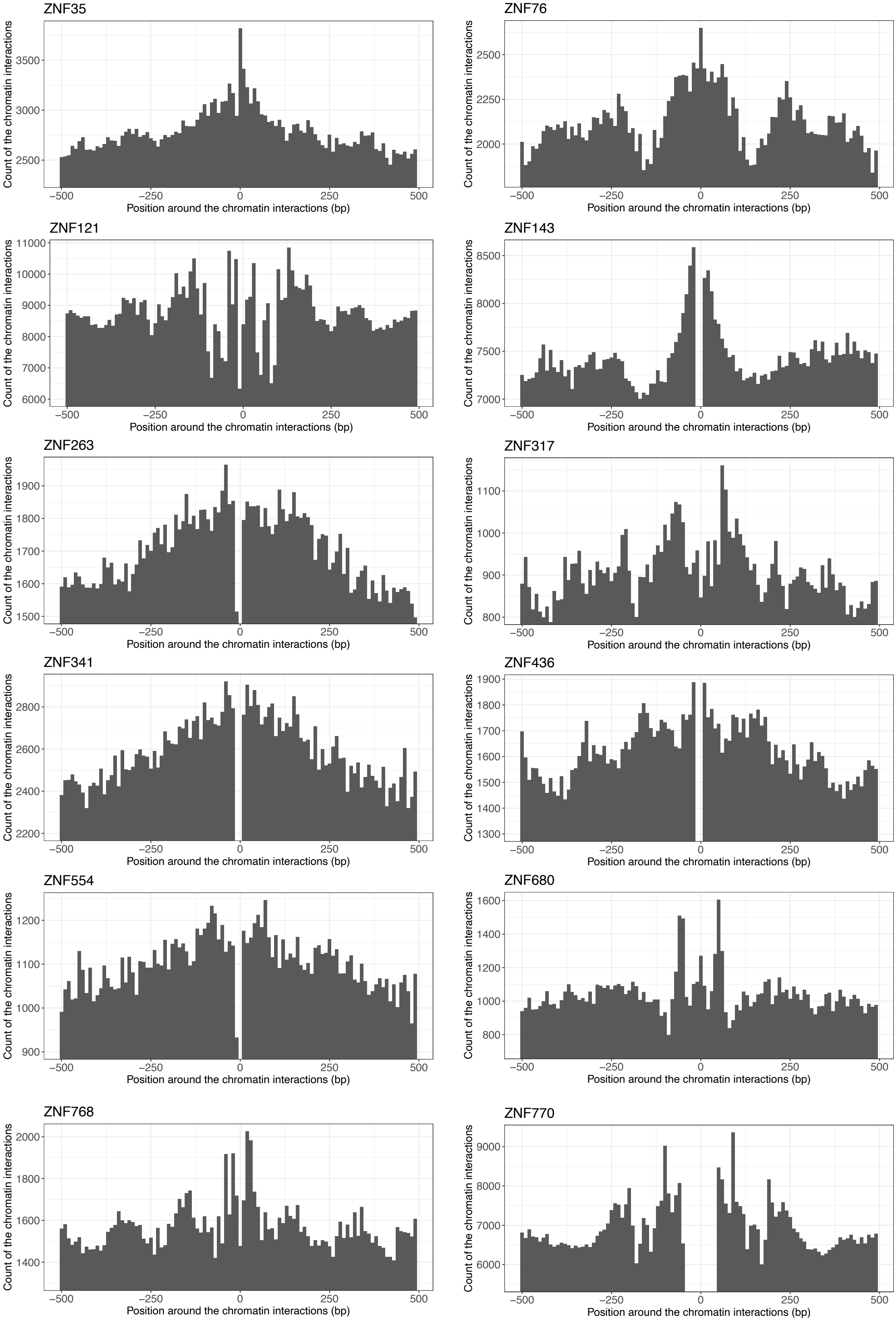
ZNF35
ZNF76
ZNF121
ZNF143
ZNF263
ZNF317
ZNF341
ZNF436
ZNF554
ZNF680
ZNF768
ZNF770
Count of the chromatin interactions
Position around the chromatin interactions (bp)
−500
−250
0
250
500

b

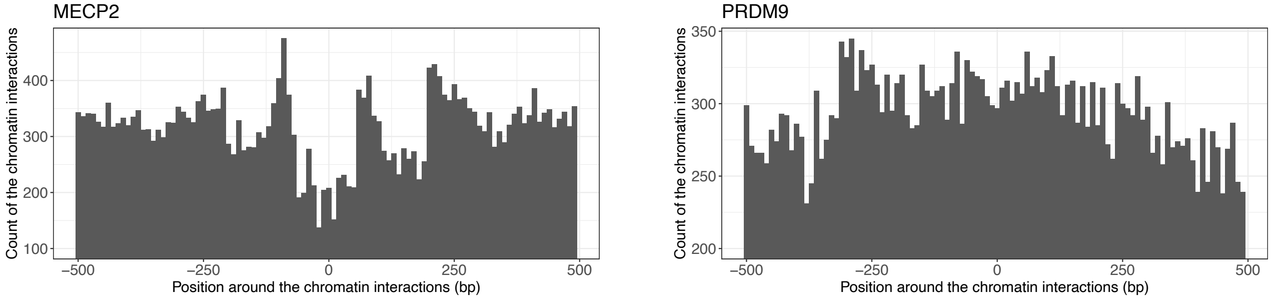
MECP2
PRDM9
Count of the chromatin interactions
Position around the chromatin interactions (bp)

c

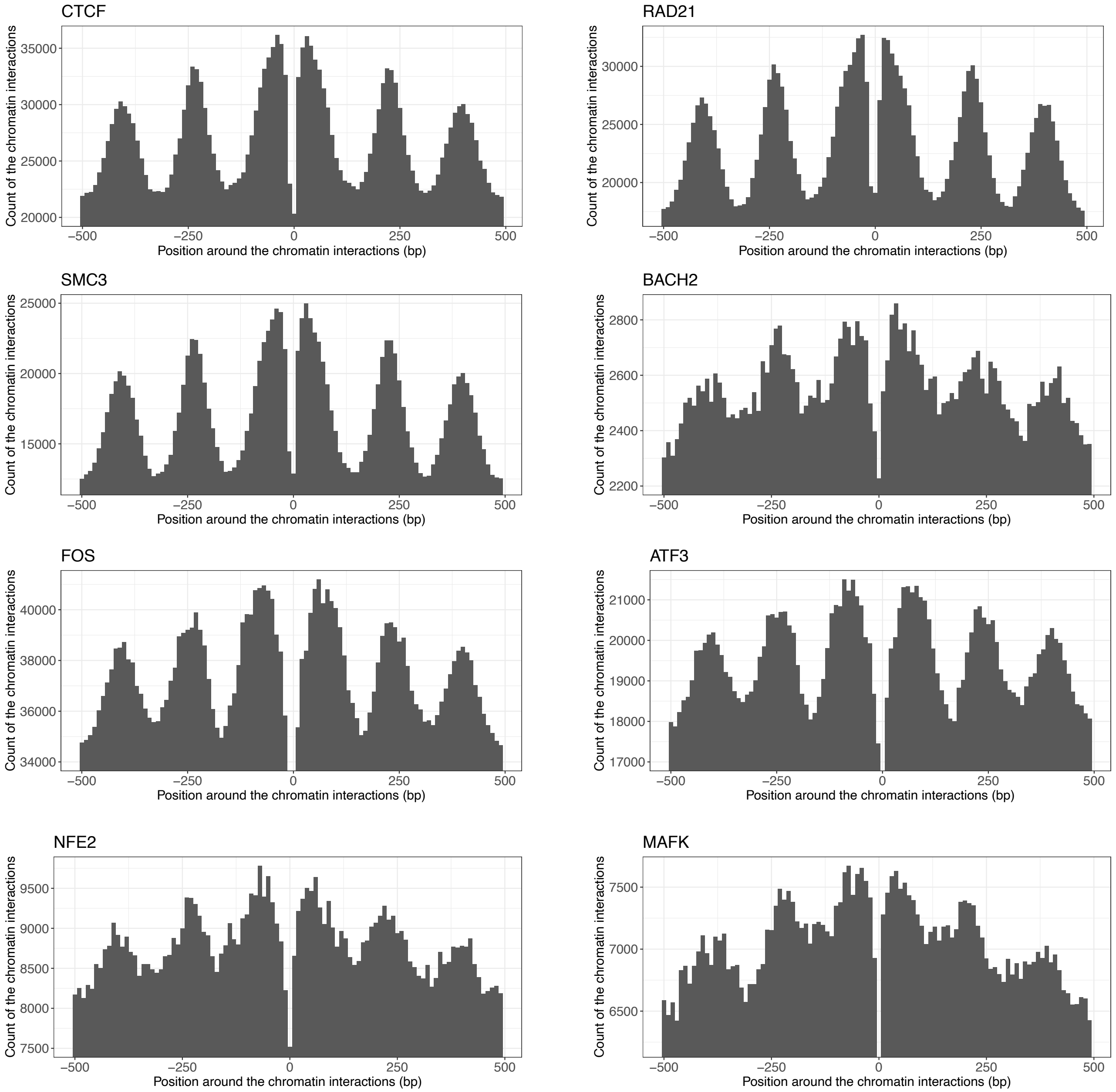
CTCF
RAD21
SMC3
BACH2
FOS
ATF3
NFE2
MAFK
Count of the chromatin interactions
Position around the chromatin interactions (bp)
