## Supplementary material for "Systematic discovery of directional regulatory motifs associated with human insulator sites": Supplementary_Note.pdf

### Results

**Most CTCF DNA-binding sites are positioned at chromatin interaction sites, supporting their key role in chromatin organization and insulator function.**

Our analysis revealed that a substantial proportion (93%) of CTCF ChIP-seq peaks from the GTRD database were found to overlap with chromatin interaction sites identified by ChIA-PET in HFF cells, with a 100-bp window used to define each interaction site. Out of a total of 2,433,551 CTCF ChIP-seq peaks and 123,396,983 chromatin interactions, 2,263,295 peaks overlapped chromatin interaction regions. These findings demonstrate that the vast majority of experimentally determined CTCF binding sites are not only present at, but are likely to be functionally engaged in chromatin interactions. Therefore, CTCF DNA-binding sites identified from experimental ChIP-seq, open chromatin profiles, and CTCF motif analysis are highly representative of those involved in chromatin interaction and insulator activity.

**Directional bias in DNA-binding site orientation is a widespread and functionally relevant feature among insulator-associated DBPs, as revealed by deep learning-based analysis at high-resolution chromatin interaction sites.**

To robustly identify insulator-associated DBPs with a regulatory role in gene expression, we focused on DNA-binding sites located within ±100 bp of chromatin interaction sites positioned more than 5 kb upstream or downstream of transcript regions (Selection Criteria S2, **Fig. S1**), utilizing the high resolution of Micro-C data. Well-characterized insulator DBPs—including CTCF, RAD21, and SMC3—displayed significant orientation-dependent differences in DeepLIFT score distributions, consistent with their established functions. Expanding this analysis to 201 DBPs, we found that 73 DBPs—including CTCF, RAD21, SMC3, ZNF143, and YY1—exhibited significant differences in DeepLIFT score distributions between distinct orientations (e.g., FR vs. RF). Additionally, 60 DBPs, including YY1 and USF1, showed significant differences even between sites with the same orientation (FF vs. RR), as shown in **Supplementary Table S1**, sheet "DBP List 2A". With more stringent criteria, comparing all four orientations (FR, RF, FF, RR) as well as non-directional sites, we observed that 23 DBPs—including CTCF, RAD21, and SMC3—maintained highly significant orientation-dependent DeepLIFT score differences among these classes. Eleven DBPs showed significant differences for same-orientation (FF vs. RR) site comparisons, while three DBPs also displayed significant non-directional site differences ($p < 10^{-3}$, **Table 1** and **Supplementary Table S2**, sheet "DBP List 2A"; FDR-adjusted $p < 0.05$, **Supplementary Table S3**, sheet "DBP List 2A"). These findings highlight that both orientation-dependent and -independent DNA-binding site biases are widespread among putative insulator-associated DBPs, supporting their functional relevance for gene regulation and chromatin architecture.

**Stratified analysis by DBP binding site number and chromatin context further validated the specificity of insulator-associated DBP prediction.**

When restricting our analysis to DBPs with 100–1,000 DNA-binding sites in HFF cells, we observed a smaller number of cases of orientation bias than among DBPs with 1,000 or more binding sites (**Supplementary Table S1**). This indicates that the detection of orientation bias is sensitive to the sample size of binding sites available for statistical analysis. Recognizing that chromatin interactions comprise various types—including enhancer–promoter interactions (EPIs), protein-protein interactions, and general chromatin looping—we sought to increase specificity by excluding DNA-binding sites that might be associated with active regulatory elements. To increase specificity, we applied ChIP-seq data for promoter- and enhancer-associated histone marks (H3K4me3, H3K27ac, and H3K4me1) in HFF cells, defining two criteria: (1) analyses omitting chromatin interaction data (Selection Criteria S3) and (2) analyses focusing on chromatin interactions spanning more than 5 kb upstream and downstream of a transcript (Selection Criteria S4). For both criteria, analyses were further stratified by DBPs with at least 1,000 binding sites and those with 100–1,000 sites (**Supplementary Tables S1–3**). Importantly, using DNA-binding sites that did not overlap these histone marks, we successfully identified several known insulator-associated DBPs previously reported in the literature, such as BCL6, FOXA3, HNF4A, MYB, USF1, and ZNF143 (see Identification and classification of insulator-associated DBPs; **Supplementary Table S2**, sheet “DBP List 1,2”). These findings demonstrate the robustness of our prediction approach in identifying functionally relevant insulator-associated DBPs, even under stringent genomic and epigenomic selection criteria.
