## Supplementary figures and images for "Systematic discovery of directional regulatory motifs associated with human insulator sites"

### Supplementary_Fig5.pdf

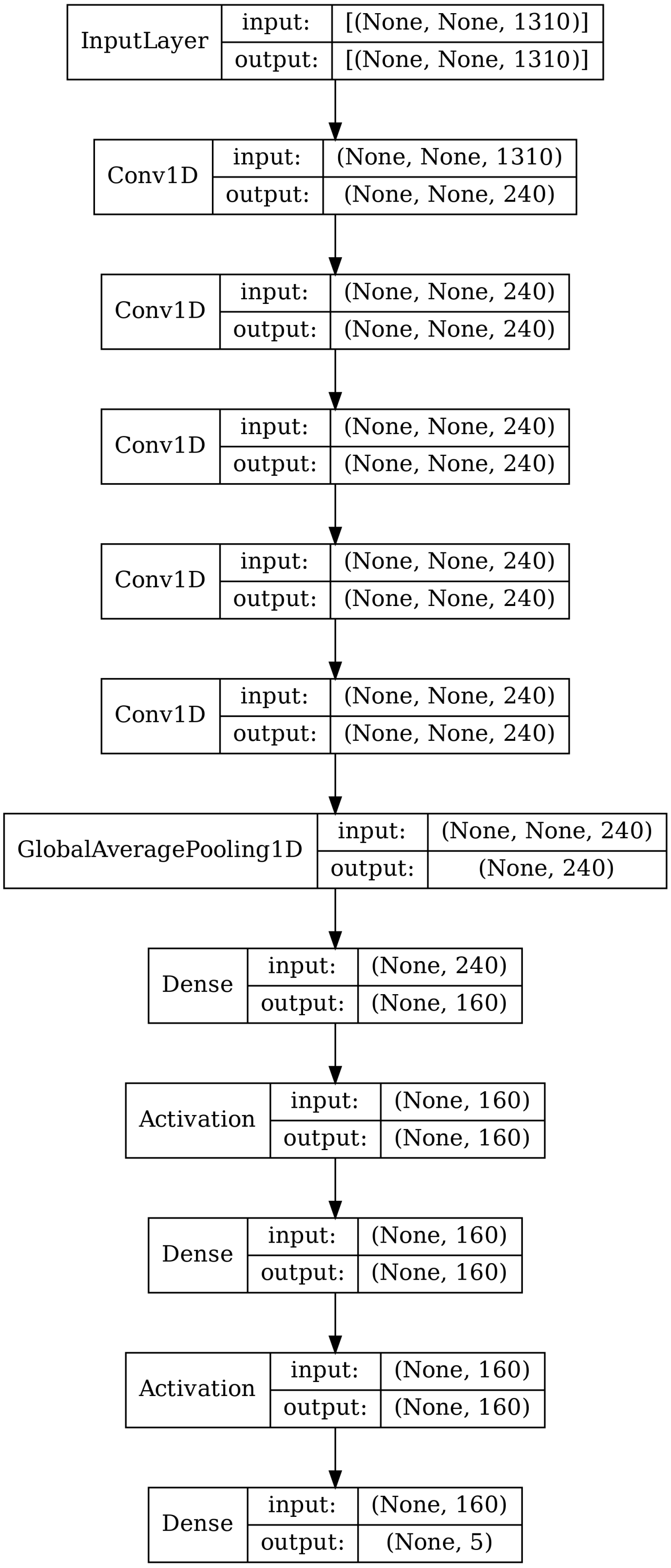


Supplementary Fig. 5. Schematic structure of the deep neural network
